# Characterising genome composition and large structural variation in banana varietal groups

**DOI:** 10.1101/2023.06.08.544197

**Authors:** Janet Higgins, Jaime Andrés Osorio-Guarín, Carolina Olave-Achury, Deisy Lisseth Toloza-Moreno, Ayda Enriquez, Federica Di Palma, Roxana Yockteng, José J. De Vega

## Abstract

**Background:** Bananas and plantains (*Musa* spp.) are one of the most important crops worldwide. The cultivated varieties are vegetatively propagated, and their diversity is essentially fixed over time. Nevertheless, millennia of diversification and selection have led to hundreds of edible varieties. *M. acuminata* and *M. balbisiana* respectively provided the A and B subgenomes that mostly constitute these varieties. Here we aimed to characterise chromosomal exchanges and structural variation among lineages to understand shared foundational events and identify sources of allelic diversity in introgressed loci for genetic improvement.

**Methods:** We identified clonal somatic groups among 188 banana and plantain accessions introduced for cropping in Colombia, using admixture, principal component, and phylogenetic analyses. We established a new alignment-based metric, named *Relative Averaged Alignment* (RAA), to infer subgenome composition (AA, AAB, etc.). We later used comparisons in read coverage along conserved chromosomal windows between the A, B, and S subgenomes to identify introgressions.

**Results:** In our panel, we identify ten varietal groups composed of somatic clones, plus three groups of tetraploid accessions. We demonstrated RAA can be used to infer subgenome composition in the total genome and individual chromosomes. We identified 20 introgressions, several newly reported, among the AAB and ABB varieties. We did not observe B-donor introgression in any AA/AAA varietal groups. We identified variation in length in at least two introgressions, a B-donor introgression in chromosome 7 between the “Maoli” and a “Popoulu” subdivisions, and an S-donor (*M. schizocarpa*) introgression in chromosome 2 in four varietal groups with different compositions (AAA, AAB, ABB, and AA).

**Conclusions:** The extensive distribution of introgressions and the variation in the length of some introgressions between varieties support that the emergence of many varieties can be attributed to intricate founding events, which encompassed multiple instances of hybridisation and subsequent residual backcrossing. We also showed the contribution of *M. schizocarpa* to four cultivated varieties, and proposed subdivision-specific intergenomic recombination in chromosome 7 between subgroups Maoli and Popoulu plantains. Introgressed loci over these 20 introgressions likely provide an extensive resource of allelic diversity to further explore their contribution to disease resistance, climatic adaption, etc. and potential for exploiting in breeding and genome editing.

## BACKGROUND

Bananas, part of the *Musa* genus, are large herbaceous plants grown in tropical and subtropical regions of Southeast Asia, Africa, and America belonging to the Musaceae family. Bananas are one of the most important crops cultivated worldwide, with annual production exceeding 124 million tonnes in 2021 (FAO 1997). However, despite hundreds of banana cultivars worldwide, only a few are grown commercially for large-scale production, with the main commercial banana being triploid somatic clones from the Cavendish subgroup.

The domestication and selection of seedless fruit have resulted in the fixation of parthenocarpy and female sterility in cultivated bananas. Therefore, the crop is primarily vegetatively propagated, and the diversity of cultivated bananas is essentially fixed over time. The introduction of genetic diversity is limited to the somatic accumulation of mutations, somaclonal variation introduced via tissue culture, breeding crosses between sexual (usually diploid) accessions, or in recent years, genome editing. Nevertheless, millennia of diversification of wild genotypes and human selection of hybrids have led to the current existence of hundreds of edible banana and plantain varieties (Heslop-Harrison and Schwarzacher 2007).

The origin of cultivated bananas is believed to involve four genetic pools, mainly through natural inter(sub)specific hybridisation with variable levels of contribution: *Musa acuminata* Colla and *M. balbisiana* Jacq., from various subspecies, are the contributors of the named A and B subgenomes, respectively. In addition, some cultivars show minor contributions from *M. schizocarpa* N.W.Simmonds (S subgenome), and *M. textilis* Nee and its relatives in the *Musa* section *Callimusa* (T subgenome) (Heslop-Harrison and Schwarzacher 2007; D’hont *et al*. 2012). Based on morphology and geographical distribution, *M. acuminata* has been divided into nine subspecies, of which *M. acuminata* subsp. *banksii*, *zebrina* and *malaccensis* appear to be the main origins of A subgenomes observed in cultivated banana varieties (Perrier *et al*. 2011, 2019; Christelová *et al*. 2017).

Cultivated bananas have been classified into groups based on qualitative morphological descriptors and genome composition (AA, AB, AAA, AAB, ABB) (Simmonds and Shepherd 1955). Among them, the most common and widespread are the allotriploids, such as the exporting commercial Cavendish varieties (AAA) or the cooking plantains (AAB). Edible diploid bananas (AA, AB) are also cultivated, especially in subsistence farming systems. Given their socioeconomic importance, over 6,800 *Musa* accessions are currently managed in 30 collections (Ruas *et al*. 2017; Houwe *et al*. 2020), with a large collection maintained at the International Musa Germplasm Transit Centre (ITC), comprising more than 1,600 accessions (Houwe *et al*. 2020). Its genetic diversity has been well-characterised using several genotyping methods, including SSR, ISSR, RAM and SCoT markers (Perrier *et al*. 2011; Florez *et al*. 2012; Christelová *et al*. 2017; Igwe *et al*. 2021), flow-cytometric analysis to determine ploidy level (Christelová *et al*. 2017), and chromosome painting to analyse karyotyping (Simonikova *et al*. 2019, 2022). However, genetic information is still limited due to polyploidy, parthenocarpy and complexity inherent in sample collection.

Recurrent chromatin exchanges between homoeologous chromosomes have been described, tentatively originating via following backcrosses between hybrids with residual fertility and the parental donors (Baurens *et al*. 2019; Cenci *et al*. 2021). Evidence suggests that most varietal clonal groups may be the product of complex multiple hybridisation events. There is evidence of nine different hybridisation events that gave rise to ABB varieties (Cenci *et al*. 2021). Questions remain concerning the large structural variation among *Musa* spp., particularly following the growing evidence that most varietal clonal groups may be the product of complex multiple hybridization events (Baurens *et al*. 2019; Cenci *et al*. 2021).

The objectives of the present study were to (i) clarify the extent of and diversity within banana varietal clonal groups, i.e., the level of somatic variation within groups, and (ii) to identify lineage-specific large structural variation and chromosomal exchanges between homoeologous chromosomes. These objectives aim to bring new insights into the origin of banana varieties, such as shared foundational events and related ancestry, and clarify regions where genome composition deviates from that generally described for the varietal group, because these loci constitute untampered diversity despite having been fixed for a long time. This information can inform breeding and provide candidate targets for genome editing.

## METHODS

### DNA extraction and sequencing

A total of 190 accessions from the *in-situ* banana collection managed by AGROSAVIA in its research centre in Palmira, Colombia (3.51424, −76.3158), were used for this study (Table S1, additional file 1). The passport information is available in *MGIS: Musa Germplasm Information System* (Ruas *et al*. 2017) as “COL004”. This diversity panel represents the diversity of banana introduced in Colombia and is actively maintained. Genomic DNA of the accessions was extracted from liquid nitrogen macerated young leave material using the DNeasy Plant Mini Kit (QIAGEN, Germany) and shipped to the *Genomic Pipelines* sequencing service at the Earlham Institute (Norwich, UK), where DNA libraries were constructed and assessed using a modified version of *Illumina*’s “Nextera DNA library Prep” protocol, known as “Low Input Transposase Enabled (LITE)” protocol (Beier *et al*. 2017; Perez-Sepulveda *et al*. 2021), and sequenced in an *Illumina* Novaseq S4 lane (150bp paired-end reads) aiming for ∼7X average depth per accession.

### Read alignment and relative averaged alignment (RAA) metric

Raw reads in FastQ format were pre-processed using Trim Galore v0.5 (Krueger 2015) with the options for *Illumina* paired reads and to remove the *Nextera* adaptors, any bases with quality under 20, and a minimum read length of 80bp. Processed reads were aligned using BWA MEM v0.7.17 (Li 2013), with the options −M and −R to define read-groups, against the four genome references.

Several banana reference genome assemblies are available (Droc *et al*. 2013). We used the high-quality chromosome level assemblies of a *M. acuminata* doubled-haploid cv. Pahang accession, a *M. balbisiana* cv. Pisang Klutuk Wulung accession (Wang *et al*. 2019), and a *M. schizocarpa* wild accession (Belser *et al*. 2018). These chromosomal-level assemblies are respectively referred to as “A-genome reference”, “B-genome reference”, and “S-genome reference” in the paper. The length of the reference for the A-genome, B- genome and S-genome references was 479.219 Mb, 457.198 Mb, and 525.284 Mb, respectively. In addition, we used an unpublished *Nanopore* long-read assembly of accession ITC0643 from the Bluggoe subgroup (ABB) kindly made available to us (M. Rouard, *per. comm.*).

Coverage and alignment statistics were obtained using Samtools flagstat v1.7 (Li *et al*. 2009) for the complete genome and separately for each of the 11 chromosomes in each reference. The “relative averaged alignment” (RAA) is a normalised percentage of properly paired reads in a sample and reference that accounts for variation in sample quality (PCR duplications, DNA quality, etc) and differences in the genetic distance between varieties and the reference (reference bias). RAA was calculated by dividing the percentage of properly paired reads from a sample in a reference by a weight factor (in the range of 0.95- 1.05). The weight factor was obtained by averaging the ratios in each of the reference genomes between the properly paired reads in the sample and variety cluster (example in Table S2). RAA per chromosome was similarly calculated except for each chromosome’s alignment statistics instead of the total genome.

### SNP calling and population structure analysis

SNP calling was carried out against the A-genome and B-genome references. The alignment BAM files were sorted using Samtools v1.7, and duplicate reads were marked using Picard tools v1.128. SNP calling was done using GATK HaplotypeCaller v3.7.0 (McKenna *et al*. 2010) using all the BAM files as input (multisample mode). The resulting VCF files were filtered for bi-allelic SNP calls with a minimal quality value of 100 and a read depth of over 10 and under 300 reads using BCFtools v1.9 (Danecek *et al*. 2021). Later, GATK SelectVariants v3.7.0 was used to remove SNP sites with over 30% missing samples, and finally, SNP sites were filtered for MAF 1% using BCFtools.

The population structure of the diversity panel was estimated using STRUCTURE v2.3.4 (Pritchard *et al*. 2000). SNP calls were transformed to diploid (0/0, 0/1, and 1/1). Heterozygous calls (0/1) were changed to missing (./.) if either allele was supported by less than two reads. The files were then thinned to one SNP site within 50 bps (--thin 50) using VCFtools v0.1.13 (Danecek *et al*. 2011). The admixture model was used with a burn-in period length of 10,000 and 50,000 MCMC iterations. Twenty independent runs were performed for each K from 2 to 10. Ten replicated Q-matrices belonging to the largest cluster were aligned using the R package POPHELPER v2.2.7 (Francis 2017), and then merged using CLUMPP v1.1.2 (Jakobsson and Rosenberg 2007). Delta K (ΔK) was estimated using the Evanno method (Evanno *et al*. 2005) within POPHELPER.

Principal component analysis (PCA) was carried out using Tassel v5.2.41 (Bradbury *et al*. 2007). A neighbour-joining (NJ) phylogenetic tree was built with SNPrelate (Zheng *et al*. 2012) using identity-by-descent (IBS) and hierarchical clustering (functions snpgdsIBS and snpgdsHCluster), and plotted with iTOL v6 (Letunic and Bork 2021).

### Relative read coverage to synthetic AB and AS references

Trimmed reads were aligned using BWA-MEM v0.7.17, using the options −M and −k 35, to the A and B-genomes concatenated, or the A and S-genome concatenated. BAM files were sorted and duplicated reads were removed, as before. Uniquely mapped reads were obtained by excluding reads with the tags ’XA:Z:’ and ’SA:Z:’, and further filtered to retain only properly mapped paired reads (-f 0×2). BEDtools genomeCoverageBed and BEDtools map v1.7 (Quinlan and Hall 2010) were used to obtain the median read coverage or read depth for each 100 Kbp window. All 100 Kbp windows the A-genome and B-genome were aligned to each other using minimap2 v2.22 (-x asm10) to identify homologous windows (Li 2018). Because of the conserved sequences between the A- and B-genomes, a “background coverage” was observed between genomes, e.g. a low proportion of reads from AA/AAA accessions consistently aligned in the B-reference.

The relative coverage between the A-genome and B-genome was plotted using R, and the library ggplot2 (Wickham 2016). Two plots were produced in parallel to account only for the regions conserved and represent the differences in chromosome length between genomes A and B, one containing all the windows/regions in the A genome and the conserved homologous windows in the B genome below (B aligned to A), and another containing all the windows in the B genome and the conserved windows in the A genome below (A aligned to B). Coverage was normalised by dividing the window coverage by the chromosome average coverage to obtain values in the 0-2 range. The A-genome is always plotted in blue and coverage in the B-genome is always plotted in red. This analysis was completed in individual samples and on each varietal clonal group by merging the BAM files from the group’s accessions with Samtools v1.7 (Li *et al*. 2009). The same analysis was completed using the A-genome and S-genome combined.

## RESULTS

### Genotyping of the diversity panel

The average coverage (read depth) per accession was 7.7, 7.4, and 6.8X, against the A, B and S genomes, respectively. Two samples failed during sequencing, reducing the panel to 190 accessions for analysis. The average percentage of properly aligned paired reads (at the right insert length and read orientations) was 81.8, 70.9, and 76.0 % against the A, B, and S genomes, respectively. We also used a genome assembly from an ABB cultivar (587.01 Mb); where the coverage per accession was 6.0, and the average percentage of properly aligned paired reads was 80.1% (Table S3).

SNVs were called against the A-genome and B-genome, as these are the main donor genomes in cultivated bananas, for population analysis. 39,518,945 variants were obtained against the A-genome reference, of which 35,038,465 were SNPs. 33,401,534 of these SNPs were biallelic. The equivalent metrics of the B-genome were 34,104,907 variants, 29,625,724 SNPs, of which 28,161,490 were biallelic. After filtering, we obtained 187,133 and 220,451 SNP loci against the A-genome and B-genome, respectively. These datasets were used for coverage, principal component, and phylogenetic analyses. These datasets were further thinned by physical distance (50bp) and allele frequency (1 %) into 35,246 and 42,745 SNP sites, respectively, for admixture (genetic ancestry) analysis.

### Delimitation of varietal clonal clusters using admixture analysis

A set of 35,246 SNP loci called against the A-genome was used to place 151 accessions into 13 genetic clusters based on five genetically distinct ancestral founders (K=5) using STRUCTURE analysis (Fig. 1). We ran the analysis for 2 to 10 distinct sources (K values) based on an estimation of recognised genetic groups. The Evanno ΔK method indicated that the most likely value of K was 5 (Fig. S1), after which no further meaningful genetic clusters were detected (Fig. S2). The STRUCTURE analysis was repeated using a set of 42,745 SNP loci called against the B-genome. Again, the Evanno ΔK method indicated that the most likely value of K was 5 (Fig. S1), and again no further varietal groups were resolved until K = 6 (Fig. S3). The partitioning between the Cavendish and Gros Michel AAA clusters was only observed against the A-genome reference (Fig. 1), while the separation between the Bluggoe ABB and Pelipita ABB clusters was more clearly obtained against the B-genome (Fig. S4).

**Figure 1:**
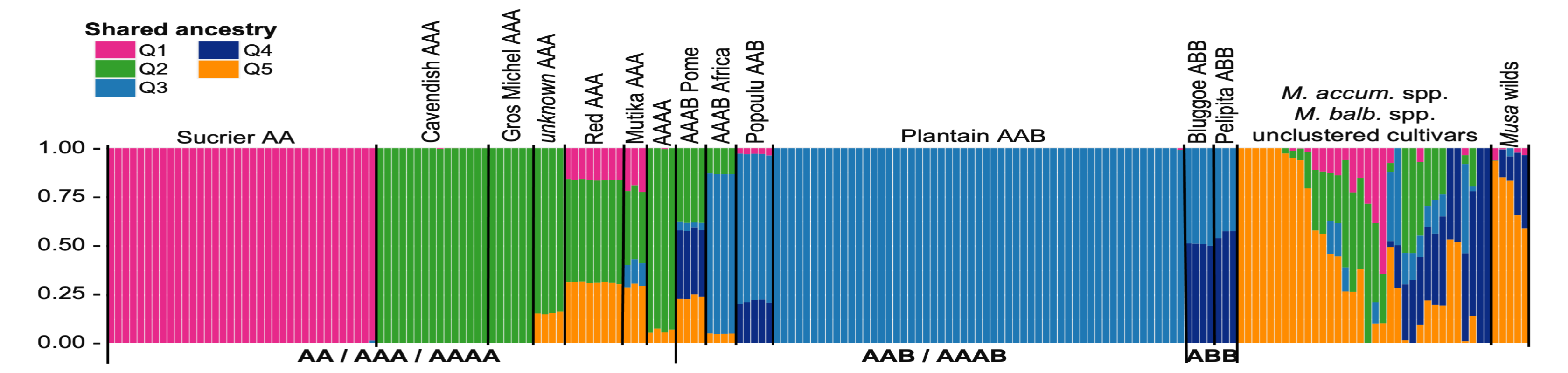
Admixture analysis of the genetic ancestry inferred in the complete set of 190 *Musa* accessions. Each accession is represented by a stacked column partitioned by the proportion of the ancestral genetic component, where each identified ancestral genetic component is represented by a different colour. Genetic composition was used to assign accessions to groups.

The 13 genetic clusters obtained were later labelled using the passport information registered in MGIS (Ruas *et al*. 2017) for the accessions in the AGROSAVIA genebank (COL004), which enabled all the genetic clusters to be associated with a variety, except for one cluster that was labelled “unknown AAA” (Table 1). Ten genetic clusters corresponded to clonal clusters, i.e. somatic clones from a lineage established in the same event. The remaining three genetic clusters were synthetic tetraploids generated in breeding crosses.

**TABLE 1.** Summary of the 13 genetic clusters identified in the diversity panel studied.

| Cluster name | Subgroup | Composition | Accessions |
| --- | --- | --- | --- |
| Sucrier AA | Sucrier | AA | 36 |
| Cavendish AAA | Cavendish | AAA | 15 |
| Gros Michel<br>AAA | Gros Michel | AAA | 6 |
| *Unknown* AAA | - | AAA | 4 |
| Red AAA | Red | AAA | 8 |
| Mutika AAA | Mutika/Lujugira | AAA | 3 |
| AAAA | Breeding<br>material | AAAA | 4 |
| AAAB Pome | Breeding<br>material | AAAB | 4 |
| AAAB Africa | Breeding<br>material | AAAB | 4 |
| Popoulu AAB | Maia<br>Maoli/Popoulu | AAB | 5 |
| Plantain ABB | Plantain | AAB | 55 |
| Bluggoe ABB | Bluggoe | ABB | 4 |
| Pelipita ABB | Pelipita | ABB | 3 |
| Wild relatives | - | - | 5 |
| Unclustered* | - | - | 34 |

We did not place 39 accessions into clusters, either single representatives of *M. acuminata* or *M. balbisiana* subspecies, or cultivars from varieties poorly represented in our panel, as we required at least three samples to establish a cluster. Among these 39, five were annotated as wild bananas in MGIS (Table S1).

The five identified ancestry sources were named Q1 to Q5, corresponding to phylogenetically distinct ancestries (Fig. 1). The clusters “Cavendish AAA”, “GrosMichel AAA”, and “unknown AAA” shared A-genome ancestry (Q2). The A-genome donor in “Sucrier AA” was distinguishable from the previous (Q1). Based on the ancestry and passport associated with the unclustered *M. accuminata* and *M. balbisiana* accessions, Q5 is a third A-genome ancestry contributing to the “Red AAA”, “unknown AAA”, and “Mutika AAA” clusters but absent in the other AA/AAA groups. The B-genome ancestry named Q4 was the major component in the ABB clusters, namely “Bluggoe ABB” and “Pelipita ABB”, but was fully absent in plantains (AAB), since the cluster “Plantain AAB” evidenced a homogenous and independent B-genome origin (Q3). This genetic cluster included the African-origin Plantain accessions (De Langhe, 2015) in our study. The cluster “Popoulu AAB” (with the “pacific plantains”) was an admixture of both Q4 and Q3. Remarkably, the “Mutika AAA” cluster evidenced a minor presence of Q3 ancestry, and it was generally highly admixed from four of the ancestries. The synthetic tetraploid clusters directly evidenced the genetic composition of their contemporary parental crosses reported in MGIS.

### Relation among the genetic clusters using principal component and phylogenetic analysis

Both PCA (Fig. 2) and phylogenetic tree (Fig. 3) showed that the genetic clusters were generally placed together based on the number of B genomes, i.e. AA/AAA/AAAA, then AAB clusters, and finally ABB clusters were more distant.

**Figure 2:**
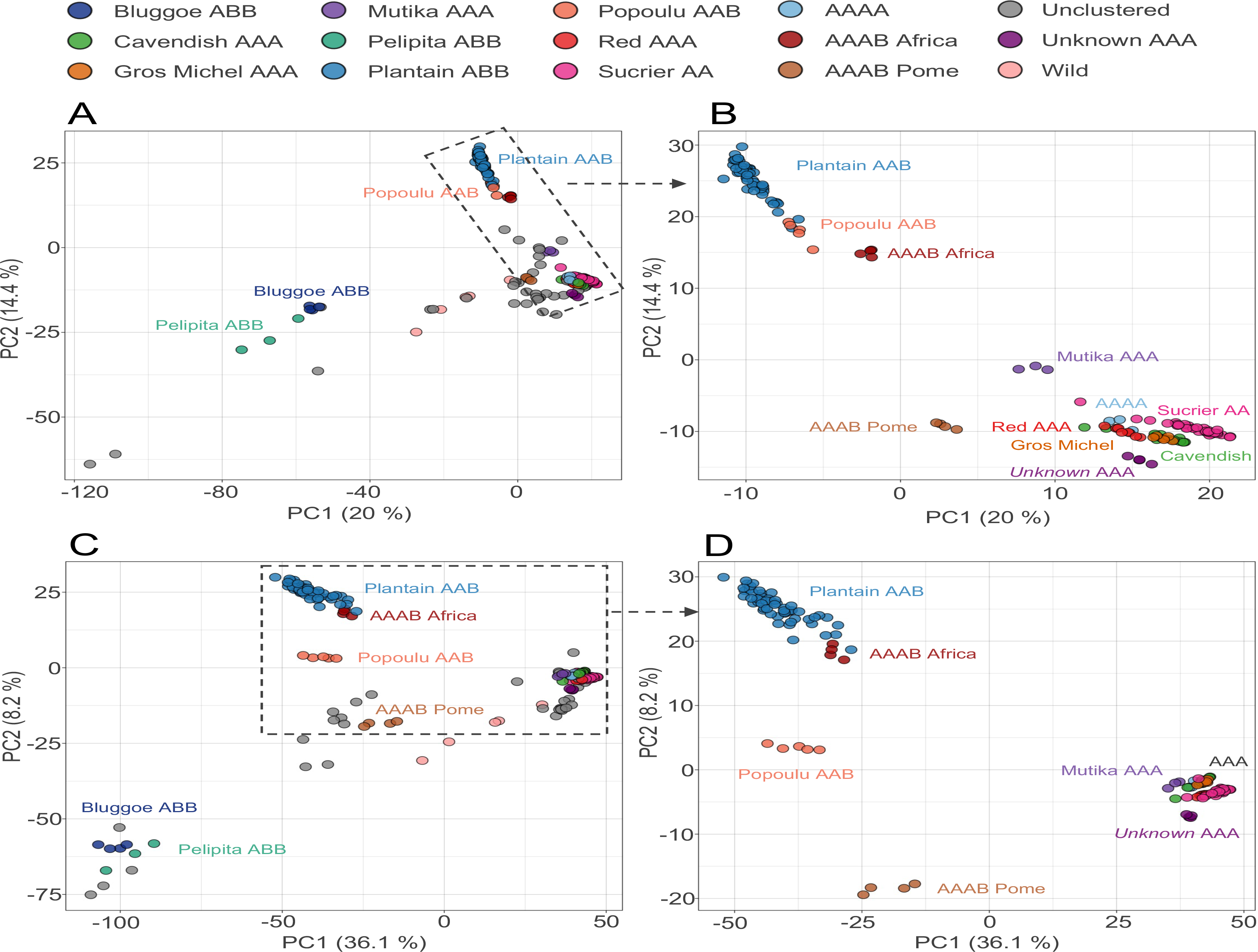
Population structure by Principal Component Analysis (PCA) using the top two principal components to separate the 188 accessions, which were coloured by genetic clusters. (A & B) SNPs called against the A-genome. (C & D) SNPs called against the B-genome. (B & D) Expanded PCA for the 142 accessions in the 11 clusters excluding the ABB groups.

**Figure 3:**
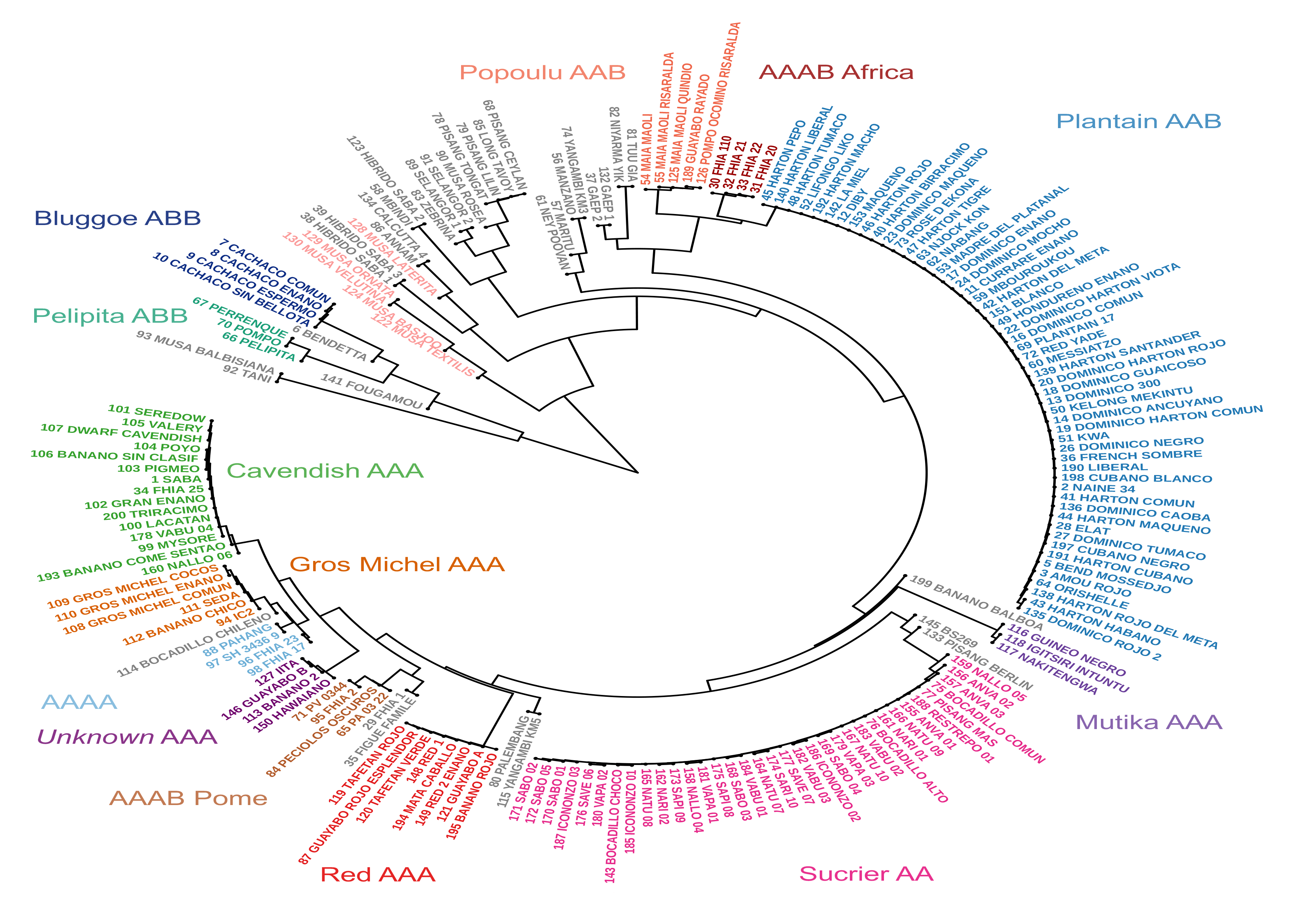
Phylogenetic tree using hierarchical clustering of the complete set of 190 accessions separated the accessions in similar divisions to the PC and admixture analyses and evidence groups of somatic clones.

“Cavendish AAA”, “Red AAA”, “Gros Michel AAA”, “Sucrier AA” and the tetraploid “AAAA” genetic clusters, with no B genome, were placed closely (Fig. 2 and 3). In the phylogenetic tree, “Sucrier AA” was in a different branch than the “AAA/AAAA” accessions. “Mutika AAA” was distant from the other “AAA/AA” in both PC and phylogenetic analyses. While the “AAAB Pome” cluster was clearly separated in the PCA, it was embedded with the AAA accessions in the phylogenetic tree. The “Popoulu AAB” and “Plantain AAB” overlapped when analysed using the A-genome as reference (Fig. 2B), but could be separated when using the B-genome as reference (Fig. 2D). While the cluster of synthetic tetraploids “AAAB Africa” overlapped with its B-progenitors, the other cluster of synthetic tetraploids “AAAB Pome” formed a well-separated group from any other in the B-genome reference, because of its different B-donor ancestor (Fig. 2C, 2D). Two unclustered accessions, “FHIA 1” (sample 29) and the Pome cultivar “Figue Famile” (sample 35) were placed close to the “AAAB Pome” cluster (Fig. 2C).

### Inferring subgenome composition based on alignment metrics

The RAA was calculated for each sample and plotted by genetic cluster (Fig. 4). For comparison, the percentage of aligned paired-reads per sample and reference before RAA normalisation are shown in Fig. S5, and decomposed by genome reference in Fig. S6.

**Figure 4:**
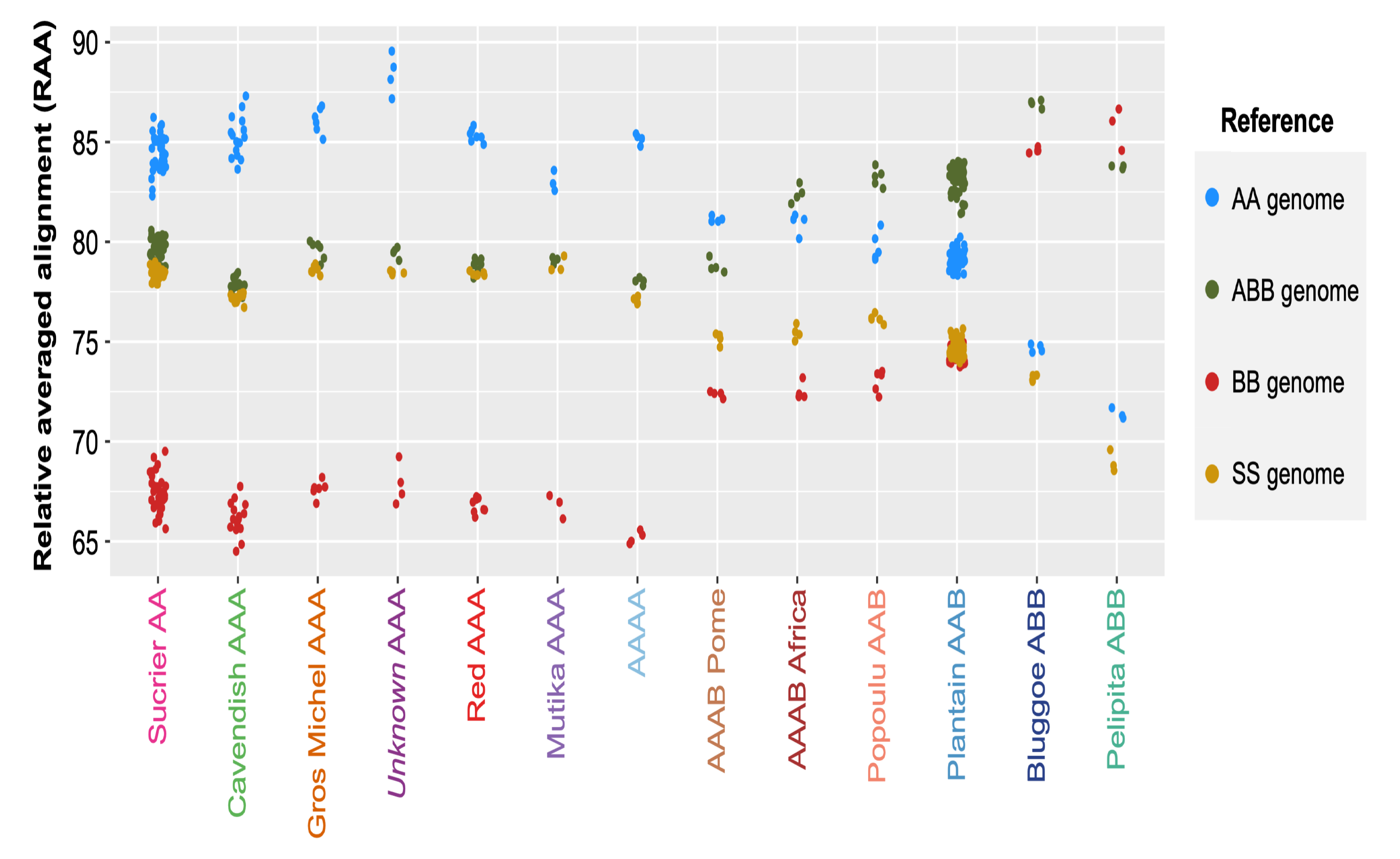
Relative averaged alignment (RAA) metric per accession (dots) grouped in 13 clusters against four reference genomes (colours) representing the donor A, B and S contributors to cultivated banana, plus a haplotype-resolved long-read assembly of a hybrid ABB accession.

The AA/AAA/AAAA accessions showed (Fig. 4) the highest RAA to the A-genome (blue dots) and a significantly lower RAA to the B-genome (red dots), while RAA to the ABB-genome (green dots) and the S-genome (yellow dots) were similar. The RAA to the A-genome was higher in the “unknown AAA” cluster than in the other AA/AAA/AAAA accessions, and lower in the “Mutika AAA”. This supports a higher contribution of *M. acuminata* subsp. *malaccensis* (A-genome reference) to the “unknown AAA” accessions. The RAA decreased for the A-genome and increased for the B-genome for accessions with a B-genome contribution (AAB, AAAB, ABB). The “Plantain AAB” had a higher RAA to the B-genome than the “Popoulu AAB”. The “Bluggoe ABB” accessions had the highest alignment rate to the ABB-genome and not the B-genome since the former was generated from an accession from the ‘Bluggoe’ subgroup. “Pelipita ABB” were the only cluster with the highest RAA to the B-genome.

The RAA to each of the reference genomes was also calculated for each of the individual 11 chromosomes (Fig. 5) to clarify the contribution from each A, B and S-genome donor. We decided not to normalise RAA by chromosome lengths, as results could be plotted and interpreted together without adding an extra transformation. Most chromosomes in a genetic cluster showed similar RAA values, evidenced by similar patterns in figure 5. Deviating from the general pattern, chromosome 2 in “Sucrier AA”, “Red AAA”, “Popoulu AAB”, and “Mutika AAA” showed the highest alignment rate to the S-genome (instead of to the A-genome); chromosome 7 in “Mutika AAA” and “Popoulu AAB” showed a notably higher RAA to the S-genome than to the A-genome; and chromosome 7 in “Plantain AAB”, “Bluggoe ABB” and “Pelipita ABB” showed the highest RAA to the B-genome. RAA values between “AAAB Pome” and “AAAB Africa” were very similar except in chromosome 7 (Fig 5). Notably, the clusters “unknown AAA” and “GrosMichel AAA” did not differ (Fig. 5).

**Figure 5:**
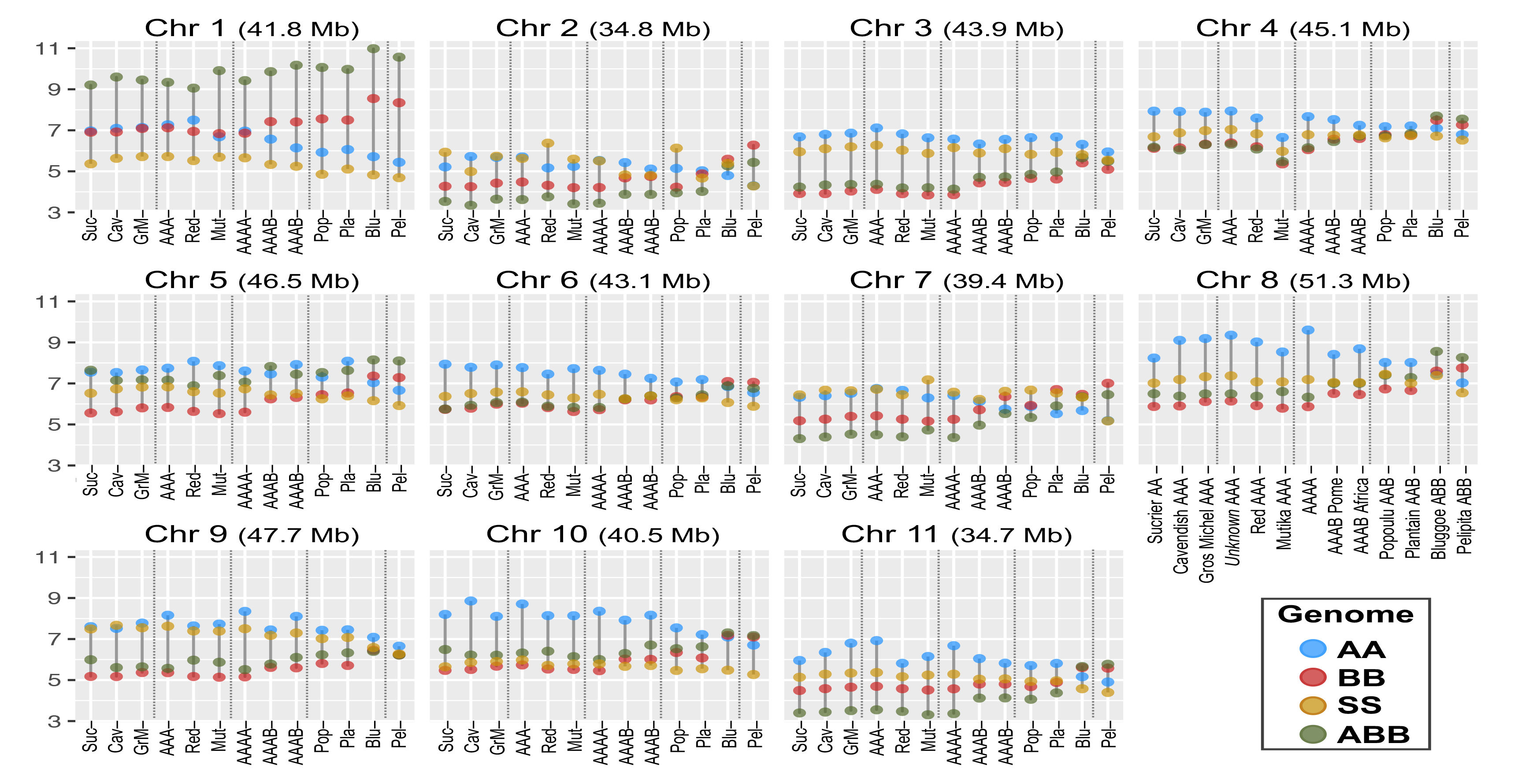
Relative averaged alignment (RAA) metric per accession (dots) grouped in 13 clusters against each of the 11 base chromosomes of the four reference genomes (colours) representing the donor A, B and S contributors to cultivated banana, plus a haplotype-resolved long-read assembly of a hybrid ABB accession.

The previous RAA metrics were also calculated in each individual accession to confirm somatic clones. These multiple plots are available in additional file 2. RAA values to each reference genome were similar among individual accessions in a genetic cluster, as expected among somatic clones. Exceptions were sample 142 (accession “LAMIEL”) within the “Plantain AAB” cluster, particularly in chromosome 4, and sample 66 (accession “PELIPITA”) in the “Pelipita ABB” cluster, particularly in chromosome 9. While the tetraploid genetic clusters (AAAA/AAAB) were not clonal, RAA values among accessions within these clusters were also very close to each others.

### Contribution from the A and B genomes along chromosomes

Using simultaneous alignments to the A and B-genome references, we could compare changes in read depth between A and B references (named “relative coverage”) and estimate the changes in donor contribution along chromosomal regions.

The relative coverage for the AAB and ABB genetic clusters are shown in Fig. 6 and additional file 3. The “Plantain AAB” cluster, which includes the Africa-origin plantains (De Langhe, 2015), evidenced a 2:1 proportion between the A and B genomes (AAB) in all chromosomes except in chromosome 7 (Fig. 6A). The proportion in chromosome 7 was 1:2 (ABB) along the whole chromosome. In addition, several A-donor introgressions, evidenced by 3:0 ratios (AAA), were observed in chromosomes 4, 6, 8, 9 and 10 (highlighted in boxes in figure 6A).

**Figure 6:**
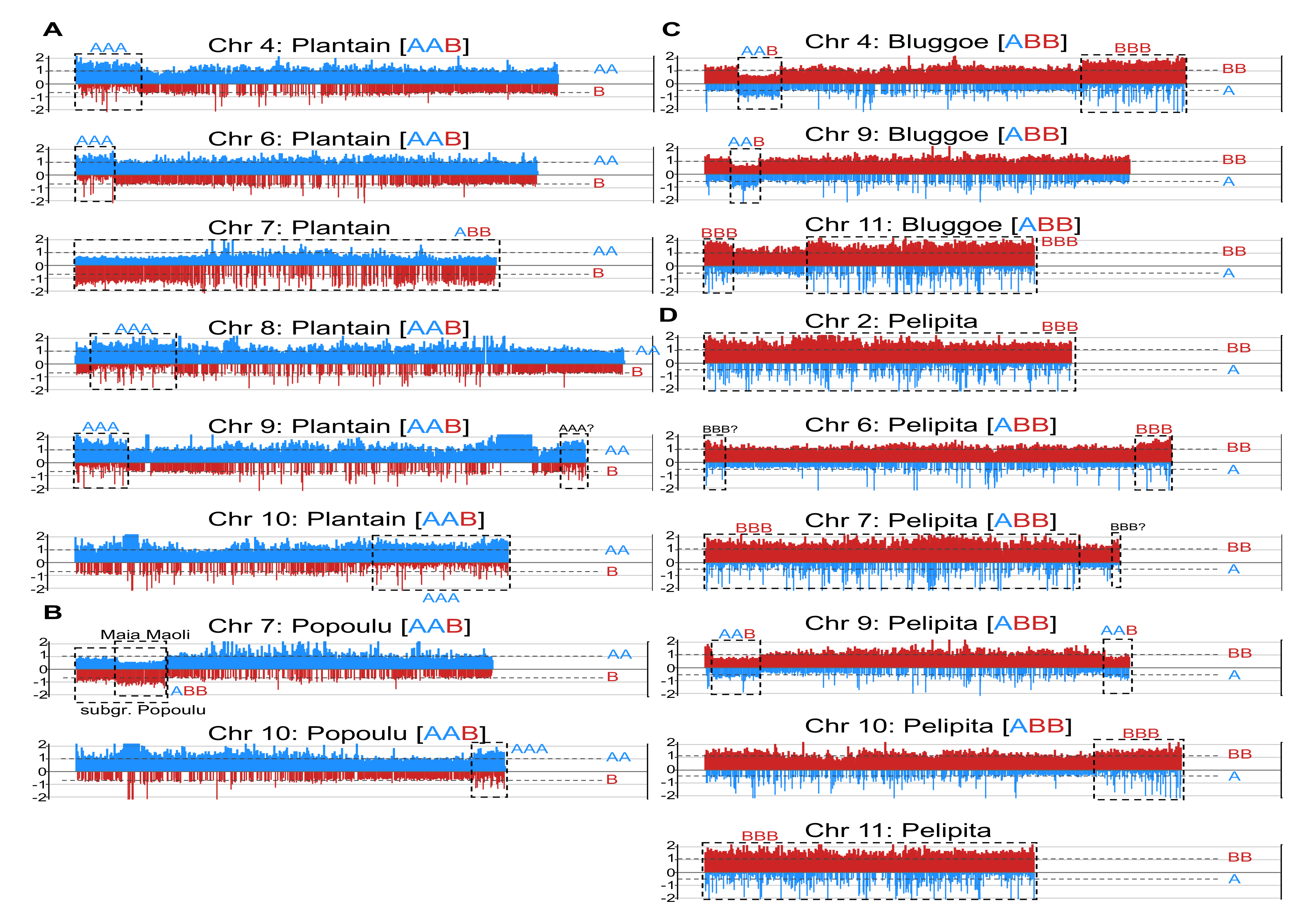
Introgression (boxes) identified in the four AAB and ABB clusters (A-D) based on relative read coverage or depth to the A and B-genome references. The A-genome is always plotted in blue and coverage in the B-genome is always plotted in red. Two plots were produced in parallel to account only for the regions conserved, either containing all the windows/regions in the A genome and the conserved homologous windows in the B genome below (blue on top of red) or the inverse (red on top of blue). Coverage was normalised by dividing the window coverage by the chromosome average coverage to obtain values in the 0-2 range. Because of the conservation between the A and B- genomes, a “background coverage” was always observed between genomes.

The “Popoulu AAB” cluster (Fig. 6B), which included the Pacific-origin plantains (De Langhe, 2015), had a 2:1 (AAB) proportion along all chromosomes, except for an A-donor introgression at the end of chromosome 10, evidenced by a 3:0 coverage ratio (AAA), and a B-donor introgression at the start of chromosome 7, evidenced by a 1:2 coverage ratio (ABB). This B-donor introgression at the start of chromosome 7 started at the beginning of chromosome 7 in accessions 126 and 189, but was shorter and started 1/8th within the chromosome 7 in the three “Maia maoli” accessions in the genetic cluster (accessions 54, 55 and 125). These accessions are individually plotted in figure S7.

The “Bluggoe ABB” cluster (Fig. 6C) had a 1:2 ratio (ABB) along most chromosomes, except for five introgressions: three B-donor introgressions, evidenced by a coverage ration of 0:3 (BBB), in chromosomes 4 and 11 (x2); and two A-donor introgressions, evidenced by a 2:1 coverage ratio (AAB), in chromosomes 4 and 9.

The “Pelipita ABB” cluster had the expected 1:2 (ABB) coverage ratio, except the complete chromosomes 2, 7 and 11 showed a 0:3 (BBB) coverage ratio (Fig. 6D). These full chromosomal rearrangements were identified in all three accessions in the cluster. In addition, two A-donor introgressions, evidenced by 2:1 coverage ratio (AAB), were noticed in chromosome 9, and two B-donor introgression in chromosomes 6 and 10 (Fig. 6D).

The relative coverage for the synthetic tetraploids were consistent with their progenitors’ composition, as shown in figure S8 (AAAB clusters) and Fig. S9 (AAAA cluster). All 11 chromosomes in “AAAB Pome” had a coverage ratio consistent with an AAAB composition, i.e. there was no evidence for introgressions. However, chromosome 7 in the “AAAB Africa” cluster showed a 2:2 ratio (AABB), which was not observed in the “AAAB Pome” cluster (Fig. S8). This is similar to the introgression observed in the “Plantain AAB” cluster, indicating the ancestral origin via parental contribution of the observed introgression.

The relative coverage for the AA/AAA genetic clusters did not evidence B-donor introgressions, as shown in additional file 4. We also generated plots for the individual accessions to identify any rearrangements specific to one individual (additional file 5).

### Contribution of A and S genome along chromosomes

Simultaneous alignment to the A and S genomes revealed an introgression from the S-genome on the first half of chromosome 2 in the “Red AAA”, “Popoulu AAB”, “Bluggoe ABB” and “Sucrier AA” genetic clusters (Fig. 7). No further intergenomic recombinations were found in any other chromosome or genetic cluster (Additional file 6). The coverage and length of this introgression in chromosome 2 were not similar among clusters, but overall, the result suggests these genetic clusters originated from closely related A-donor ancestors.

**Figure 7:**
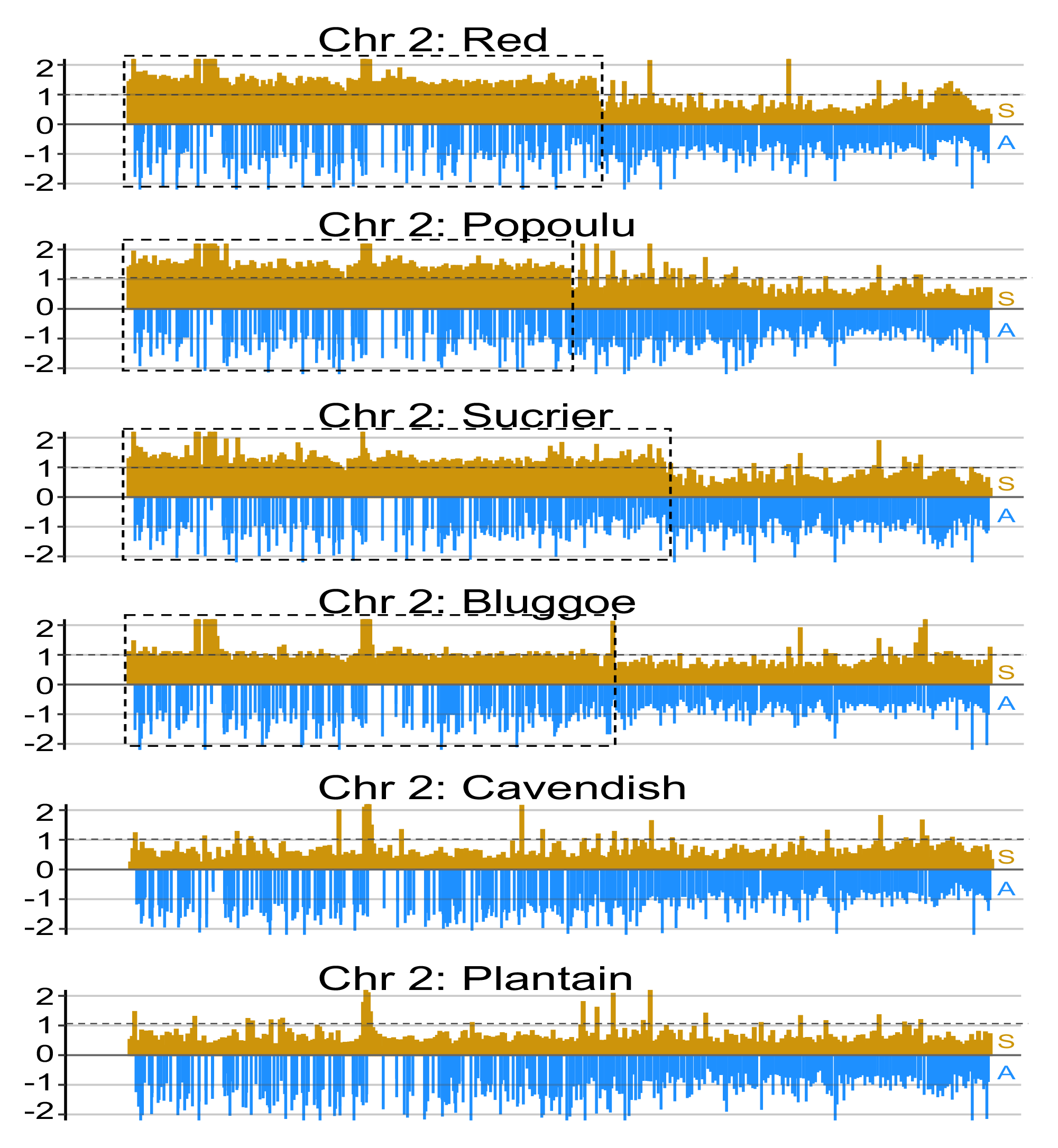
Introgression (boxes) in chromosome 2 in four varieties with different compositions based on relative read coverage or depth to the A and S genome references (blue and yellow colours, respectively). Two plots were produced in parallel containing all the windows/regions in the S genome and the conserved homologous windows in the A genome below (yellow on top of blue). Coverage was normalised by dividing the window coverage by the chromosome average coverage to obtain values in the 0-2 range.

In addition, we calculated the average read depth (coverage) between A and S for each chromosome (Table S4). The “Mutika AAA” cluster had the lowest A/S coverage ratio over all the chromosomes compared to the other clusters, except in chromosome 2, where the four clusters containing the introgression had the lowest ratio.

## DISCUSSION

To verify the extent of the varietal groups in our diversity panel, our initial analyses established the relation between banana accessions, which showed phenotypic diversity before genotyping analysis. Banana crops are vegetatively propagated, so their diversity is essentially fixed over time, and the genotypes in a varietal group putatively share a common origin and are somatic clones of each other (Heslop-Harrison and Schwarzacher 2007). Since the genetic diversity of the varietal groups is fixed over time, the phylogeny observed between the varietal clusters reflects the genetic distance between the donor ancestors that established each clonal lineage. Nevertheless, it was unsurprising that we found a genetic variation despite the absence of sexual reproduction: millennia of diversification of wild genotypes and human selection of hybrids have led to hundreds of edible banana varieties (Christelová *et al*. 2017).

We later characterised subgenome composition and identified introgressions in each varietal clonal cluster and individual accession. We particularly highlighted variety-specific chromosomal exchanges and structural variation because these results clarify shared foundational events between lineages and constitute a source of allelic diversity (e.g., in resistant alleles). Our results on composition and sizeable structural variation among lineages add to the growing evidence (Baurens *et al*. 2019; Cenci *et al*. 2021) that most clonal lineages are likely the product of several hybridisation and backcrossing events.

### Defining clonal varietal groups in a panel representative of the global banana diversity

We genotyped most of the accessions held by AGROSAVIA in Palmira, Colombia, using short-read whole-genome sequencing. This is a comprehensive collection of introduced varieties for cropping, containing a good range of cultivars, subspecies, and wild relatives with observed phenotypic diversity. However, our population structure analysis using principal component, phylogenetic and admixture analysis evidenced extensive redundancy among accessions.

We divided 151 accessions into 13 genetic clusters, containing between three and 55 accessions, based on five genetically distinct ancestral founders using STRUCTURE analysis (Fig. 1). The “admixture model” of STRUCTURE assumes that each individual has a genetic make-up from one or more of K distinct ancestral donor sources. In clonally propagated species, such as bananas, the K distinct sources reflect the shared ancestry, i.e. the relationship among the donors involved in establishing each clonal lineage. 39 of the 190 accessions were unclustered because they were either wild (5 accessions) or members of a clonal lineage poorly represented in our diversity panel (34 accessions).

Passport information was used to label each genetic cluster to a specific cultivar variety and define *clonal varietal groups*. We could not link one genetic cluster to a banana variety, because the passport data was inconclusive, and therefore we referred to it as “unknown AAA” in downstream analysis. Three clusters corresponded with the synthetic tetraploids from breeding programmes, which grouped based on the composition of the paternal varieties that originated them. Common cultivars were still absent in our panel (e.g. Ambon, Rio, Orotava, etc). Since accessions held in genebanks other than the International Musa Germplasm Transit Centre (ITC) have not been genotyped (Rouard *et al*. 2021), we verified or corrected the passport information based on the genetic analysis in Table S1.

### Alignment metrics to identify subgenome composition

We established a new method to quantify the relative alignment from each accession to three reference banana genomes, which are representative of the A, B and S genome donors. We called this normalised alignment metric “Relative averaged alignment” (RAA) The RAA accounts for the technical variation between samples and reference bias, ie. The phylogenetic distance between a variety and a genome reference. The RAA allowed us to identify subgenome composition (AA, AB, AAA, AAB, ABB, etc.) and compare with the data available in MGIS (Ruas *et al*. 2017).

Later, we used comparisons of read depth (coverage) to identify introgressions. Compared to the private SNPs approach previously used for chromosome painting (Baurens *et al*. 2019; Cenci *et al*. 2021), read depth comparisons can be used in homologous regions between the A and B genomes, and do not require inferring SNPs exclusive for a whole species (private SNPs) from a limited number of available assemblies, which can often lead to wrong assumptions.

### Lineage-exclusive introgressions in the AAB varietal groups

Traditional starchy bananas are dominated by two geographically discrete groups of AAB cultivars (De Langhe 2015); plantains from the rainforest zone of Africa and the Maoli-Popo’ulu subgroup of plantains. The Popoulu subgroup shows maximum diversity in Melanesia and is often called Pacific Plantains. In our study, these groups correspond to the named “Plantain AAB” and “Popoulu AAB” clusters, comprising 55 and 5 accessions, respectively. Both subgroups would have their origin in Indonesia and Philippines, and would share *M. acuminata* subsp*. banksii* as A-genome donor (Perrier 2009; Perrier *et al*. 2011).

The main different we observed between these two AAB plantain clusters was a B-donor (ABB composition) exchange of the complete chromosome 7 in “Plantain AAB” that was not present in “Popoulu AAB” (Fig 6). This ABB composition in chromosome 7 in “Plantain AAB” had been previously described in a single sample, “French Clair”, representative of the French division of African Plantains (Baurens *et al*. 2019).

We also observed a B-donor introgression (ABB) at the start of chromosome 7 in “Popoulu AAB” not previously described (Fig 6). This B-donor introgression (ABB) at the start of chromosome 7 was shorter in the three “Maia Maoli” accessions (samples 54, 55 and 125) compared to the other two accessions in the cluster (Fig. S7). The Maoli-Popo’ulu subgroup of plantains can be divided into “Maoli” and a “Popoulu” subdivisions based on the shape of the fruit (Ploetz 2007). We propose that this B-donor introgression (ABB) at the start of chromosome 7 is exclusive of the Maoli-Popo’ulu subgroup and different between the two subdivisions. We hypothesise that the accessions in the “Maia Maoli” subdivision lost the first half of the introgression in a subsequent backcrossing event.

In addition, we observed five A-donor telomeric introgressions (AAA) in African (“Plantain AAB”) that were not present in Pacific (“Popoulu ABB”) plantains. These AAA introgressions were at the start of chromosomes 4, 6, 8, and 9; and at the end of chromosomes 9 and 10 (Fig. 6). All these regions showed an AAA composition instead of AAB (Fig 6). The genome proportion plots by individual sample and chromosome (Additional File 2) evidence these introgressions are similar in all “Plantain AAB” except for cultivar Lamiel (Sample 142), which notably showed a B-donor introgression (ABB) in the second half of chromosome 4 that was not observed in any other plantain accession. Variation in a single accession was uncommon (Additional File 2). We hypothesise that genetic diversity observed in a single accession could be associated with somaclonal variation associated with *in vitro* propagation of germplasm (Simonikova *et al*. 2019, 2022; Van den houwe *et al*. 2020).

### Full chromosome exchanges and introgressions in the ABB varietal groups

The introgressions in the ABB varietal groups, the “Bluggoe ABB” and “Pelipita ABB” clusters in our study, have already been described (Cenci *et al*. 2021). Differences between these two ABB groups can be confirmed in chromosomes 2 and 7 based on relative alignments (Fig. 5), but not chromosome 11 because B-donor exchanges were present in both groups. We confirmed homologous exchanges in all “Pelipita ABB” accessions resulting in the entire chromosomes 2 and 11, and most of chromosome 7, all resulting in BBB composition instead of ABB (Fig 6). We also observed two A-donor introgressions (AAB) in chromosome 9, and three B-donor introgressions (BBB) in chromosomes 6 and 9. All these introgressions were noticed in all “Pelipita ABB” accessions (Fig 6) and previously reported by Baurens et al. (2019). A B-donor introgression at the end of chromosome 7 (BBB) in a Pelipita cultivar has been described (Baurens *et al*. 2019), which we did not observe since the second-arm interstitial region in chromosome 7 showed the expected ABB composition. Notably, three ABB accessions did not cluster with the others on the PCA analysis (Fig. 2) and our sample 66 showed a slightly different alignment pattern on chromosomes 9 and 10 (additional file 2). However, read coverage was not sufficient to confirm recombination in the individual accessions.

Further studies will need to clarify any further variation in outlying “Pelipita ABB” accessions. In “Bluggoe ABB”, we observed two A-donor introgressions (AAB) in chromosomes 4 and 7, and three B-donor introgressions (BBB) in chromosomes 4 and 11. These introgressions were previously reported (Baurens *et al*. 2019). While the entire chromosome 11 had a BBB composition in “Pelipita ABB”, it did not include the complete chromosome 11 in the “Bluggoe ABB” cluster, which supports a different origin or a later backcrossing event in “Bluggoe ABB”.

### Relationship between the synthetic tetraploid groups

Three genetic clusters were synthetic tetraploid hybrids based on the MGIS passport data and our observations of alignment and coverage properties, one with AAAA composition and two with AAAB composition. We were able to find the parentage of some of these accessions (Table S5). The AAAA genetic cluster grouped with the AAA Cavendish and Gros Michel triploid groups, which agrees with the parentage of sample 98 (FHIA17).

The “AAAB Pome” tetraploid cluster was named after the parental contributors of the B genome, which is confirmed in the case of the subgr. Pome plantain “Prata ana” in sample 65. As a result of this origin, these tetraploid accessions clustered close to FHIA-1 (also with a subgr. Pome parental contributor) and the AAB subgr. Pome cultivar “Figue Famile”. Similarly, the “AAAB Africa” tetraploid cluster was named after the B-genome donor of accessions FHIA-31 and FHIA-20, namely AVP-67, a French AAB Plantain from Africa. Consequently, these accessions inherited and showed the ABB composition in chromosome 7 representative to “Plantain AAB” members (Fig. S8 and S9). All the other chromosomes showed an AAB composition.

### Relationship between of the AA/AAA varietal groups

We verified the close relation between the AA/AAA varietal groups previously described. In particular, we identified no B-genome introgression in any AA/AAA group (Fig. S7). In detail, our PCA evidenced Cavendish (AAA) and Gros Michel (AAA) overlapped (Fig. 2). The close genetic relation between Gros Michel and Cavendish has been previously evidenced by genotyping (Christelová *et al*. 2017), common Mchare donor (Raboin *et al*. 2005), and a reciprocal translocation between chromosomes 3 and 8 (Simonikova *et al*. 2019). The “Sucrier AA” and “Red AAA” clusters were close but distinguishable from the former despite Sucrier and Cavendish shared ancestry (Martin *et al*. 2020). Diploid sucrier accessions tentatively comprise a varying mosaic of *M. acuminata* subsp*. banksii, zebrina, malaccensis* and wild “Pisang Madu” banana (Martin *et al*. 2020). While we did not observe genetic variation among Sucrier accessions in our tree (Fig 3) or ancestry analysis, there were three Sucrier samples separated from the others in the PCA analysis (Fig. 2).

Generally, the Mutika cultivars (East of African highland bananas) have diversified by somatic mutation to have several end uses (Kitavi *et al*. 2016, 2020), and now harbour significant epigenetic diversity(Kitavi *et al*. 2020). However, Mutika cultivars are genetically uniform, since they likely arose from a single ancestral clone introduced from Asia into Africa that subsequently underwent population expansion by vegetative propagation (Kitavi *et al*. 2016). Mutika has been described as genetically distinct from the other triploid AAA bananas, which was also evidenced in our phylogenetic and PC analyses: Mutika cultivars distinctively contains two chromosome sets from subsp. *zebrina* and one chromosome set from subsp. *banksii* (Simonikova *et al*. 2019), and a Vε cytoplasmic type (Perrier 2009).

### Contribution of S genome to cultivated banana

*M. schizocarpa* (S genome) has contributed to cultivated banana (Heslop-Harrison and Schwarzacher 2007; D’Hont *et al*. 2012), but the nature of this contribution is not fully elucidated. *M. schizocarpa* is more closely related to *M. acuminata* than *M. balbisiana* (Li *et al*. 2010, 2013), which explains the relatively high coverage in the S-genome in all the accessions, particularly the AAA samples. However, the much-increased S-genome coverage in chromosome 2 evidenced intergenomic recombination between these subgenomes (Fig. 5), and we confirmed that S-donor introgression (Fig. 7) in four varieties: Bluggoe (ABB), Sucrier (AA), Popoulu (AAB) and Red (AAA). *M. schizocarpa* and *M. acuminata* subsp. *banksii* are sympatric in Papua New Guinea (Christelová *et al*. 2017), and AS natural hybrid cultivated bananas have been documented (Daniells 2001). These AS hybrids show a close relation with *M. acuminata* subsp. *banksia* (Carreel *et al*. 1994; Christelová *et al*. 2017). ITS analysis observed a *M. schizocarpa* S-contribution to the Mutika subgroup (Nemeckova *et al*. 2018). We did not find S-genome introgressions in the Mutika group but the alignment statistics suggested this group was less closely related to the donor of the A-genome reference, as they showed the highest RAA to the S-genome.

## Conclusions

We identified ten varietal groups composed of somatic clones, using admixture, principal component, and phylogenetic analyses, and later linked each clonal lineage to a common variety. We established a new alignment-based metric, RAA, and demonstrated it can be used in the total genome and individual chromosomes to clarify subgenome composition (AA, AAB, etc.).

We identified 20 introgressions among the AAB and ABB varieties: five A-donor and one B-donor introgressions in African plantains, one A-donor and one B-donor introgressions in Popoulu plantains, two A-donor and three B-donor introgressions in Bluggoe, and two A-donor and five B-donor introgressions in Pelipita. We confirmed some known chromosomal rearrangements for the Bluggoe and Pelipita subgroups, and showed that the main difference between the African Plantain subgroup and the Popoulu subgroup (often referred to as Pacific Plantains) was the ABB make-up of chromosome 7 in the African Plantains.

We identified variation in length in at least two introgressions, a B-donor introgression that had a different length between the “Maoli” and a “Popoulu” subdivisions, and a S-donor (*M. schizocarpa*) introgression in chromosome 2 in four varieties with different compositions (AAA, AAB, ABB, AA). This variation supports intricate founding events, which encompassed multiple instances of hybridization and subsequent residual backcrossings.

## SUPPLEMENTARY DATA

Supplementary data consist of the following:

Figure S1: Estimation of the optimal number of K using the Evanno method for K2 to K10 against the A and B genomes.

Figure S2: Admixture analysis for K4 to K8 against the A genome.

Figure S3: Admixture analysis for K4 to K8 against the B genome.

Figure S4: Comparison of the clusters identified using admixture analysis in the A vs. B genomes.

Figure S5: The unnormalised percentage of properly paired reads aligning for the 13 genetic cluster, unclustered and wild accessions for the A-genome, B-genome, ABB- genome and S-genome reference.

Figure S6: The unnormalised percentage of properly paired reads aligning for the 13 genetic cluster, unclustered and wild accessions plotted separately for the A-genome, B-genome, ABB-genome and S-genome reference.

Figure S7: Relative read coverage plots against the A and B subgenomes in chromosome 7 for the five accession in “Popoulu AAB” cluster.

Figure S8: Coverage plots showing A and B subgenome structure for the AAAB synthetic tetraploid clusters. The blue and red bars represent median coverage depth (normalised by mean) over 100,000 bp windows, for the A- and B-subgenomes, respectively.

Figure S9: Coverage plots showing A and B subgenome structure for the AAAA synthetic tetraploid clusters. The blue and red bars represent median coverage depth (normalised by mean) over 100,000 bp windows, for the A- and B-subgenomes, respectively.

Additional File 1: Constains Suplementary Tables S1-S5.

Additional File 2: RAA metrics calculated in each individual accession to confirm somatic clones.

Additional File 3: The relative coverage in the A and B subgenomes for the AAB and ABB genetic clusters.

Additional File 4: The relative coverage in the A and B subgenomes for the AA and AAA genetic clusters.

Additional File 5: The relative coverage in the A and B subgenomes for each individual accession.

Additional File 6: The relative coverage in the A and A subgenomes for all genetic clusters.

## ACKNOWLEDGEMENTS

Germplasm is held in AGROSAVIA’s collection (MGIS: COL004) and is available on request. All sequence data used in this manuscript have been deposited as study PRJEB62882 in the European Nucleotide Archive. Code from the computational analyses is available in the author’s Github repository at https://github.com/jjdevega/Structural-diversity-in-banana-cultivars. Plant accessions were obtained from AGROSAVIA’s Genebank in compliance with the national laws and international treaties. A research collaboration was developed with scientists providing the samples, and collaborators are included as co-authors, the results of the research have been disseminated among stakeholders and the broader national and international scientific community. JDV, RY, and FDP conceived and managed the project. JH, JAOG, COA and JDV completed bioinformatics analysis. JAOG, DLTM and AE selected, validated and collected the samples, and completed DNA extractions. JH and JDV wrote the manuscript with contributions from all the authors. The authors declare no conflicts of interest.

## FUNDING

JH and JDV received additional funding from the Biotechnology and Biology Sciences Research Council (BBSRC)’s Global Challenge Research Fund BB/P028098/1 and from the BBSRC Core Strategic Programme Grant (Genomes to Food Security) BB/CSP1720/1 and its constituent work package BBS/E/T/000PR9818 (WP1 Signatures of Domestication and Adaptation);

## Notes

### Competing Interest Statement

The authors have declared no competing interest.

## LITERATURE CITED

Baurens F-C, Martin G, Hervouet C, et al. 2019. Recombination and large structural variations shape interspecific edible bananas genomes. Molecular Biology and Evolution 36: 97–111.

Beier S, Himmelbach A, Colmsee C, et al. 2017. Construction of a map-based reference genome sequence for barley, *Hordeum vulgare* L. Scientific Data 4: 1–24.

Belser C, Istace B, Denis E, et al. 2018. Chromosome-scale assemblies of plant genomes using nanopore long reads and optical maps. Nature Plants 4: 879–887.

Bradbury PJ, Zhang Z, Kroon DE, Casstevens TM, Ramdoss Y, Buckler ES. 2007. TASSEL: software for association mapping of complex traits in diverse samples. Bioinformatics 23: 2633–5.

Carreel F, Fauré S, de León DG, et al. 1994. Evaluation de la diversité génétique chez les bananiers diploïdes (*Musa* sp). Genetics Selection Evolution 26.

Cenci A, Sardos J, Hueber Y, et al. 2021. Unravelling the complex story of intergenomic recombination in ABB allotriploid bananas. Annals of Botany 127: 7–20.

Christelová P, De Langhe E, Hřibová E, et al. 2017. Molecular and cytological characterization of the global *Musa* germplasm collection provides insights into the treasure of banana diversity. Biodiversity and Conservation 26: 801–824.

Danecek P, Auton A, Abecasis G, et al. 2011. The variant call format and VCFtools. Bioinformatics 27: 2156–8.

Danecek P, Bonfield JK, Liddle J, et al. 2021. Twelve years of SAMtools and BCFtools. Gigascience 10: giab008.

Daniells J; J C; Karamura, D; Tomekpe, K. 2001. Musalogue: a catalogue of *Musa* germplasm. Diversity in the genus Musa.

De Langhe E Perrier, Xavier, Donohue, Mark, Denham, Tim Paul. 2015. The Original Banana Split: Multi-disciplinary implications of the generation of African and Pacific Plantains in Island Southeast Asia. Ethnobotany Research and Applications 14: 299–312.

D’hont A, Denoeud F, Aury J-M, et al. 2012. The banana (*Musa acuminata*) genome and the evolution of monocotyledonous plants. Nature 488: 213–217.

Droc G, Lariviere D, Guignon V, et al. 2013. The banana genome hub. Database 2013.

Evanno G, Regnaut S, Goudet J. 2005. Detecting the number of clusters of individuals using the software STRUCTURE: a simulation study. Molecular Ecology 14: 2611–20.

Florez JEM, Lobo M, Arana AC, Coronado YM. 2012. Caracterización molecular de genotipos de plátano del Banco de Germoplasma de Corpoica Palmira, con uso de marcadores RAMs. Acta Agronómica 61: 28–29.

FAO. 1997. FAOSTAT statistical database. Food, Agriculture Organization of the United N. [Accessed 01/02/2023].

Francis RM. 2017. Pophelper: an R package and web app to analyse and visualize population structure. Molecular Ecology Resources 17: 27–32.

Heslop-Harrison JS, Schwarzacher T. 2007. Domestication, Genomics and the Future for Banana. Annals of Botany 100: 1073–1084.

Houwe I, Chase R, Sardos J, et al. 2020.Safeguarding and using global banana diversity: a holistic approach. CABI Agric Biosci 1.

Igwe DO, Ihearahu OC, Osano AA, Acquaah G, Ude GN. 2021. Genetic diversity and population assessment of *Musa* L.(Musaceae) employing CDDP markers. Plant Molecular Biology Reporter 39: 801–820.

Jakobsson M, Rosenberg NA. 2007. CLUMPP: a cluster matching and permutation program for dealing with label switching and multimodality in analysis of population structure. Bioinformatics 23: 1801–6.

Kitavi M, Downing T, Lorenzen J, et al. 2016. The triploid East African Highland Banana (EAHB) genepool is genetically uniform arising from a single ancestral clone that underwent population expansion by vegetative propagation. Theorical Applied Genetics 129: 547–61.

Kitavi M, Cashell R, Ferguson M, et al. 2020. Heritable epigenetic diversity for conservation and utilization of epigenetic germplasm resources of clonal East African Highland banana (EAHB) accessions. Theorical Applied Genetics 133: 2605–2625.

Krueger F. 2015. Trim galore. A wrapper tool around Cutadapt and FastQC to consistently apply quality and adapter trimming to FastQ files, 516: 517.

Letunic I, Bork P. 2021. Interactive Tree Of Life (iTOL) v5: an online tool for phylogenetic tree display and annotation. Nucleic Acids Res 49: W293–W296.

Li H. 2013. Aligning sequence reads, clone sequences and assembly contigs with BWA- MEM. arXiv preprint arXiv:1303.3997.

Li H. 2018. Minimap2: pairwise alignment for nucleotide sequences. Bioinformatics 34: 3094–3100.

Li LF, Hakkinen M, Yuan YM, Hao G, Ge XJ. 2010. Molecular phylogeny and systematics of the banana family (Musaceae) inferred from multiple nuclear and chloroplast DNA fragments, with a special reference to the genus Musa. Mol Phylogenet Evol 57: 1–10.

Li H, Handsaker B, Wysoker A, et al. 2009. The Sequence Alignment/Map format and SAMtools. Bioinformatics 25: 2078–9.

Li LF, Wang HY, Zhang C, et al. 2013. Origins and domestication of cultivated banana inferred from chloroplast and nuclear genes. PLoS One 8: e80502.

Martin G, Baurens FC, Hervouet C, et al. 2020. Chromosome reciprocal translocations have accompanied subspecies evolution in bananas. Plant J 104: 1698–1711.

McKenna A, Hanna M, Banks E, et al. 2010. The Genome Analysis Toolkit: a MapReduce framework for analyzing next-generation DNA sequencing data. Genome Research 20: 1297–1303.

Nemeckova A, Christelova P, Cizkova J, et al. 2018. Molecular and Cytogenetic Study of East African Highland Banana. Front Plant Sci 9: 1371.

Perez-Sepulveda BM, Heavens D, Pulford CV, et al. 2021. An accessible, efficient and global approach for the large-scale sequencing of bacterial genomes. Genome Biology 22: 349.

Perrier X. 2009. Combining Biological Approaches to Shed Light on the Evolution of Edible Bananas. Ethnobotany Research and Applications 7: 199–216.

Perrier X, De Langhe E, Donohue M, et al. 2011. Multidisciplinary perspectives on banana (Musa spp.) domestication. Proceedings of the National Academy of Sciences 108: 11311–11318.

Perrier X, Jenny C, Bakry F, et al. 2019. East African diploid and triploid bananas: a genetic complex transported from South-East Asia. Annals of botany 123: 19–36.

Ploetz RC Kepler, Angela Kay. 2007. Banana and plantain - an overview with emphasis on Pacific island cultivars.

Pritchard JK, Stephens M, Donnelly P. 2000. Inference of population structure using multilocus genotype data. Genetics 155: 945–59.

Quinlan AR, Hall IM. 2010. BEDTools: a flexible suite of utilities for comparing genomic features. Bioinformatics 26: 841–2.

Raboin L-M, Carreel F, Noyer J-L, et al. 2005. Diploid Ancestors of Triploid Export Banana Cultivars: Molecular Identification of 2n Restitution Gamete Donors and n Gamete Donors. Molecular Breeding 16: 333–341.

Rouard M, Sardos J, Sempéré G, et al. 2021. A digital catalog of high-density markers for banana germplasm collections. *Plants, People*, Planet 4: 61–67.

Ruas M, Guignon V, Sempere G, et al. 2017. MGIS: managing banana (Musa spp.) genetic resources information and high-throughput genotyping data. Database 2017.

Simmonds NW, Shepherd K. 1955. The taxonomy and origins of the cultivated bananas. Botanical Journal of the Linnean Society 55: 302–312.

Simonikova D, Cizkova J, Zoulova V, Christelova P, Hribova E. 2022. Advances in the Molecular Cytogenetics of Bananas, Family Musaceae. Plants (Basel) 11.

Simonikova D, Nemeckova A, Karafiatova M, et al. 2019. Chromosome Painting Facilitates Anchoring Reference Genome Sequence to Chromosomes In Situ and Integrated Karyotyping in Banana (*Musa* spp.). Front Plant Sci 10: 1503.

Van den houwe I, Chase R, Sardos J, et al. 2020.Safeguarding and using global banana diversity: a holistic approach. CABI Agriculture and Bioscience 1.

Wang Z, Miao H, Liu J, et al. 2019. *Musa balbisiana* genome reveals subgenome evolution and functional divergence. Nature Plants 5: 810–821.

Wickham H. 2016. ggplot2: elegant graphics for data analysis. Springer.

Zheng X, Levine D, Shen J, Gogarten SM, Laurie C, Weir BS. 2012. A high-performance computing toolset for relatedness and principal component analysis of SNP data. Bioinformatics 28: 3326–3328.

