## Supplementary figures and images for "Characterising genome composition and large structural variation in banana varietal groups"

### 2NAINE34cov_byA.pdf

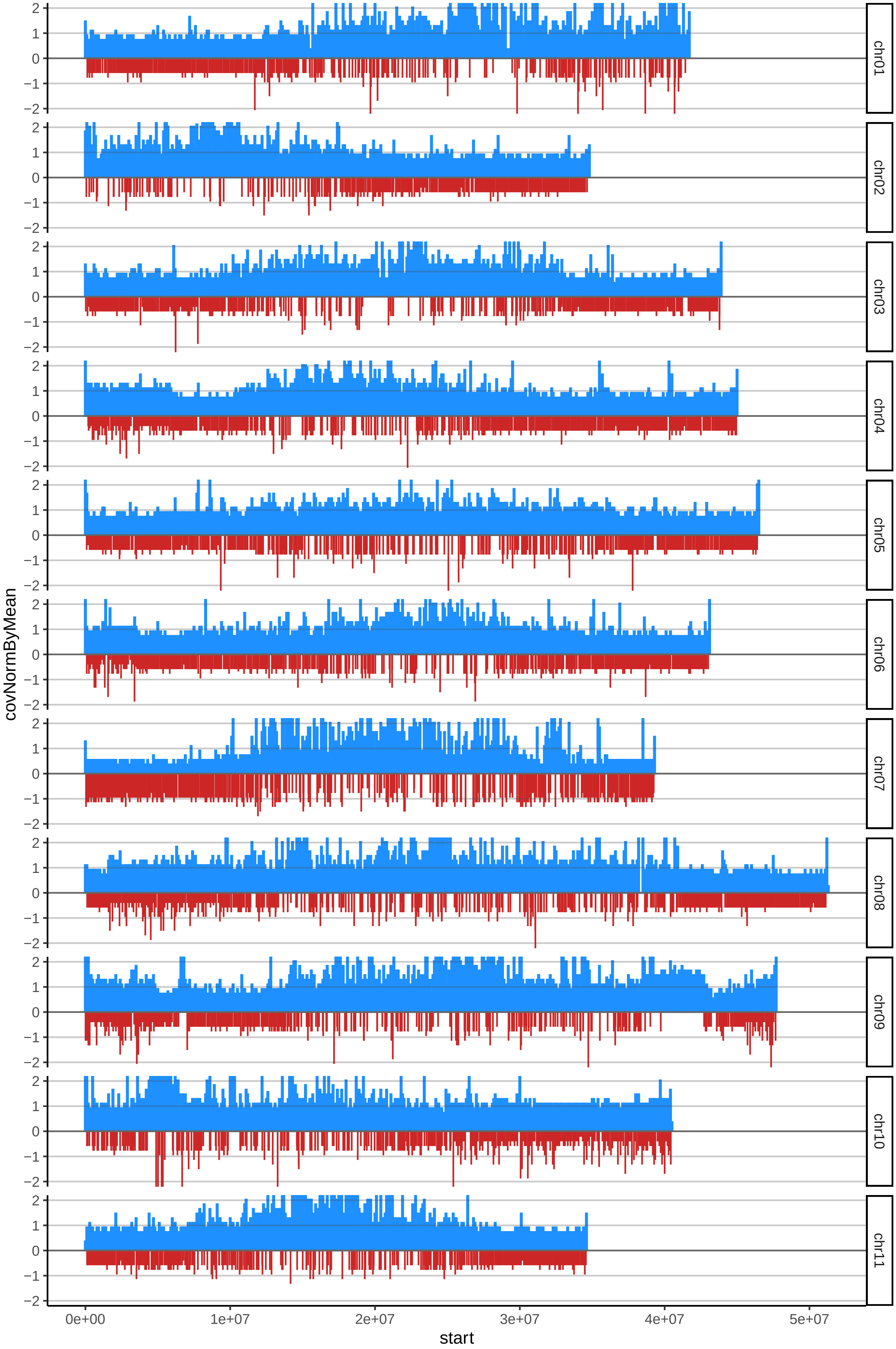

### 2NAINE34cov_byB.pdf

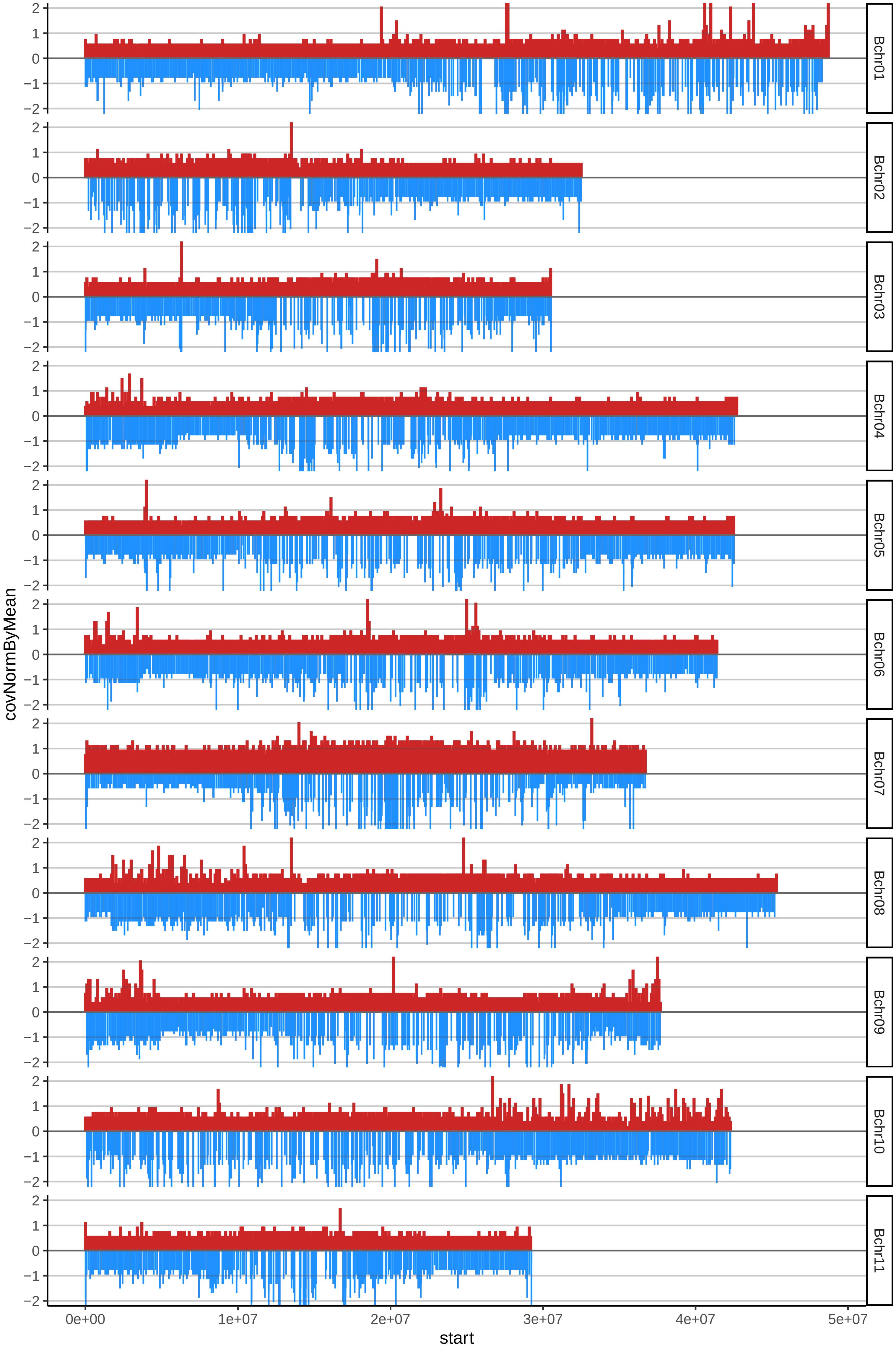

### 3AMOUROJOcov_byA.pdf

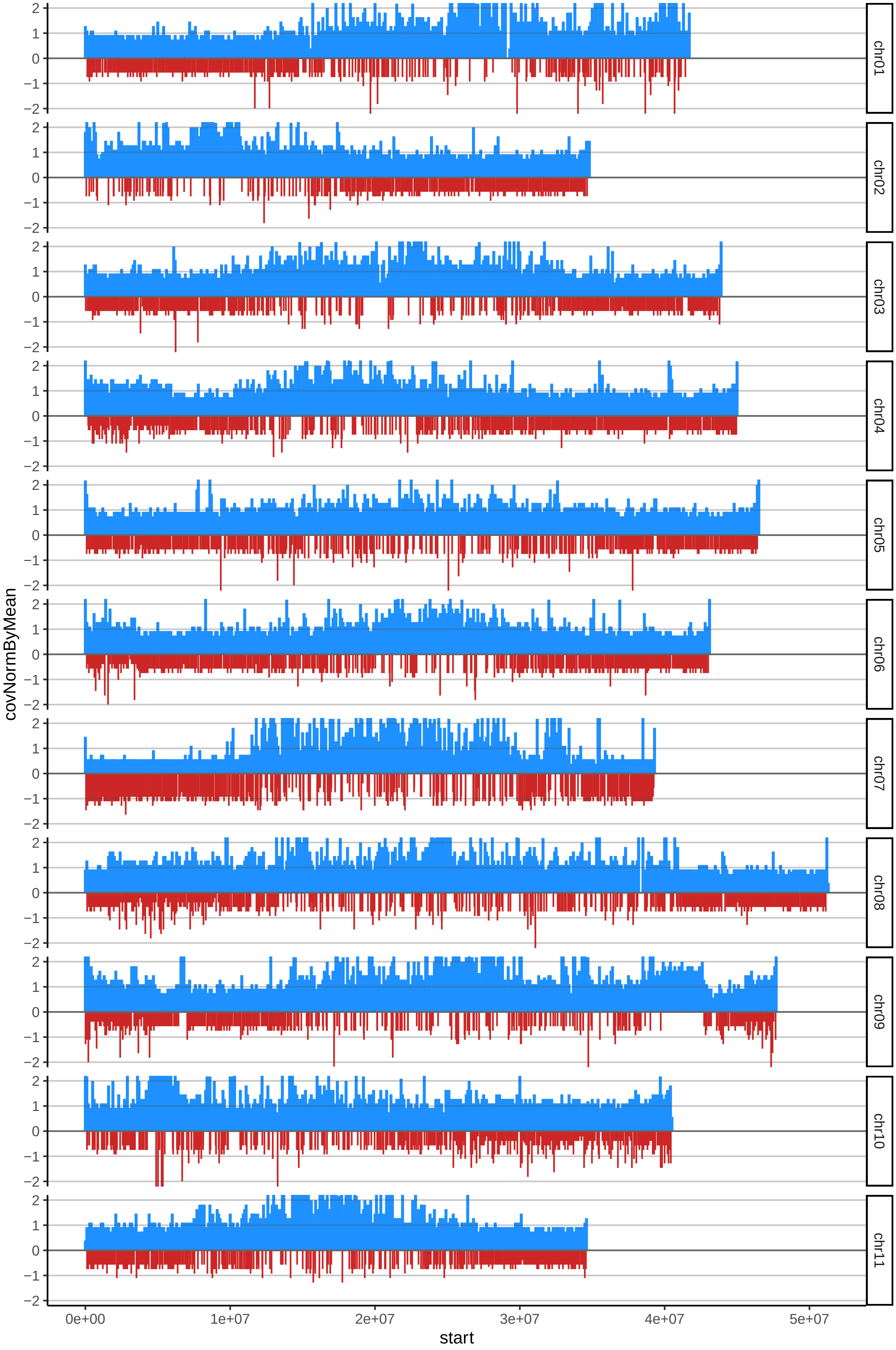

### 3AMOUROJOcov_byB.pdf

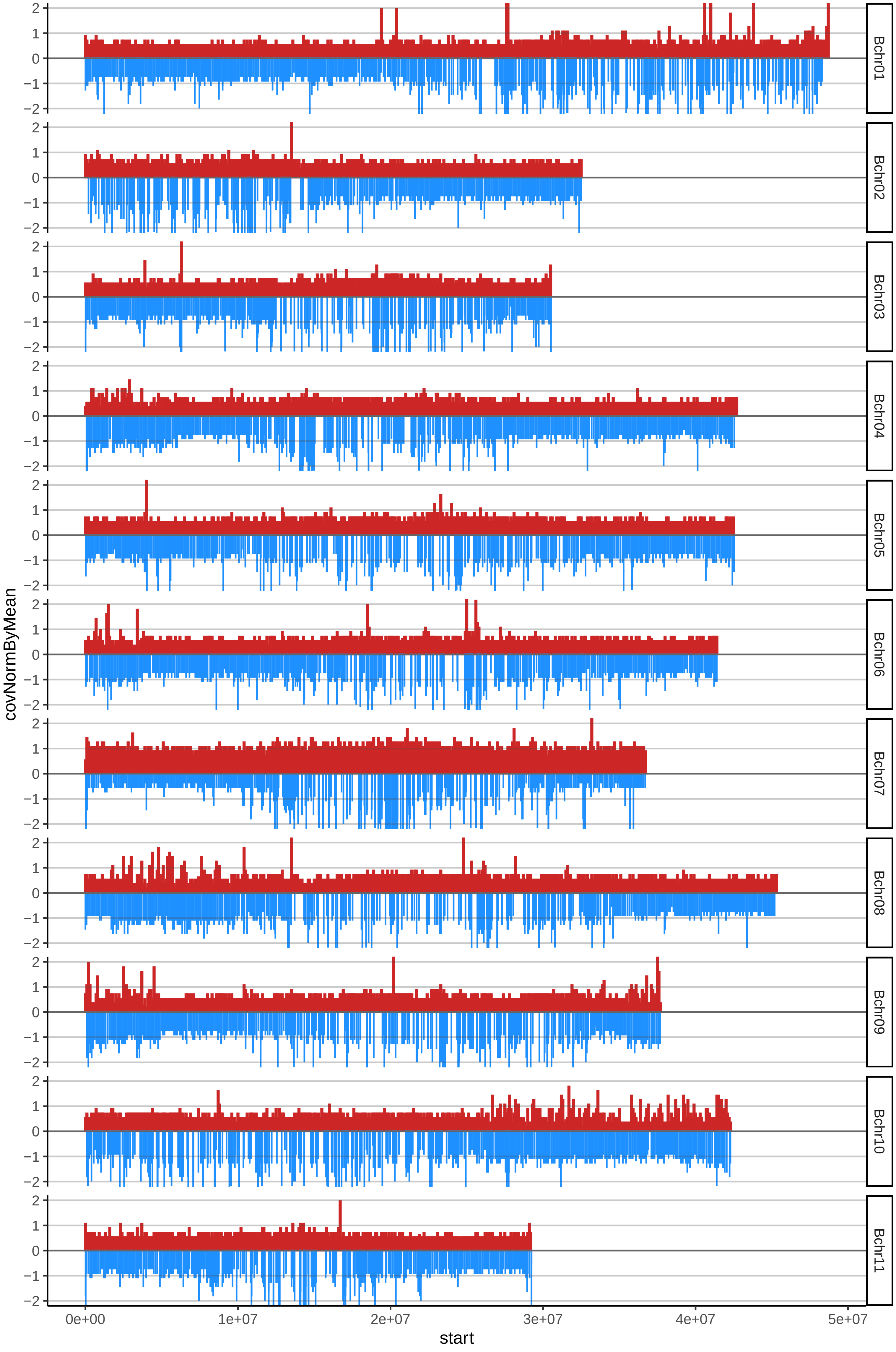

### 5BENDMOSSEDJOcov_byA.pdf

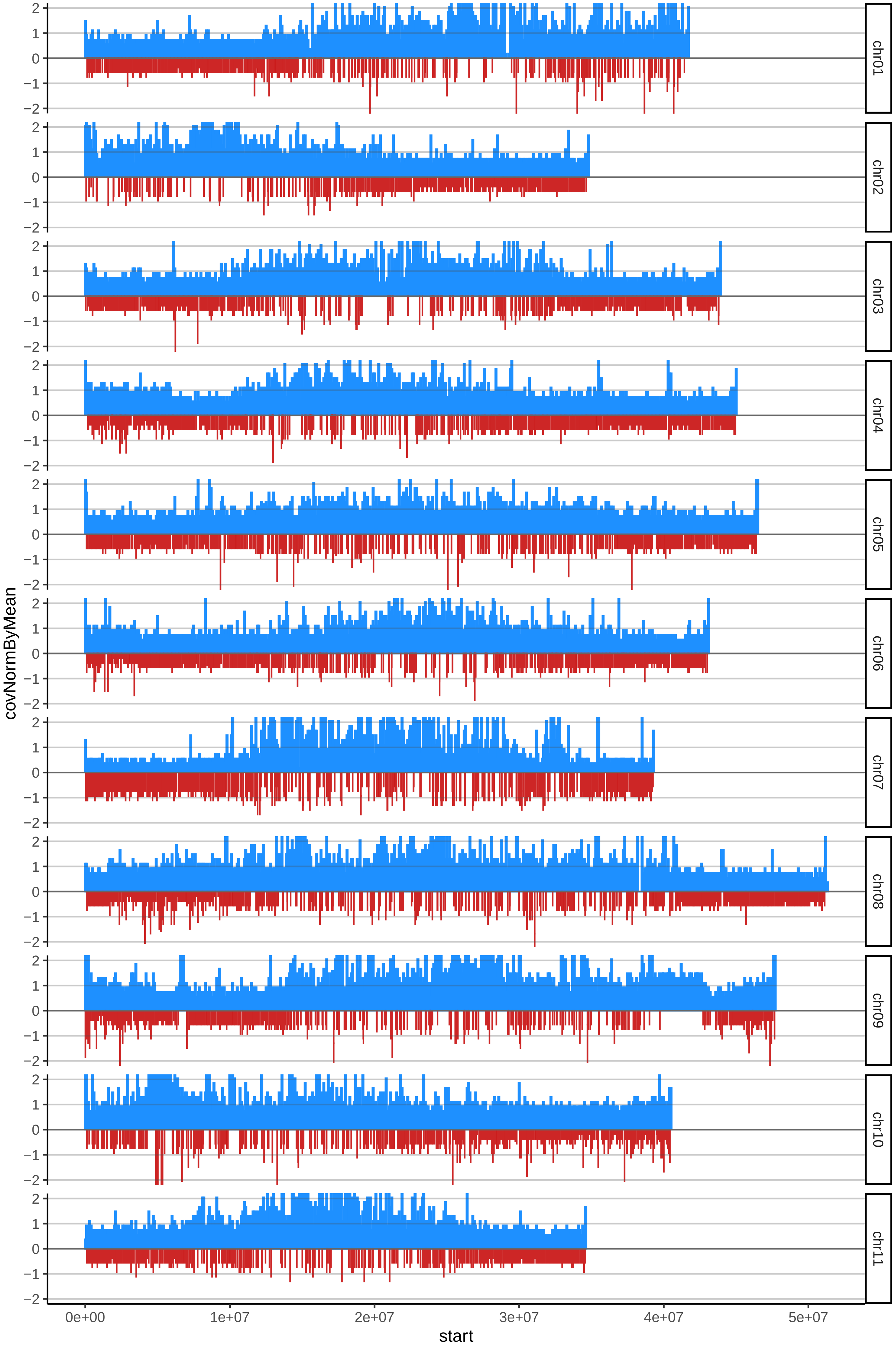

### 11CURRAREENANOcov_byA.pdf

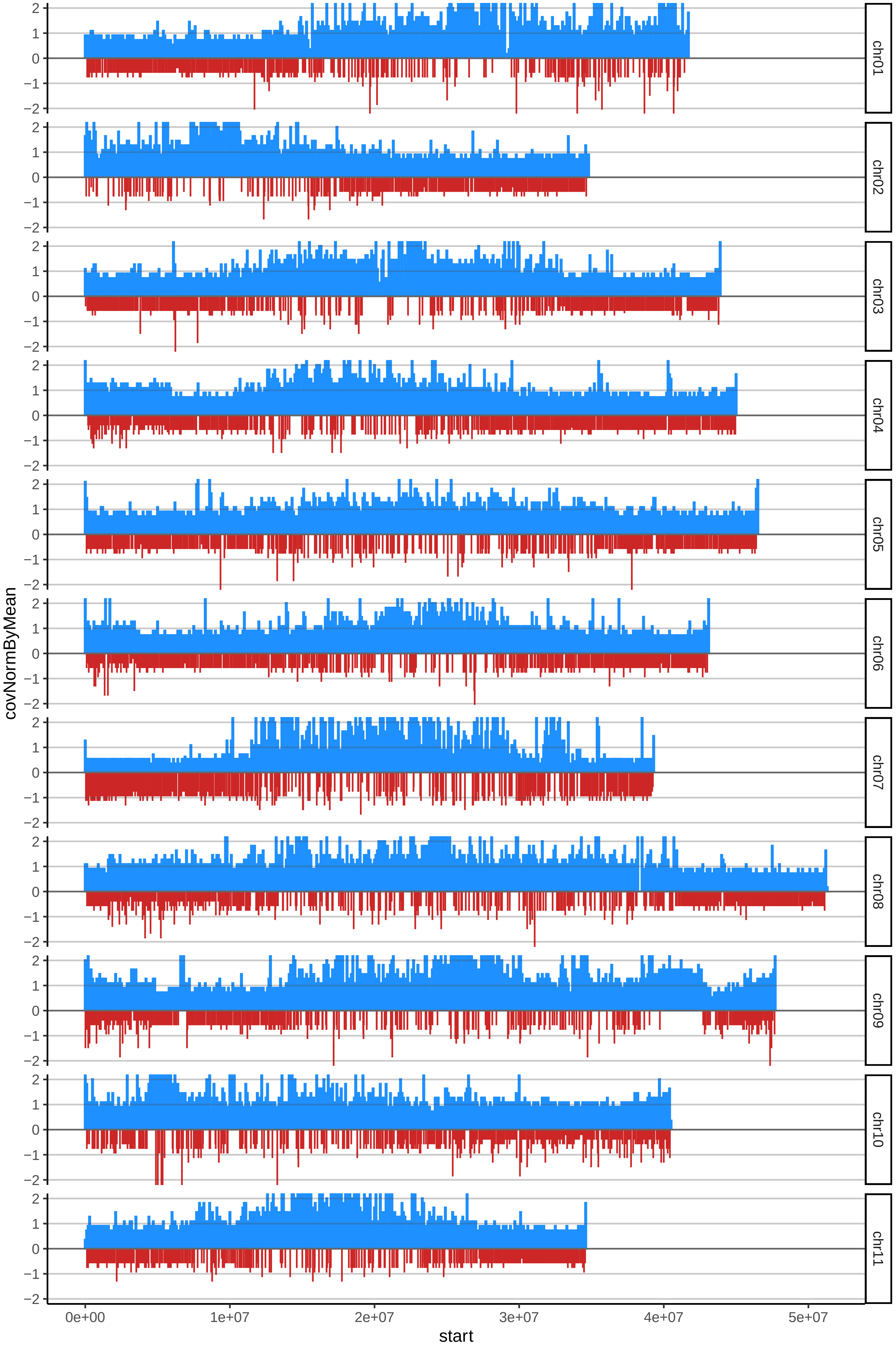

### 11CURRAREENANOcov_byB.pdf

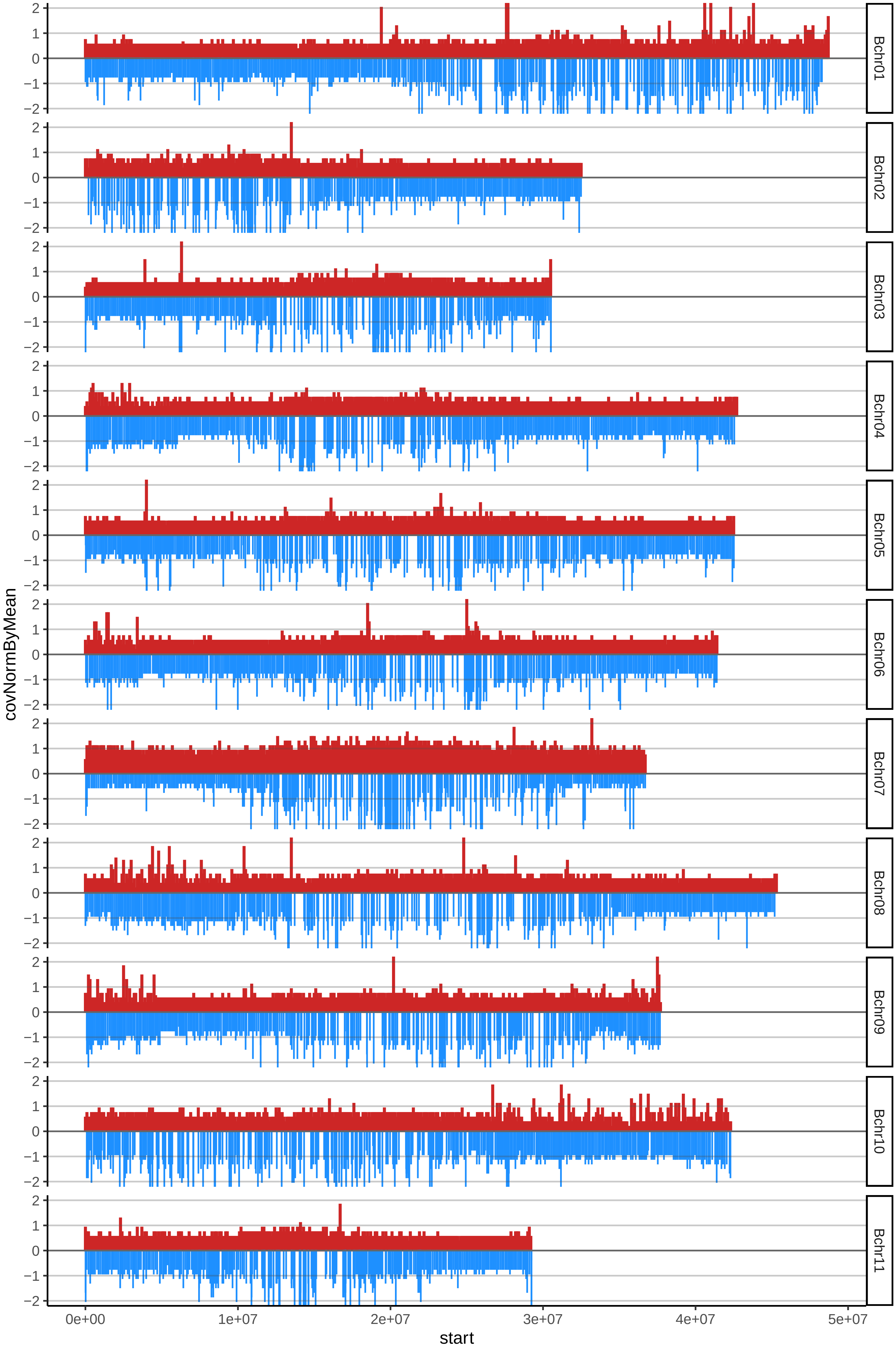

### 12DIBYcov_byA.pdf

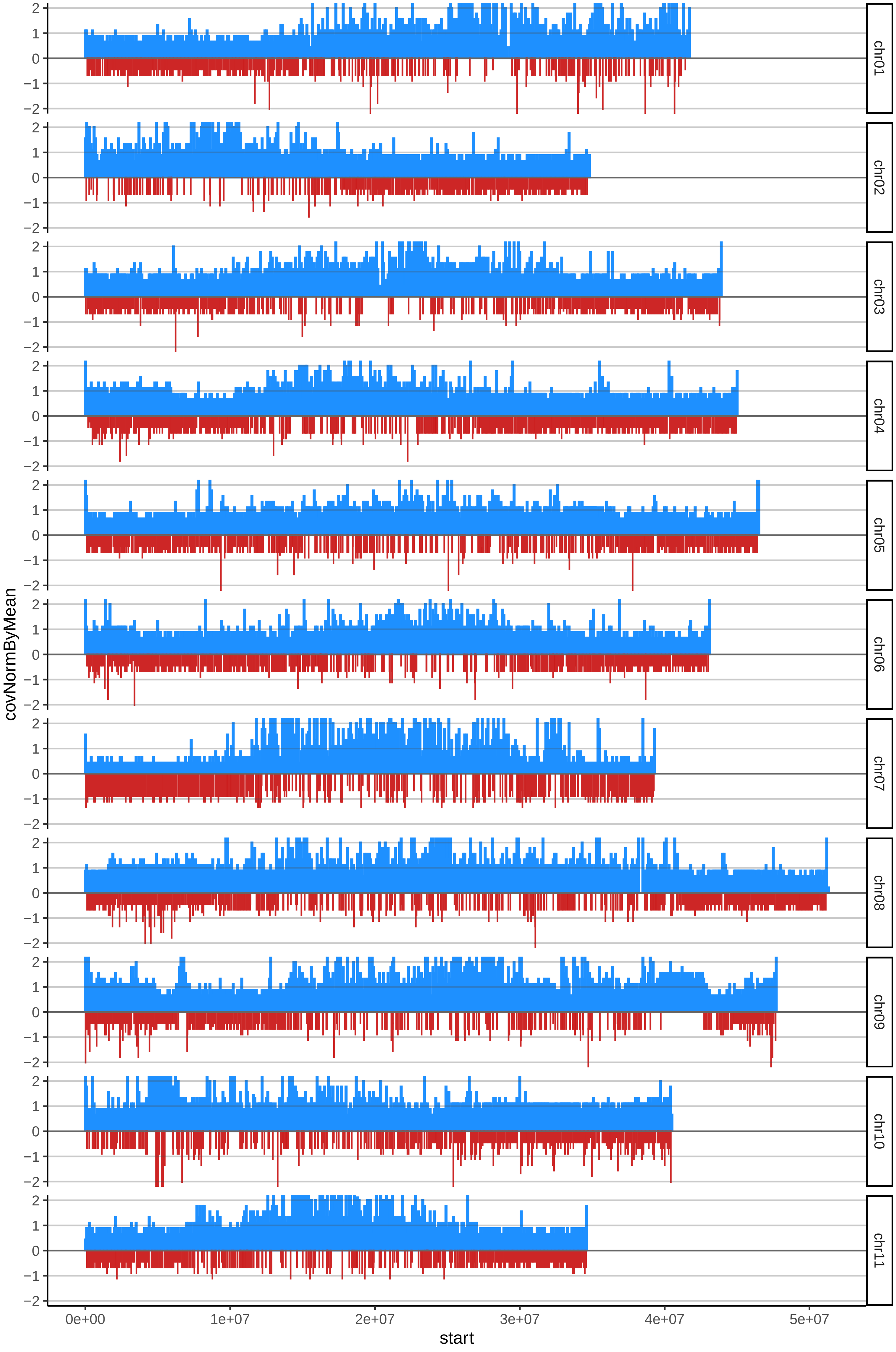

### 12DIBYcov_byB.pdf

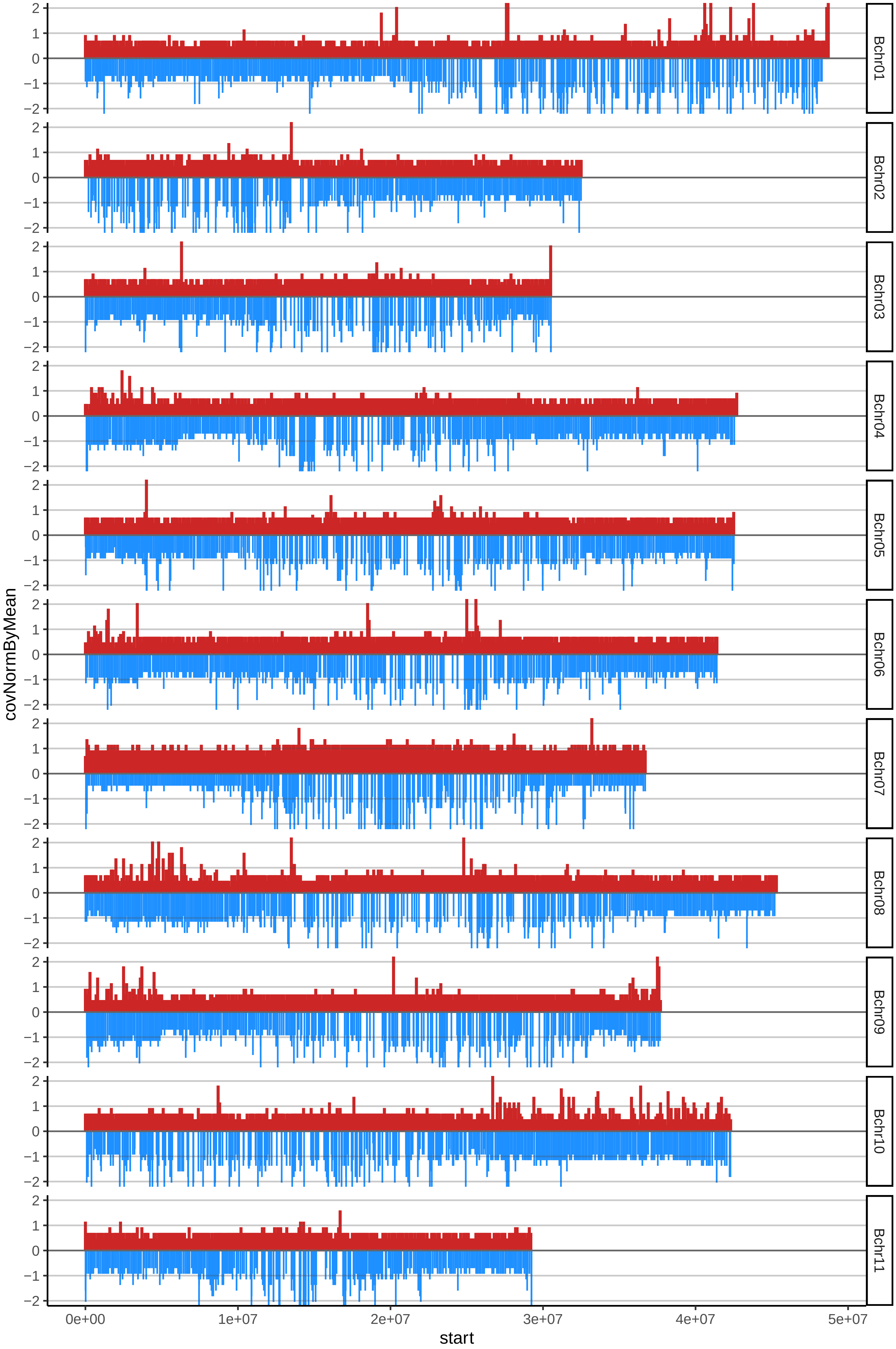

### 13DOMINICO300cov_byA.pdf

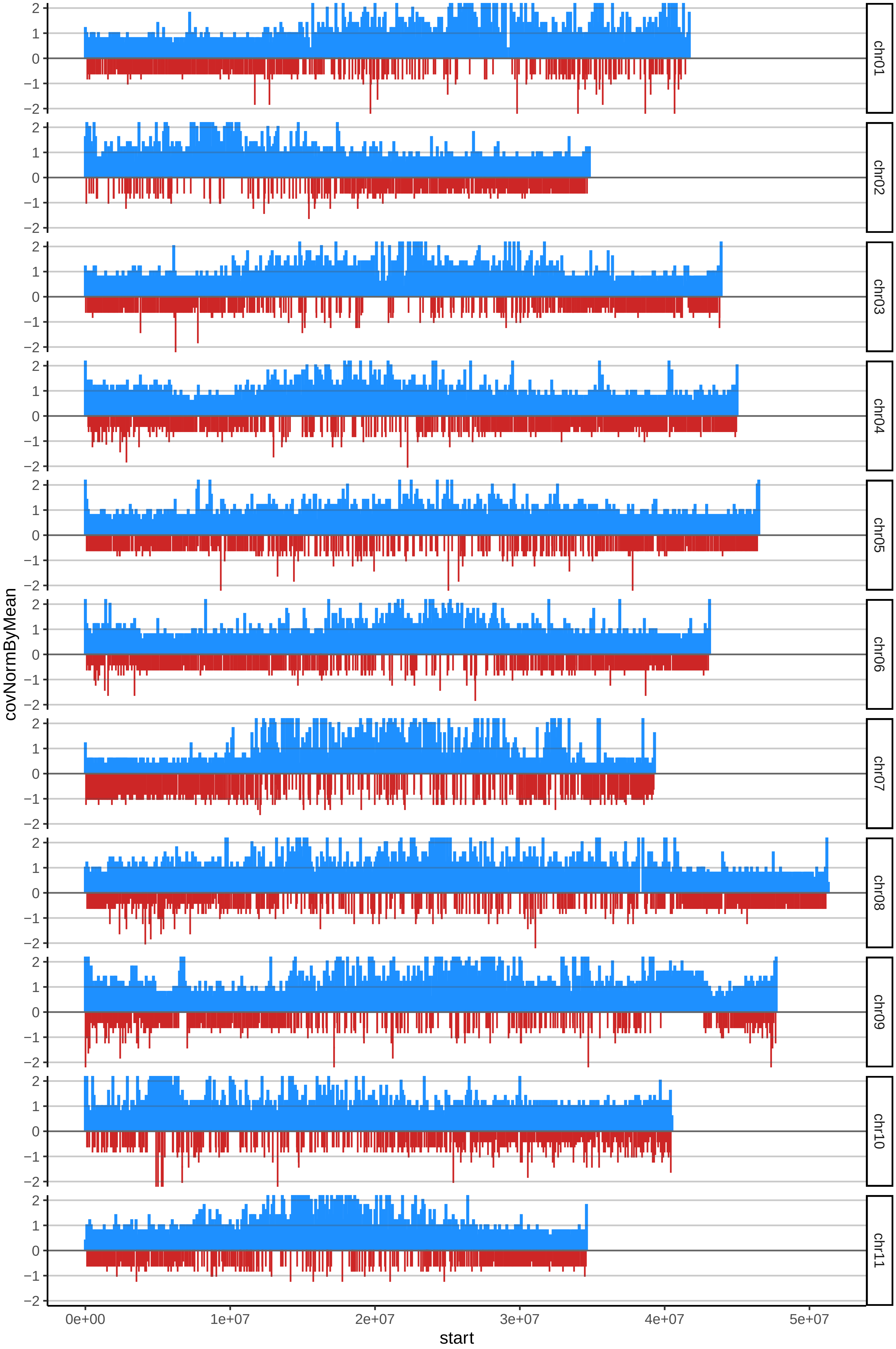

### 14DOMINICOANCUYANOcov_byA.pdf

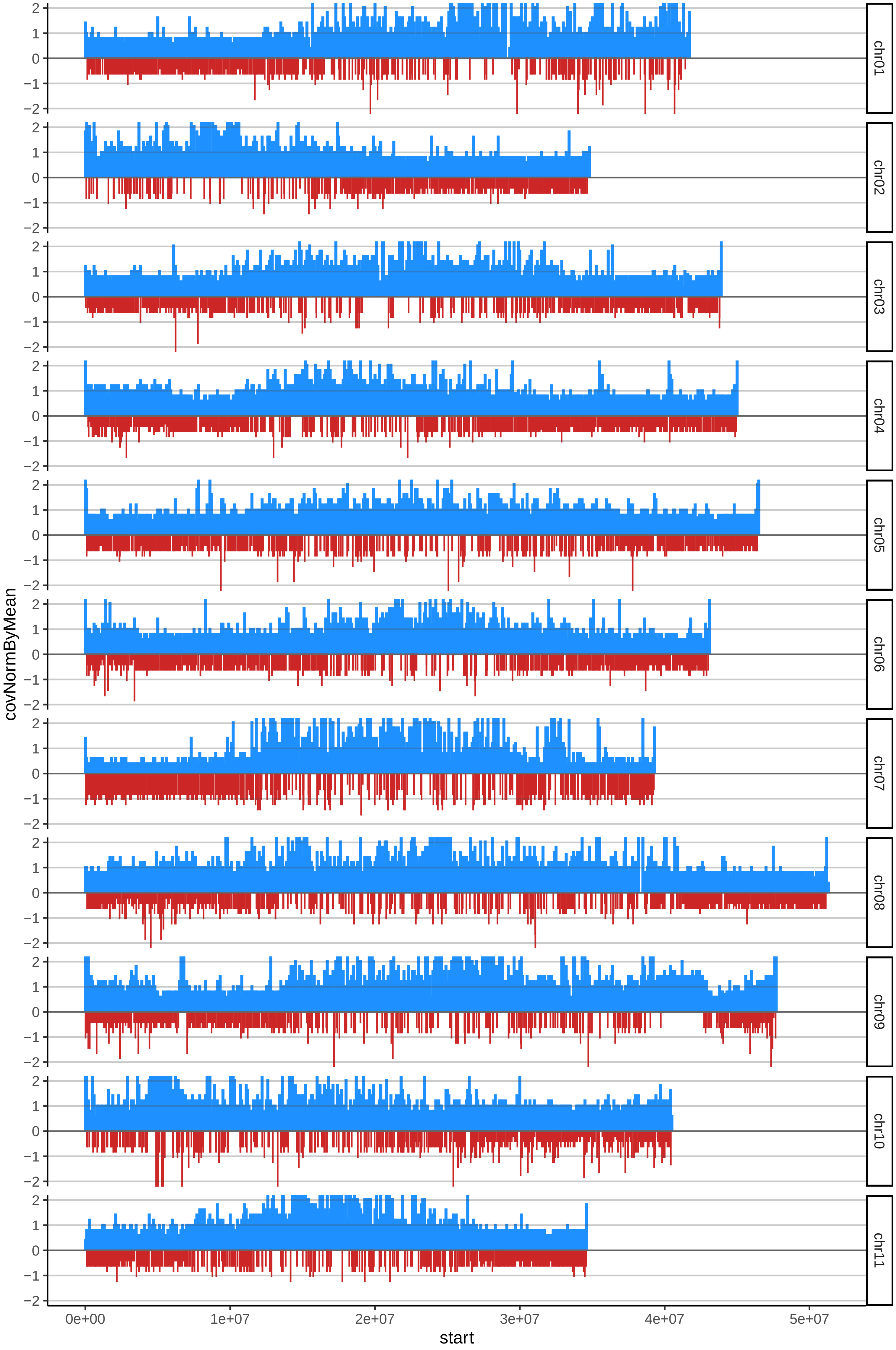

### 14DOMINICOANCUYANOcov_byB.pdf

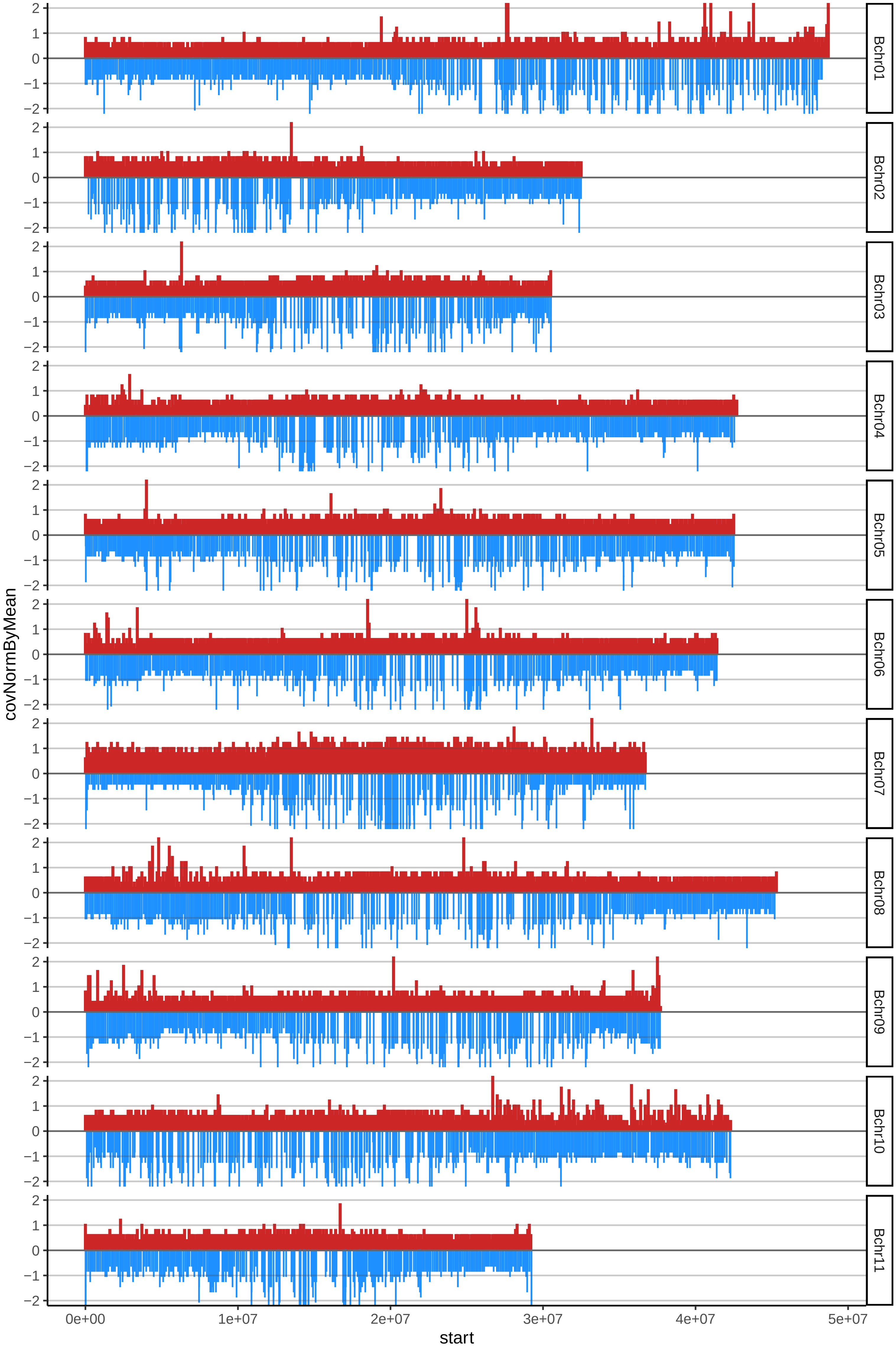

### 18DOMINICOGUAICOSOcov_byA.pdf

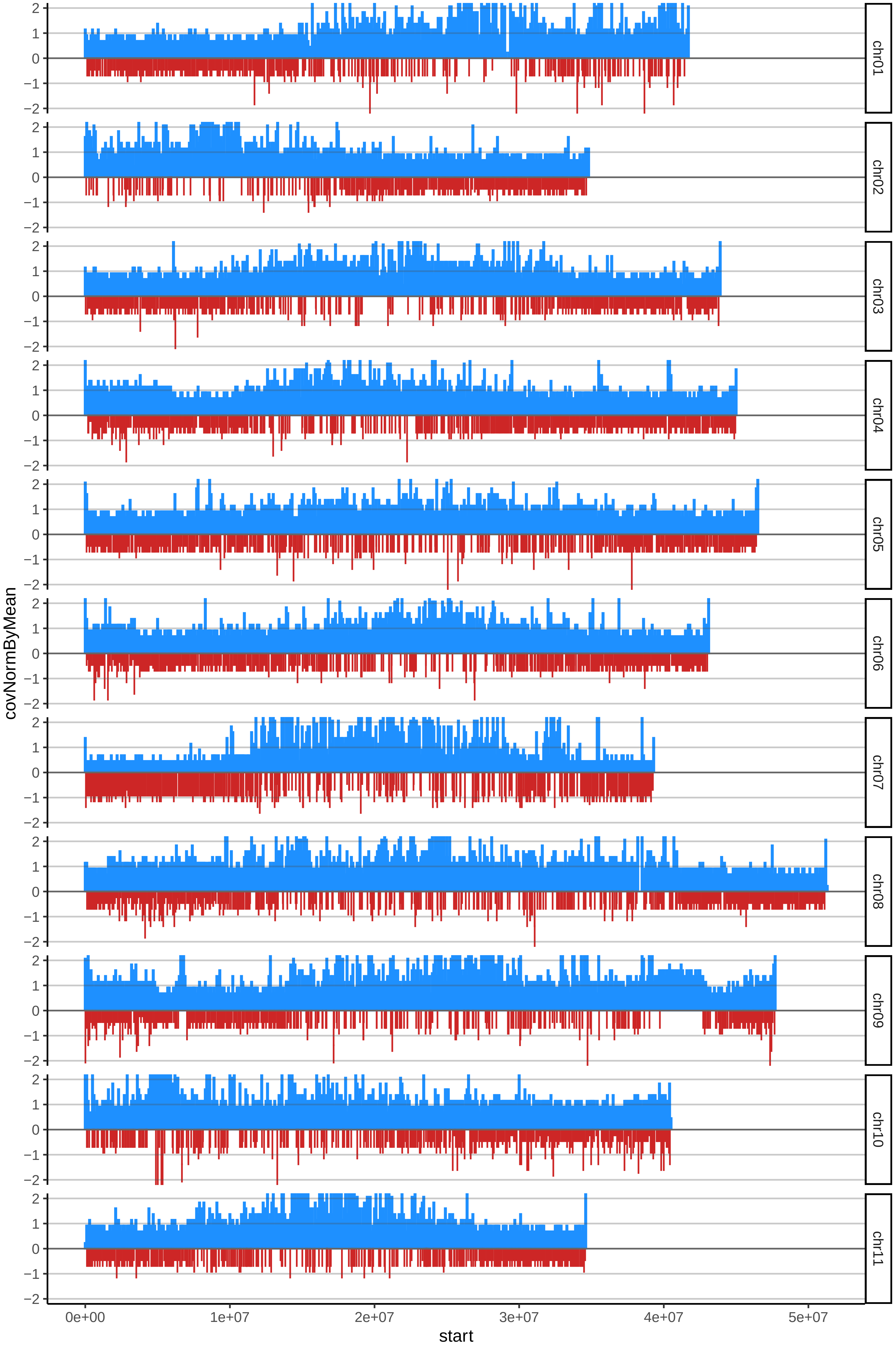

### 18DOMINICOGUAICOSOcov_byB.pdf

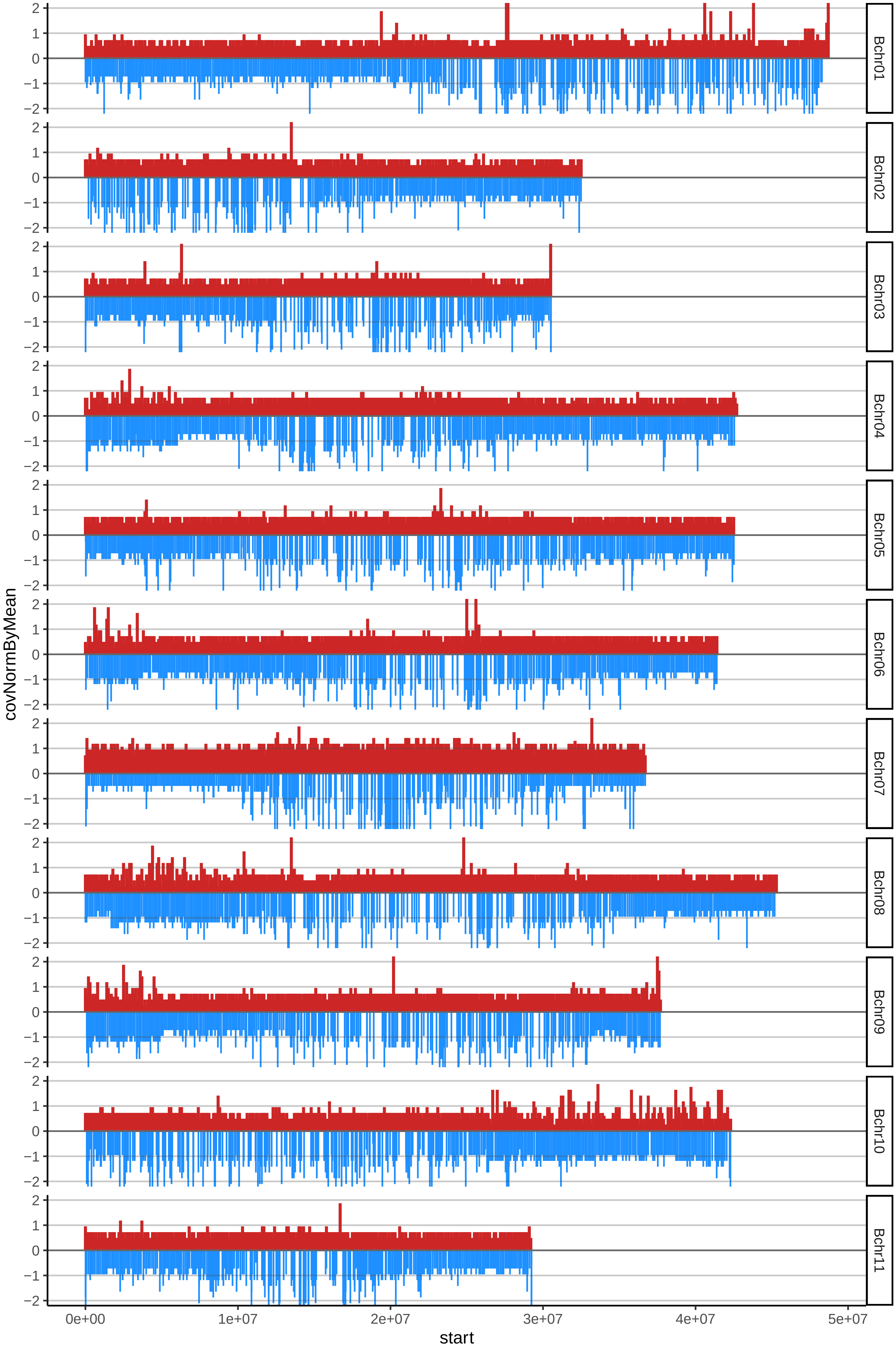

### 19DOMINICOHARTONCOMUNcov_byA.pdf

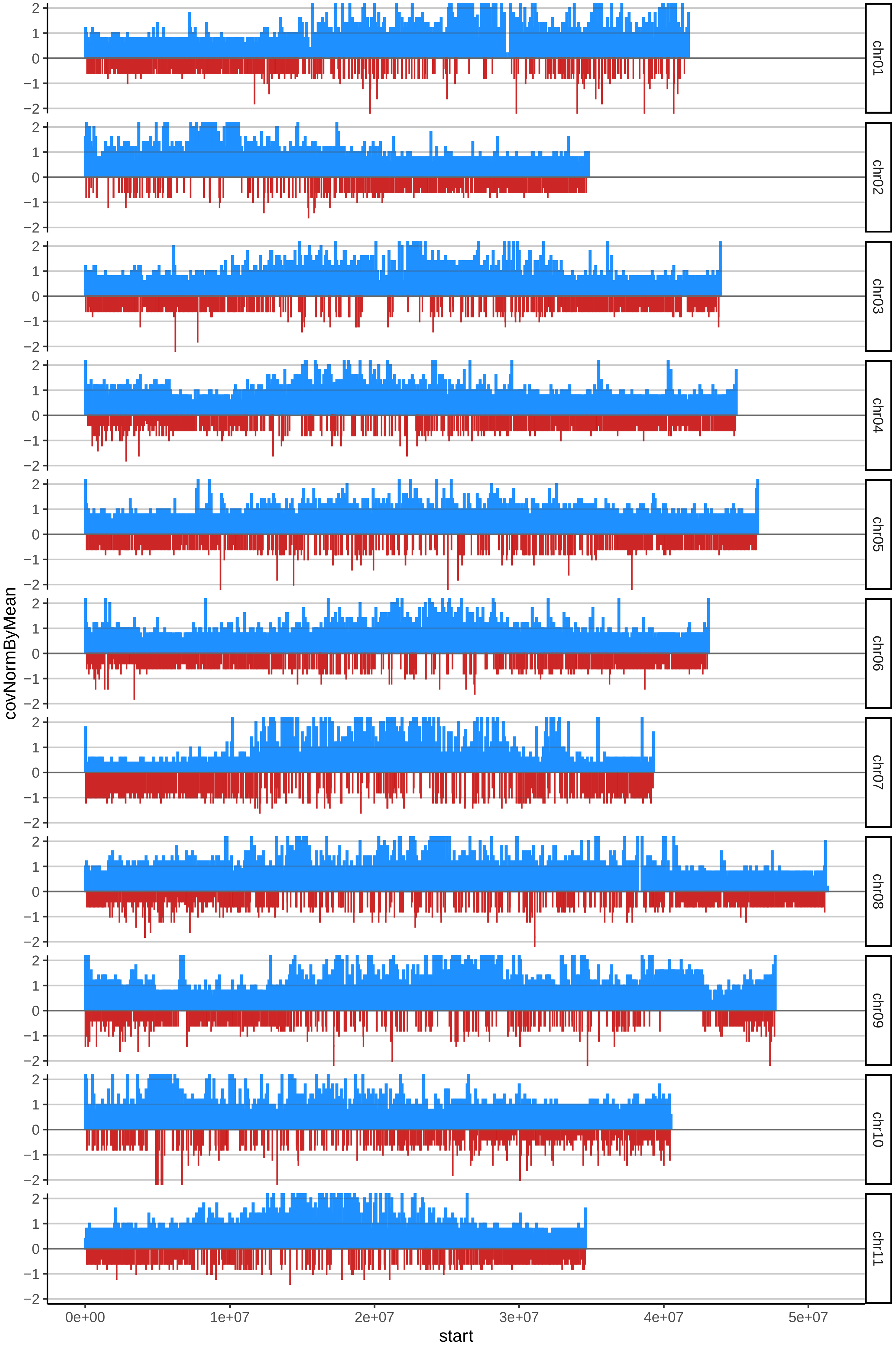

### 19DOMINICOHARTONCOMUNcov_byB.pdf

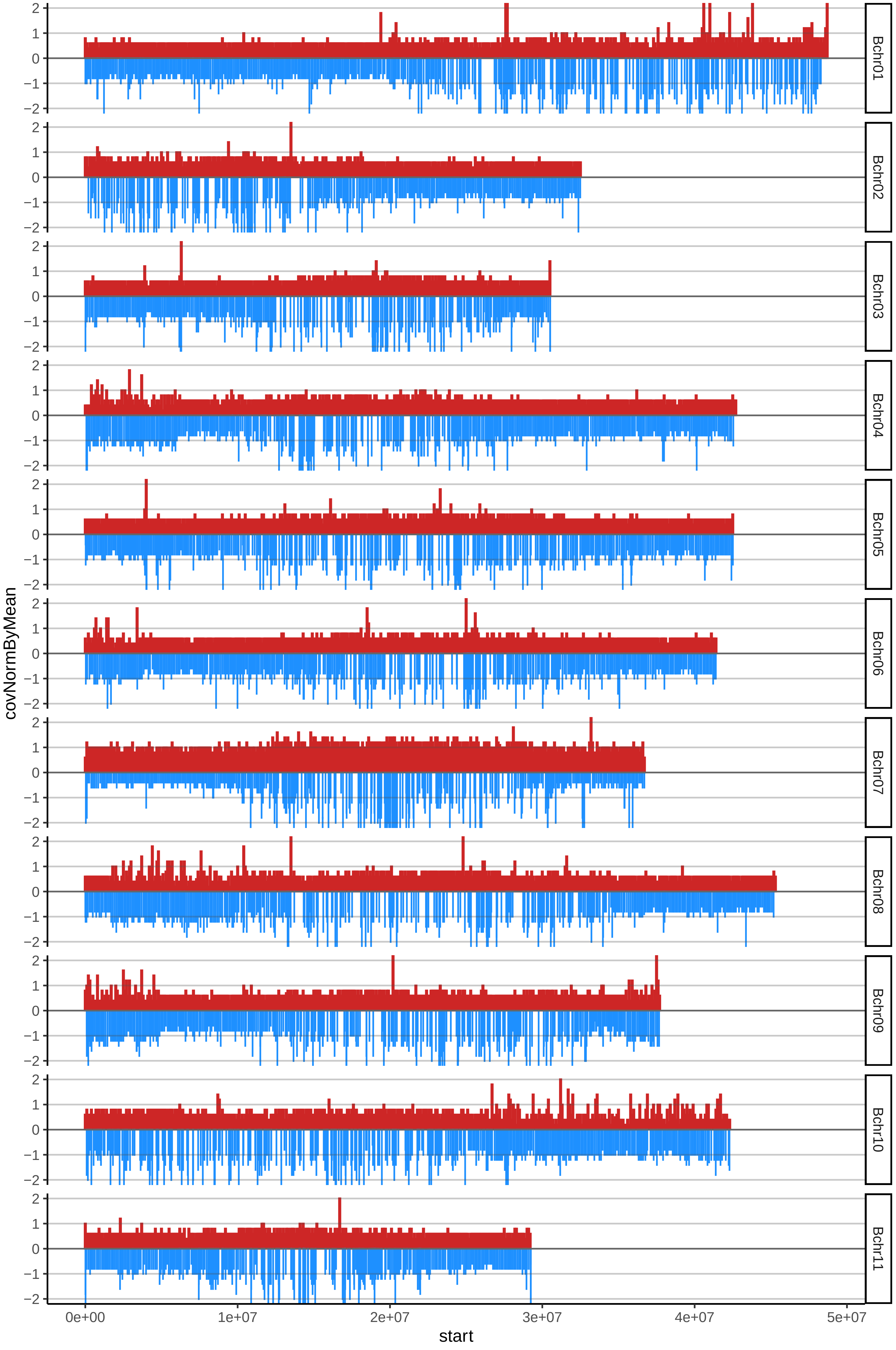

### 22DOMINICOHARTONVIOTAcov_byA.pdf

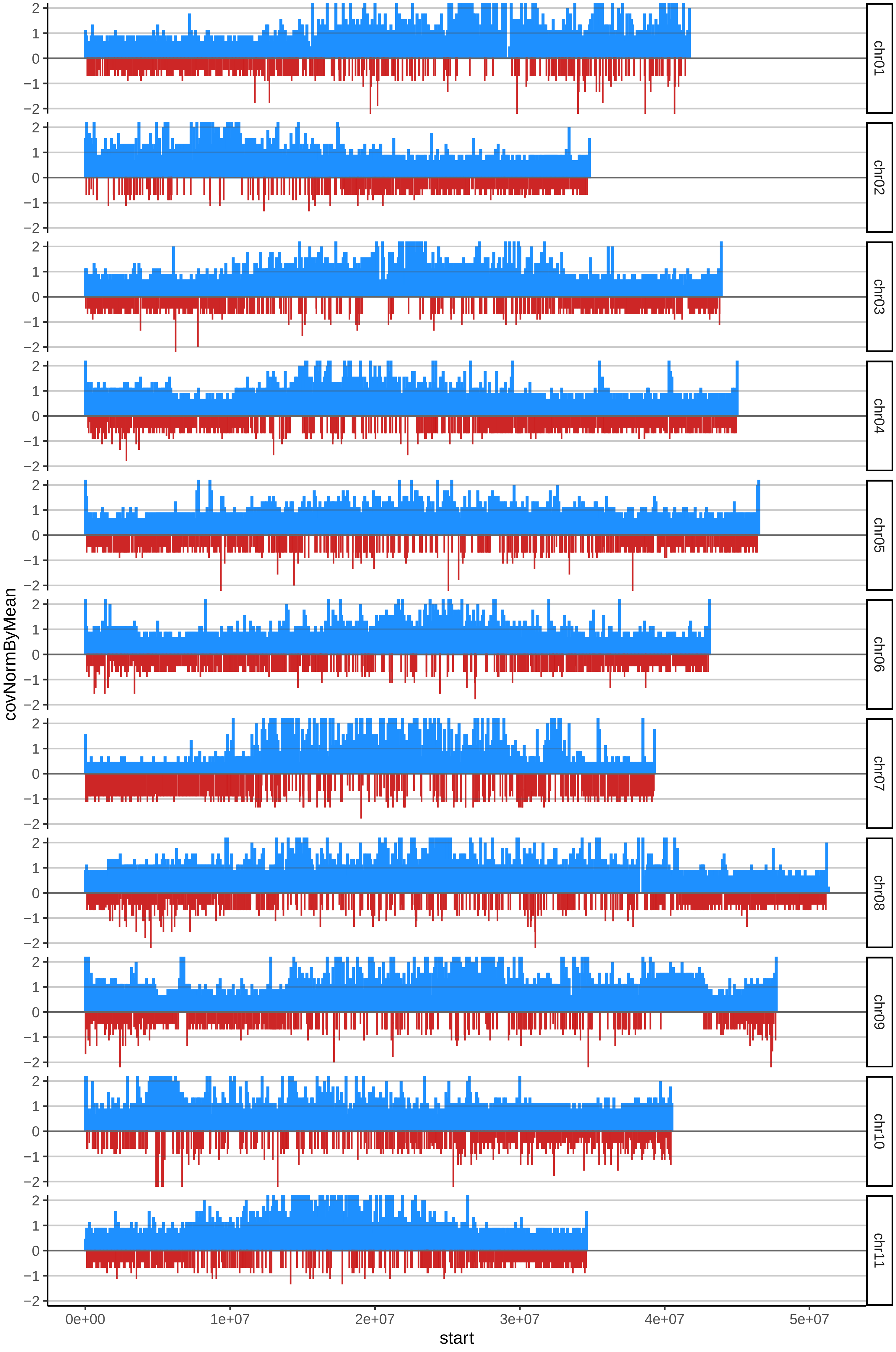

### 23DOMINICOMAQUENNOcov_byA.pdf

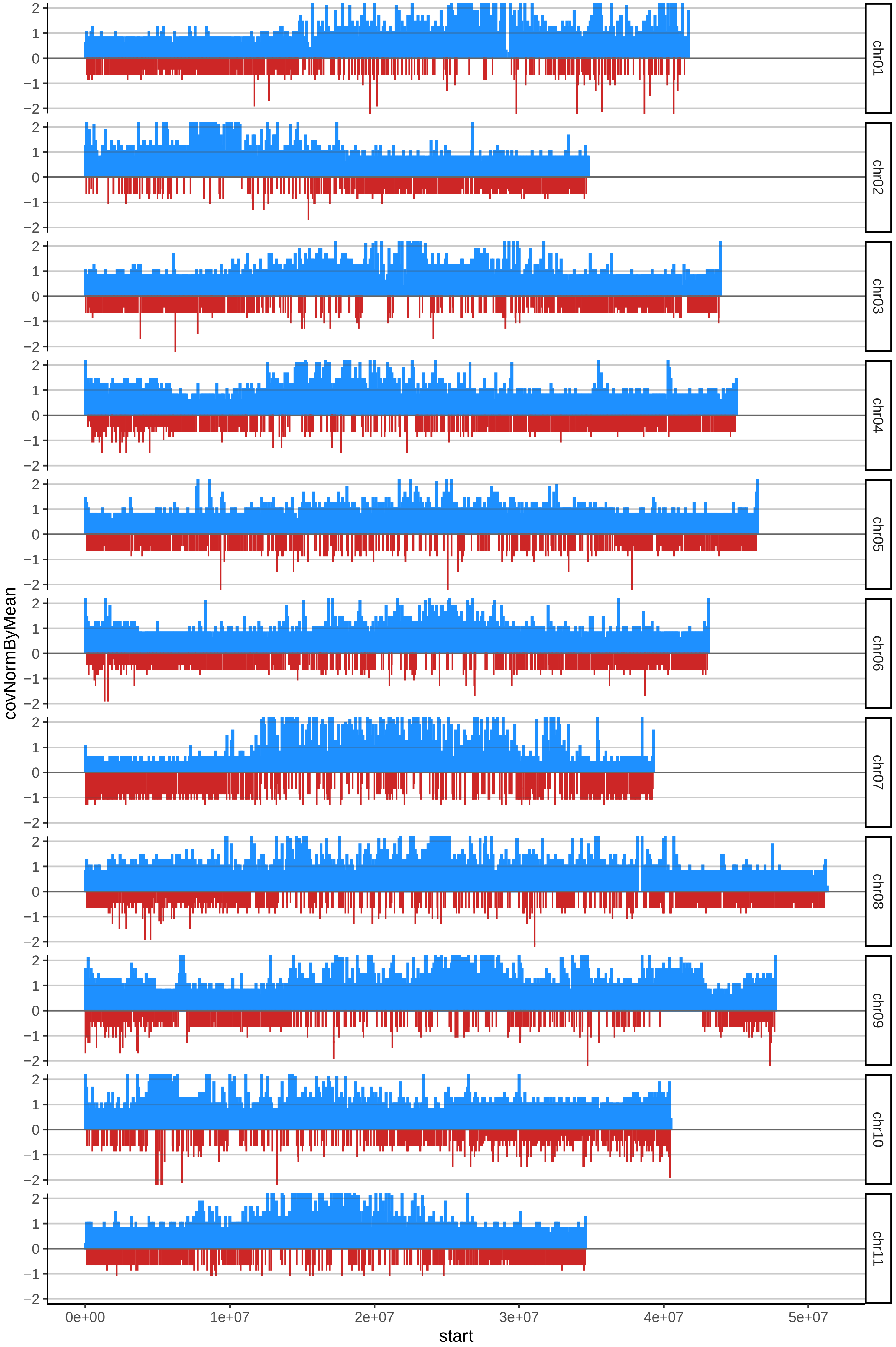

### 23DOMINICOMAQUENNOcov_byB.pdf

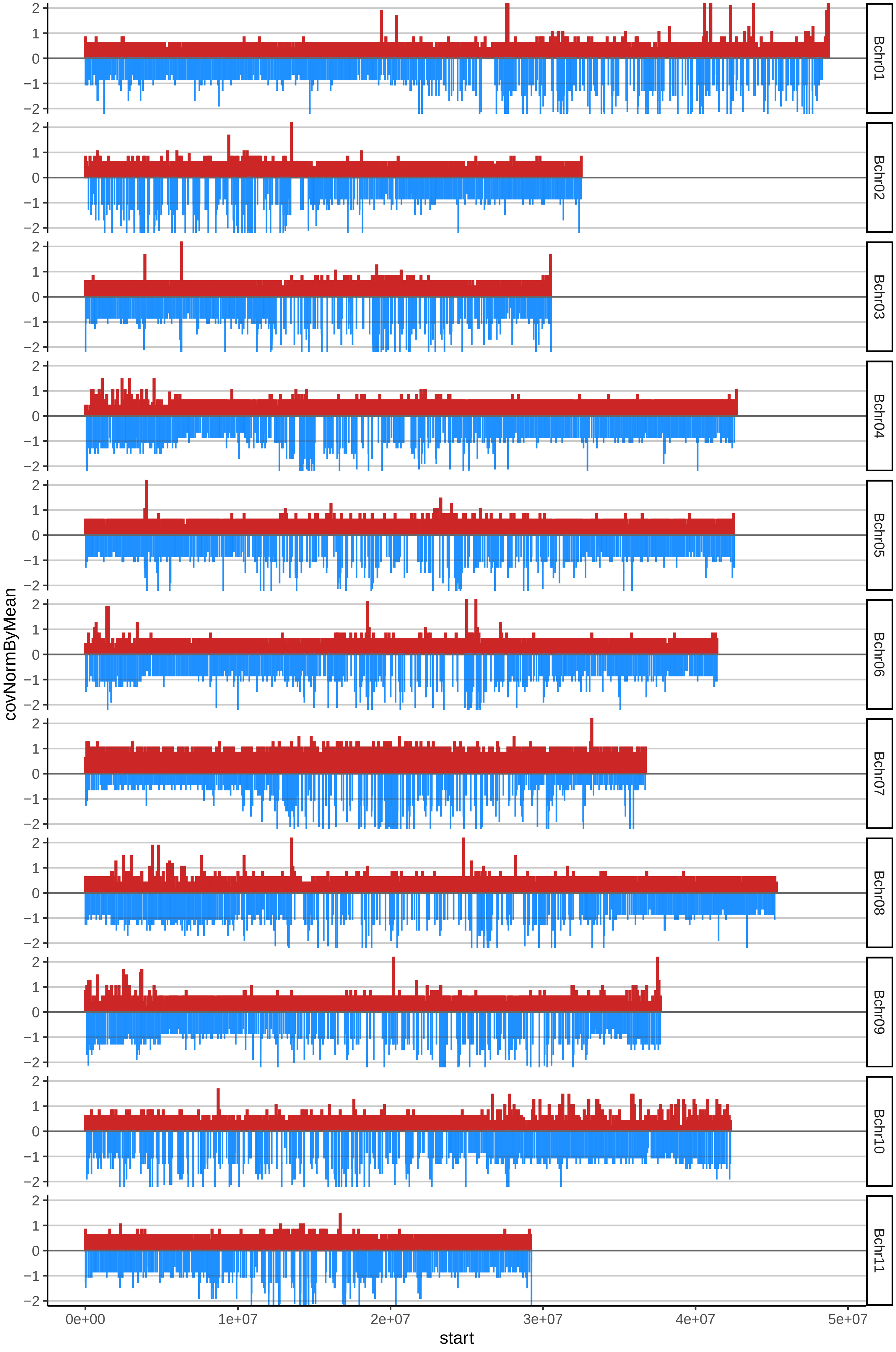

### 24DOMINICOMOCHOcov_byA.pdf

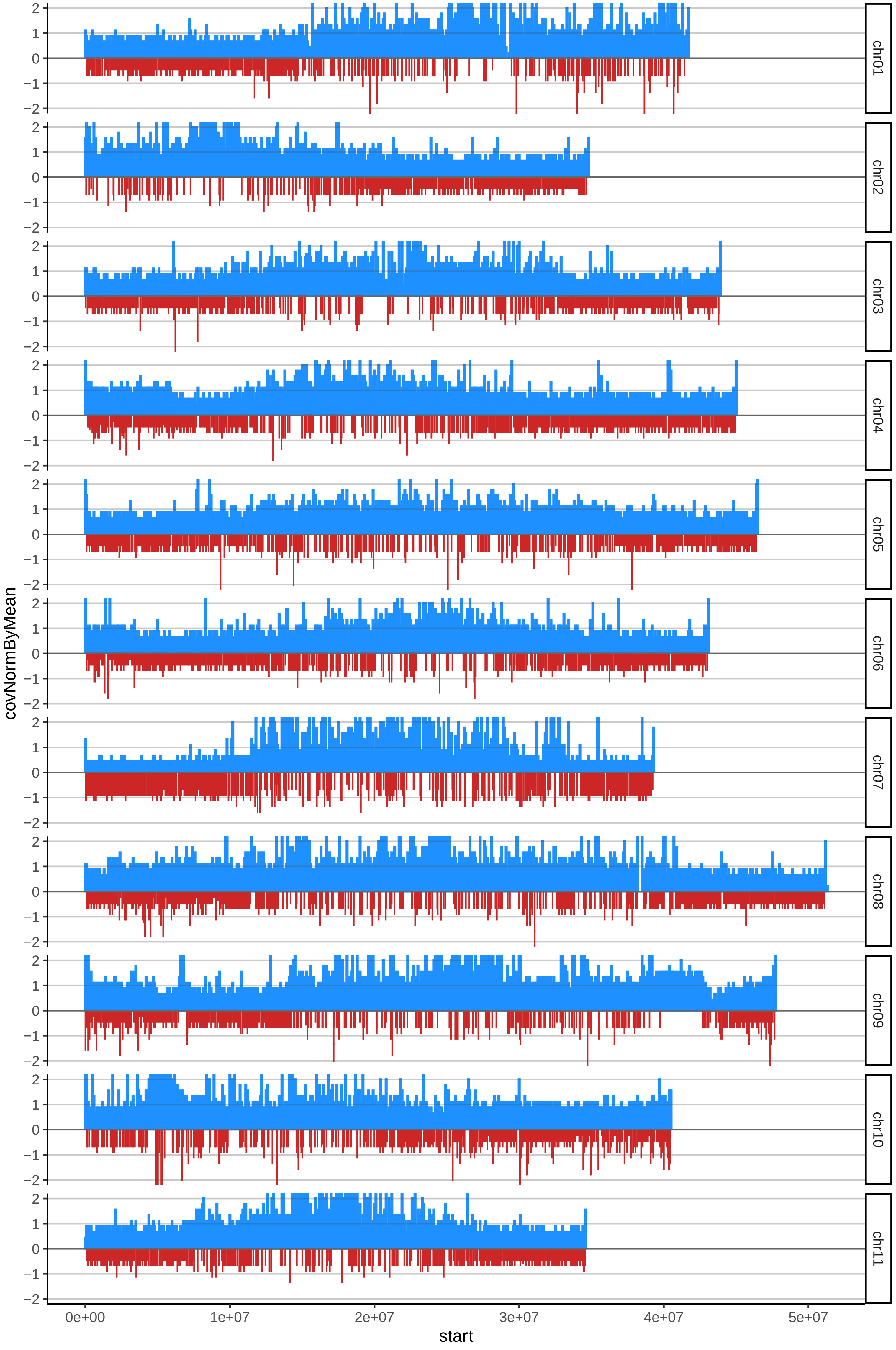

### 24DOMINICOMOCHOcov_byB.pdf

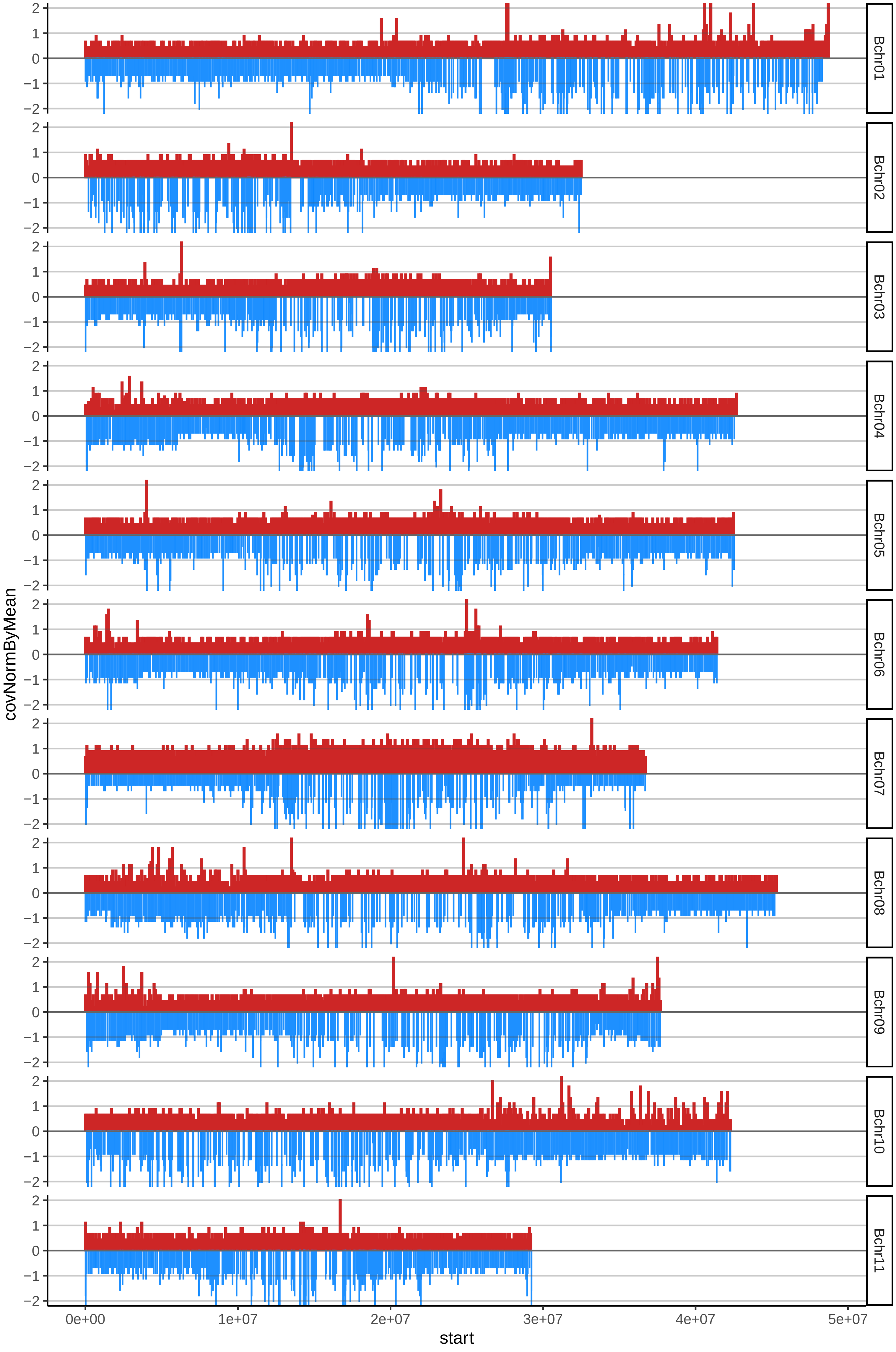

### 26DOMINICONEGROcov_byA.pdf

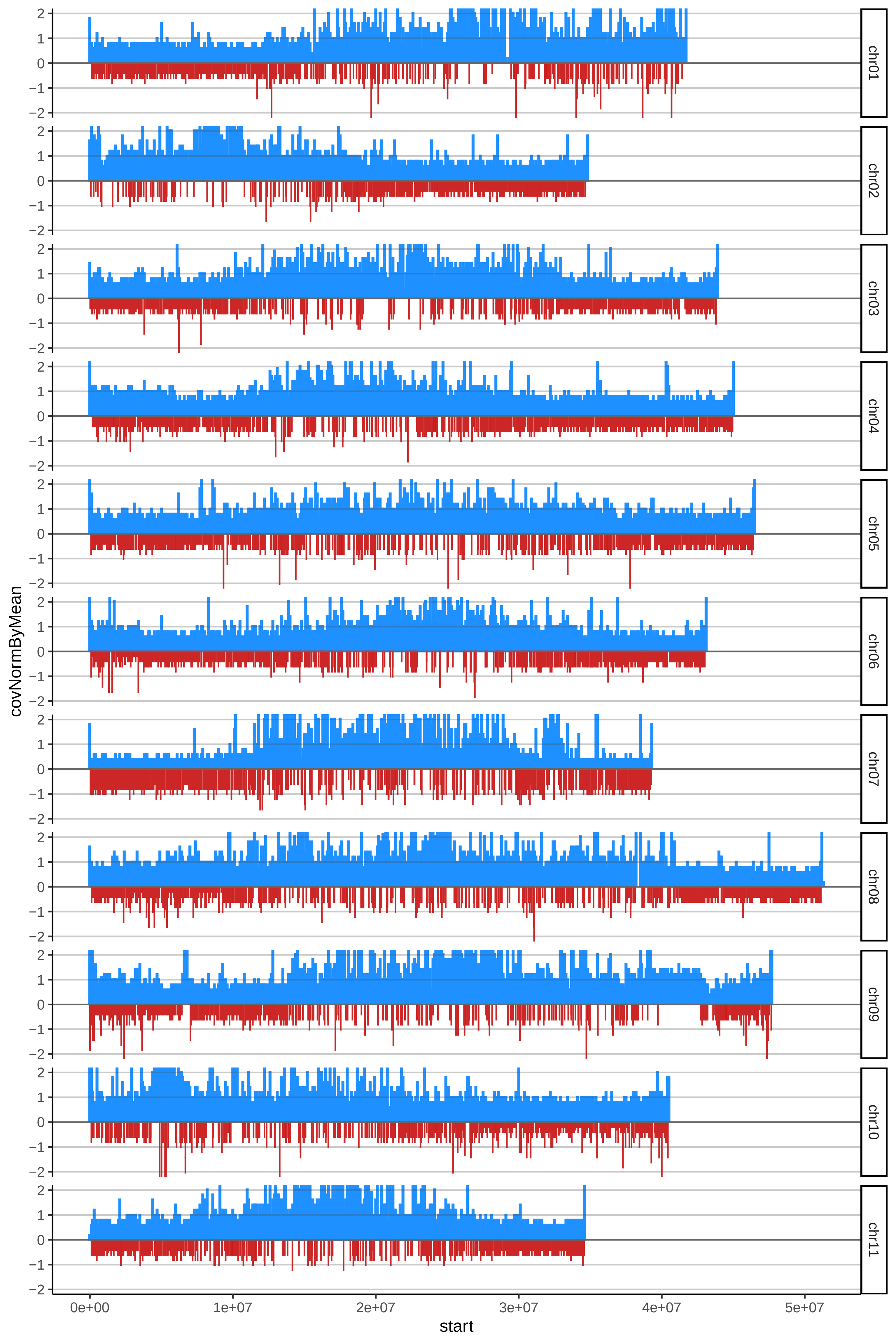

### 26DOMINICONEGROcov_byB.pdf

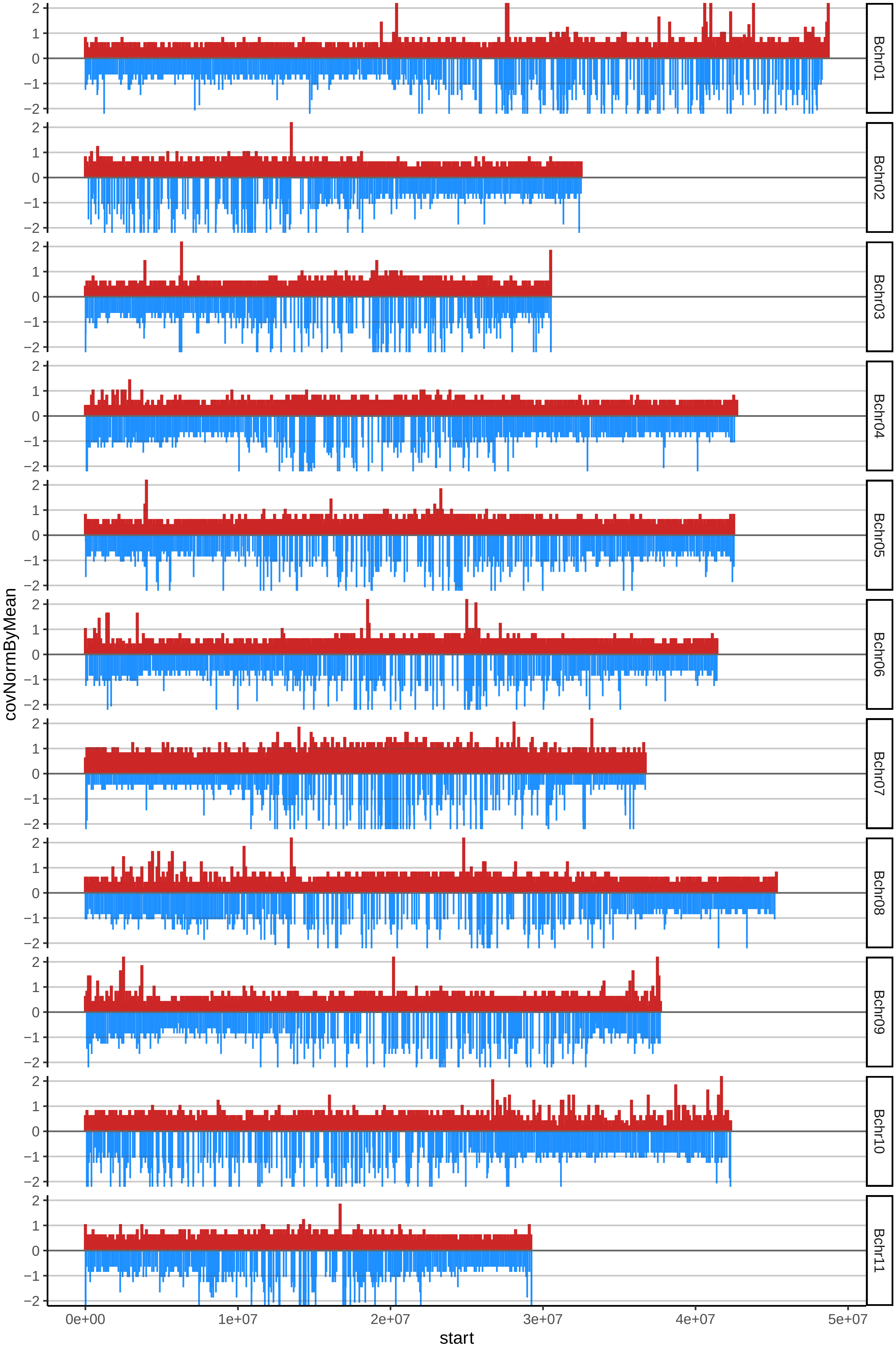

### 27DOMINICOTUMACOcov_byA.pdf

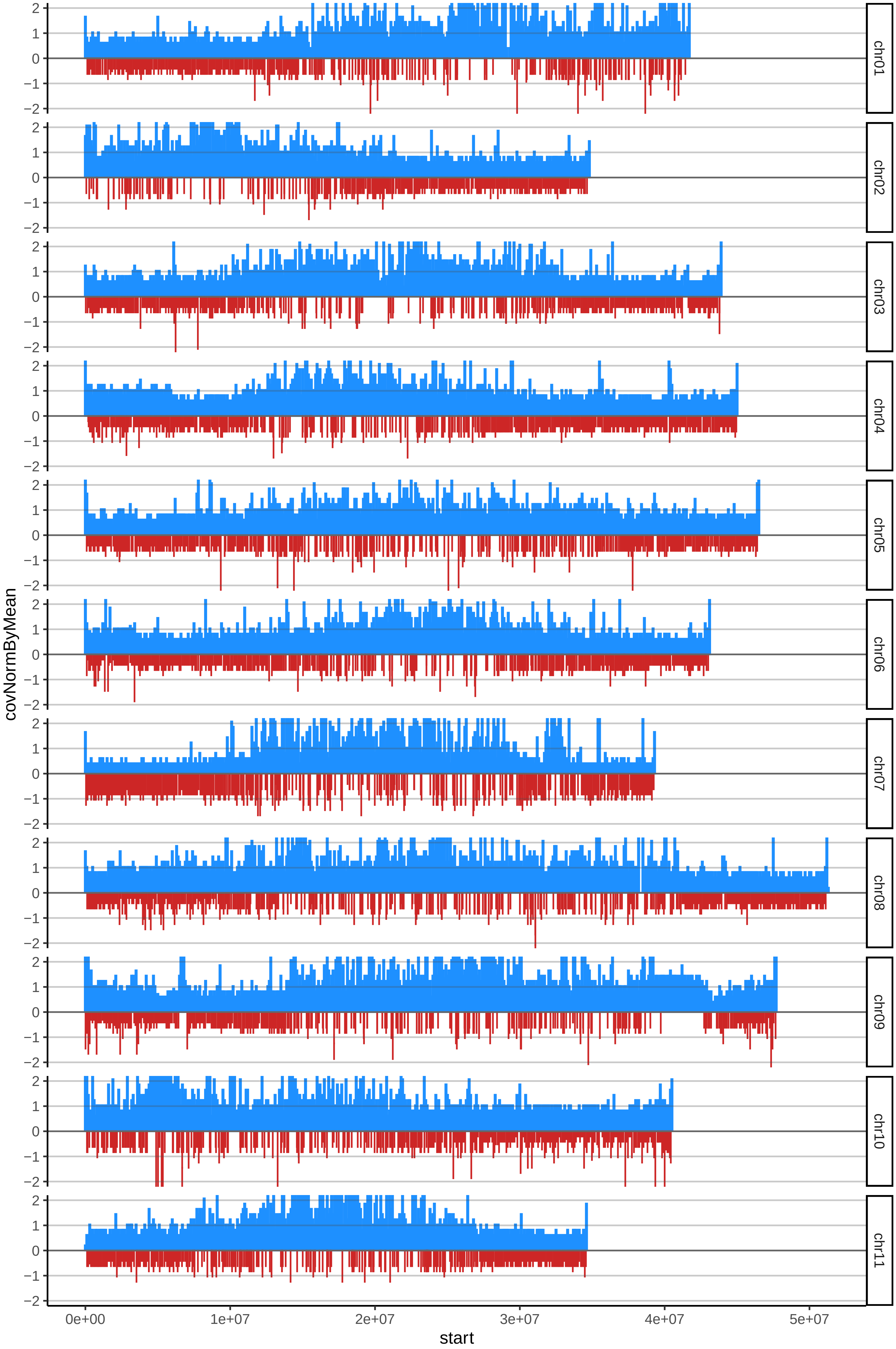

### 28ELATcov_byA.pdf

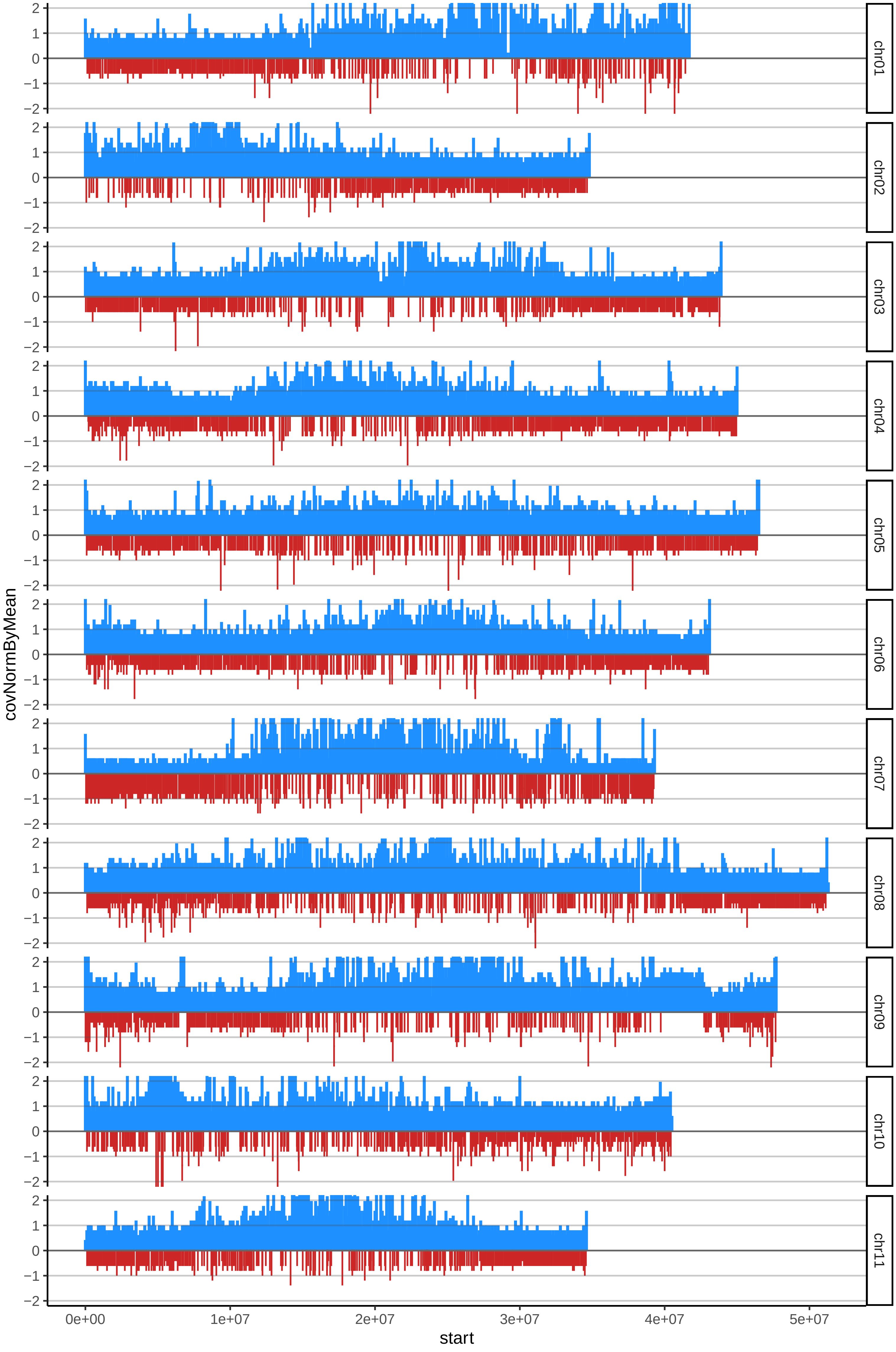

### 28ELATcov_byB.pdf

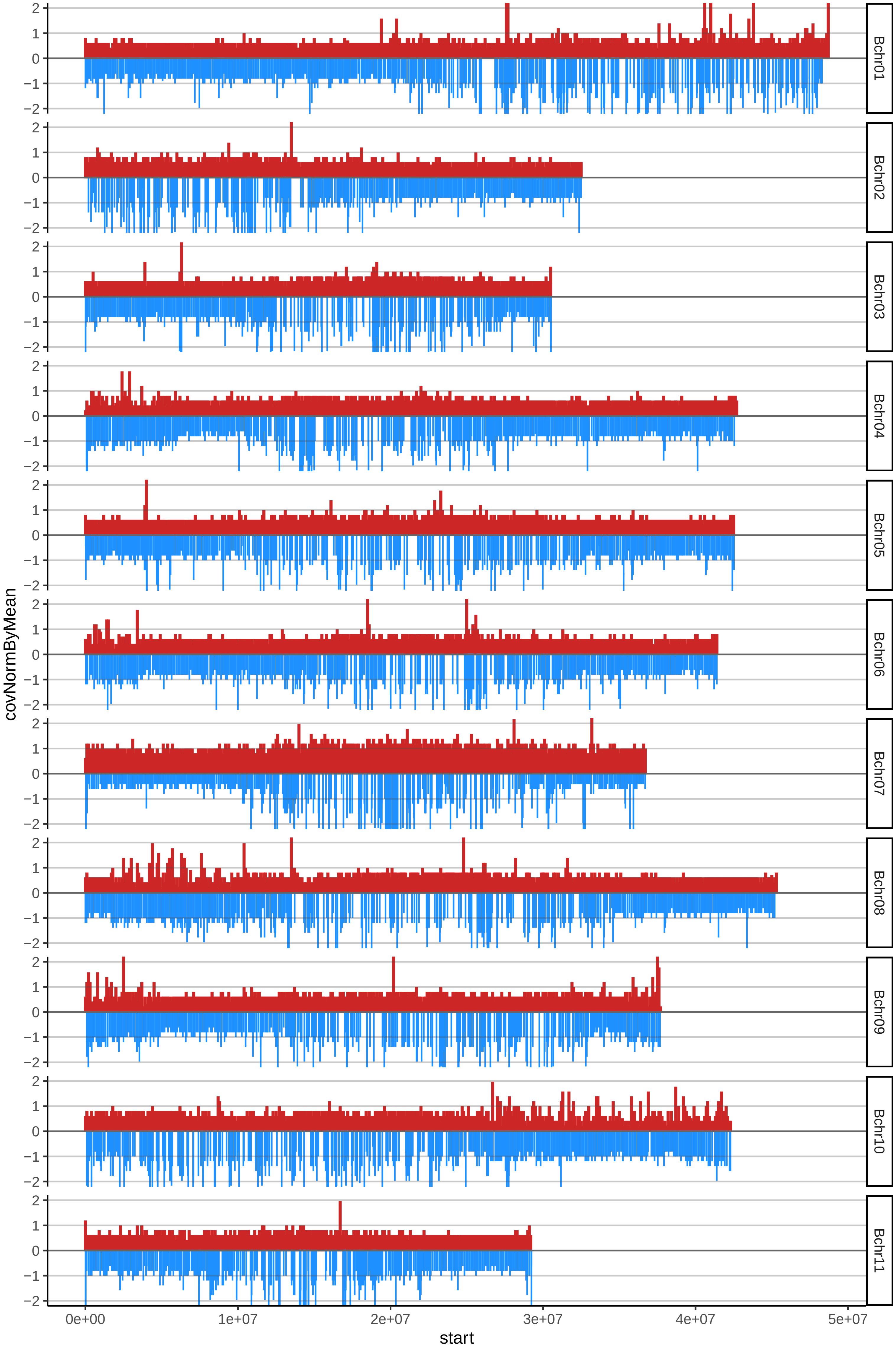

### 36FRENCHSOMBREcov_byB.pdf

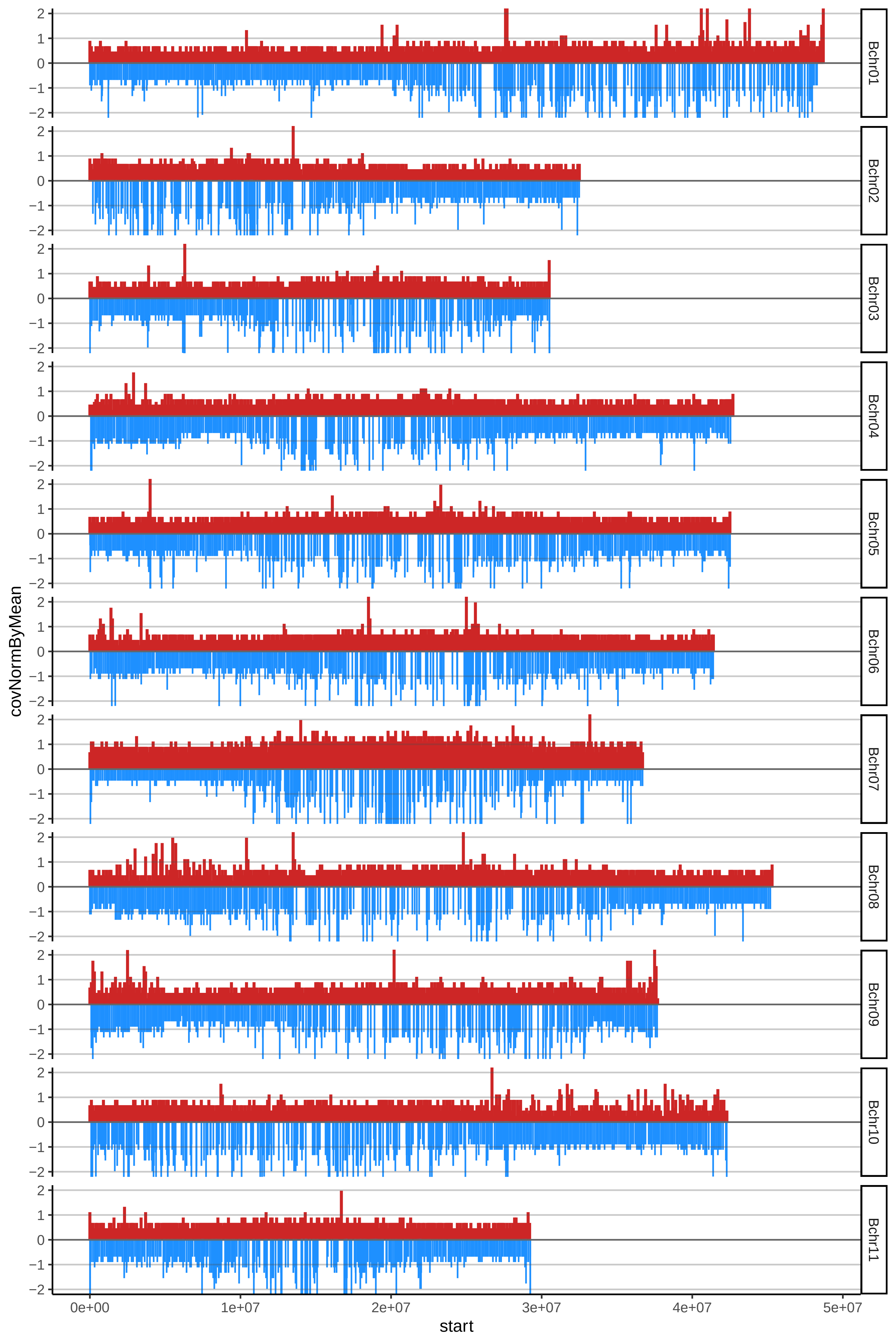

### 41HARTONCOMUNcov_byA.pdf

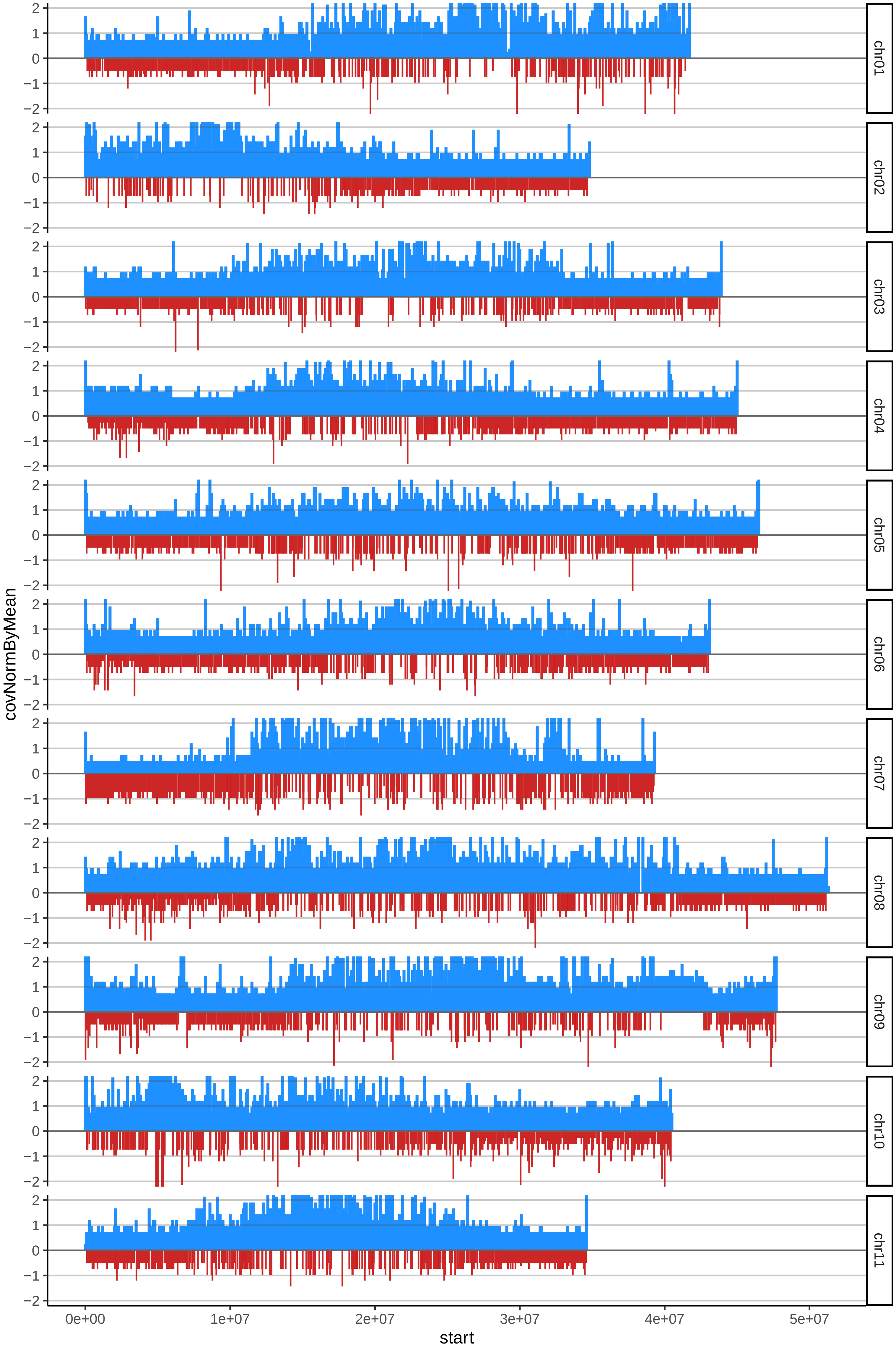

### 41HARTONCOMUNcov_byB.pdf

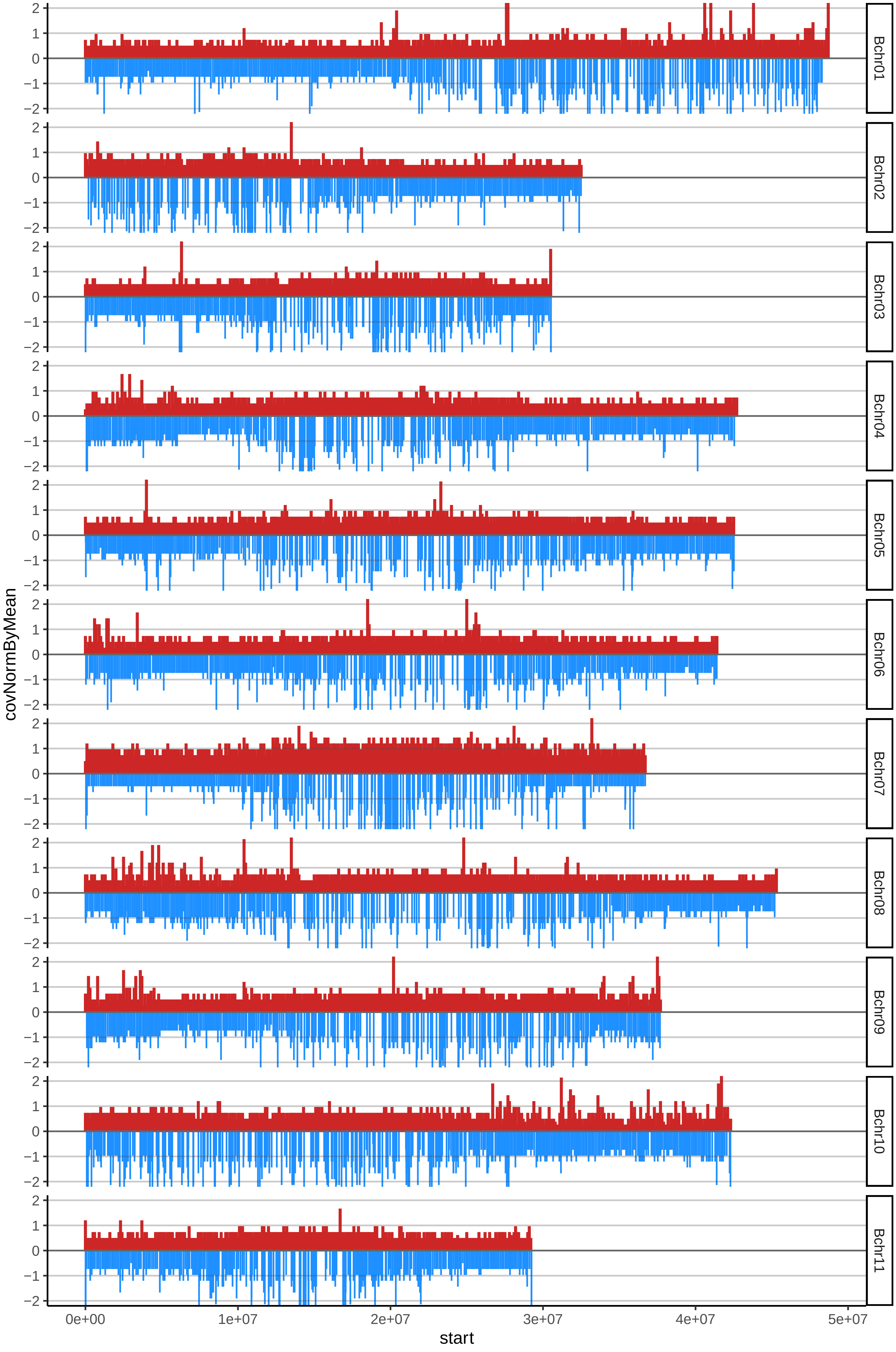

### 42HARTONDELMETAcov_byA.pdf

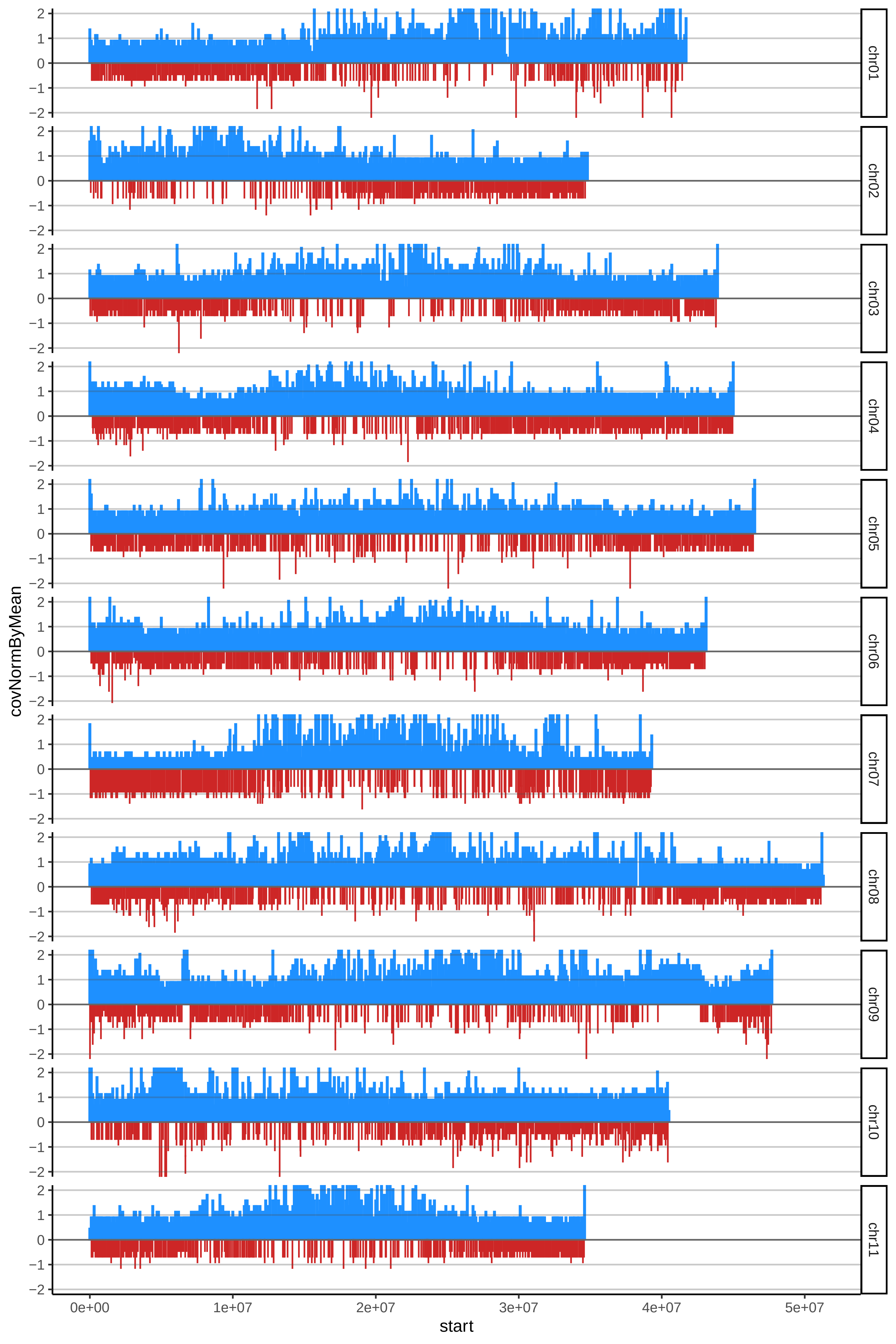

### 42HARTONDELMETAcov_byB.pdf

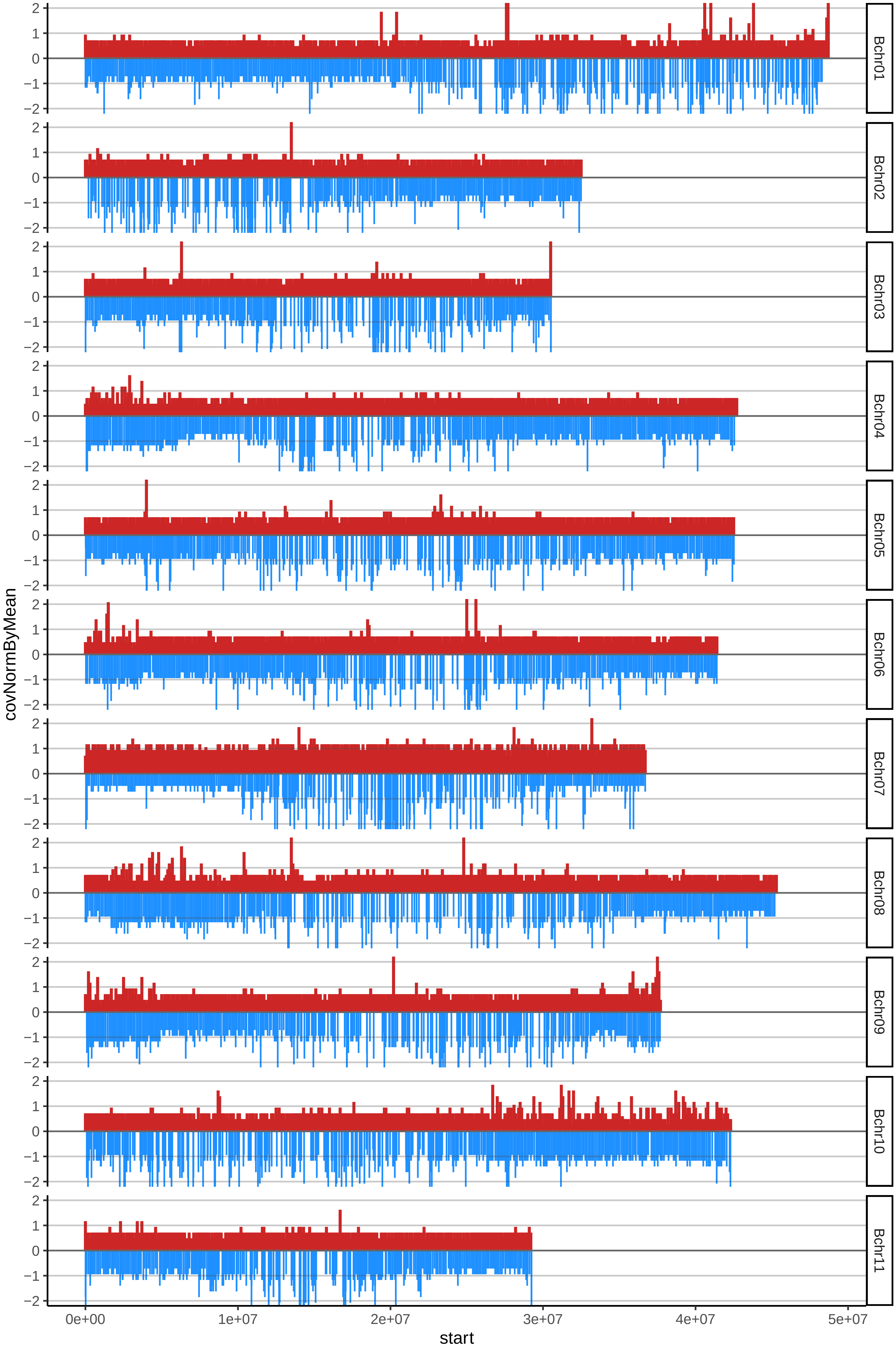

### 43HARTONHABANOcov_byA.pdf

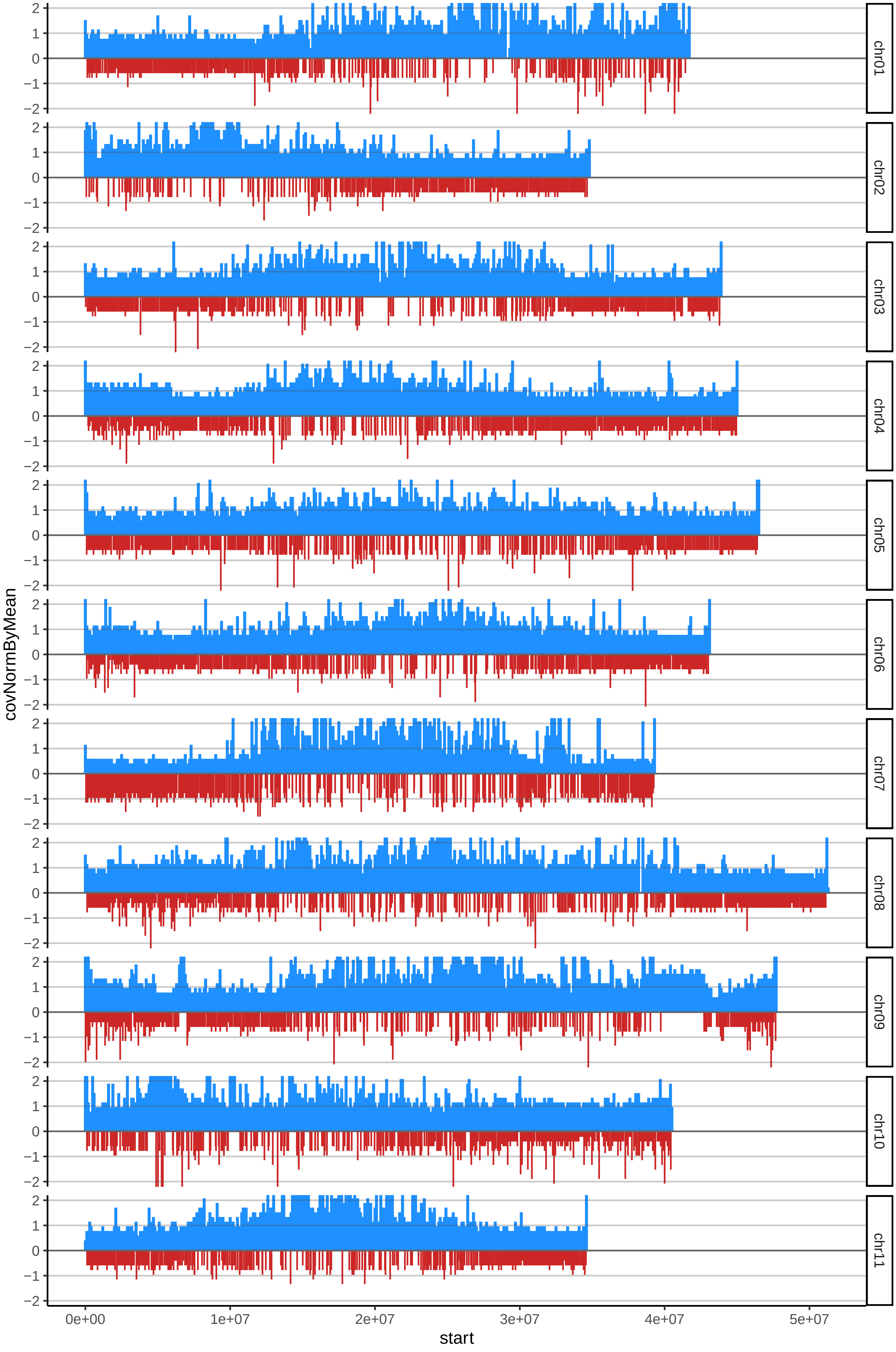

### 43HARTONHABANOcov_byB.pdf

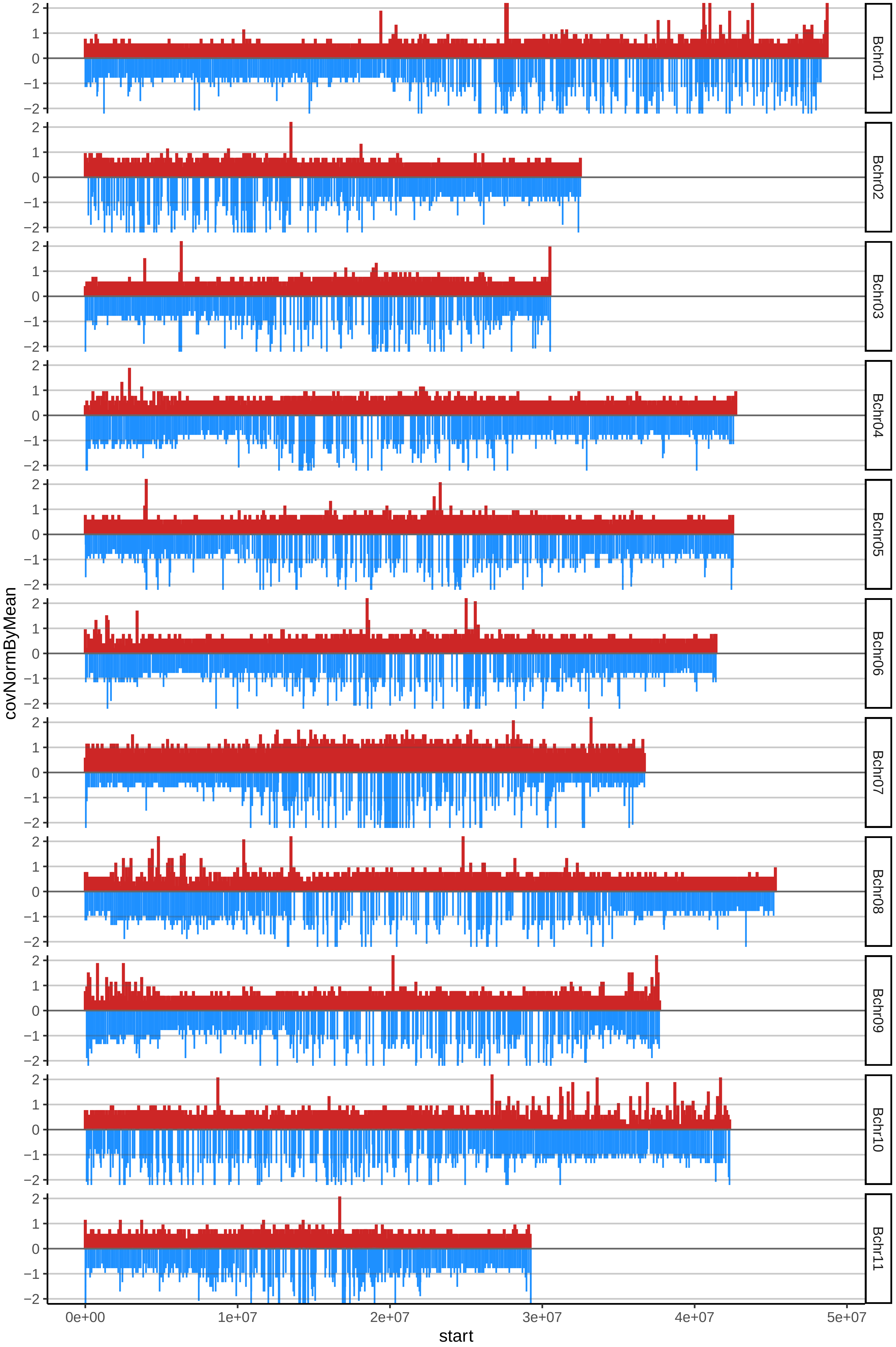

### 44HARTONMAQUEOcov_byA.pdf

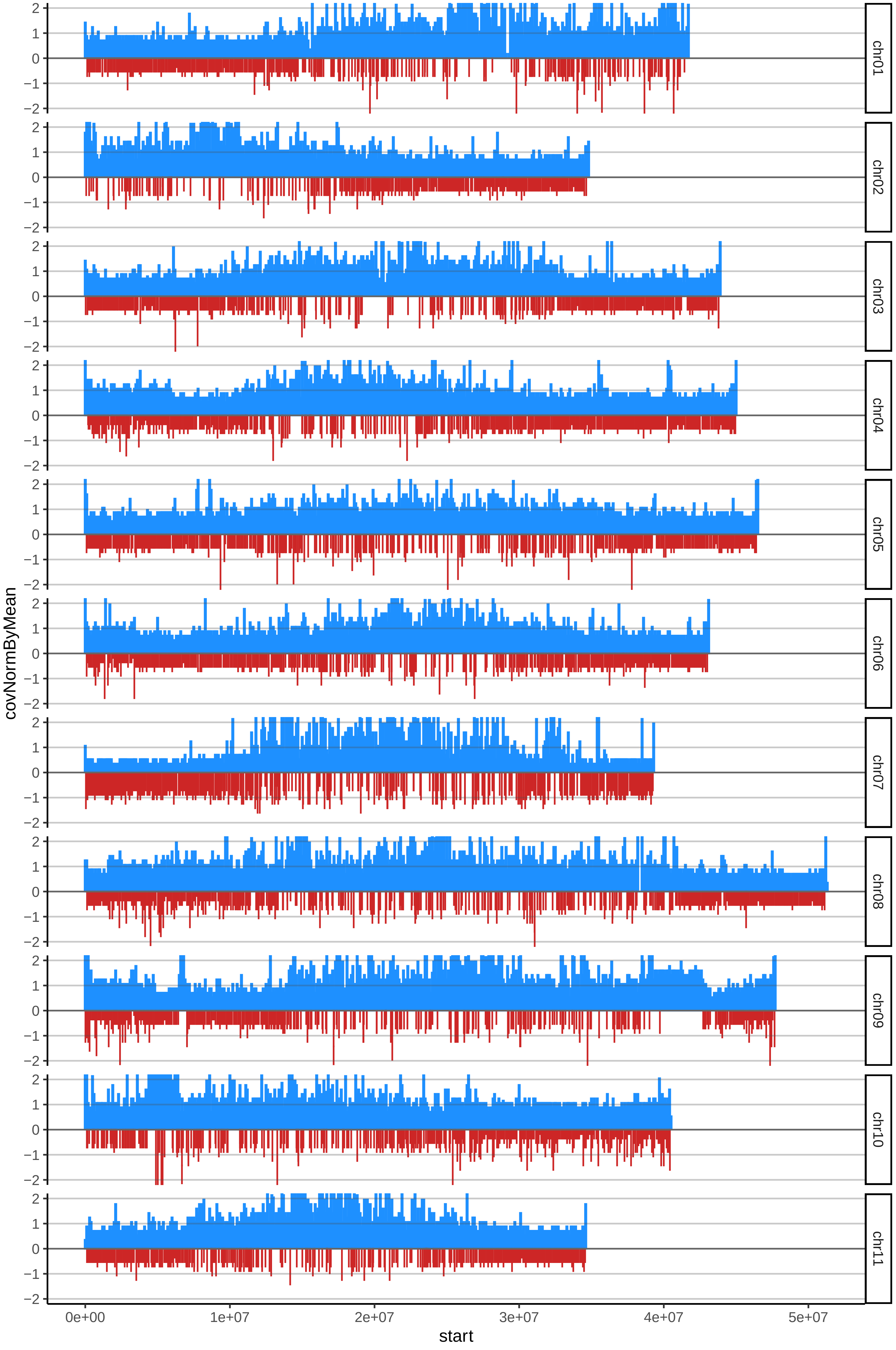

### 44HARTONMAQUEOcov_byB.pdf

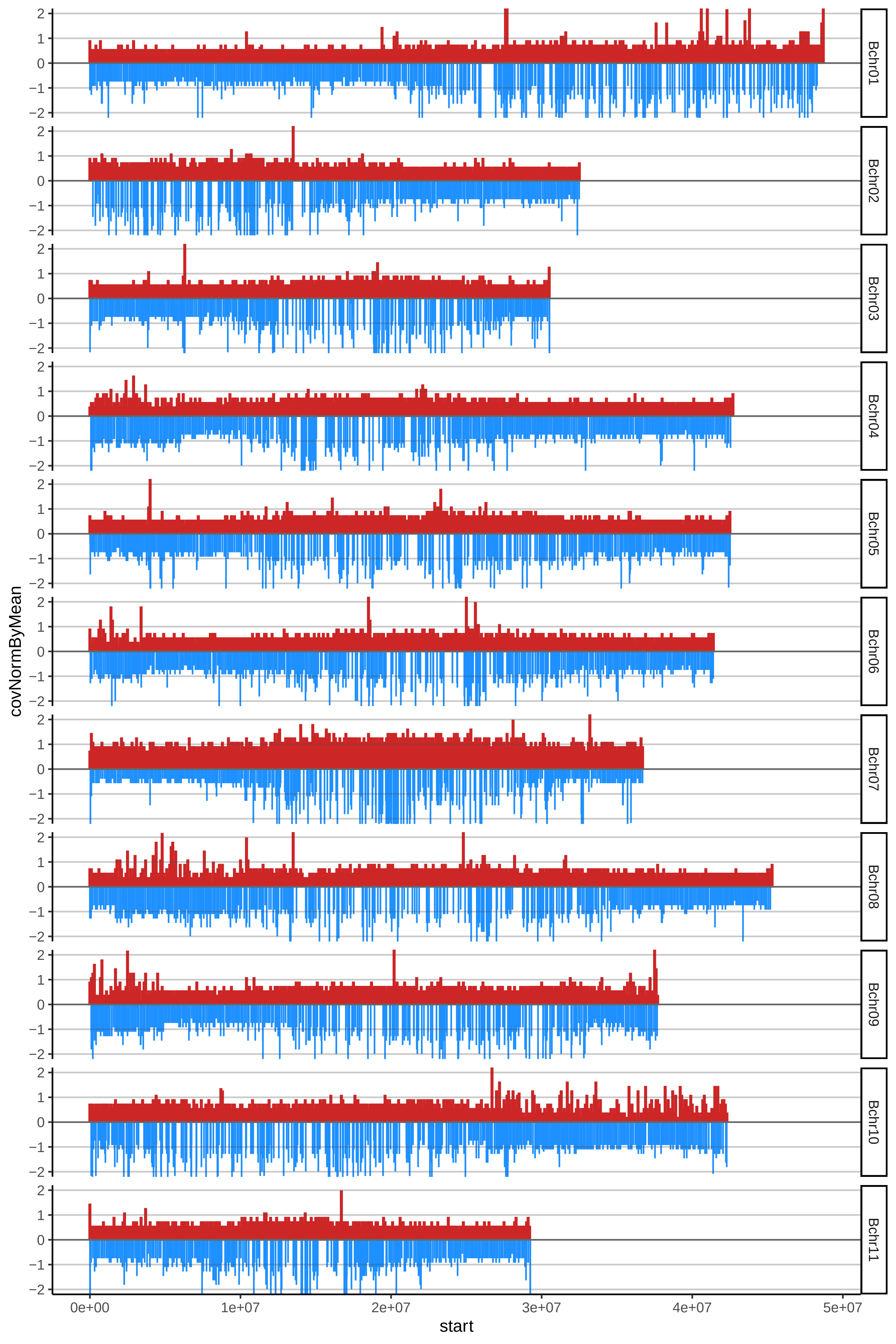

### 45HARTONPEPOcov_byA.pdf

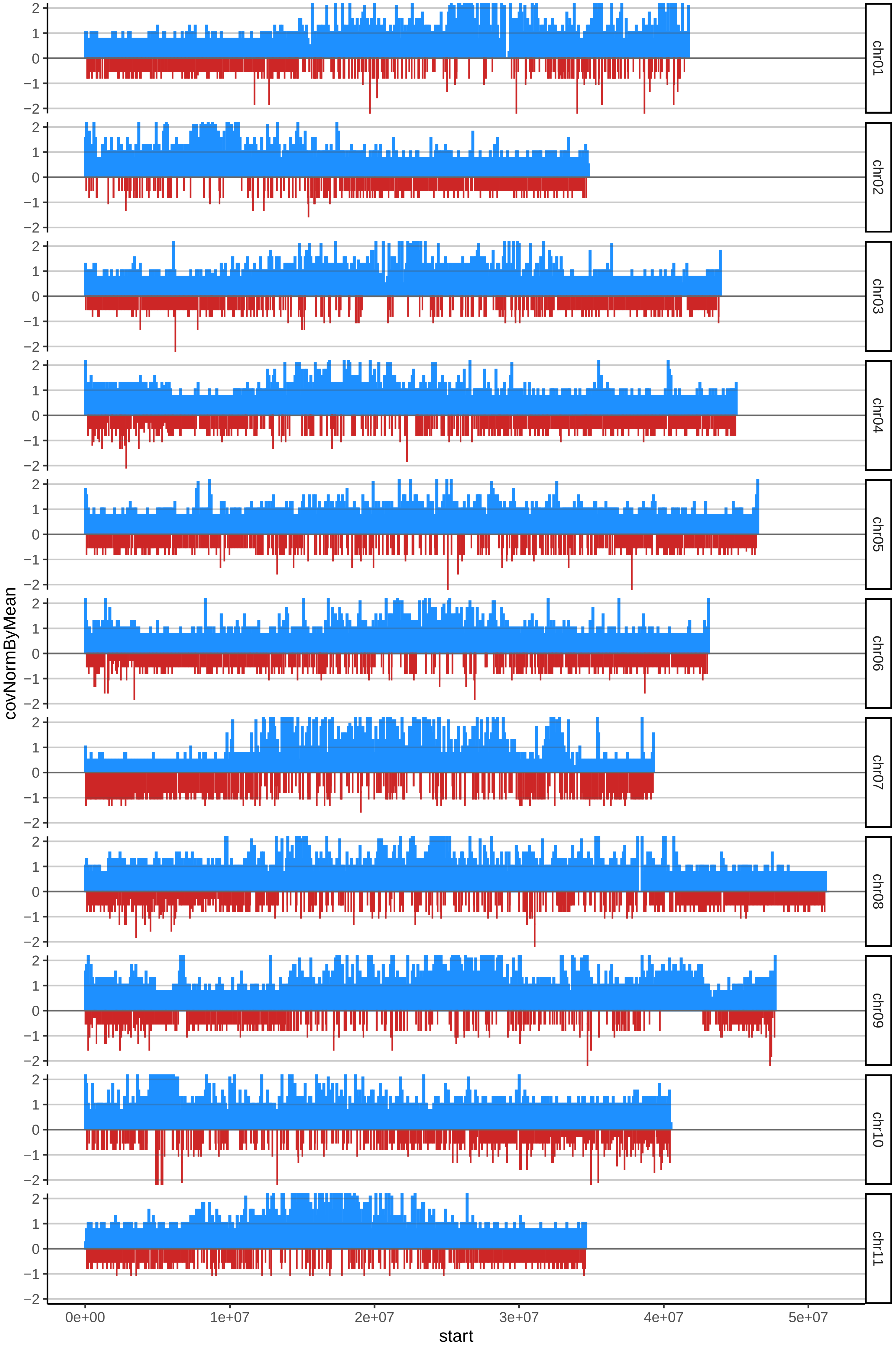

### 45HARTONPEPOcov_byB.pdf

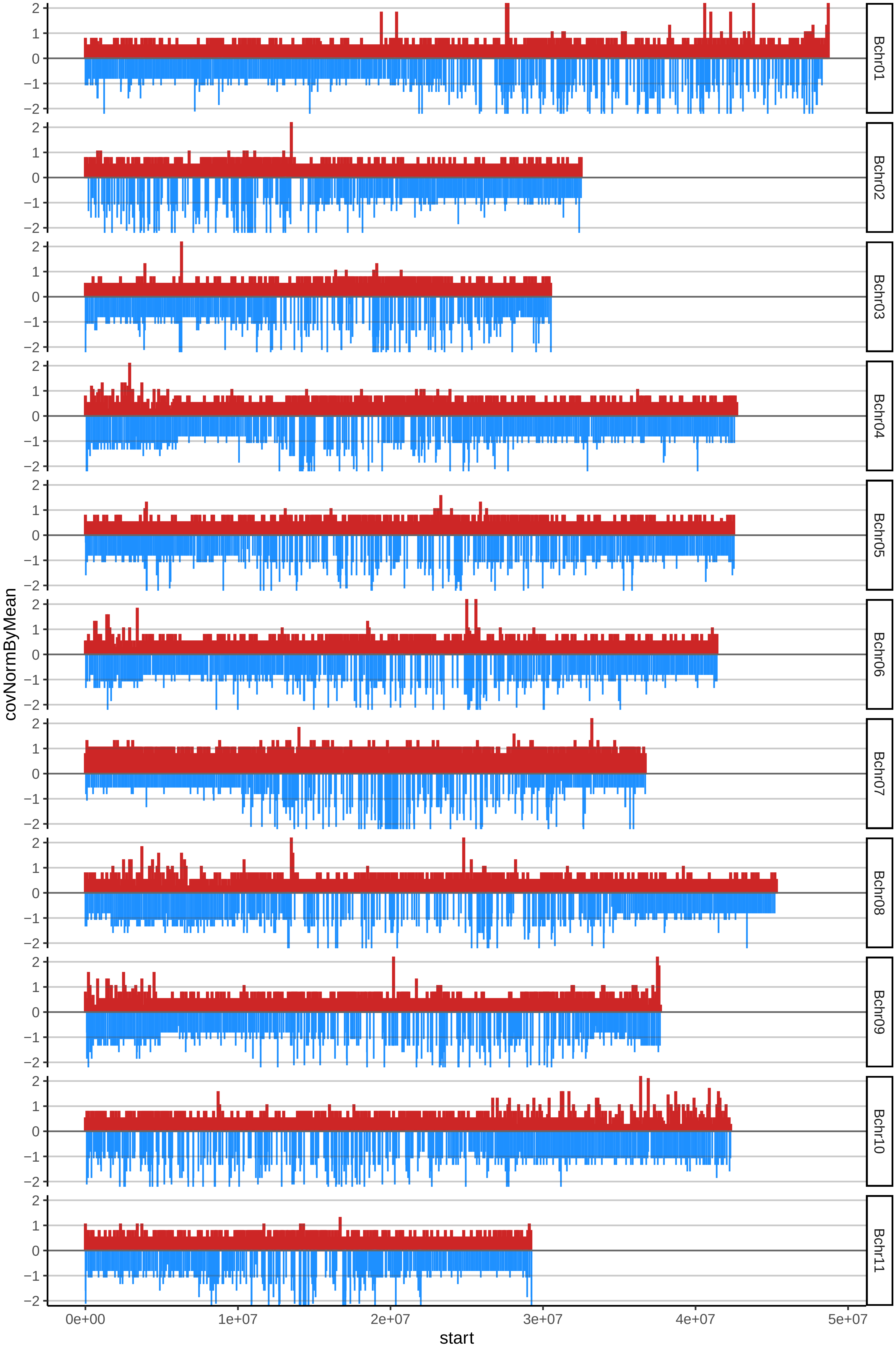

### 46HARTONROJOcov_byA.pdf

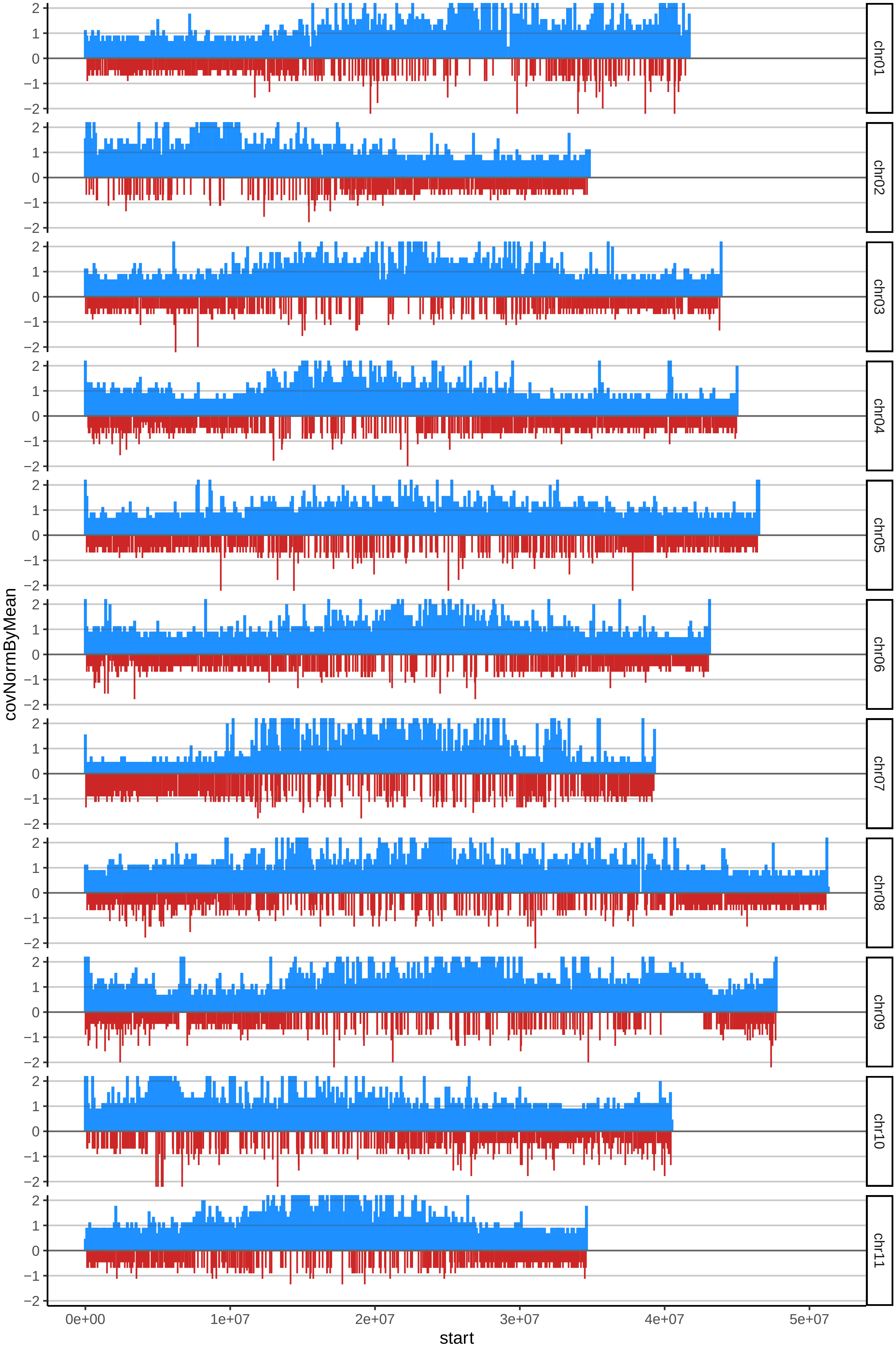

### 46HARTONROJOcov_byB.pdf

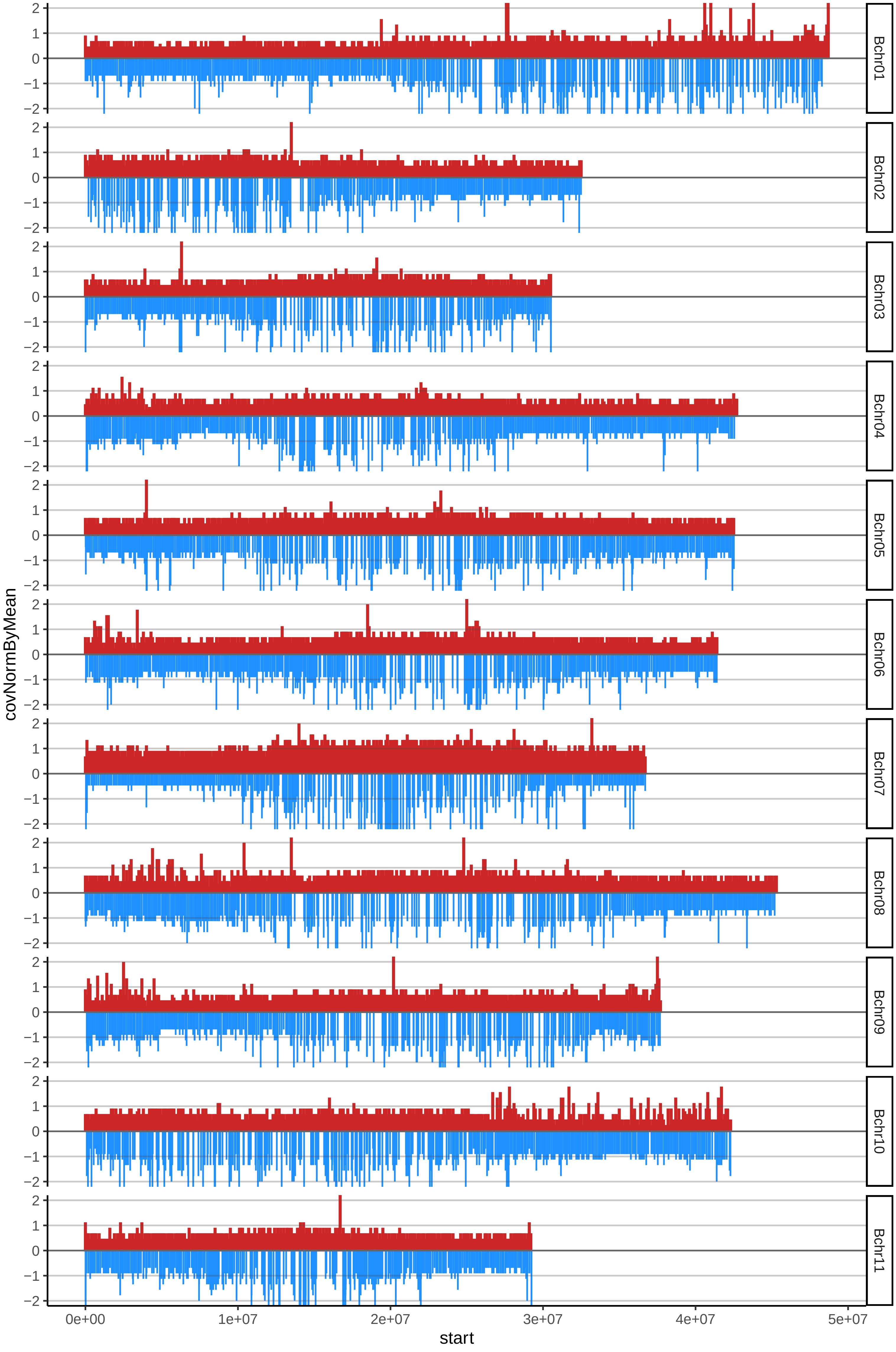

### 47HARTONTIGREcov_byA.pdf

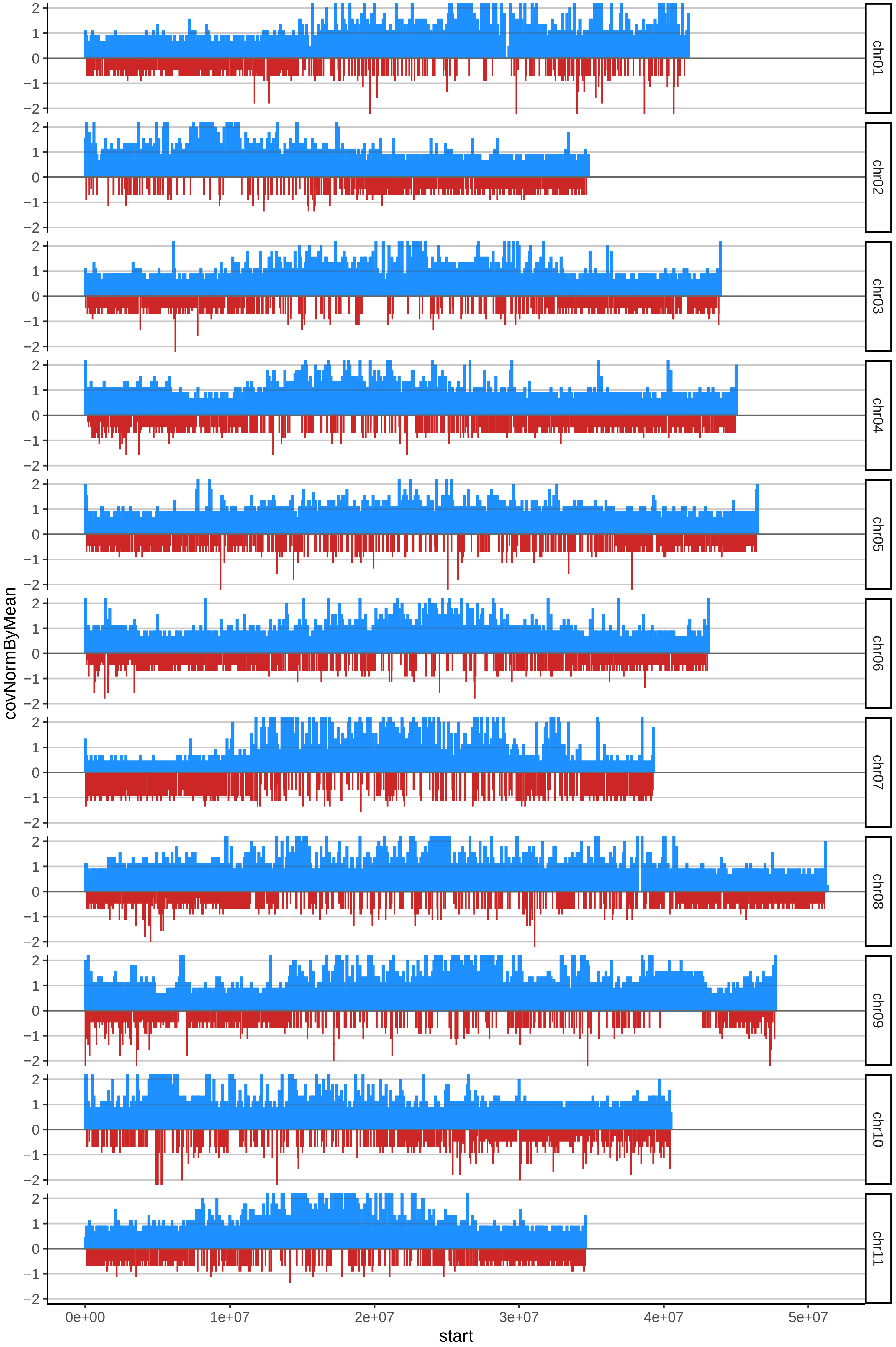

### 47HARTONTIGREcov_byB.pdf

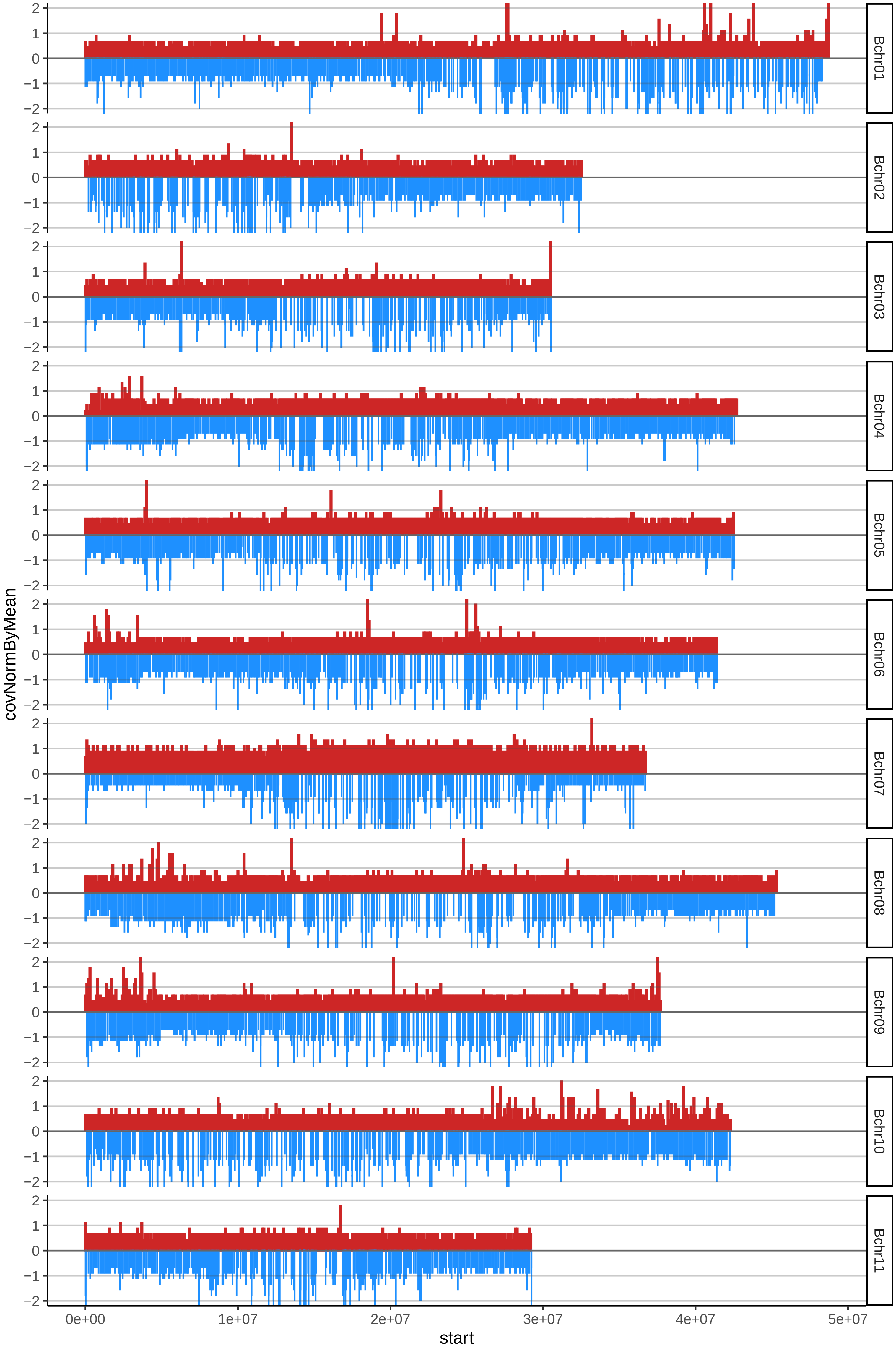

### 49HONDURENNOENANOcov_byA.pdf

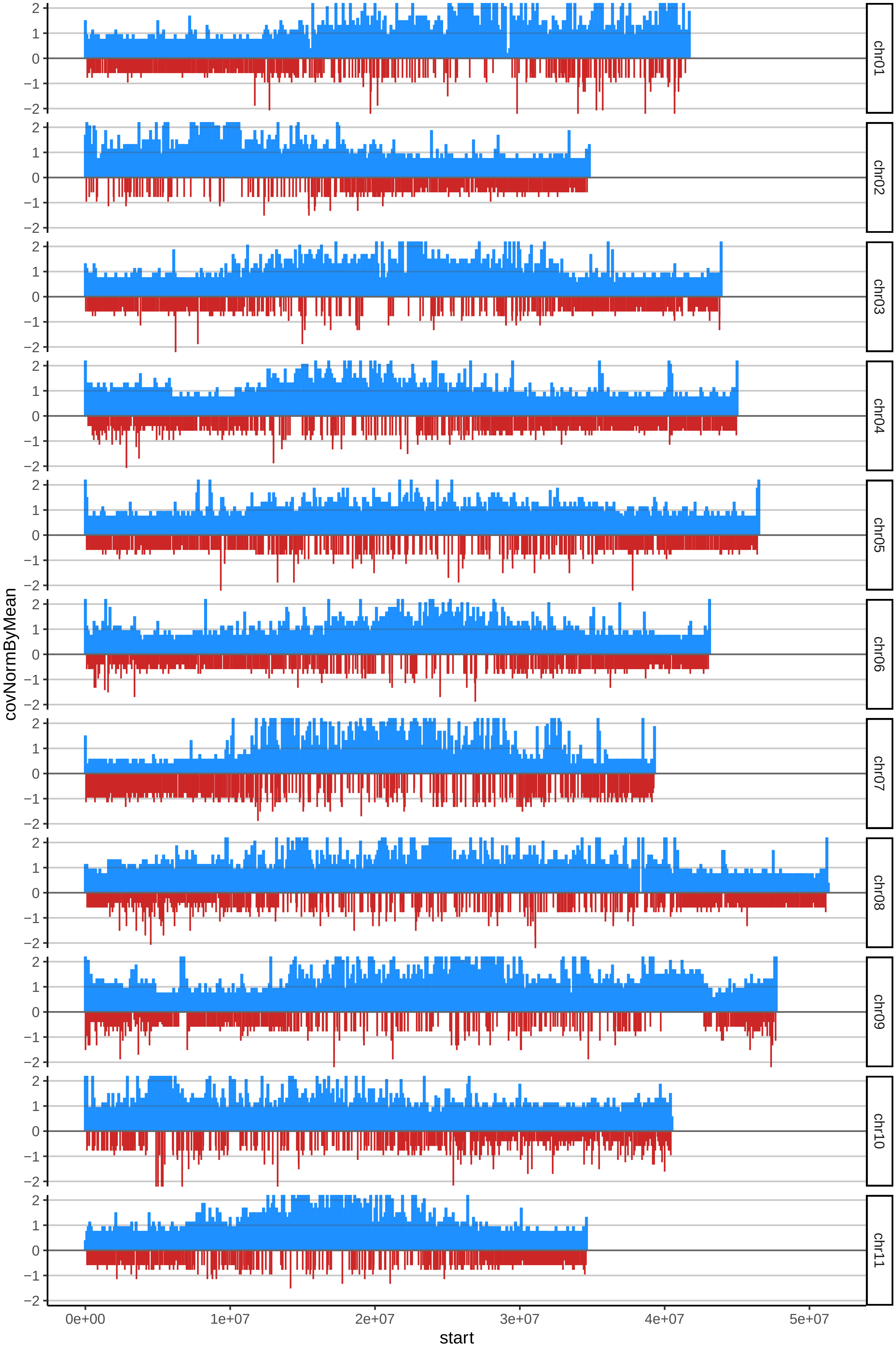

### 49HONDURENNOENANOcov_byB.pdf

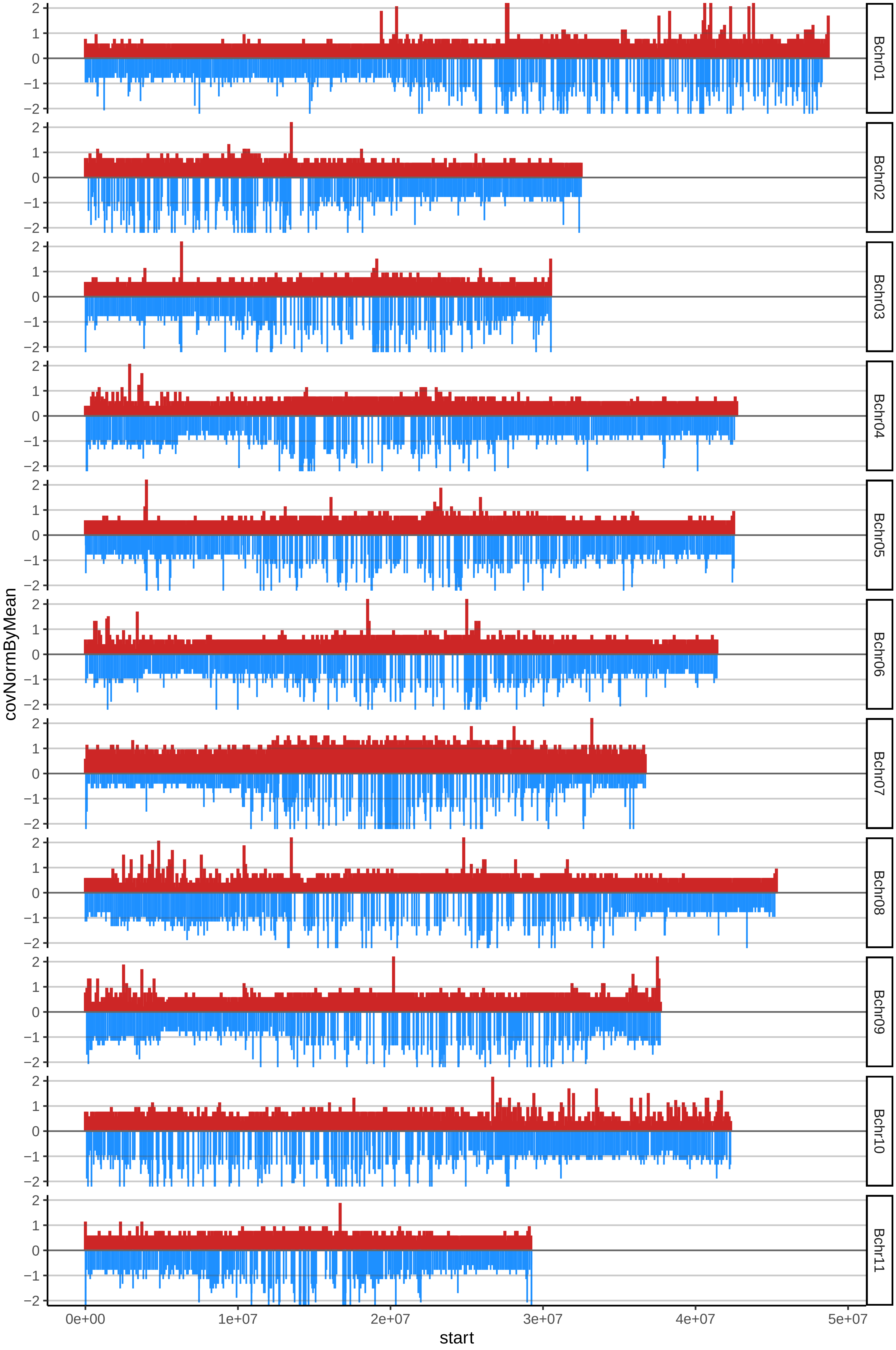

### 50KELONGMEKINTUcov_byA.pdf

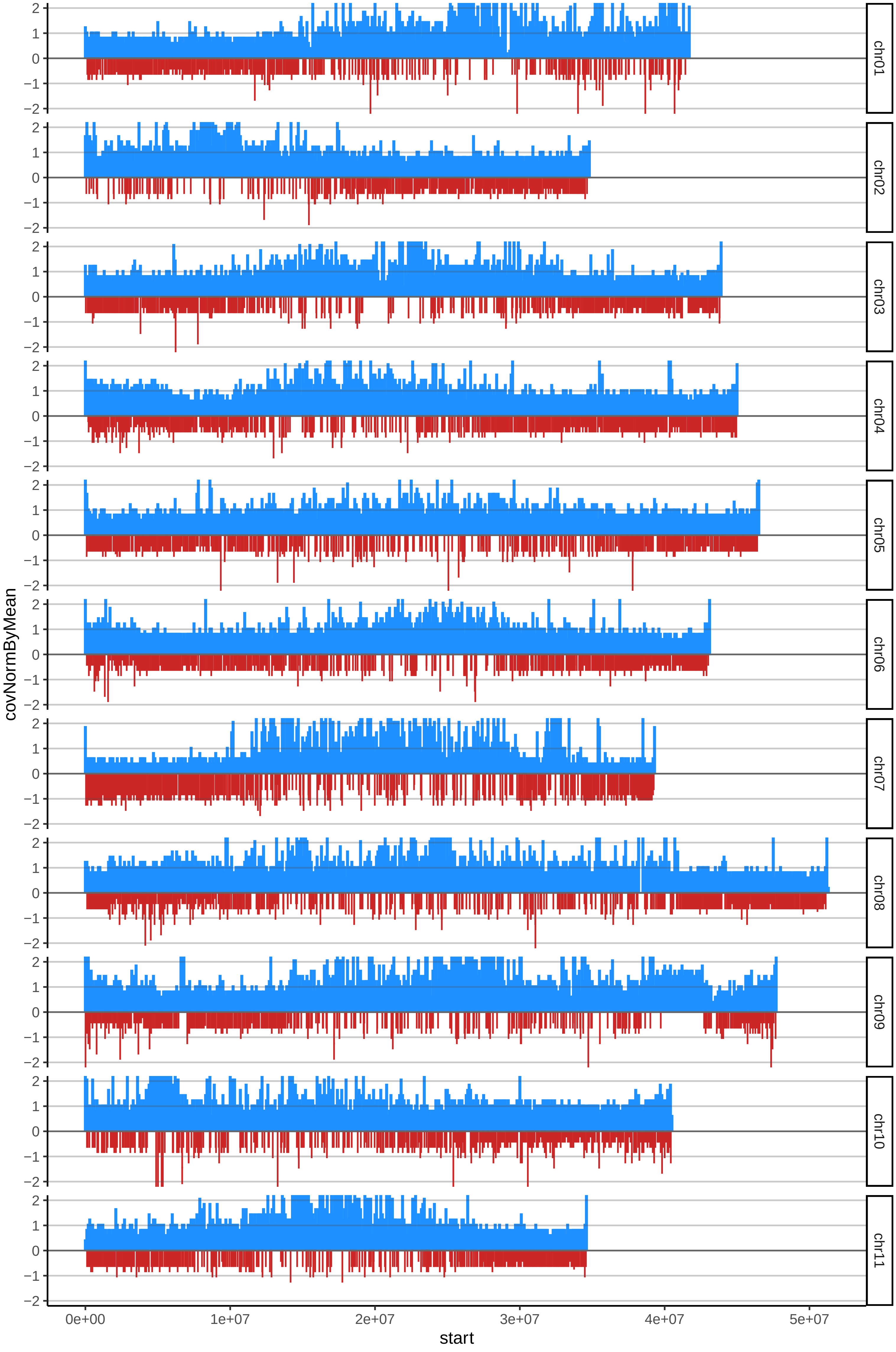

### 50KELONGMEKINTUcov_byB.pdf

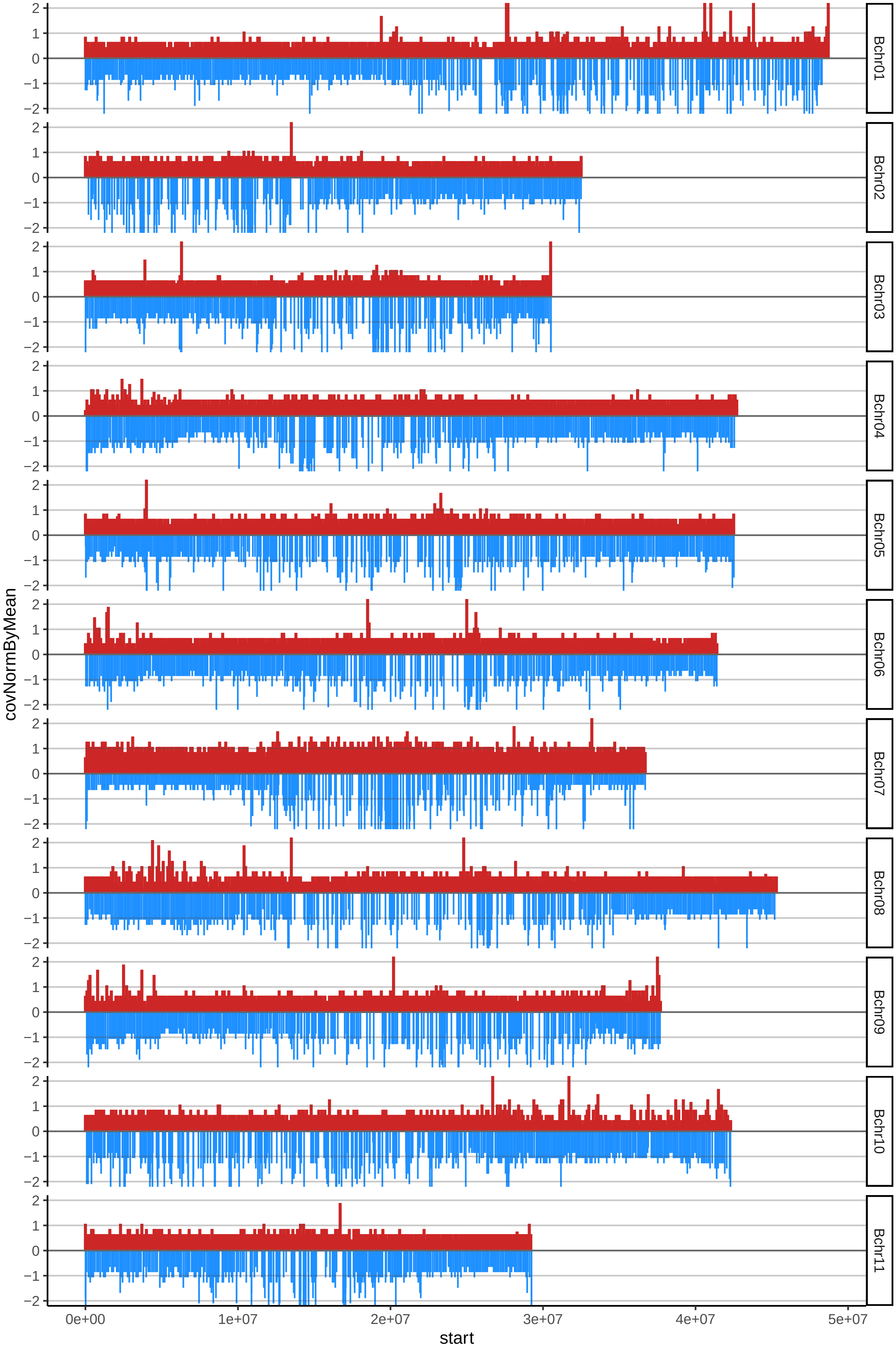

### 51KWAcov_byA.pdf

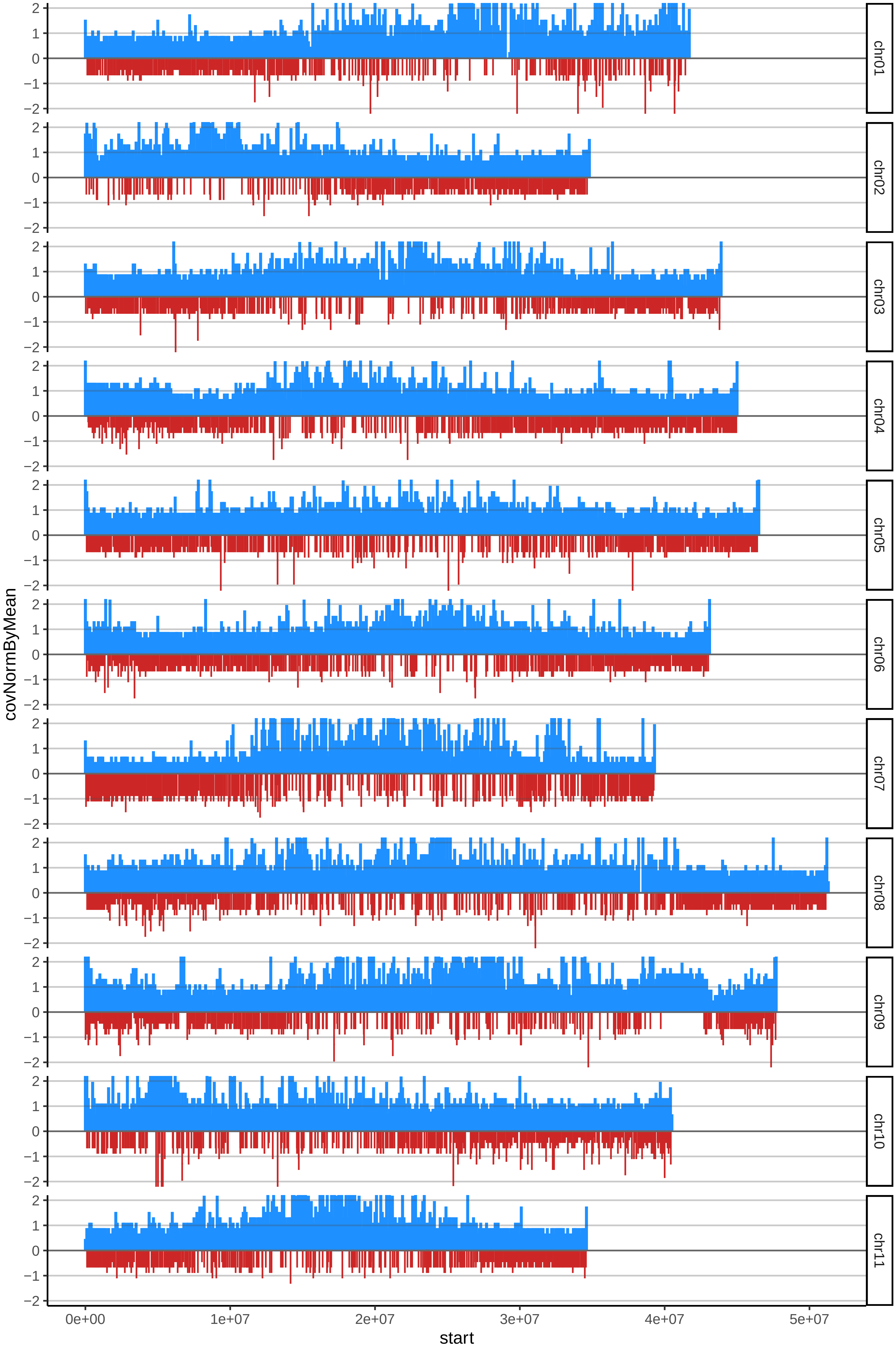

### 51KWAcov_byB.pdf

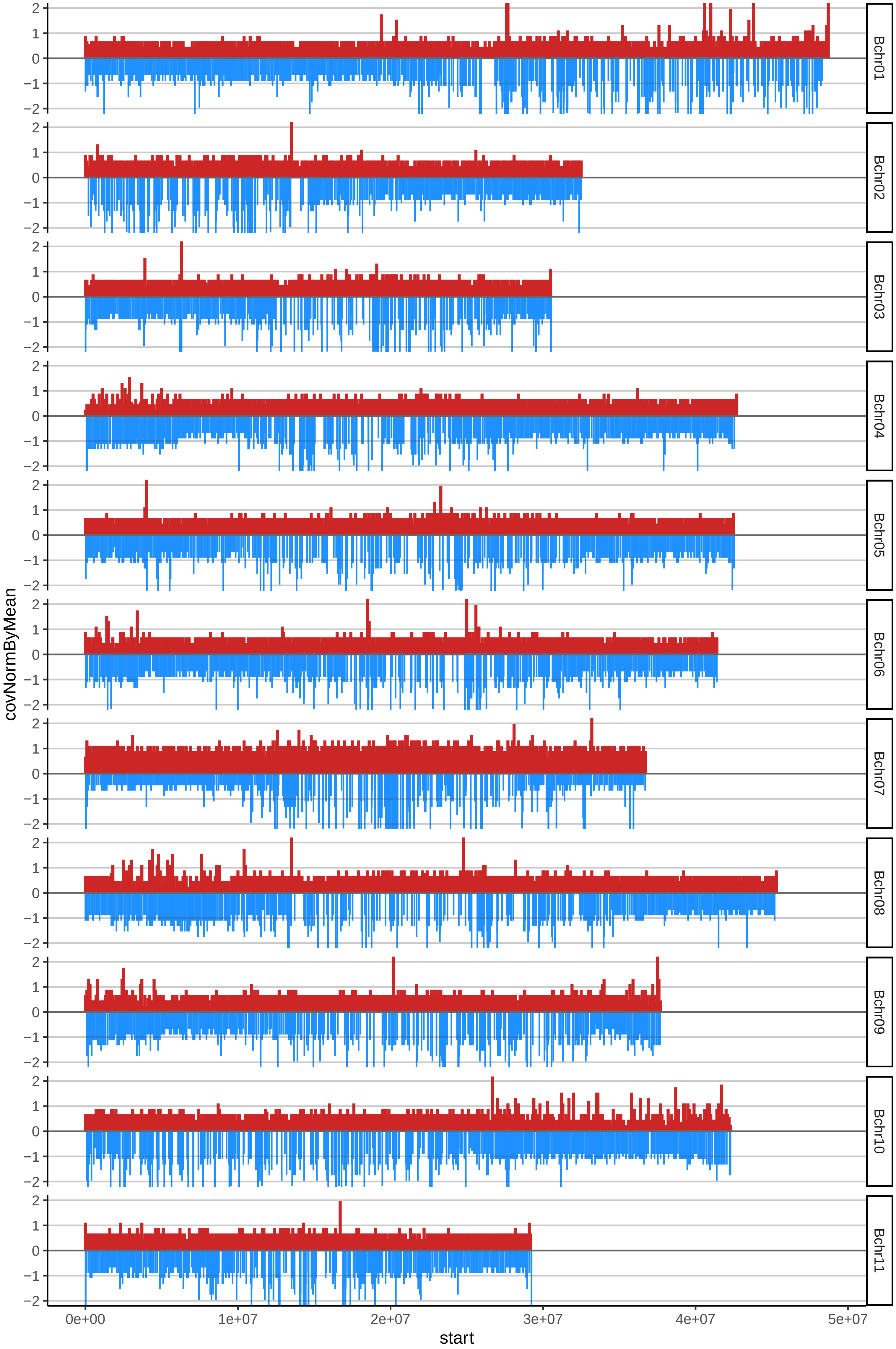

### 52LIFONGOLIKOcov_byA.pdf

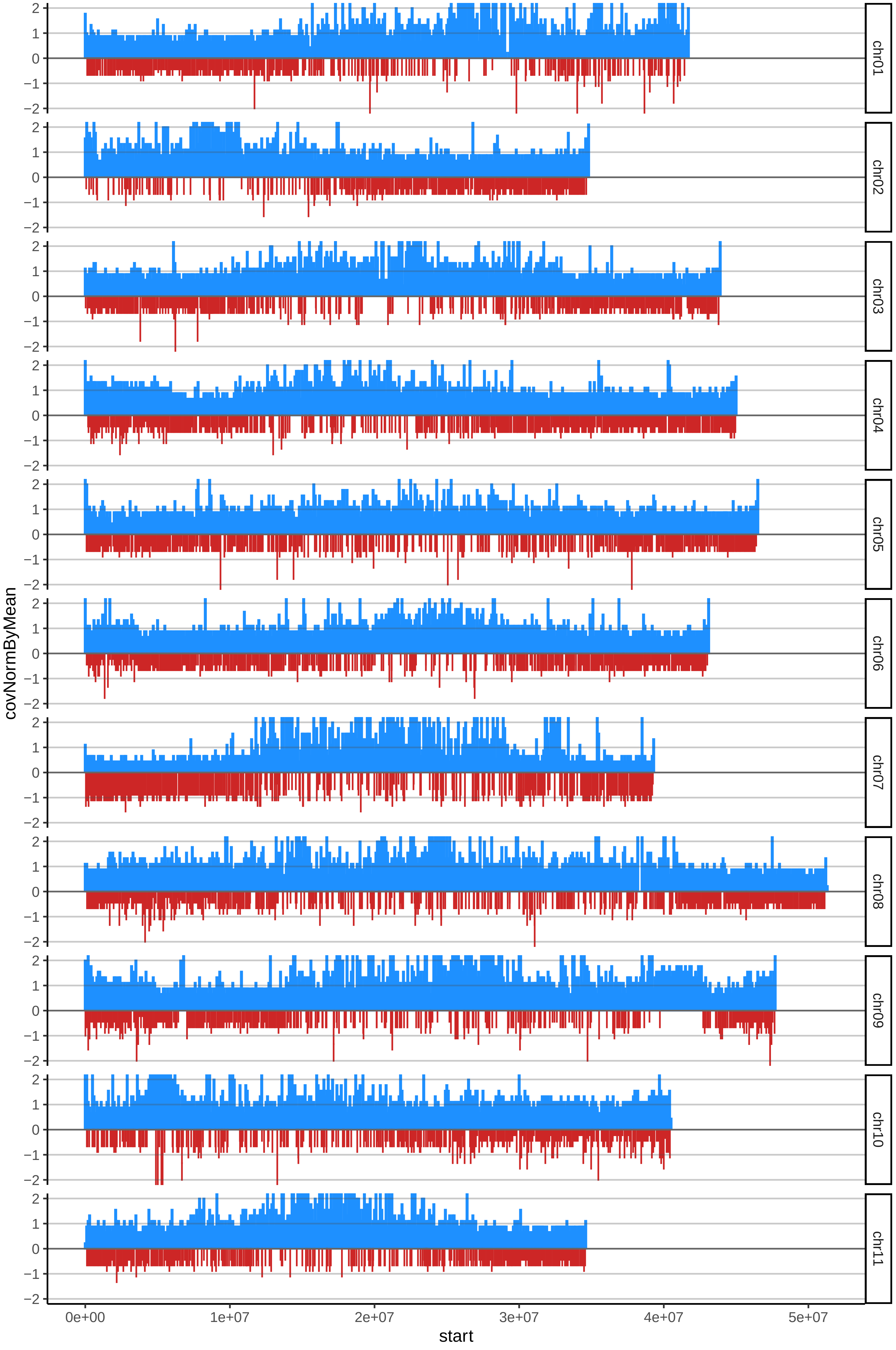

### 52LIFONGOLIKOcov_byB.pdf

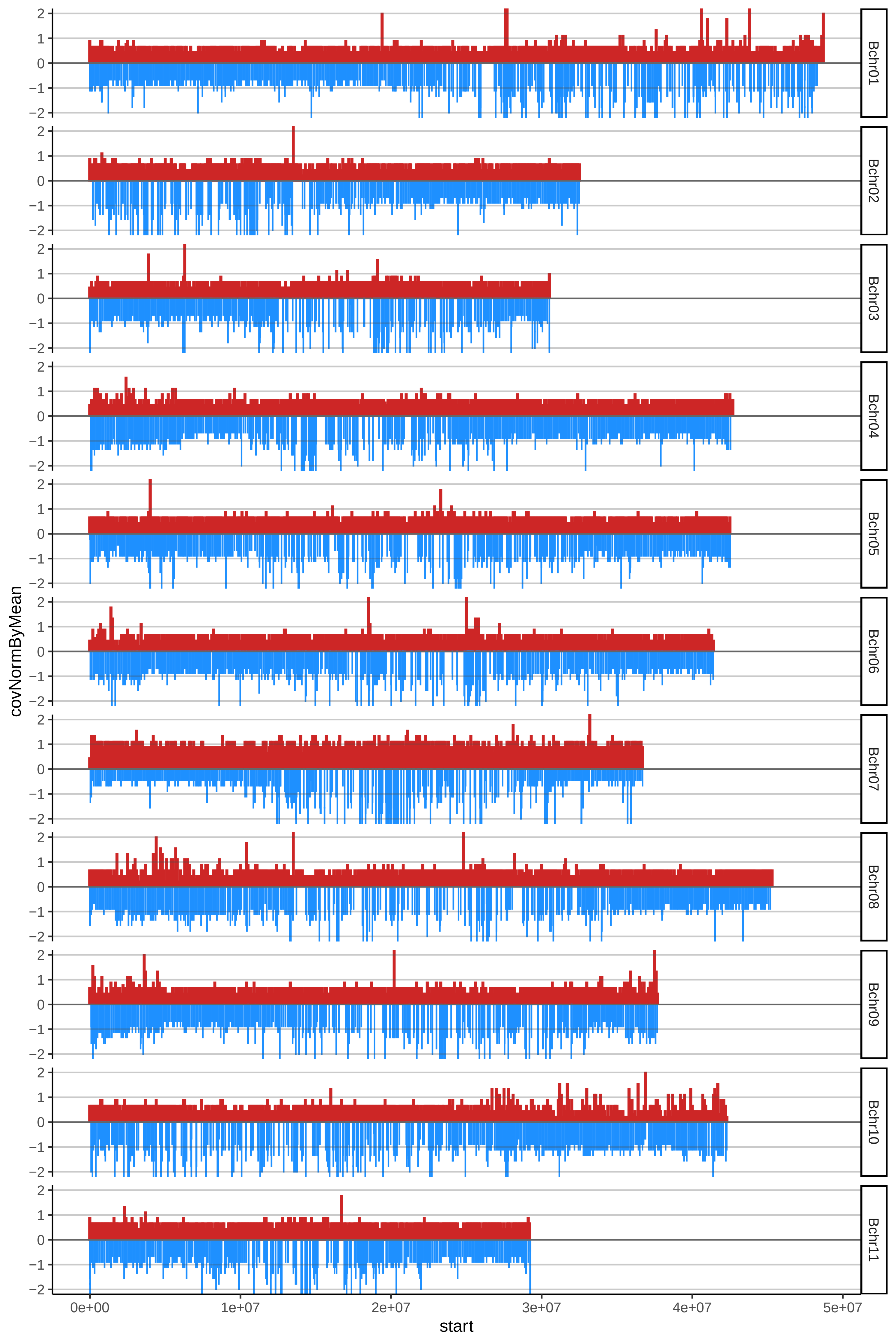

### 53MADREDELPLATANALcov_byA.pdf

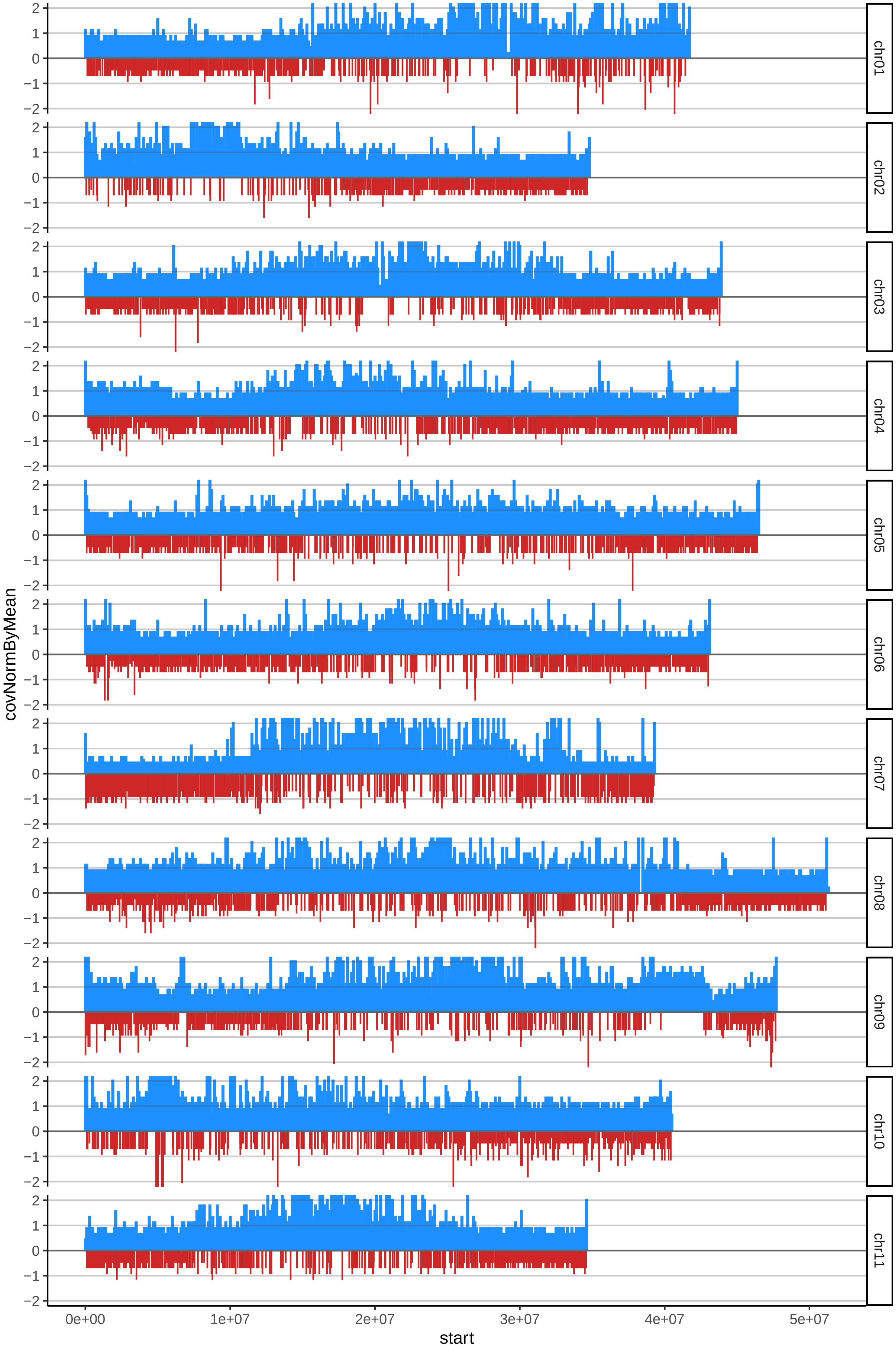

### 53MADREDELPLATANALcov_byB.pdf

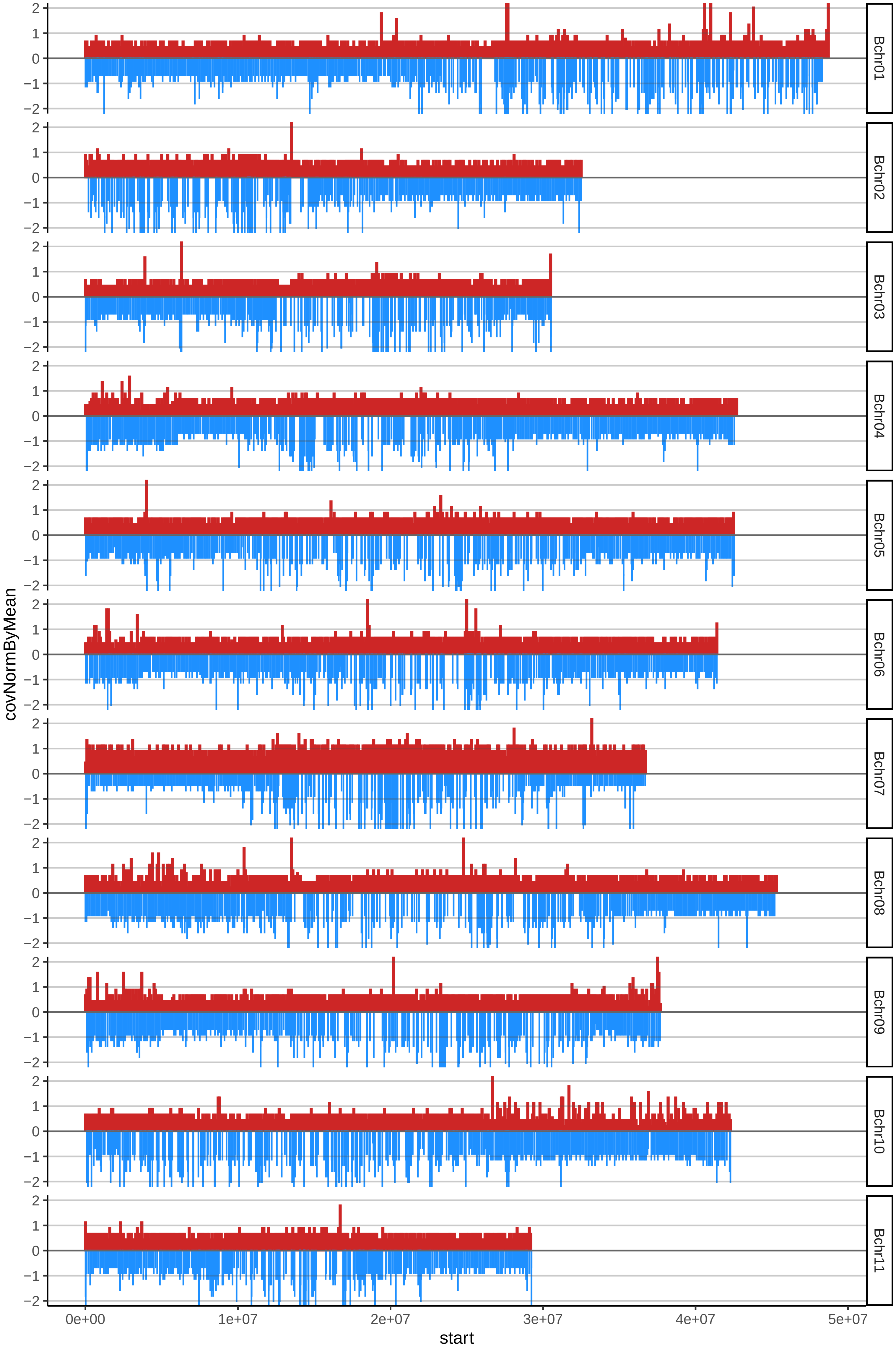

### 60MESSIATZOcov_byA.pdf

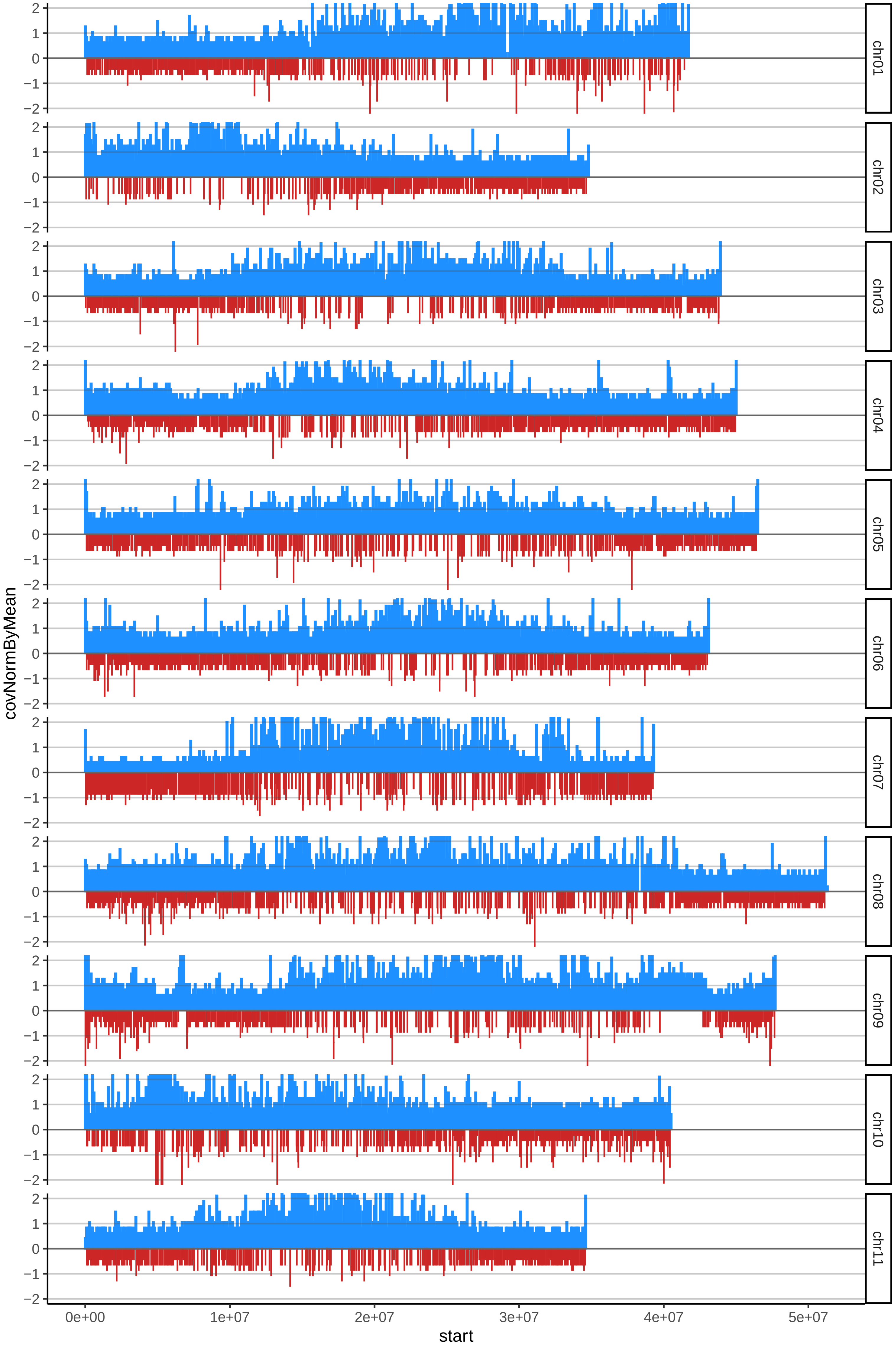

### 60MESSIATZOcov_byB.pdf

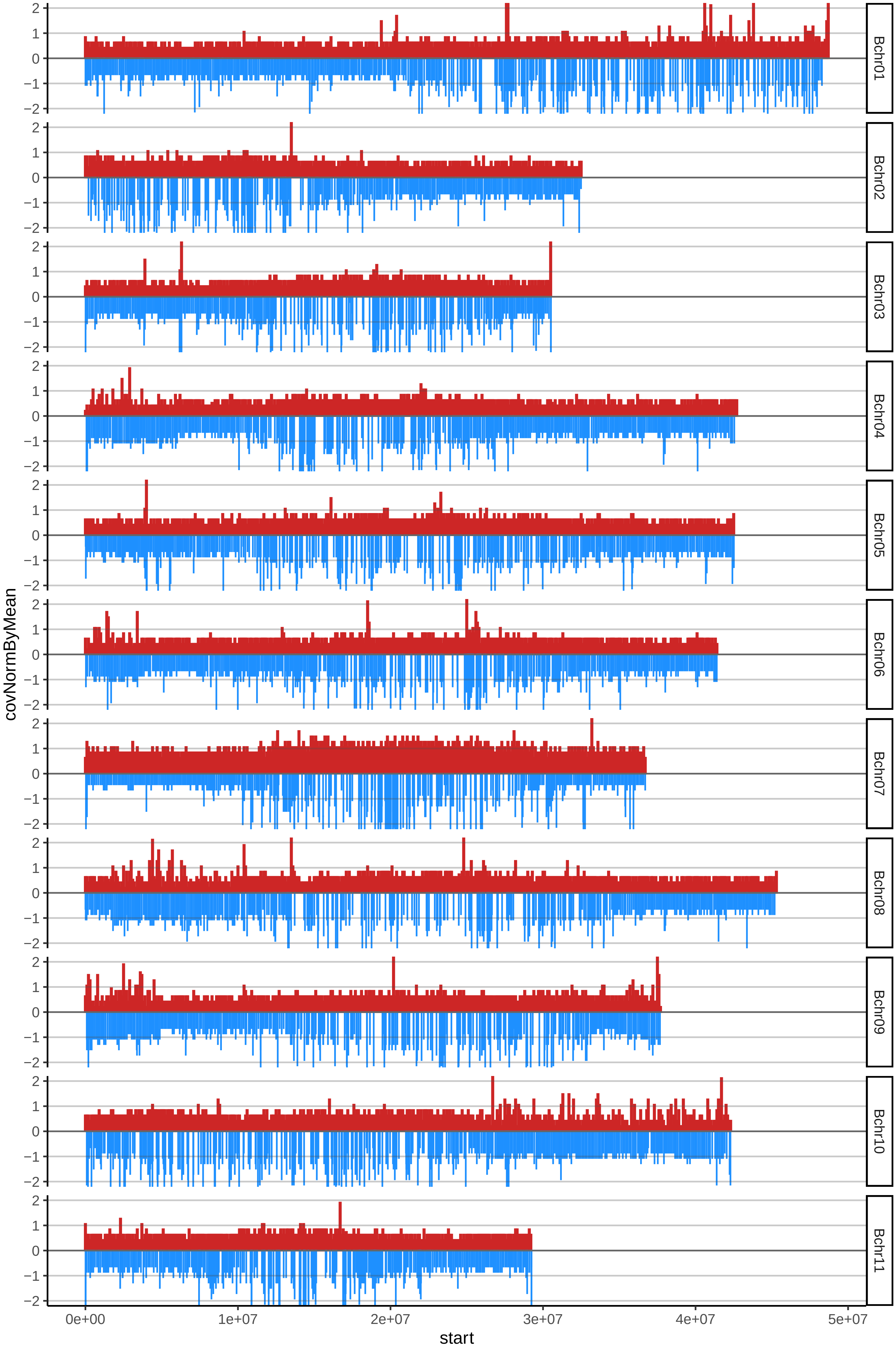

### 62NIABANGcov_byA.pdf

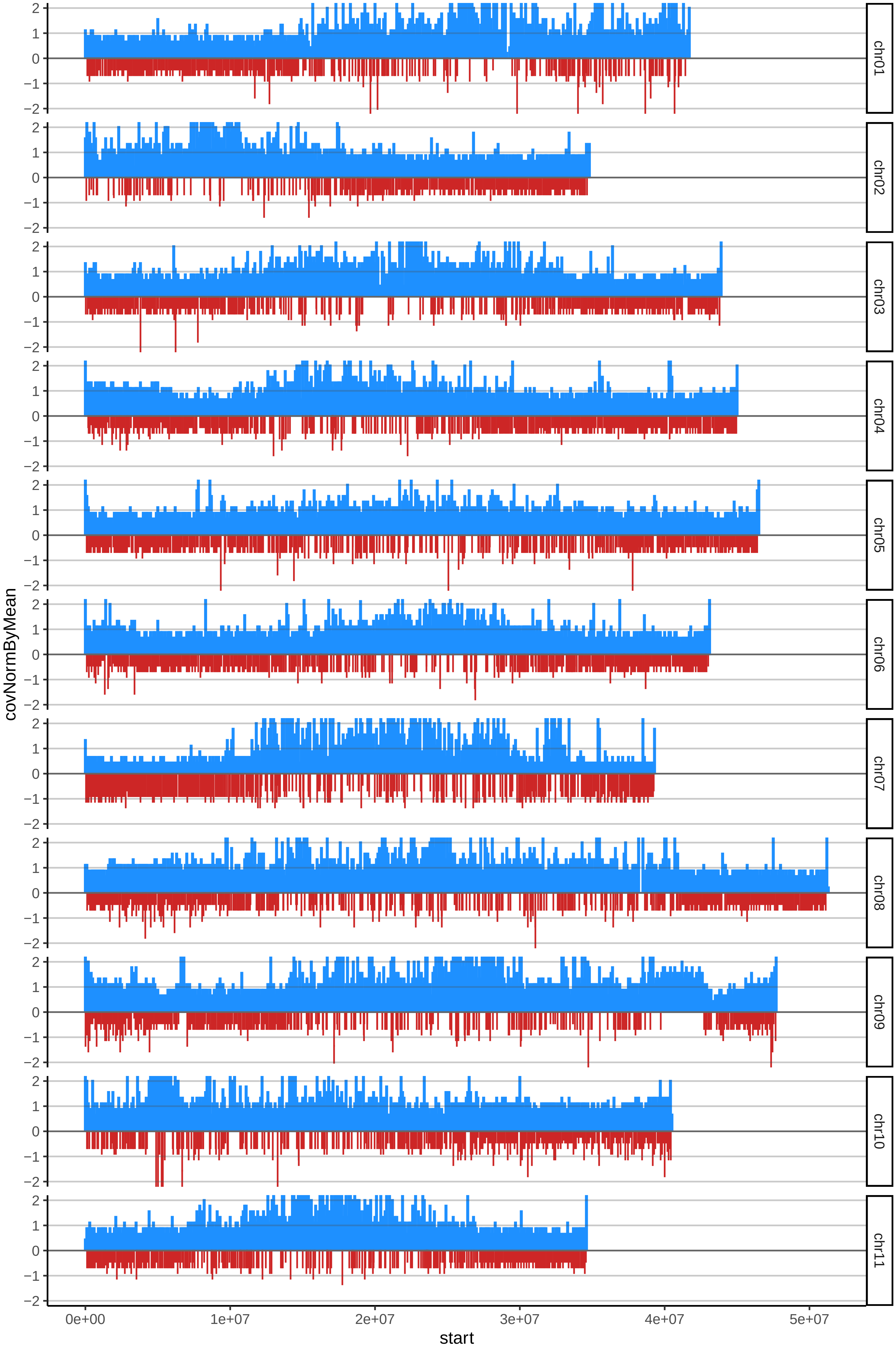

### 62NIABANGcov_byB.pdf

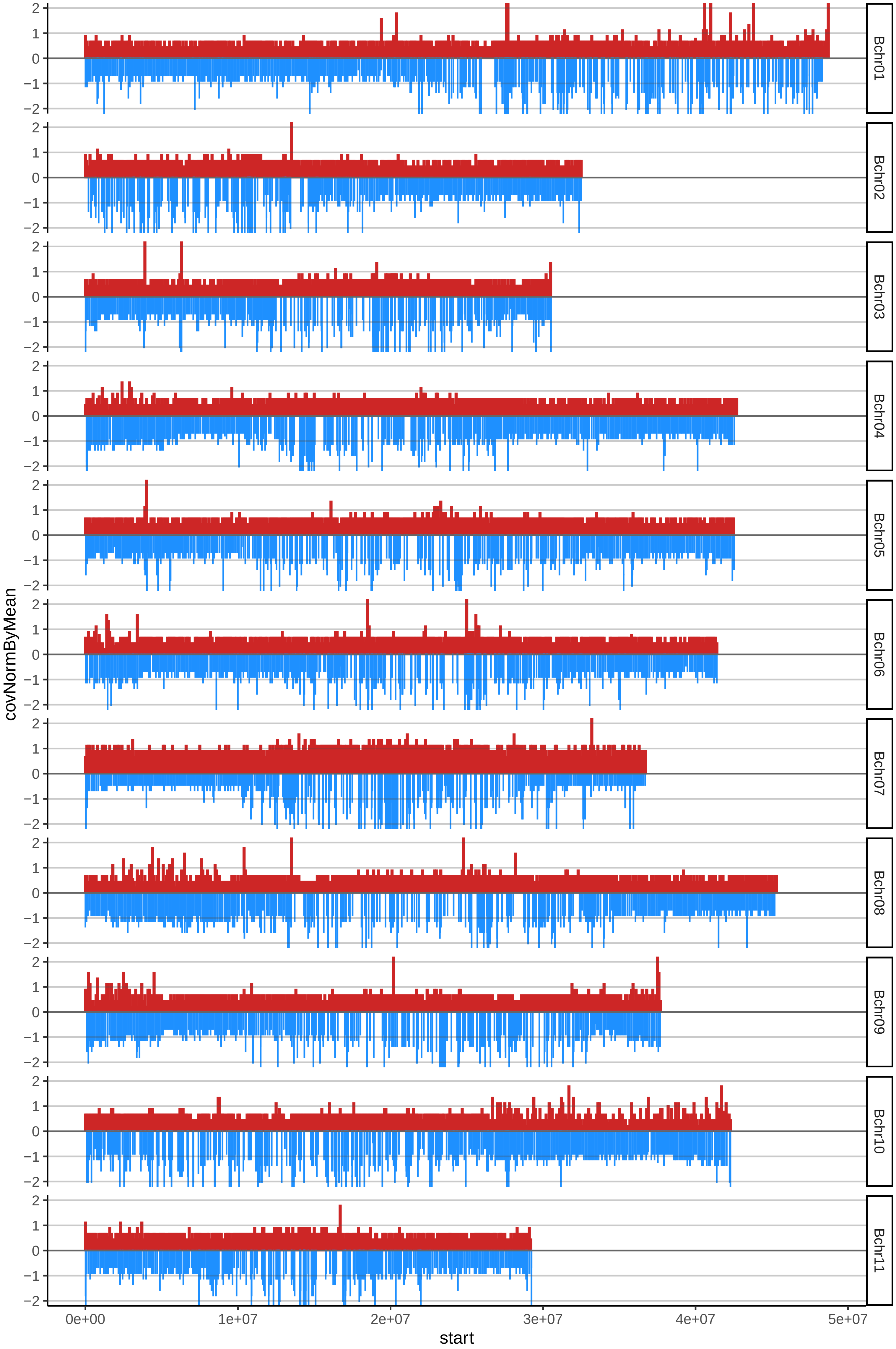

### 63NJOCKKONcov_byA.pdf

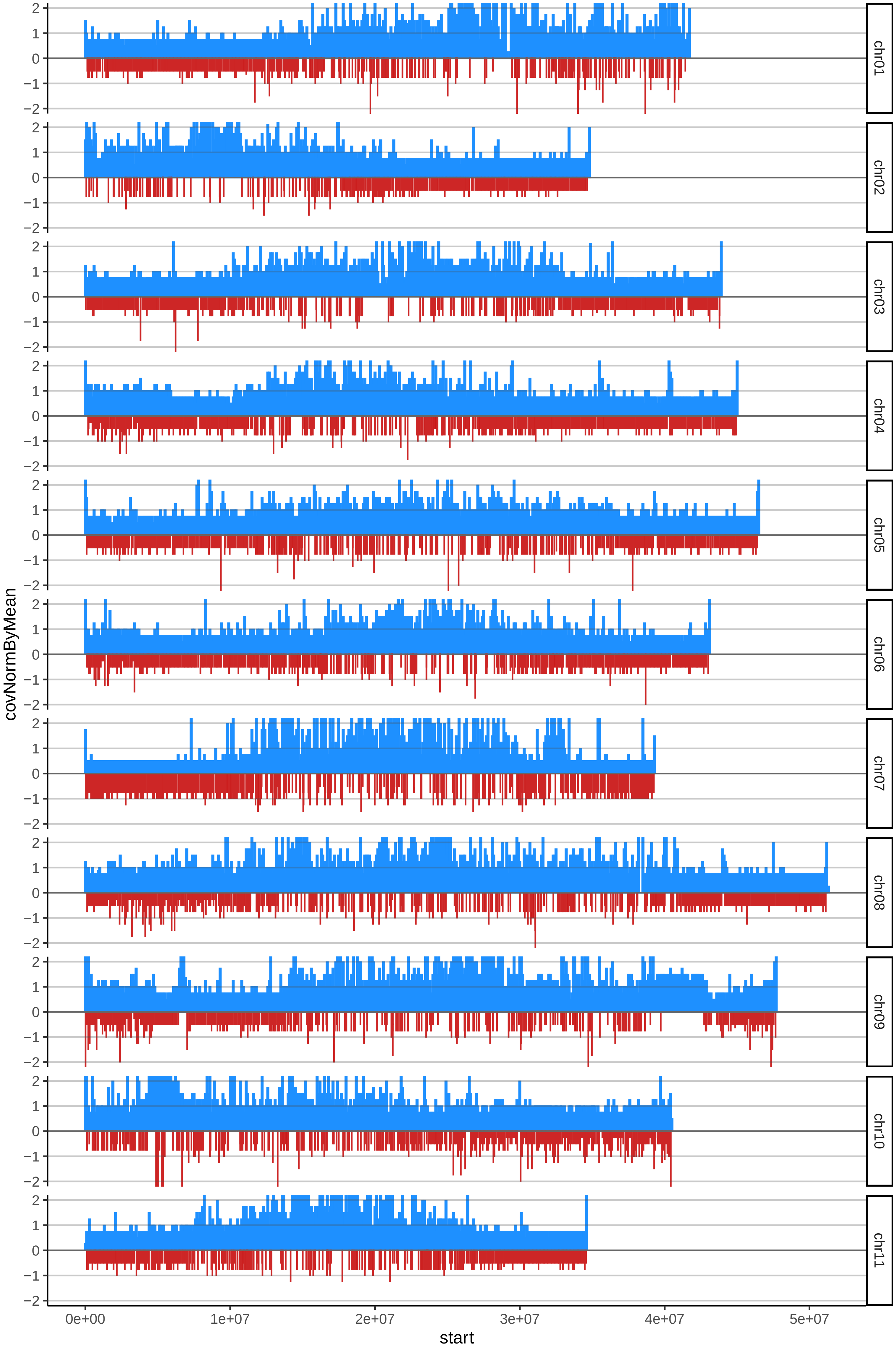

### 63NJOCKKONcov_byB.pdf

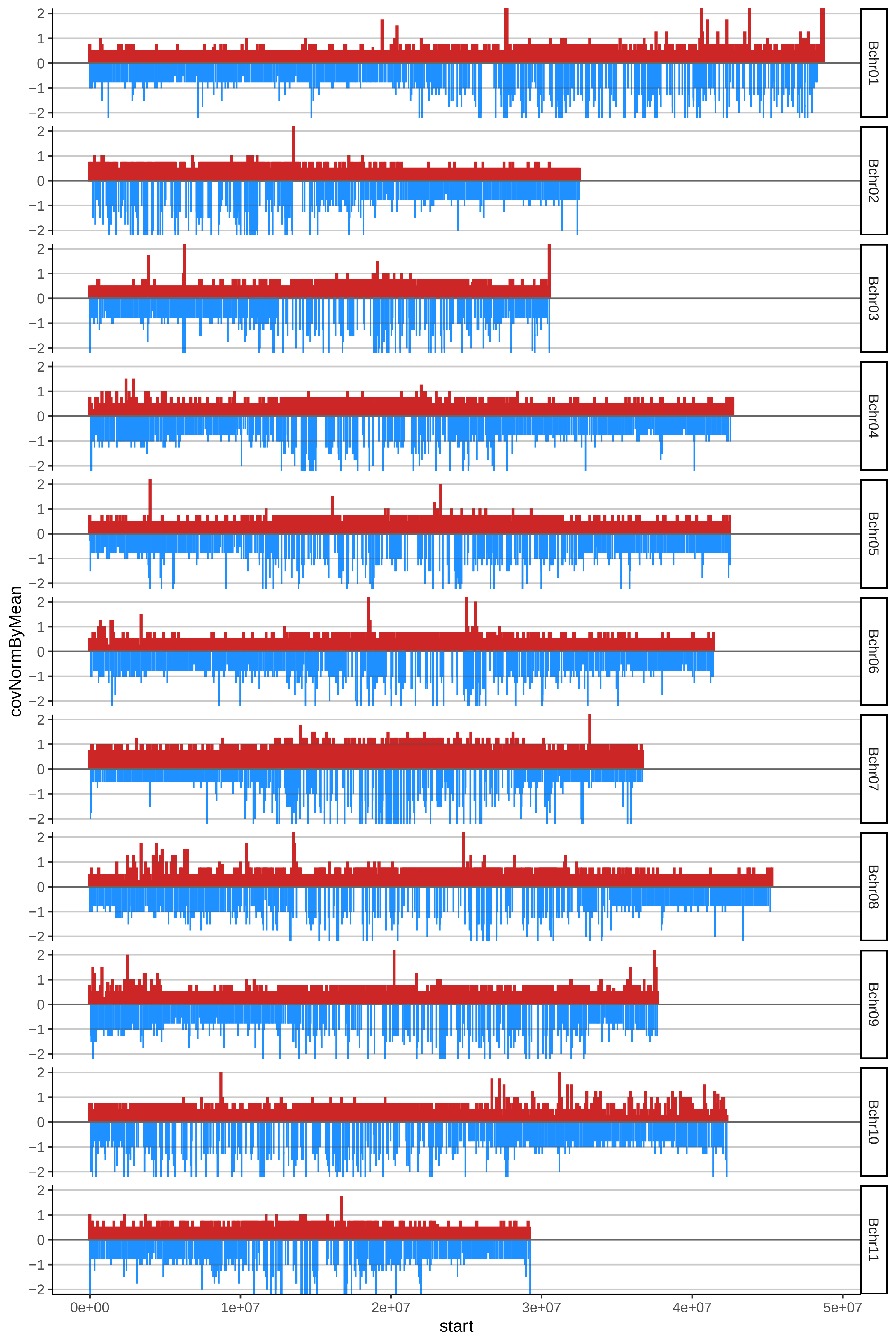

### 64ORISHELLEcov_byA.pdf

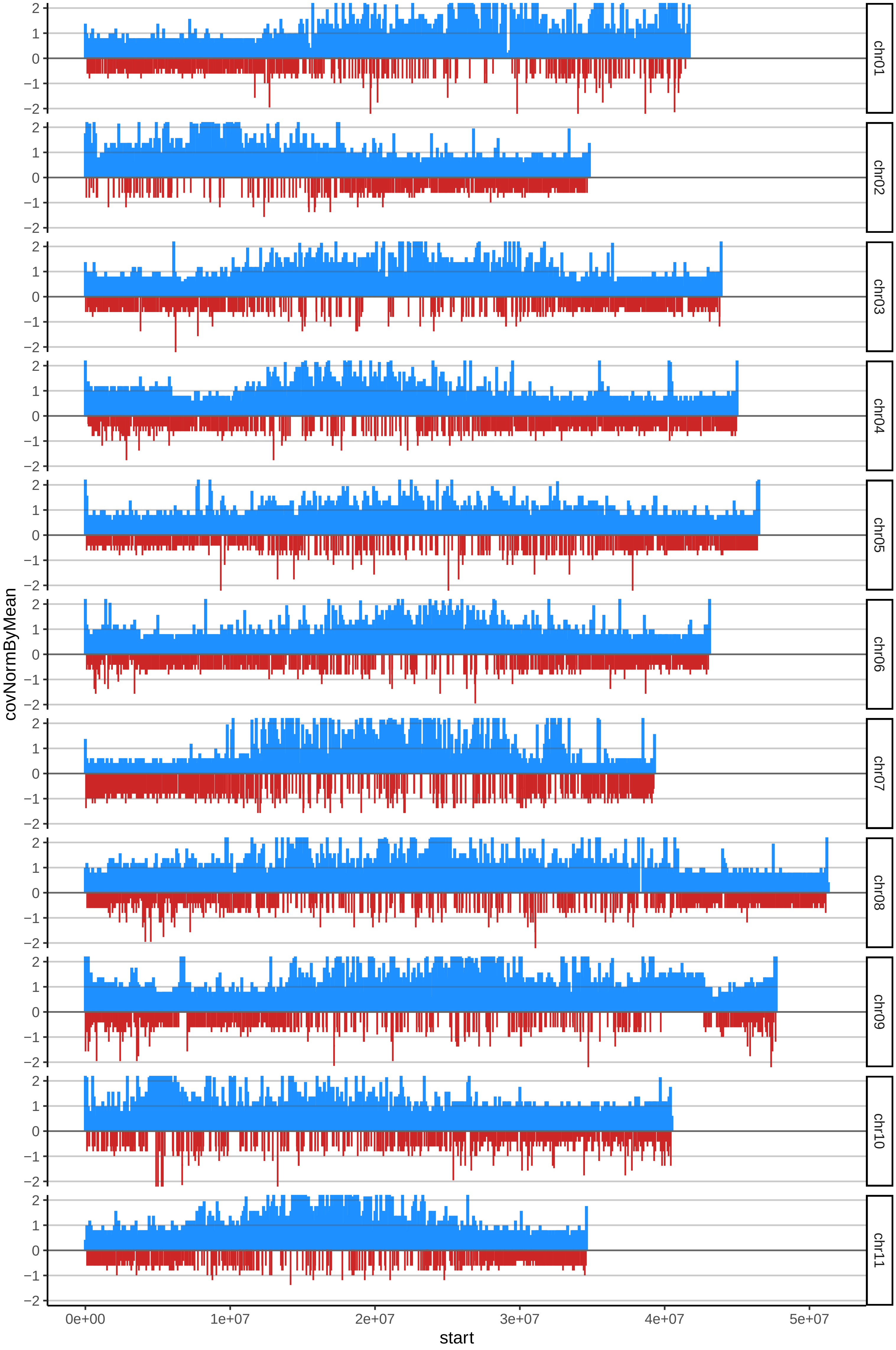

### 64ORISHELLEcov_byB.pdf

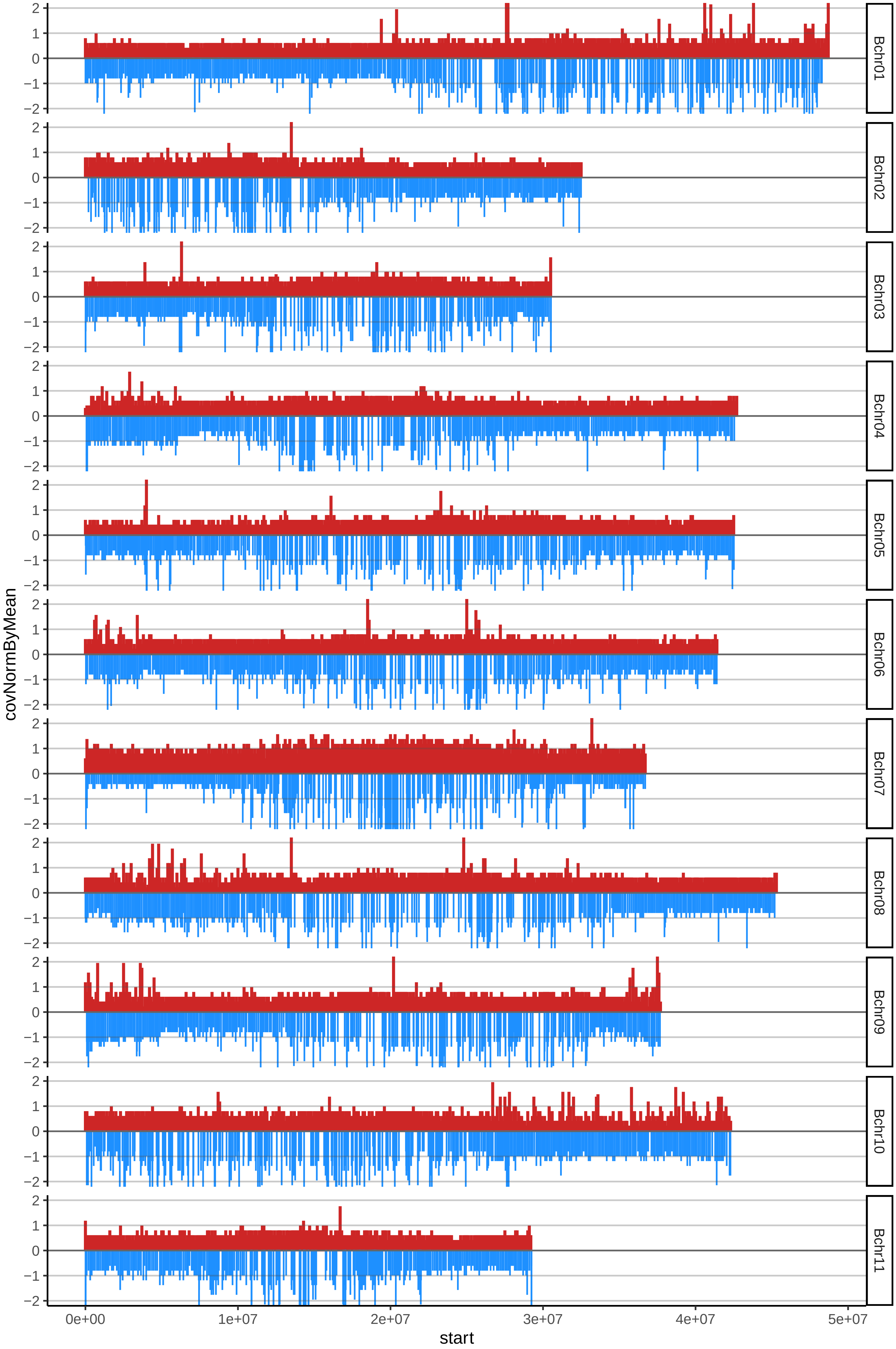

### 69PLANTAIN17cov_byA.pdf

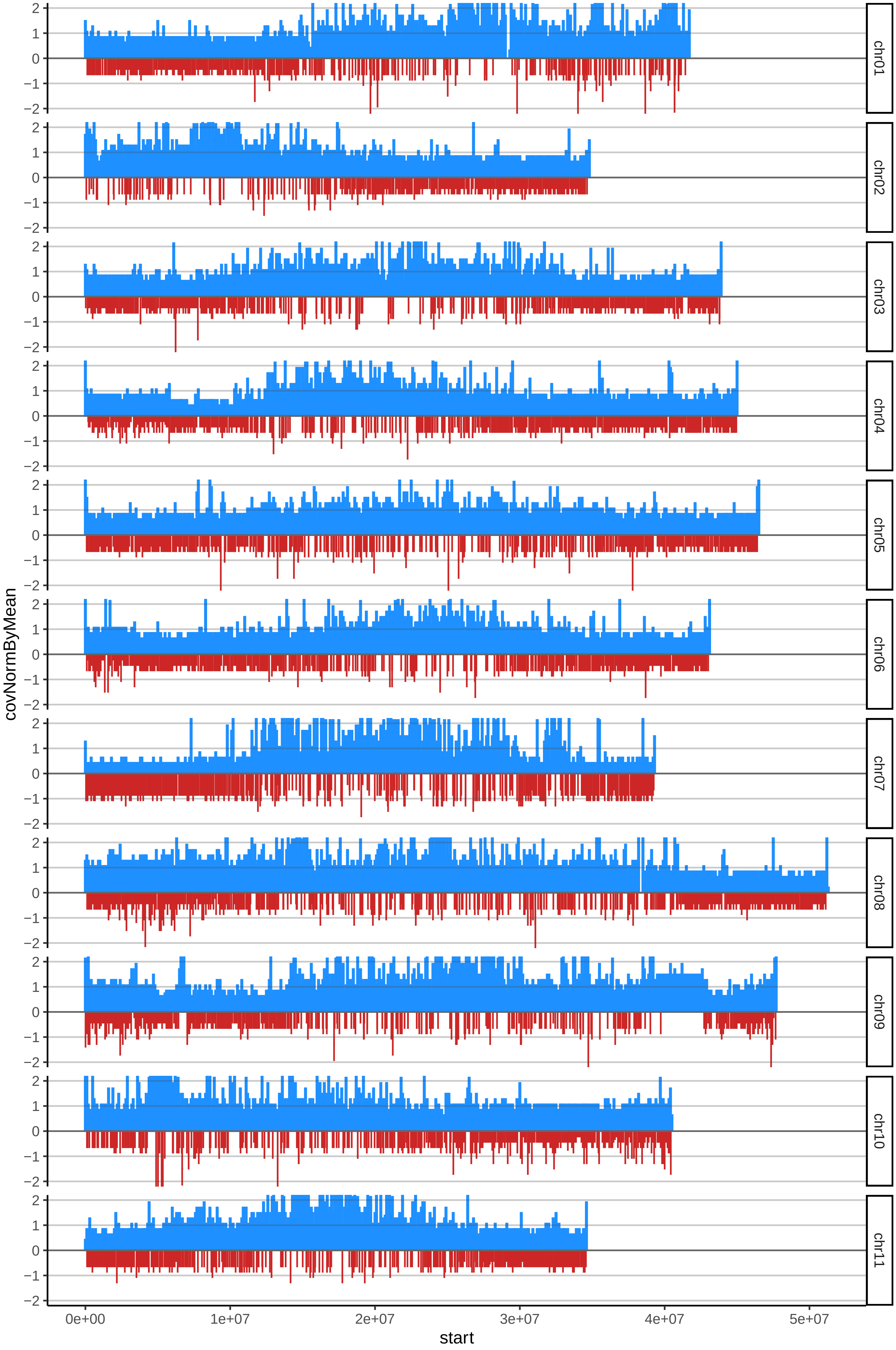

### 69PLANTAIN17cov_byB.pdf

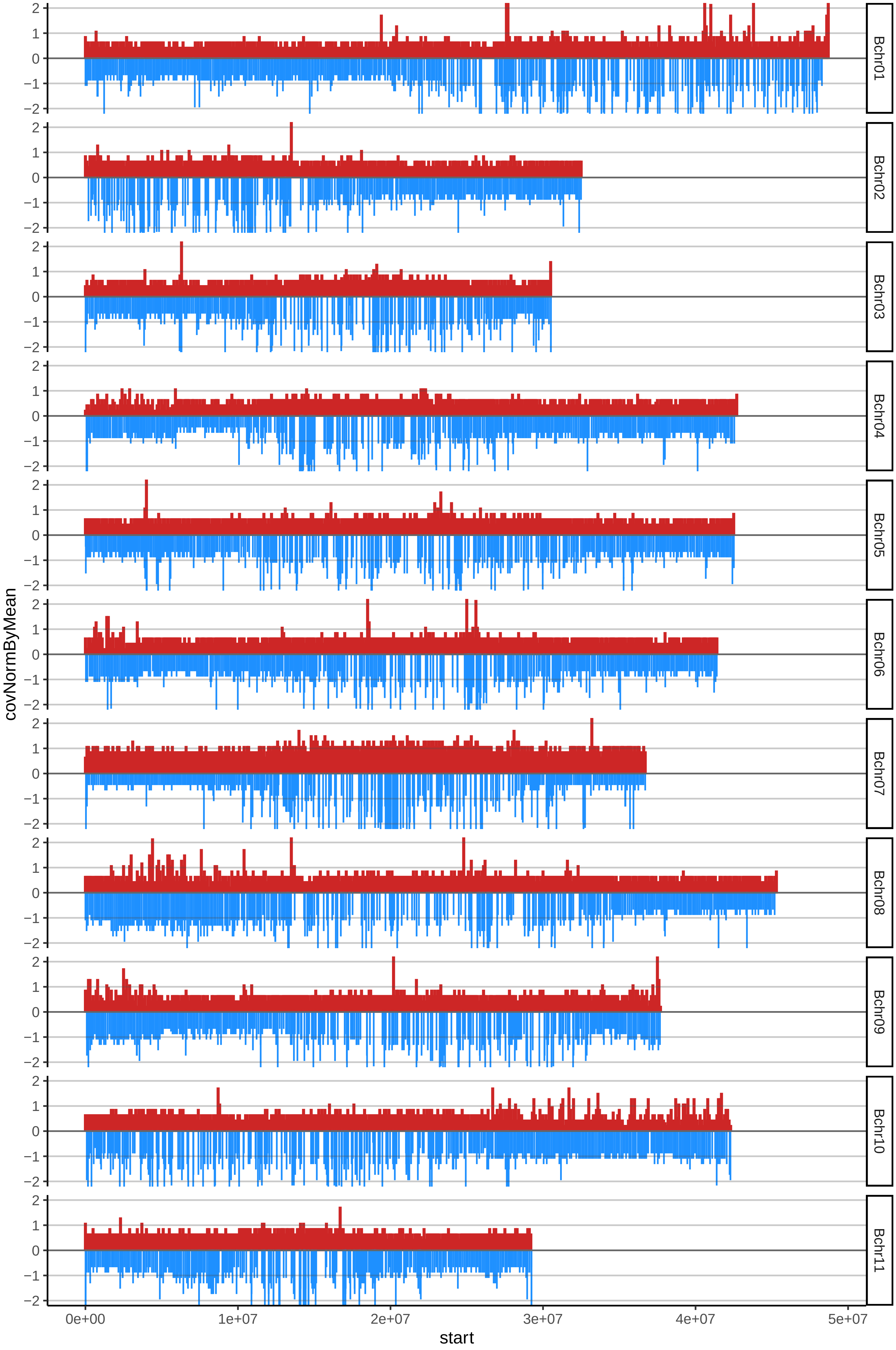

### 73ROSEDEKONAcov_byB.pdf

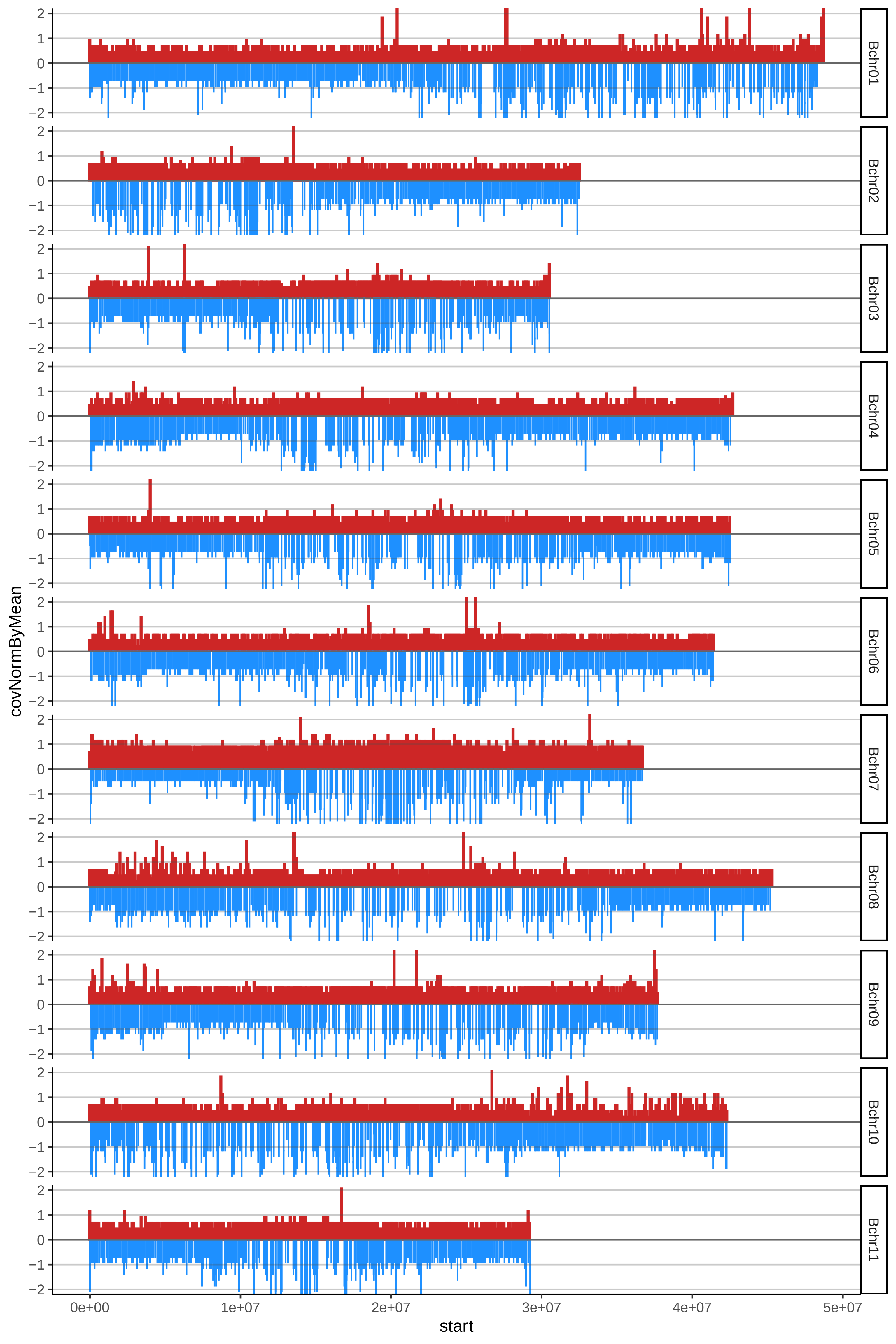

### 94IC2cov_byA.pdf

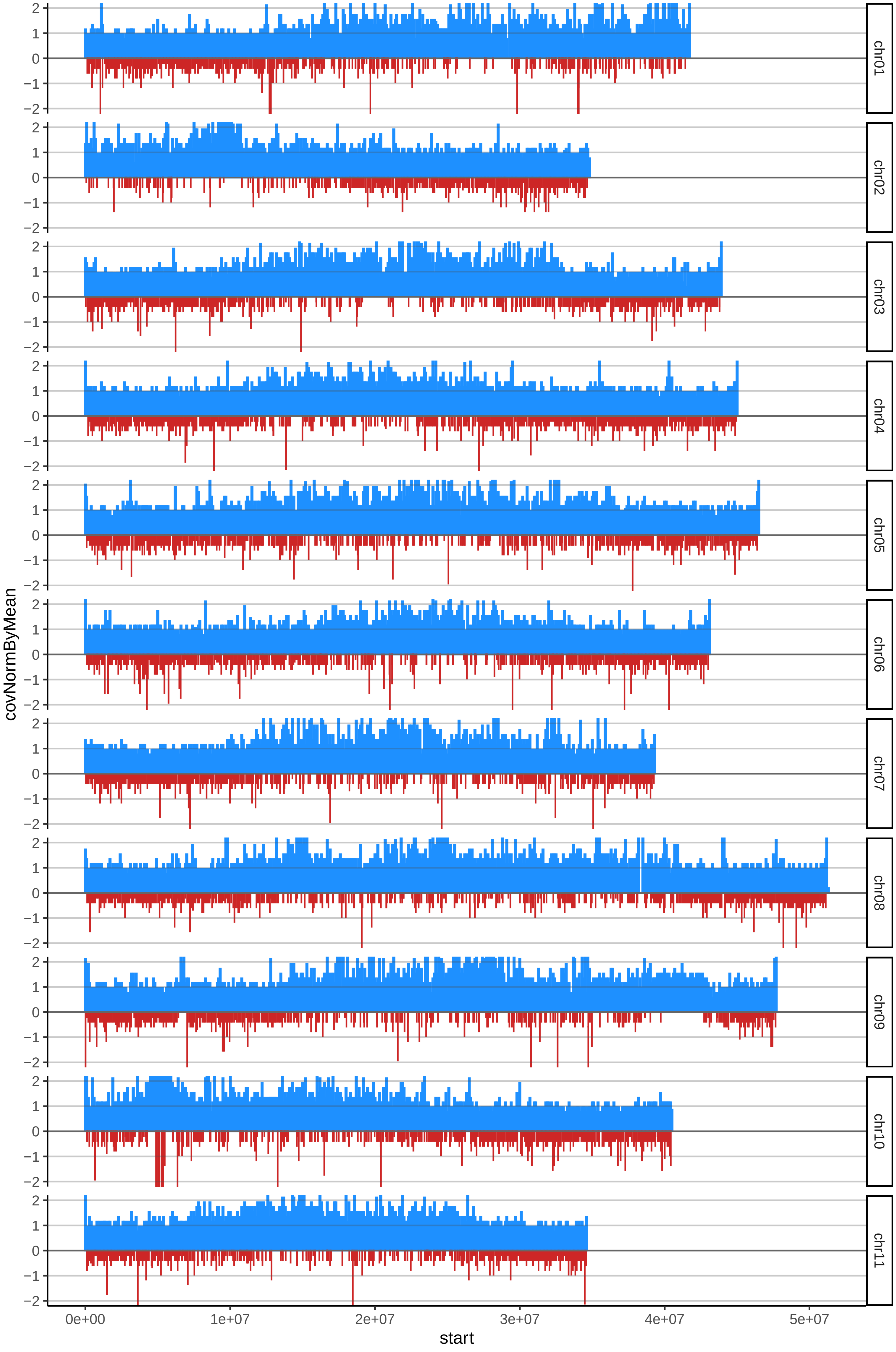

### 94IC2cov_byB.pdf

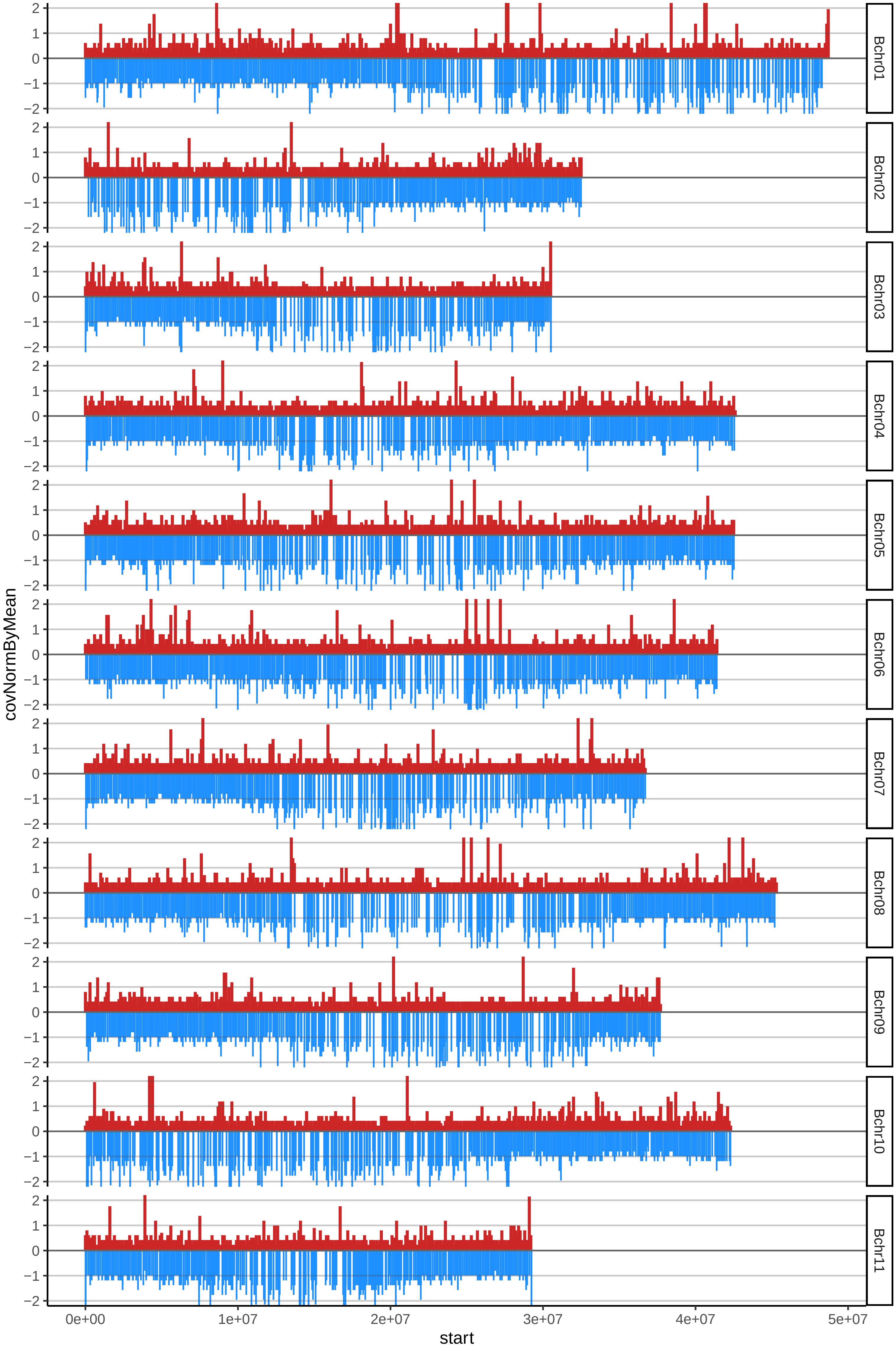

### 108GROSMICHELCOMUNcov_byA.pdf

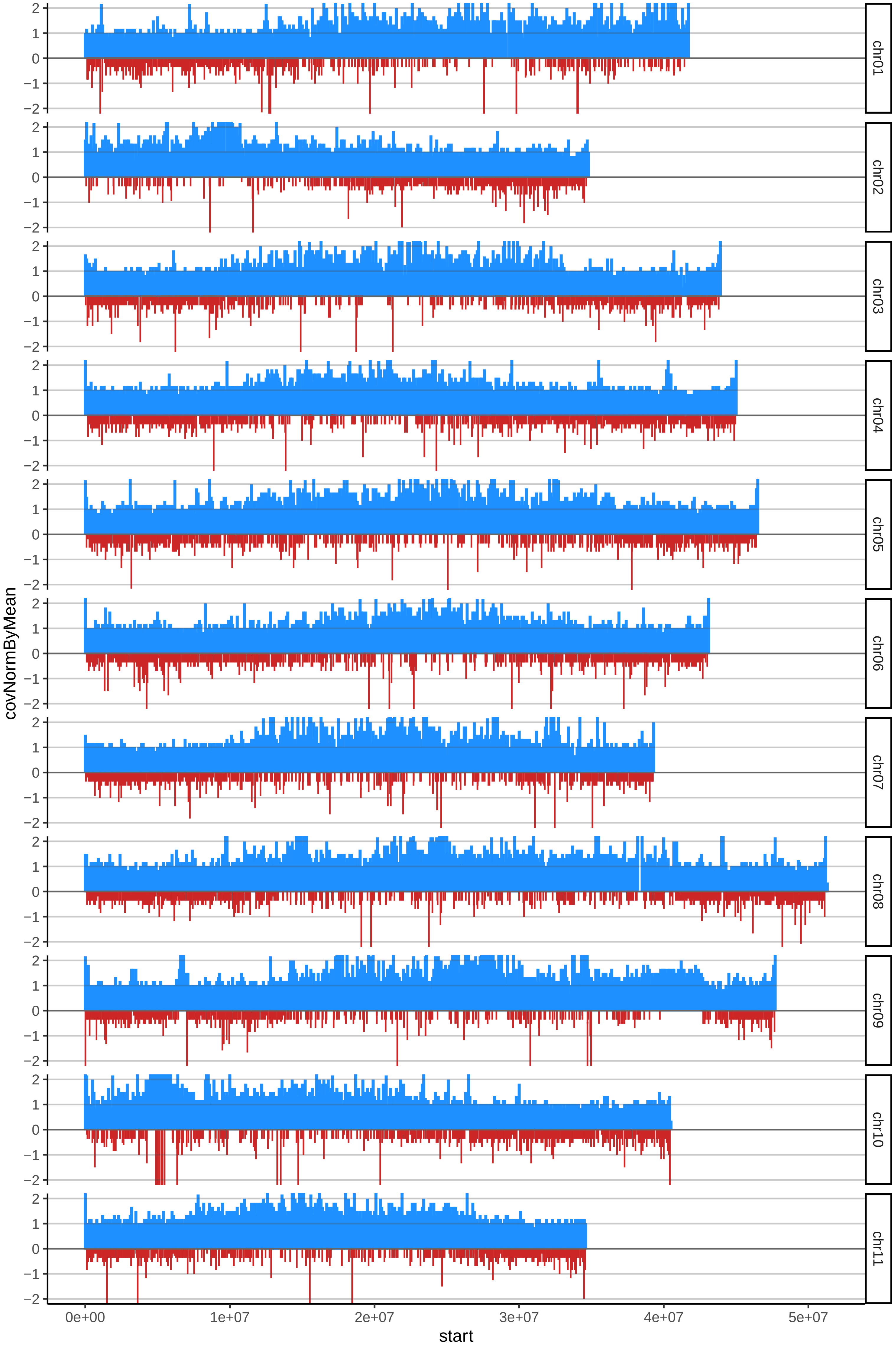

### 108GROSMICHELCOMUNcov_byB.pdf

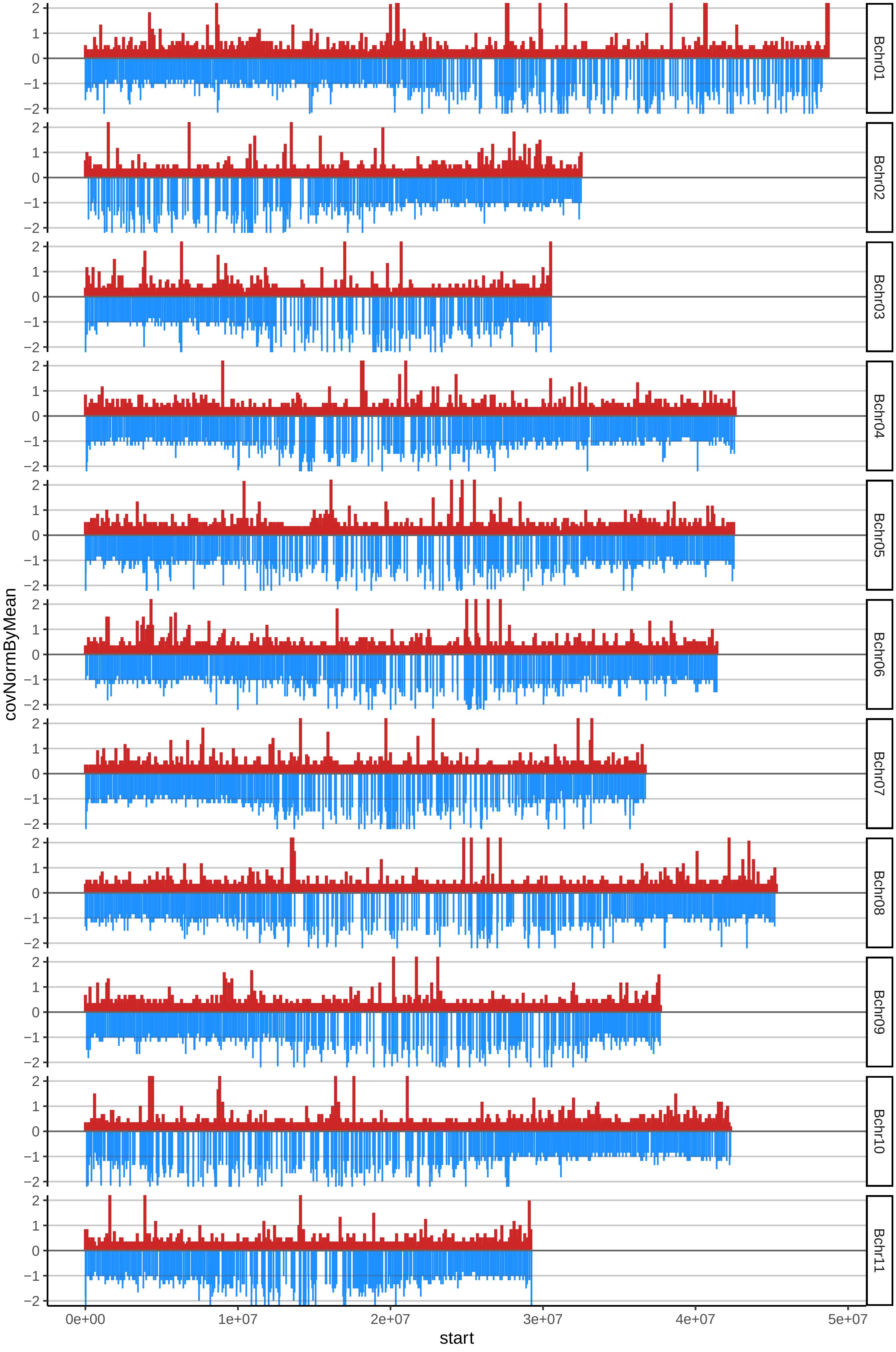

### 109GROSMICHELCOCOScov_byA.pdf

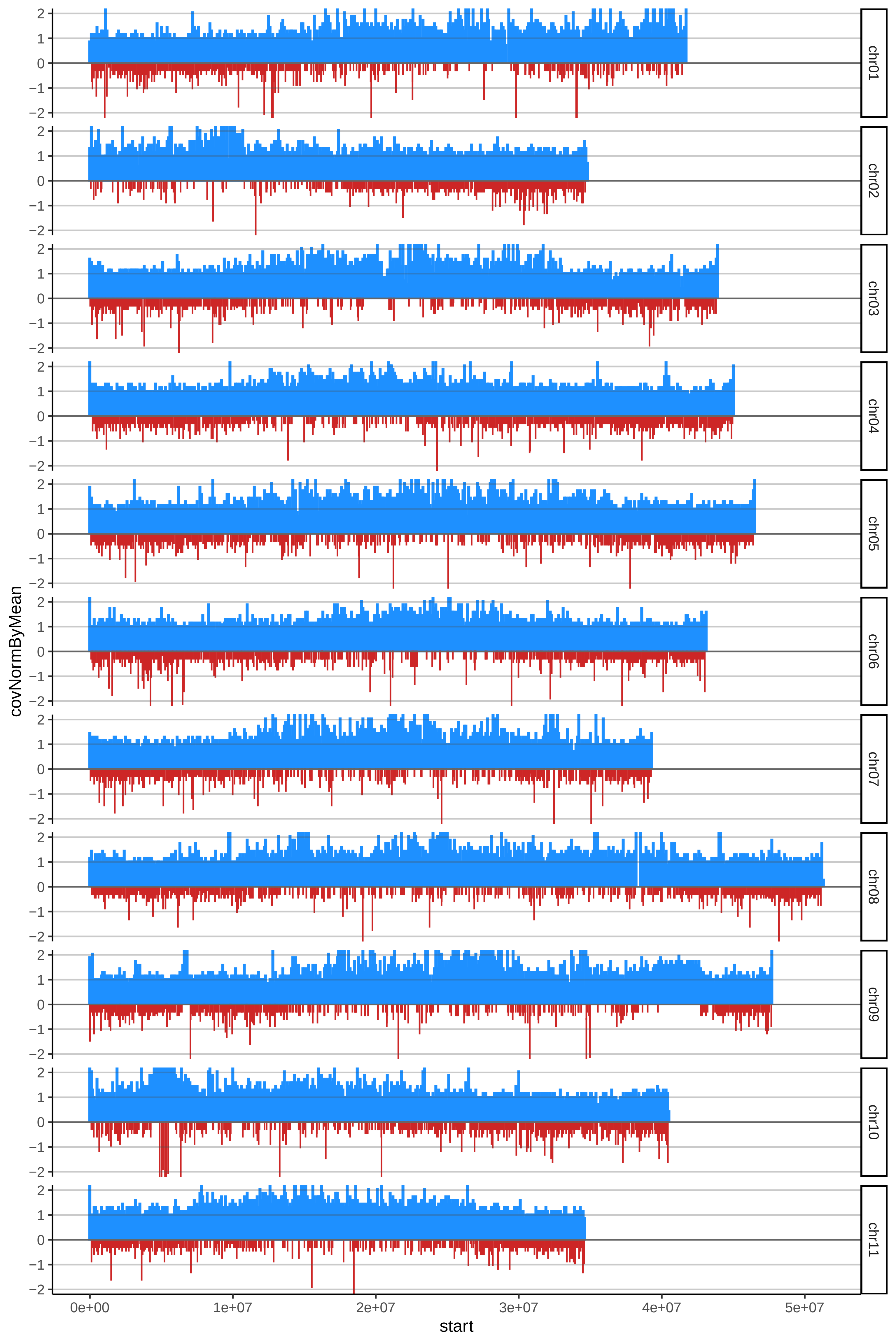

### 109GROSMICHELCOCOScov_byB.pdf

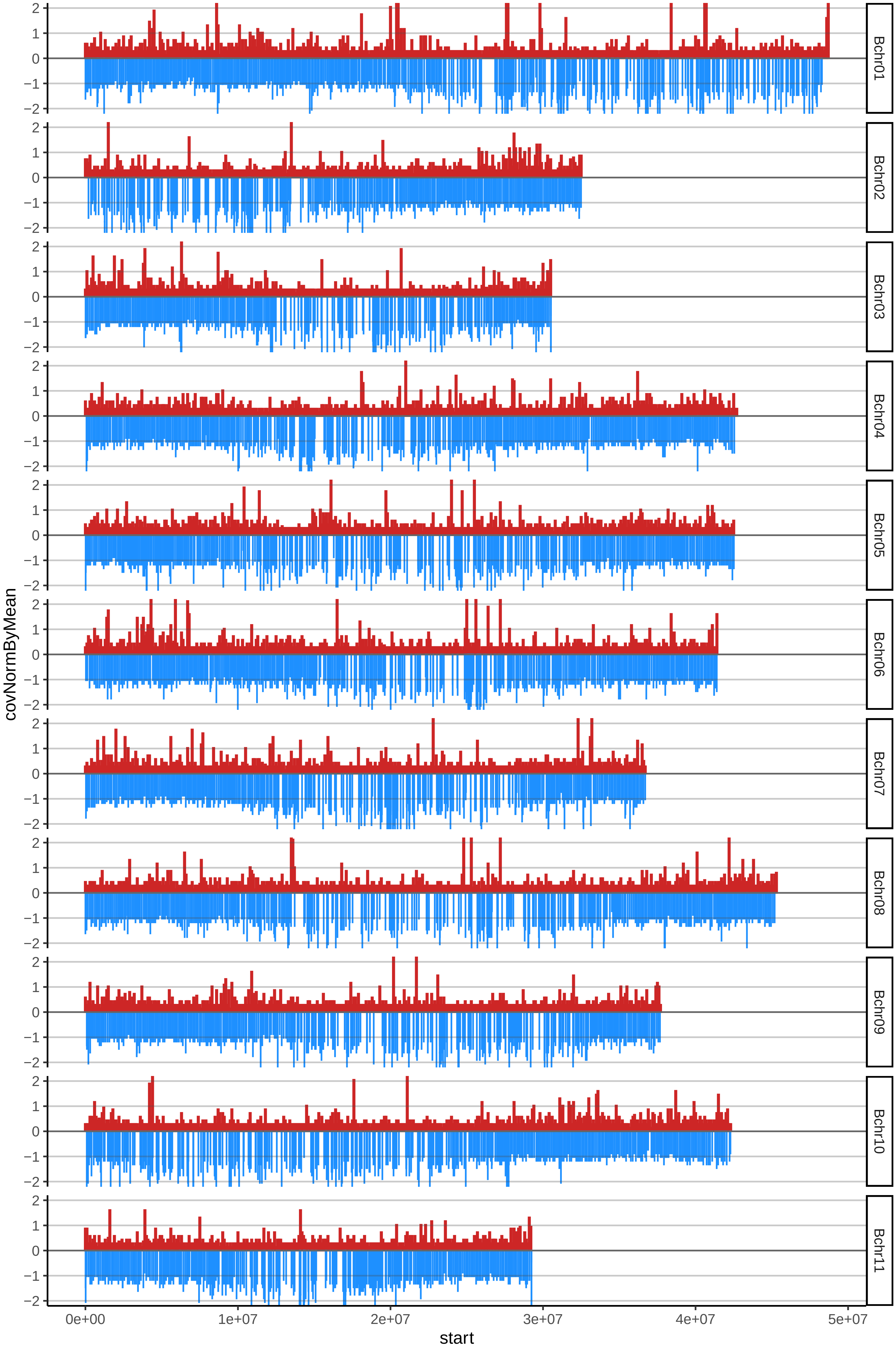

### 110GROSMICHELENANOcov_byA.pdf

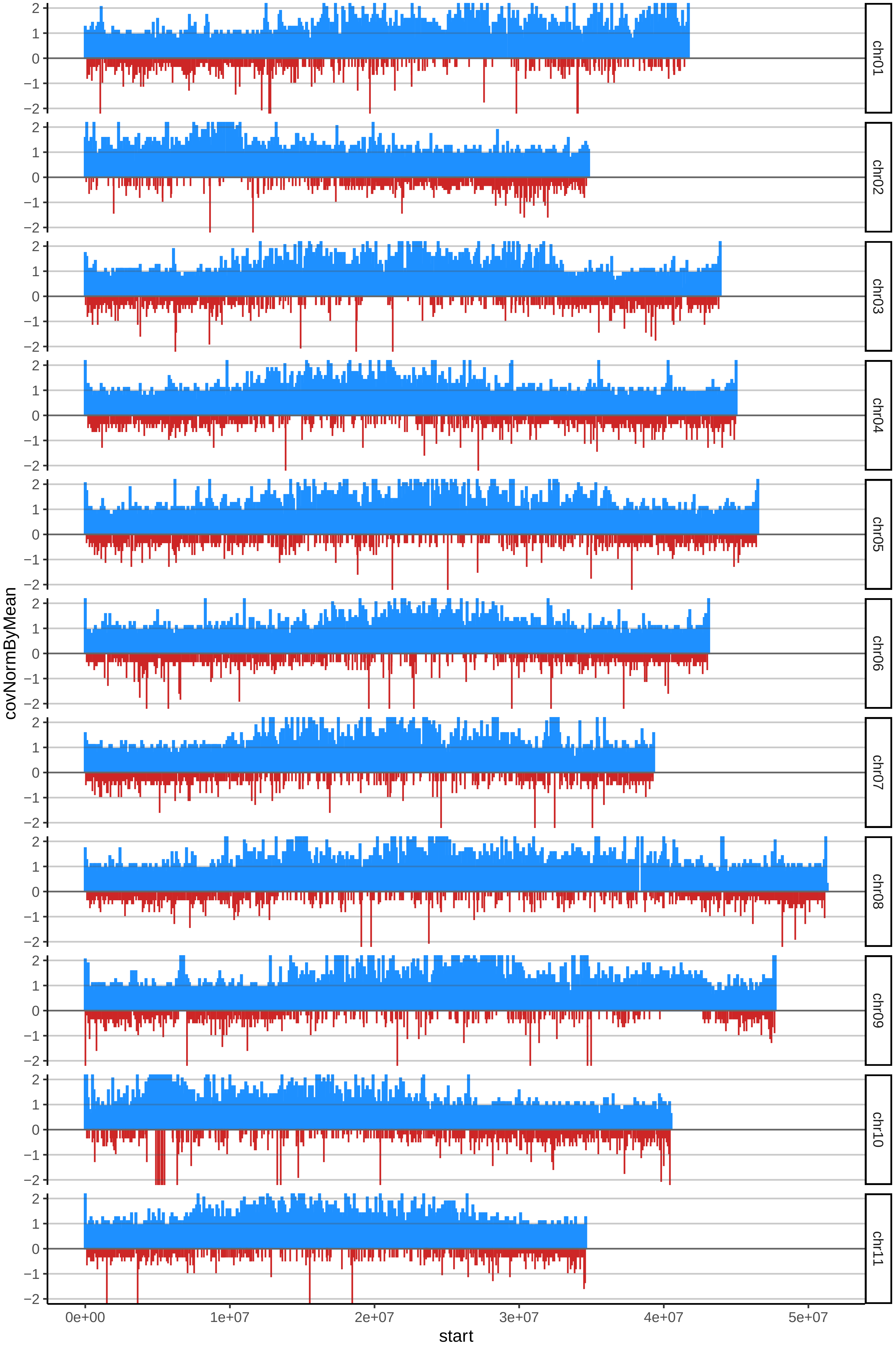

### 110GROSMICHELENANOcov_byB.pdf

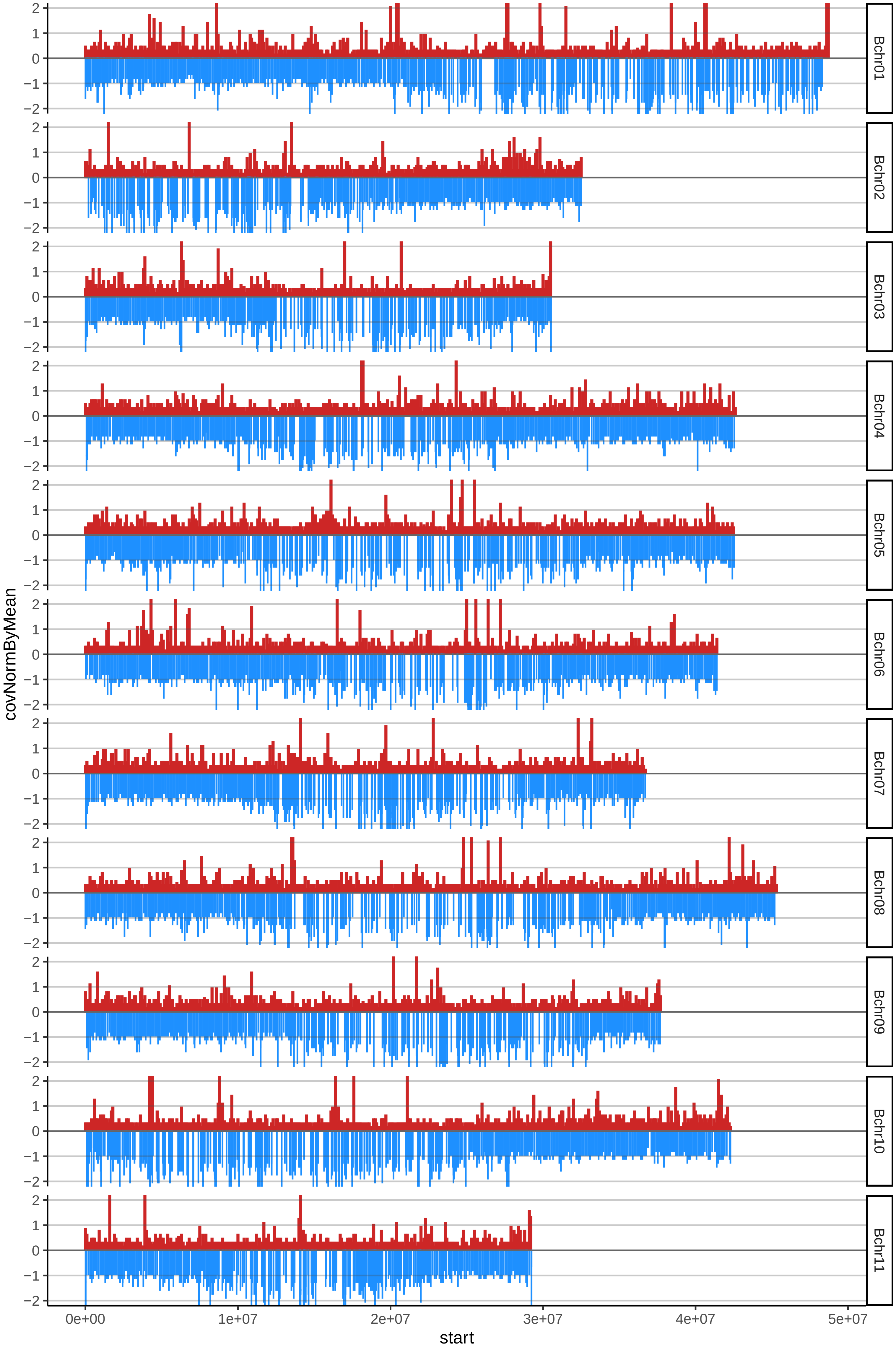

### 111SEDAcov_byA.pdf

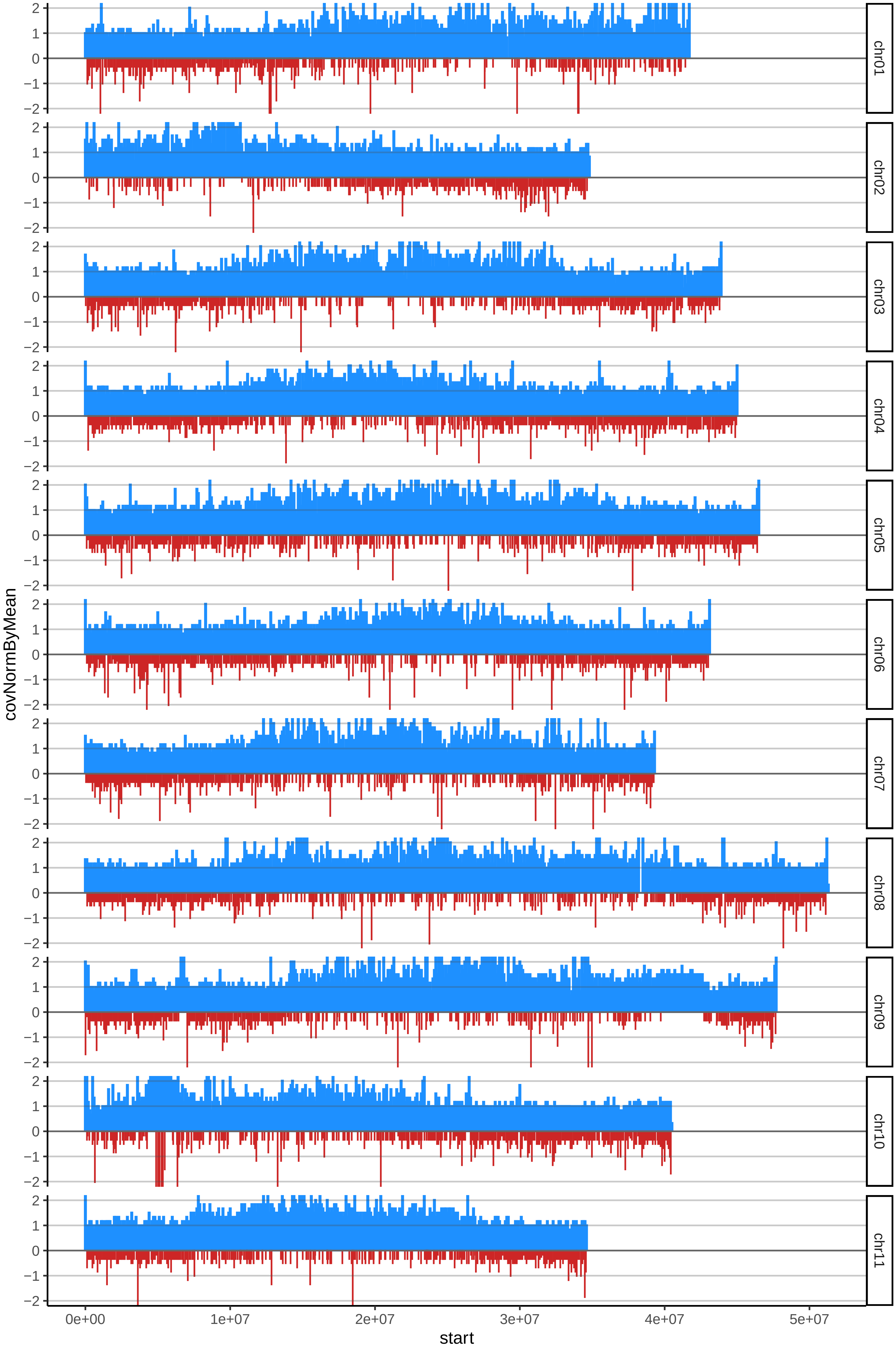

### 111SEDAcov_byB.pdf

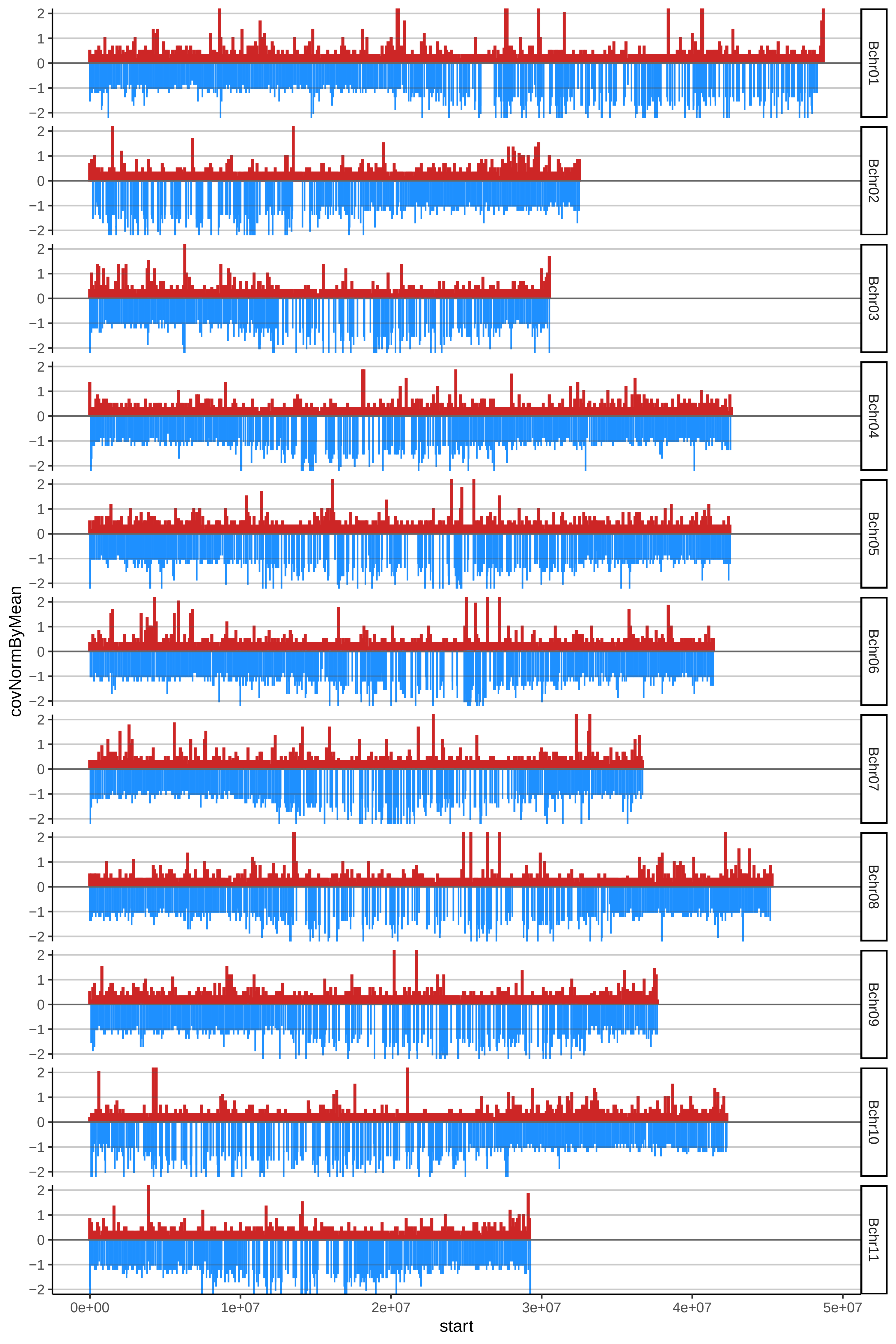

### 112BANANOCHICOcov_byA.pdf

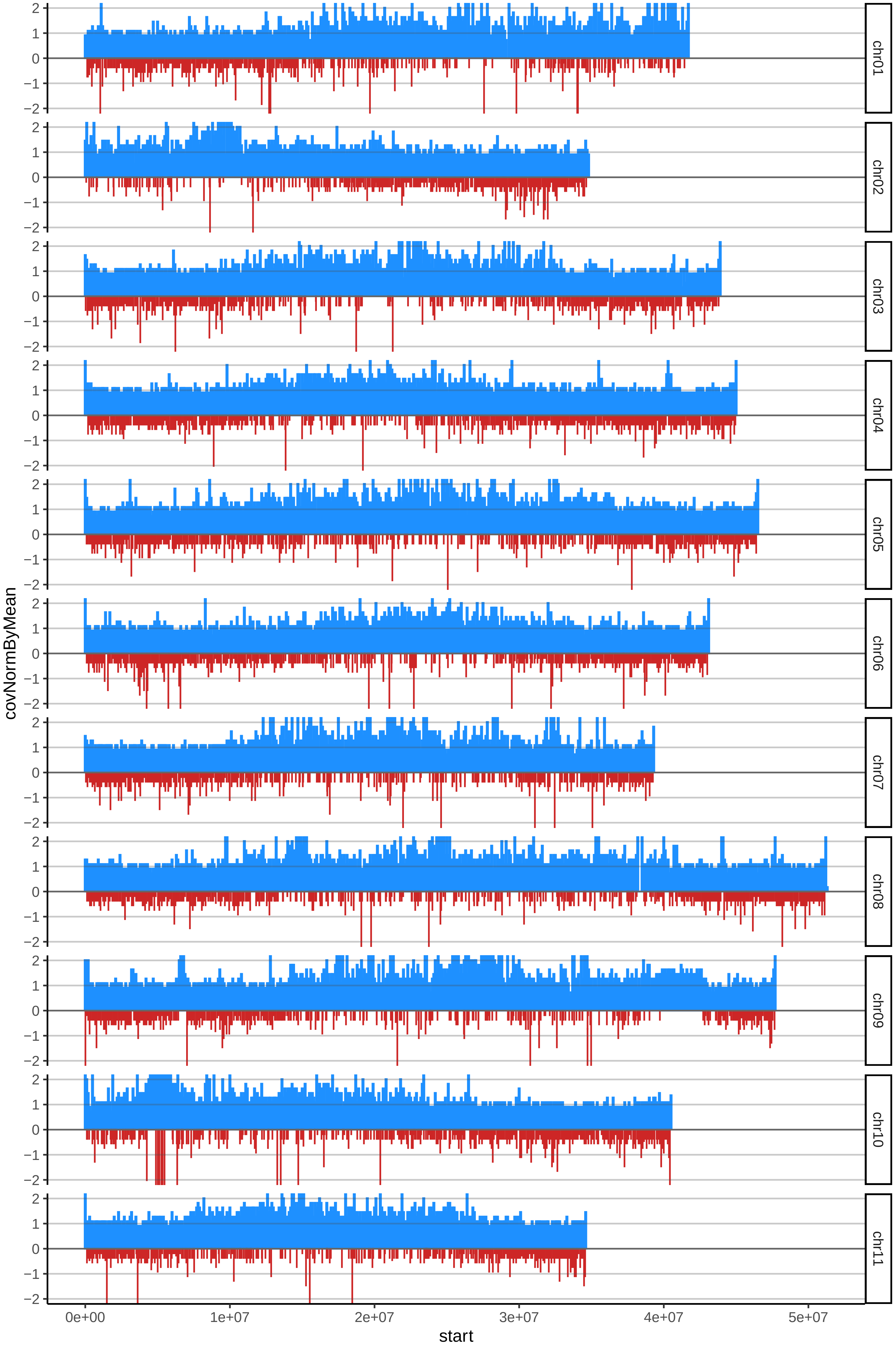

### 112BANANOCHICOcov_byB.pdf

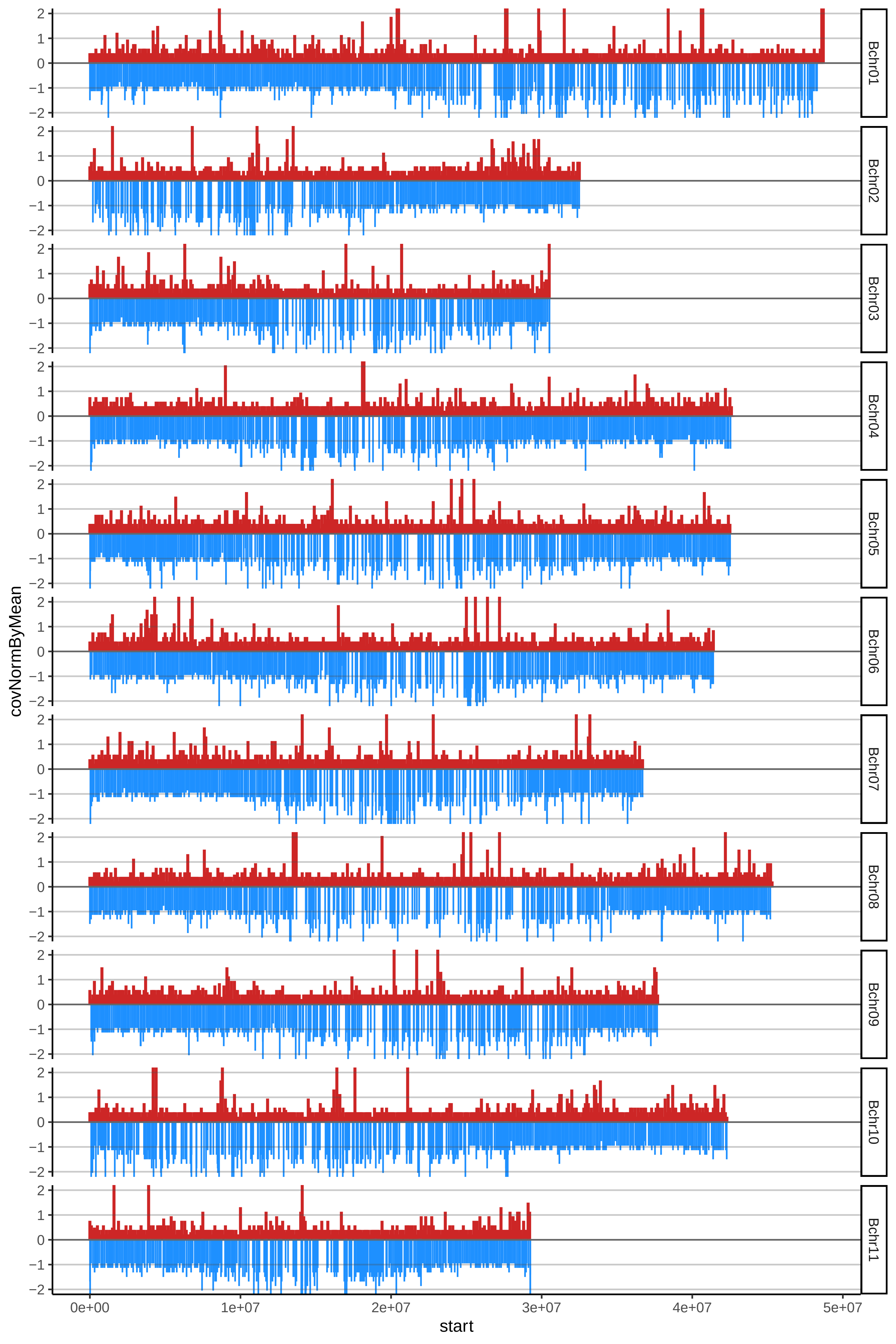

### 135DOMINICOROJO2cov_byA.pdf

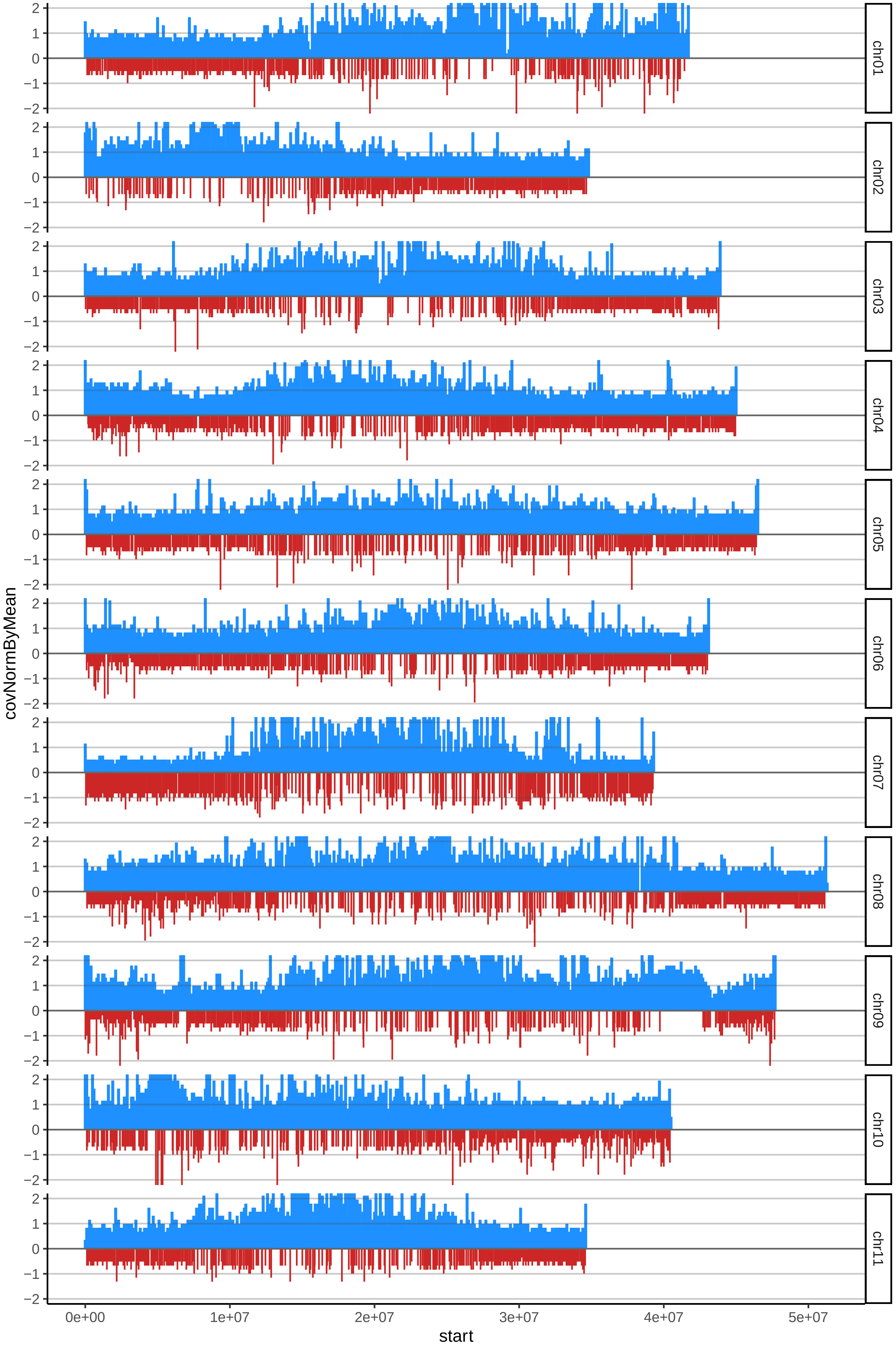

### 135DOMINICOROJO2cov_byB.pdf

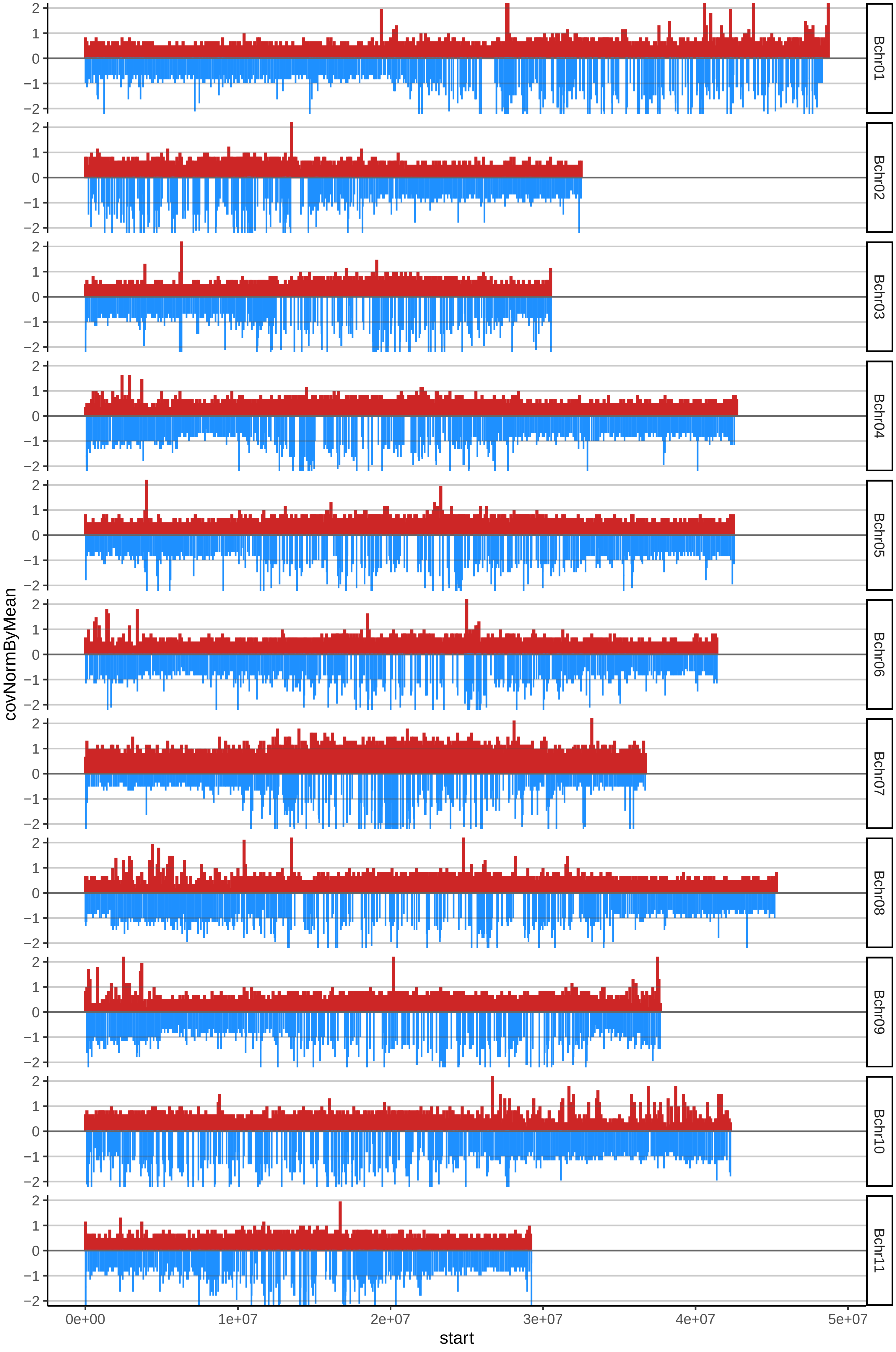

### 137AMOUVERDEcov_byA.pdf

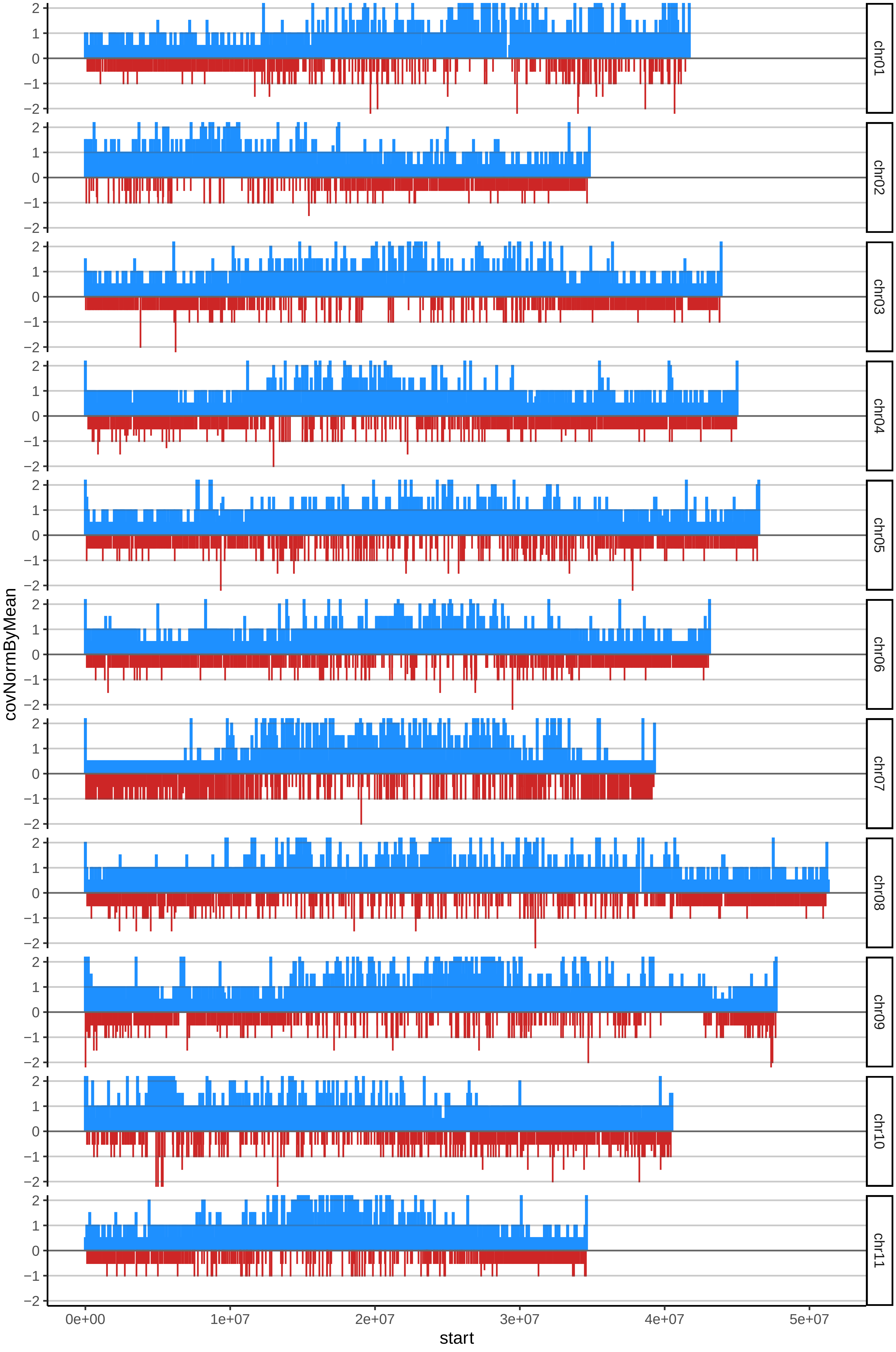

### 137AMOUVERDEcov_byB.pdf

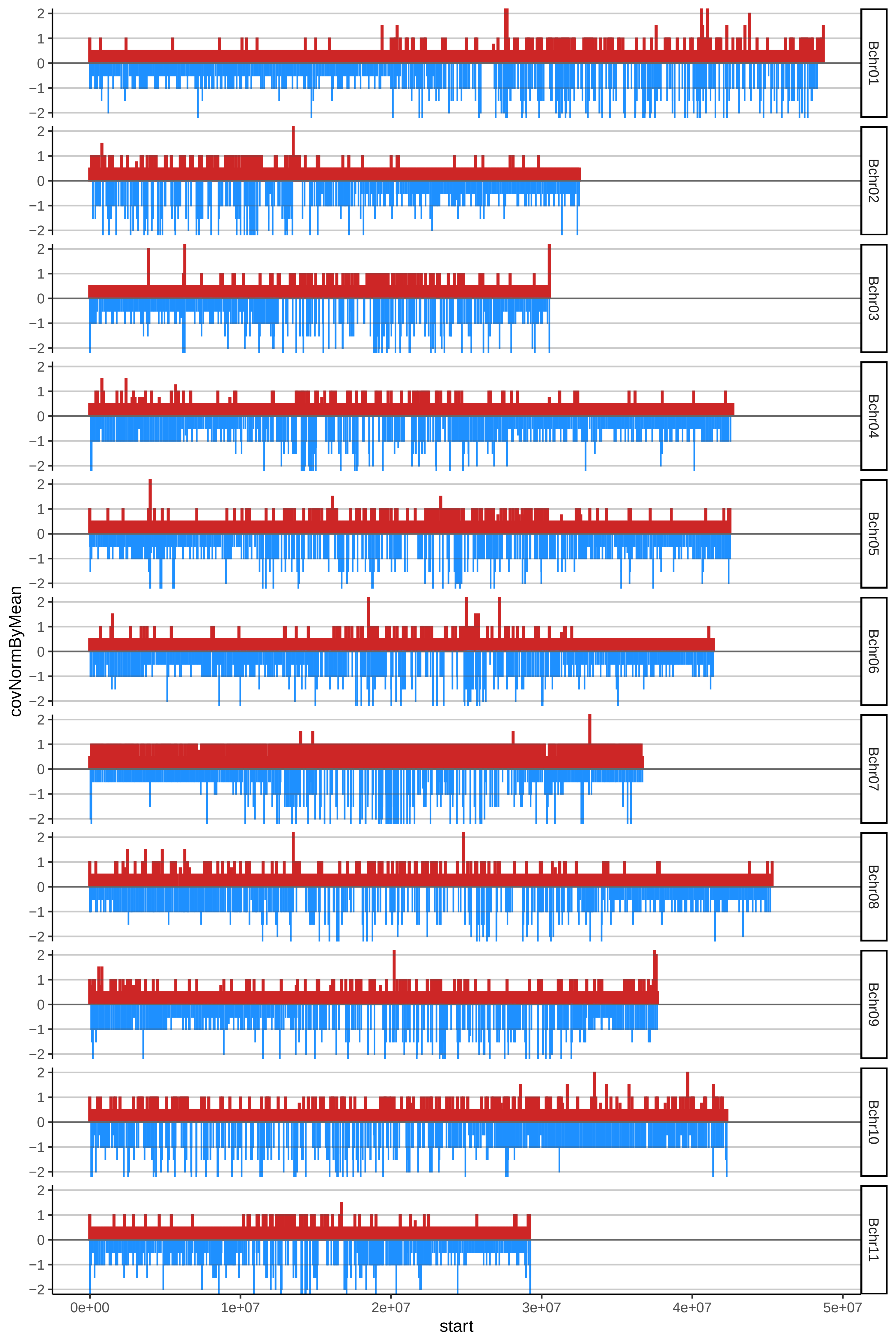

### 139HARTONSANTANDERcov_byA.pdf

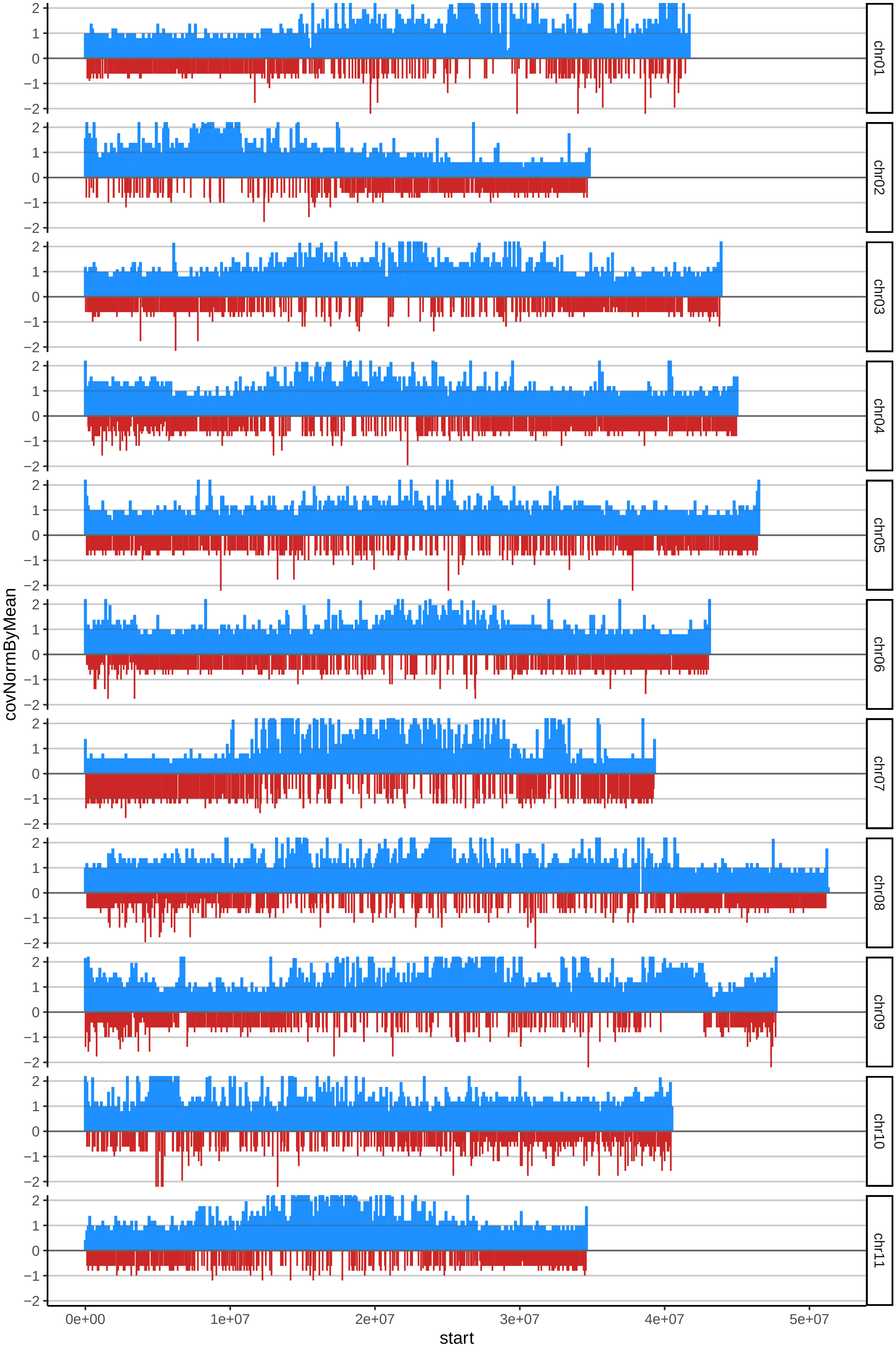

### 139HARTONSANTANDERcov_byB.pdf

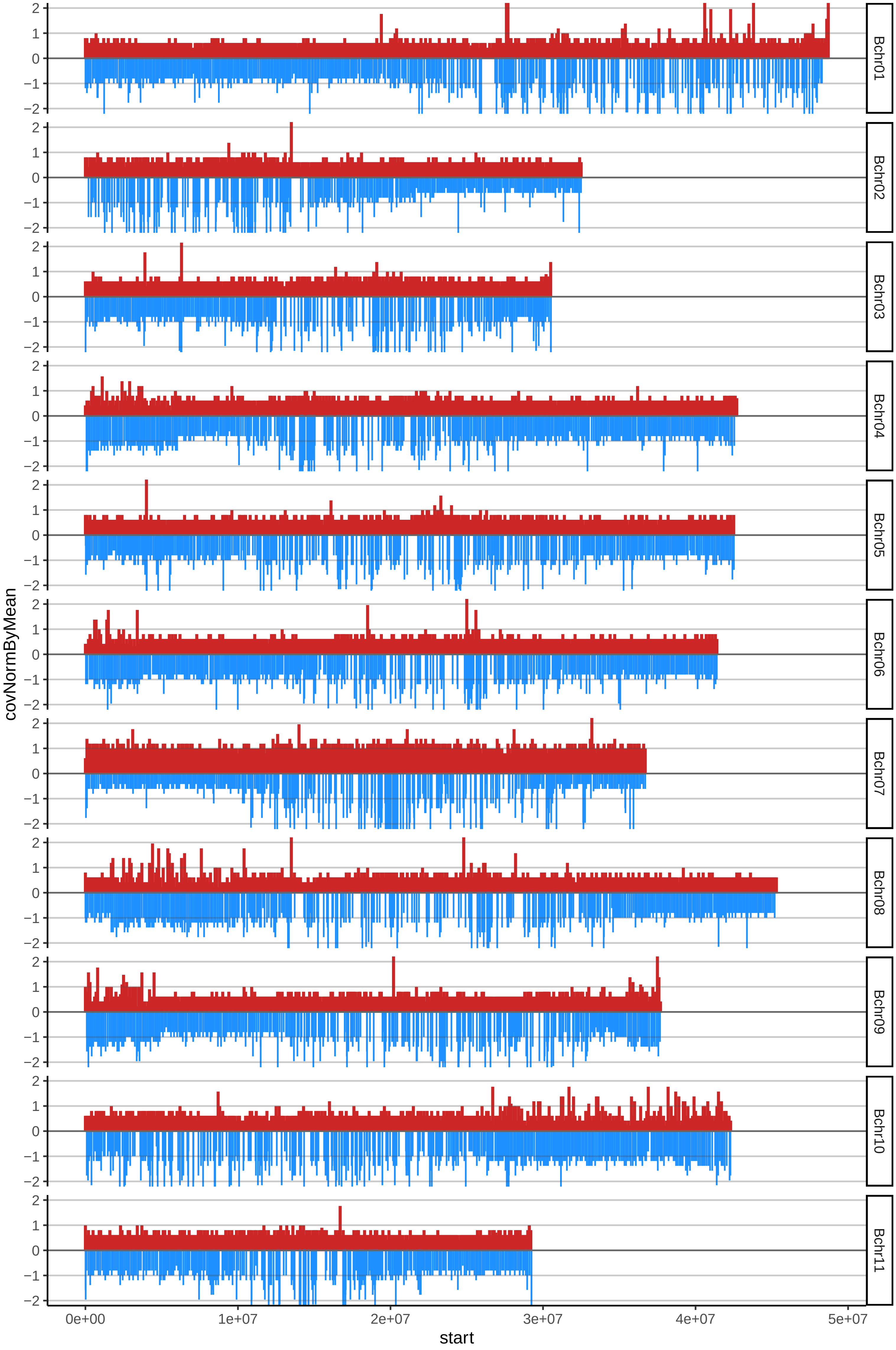

### 142LAMIELcov_byA.pdf

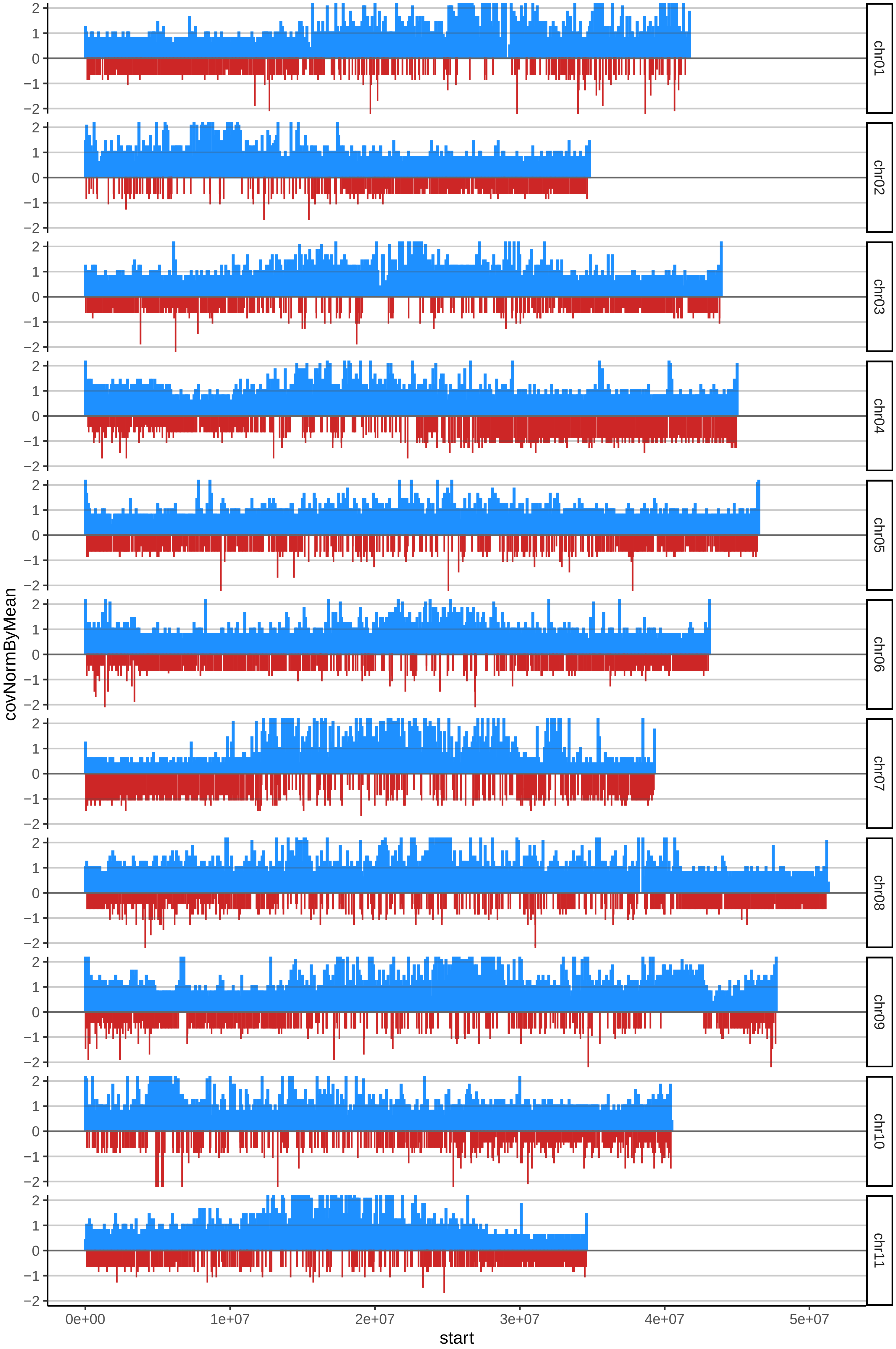

### 142LAMIELcov_byB.pdf

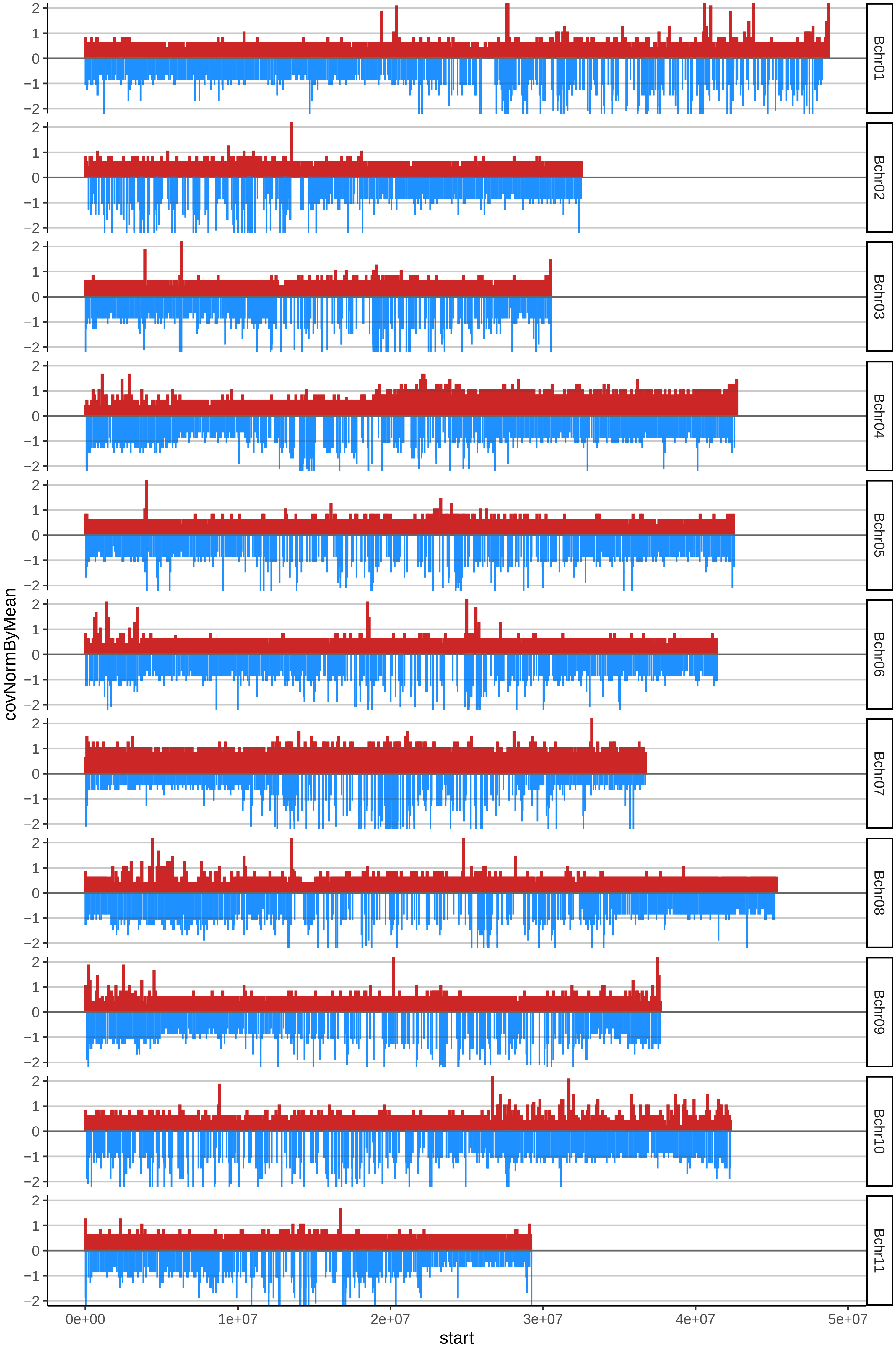

### 191HARTONCUBANOcov_byA.pdf

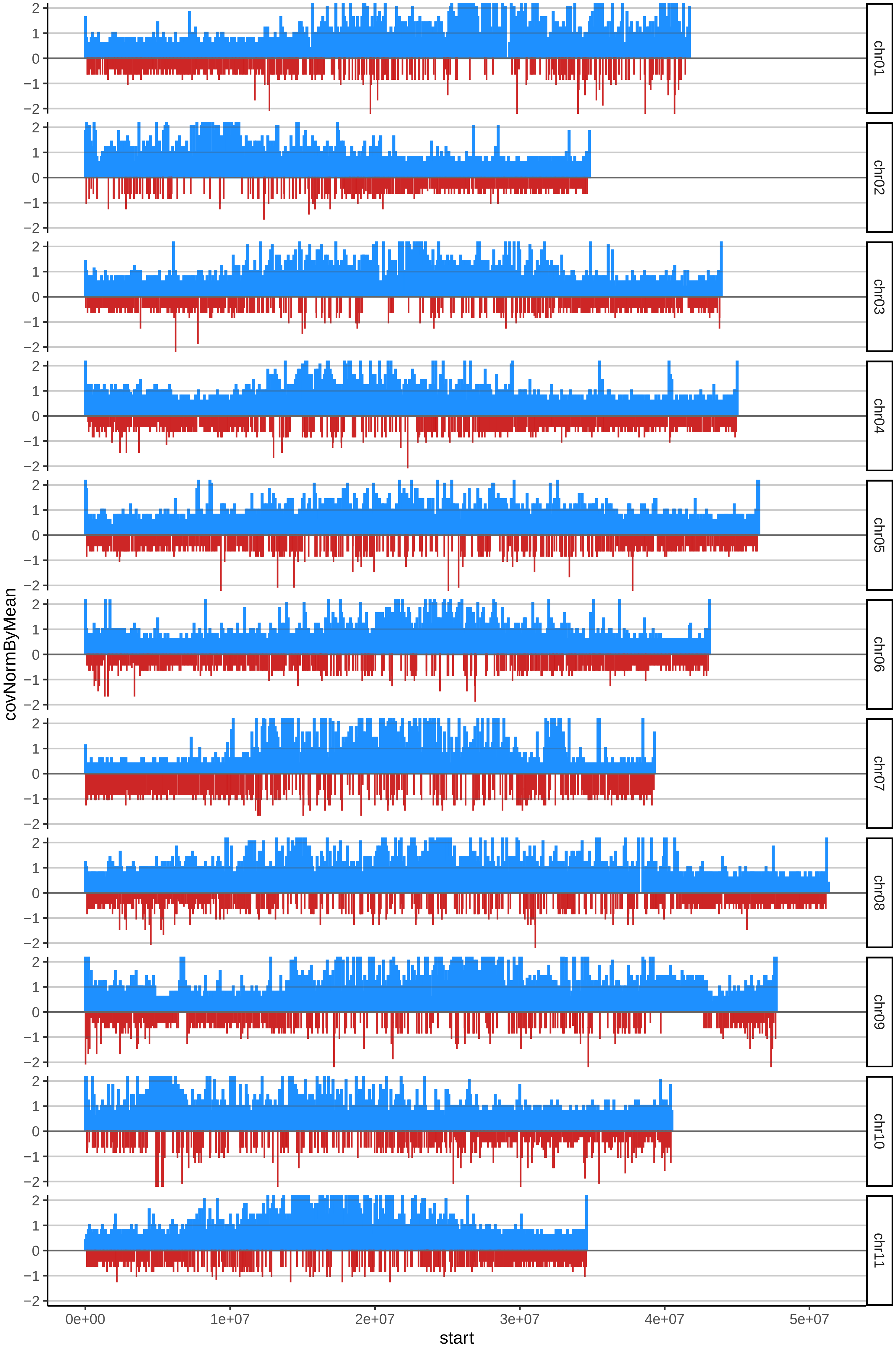

### 191HARTONCUBANOcov_byB.pdf

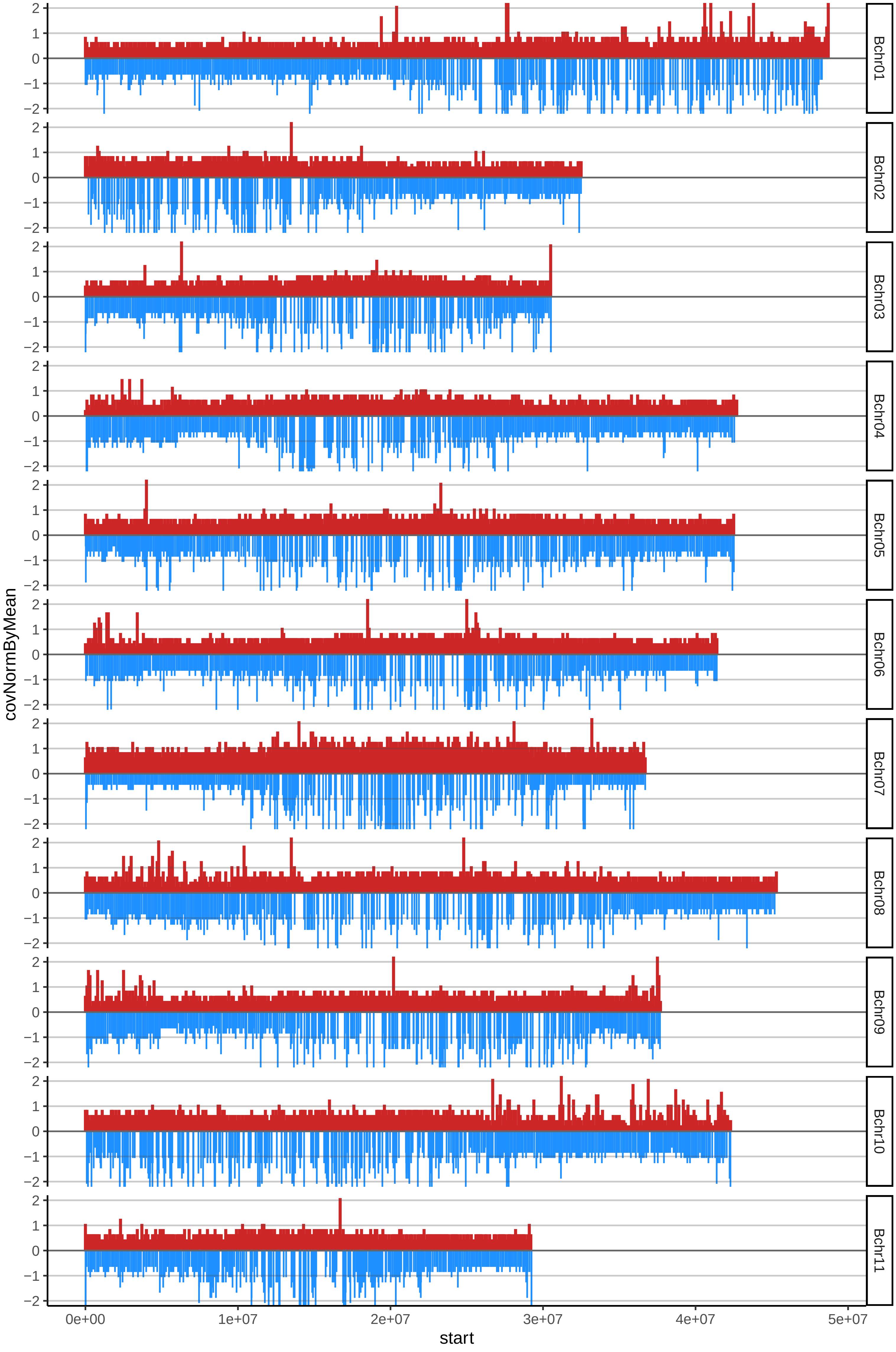

### 192HARTONMACHOcov_byA.pdf

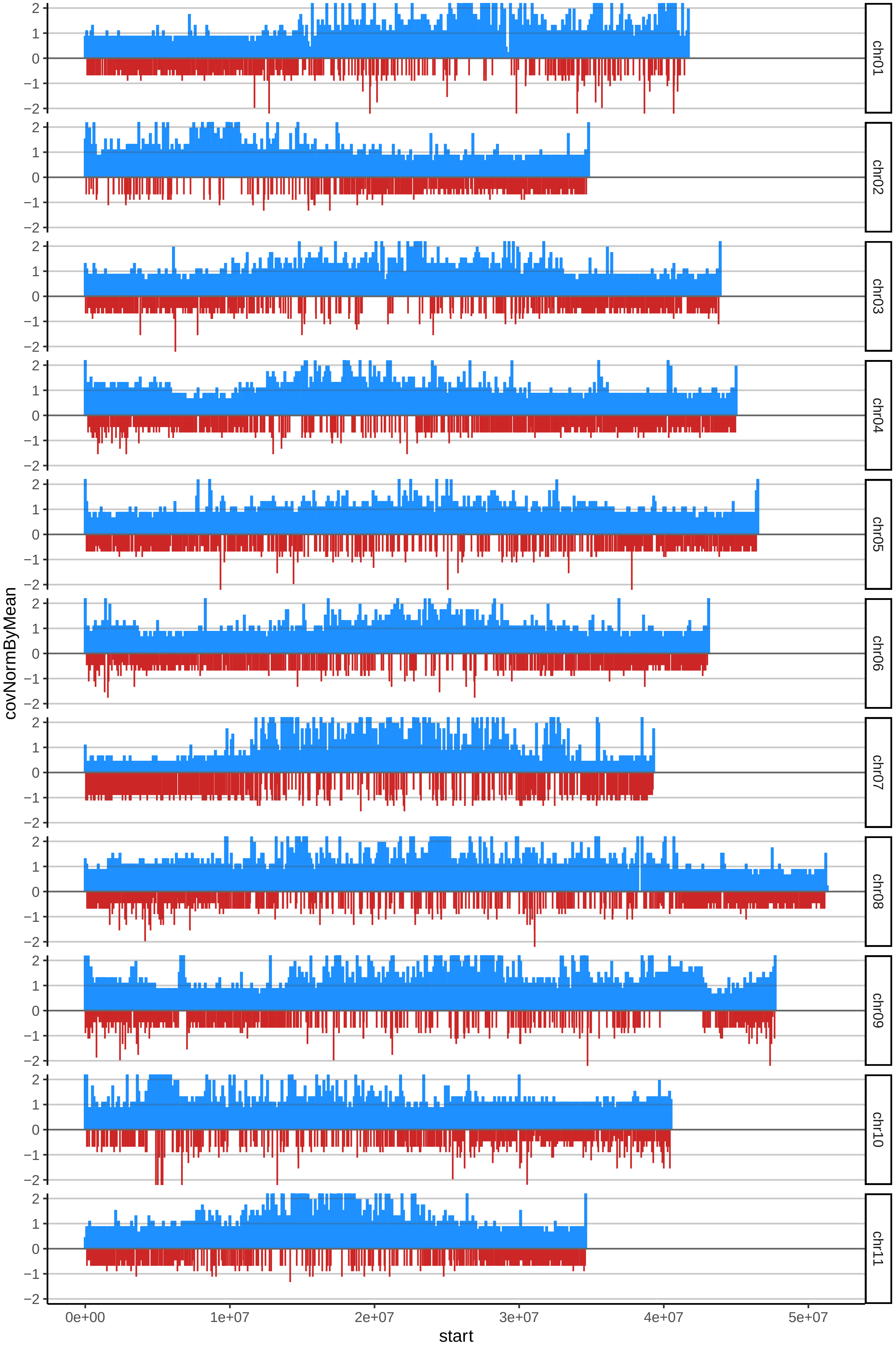

### 192HARTONMACHOcov_byB.pdf

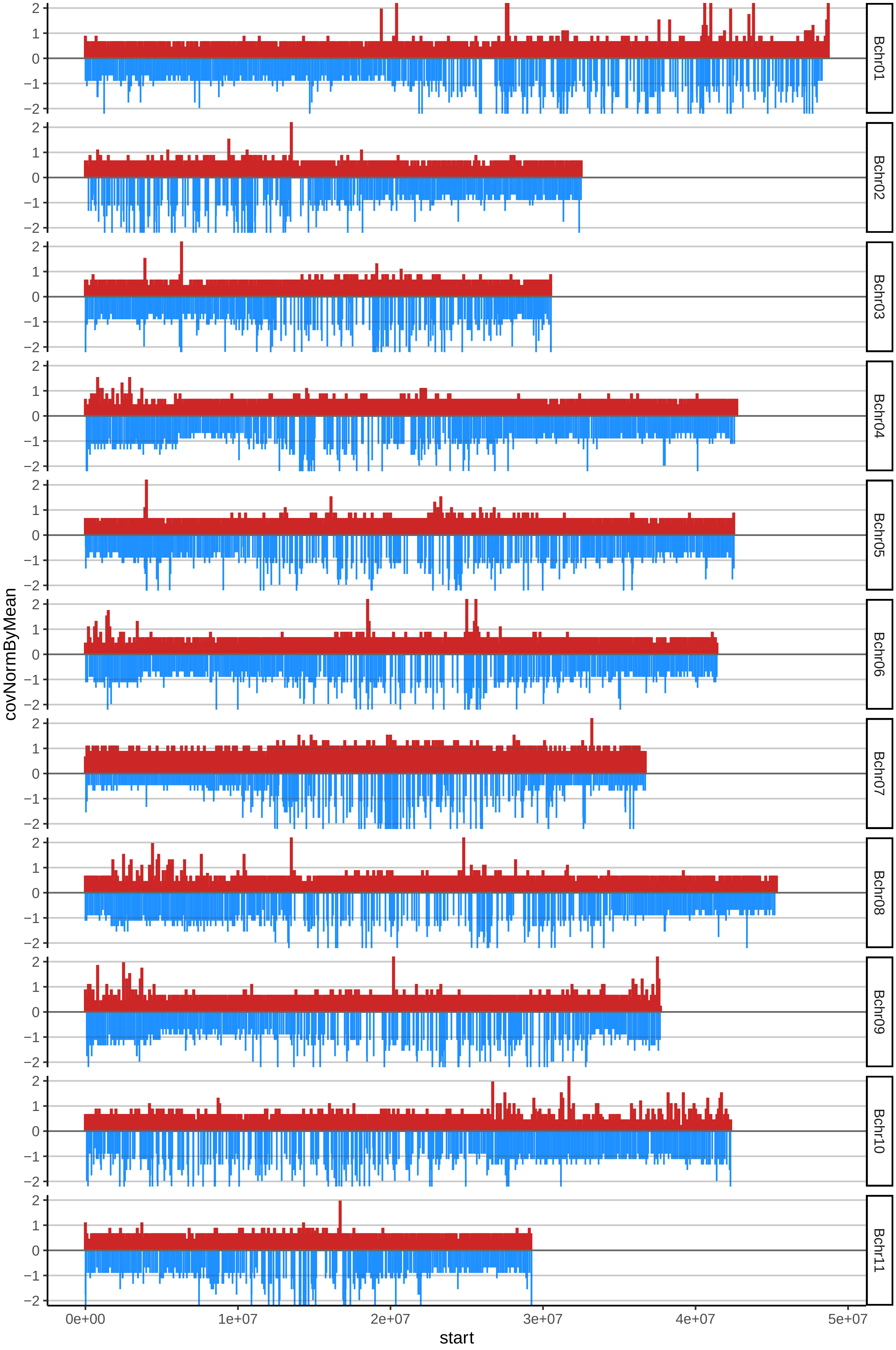

### 197CUBANONEGROcov_byA.pdf

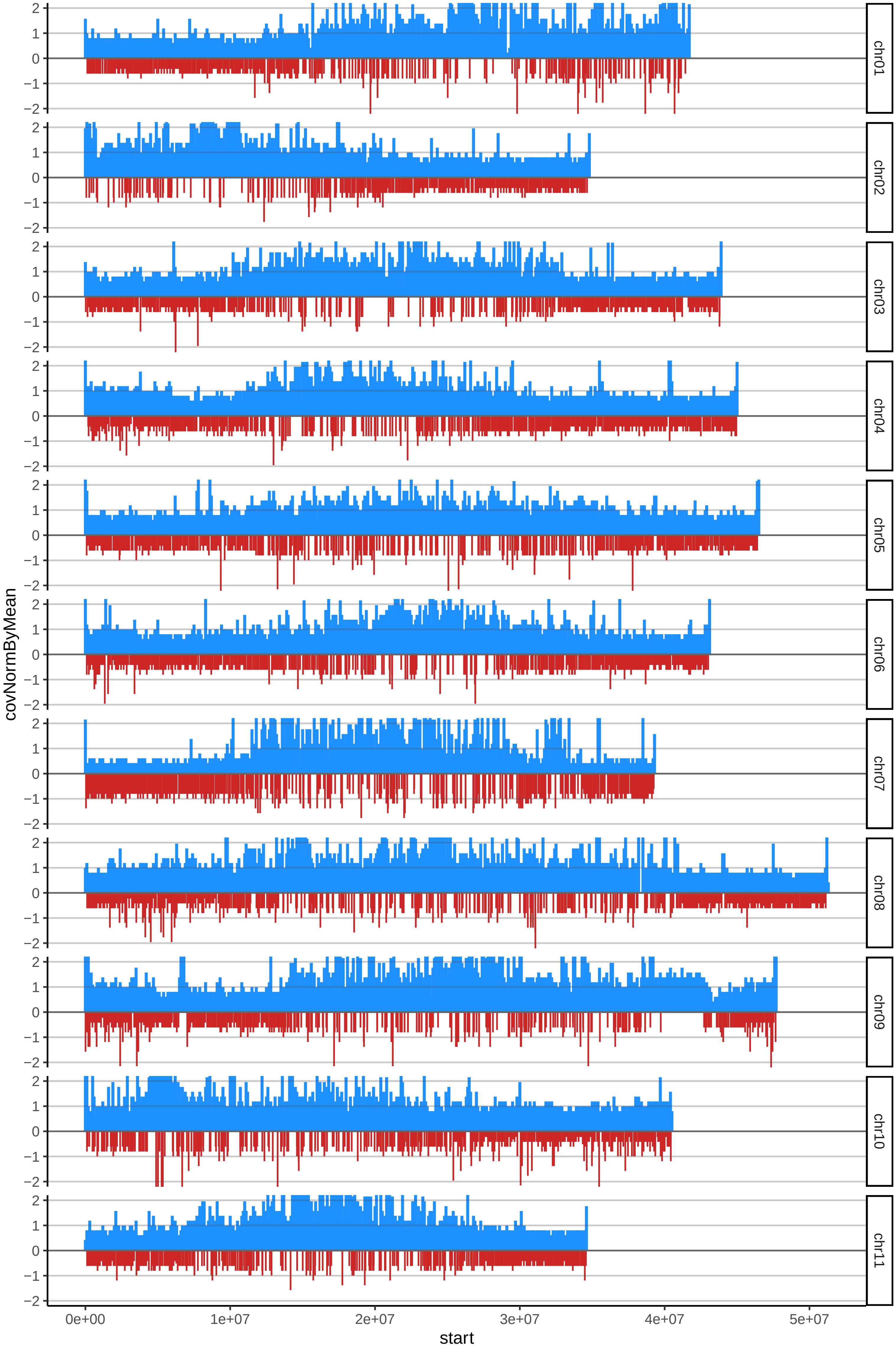

### 197CUBANONEGROcov_byB.pdf

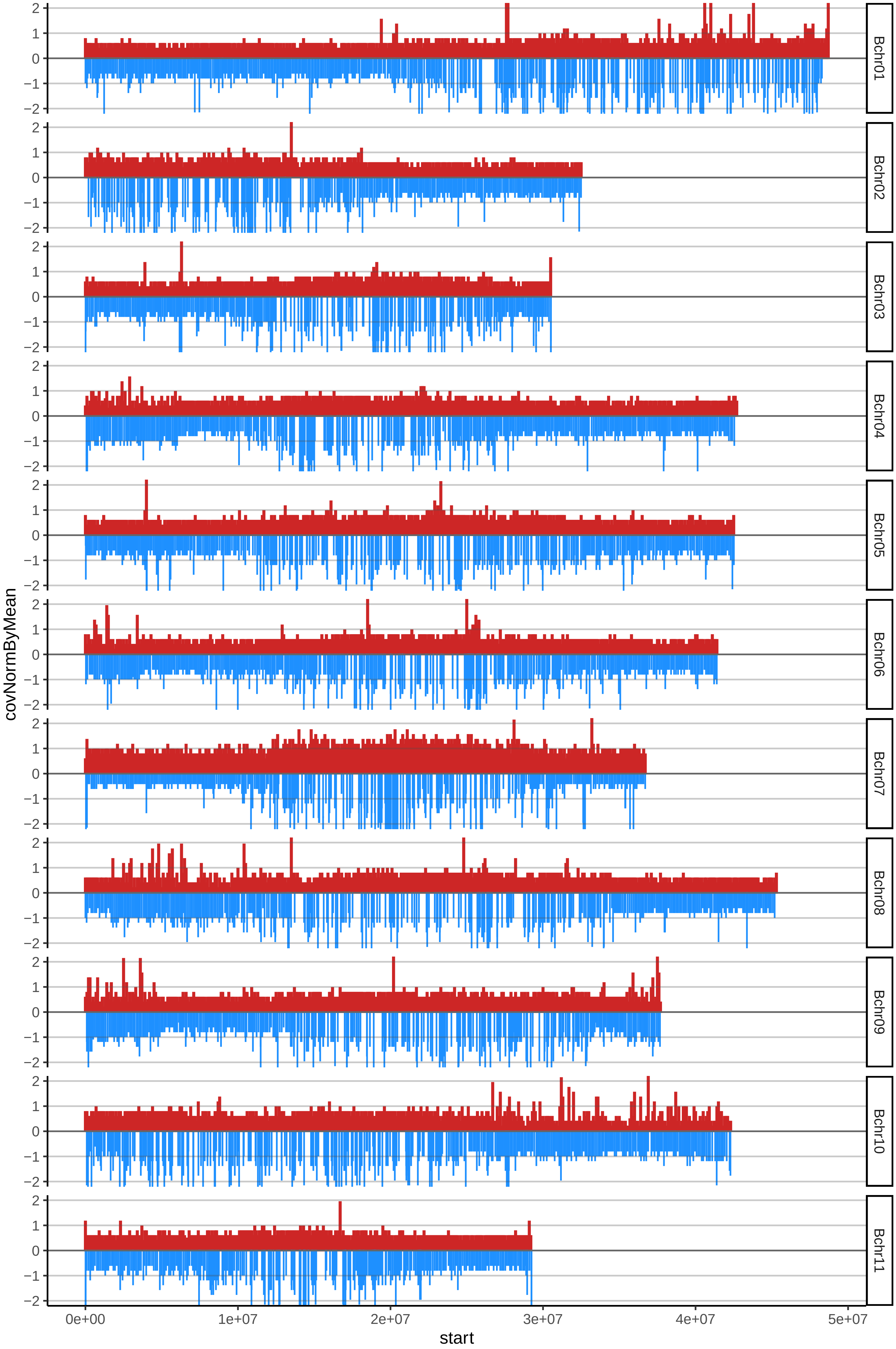

### 198CUBANOBLANCOcov_byA.pdf

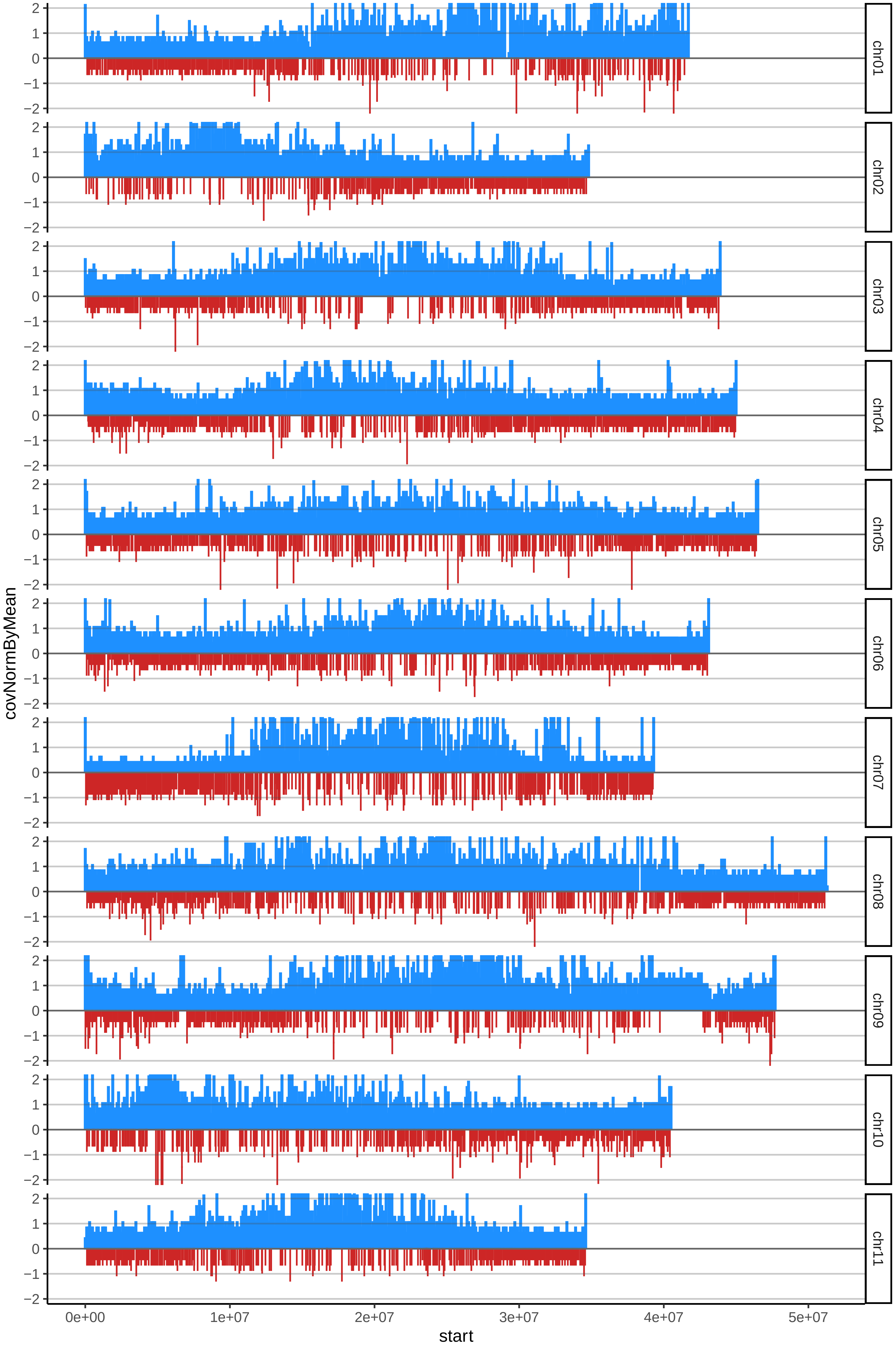

### 198CUBANOBLANCOcov_byB.pdf

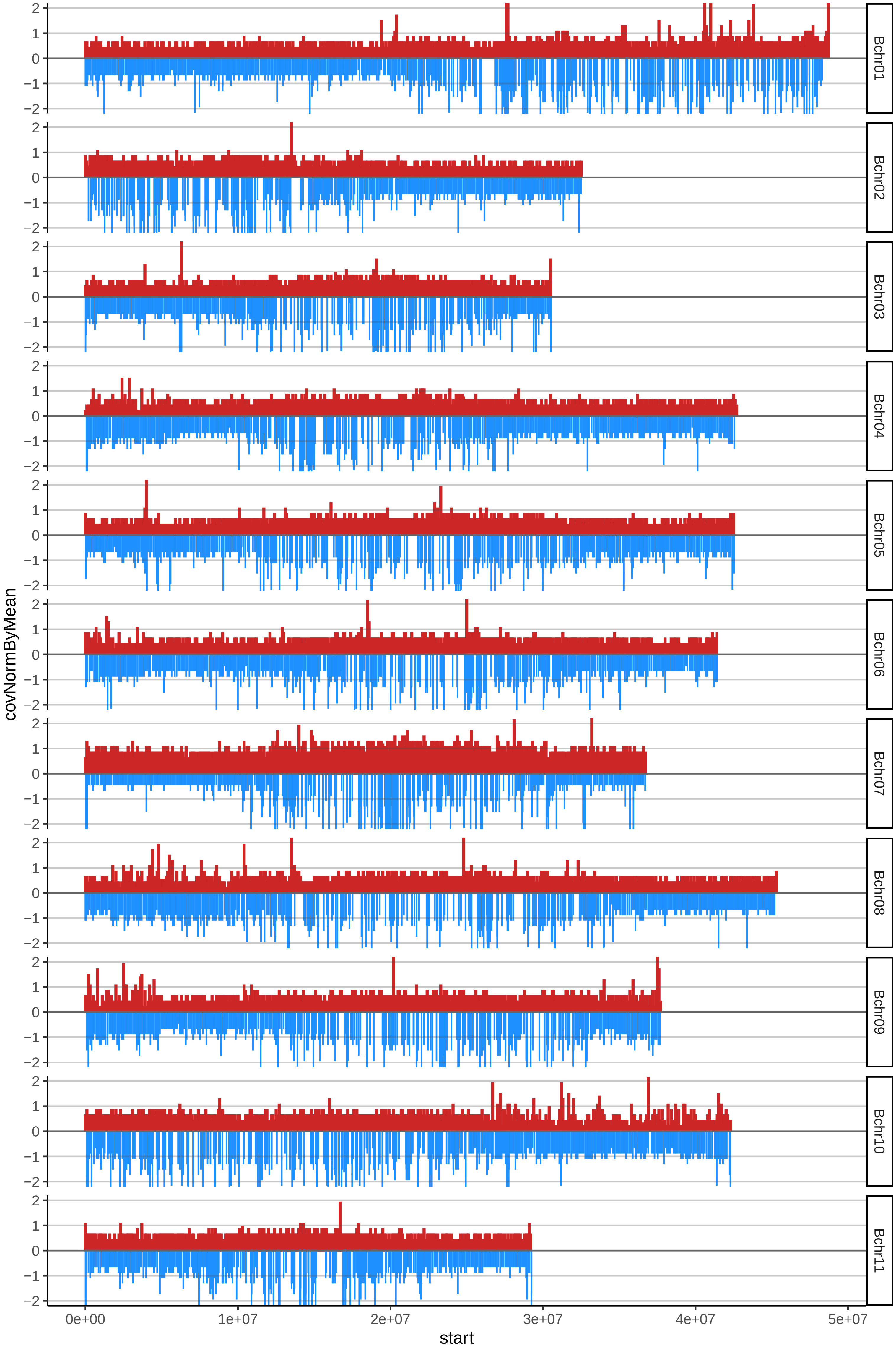

### additional file 3

A Plantain_AAB – 55 accessions

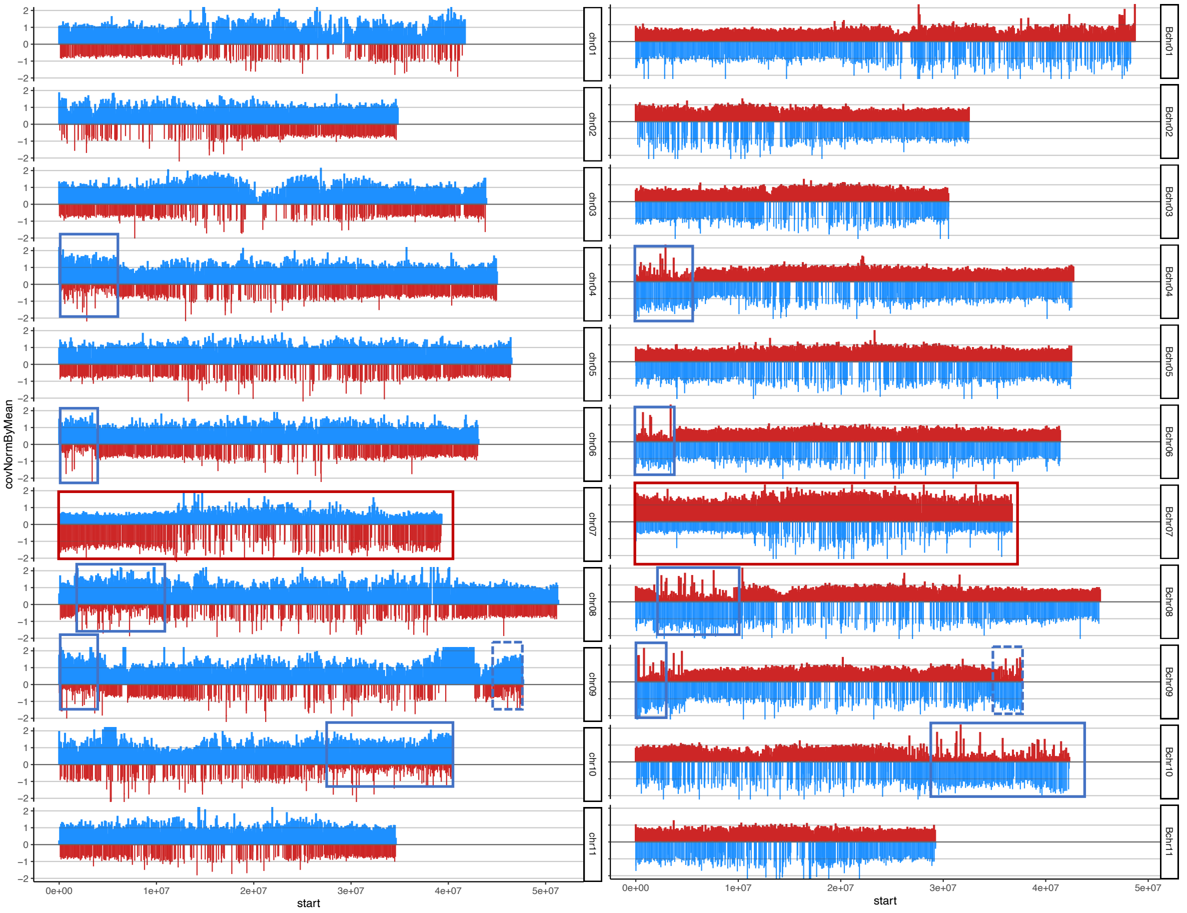


Popoulu_AAB – 5 accessions

B

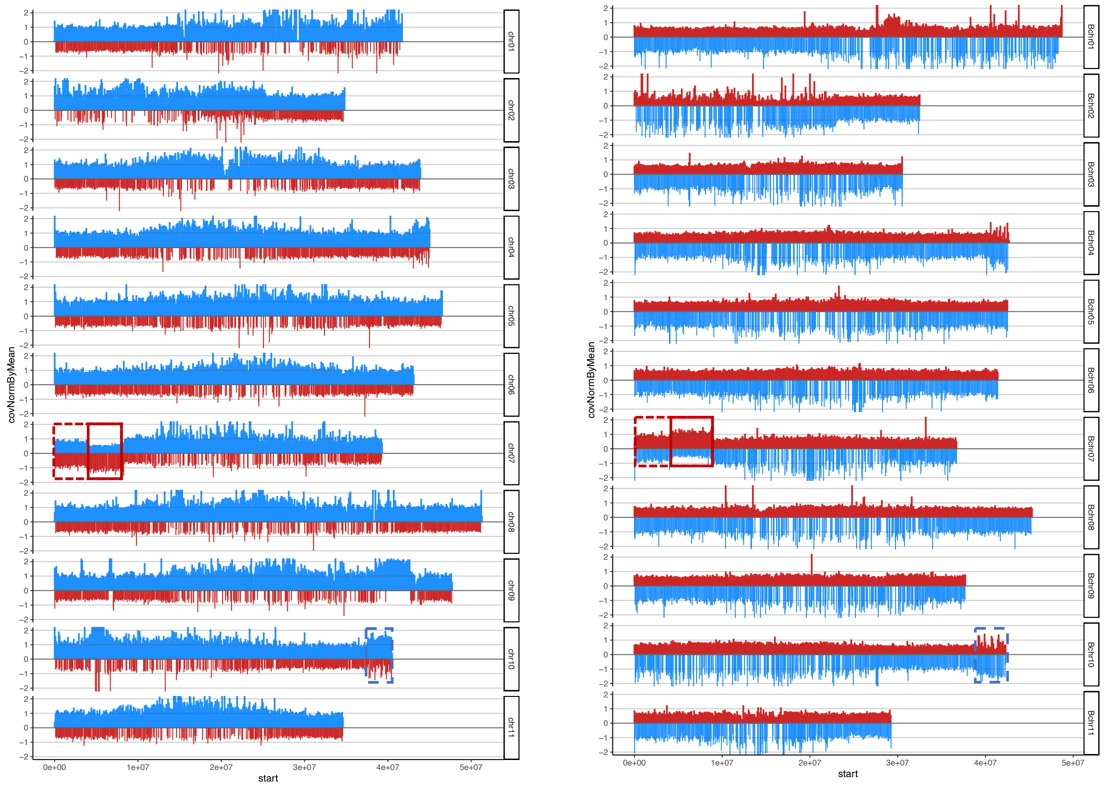


B aligned to A A aligned to B

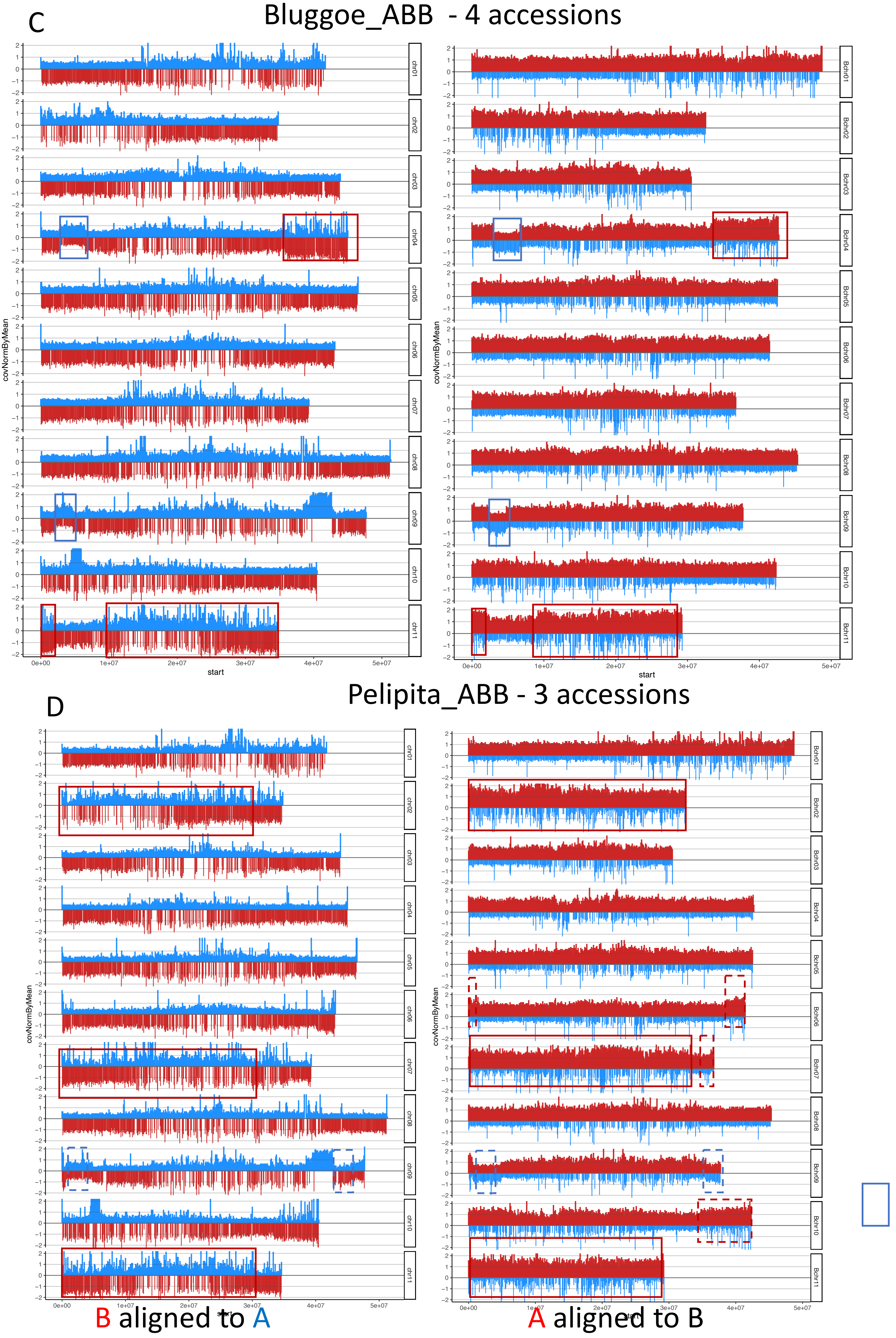

### additional file 4

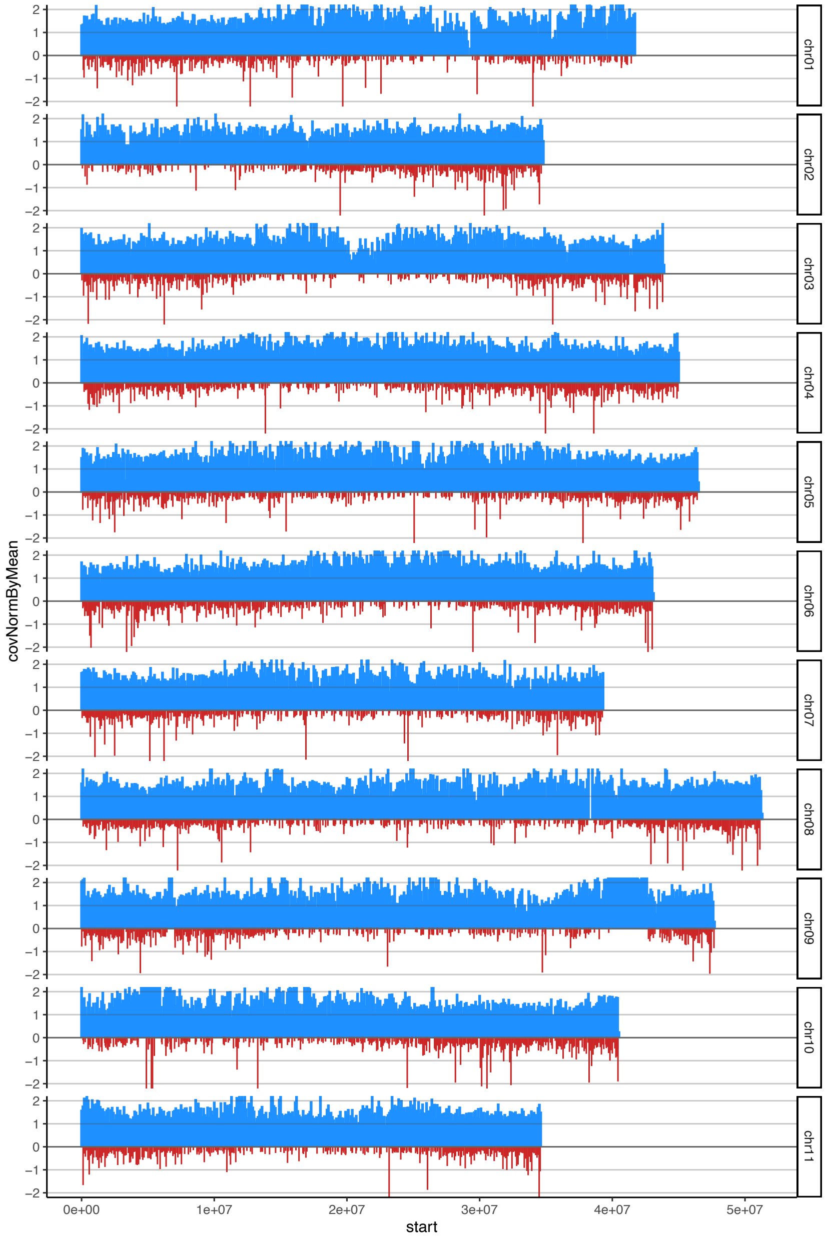

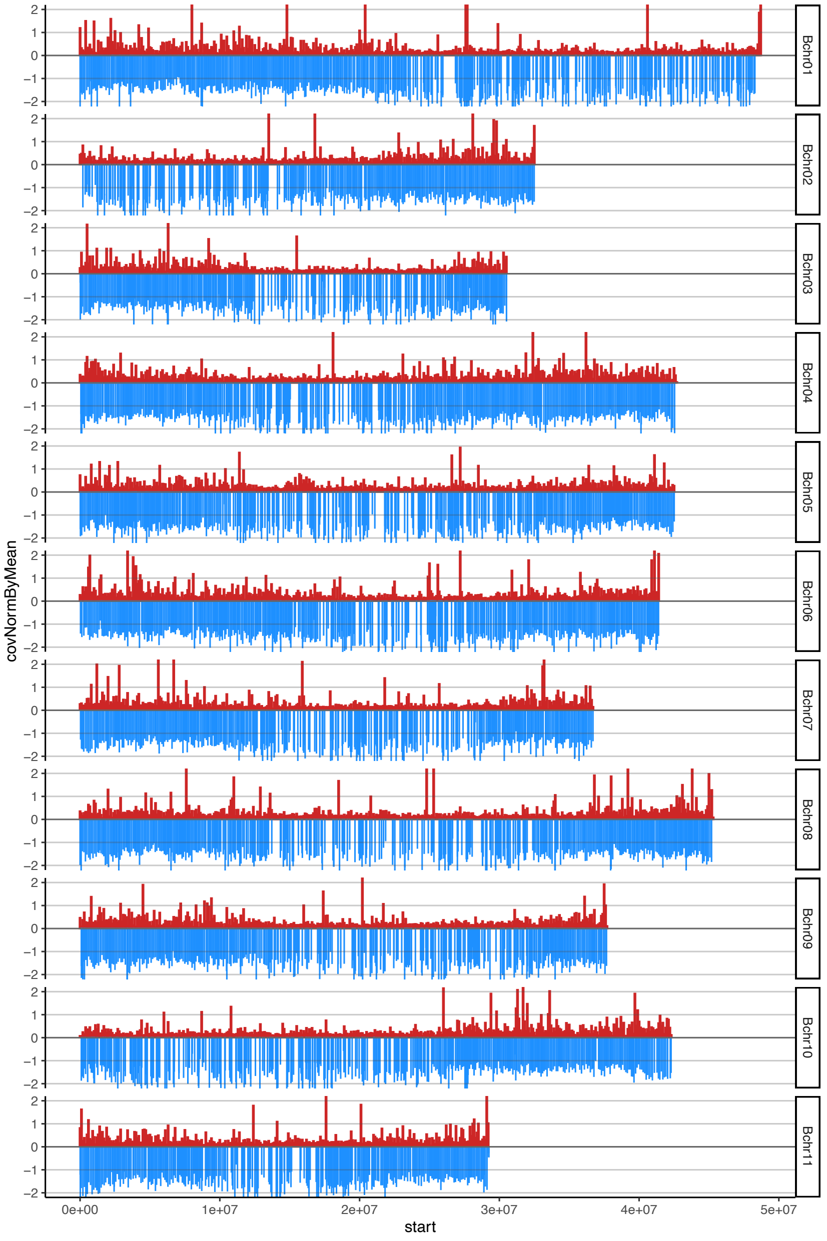

#### Cavendish _AAA

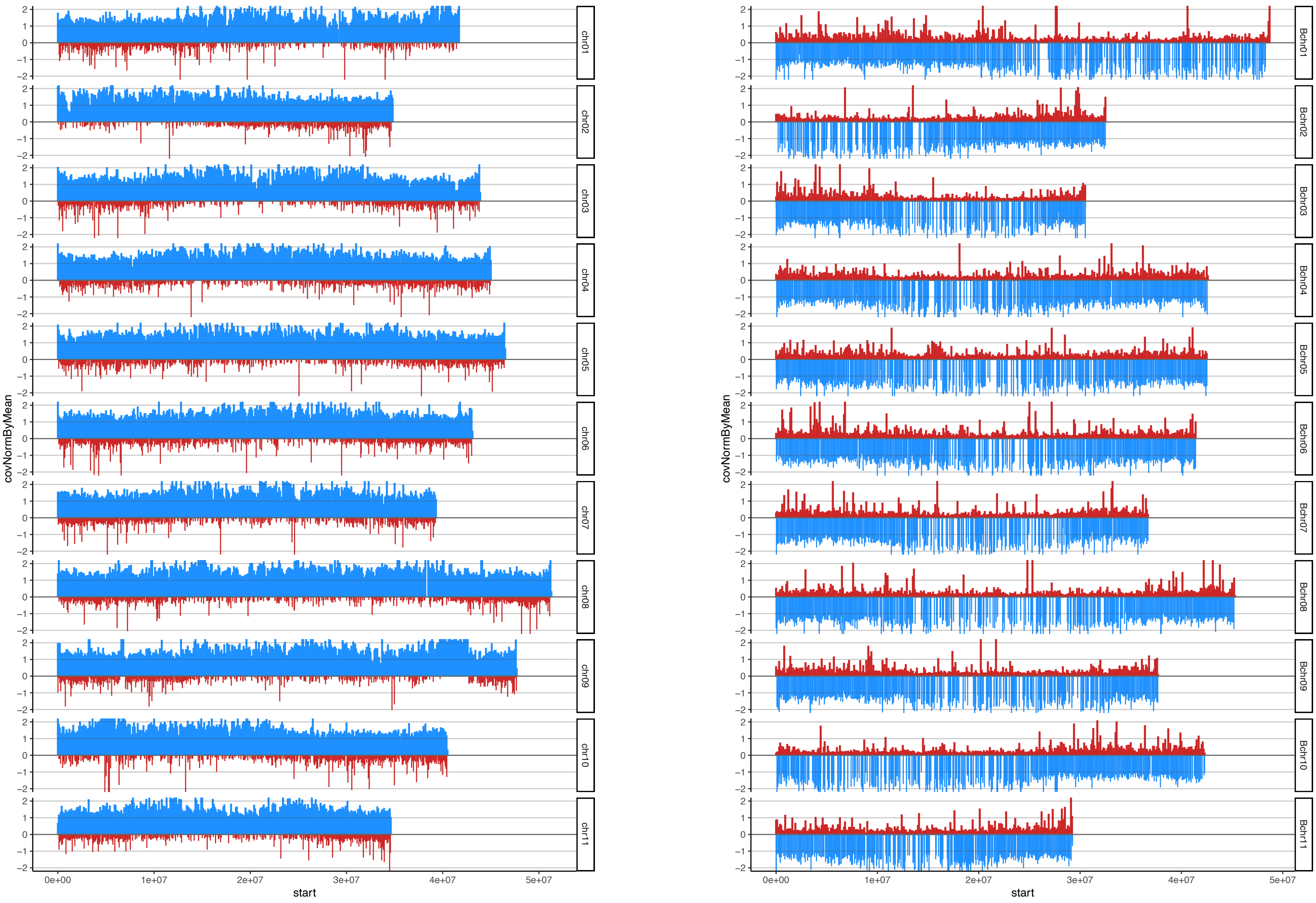

#### Gros Michel_AAA

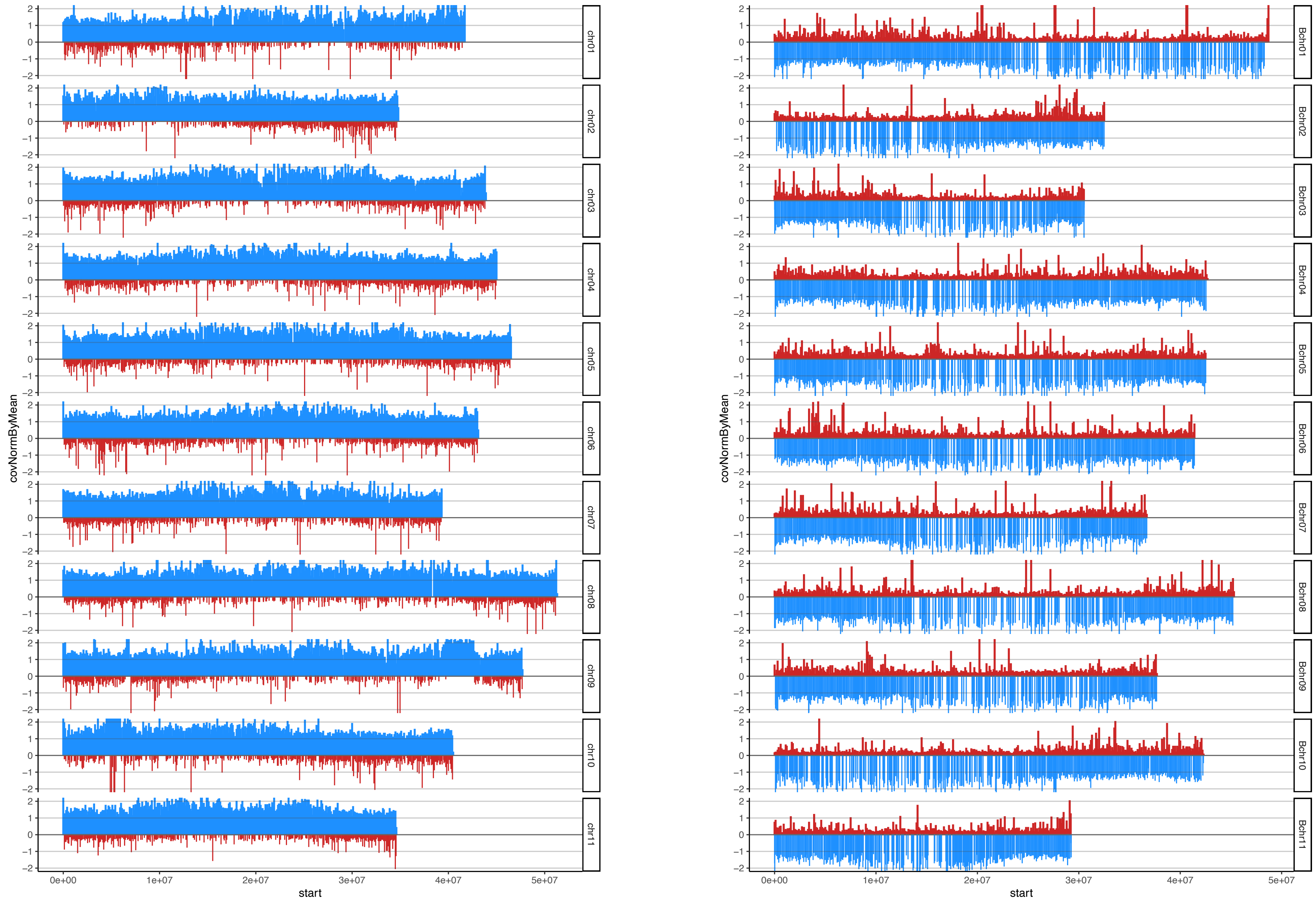

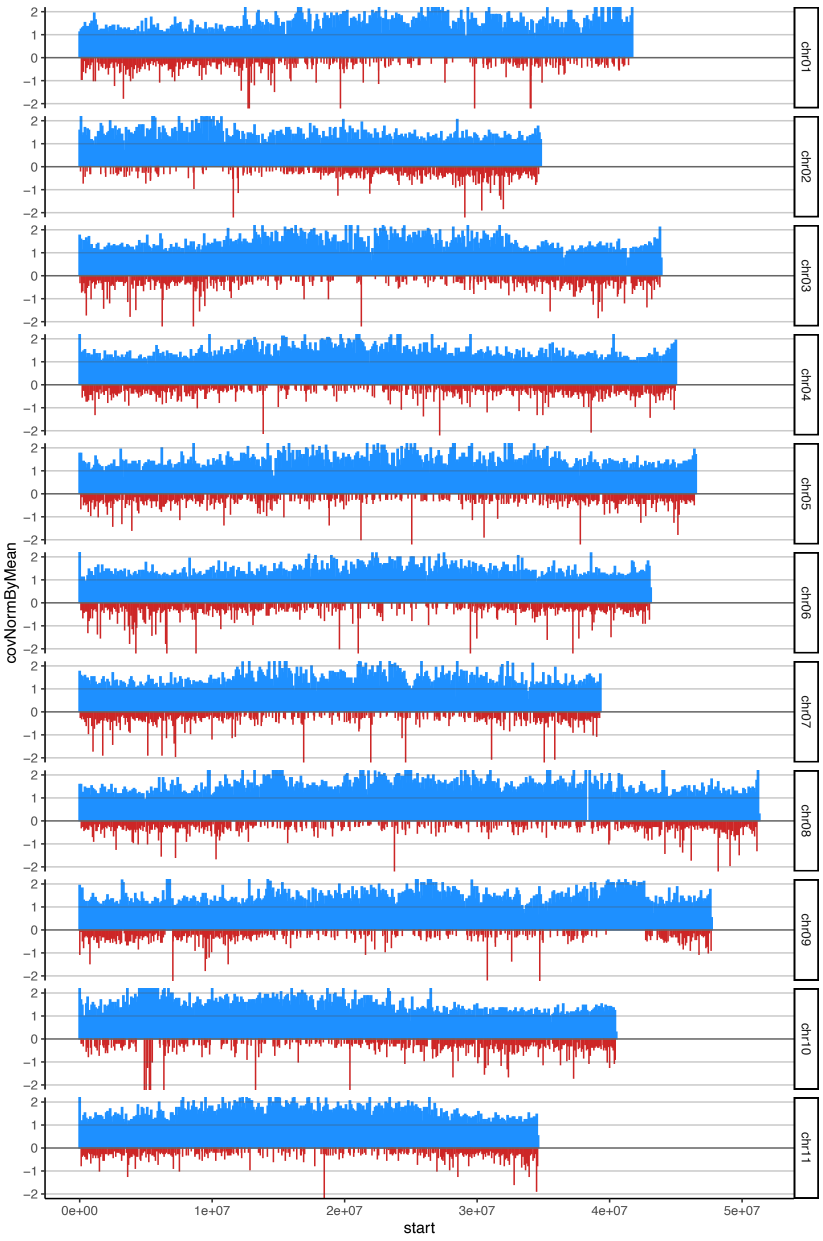

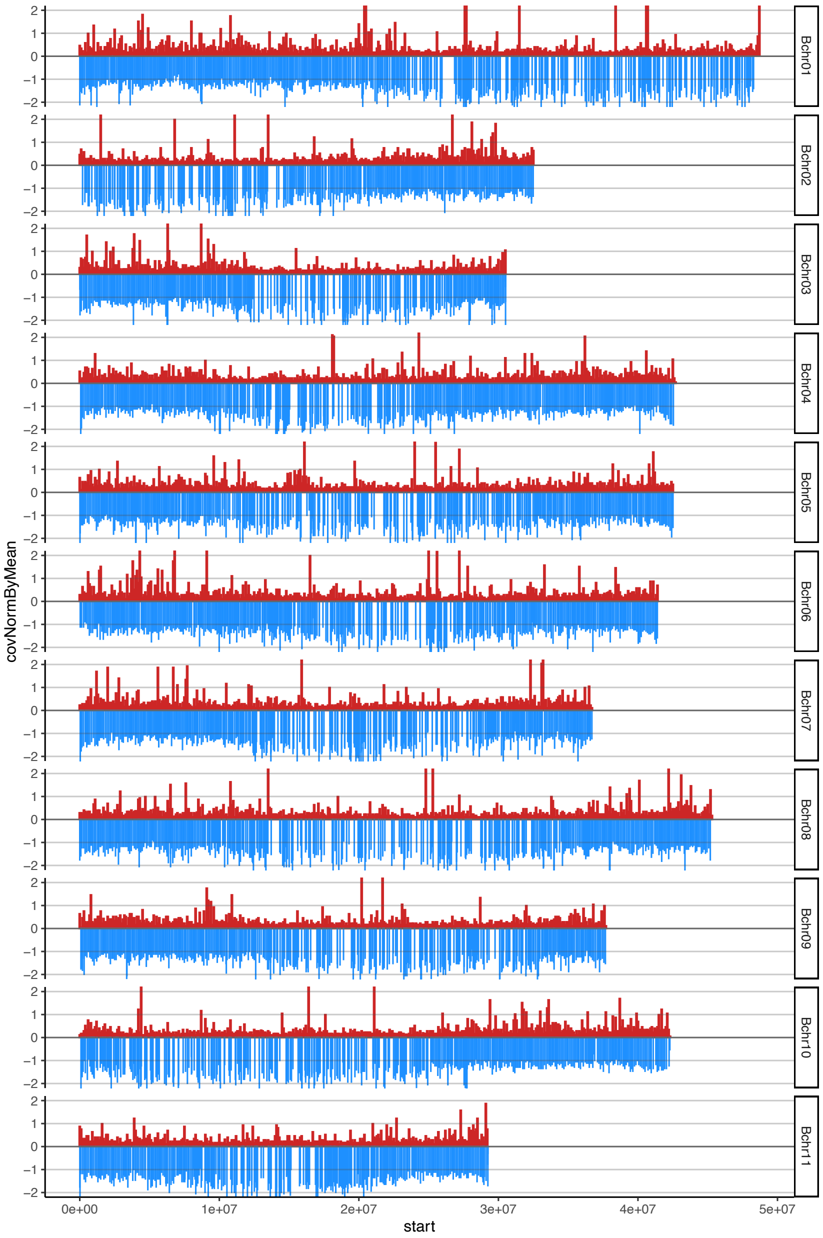

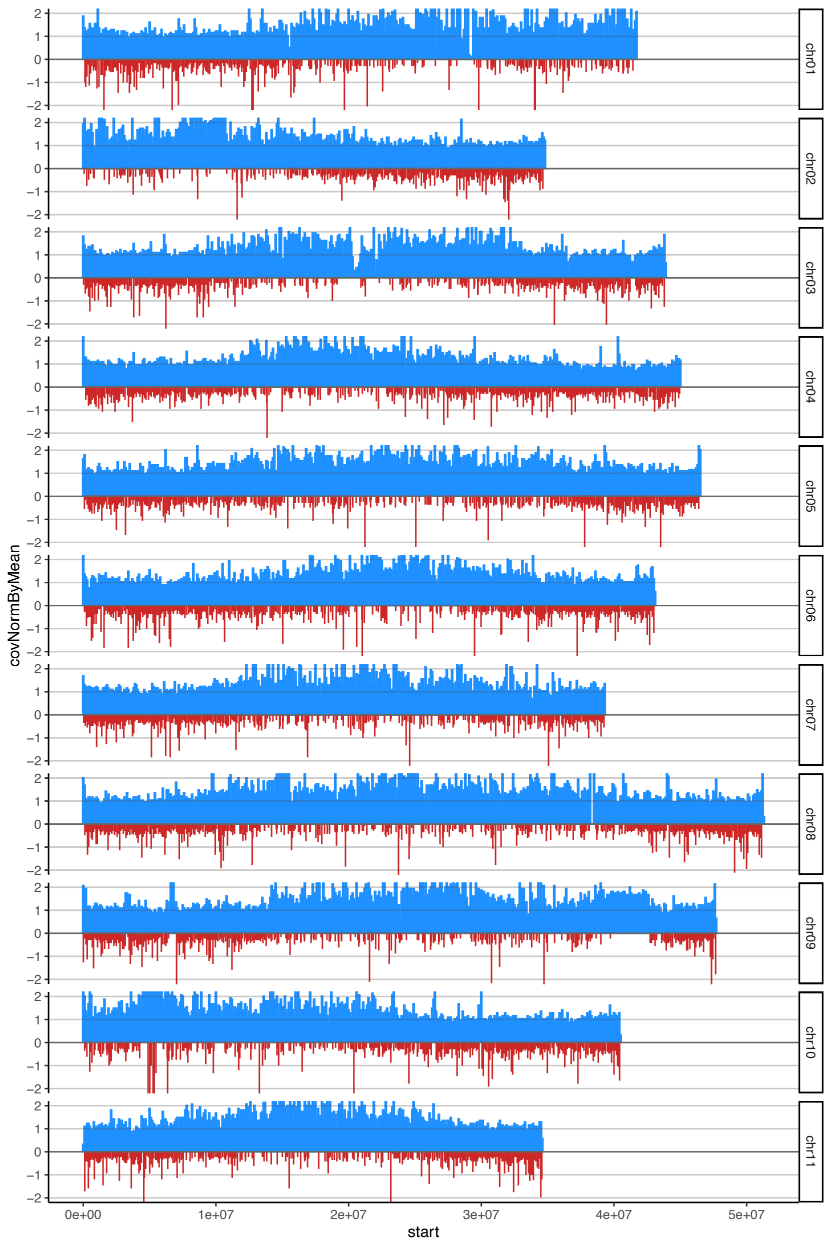

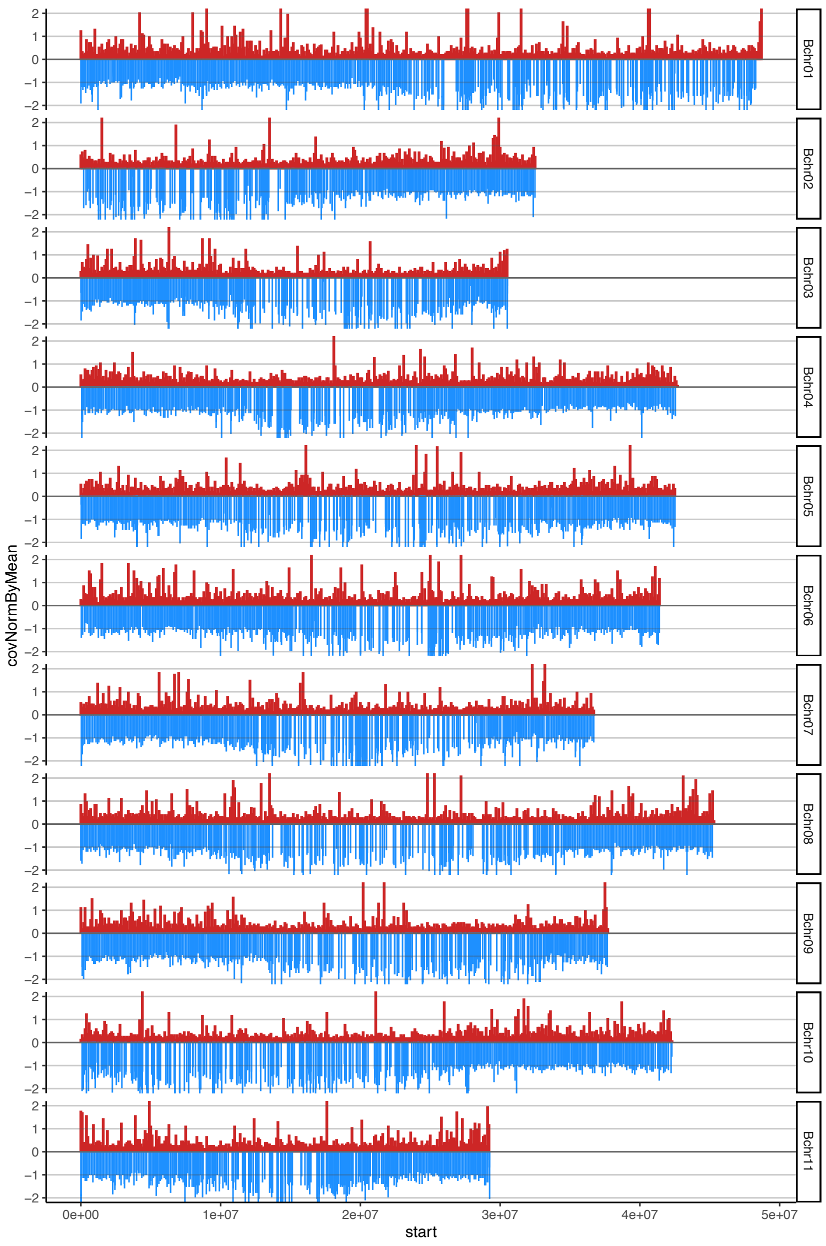

### suppl figure S1

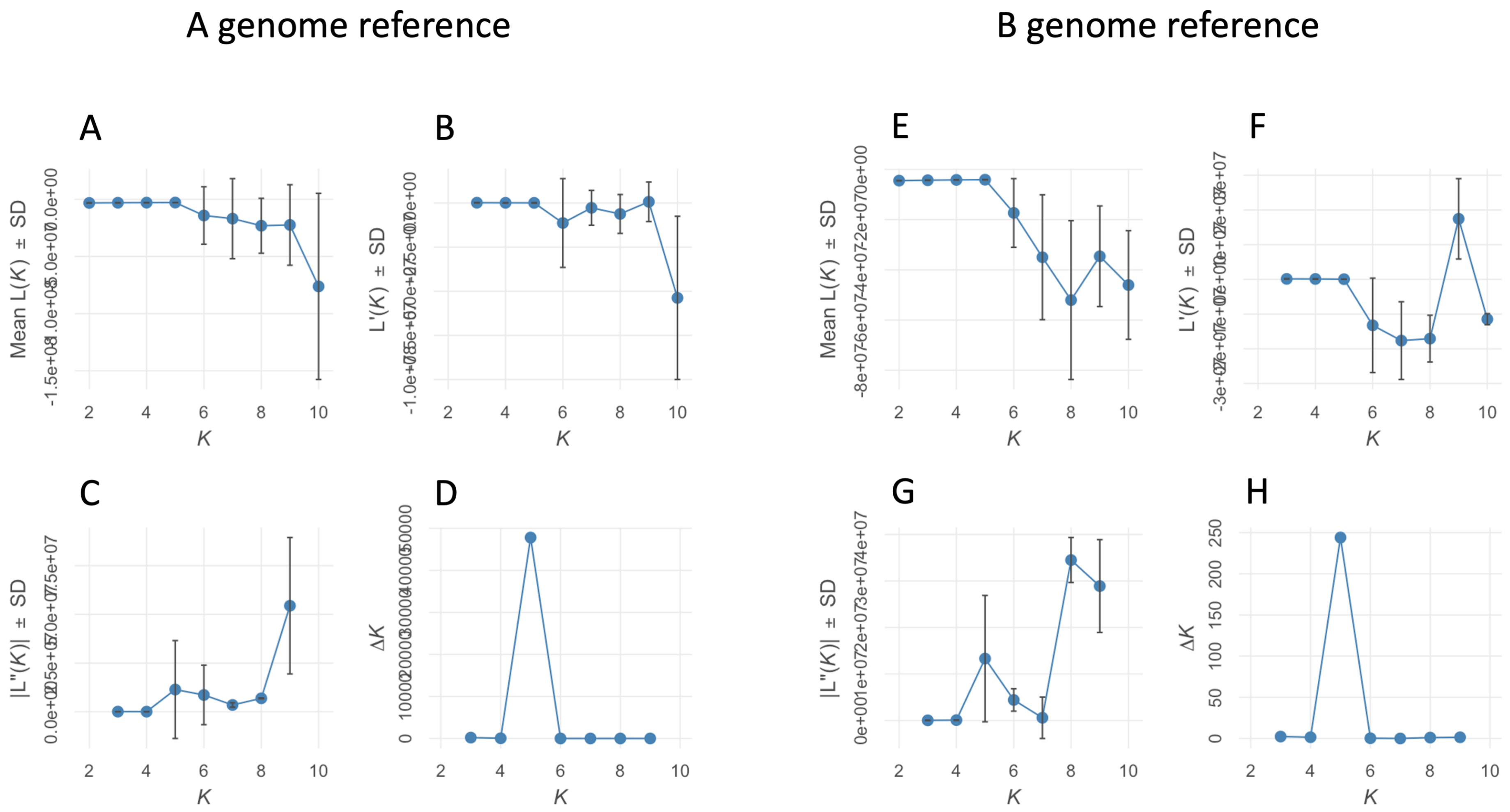

### suppl figure S2

Fig. S2. Admixture analysis for K4 to K8 against the A genome reference

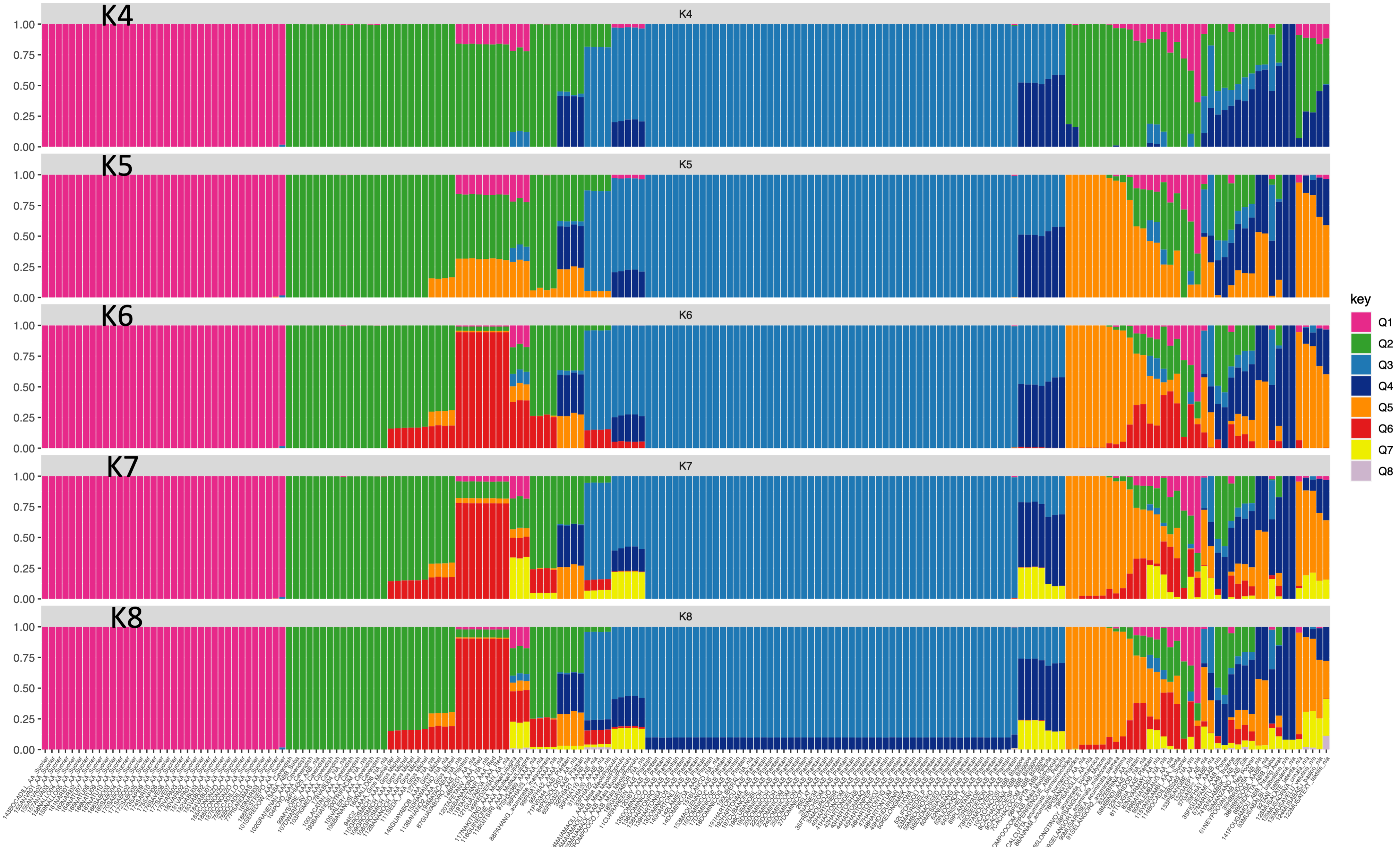

### suppl figure S3

Fig. S3. Admixture analysis for K4 to K8 against the B genome reference

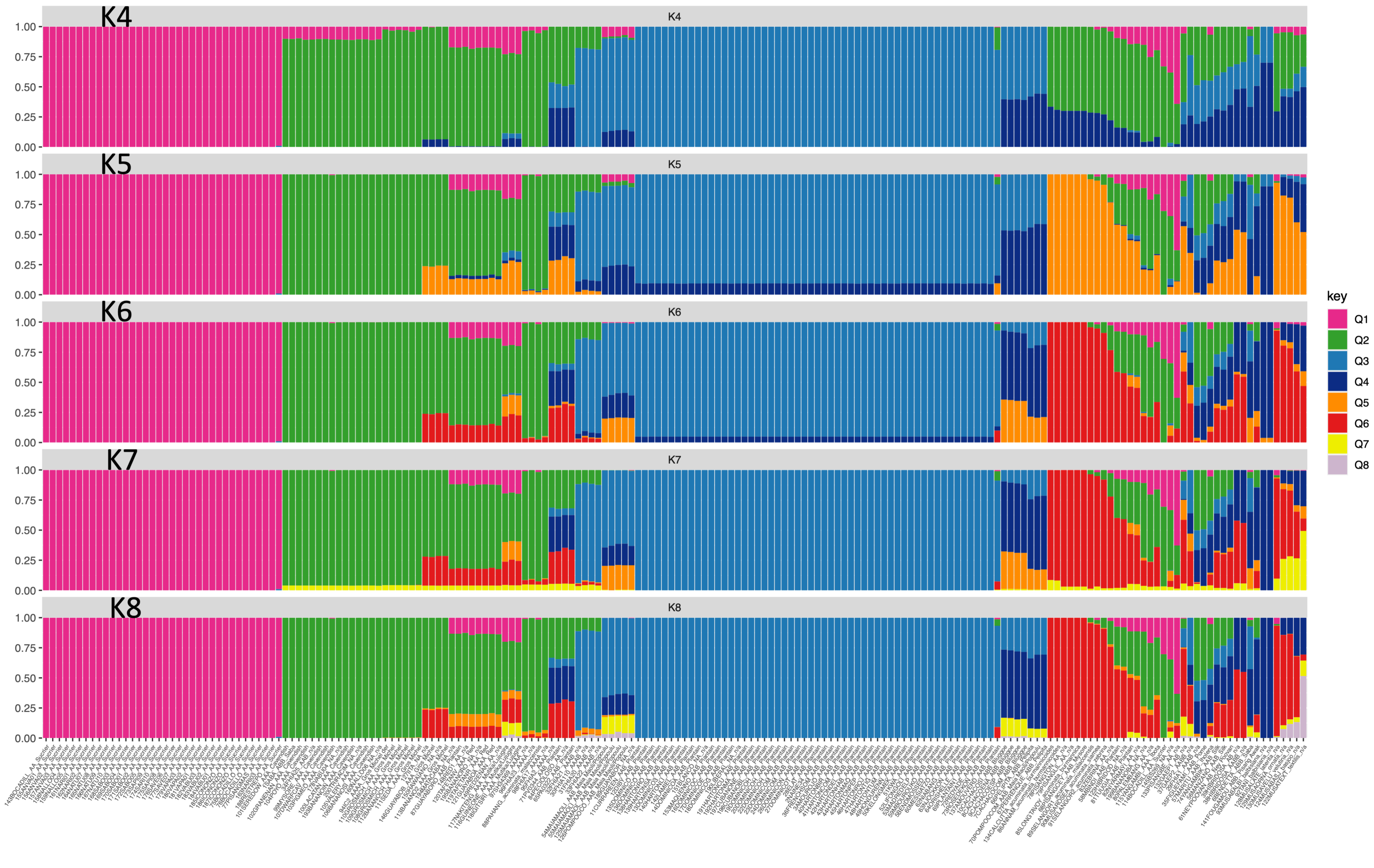

### suppl figure S5

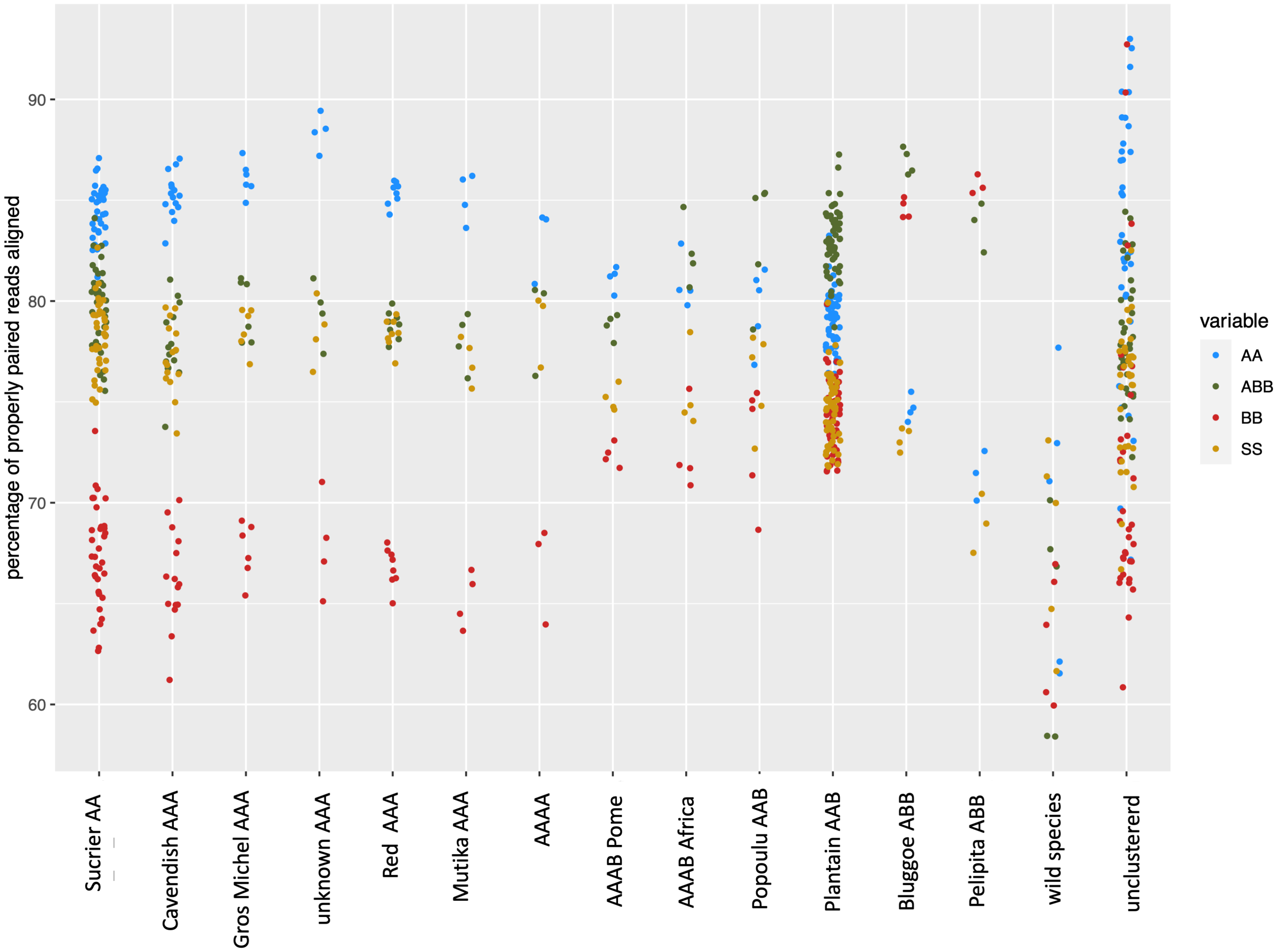

### suppl figure S6

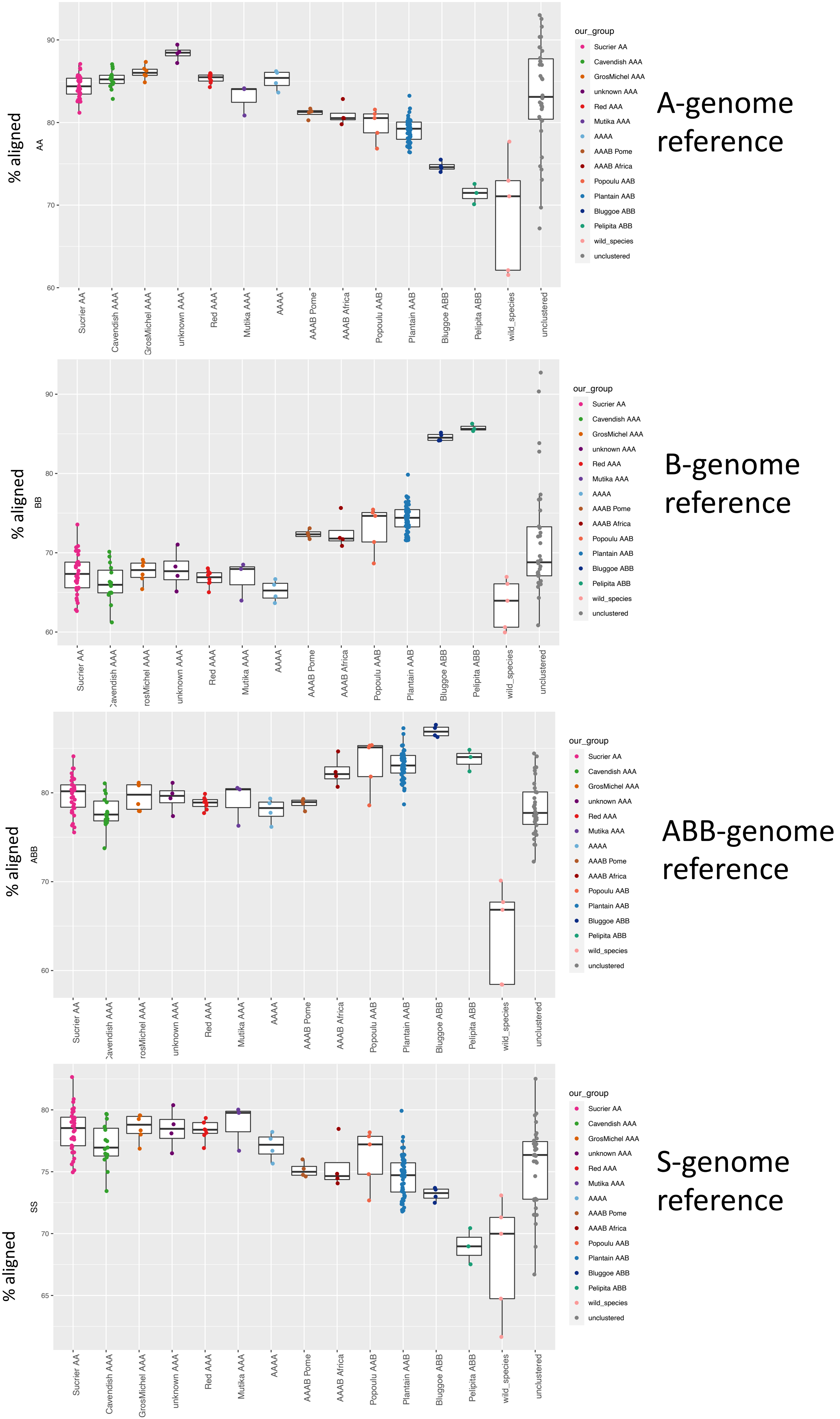

### suppl figure S7

54MAIAMAOLI

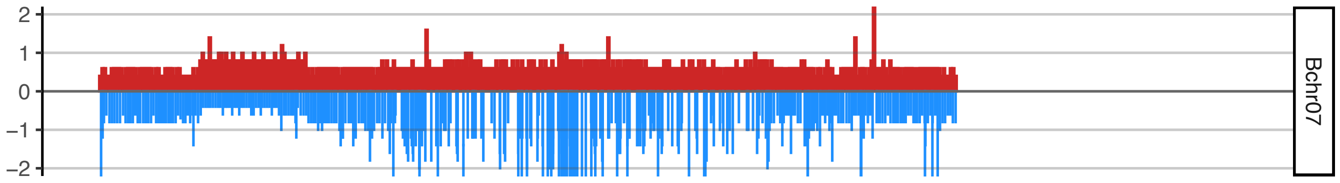


55MAIAMAOLIRISARALDA

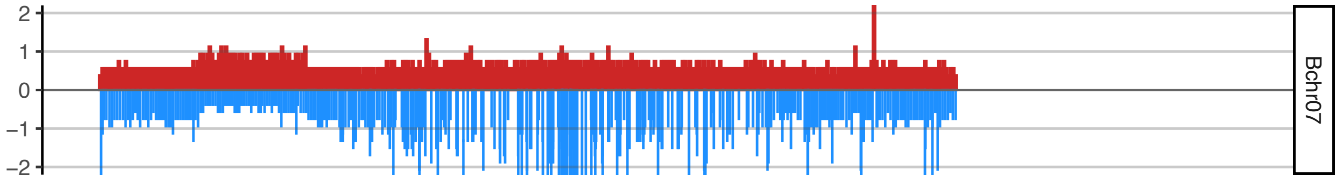


125MAIAMAOLIQUINDIO

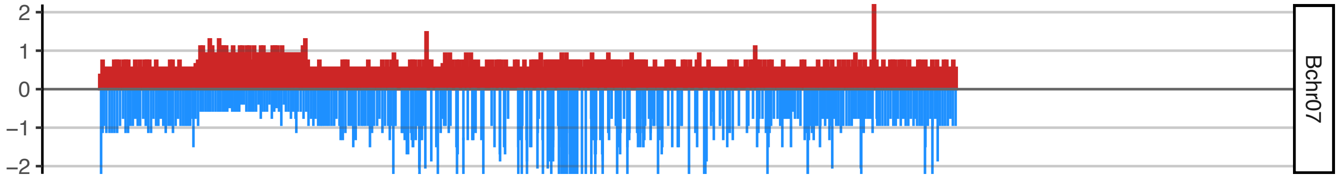


126POMPOOCOMINORISARALDA

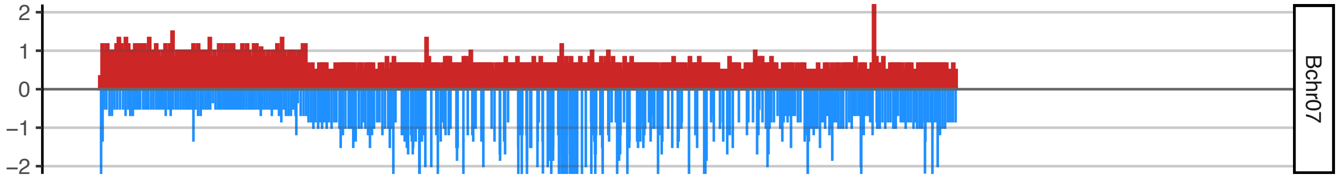


189GUAYABORAYADO

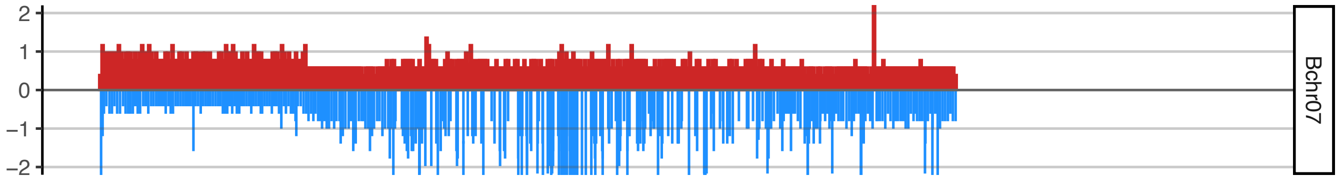

### suppl figure S8

#### AAAB_Pome

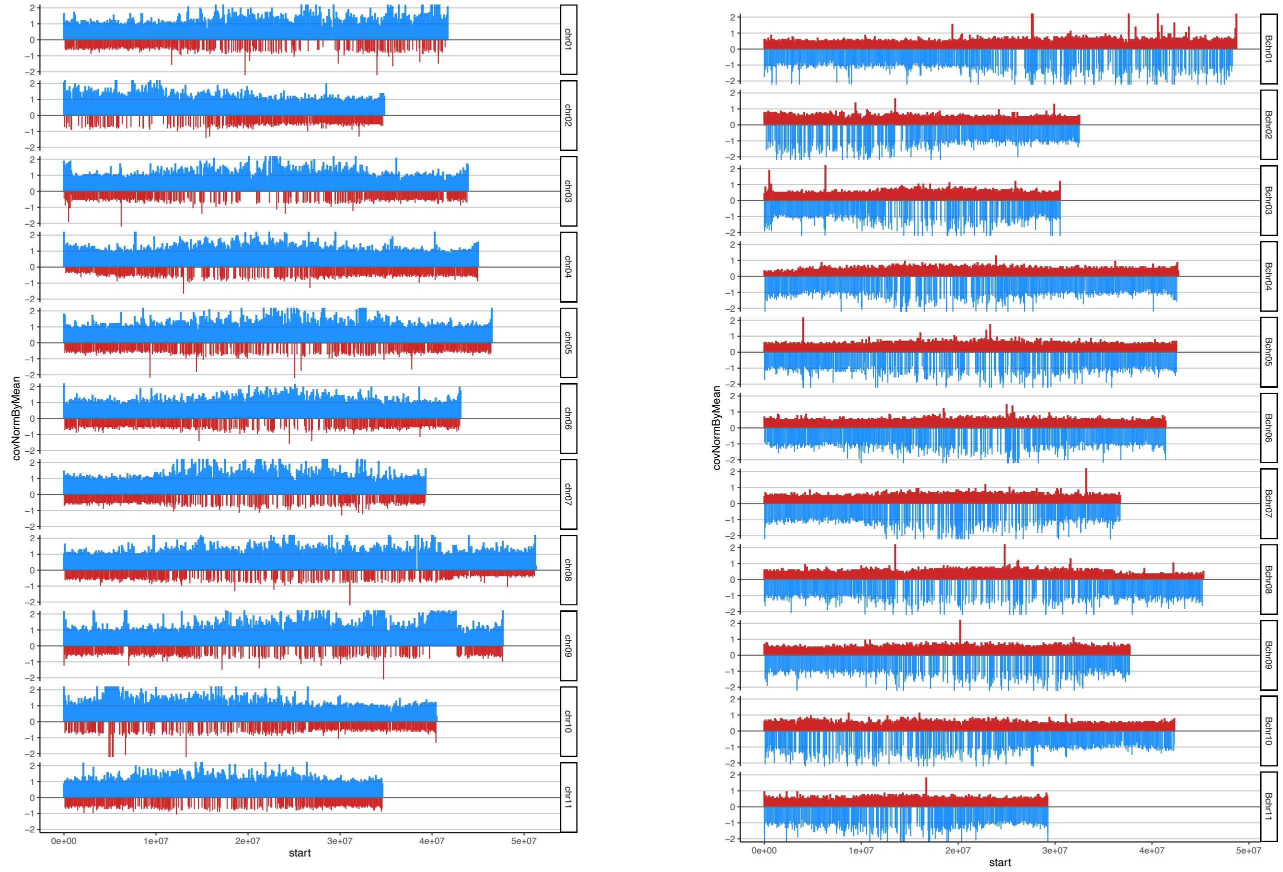


#### AAAB_Africa

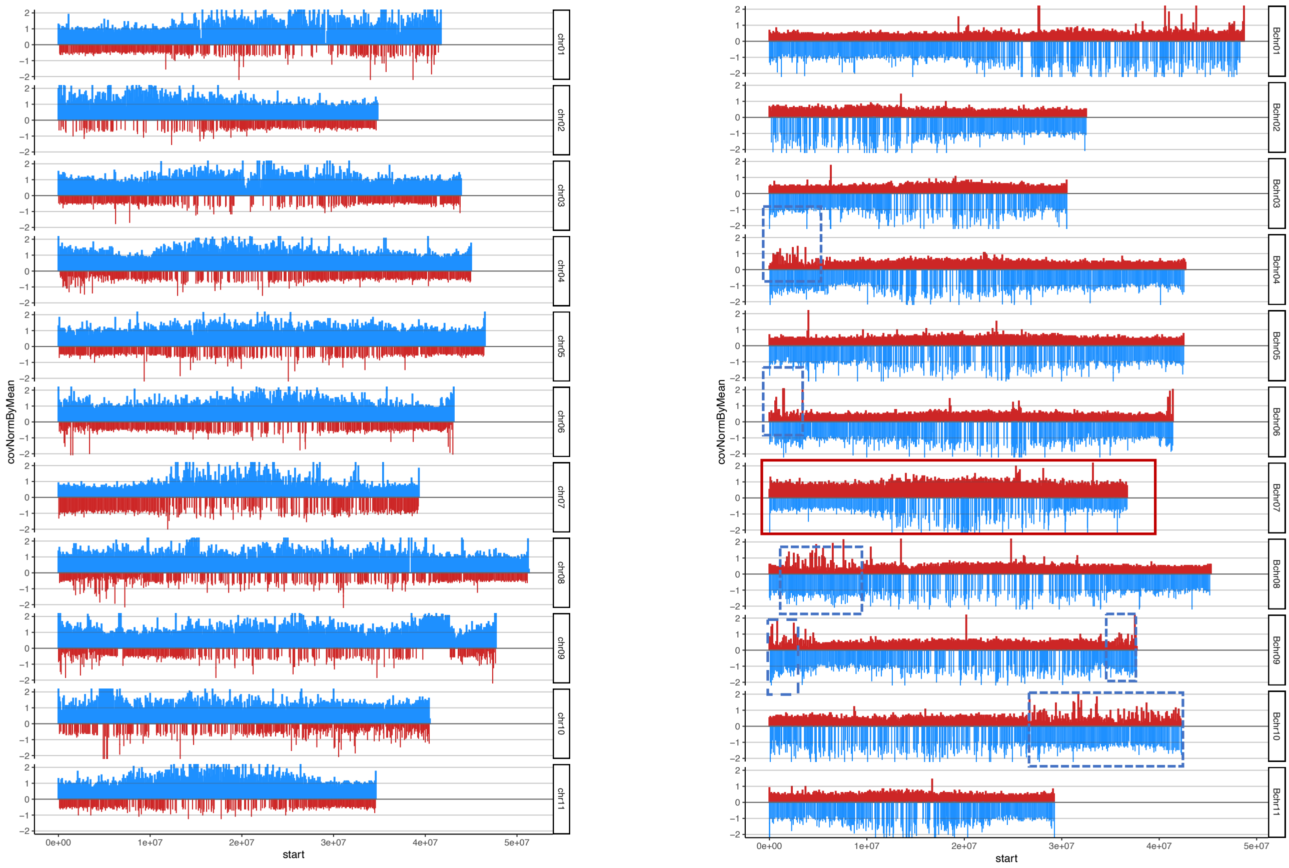

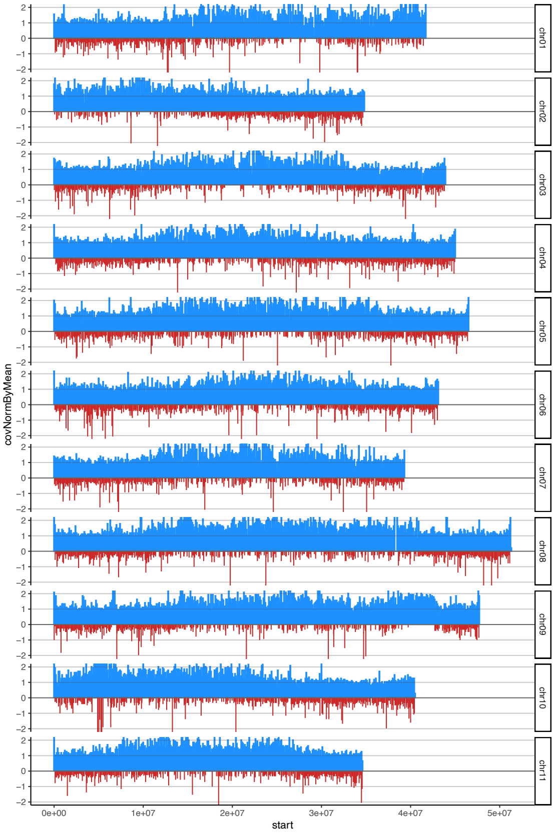

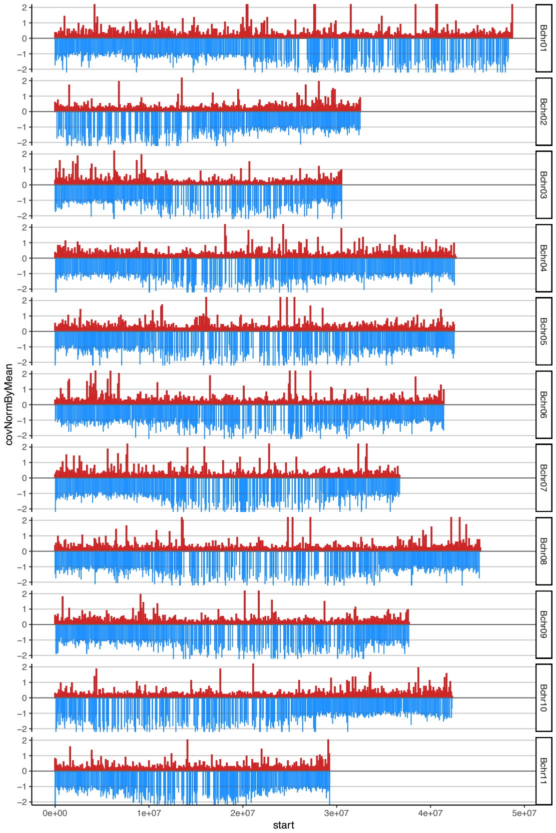

### suppl figure S9

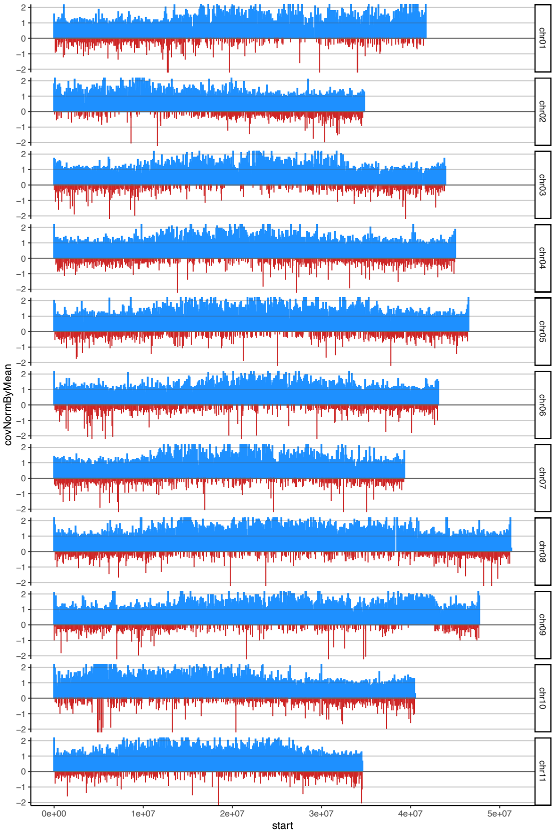

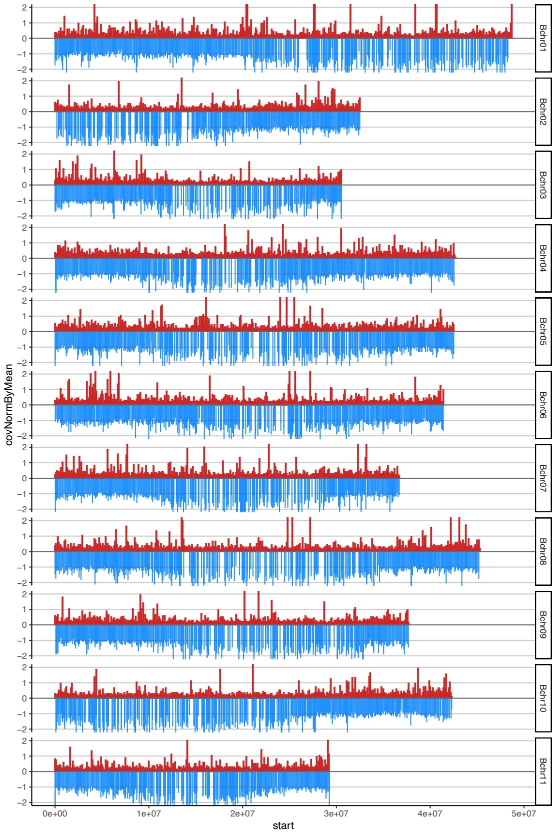
