## Supplementary material for "Characterising genome composition and large structural variation in banana varietal groups": suppl figure S4

Fig. S4. Comparison of clusters identified using admixture analysis in the A vs. B genome references

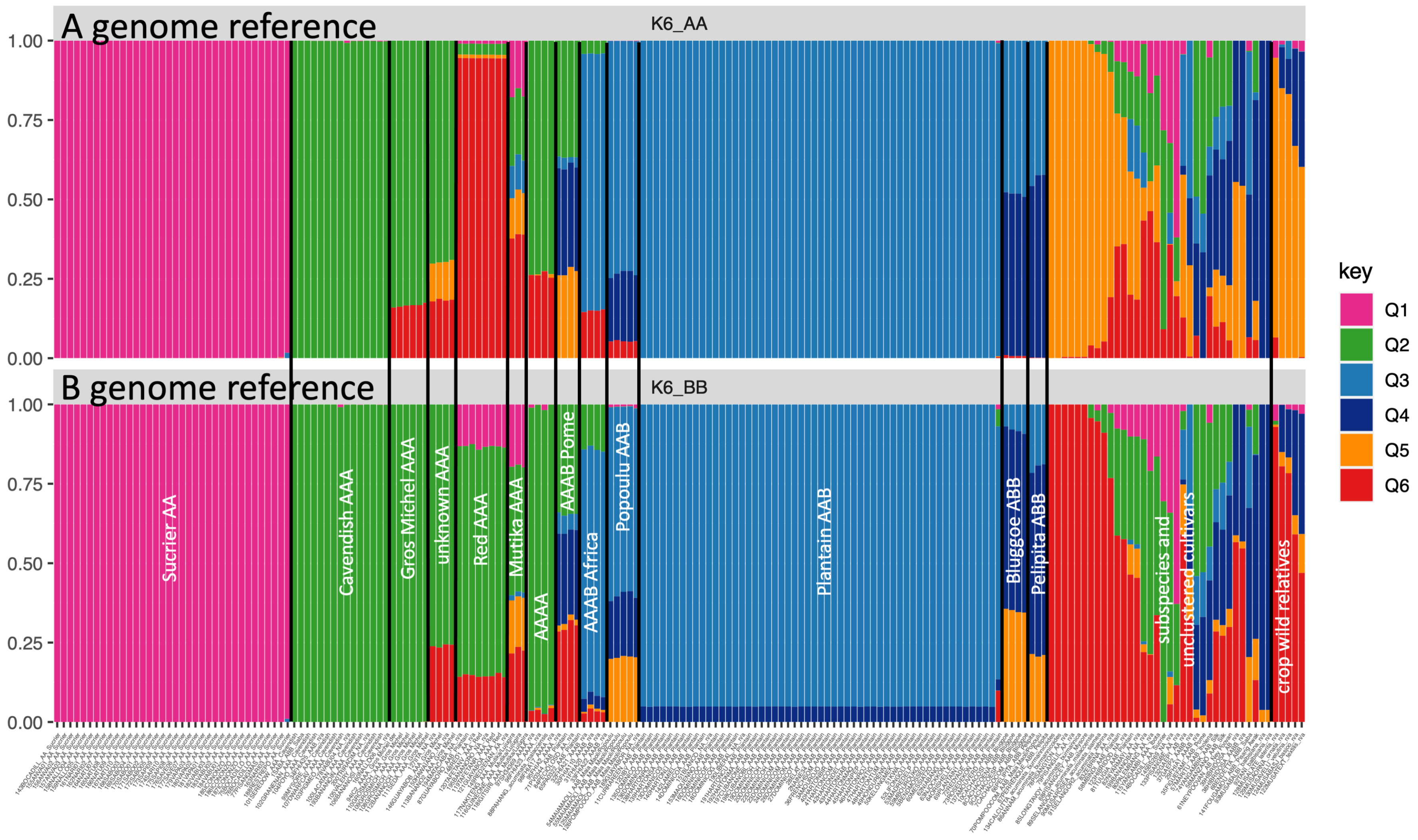
