## additional file 2 for "Characterising genome composition and large structural variation in banana varietal groups"

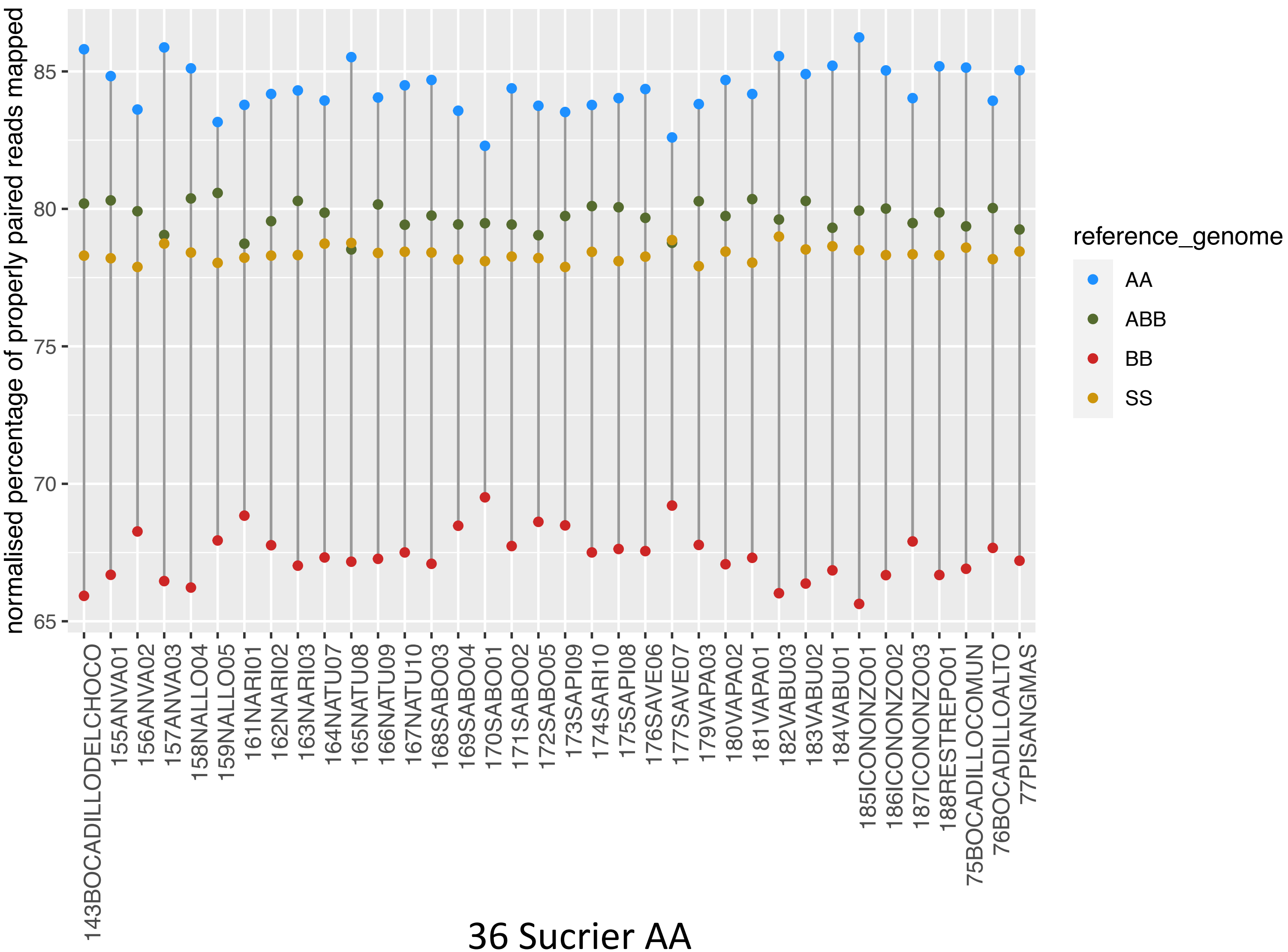


36 Sucrier AA

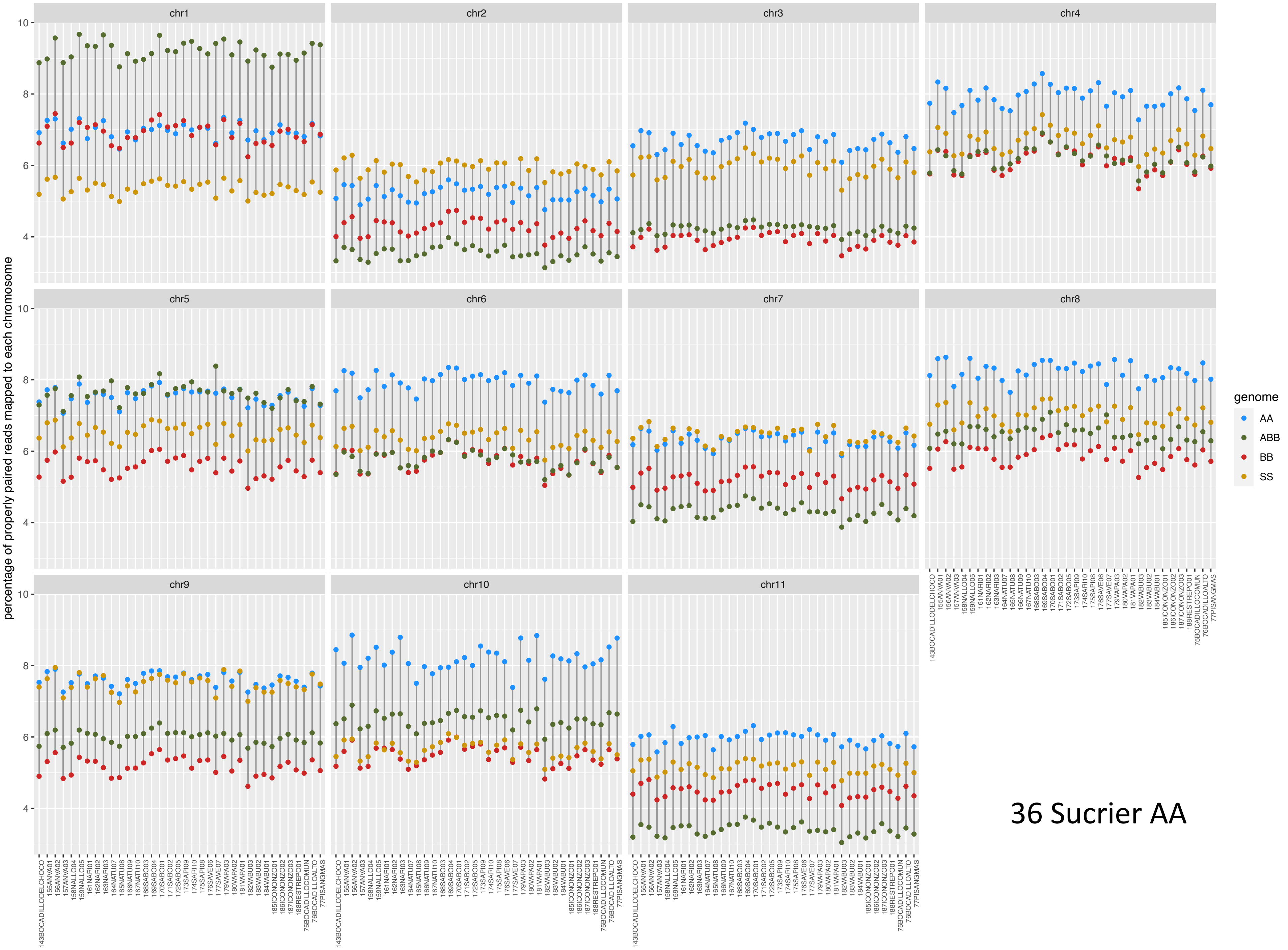


36 Sucrier AA

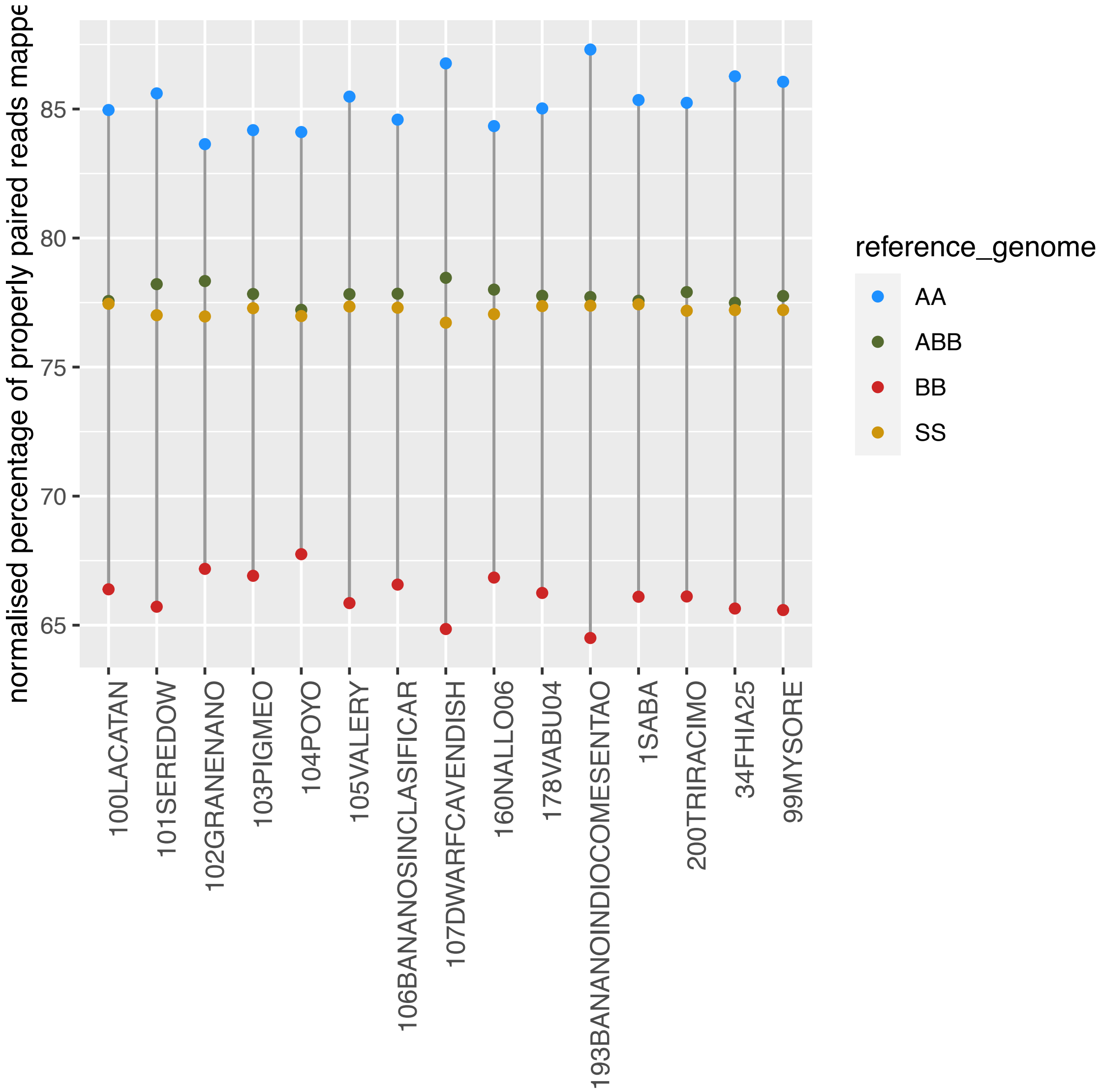


15 Cavendish

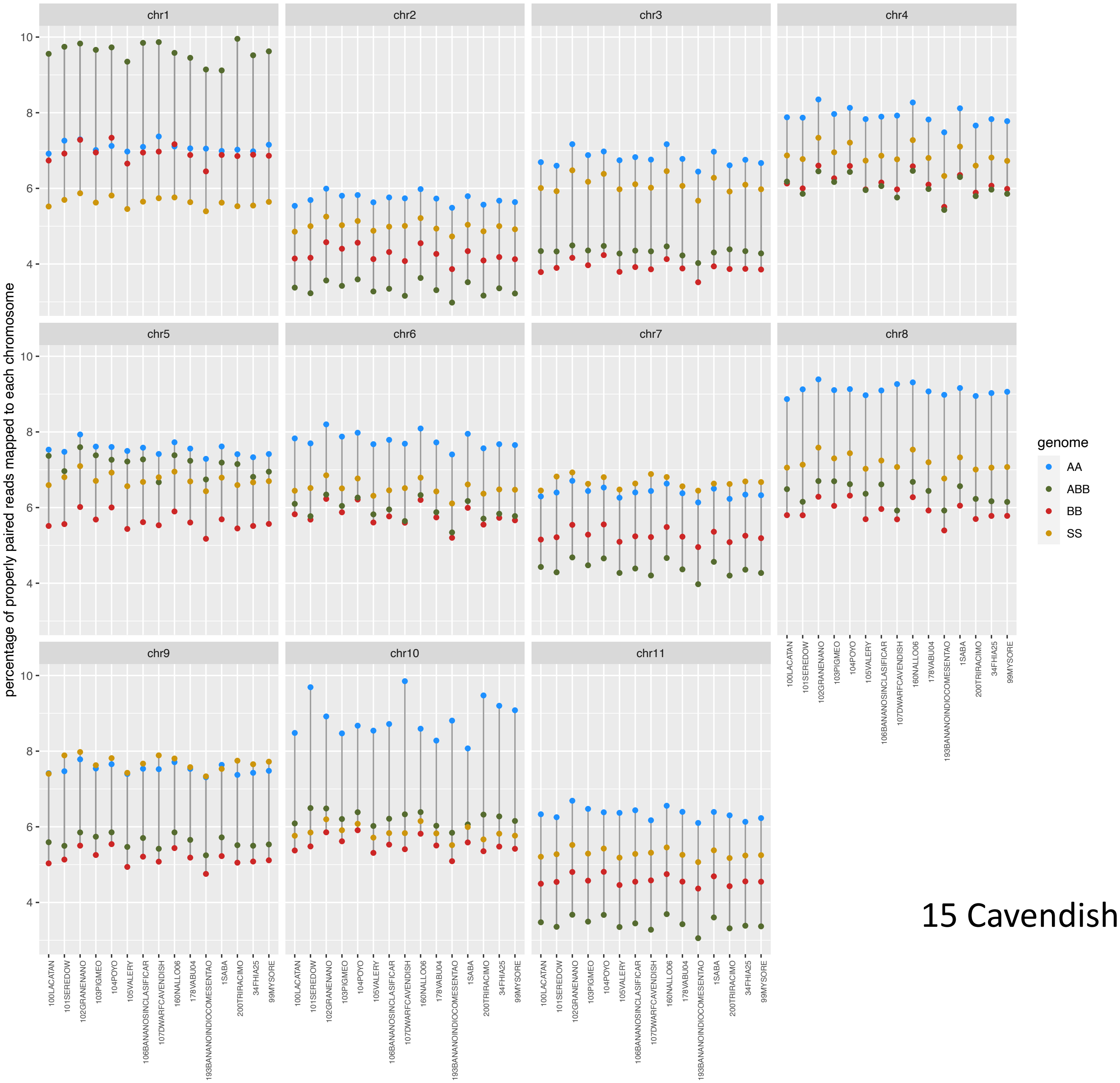


15 Cavendish AAA

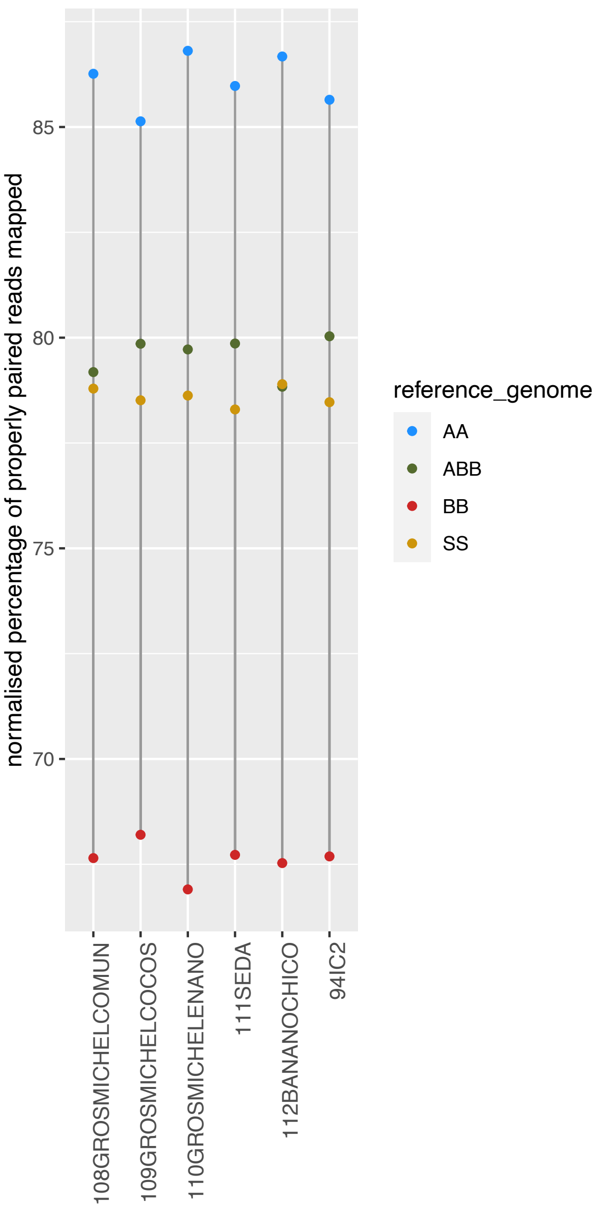


6 Gros Michel AAA

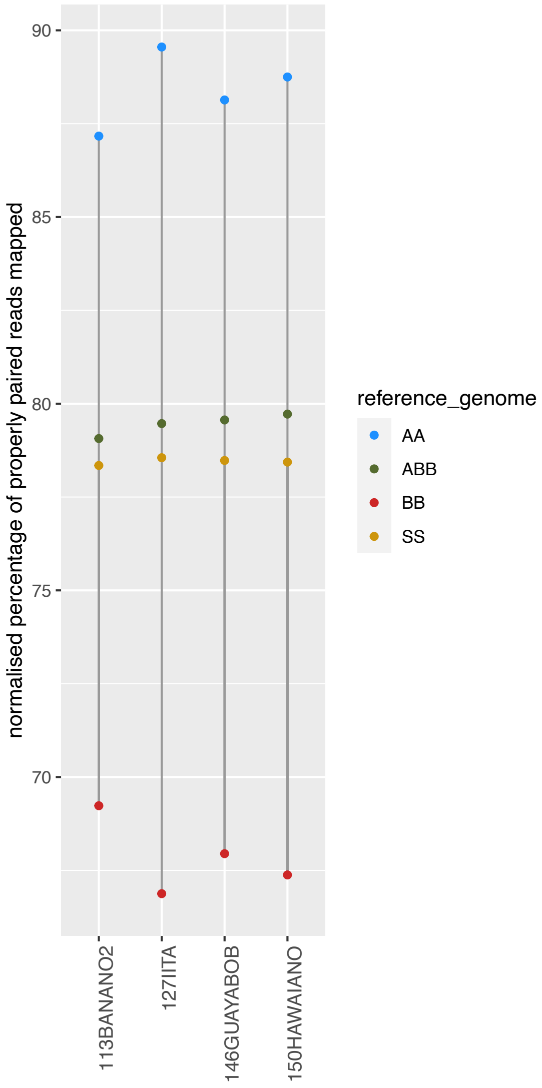


4 unknown AAA

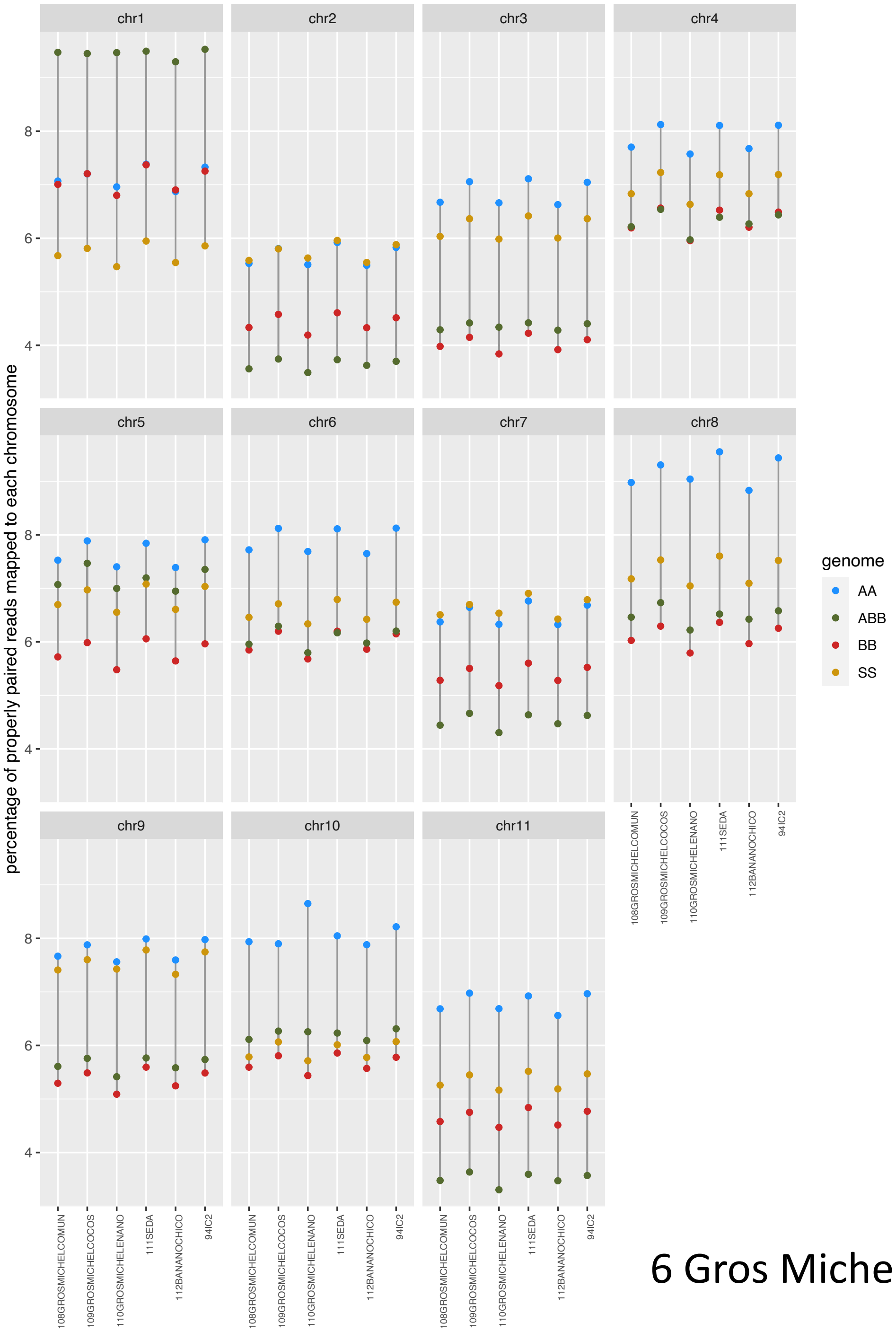


6 Gros Michel AAA

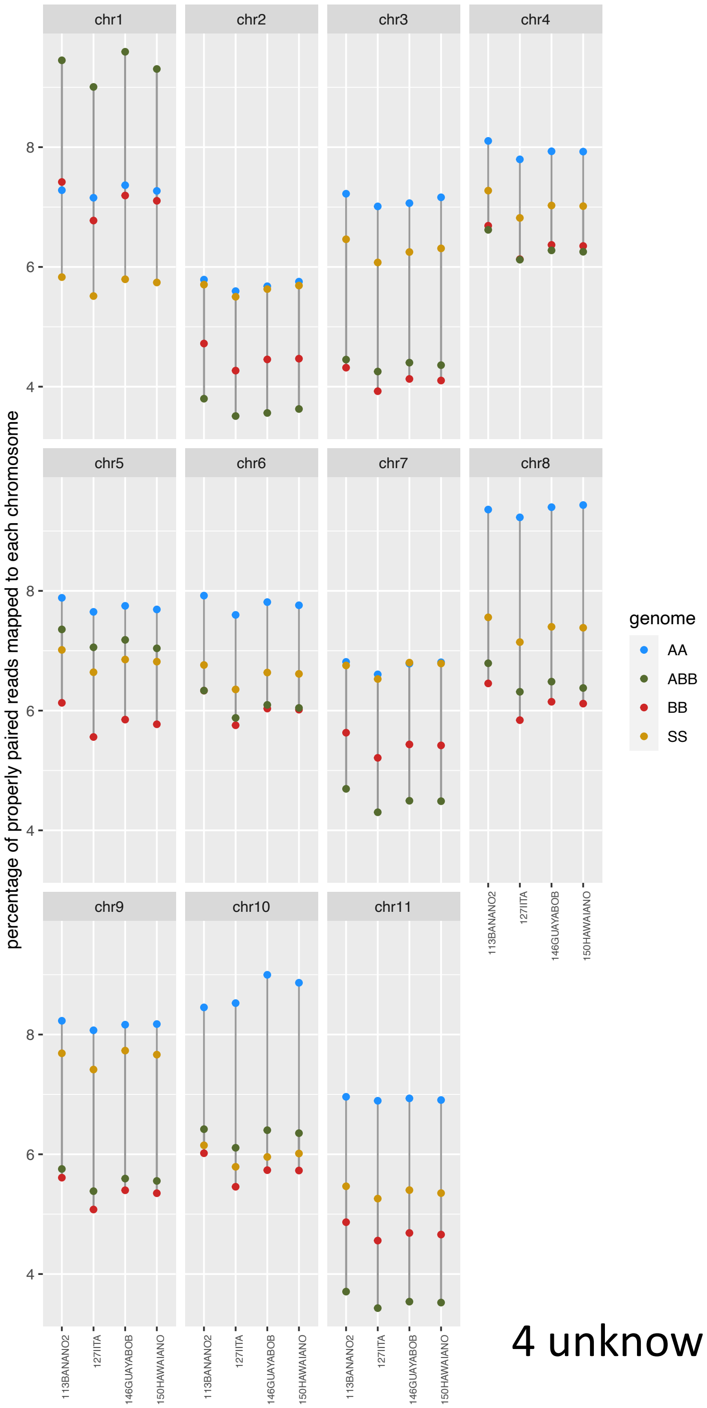


4 unknown AAA

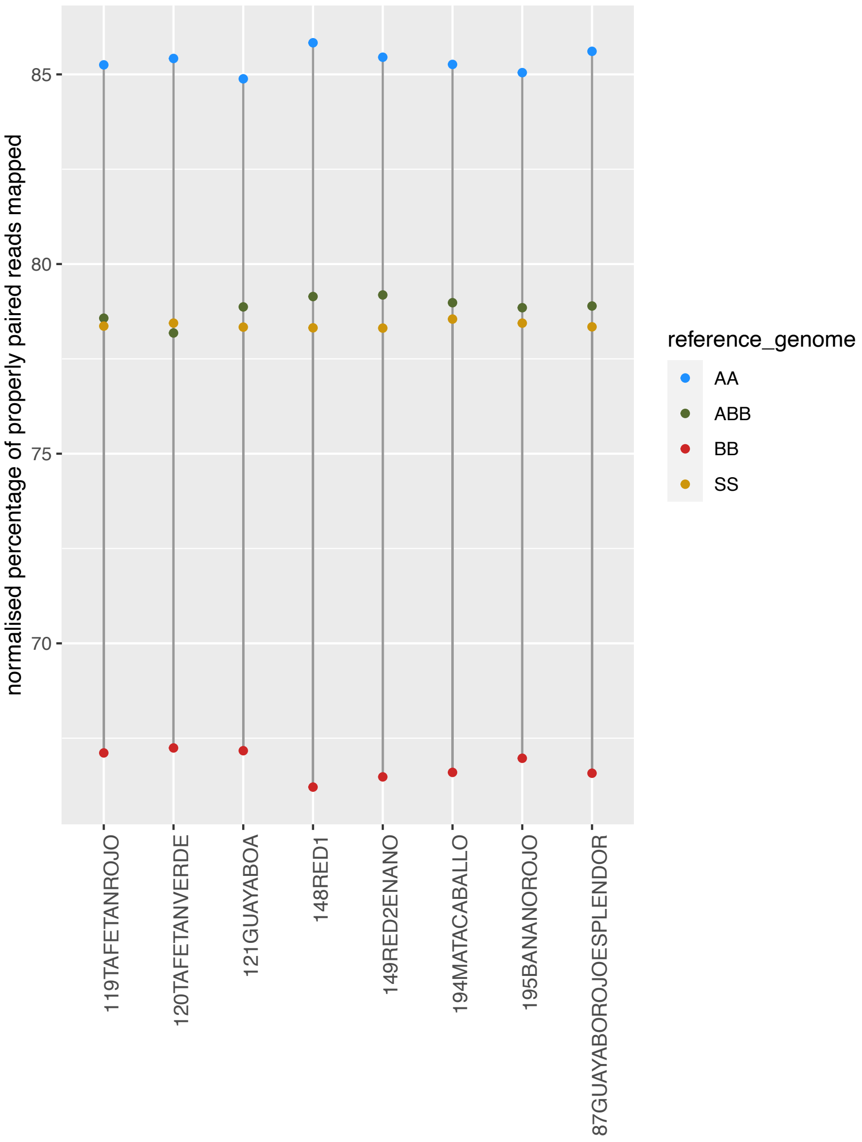


8 Red AAA

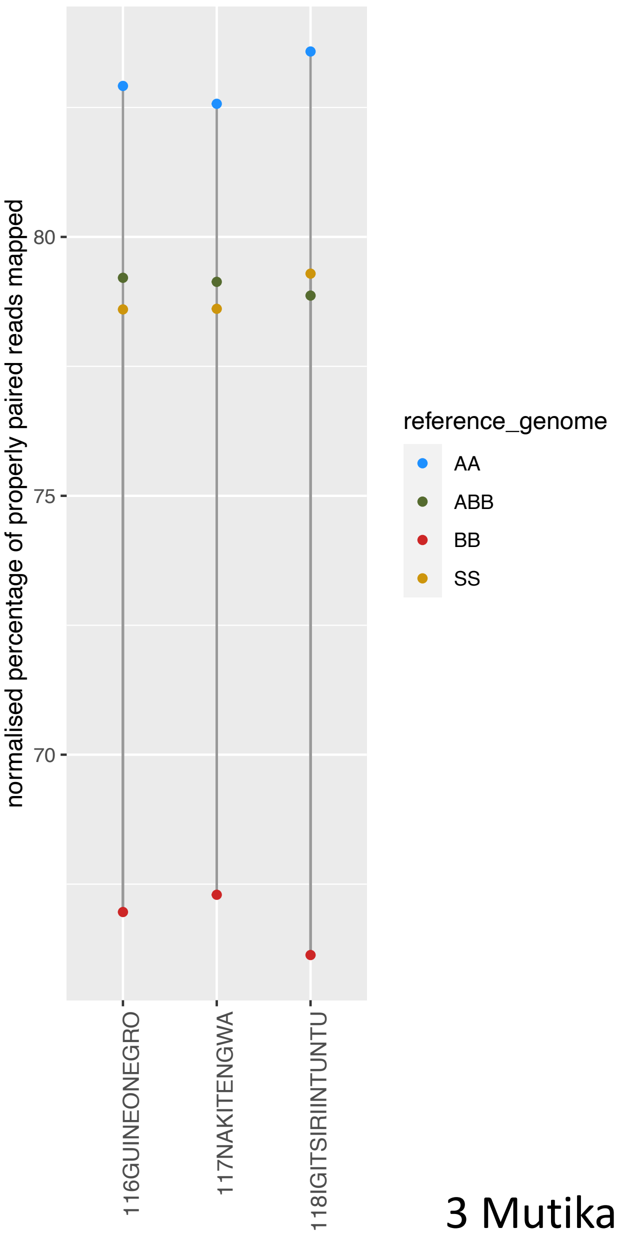


3 Mutika AAA

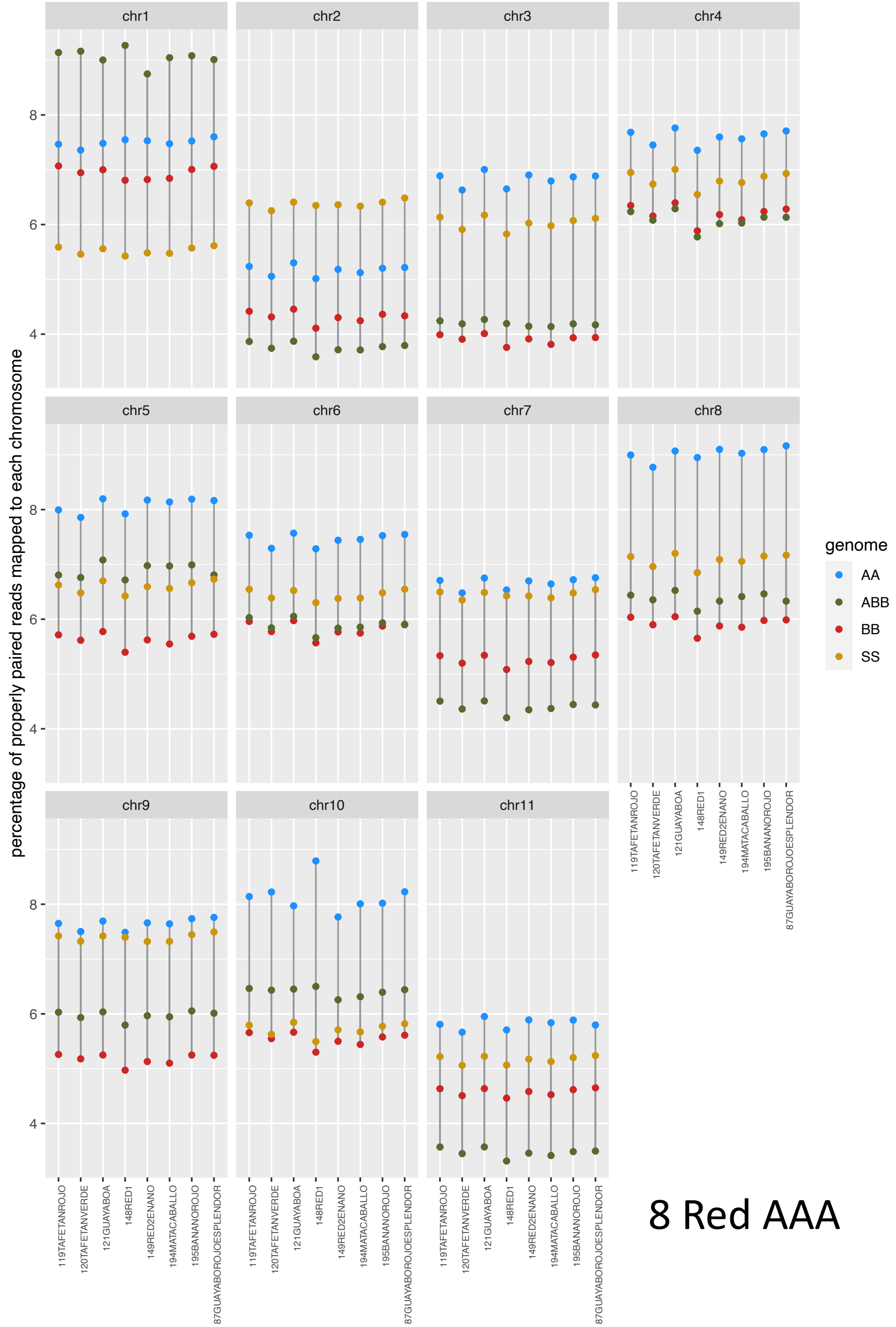


8 Red AAA

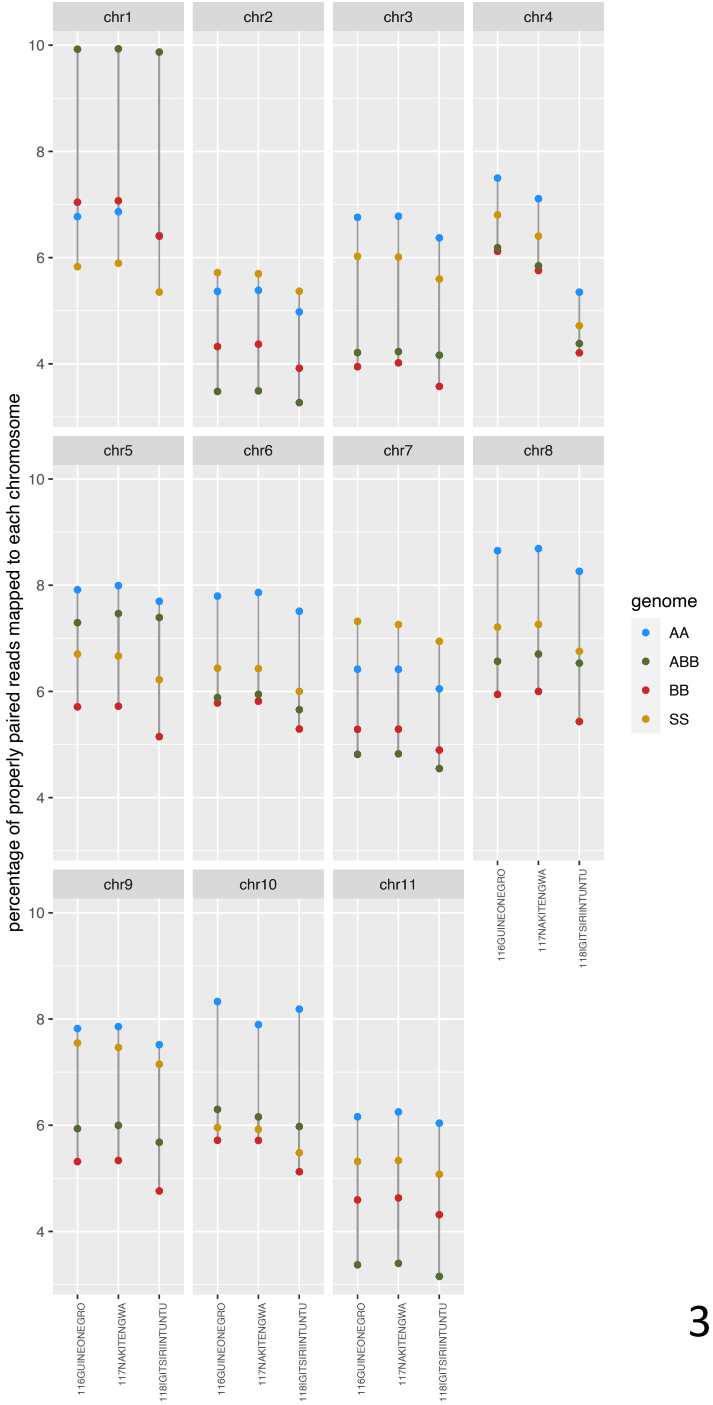


3 Mutika AAA

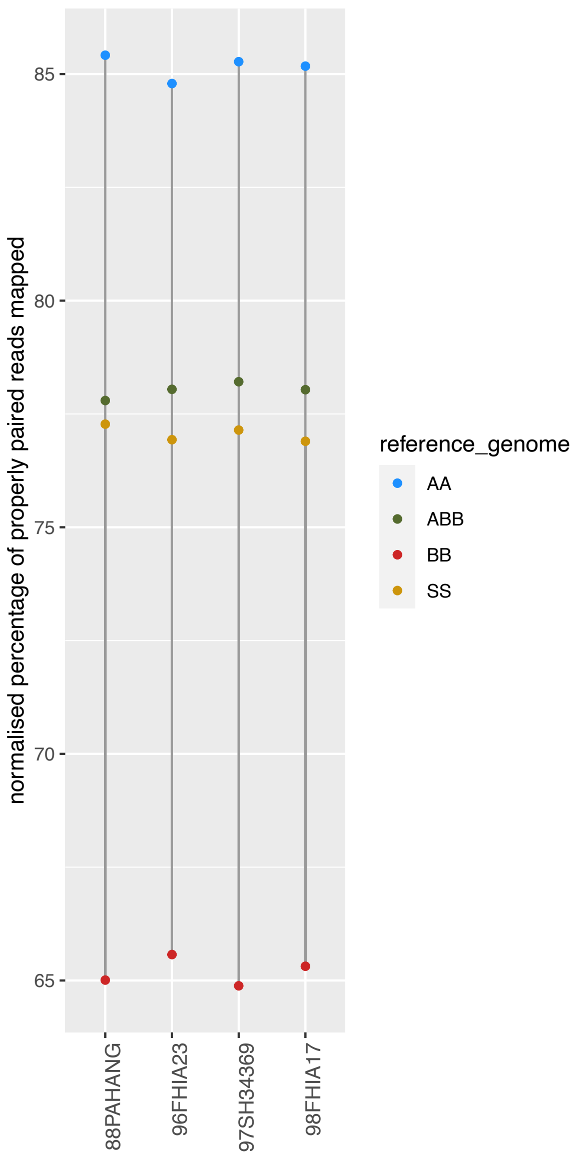


AAAA

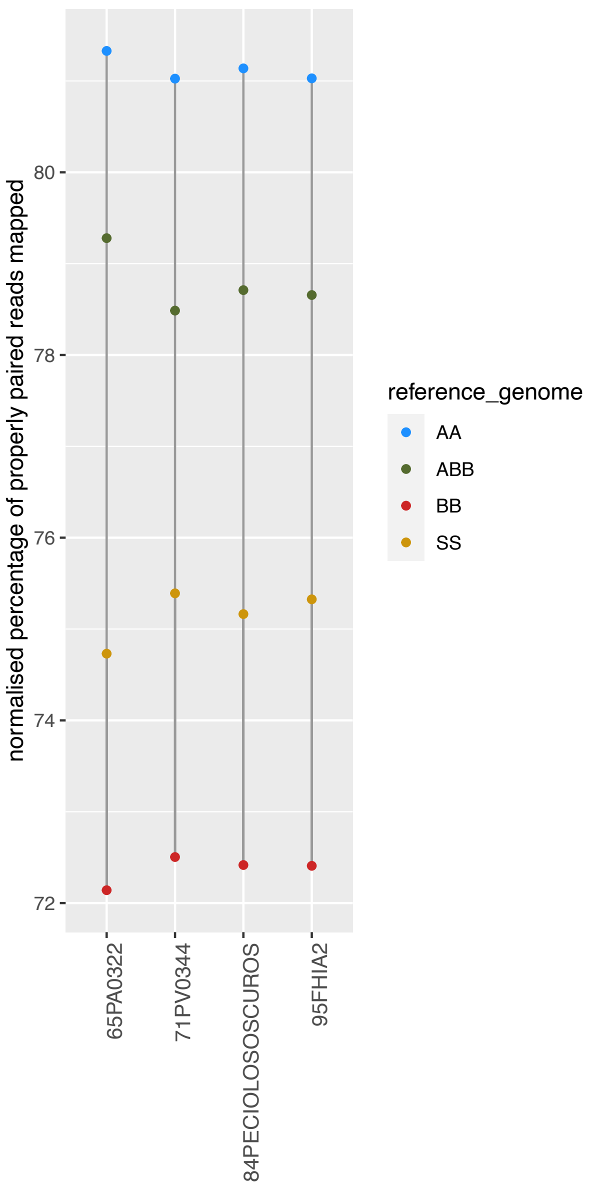


AAAB Pome

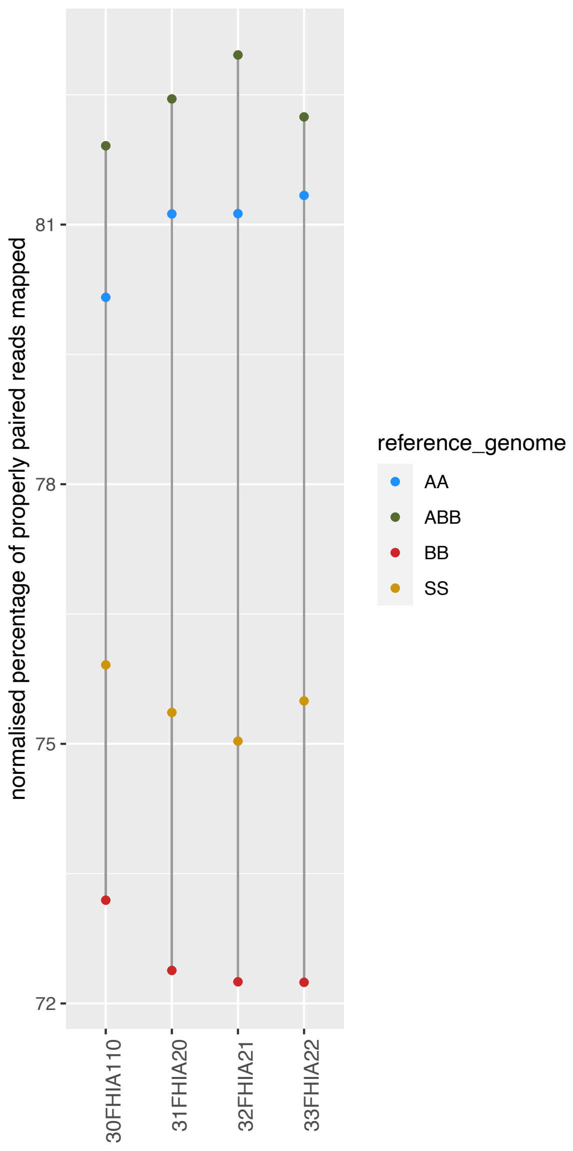


AAAB Africa

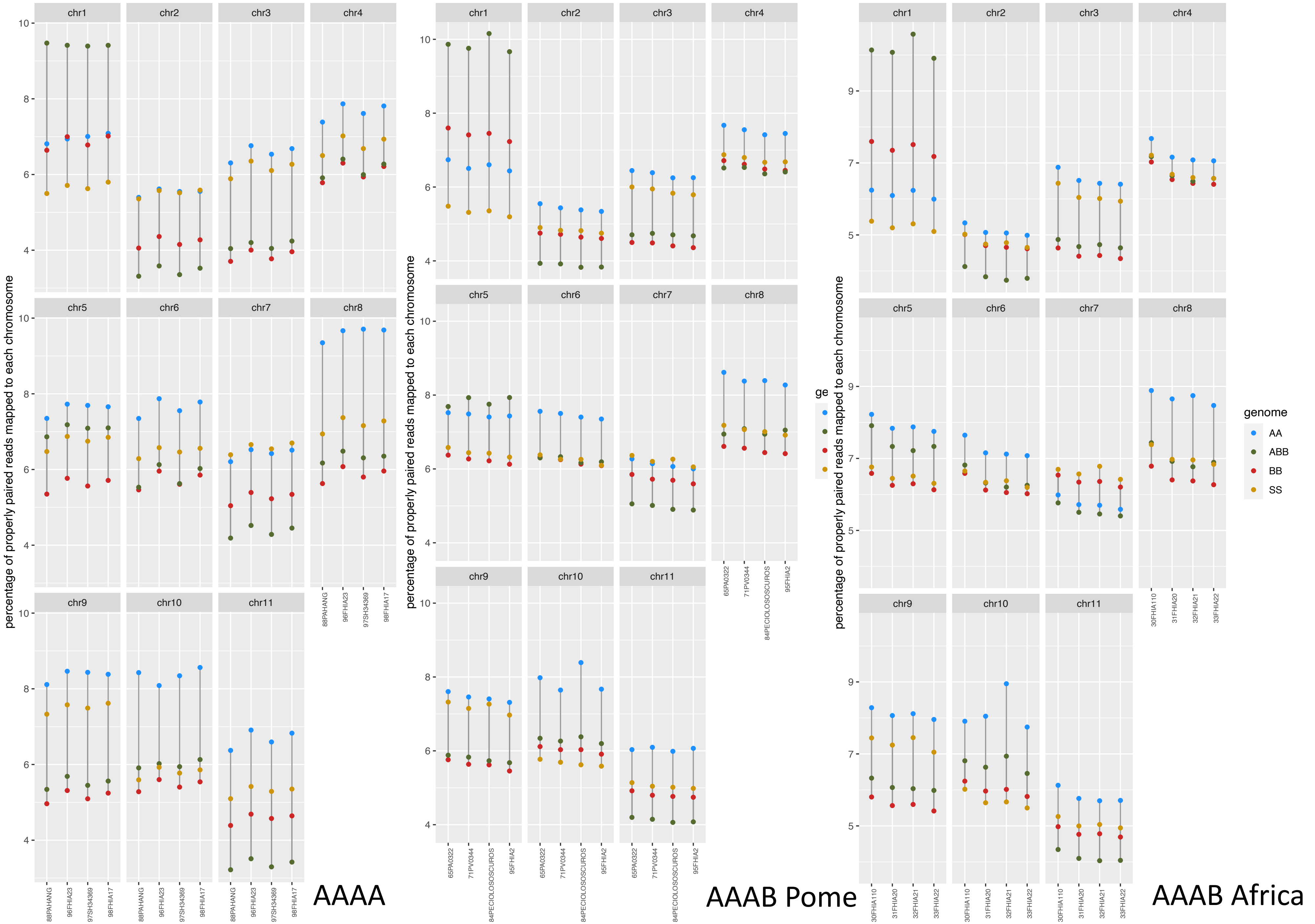
AAAA
AAAB Pome
AAAB Africa
percentage of properly paired reads mapped to each chromosome
chr1
chr2
chr3
chr4
chr5
chr6
chr7
chr8
chr9
chr10
chr11
88PAHANG
96FHIA23
97SH34369
98FHIA17
65PA0322
71PV0344
84PECIOLOSOSCUROS
95FHIA2
30FHIA110
31FHIA20
32FHIA21
33FHIA22
genome
AA
ABB
BB
SS

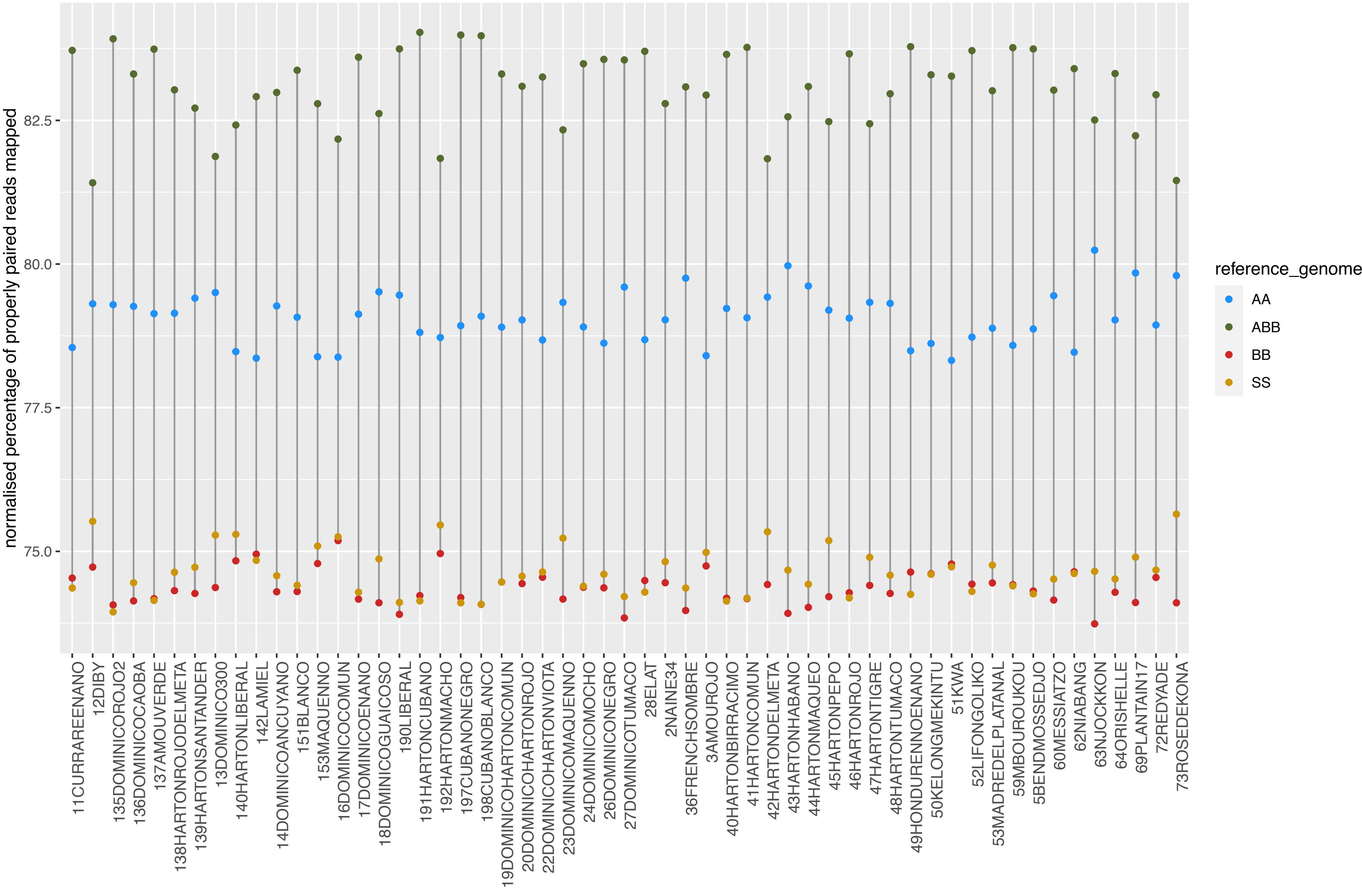


55 Plantain AAB

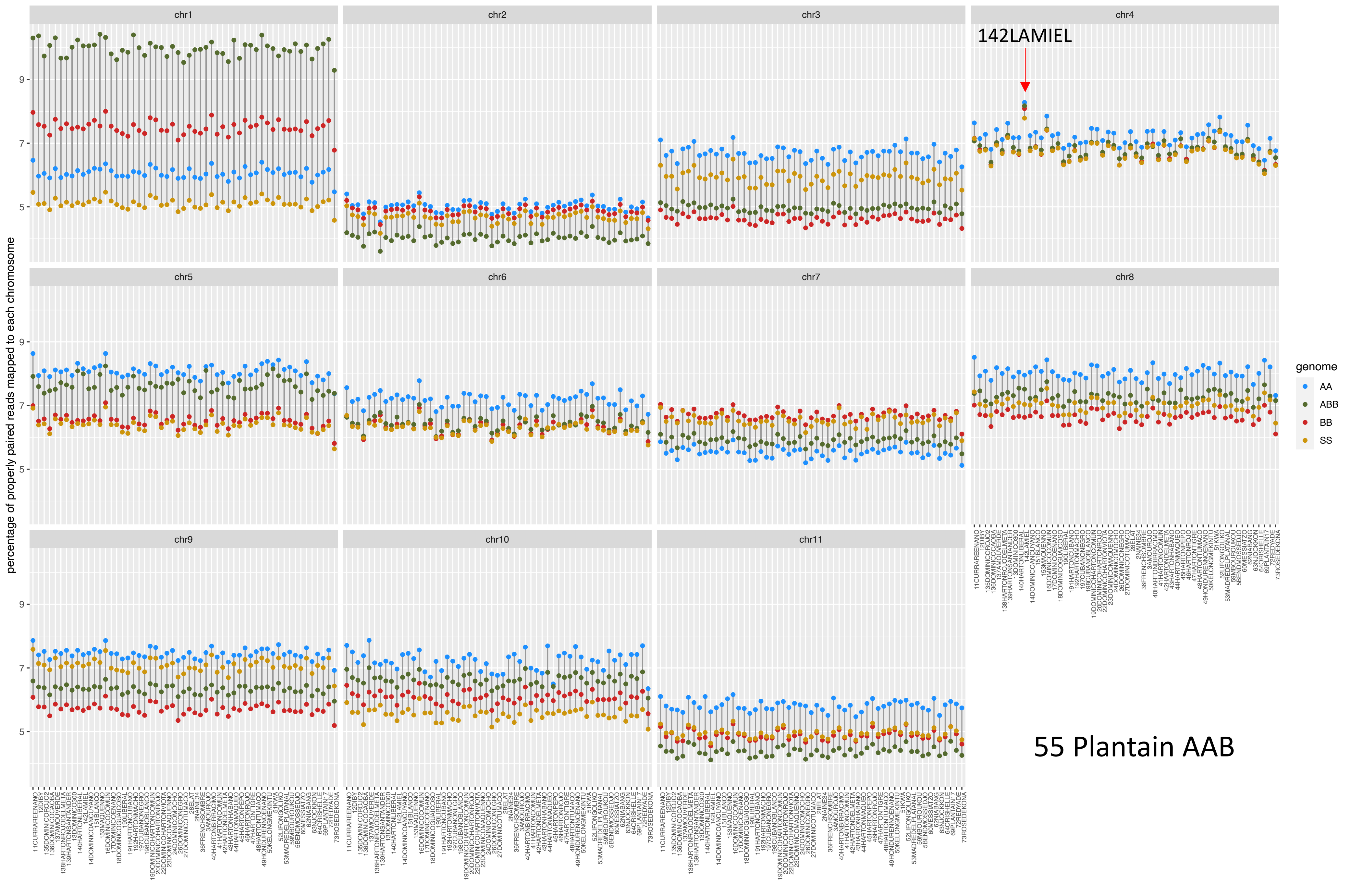
chr1
chr2
chr3
chr4
142LAMIEL
chr5
chr6
chr7
chr8
chr9
chr10
chr11
9
7
5
percentage of properly paired reads mapped to each chromosome
genome
AA
ABB
BB
SS
55 Plantain AAB

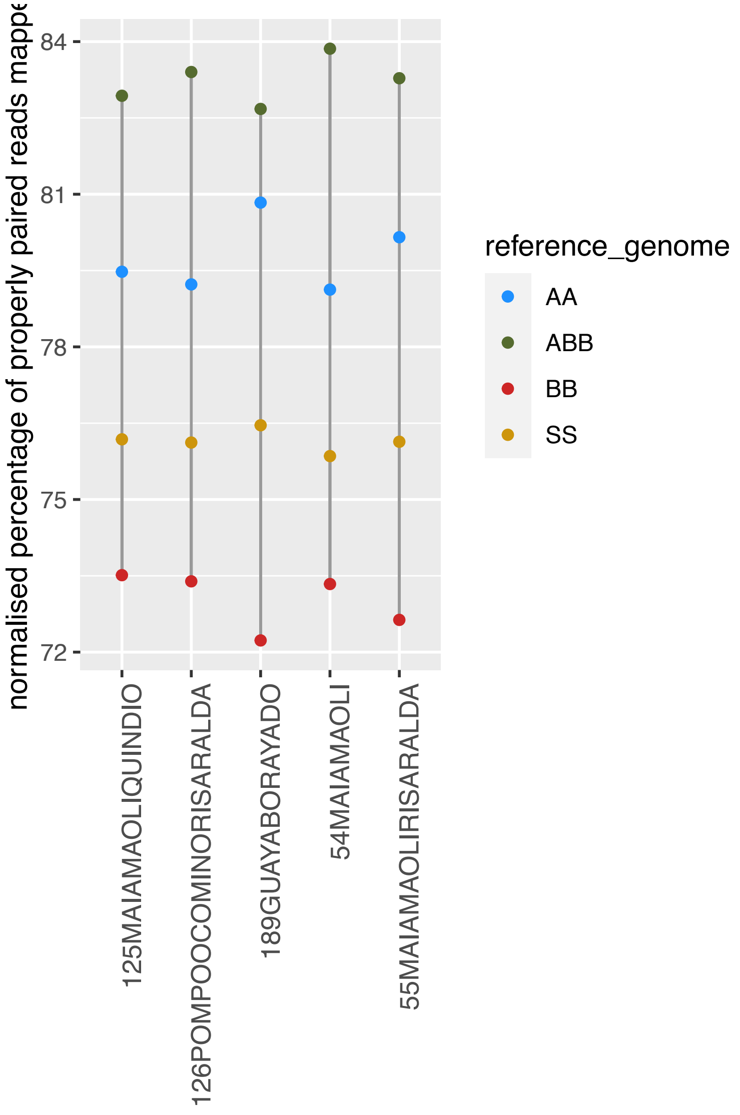


5 Popoulu ABB

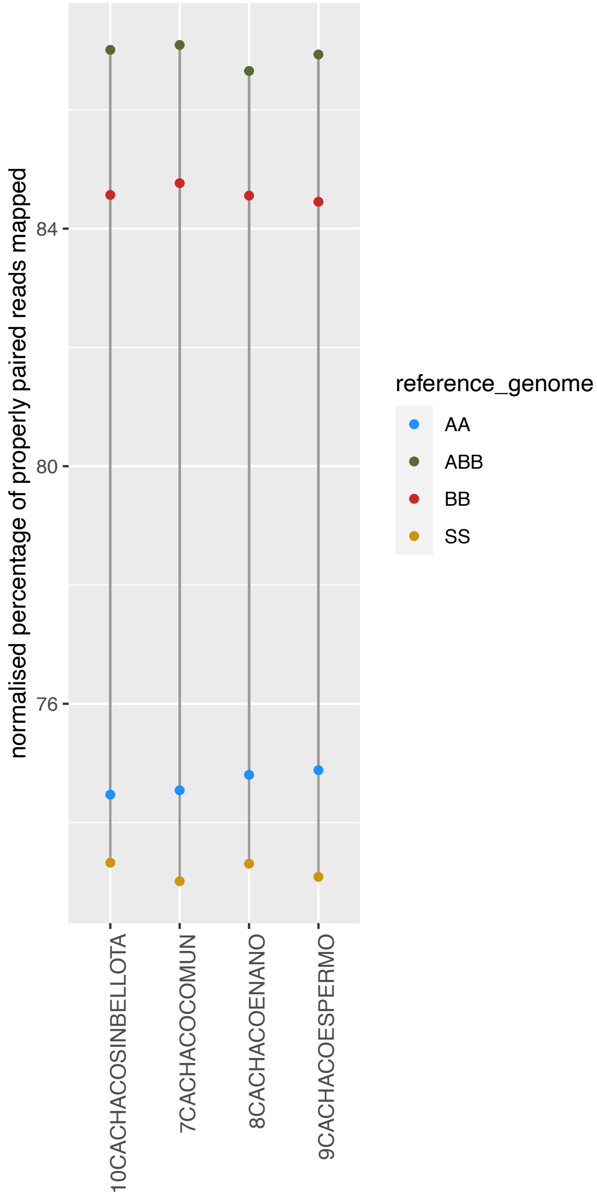


4 Bluggoe ABB

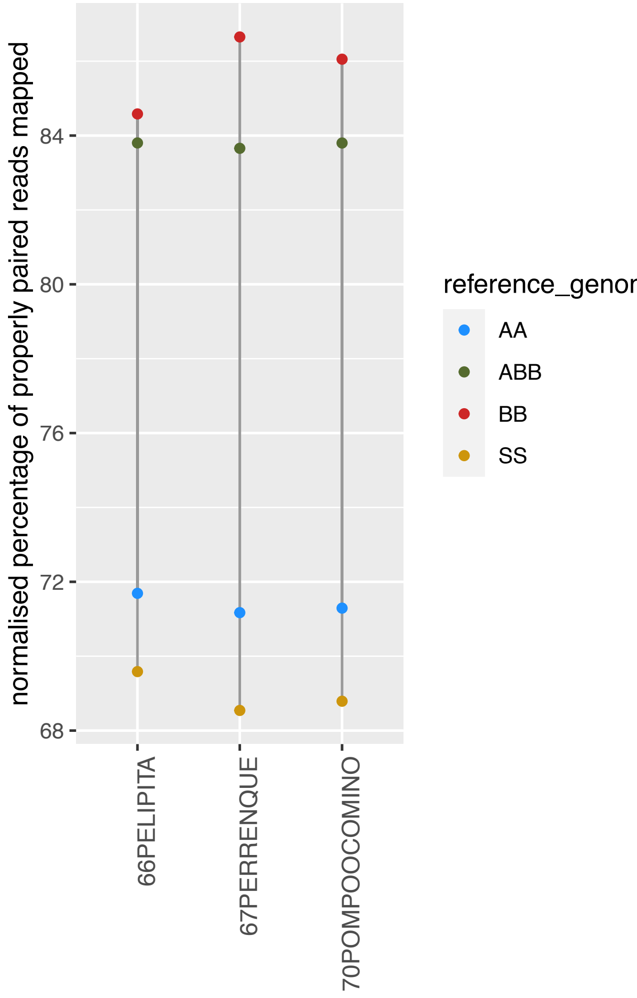


3 Pelipita ABB

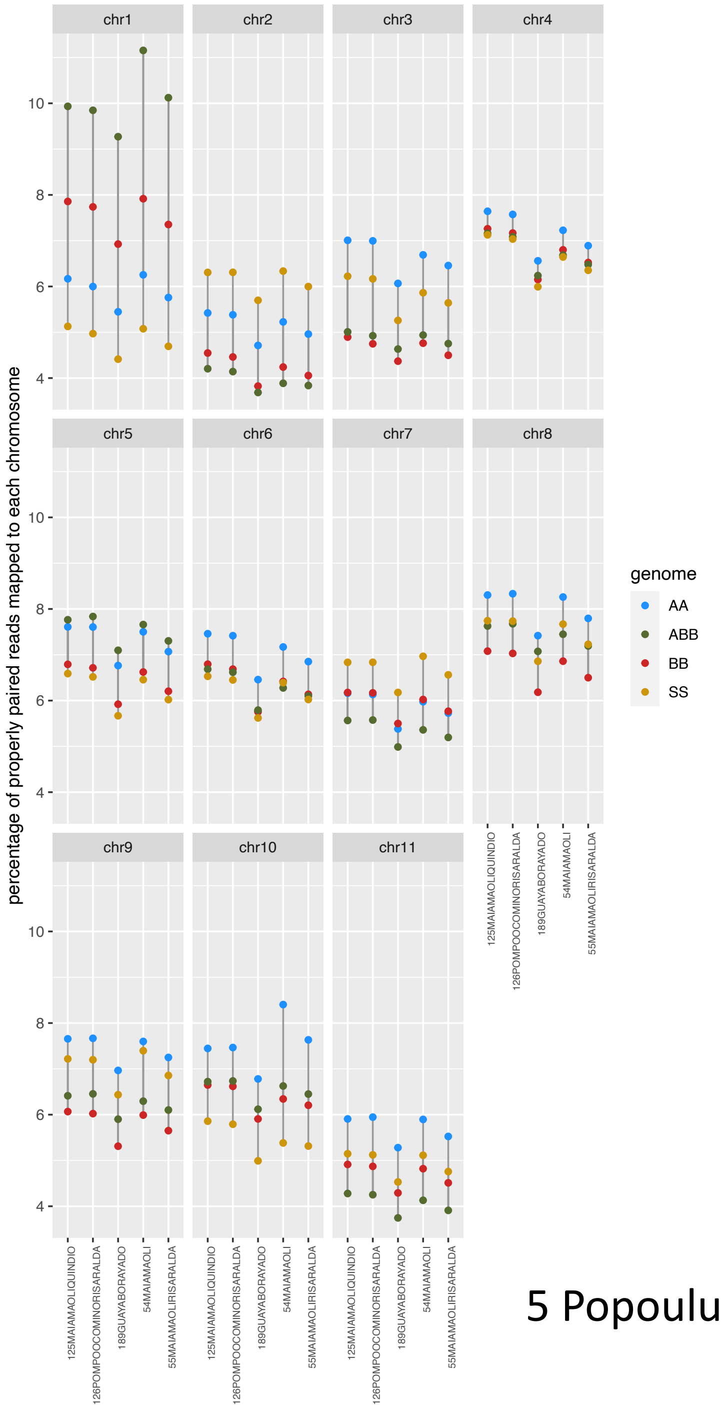


5 Popoulu ABB

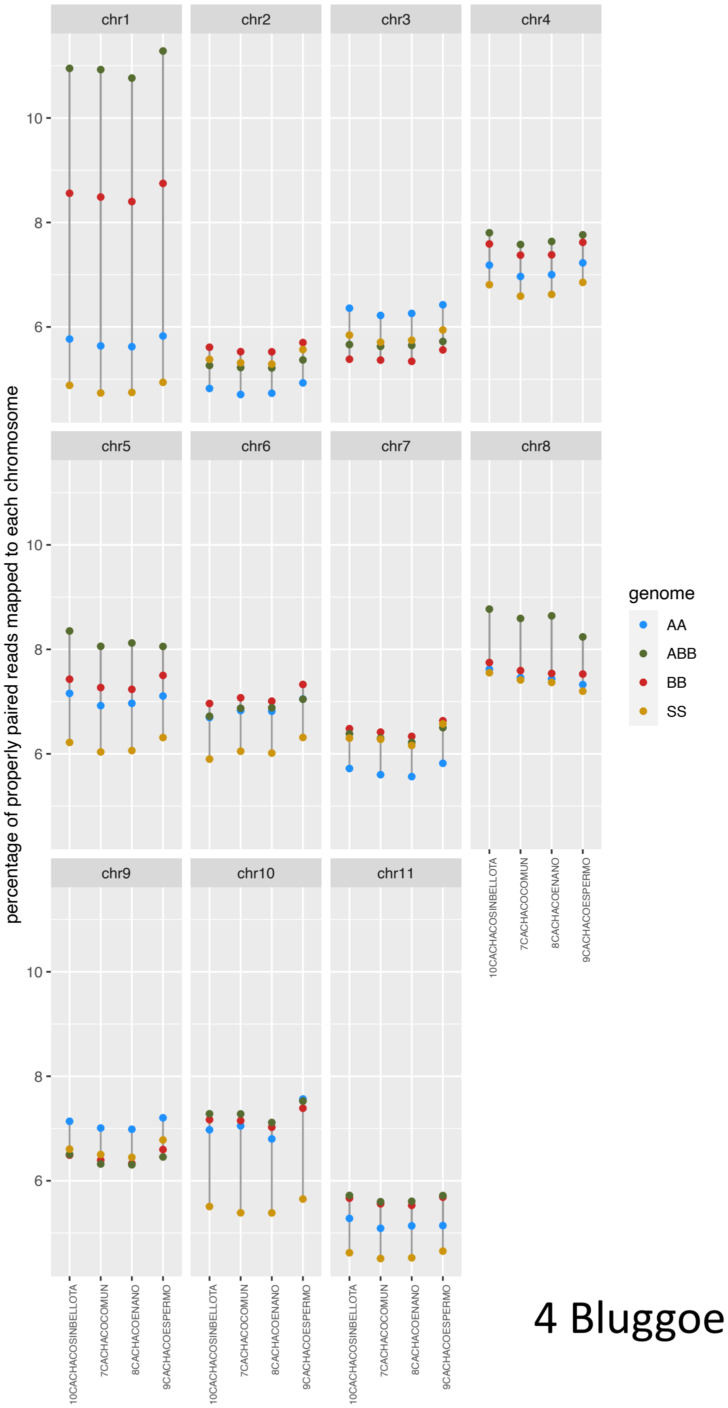


4 Bluggoe ABB

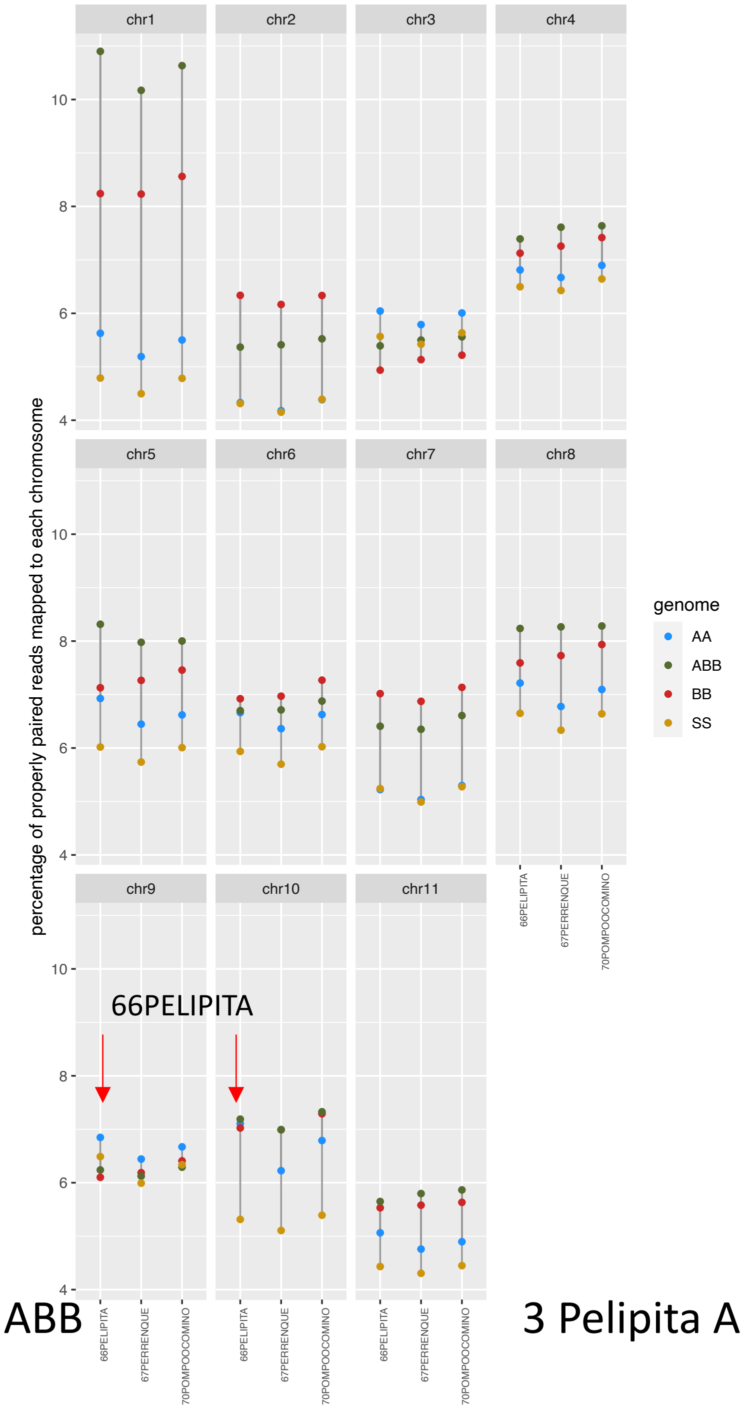


3 Pelipita ABB
