## Supplementary figures and images for "Characterising genome composition and large structural variation in banana varietal groups"

### 76BOCADILLOALTOcov_byA.pdf

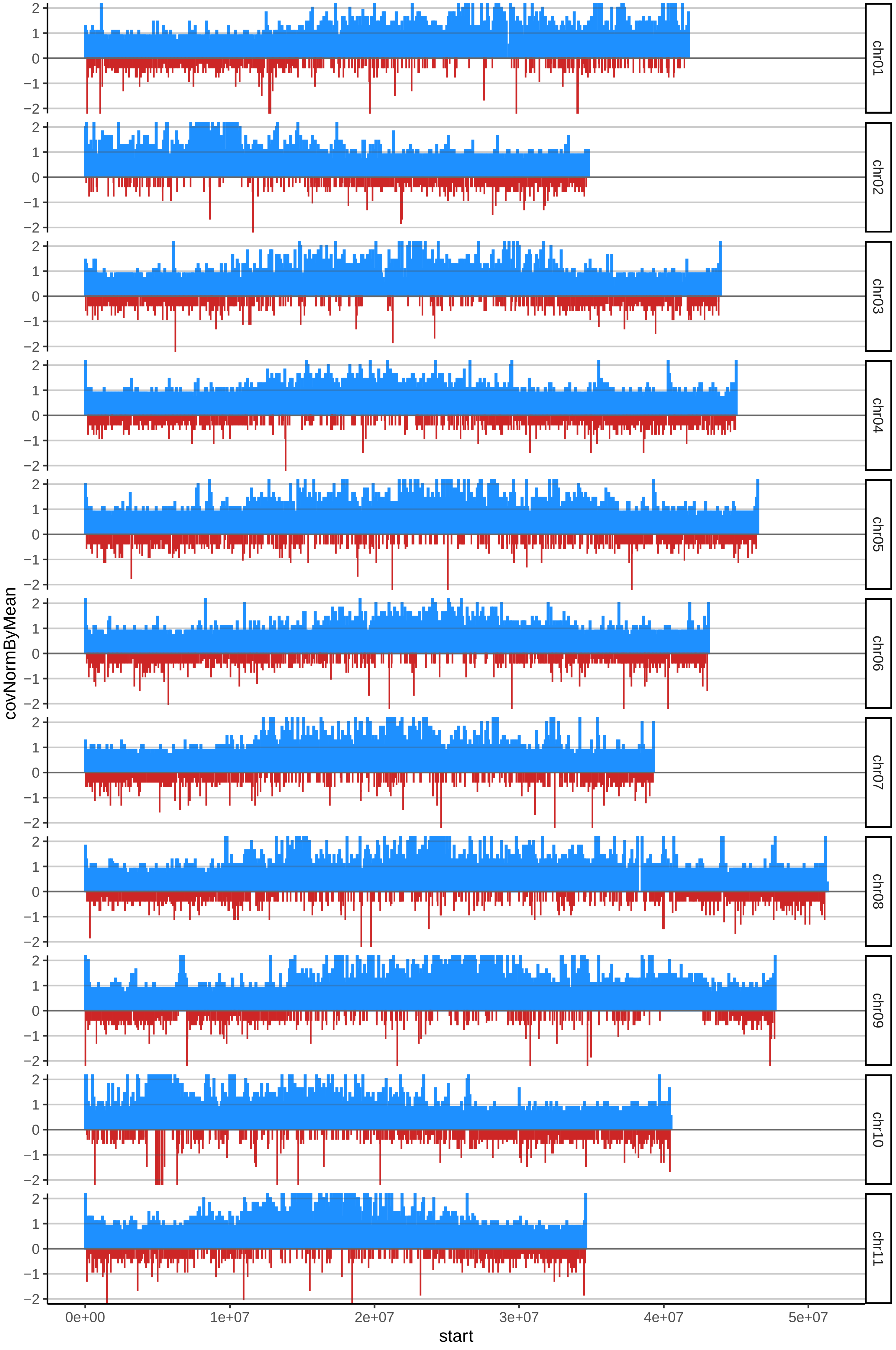

### 143BOCADILLODELCHOCOcov_byB.pdf

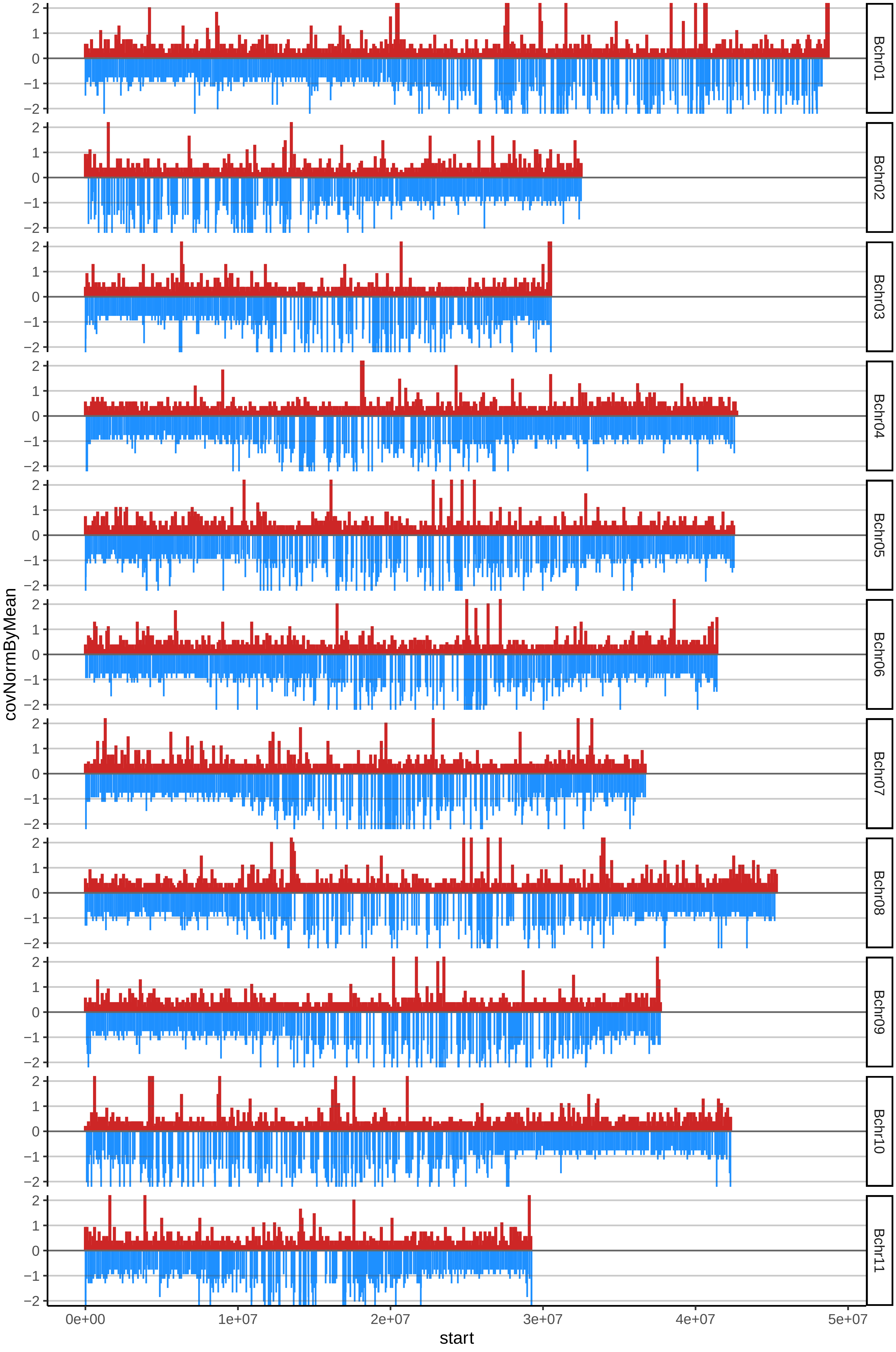

### 162NARI02cov_byA.pdf

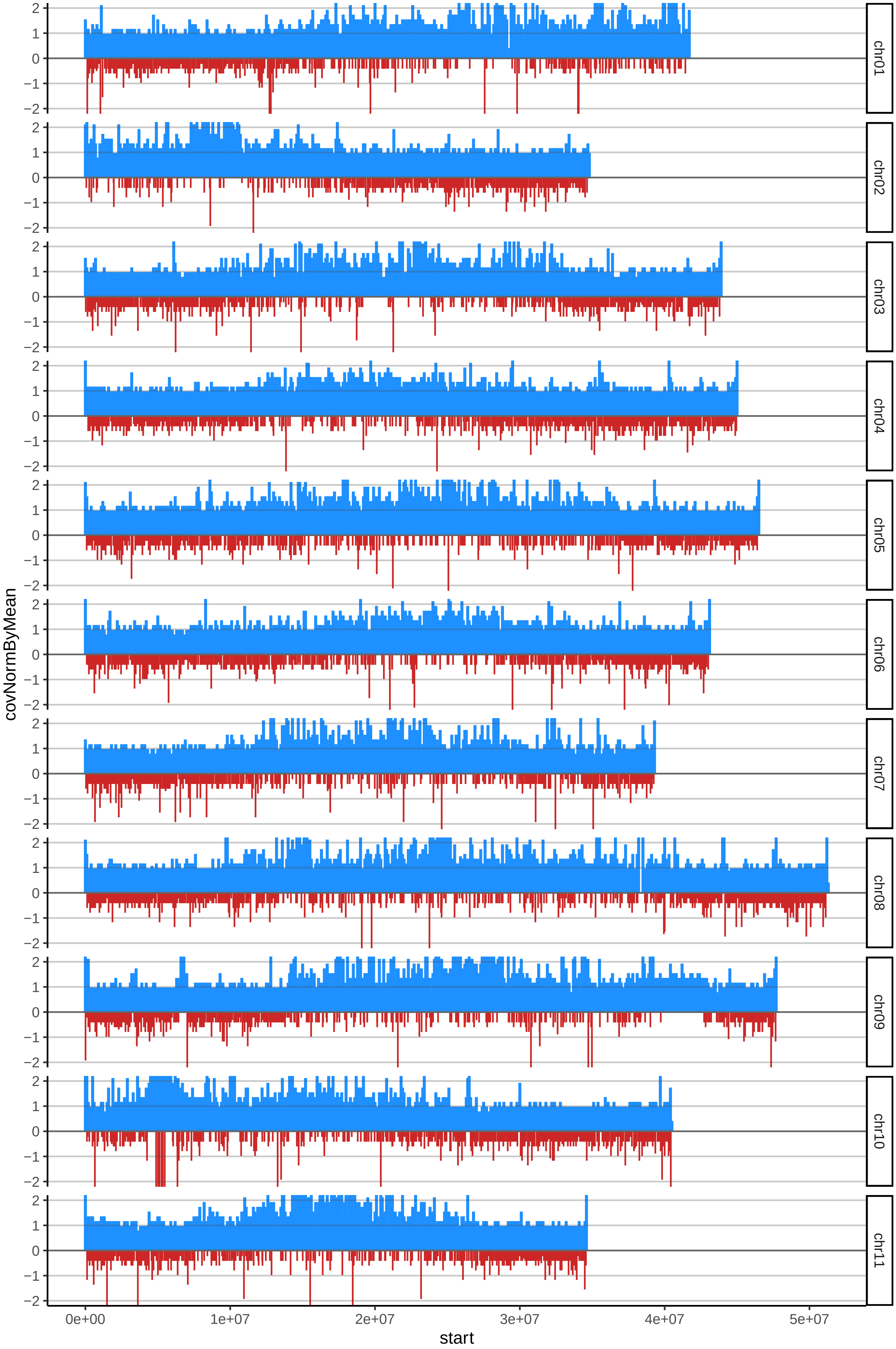

### 168SABO03cov_byA.pdf

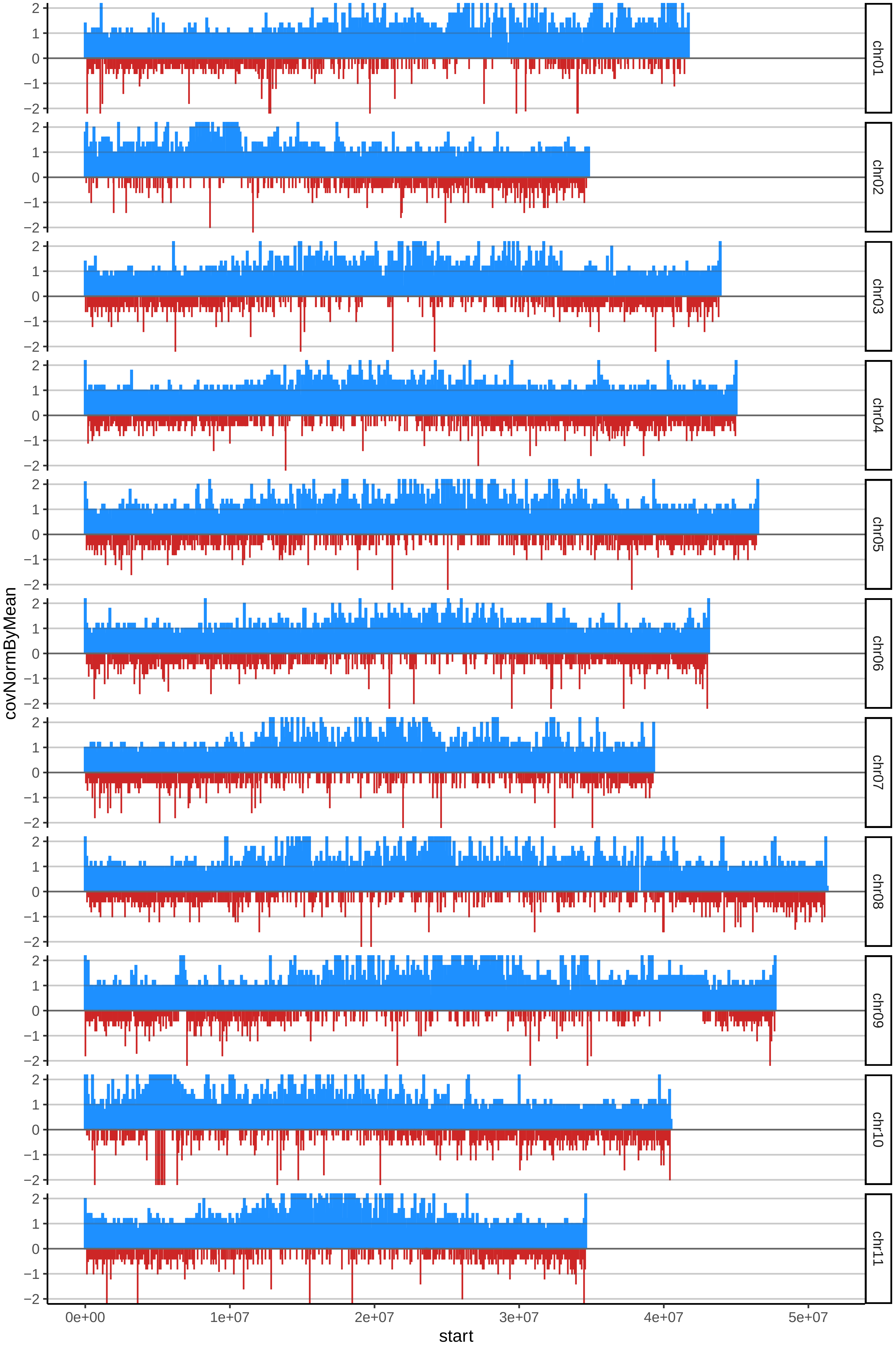

### 171SABO02cov_byA.pdf

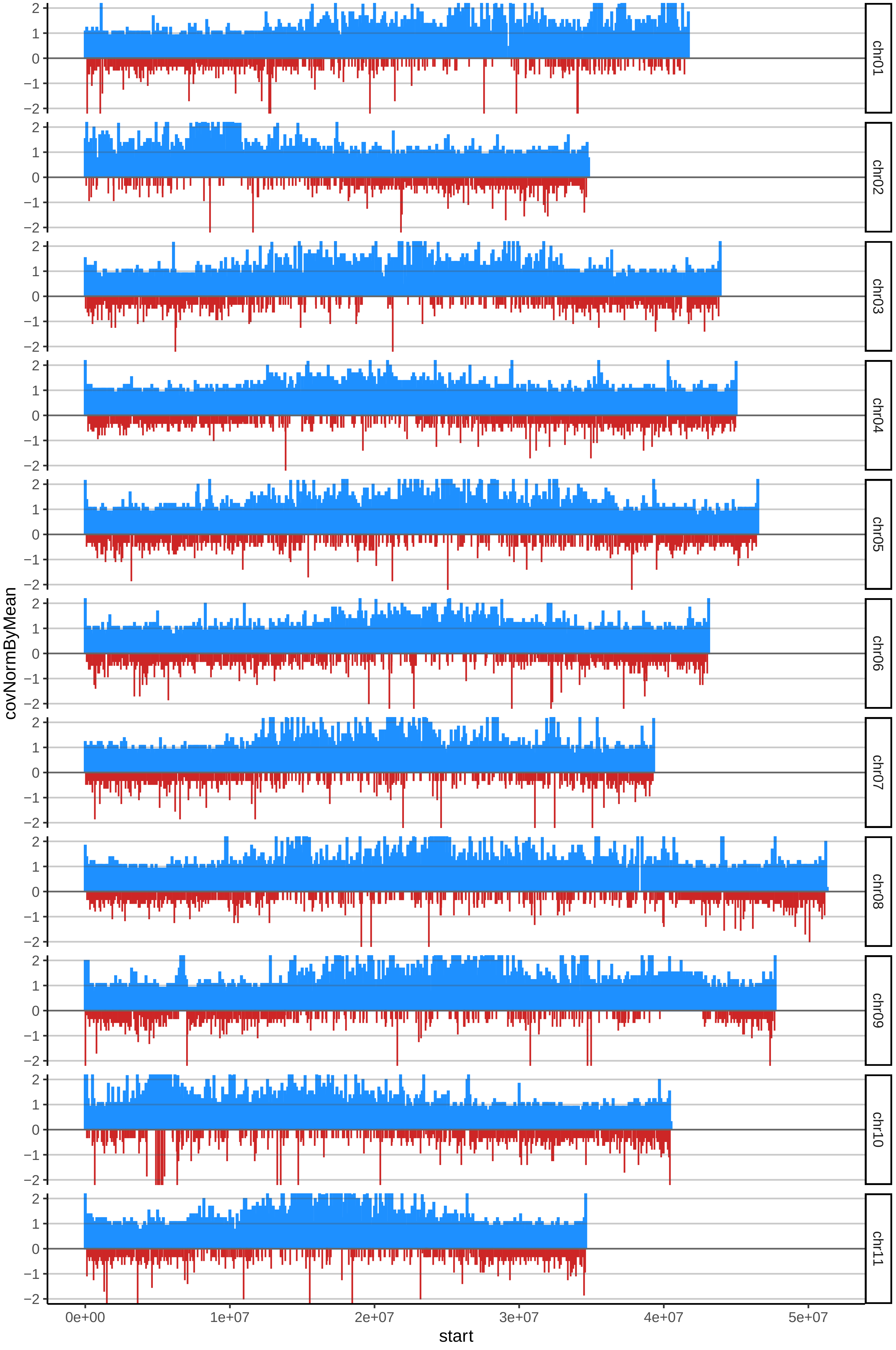

### 177SAVE07cov_byB.pdf

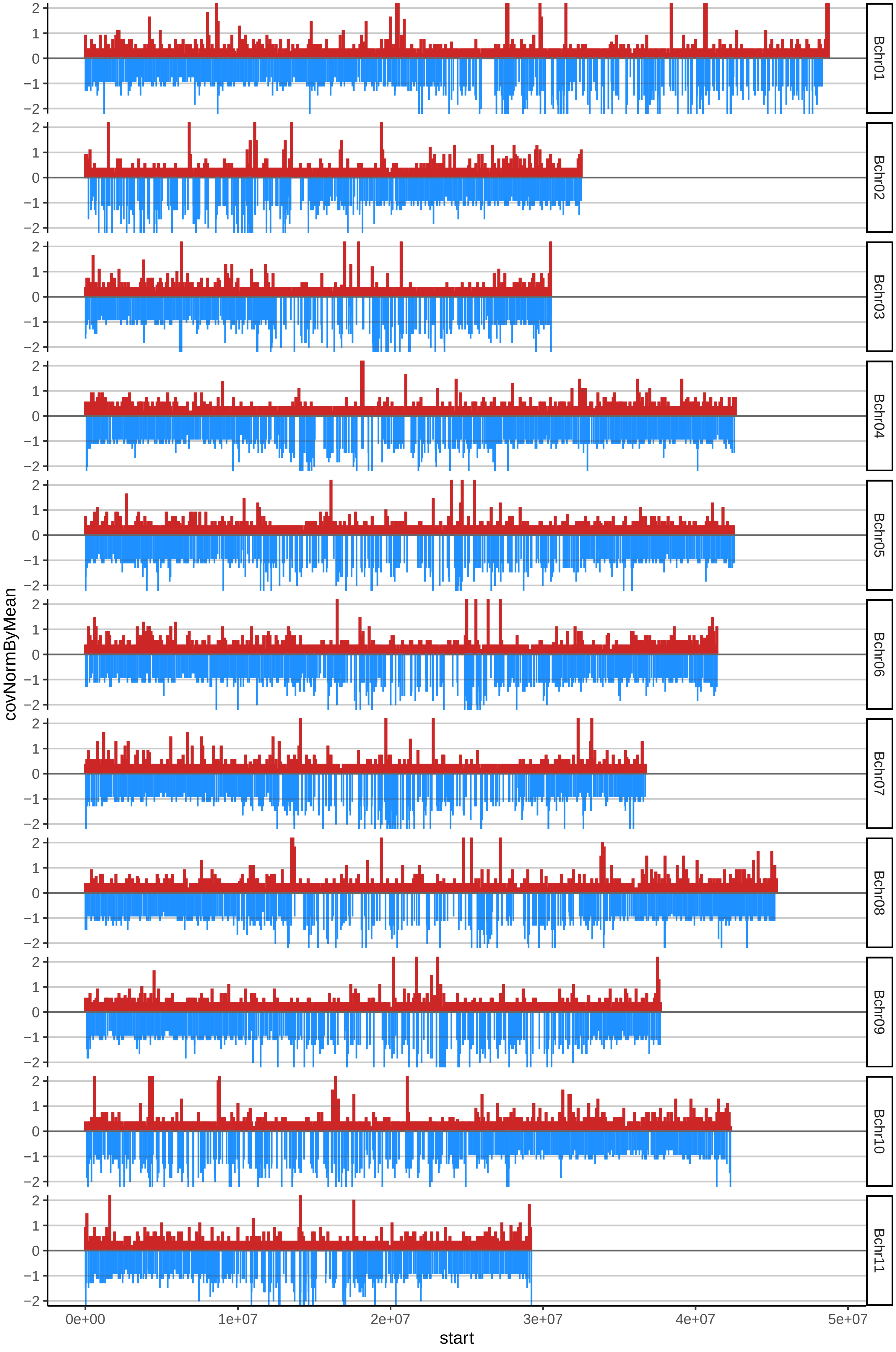

### 179VAPA03cov_byA.pdf

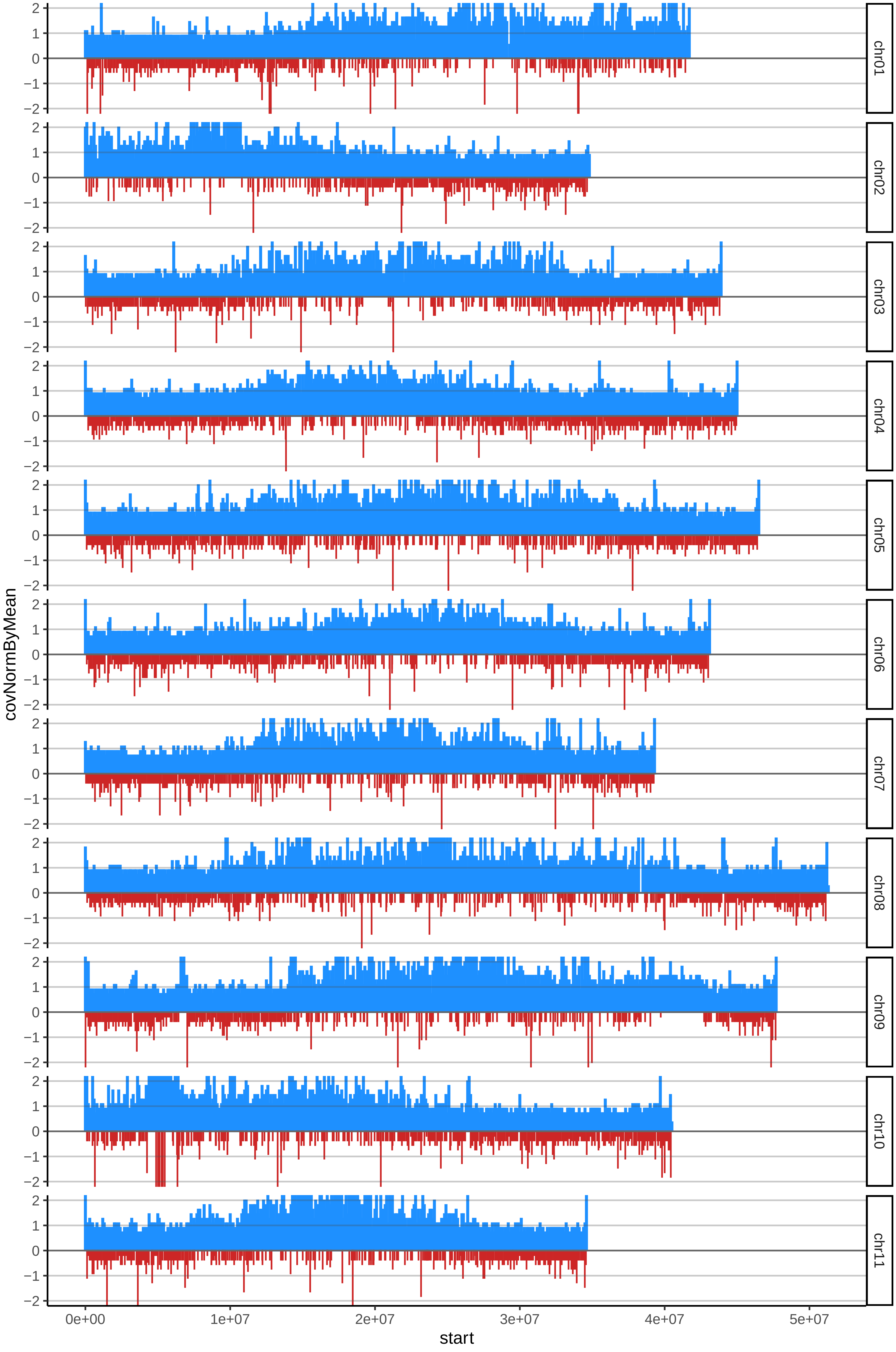

### 181VAPA01cov_byA.pdf

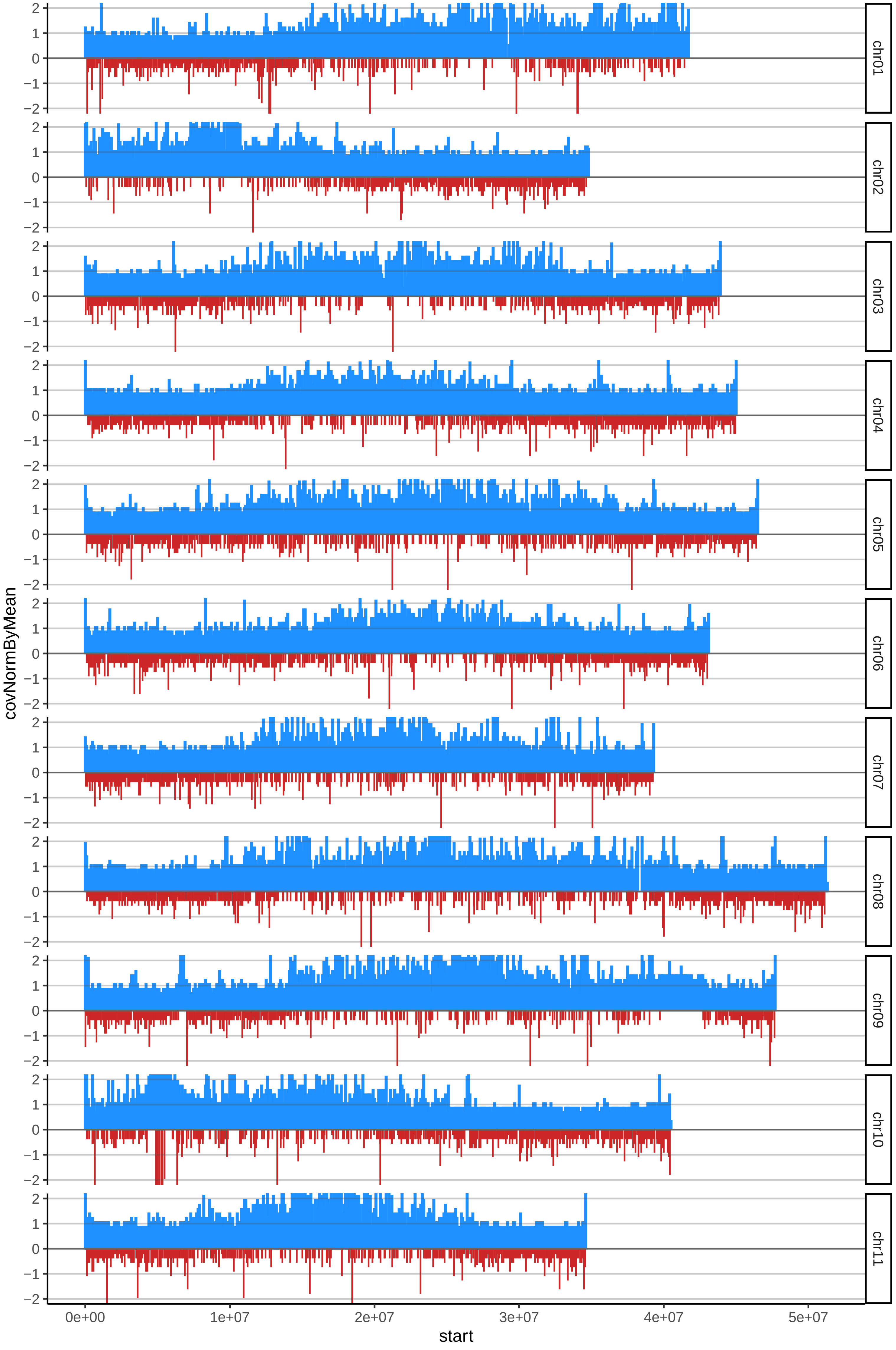

### 184VABU01cov_byB.pdf

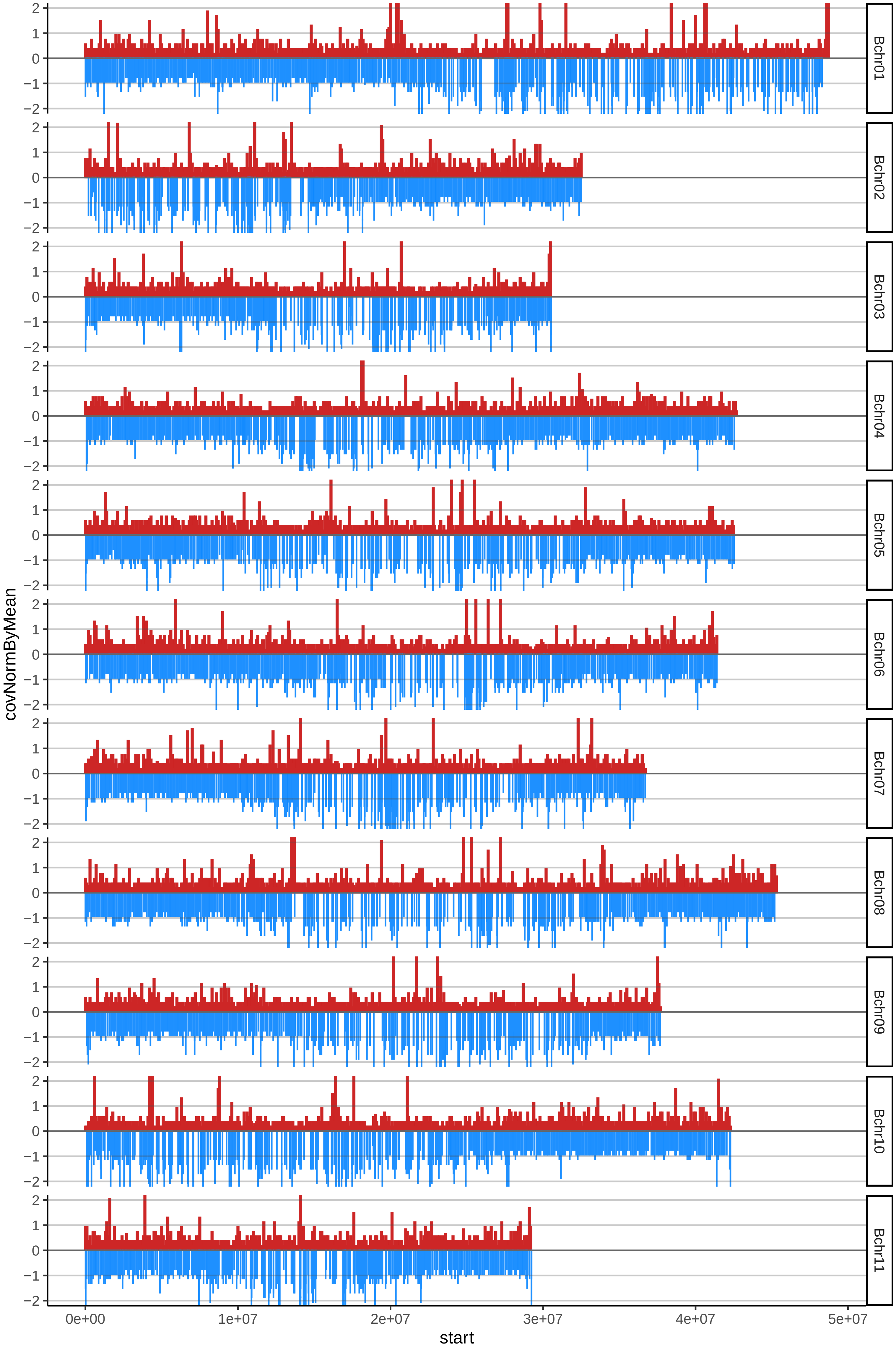

### 187ICONONZO03cov_byA.pdf

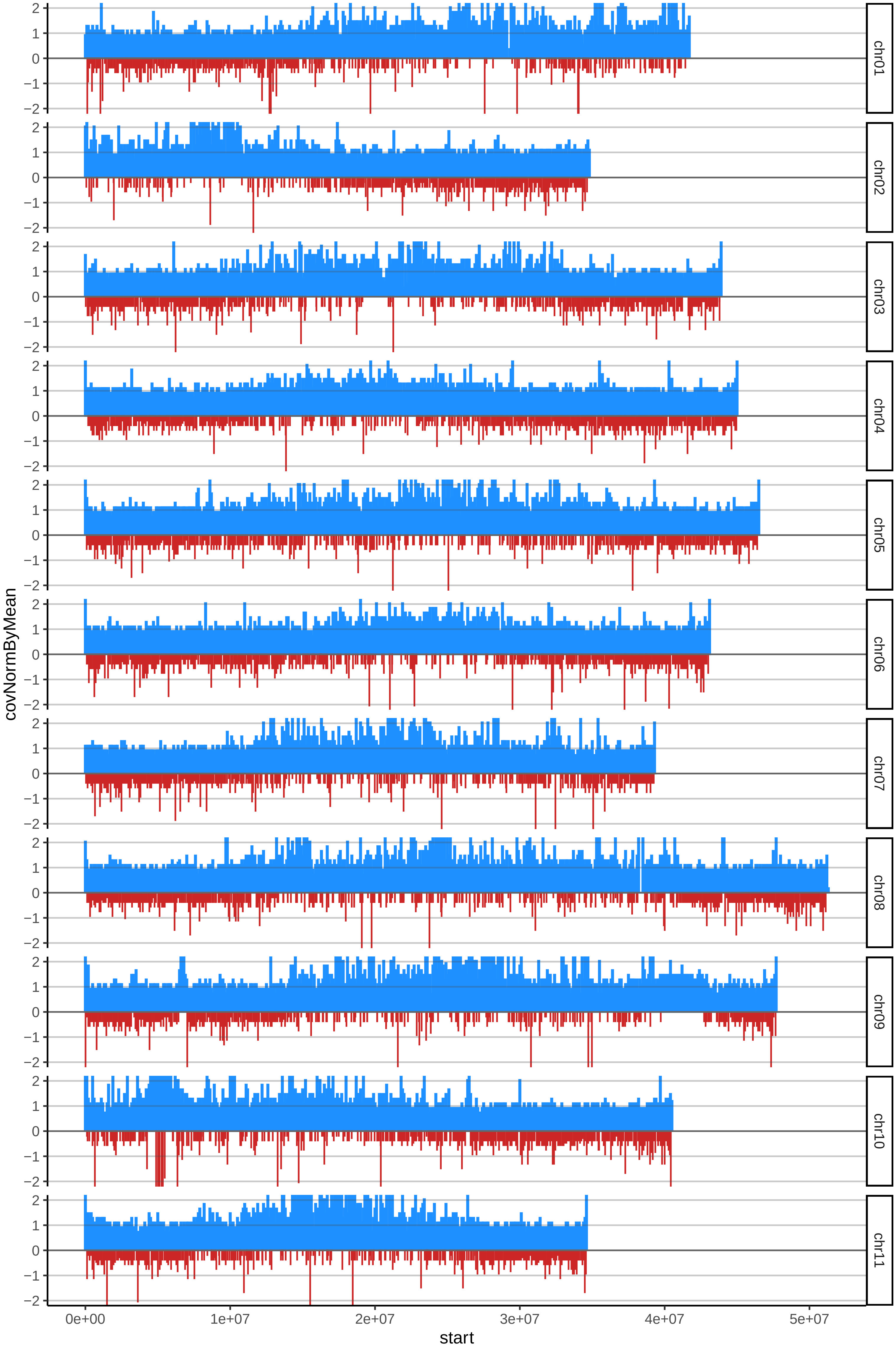
