## Supplementary figures and images for "Characterising genome composition and large structural variation in banana varietal groups"

### 1SABAcov_byA.pdf

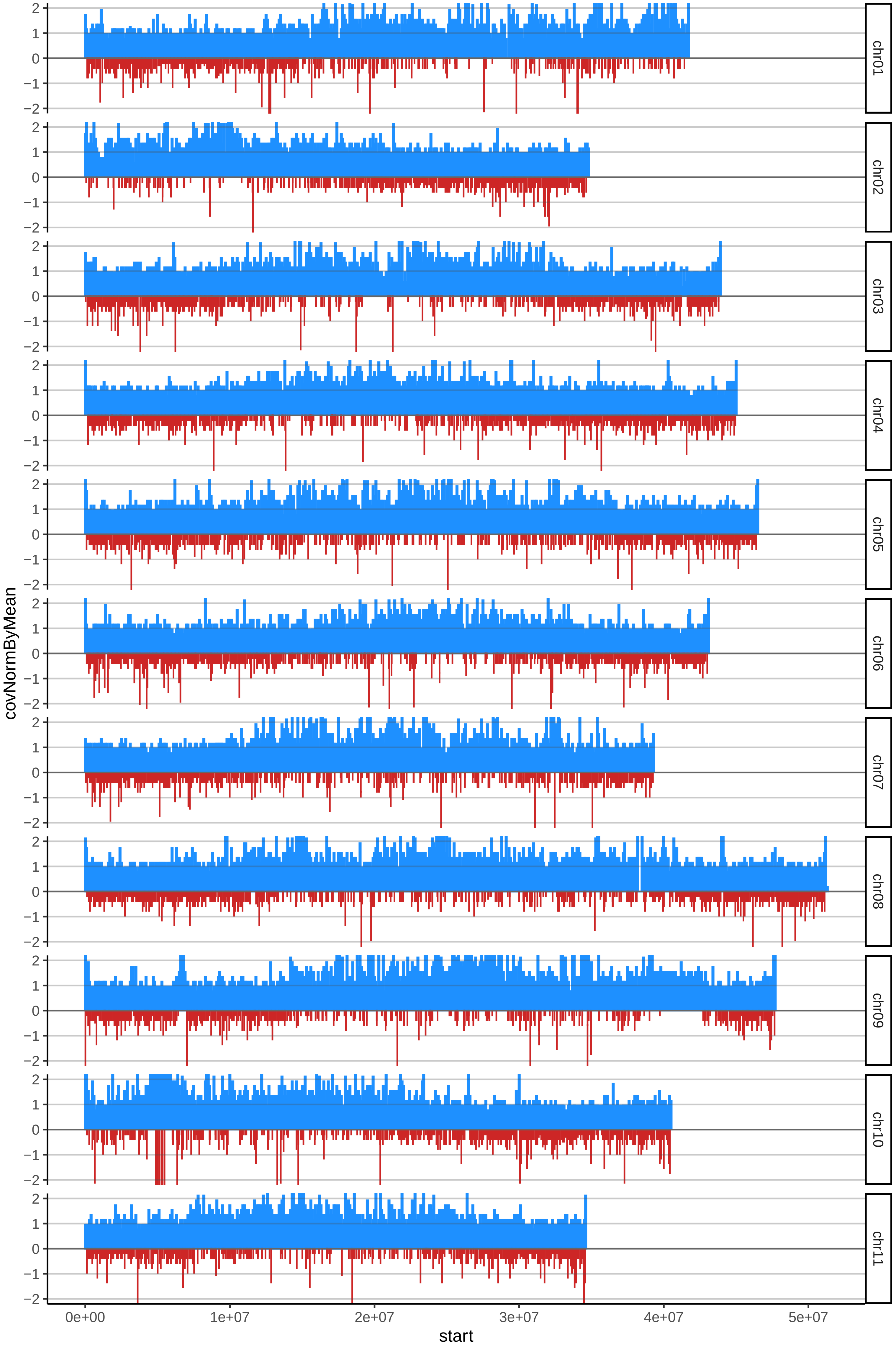

### 1SABAcov_byB.pdf

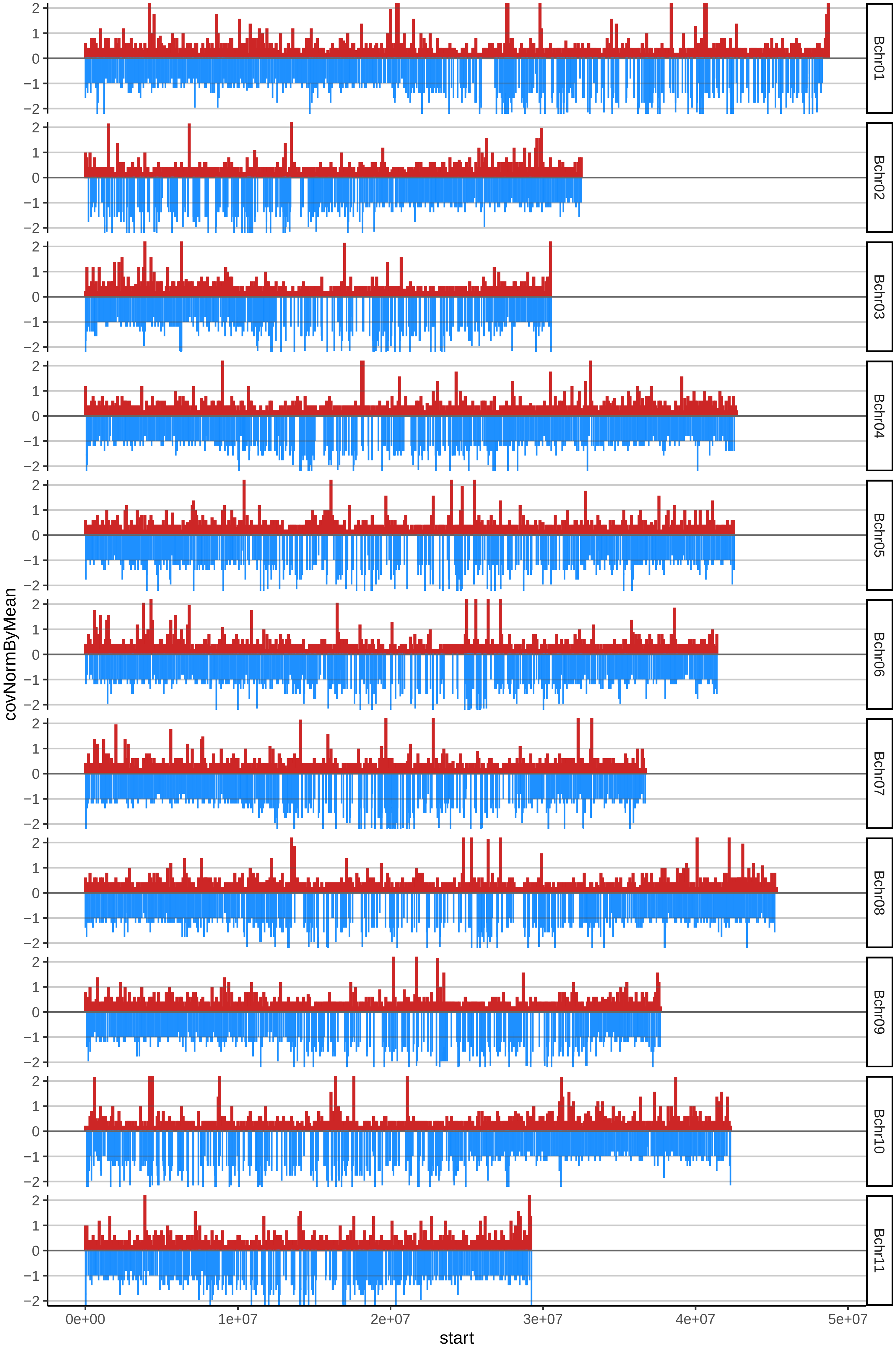

### 5BENDMOSSEDJOcov_byB.pdf

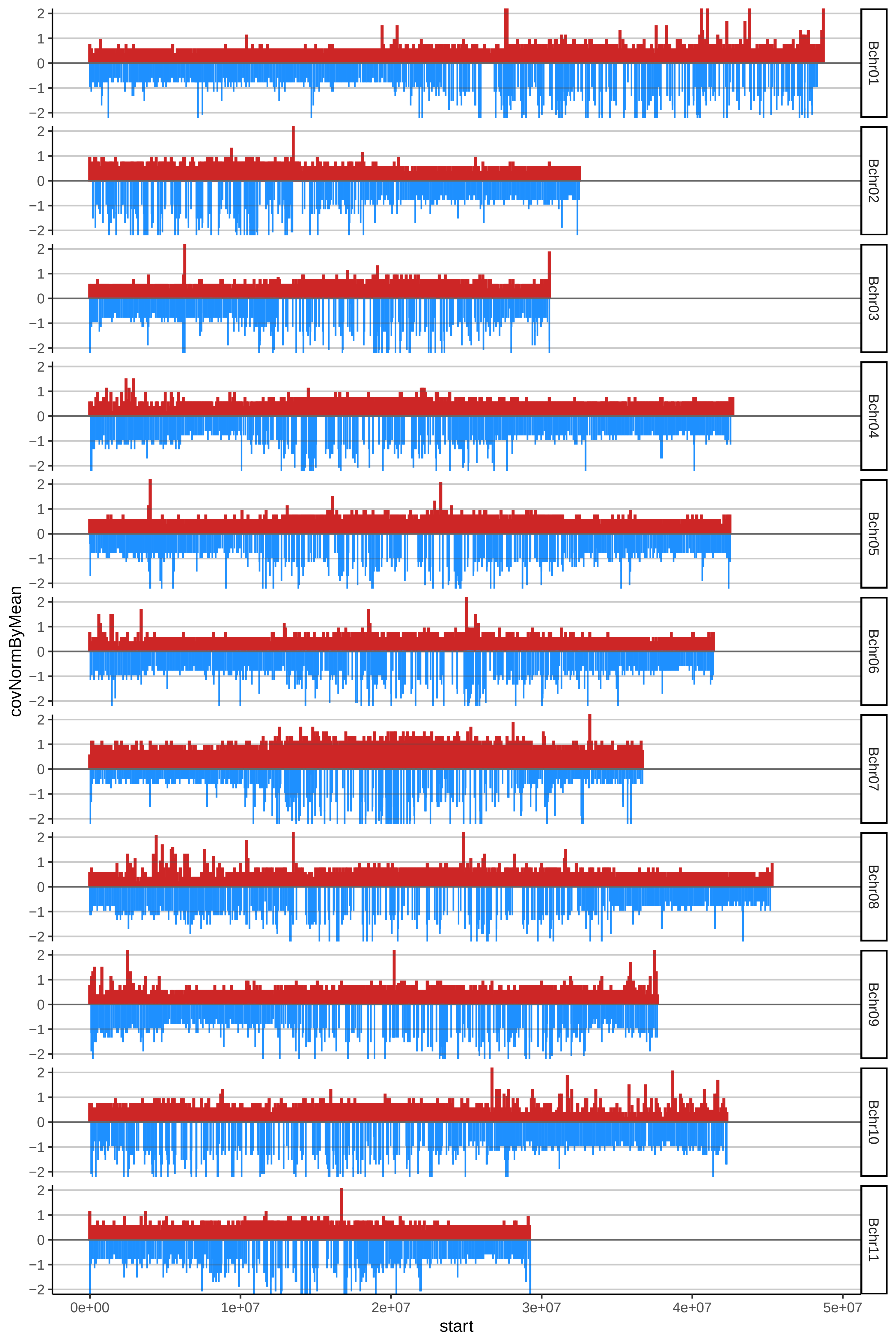

### 7CACHACOCOMUNcov_byA.pdf

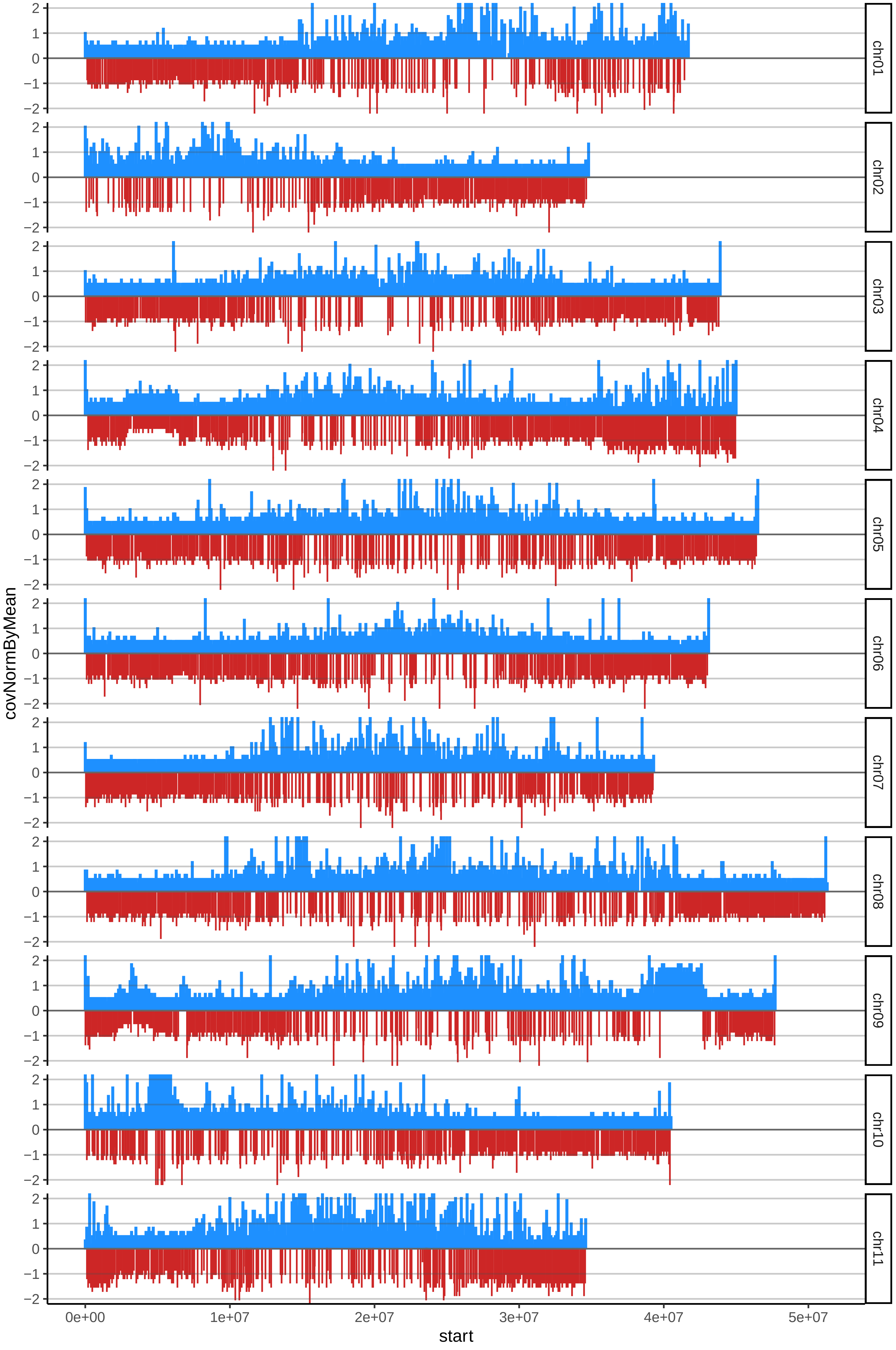

### 7CACHACOCOMUNcov_byB.pdf

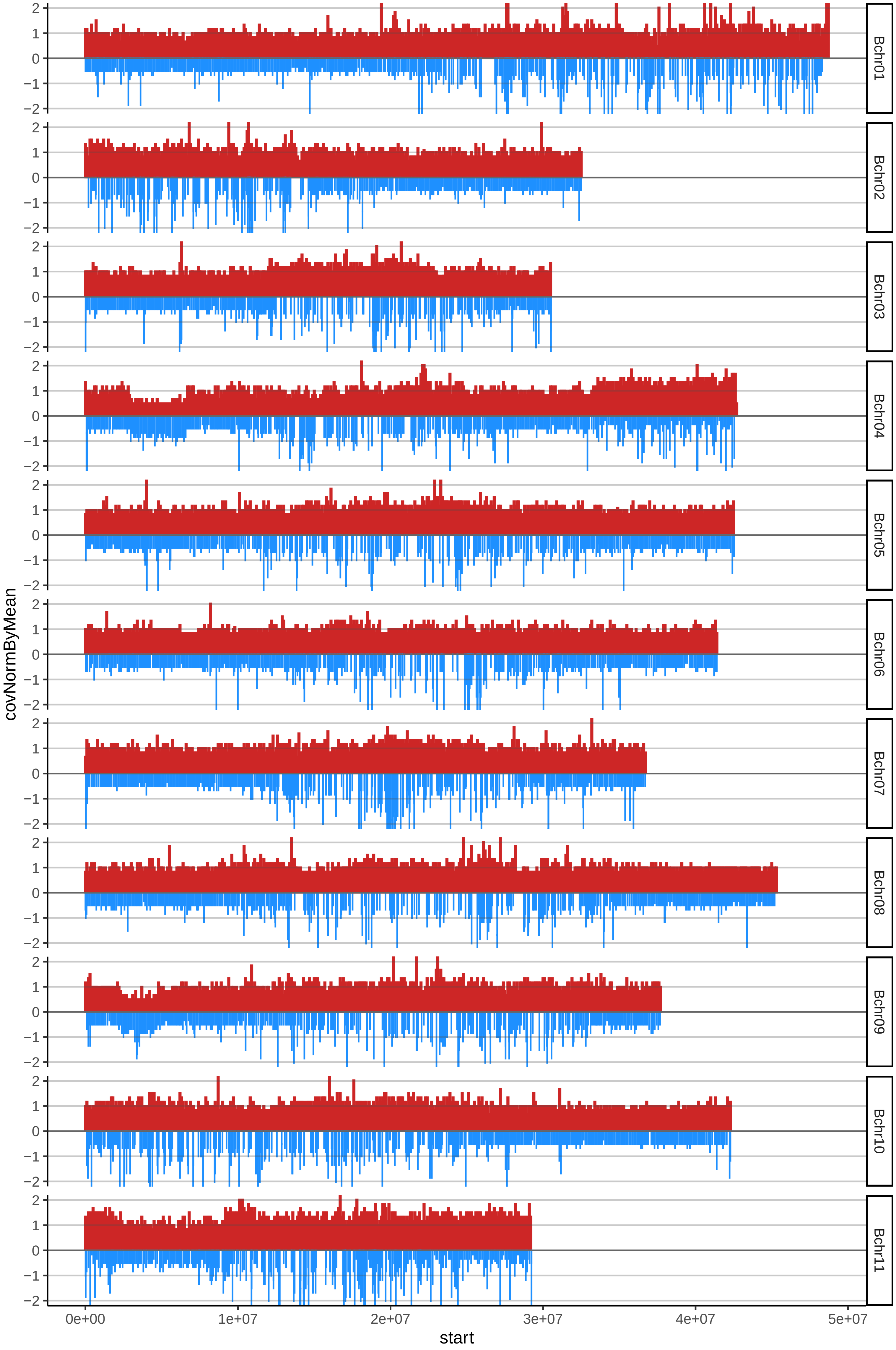

### 8CACHACOENANOcov_byA.pdf

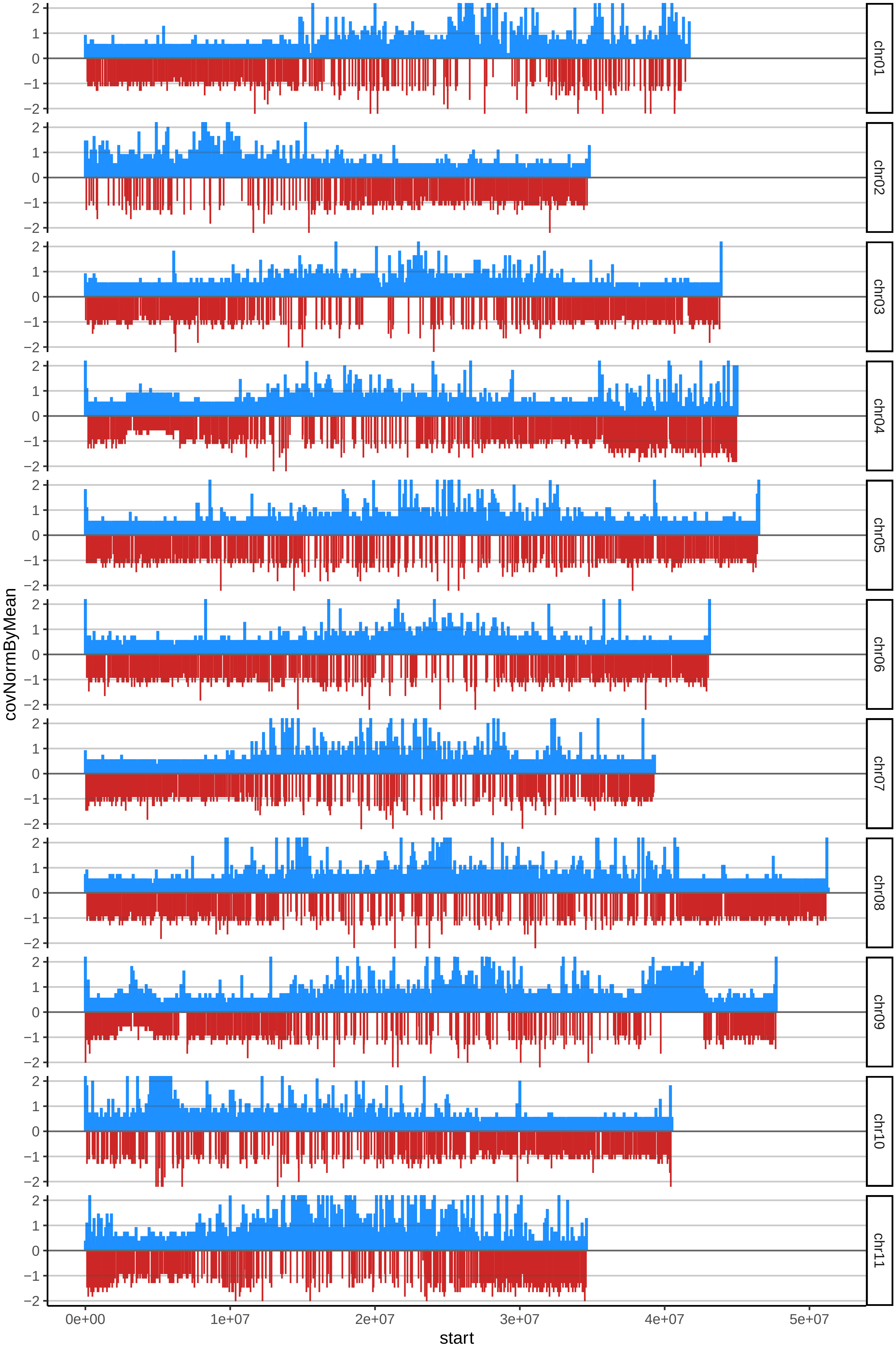

### 8CACHACOENANOcov_byB.pdf

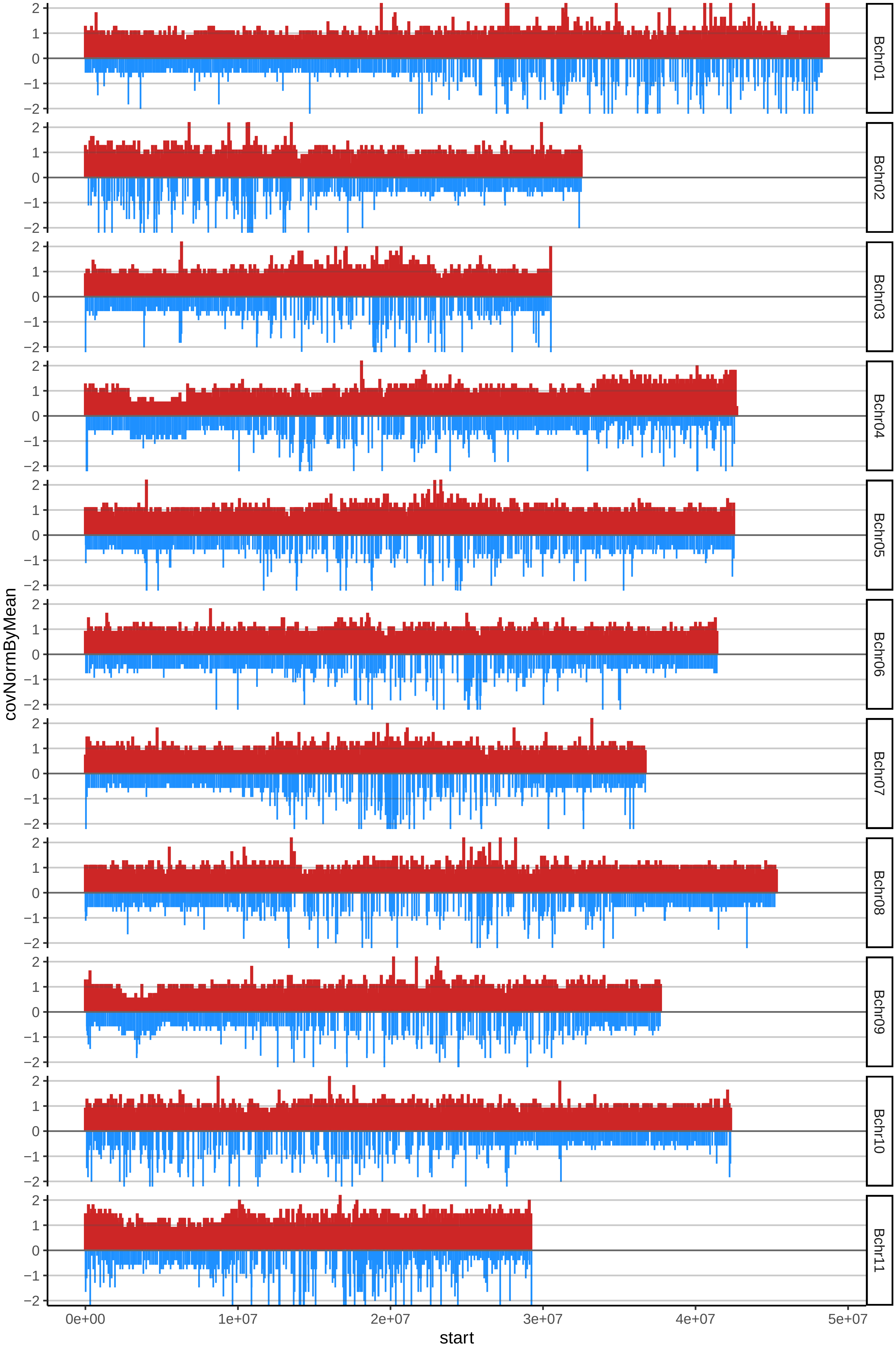

### 9CACHACOESPERMOcov_byA.pdf

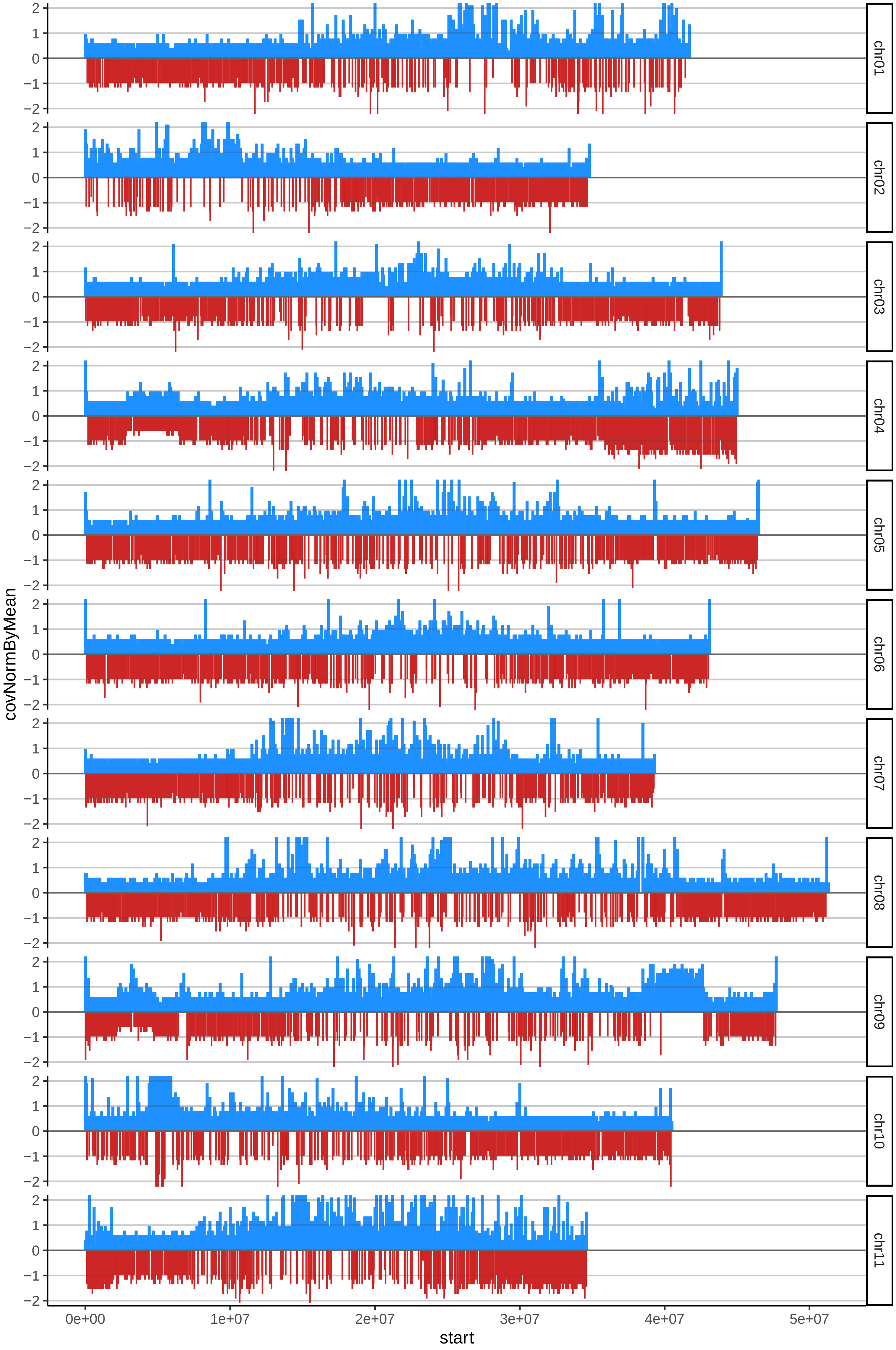

### 9CACHACOESPERMOcov_byB.pdf

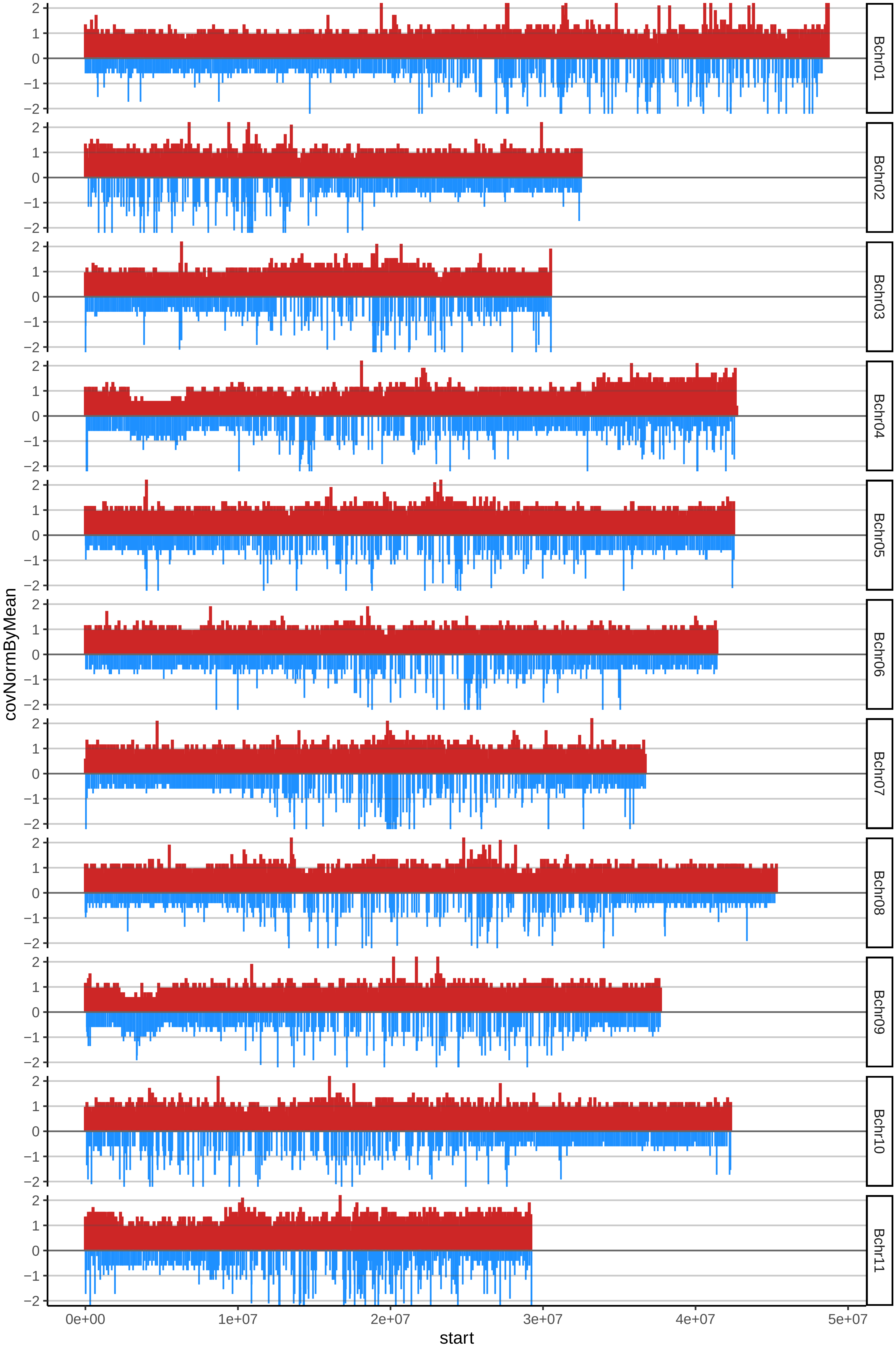

### 10CACHACOSINBELLOTAcov_byA.pdf

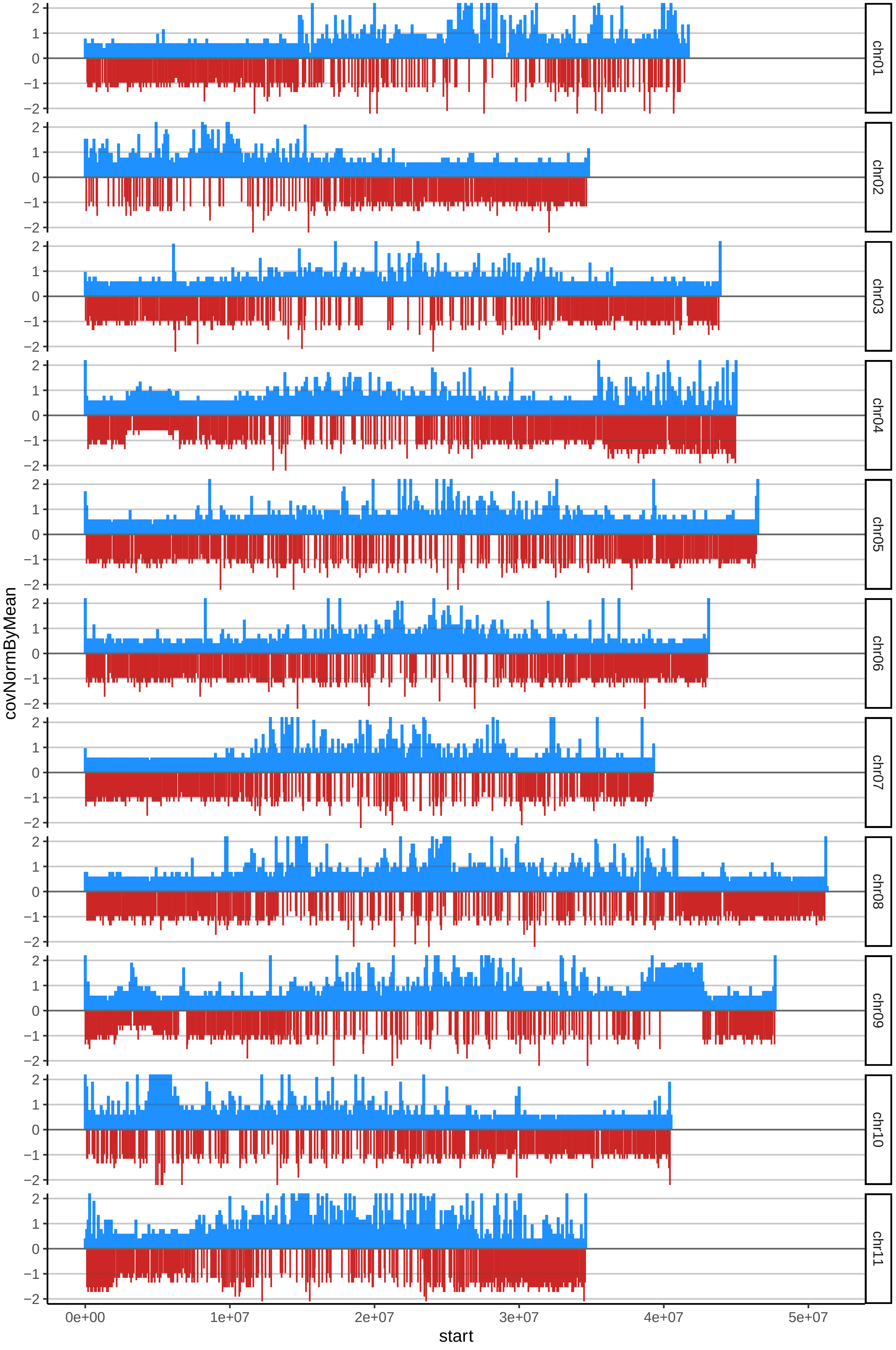

### 10CACHACOSINBELLOTAcov_byB.pdf

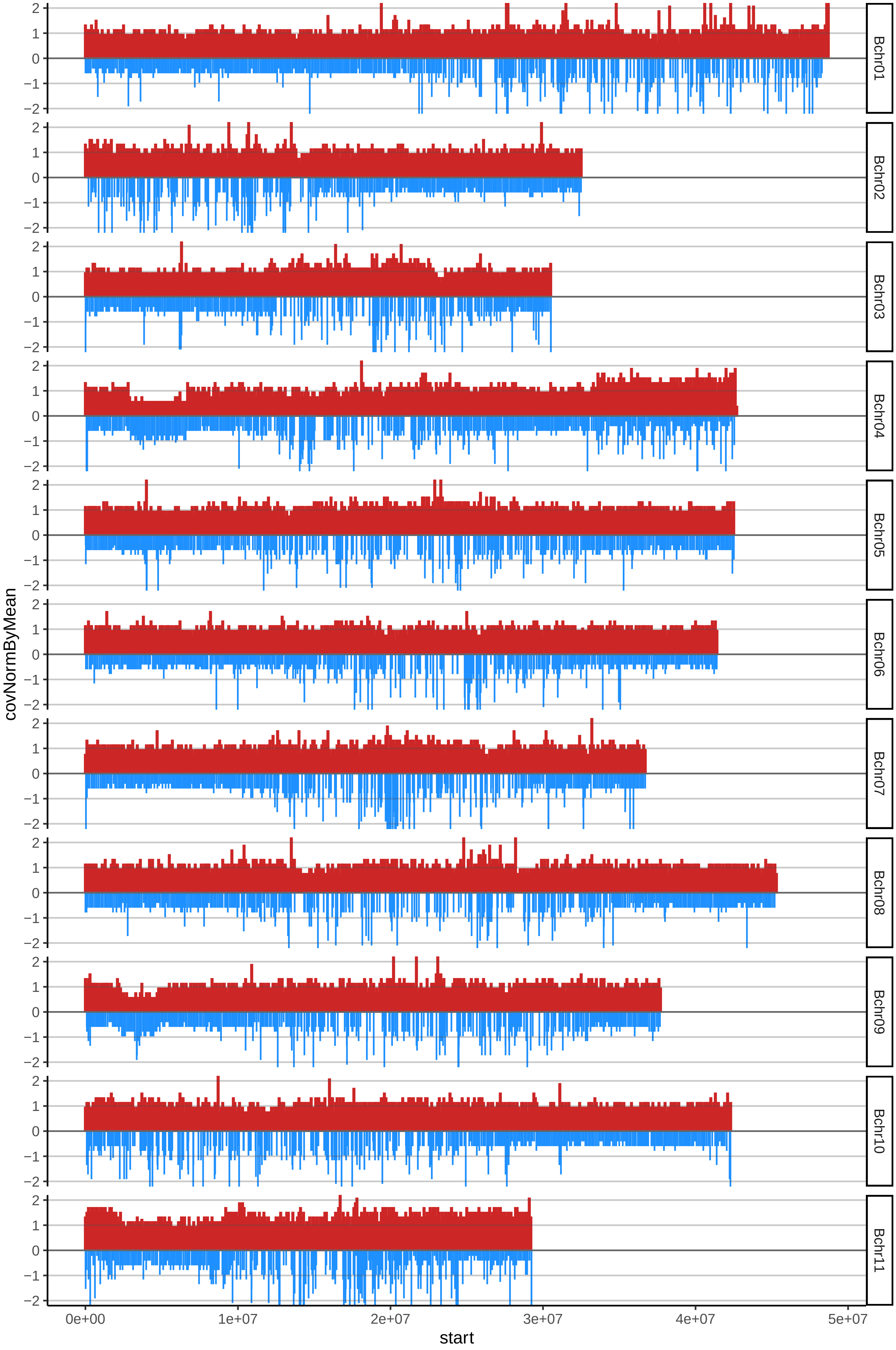

### 13DOMINICO300cov_byB.pdf

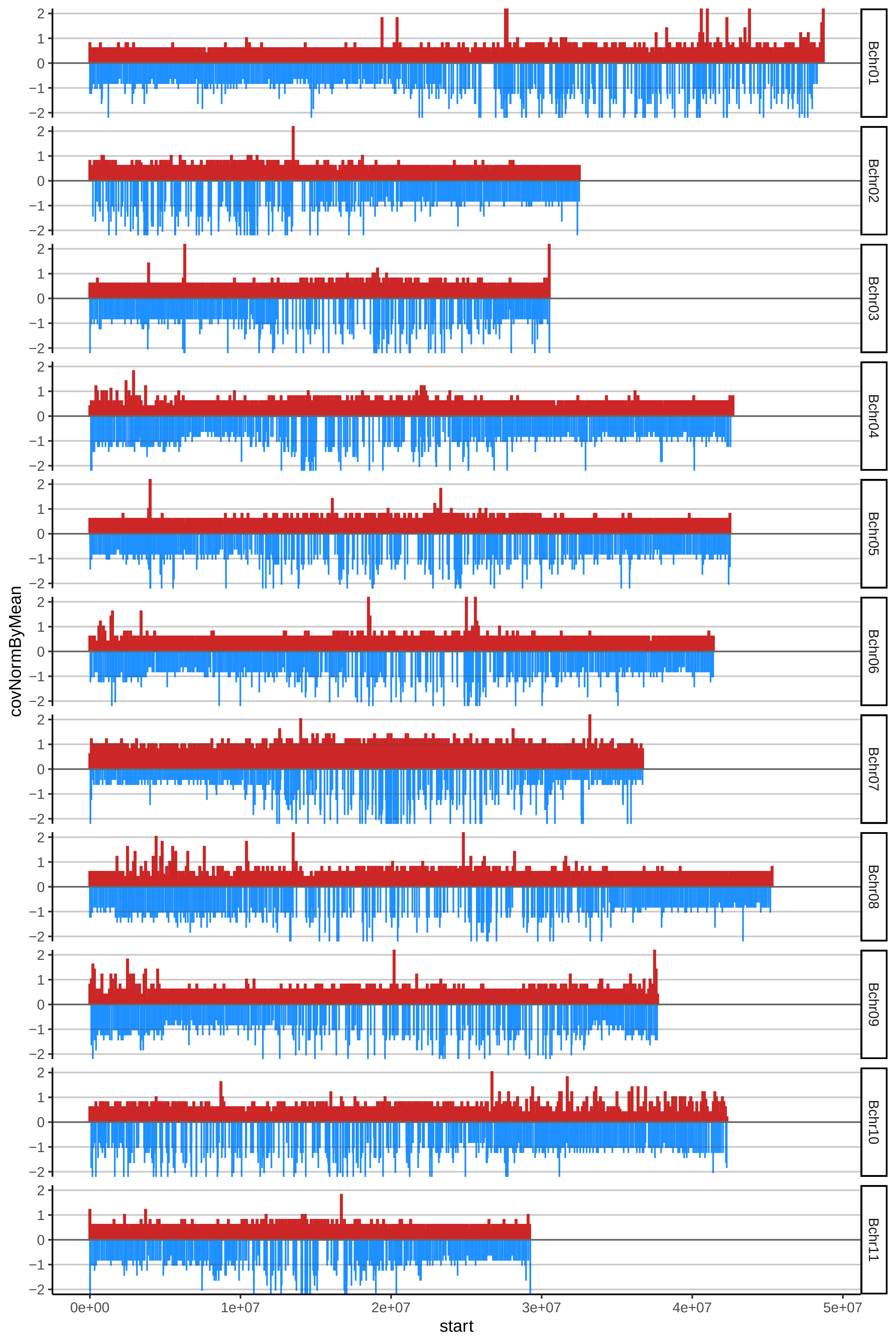

### 16DOMINICOCOMUNcov_byA.pdf

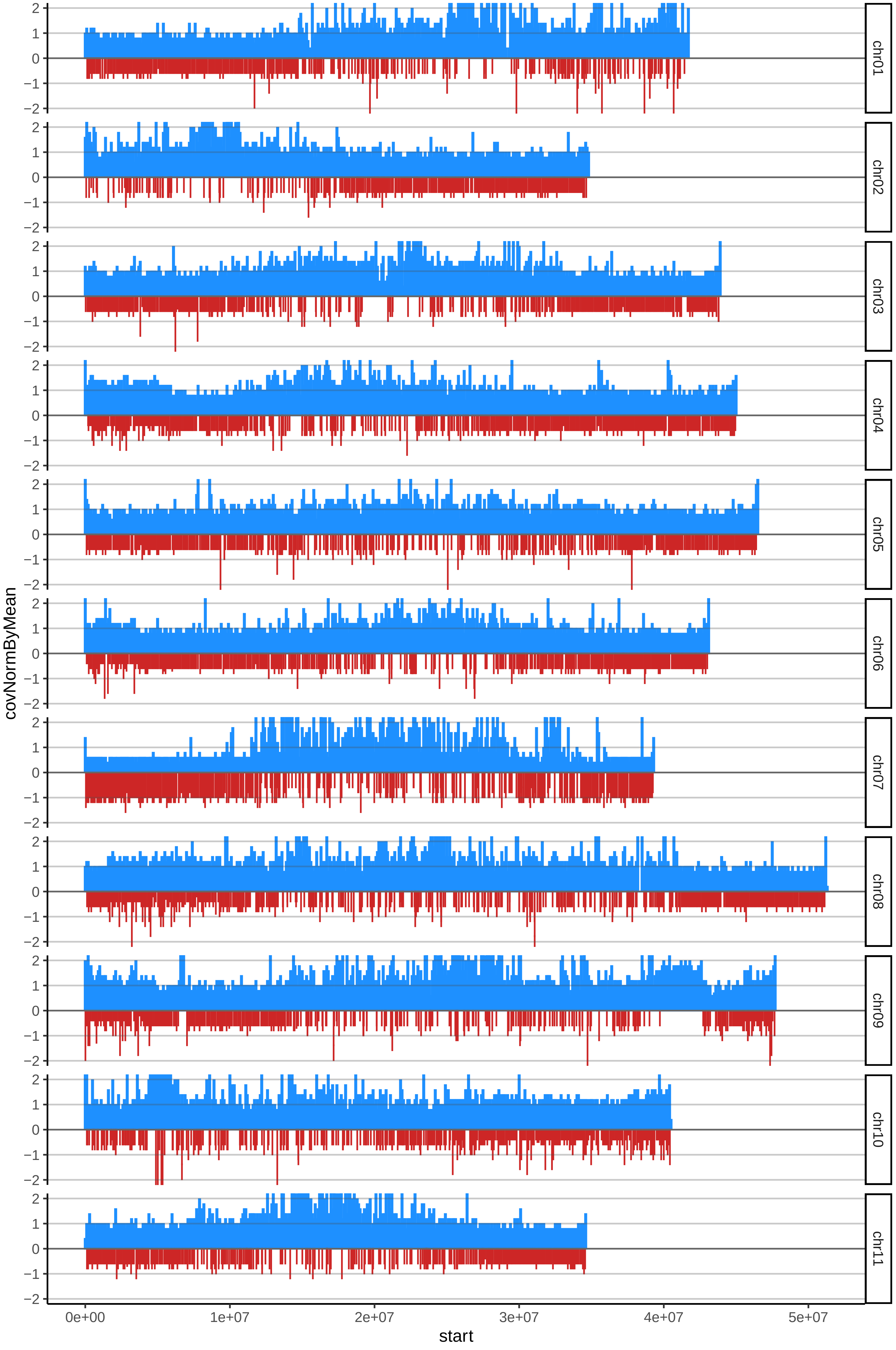

### 16DOMINICOCOMUNcov_byB.pdf

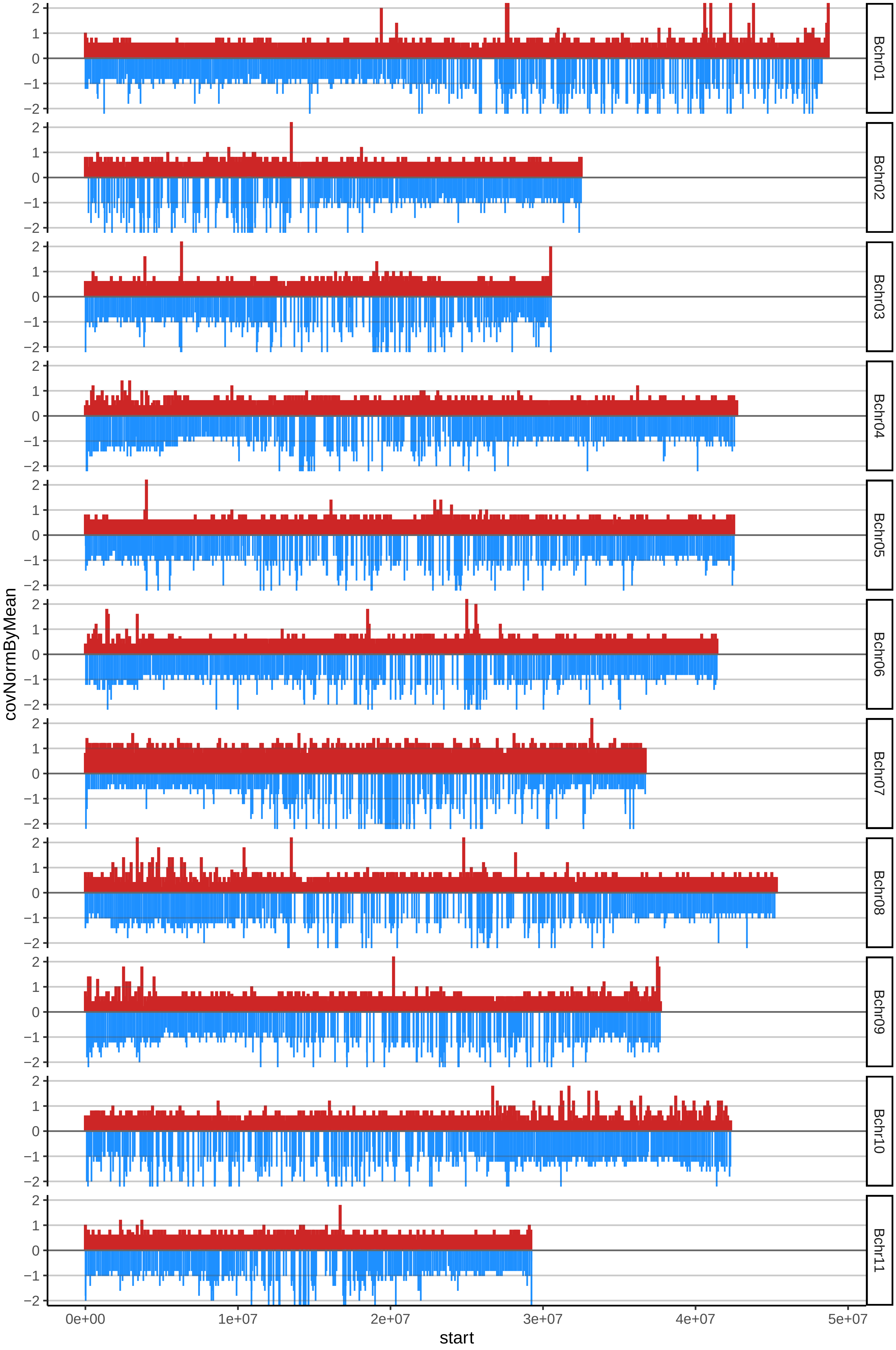

### 17DOMINICOENANOcov_byA.pdf

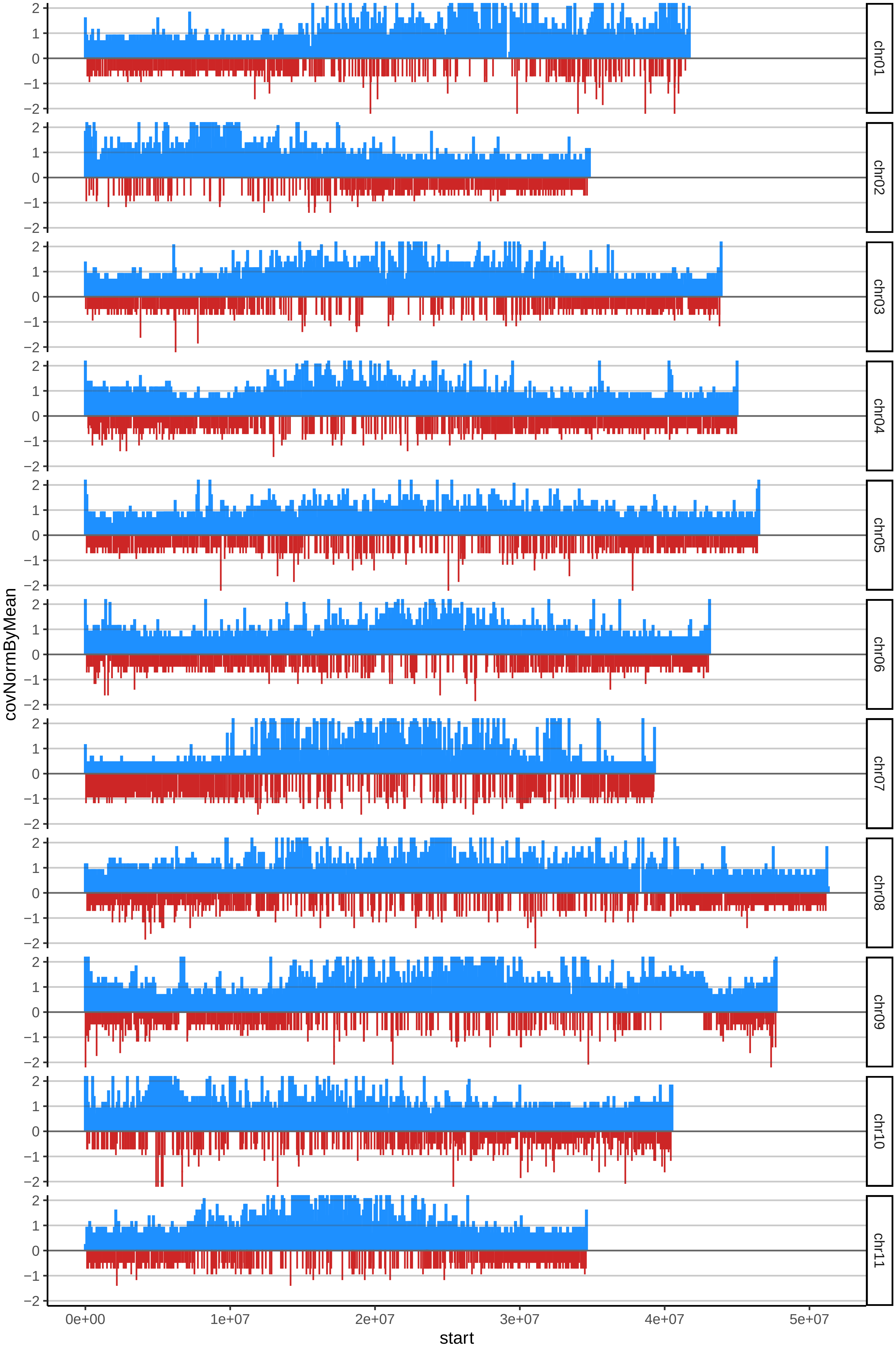

### 17DOMINICOENANOcov_byB.pdf

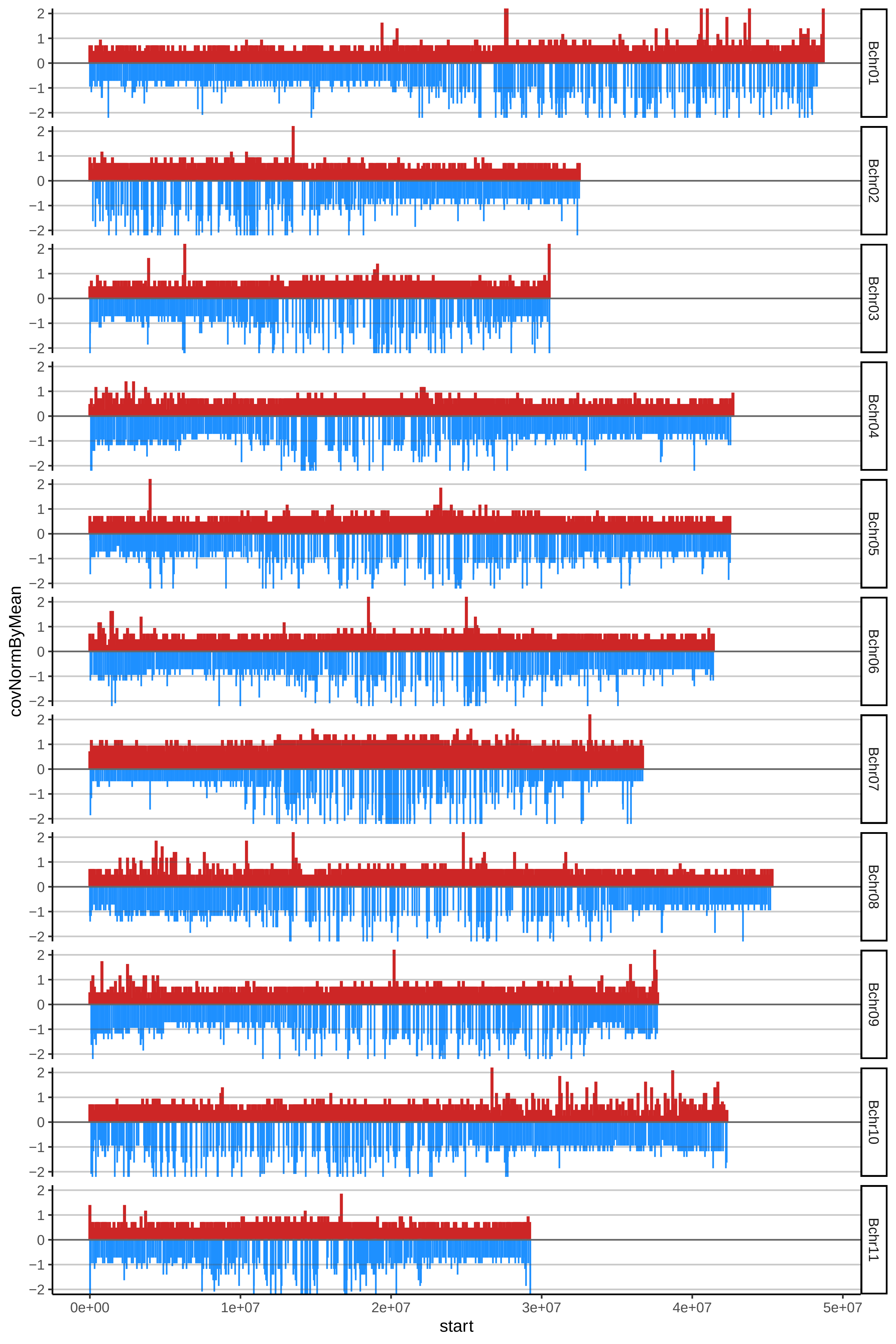

### 20DOMINICOHARTONROJOcov_byA.pdf

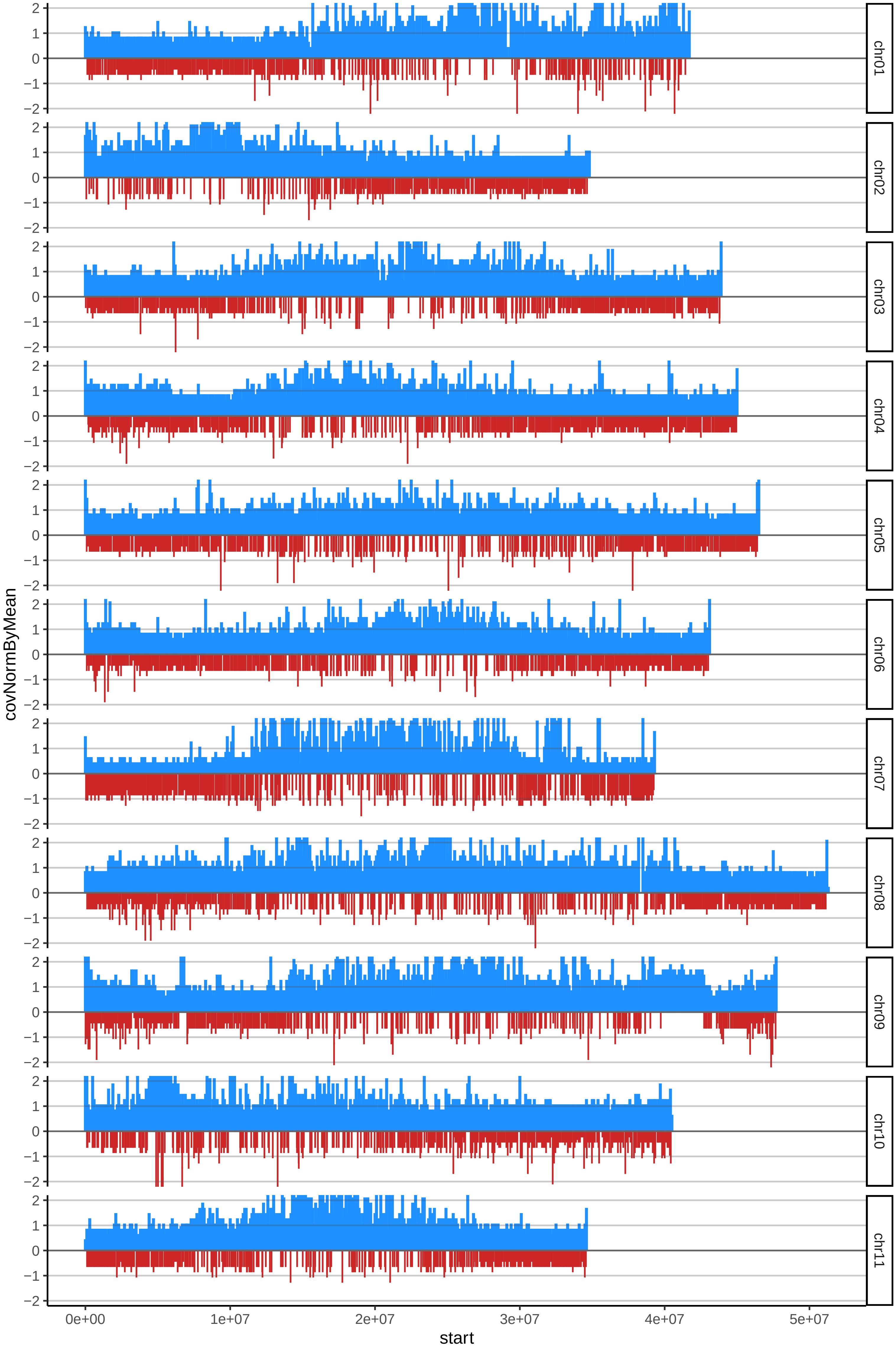

### 20DOMINICOHARTONROJOcov_byB.pdf

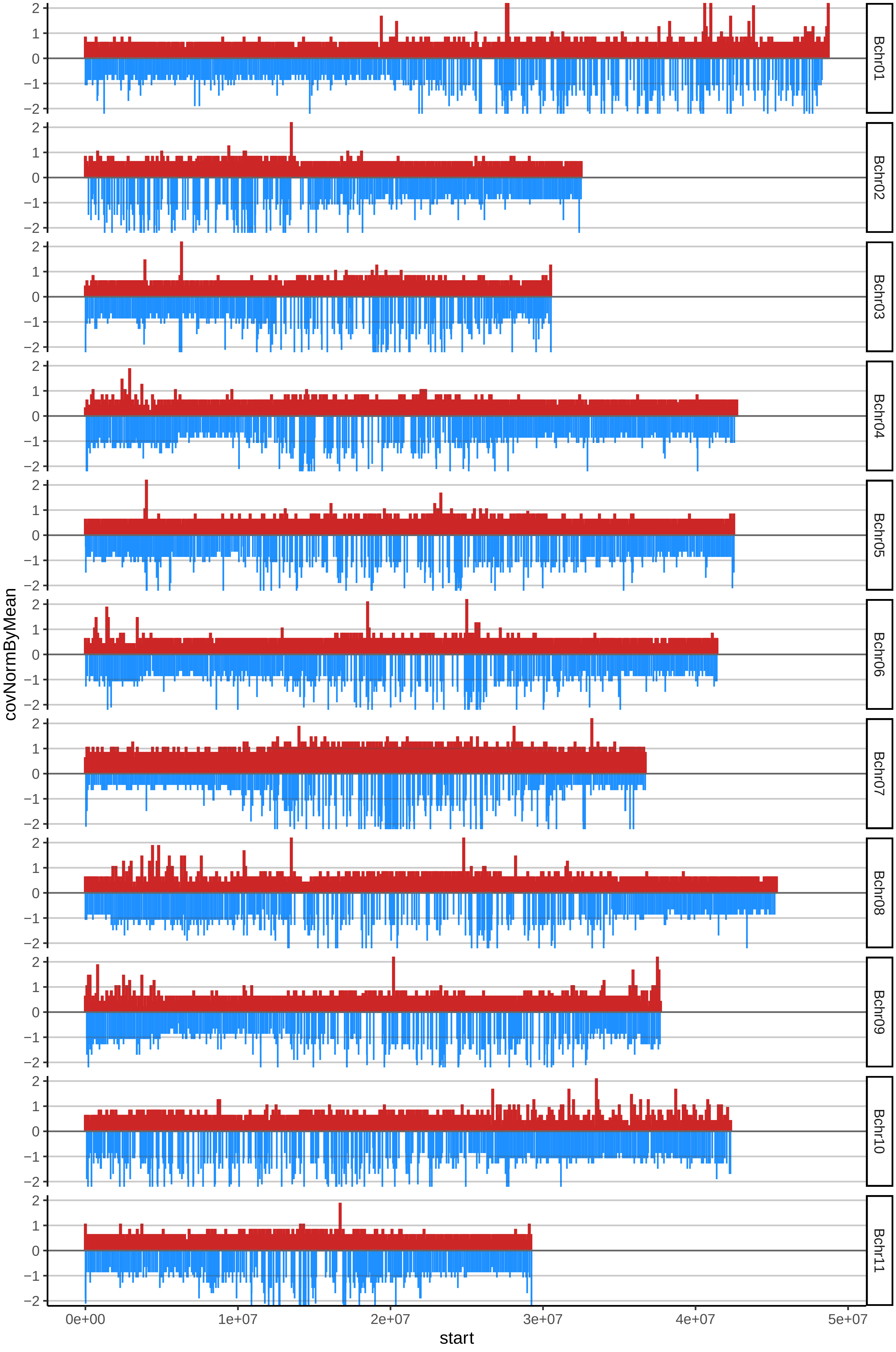

### 22DOMINICOHARTONVIOTAcov_byB.pdf

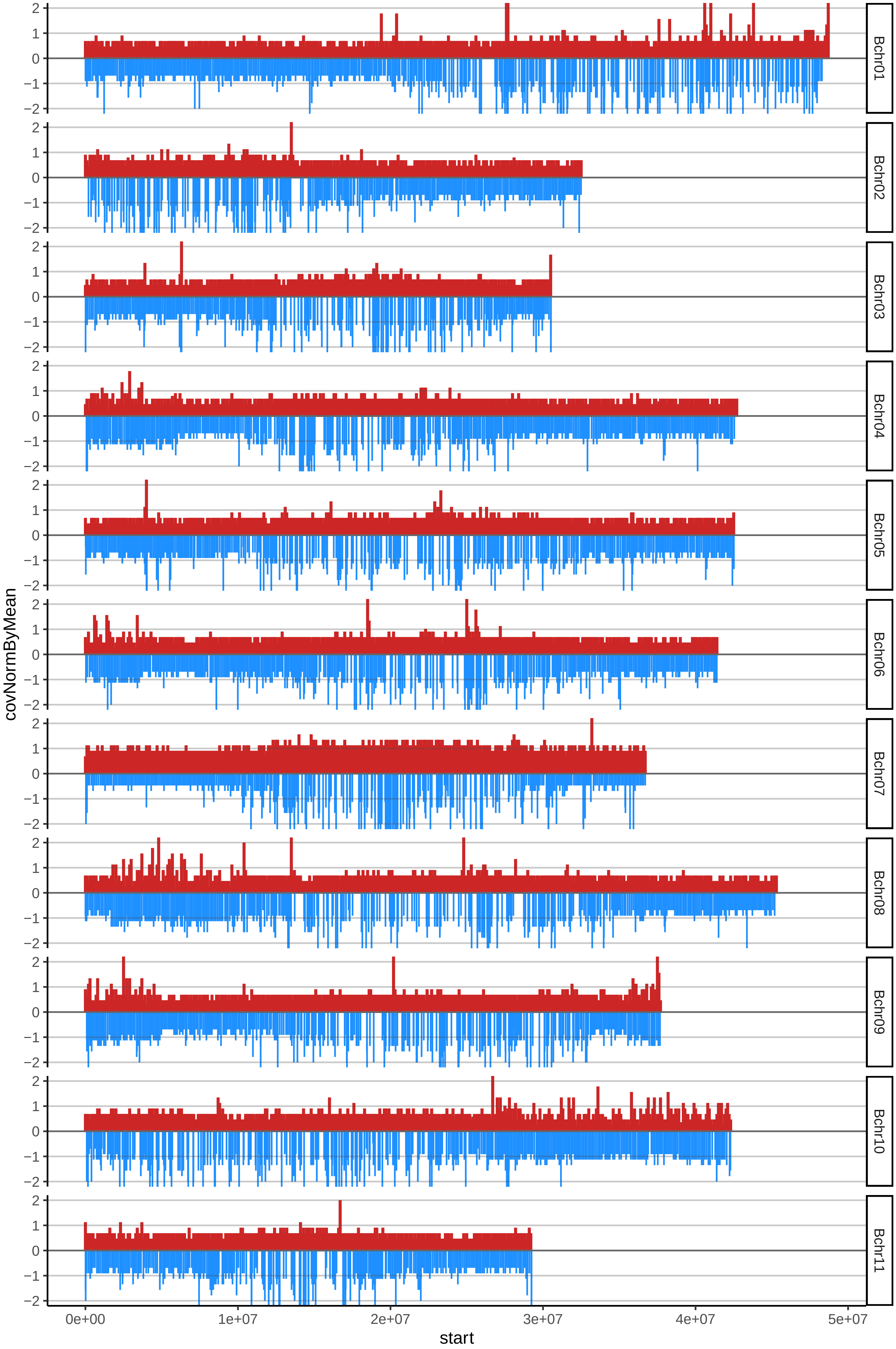

### 27DOMINICOTUMACOcov_byB.pdf

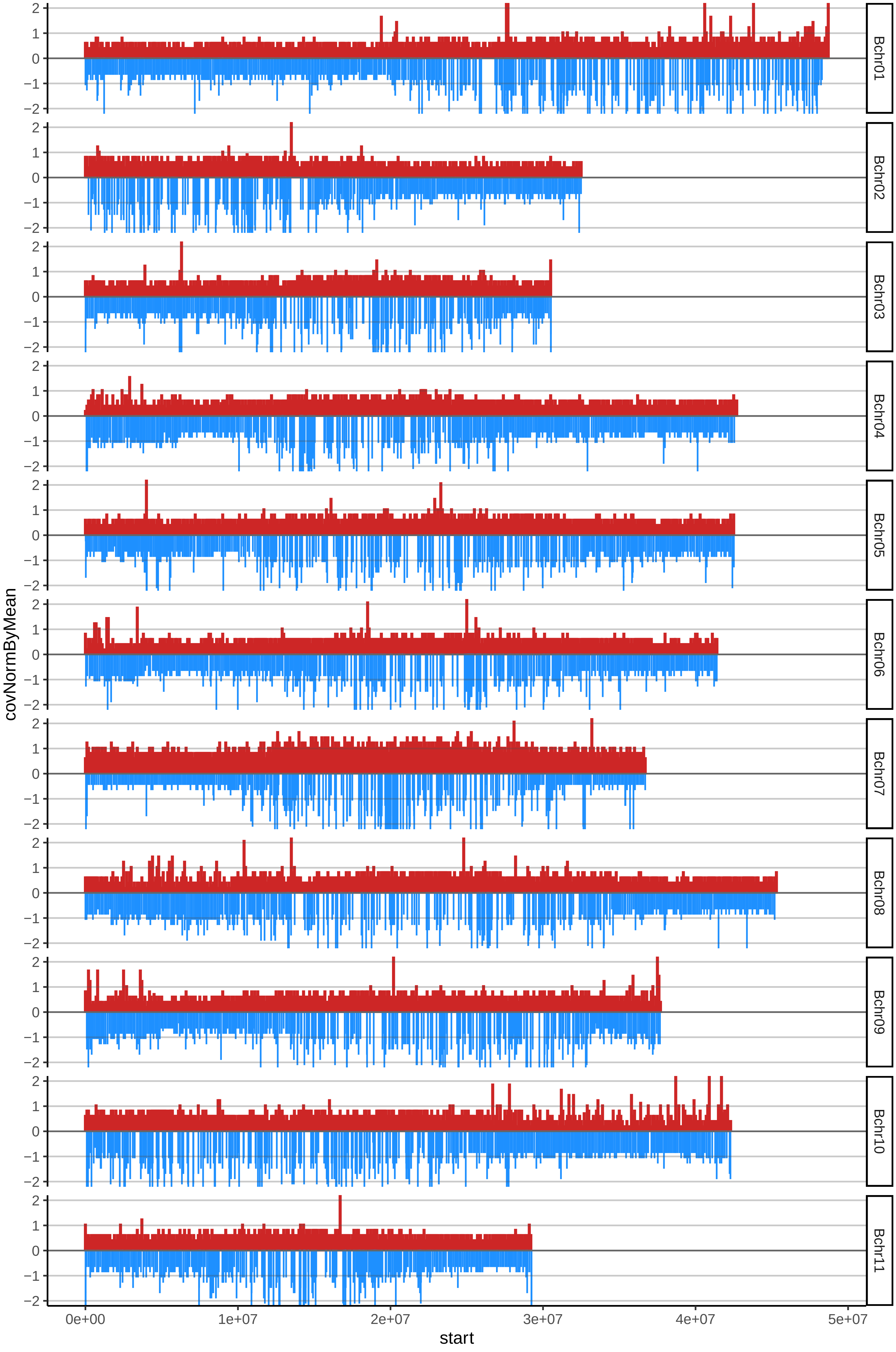

### 34FHIA25cov_byA.pdf

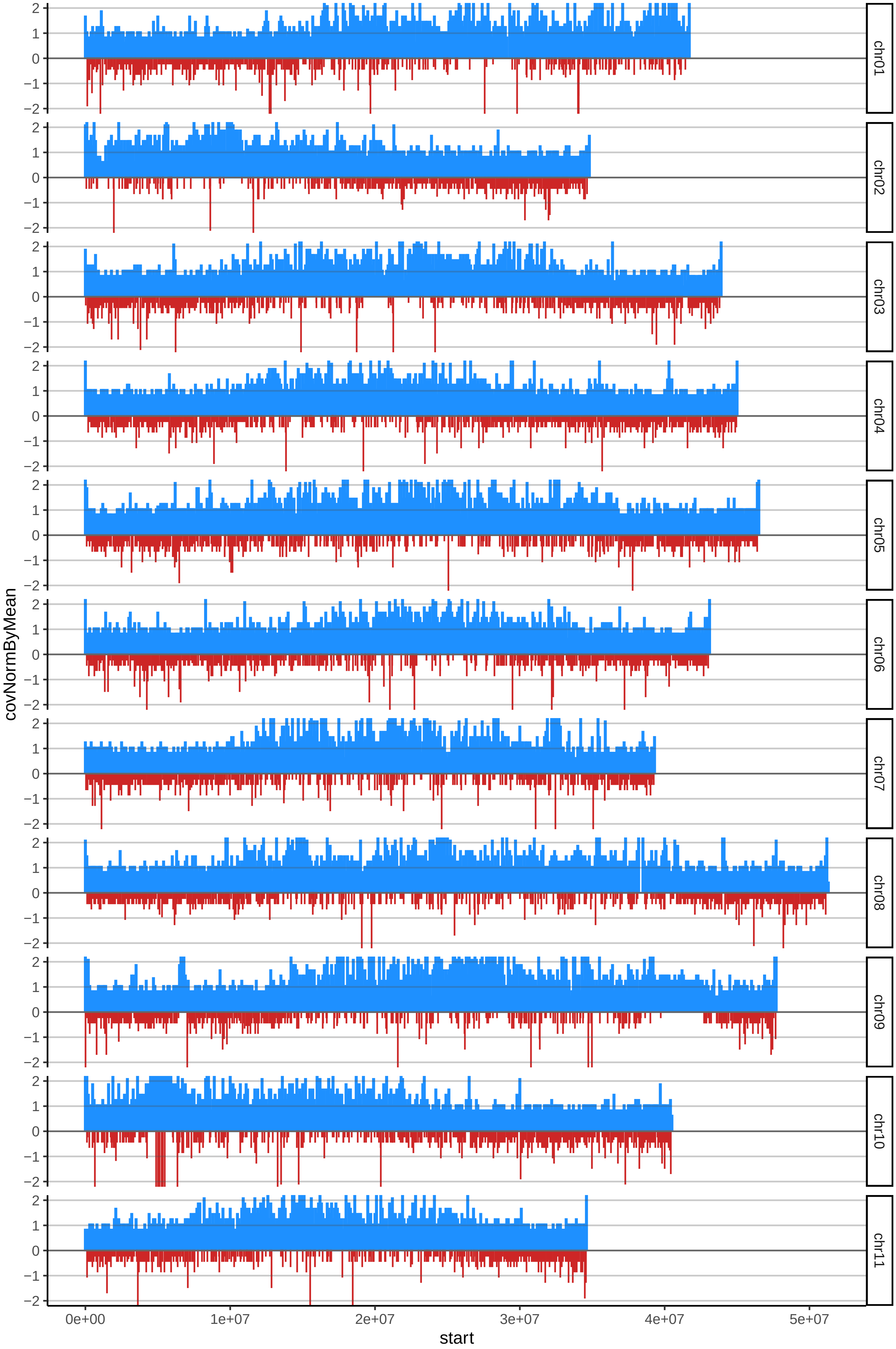

### 34FHIA25cov_byB.pdf

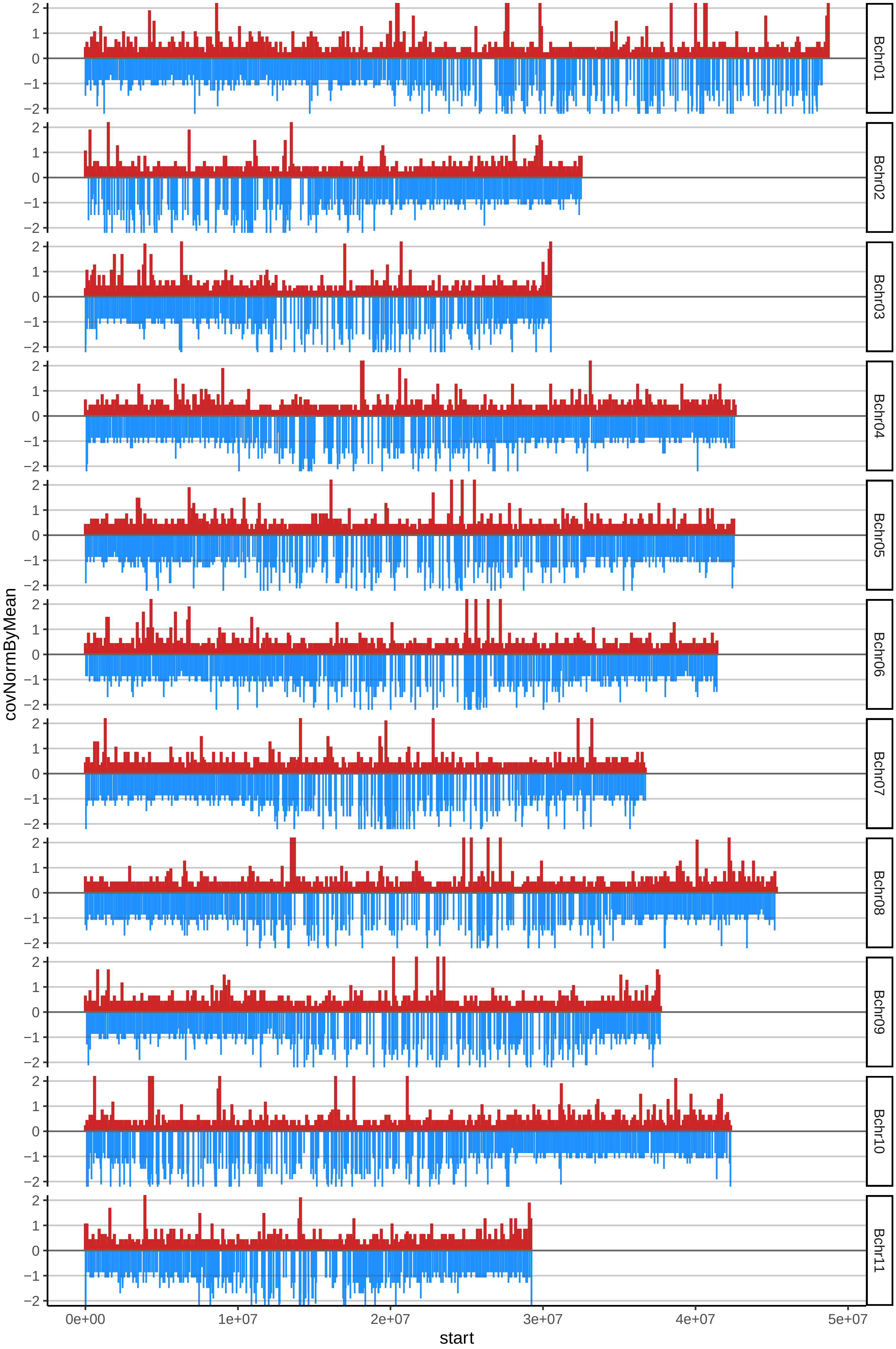

### 36FRENCHSOMBREcov_byA.pdf

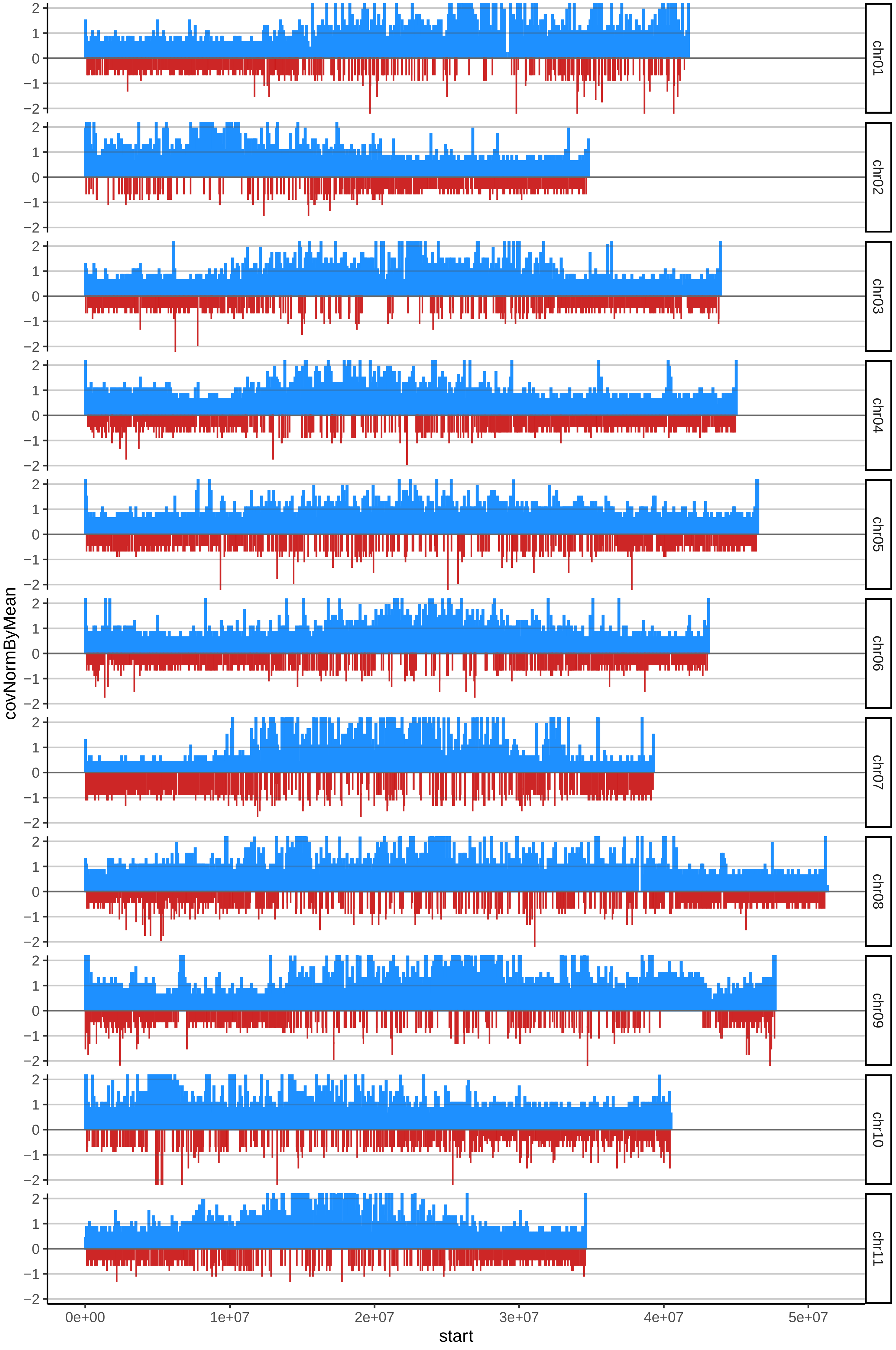

### 40HARTONBIRRACIMOcov_byA.pdf

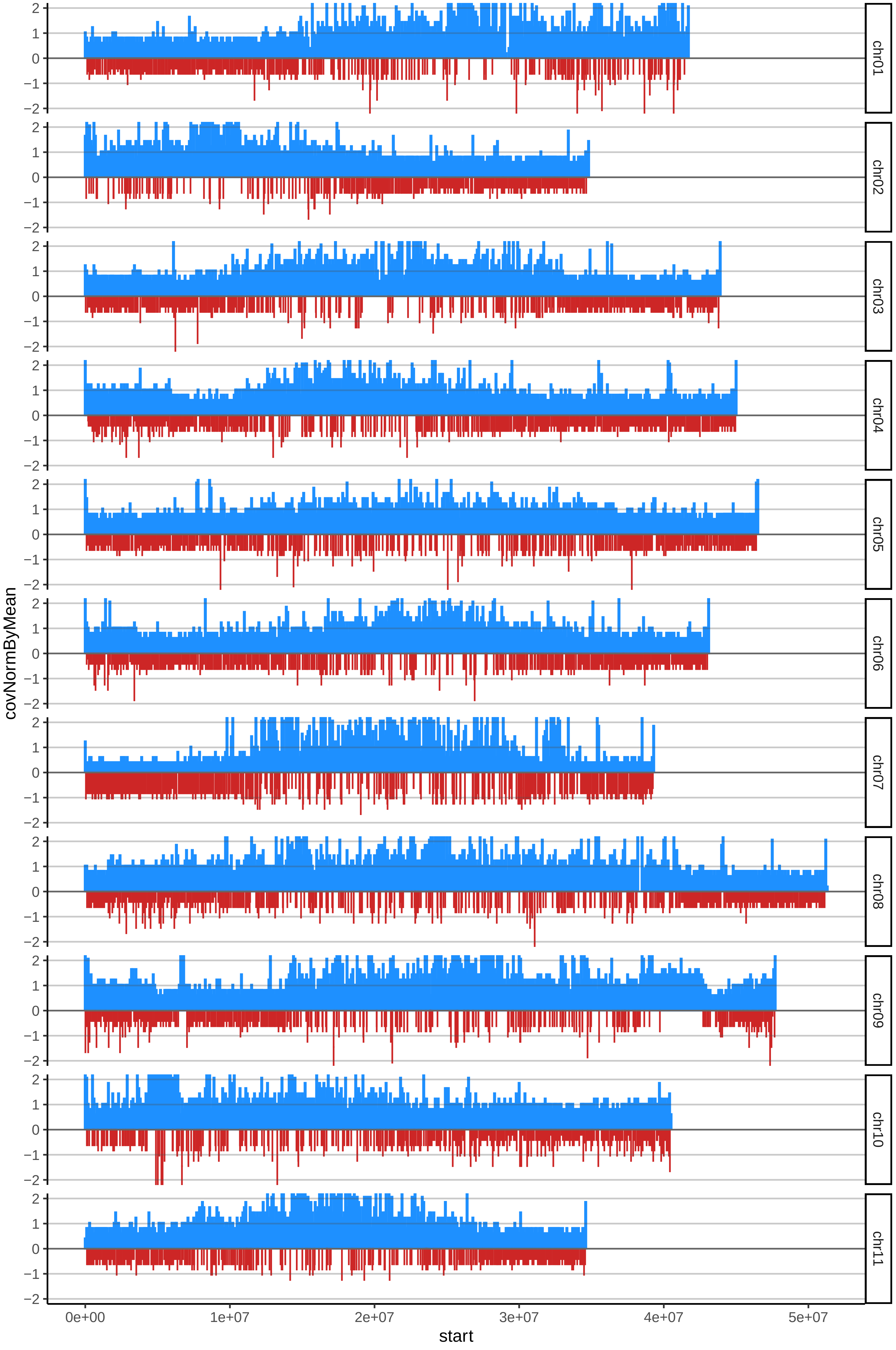

### 40HARTONBIRRACIMOcov_byB.pdf

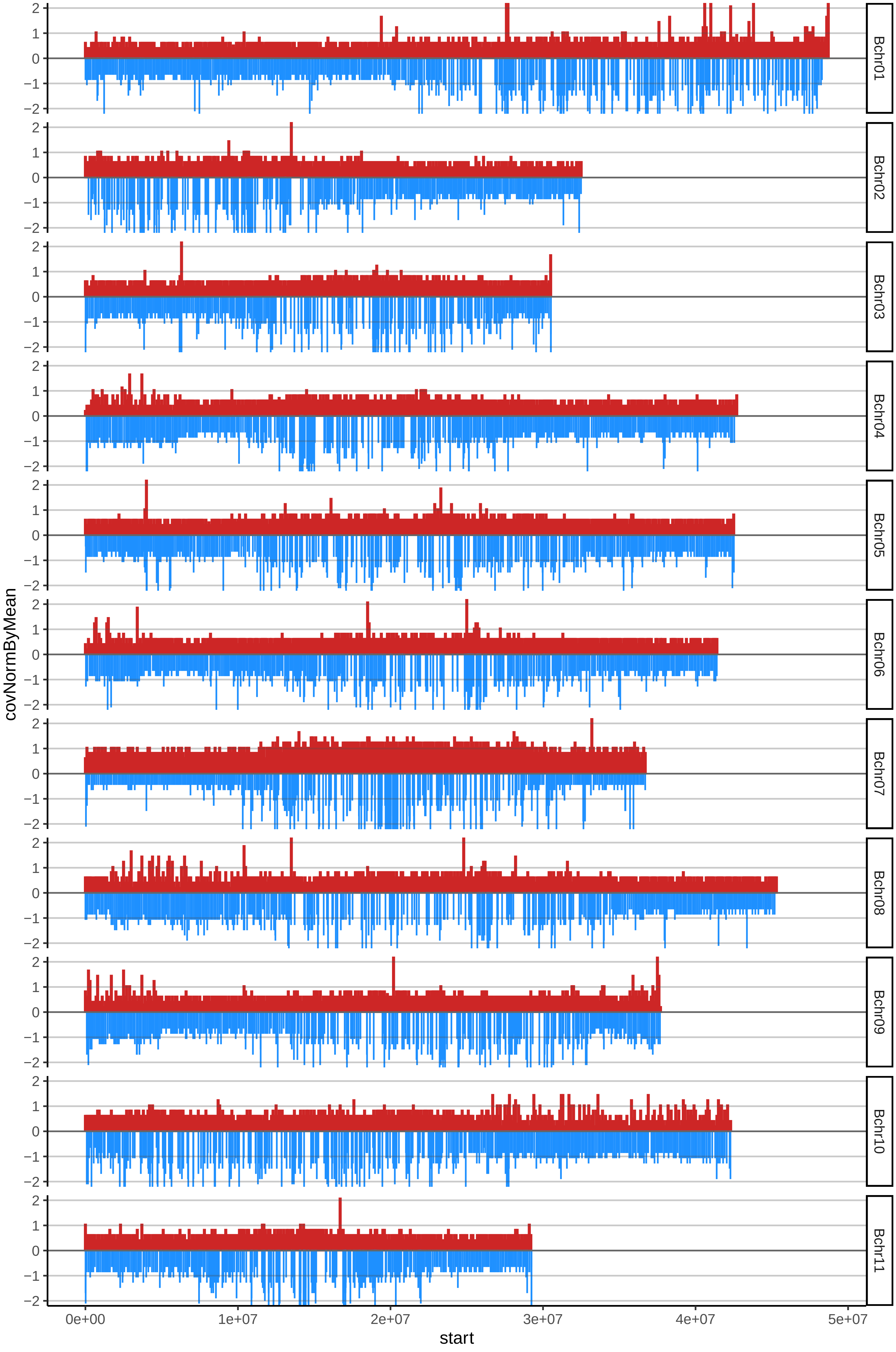

### 48HARTONTUMACOcov_byA.pdf

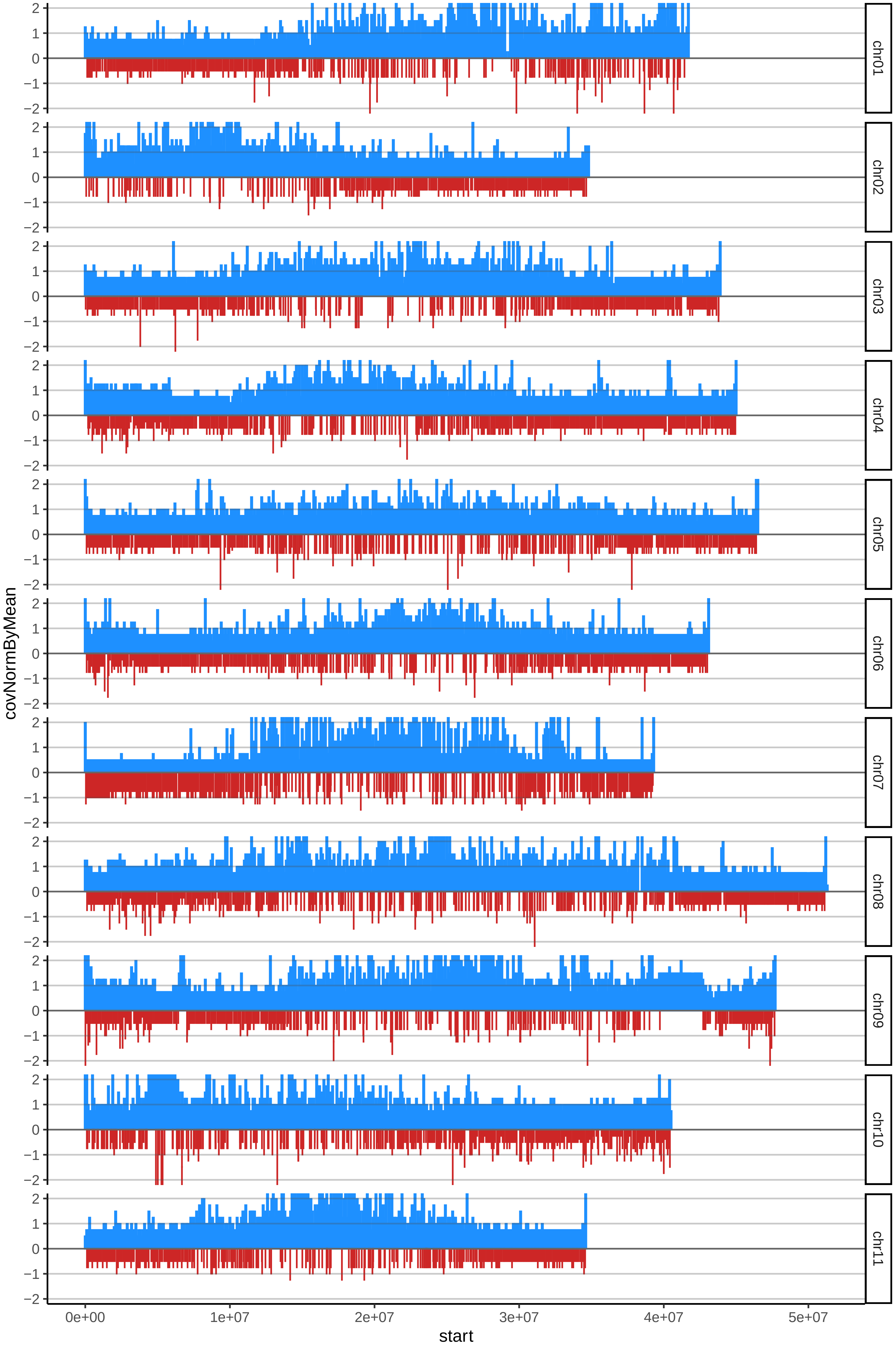

### 48HARTONTUMACOcov_byB.pdf

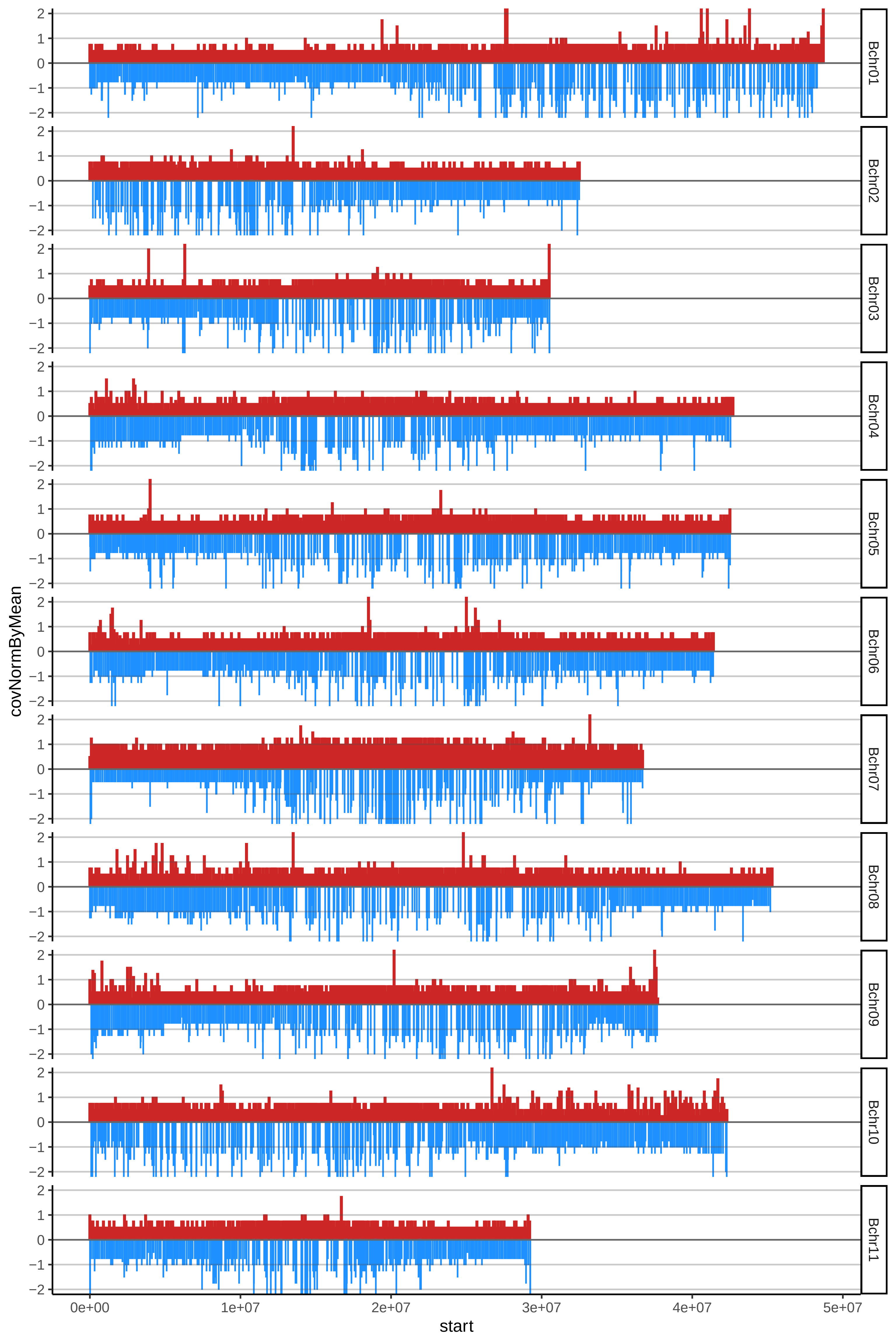

### 59MBOUROUKOUcov_byA.pdf

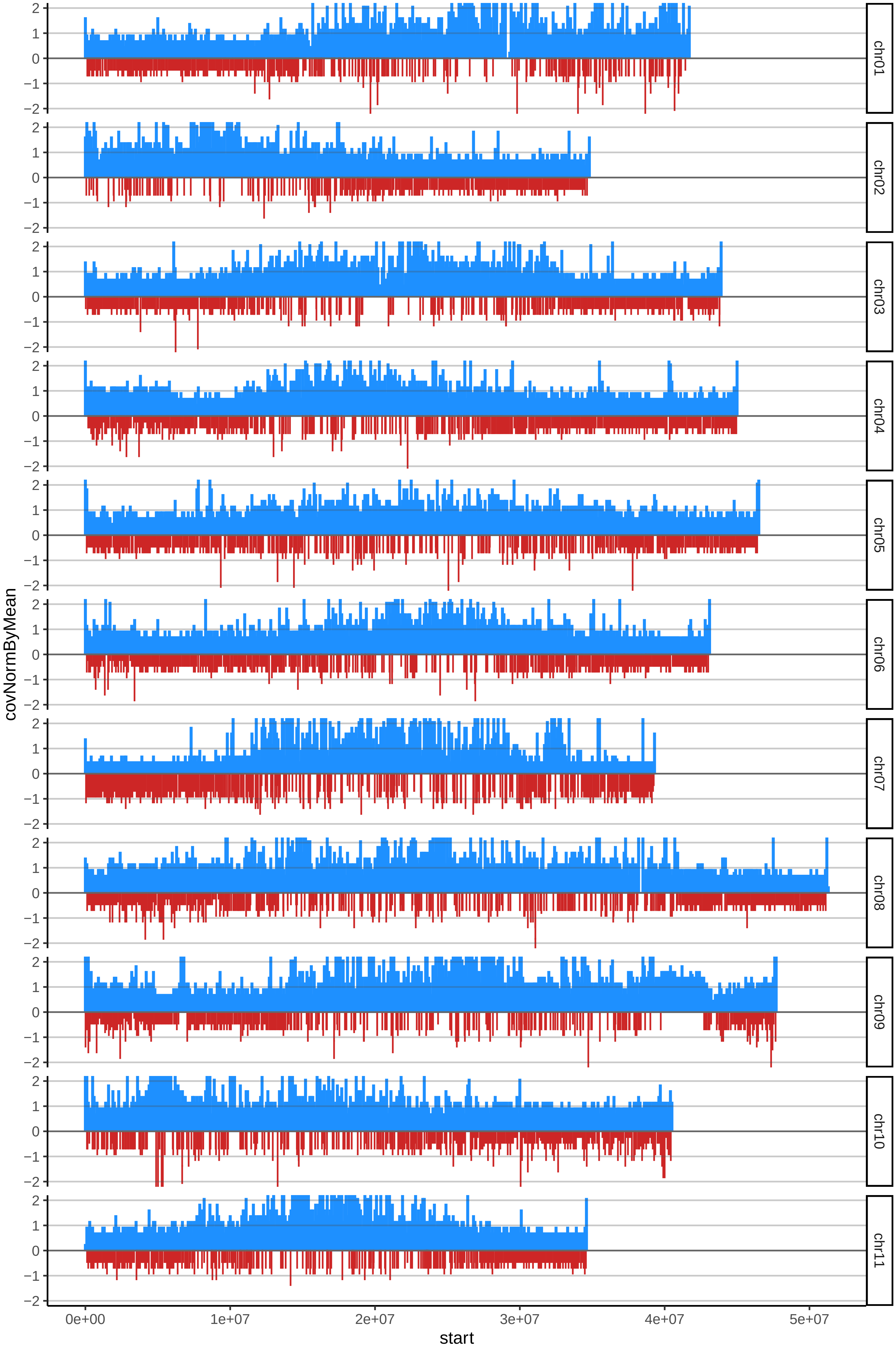

### 59MBOUROUKOUcov_byB.pdf

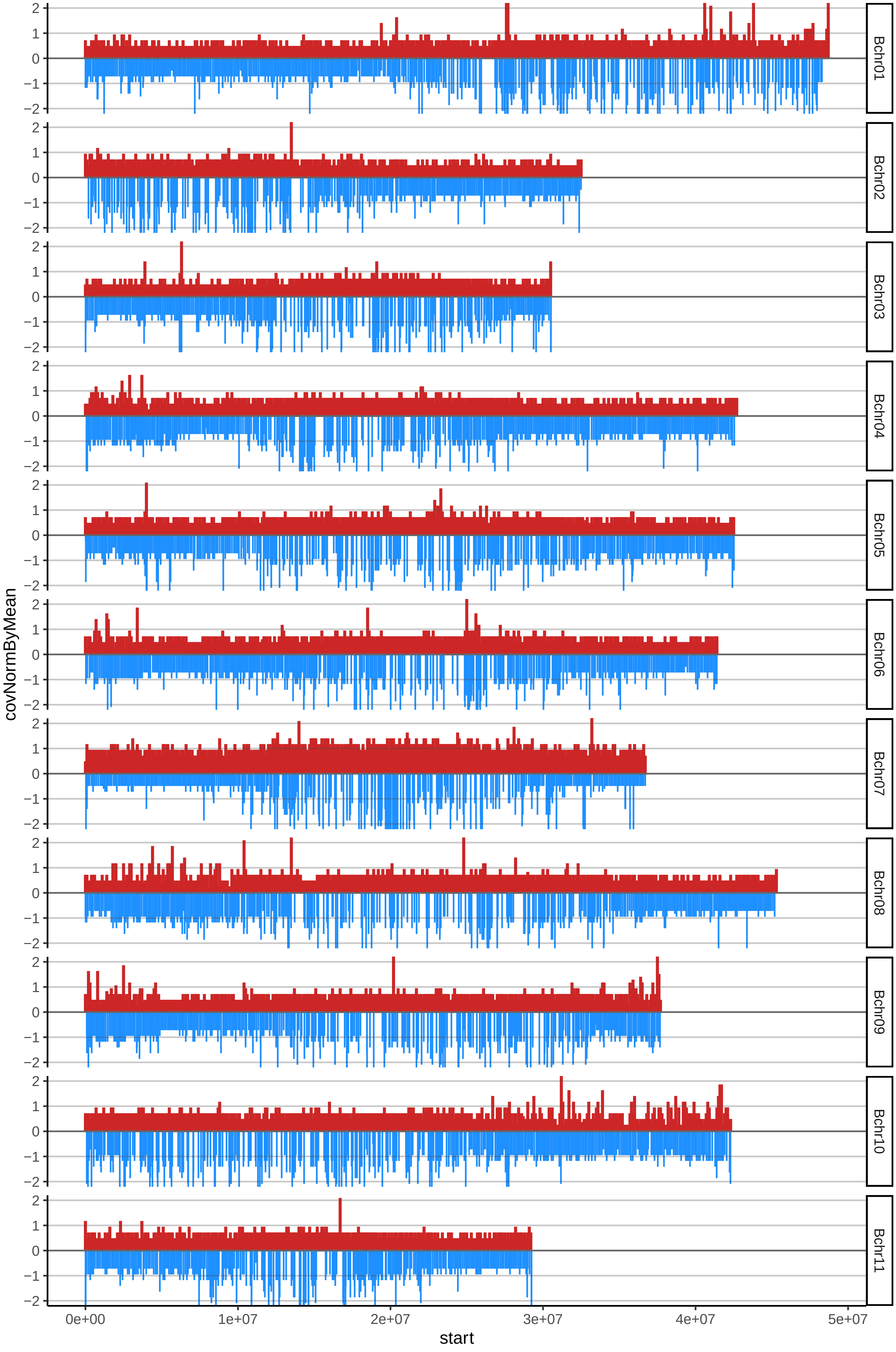

### 72REDYADEcov_byA.pdf

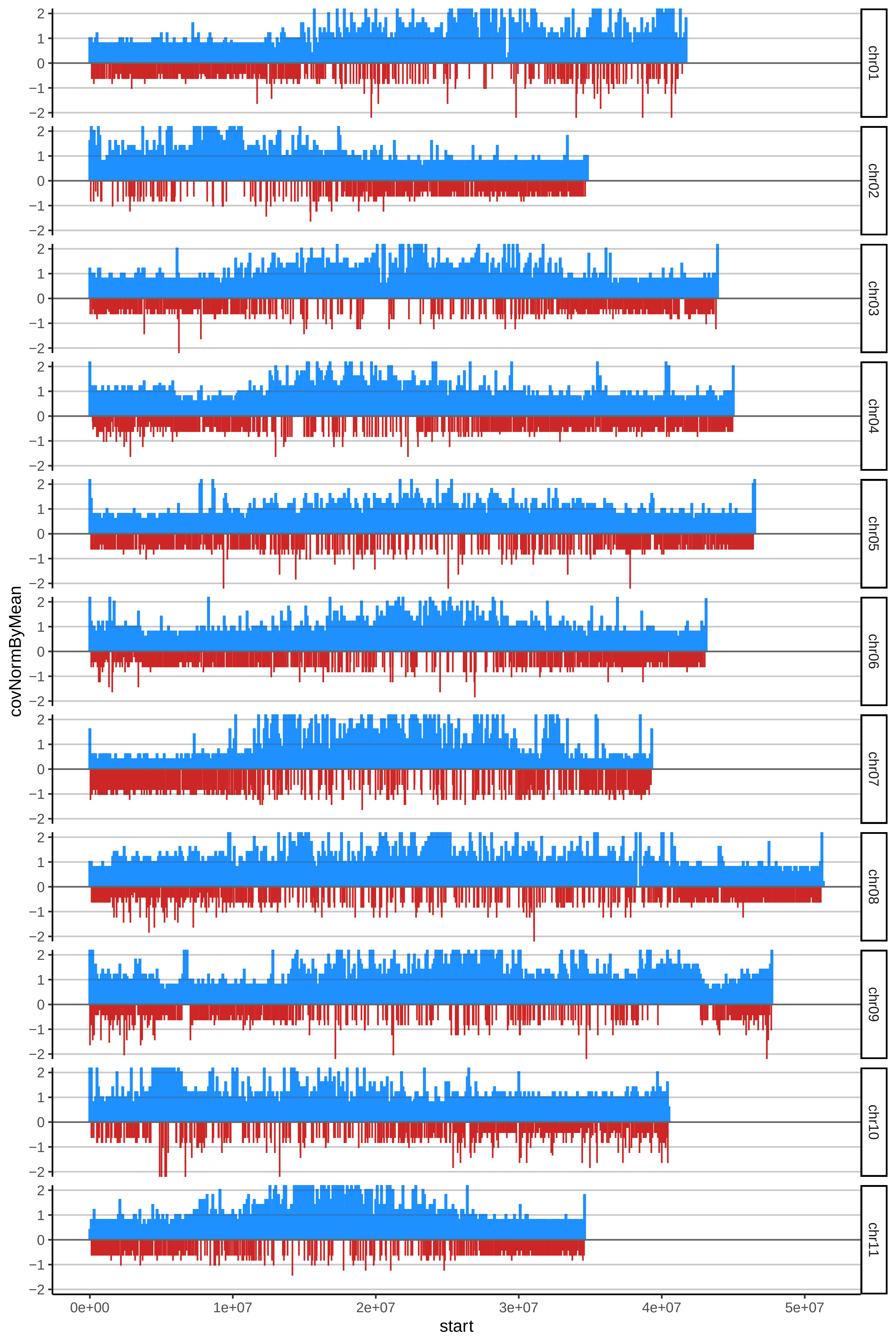

### 72REDYADEcov_byB.pdf

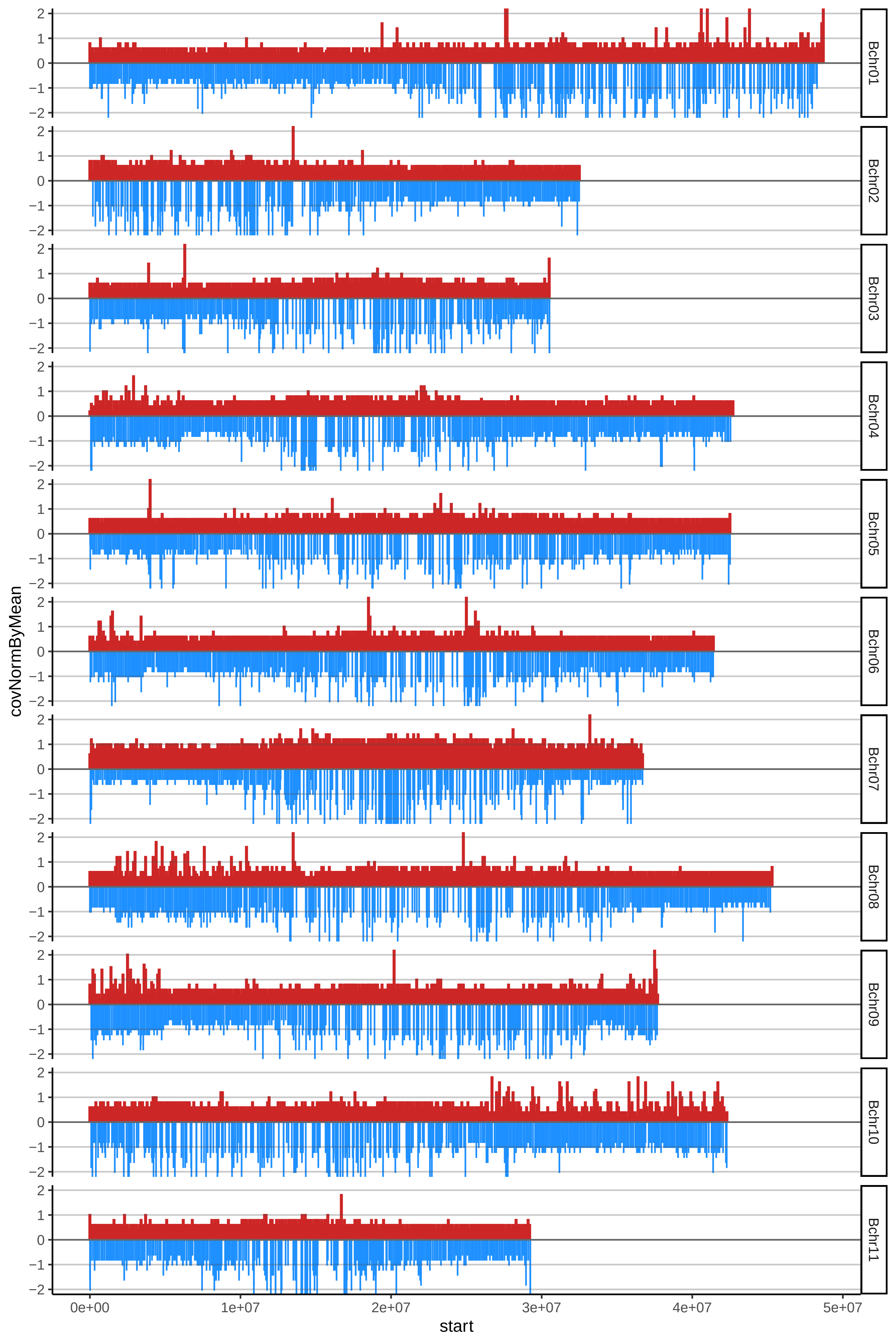

### 73ROSEDEKONAcov_byA.pdf

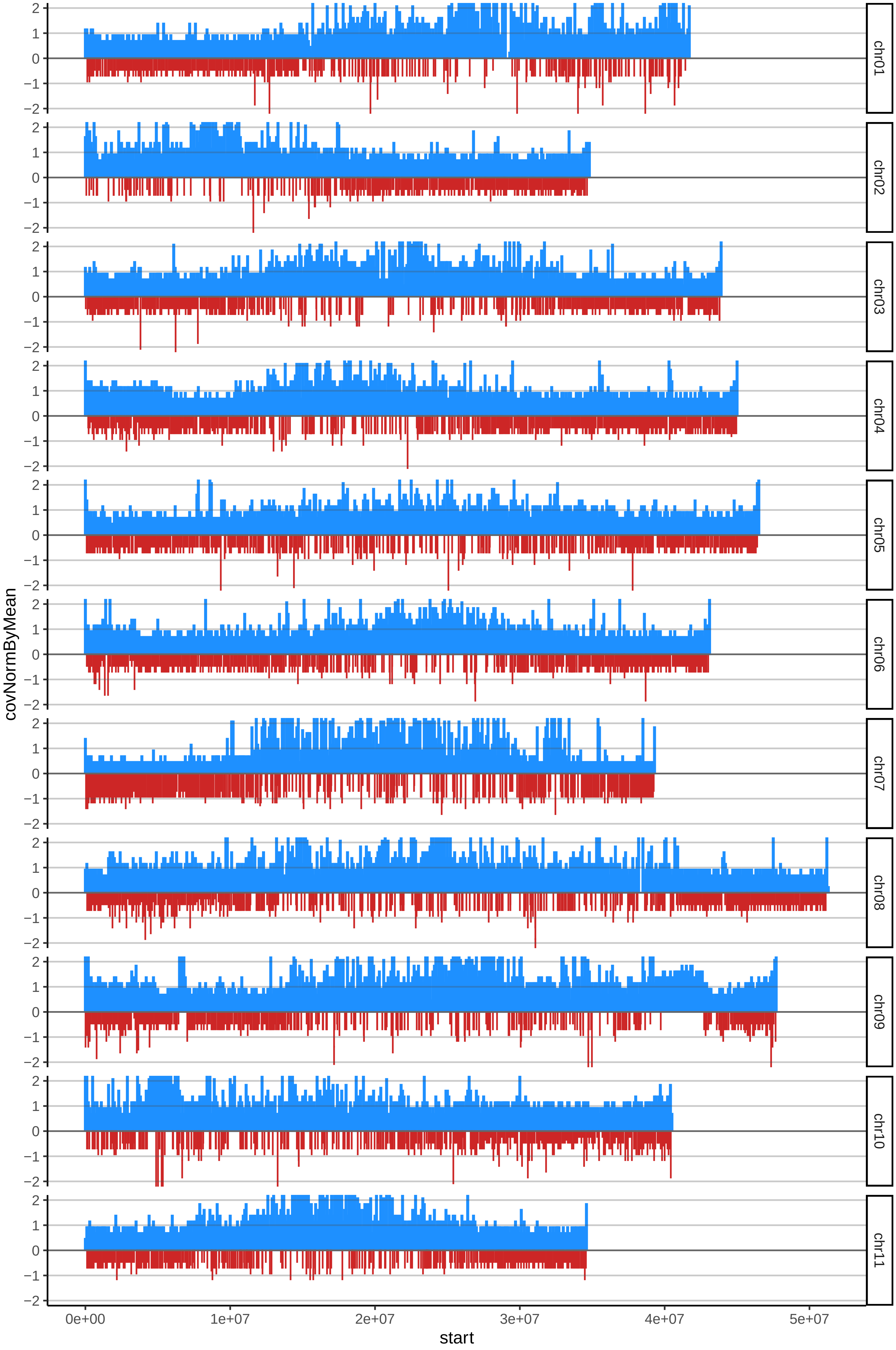

### 99MYSOREcov_byA.pdf

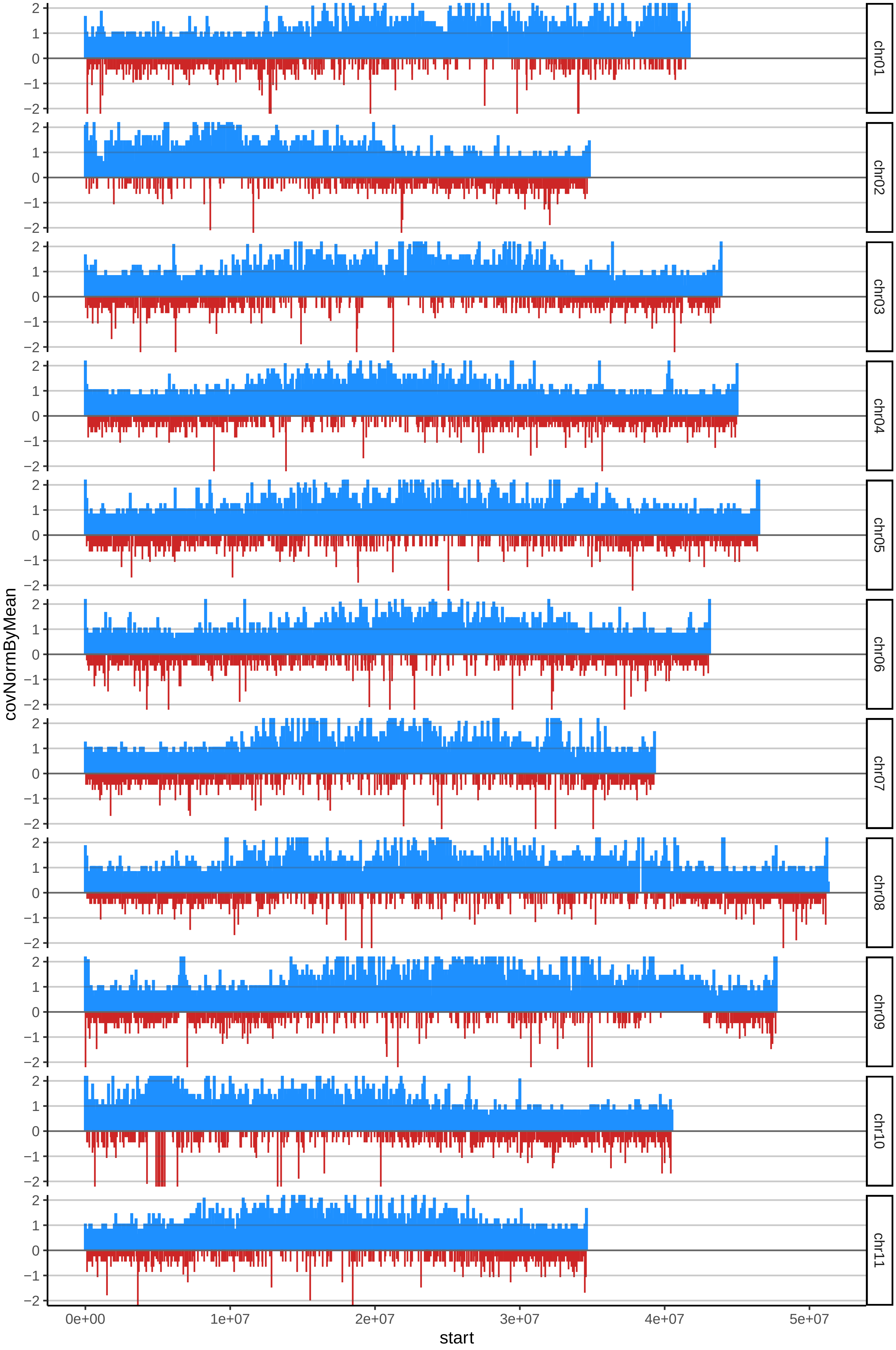

### 99MYSOREcov_byB.pdf

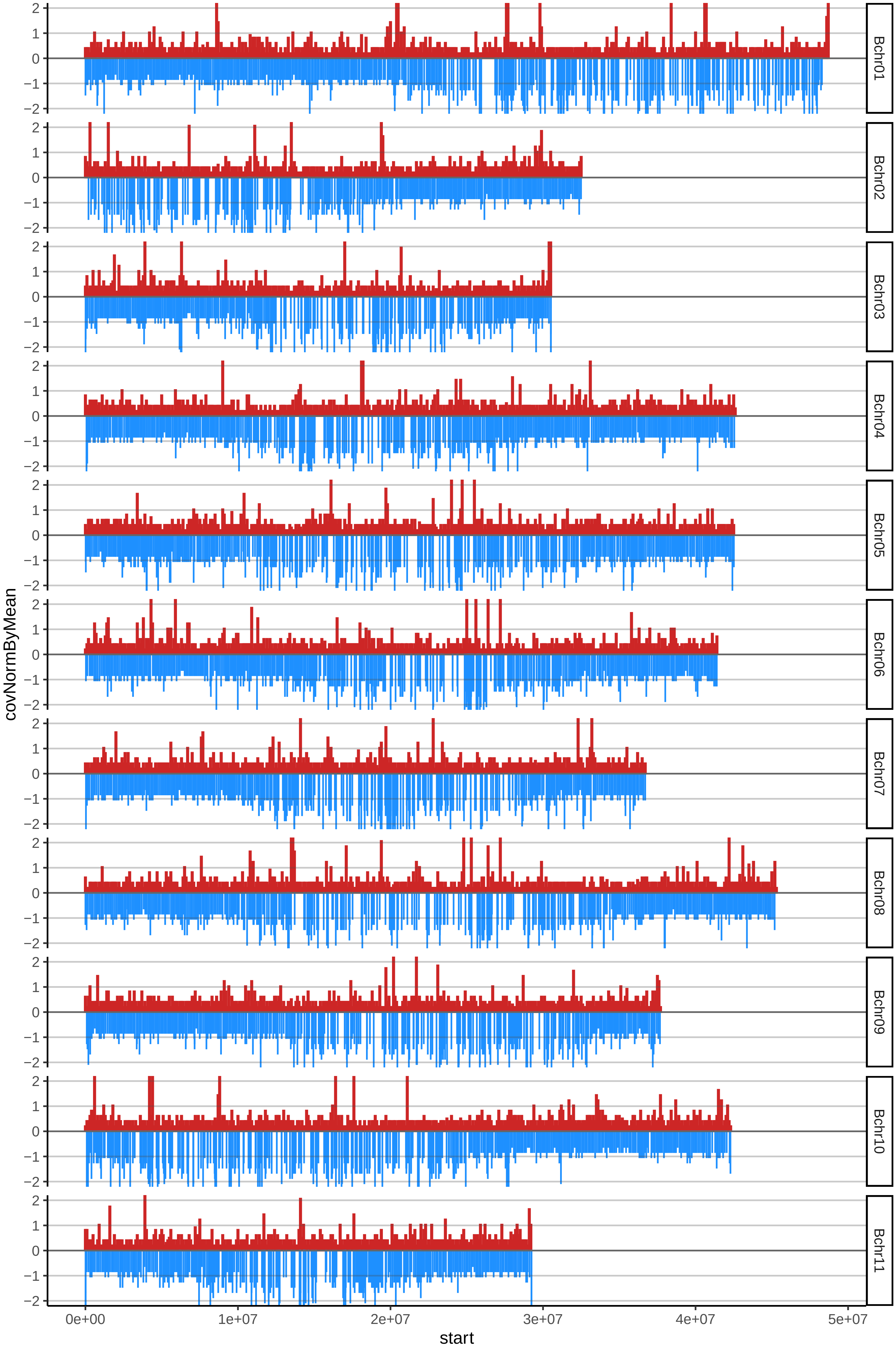

### 100LACATANcov_byA.pdf

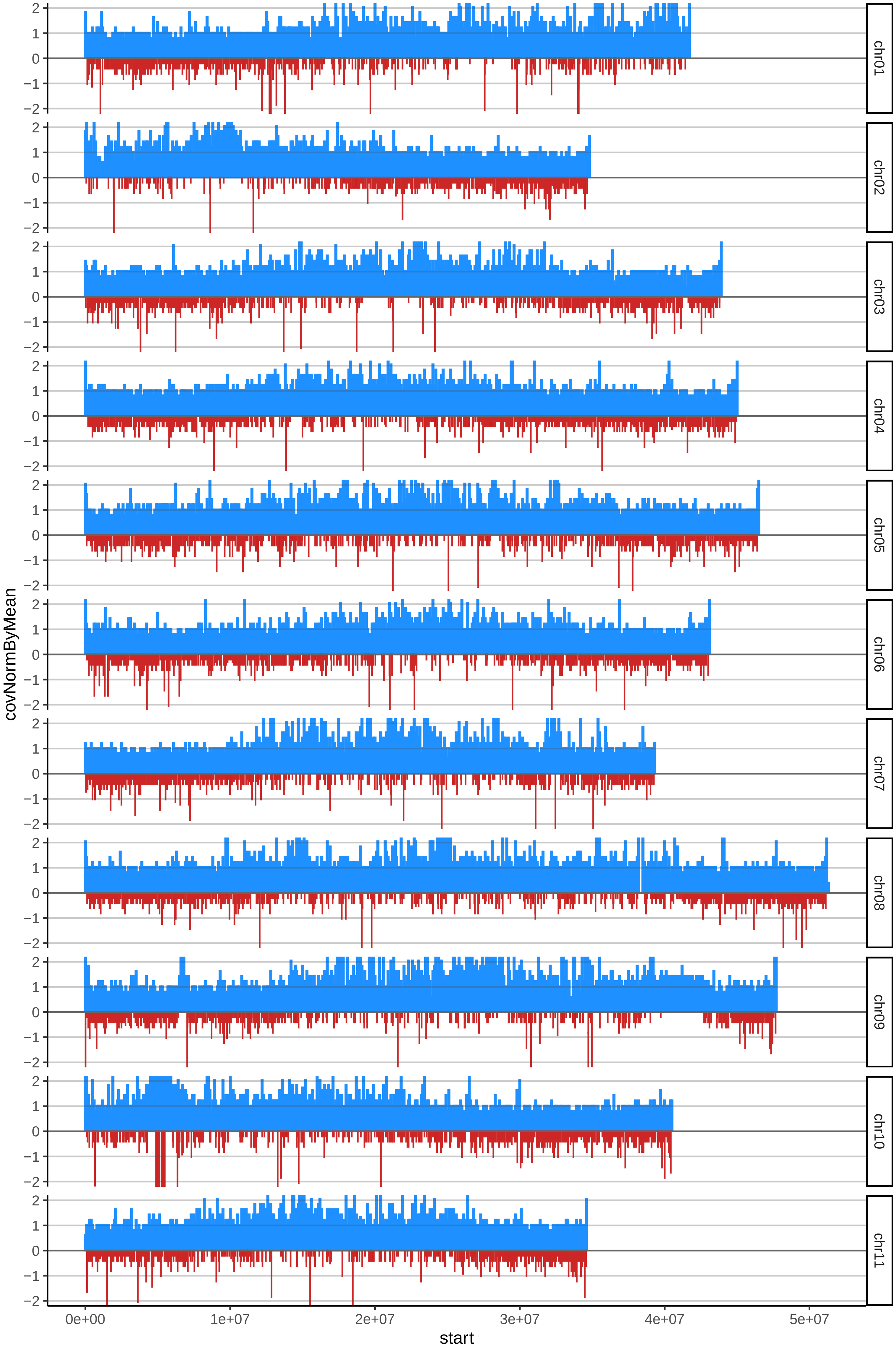

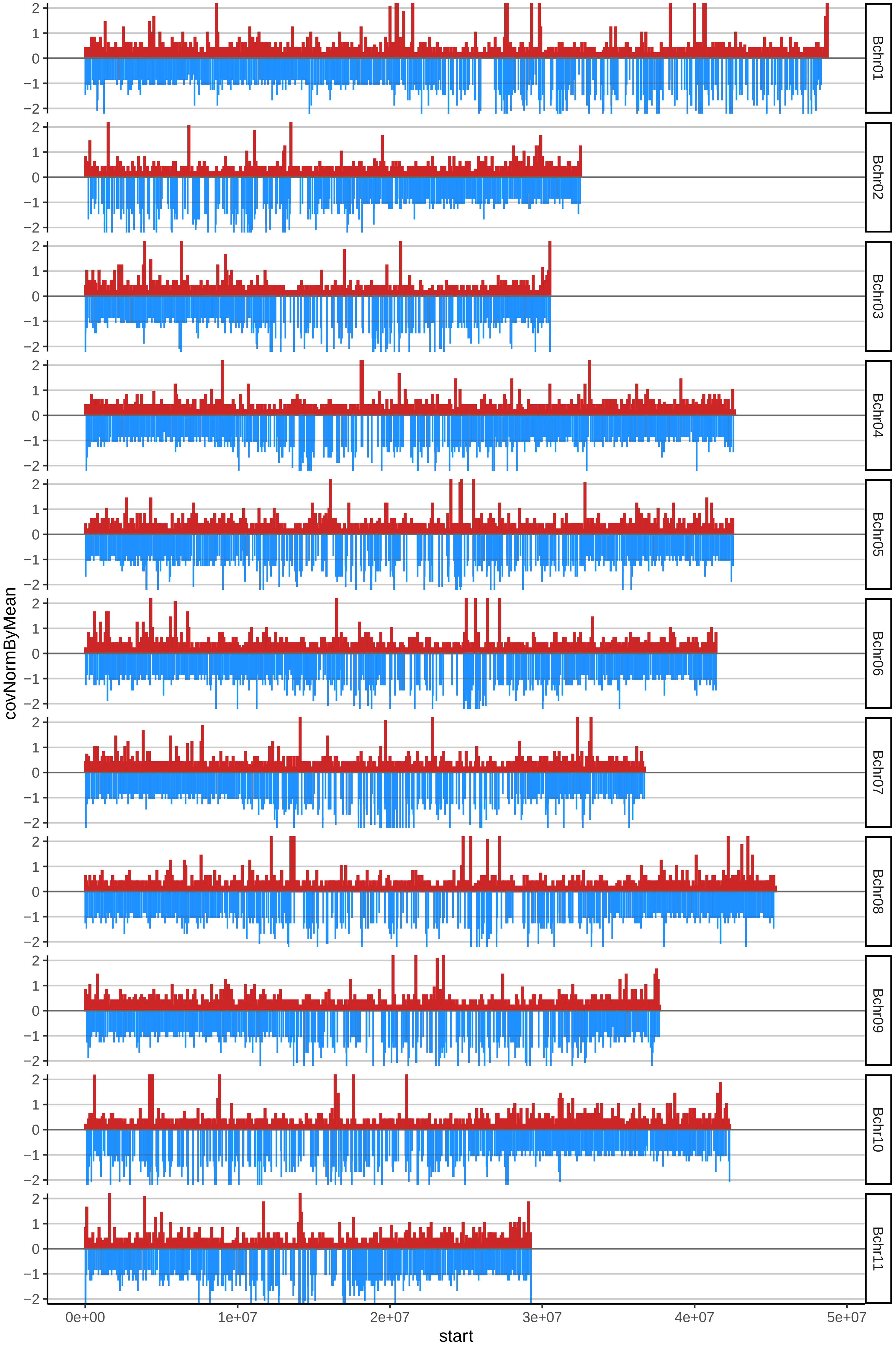

### 100LACATANcov_byB.pdf

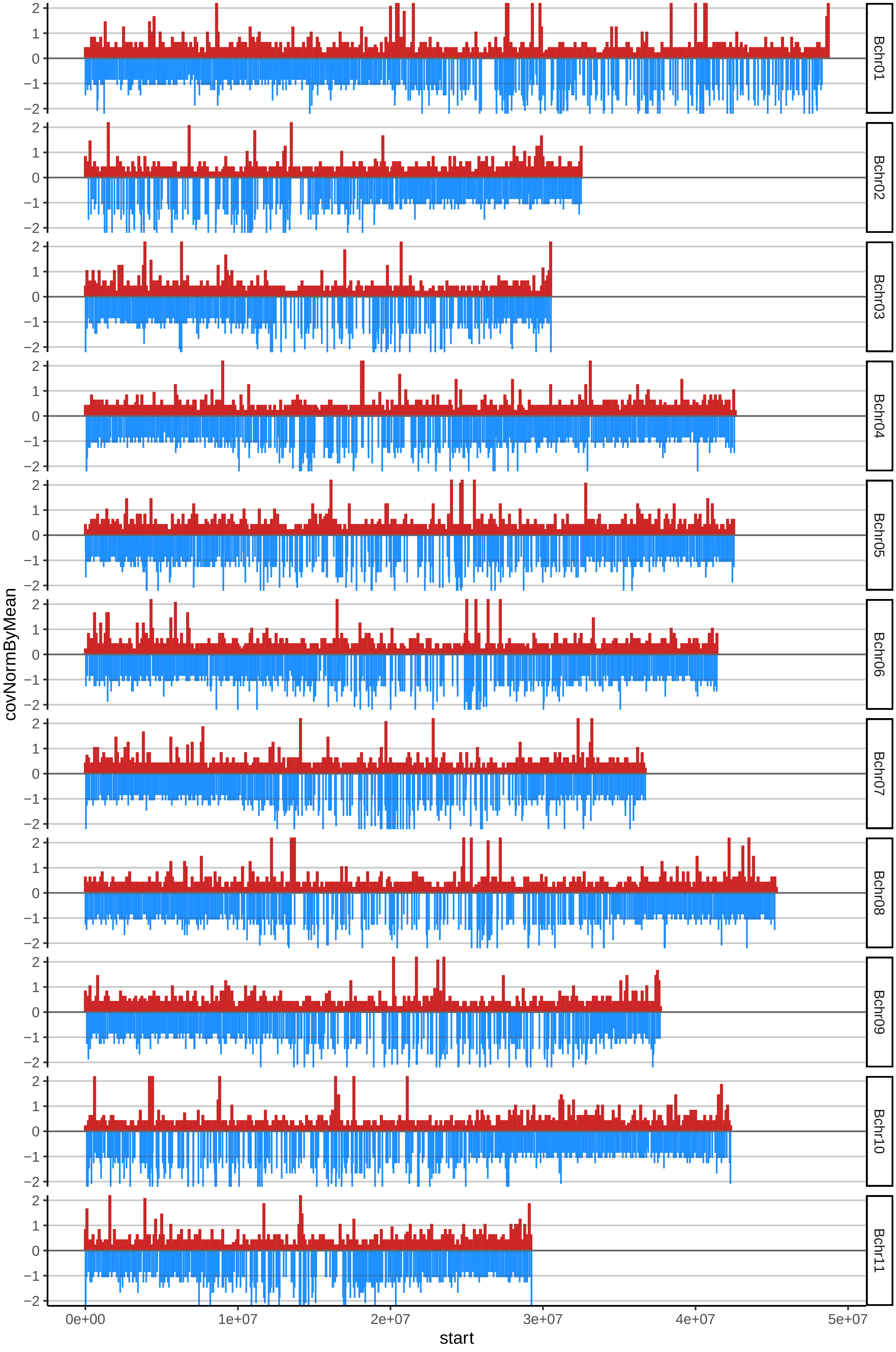

### 101SEREDOWcov_byA.pdf

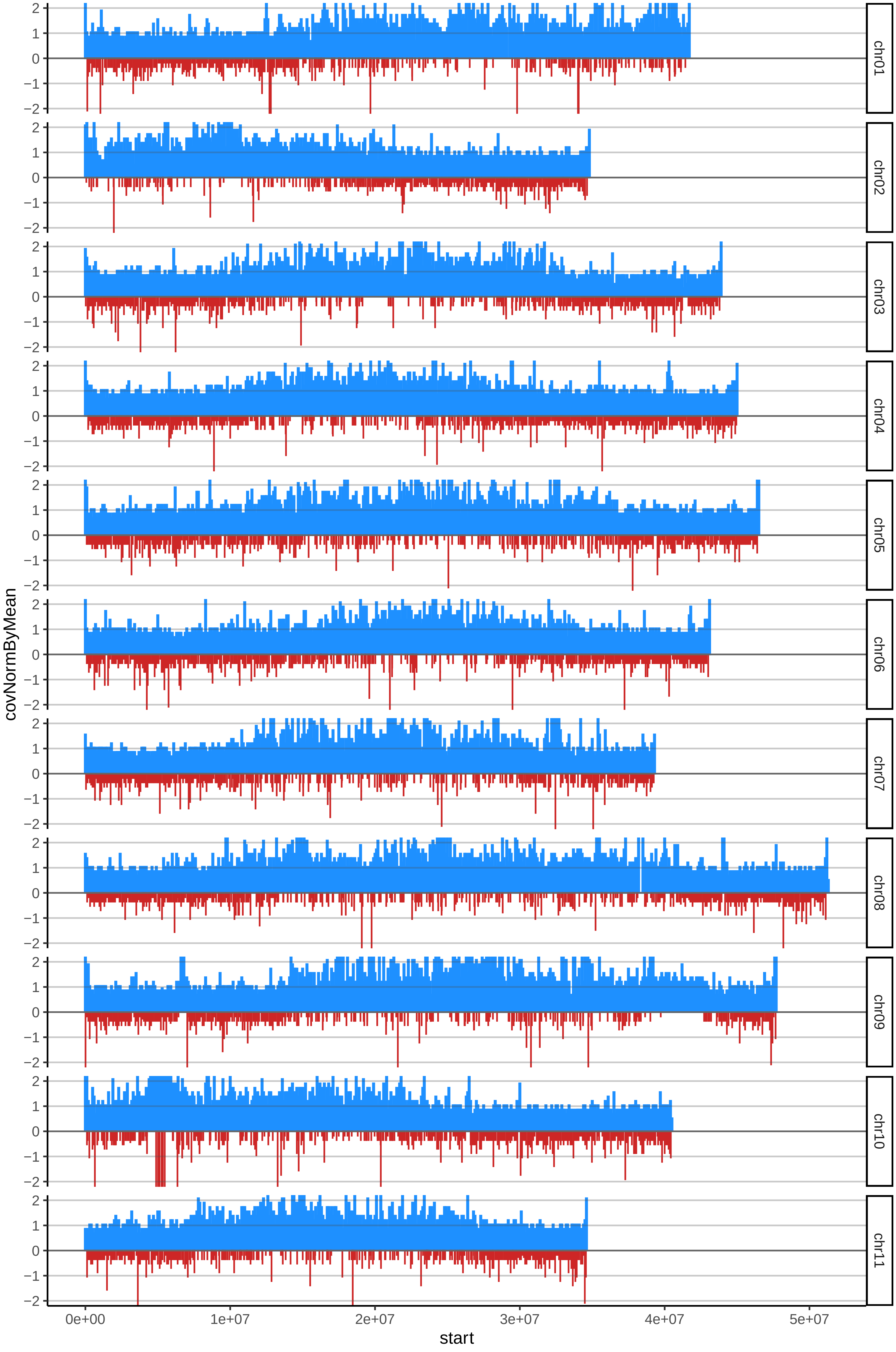

### 101SEREDOWcov_byB.pdf

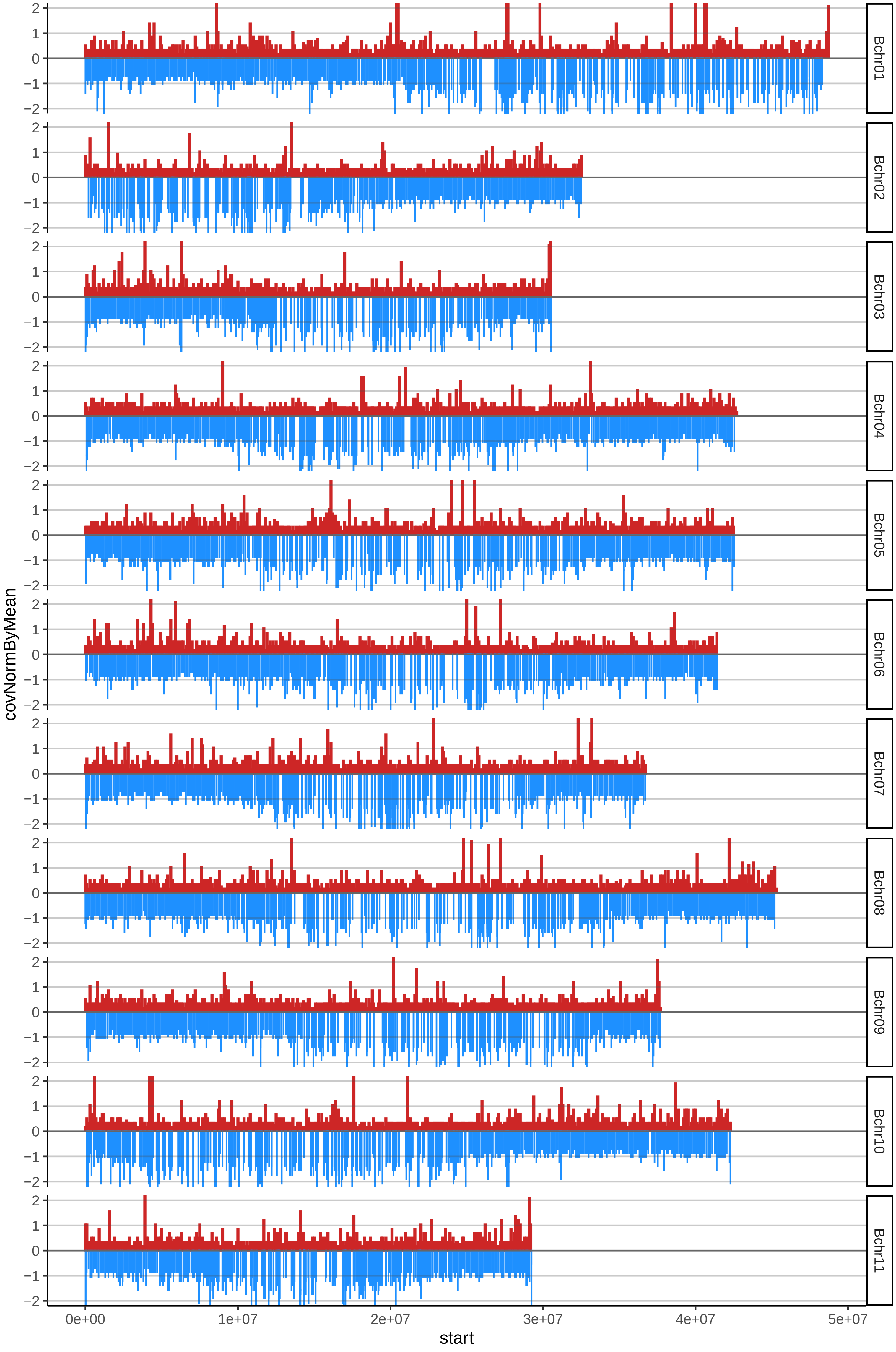

### 102GRANENANOcov_byA.pdf

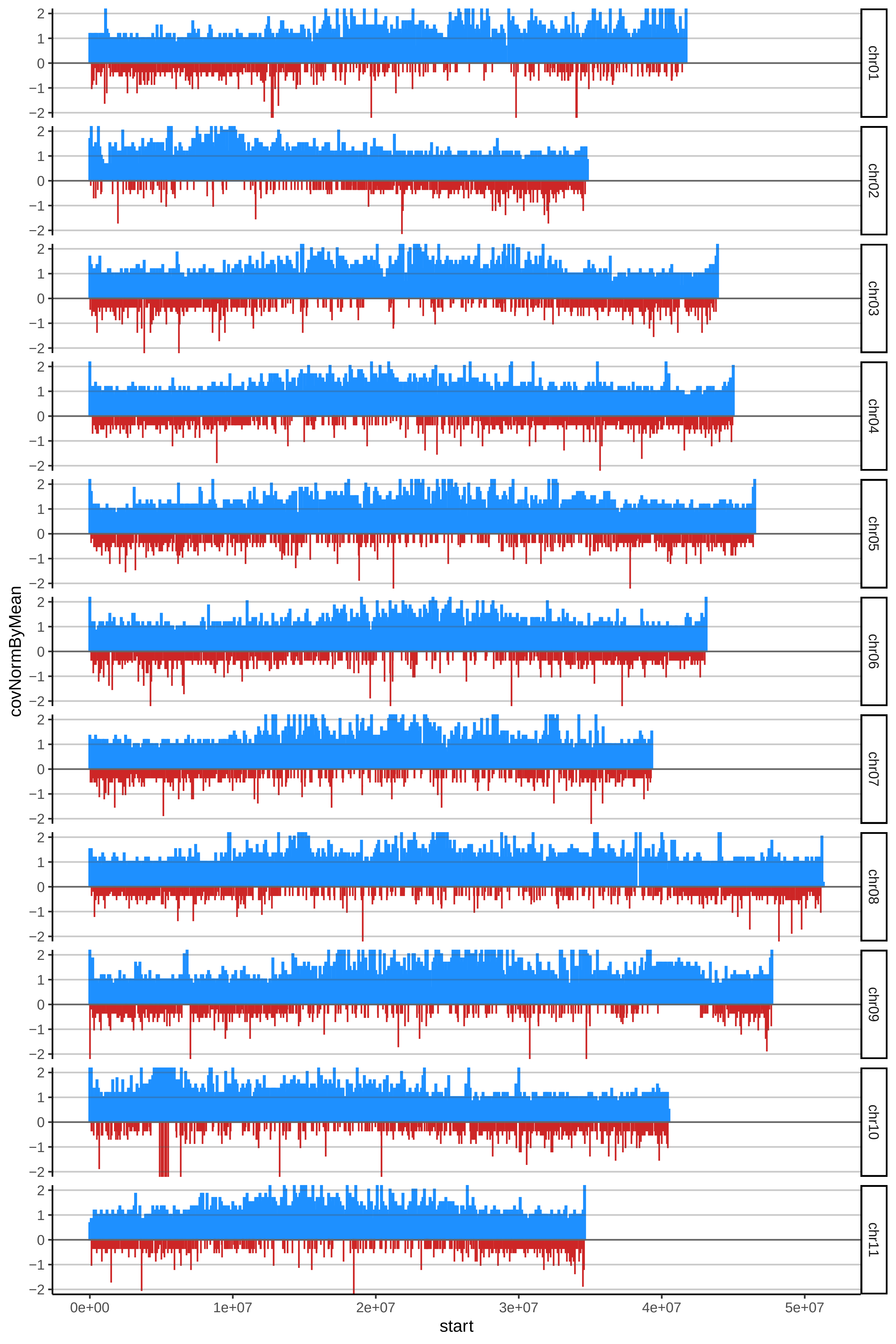

### 102GRANENANOcov_byB.pdf

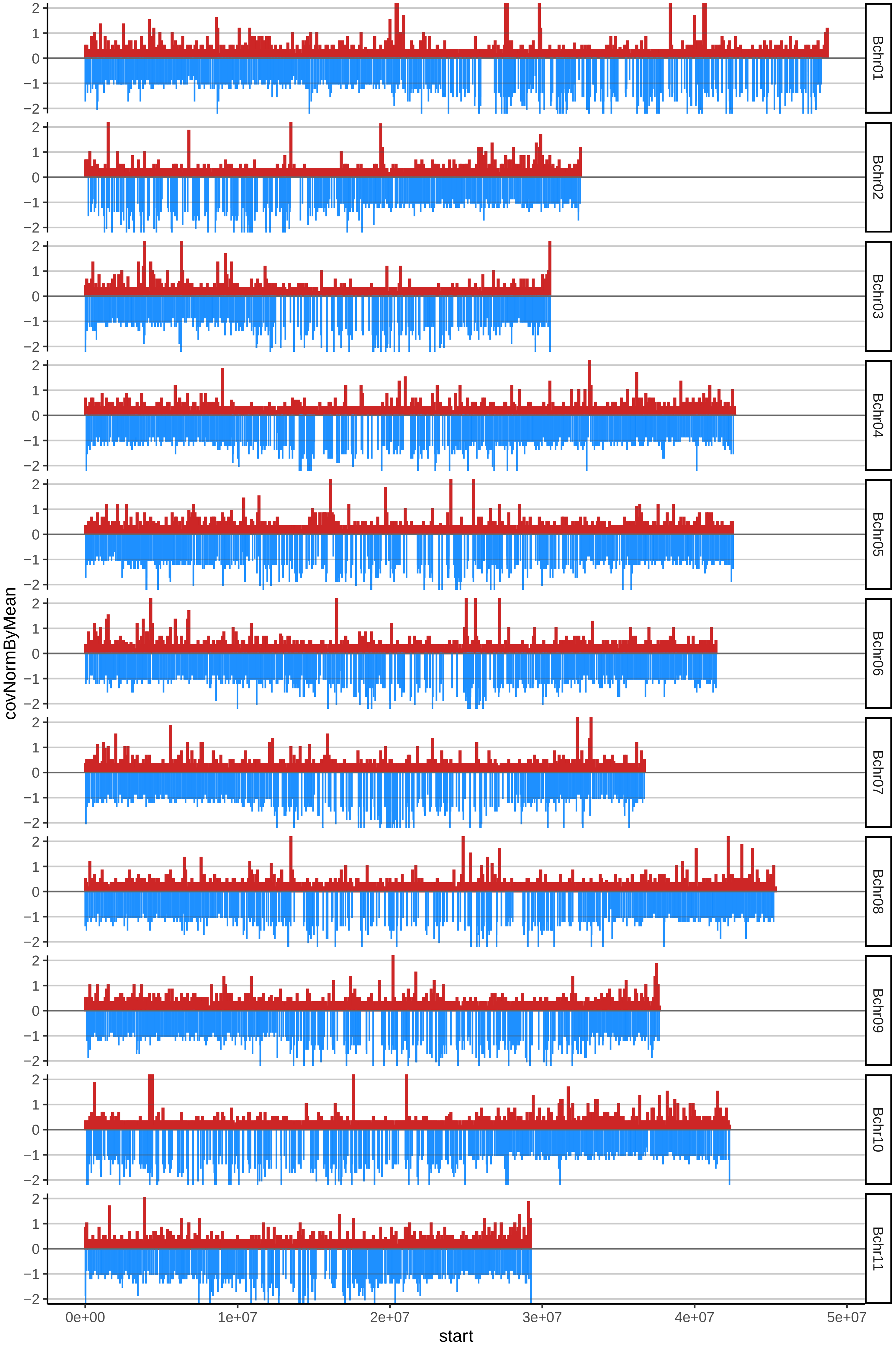

### 103PIGMEOcov_byA.pdf

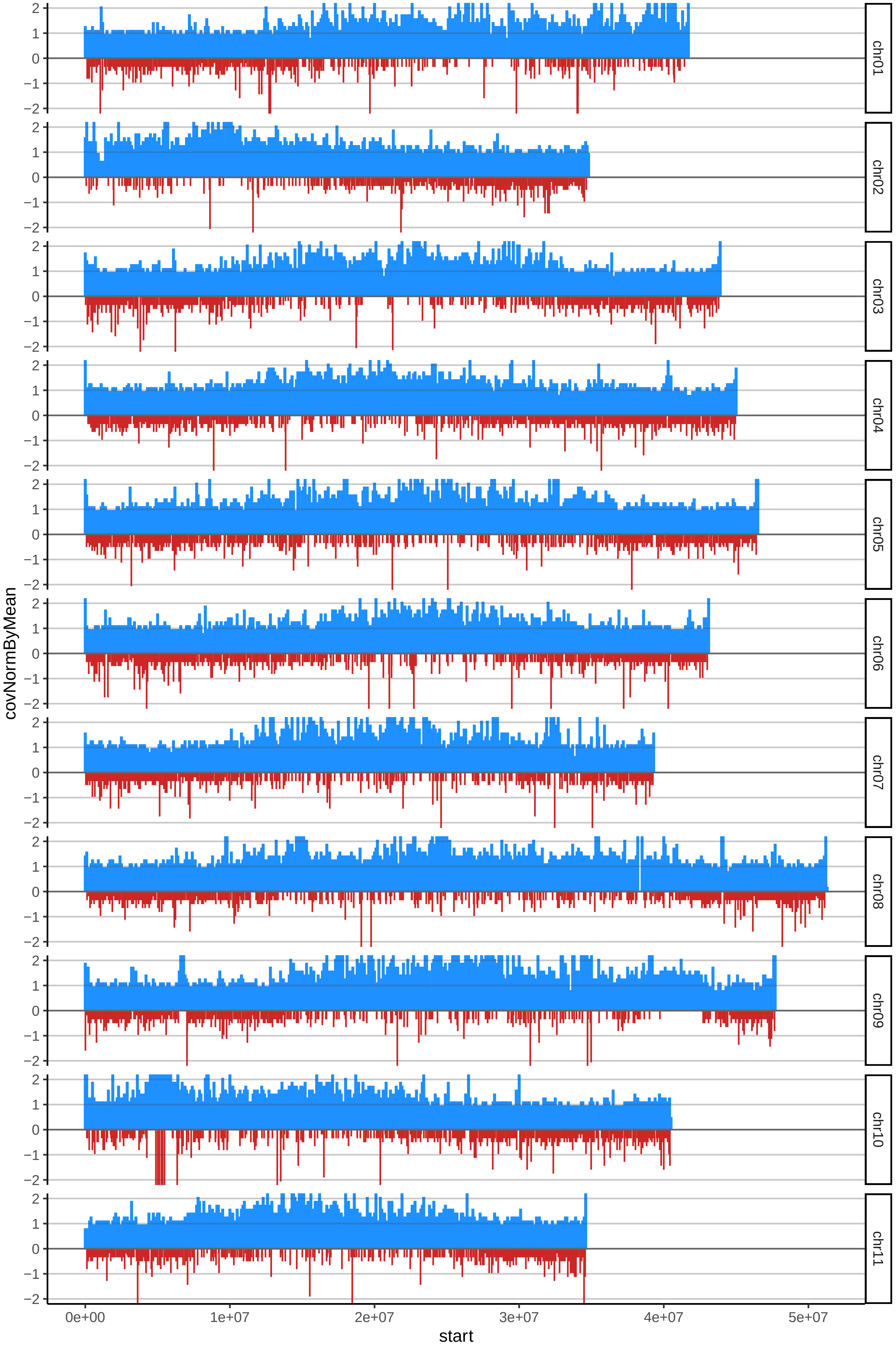

### 103PIGMEOcov_byB.pdf

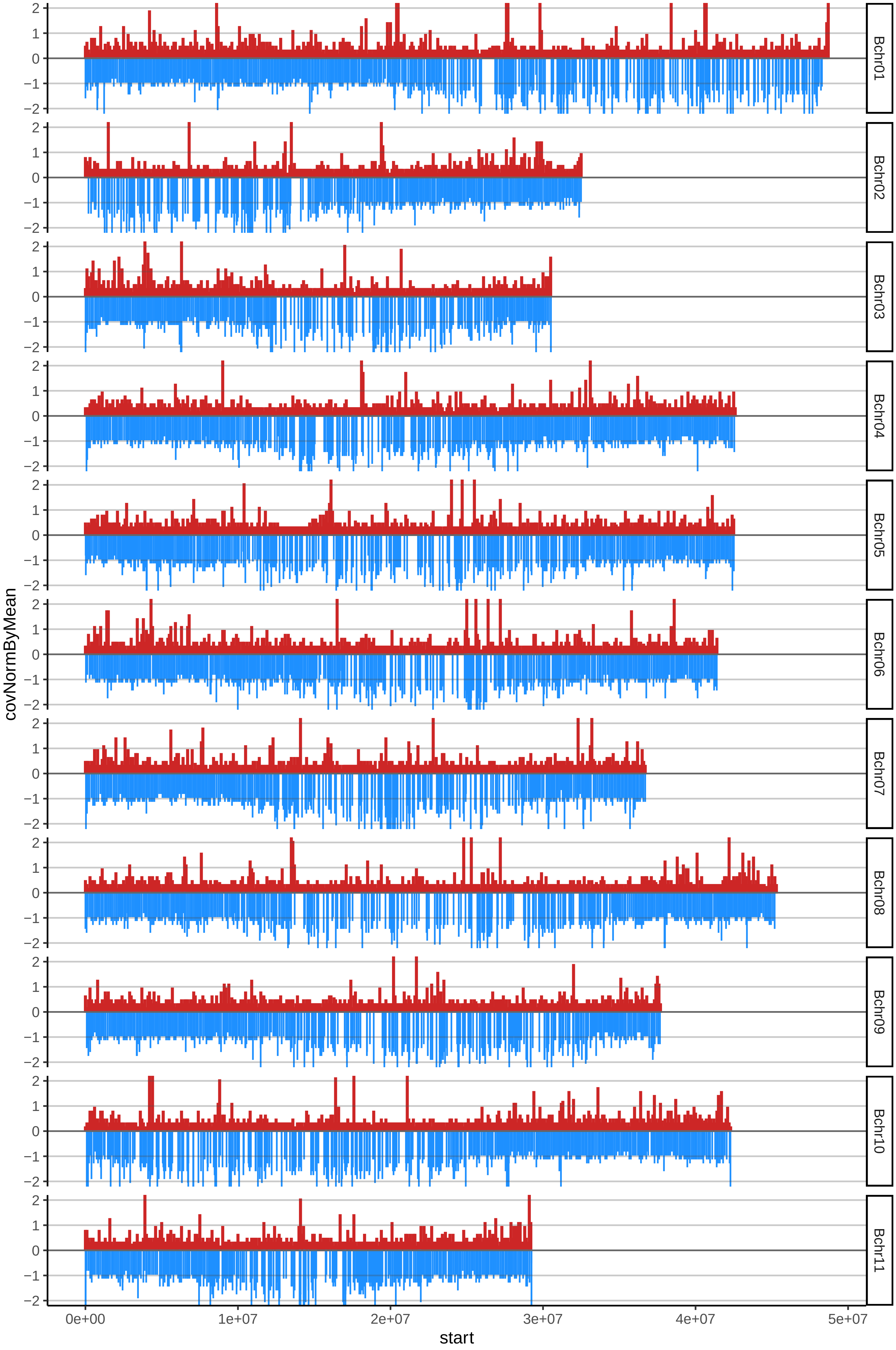

### 104POYOcov_byA.pdf

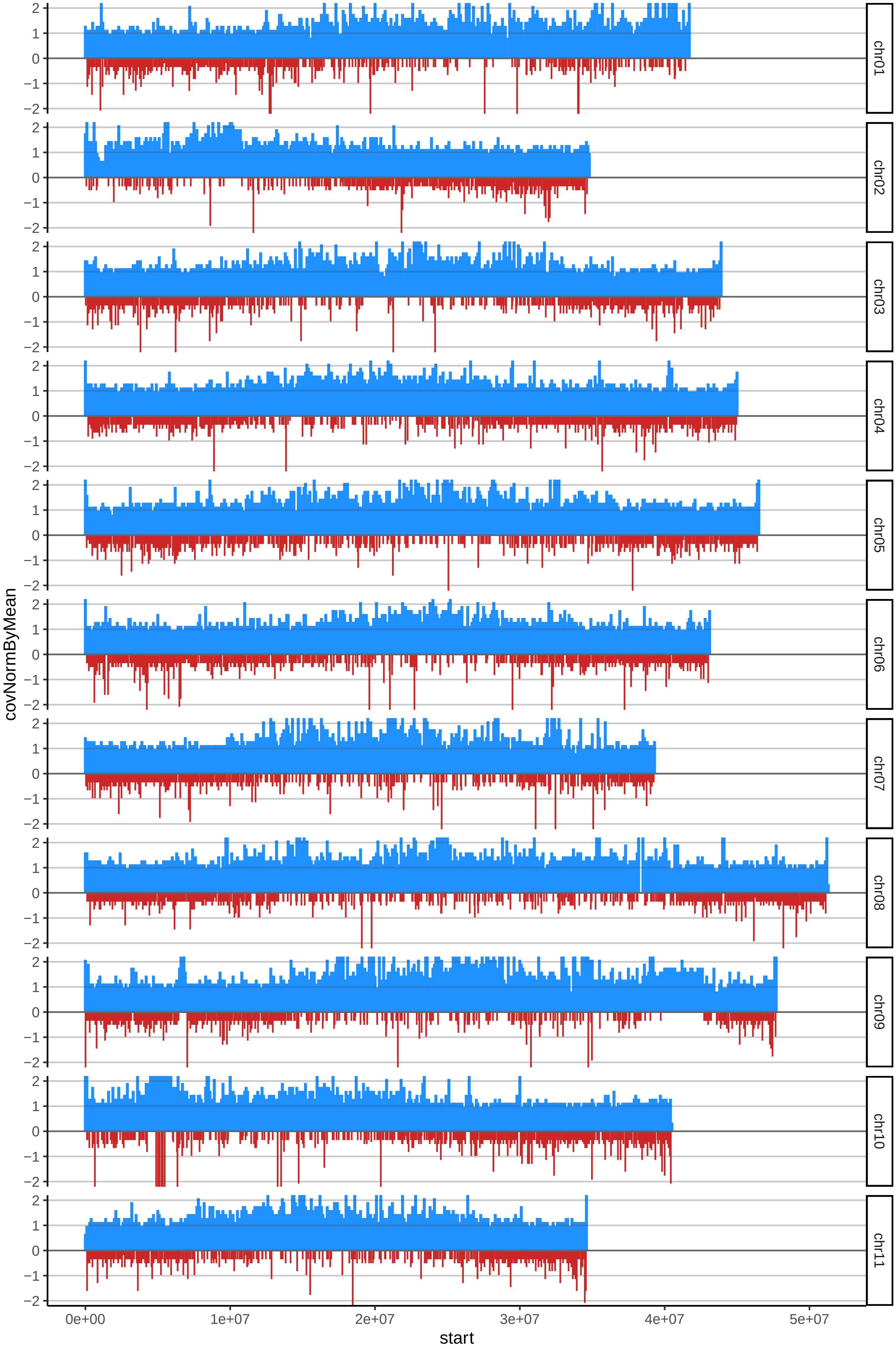

### 104POYOcov_byB.pdf

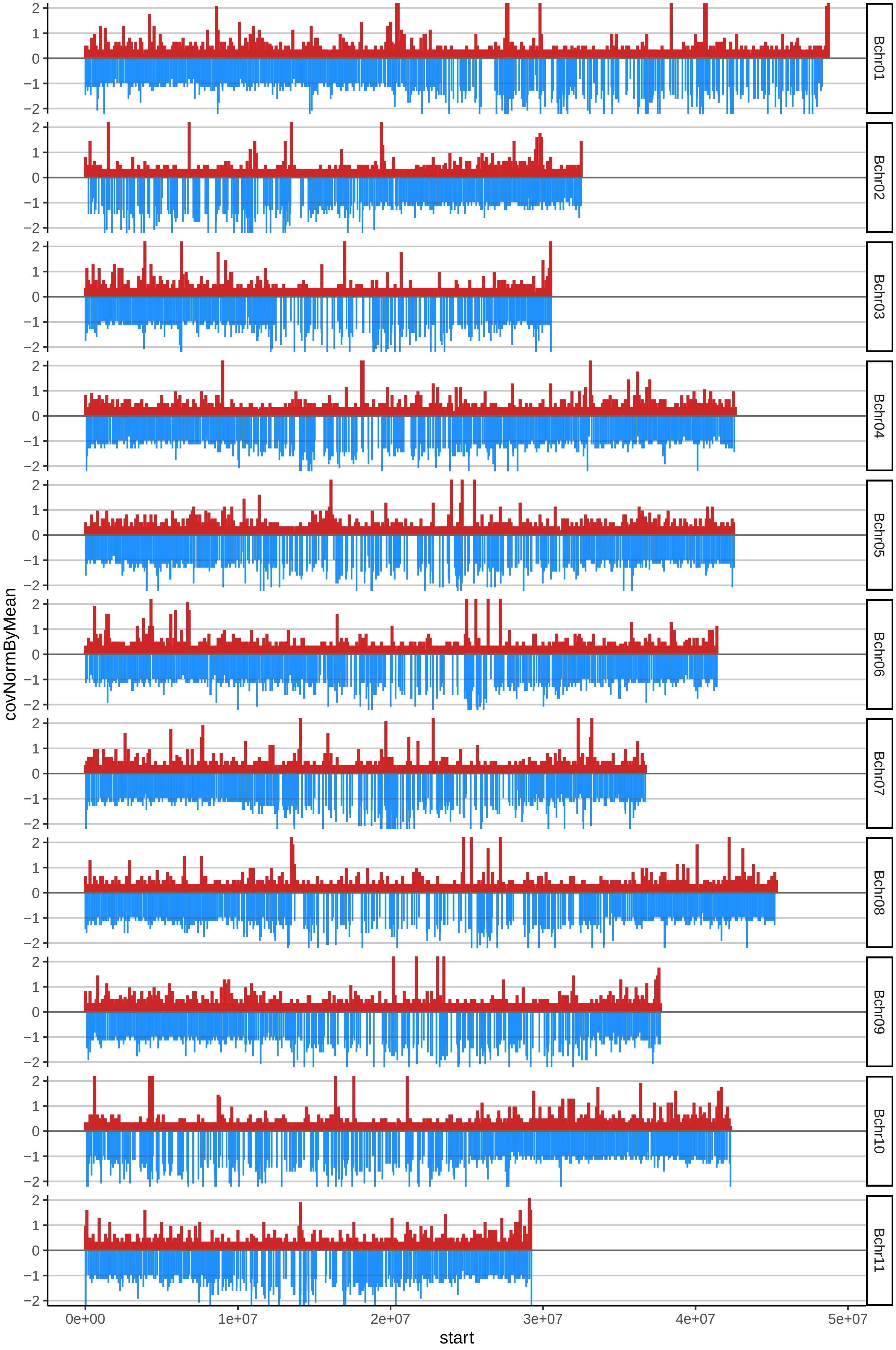

### 105VALERYcov_byA.pdf

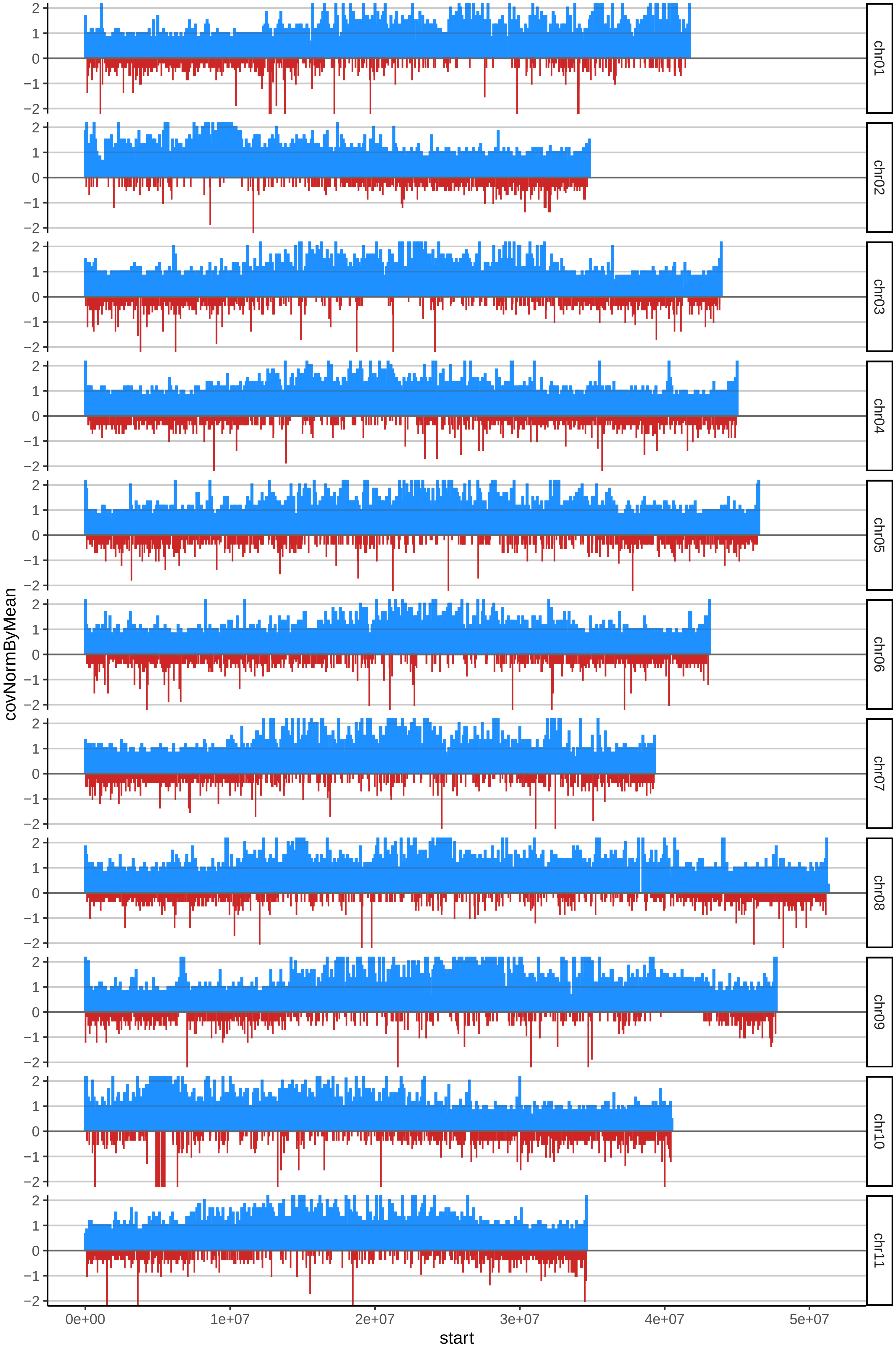

### 105VALERYcov_byB.pdf

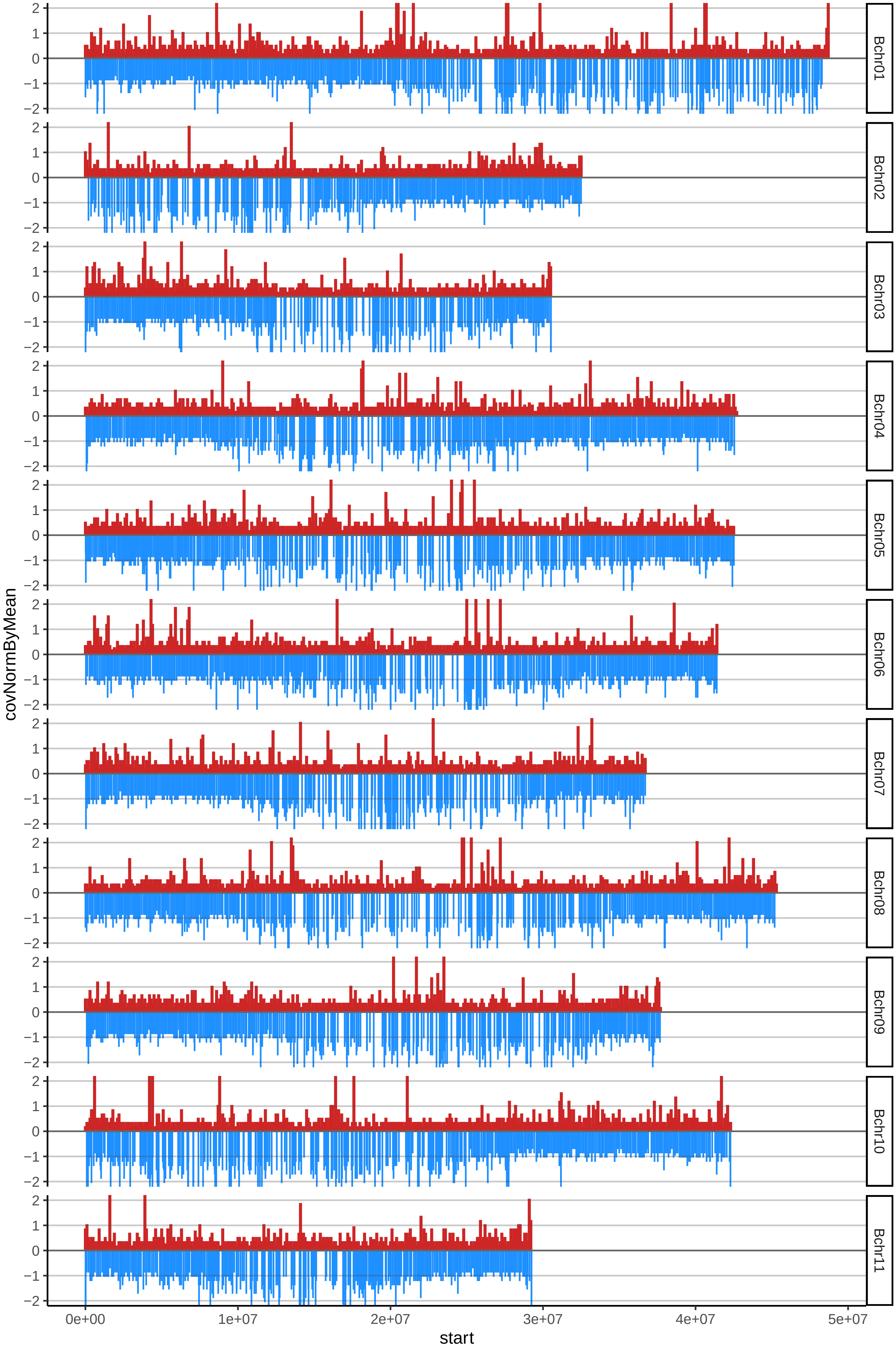

### 106BANANOSINCLASIFICARcov_byA.pdf

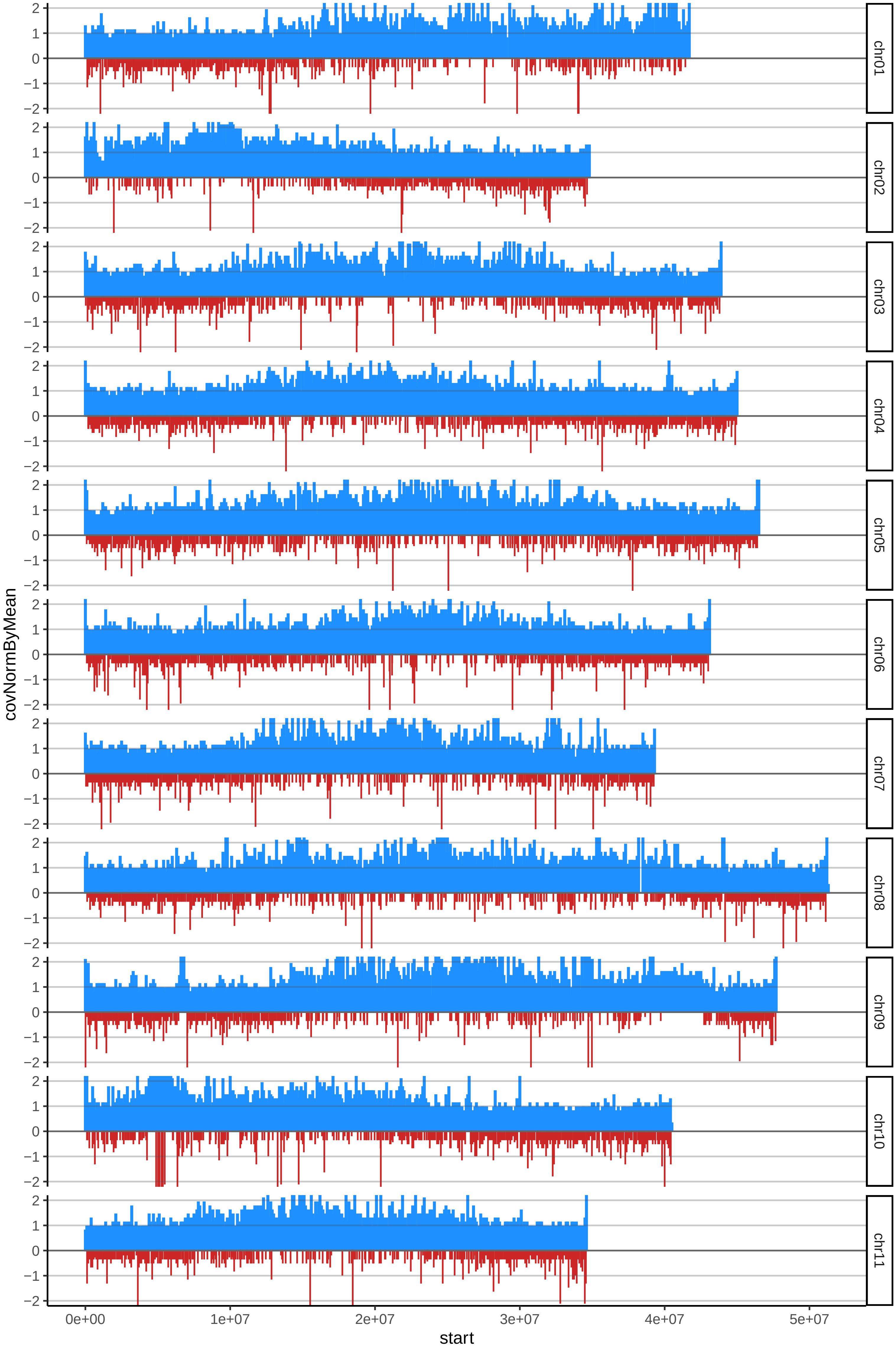

### 106BANANOSINCLASIFICARcov_byB.pdf

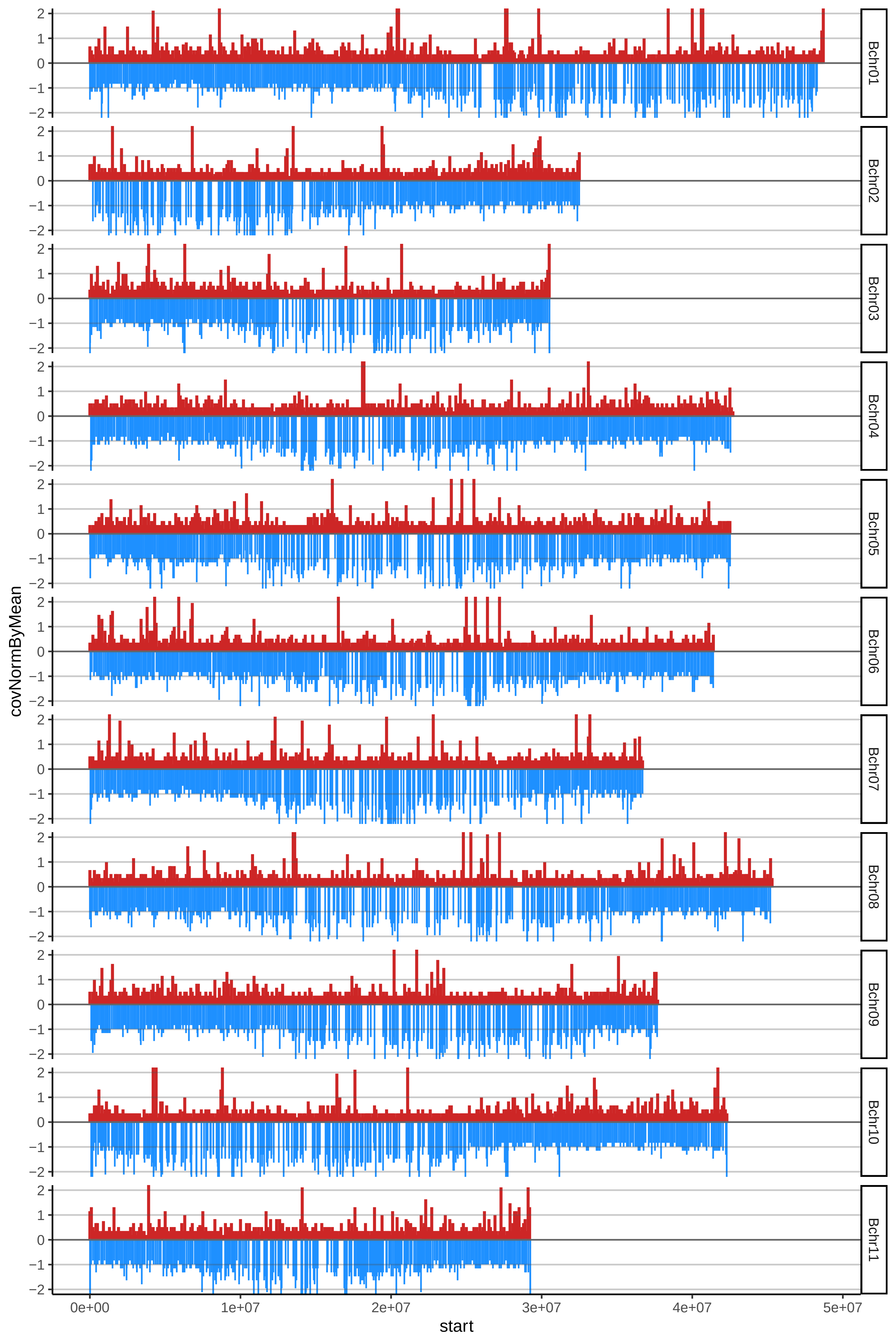

### 107DWARFCAVENDISHcov_byA.pdf

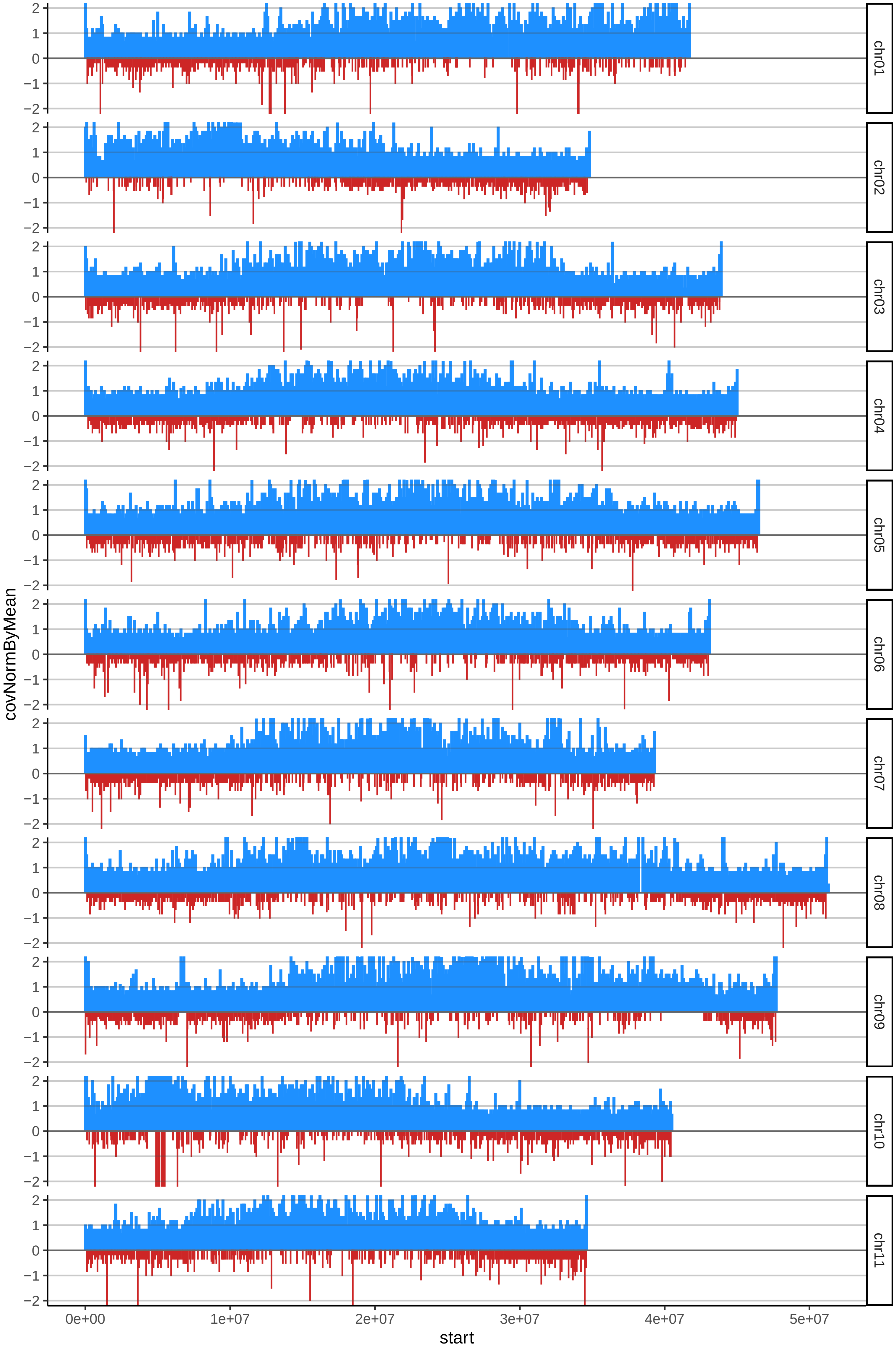

### 107DWARFCAVENDISHcov_byB.pdf

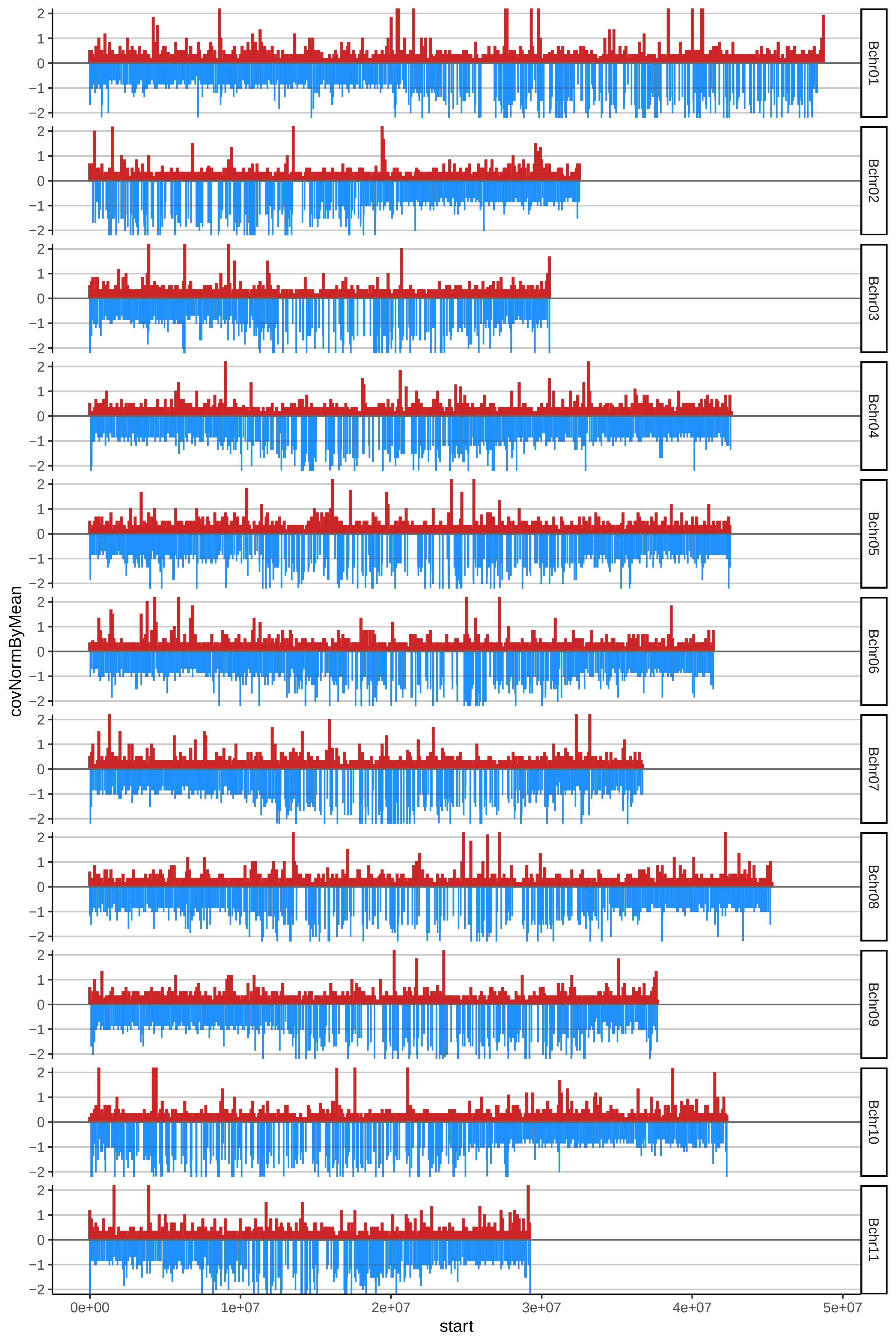

### 113BANANO2cov_byA.pdf

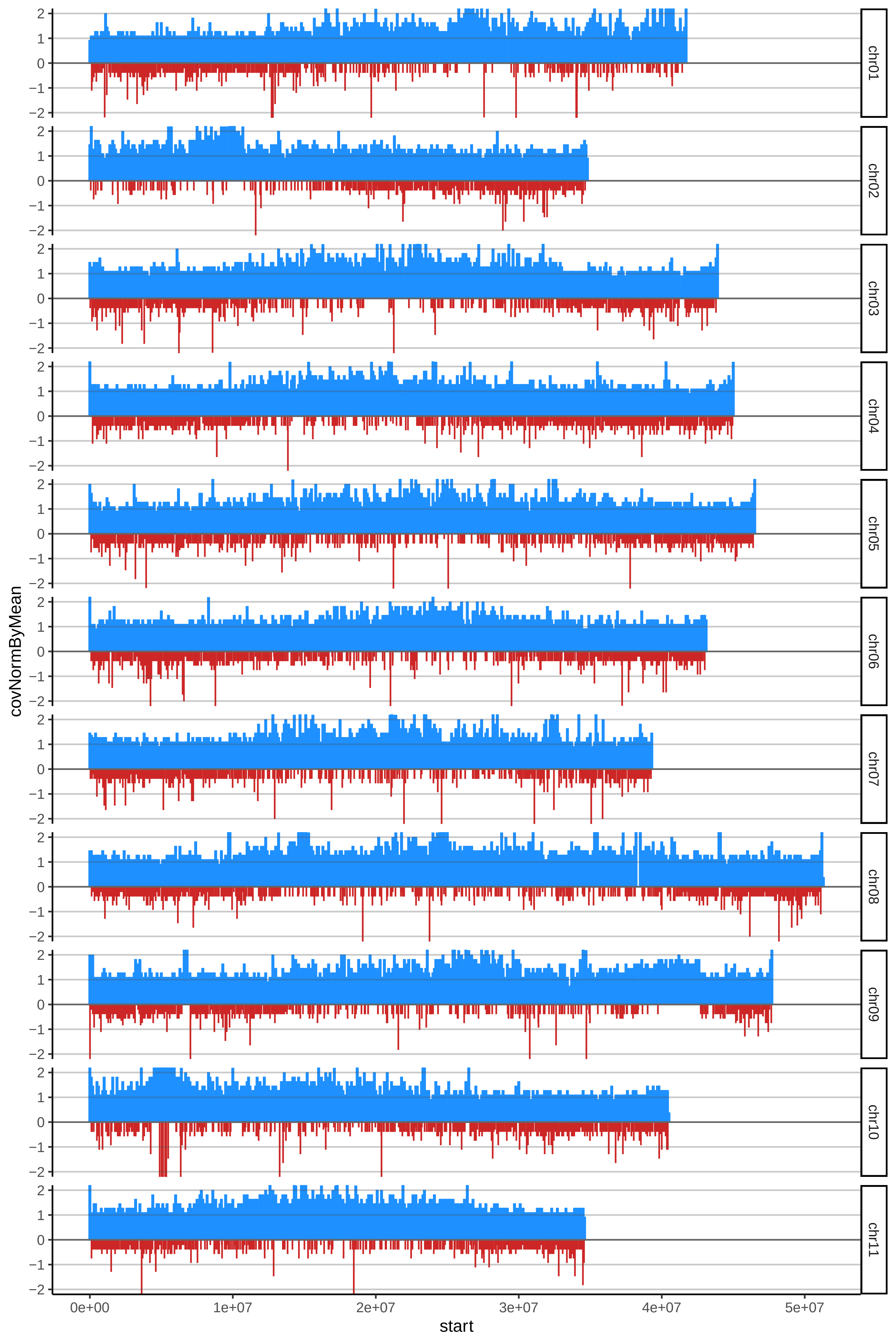

### 113BANANO2cov_byB.pdf

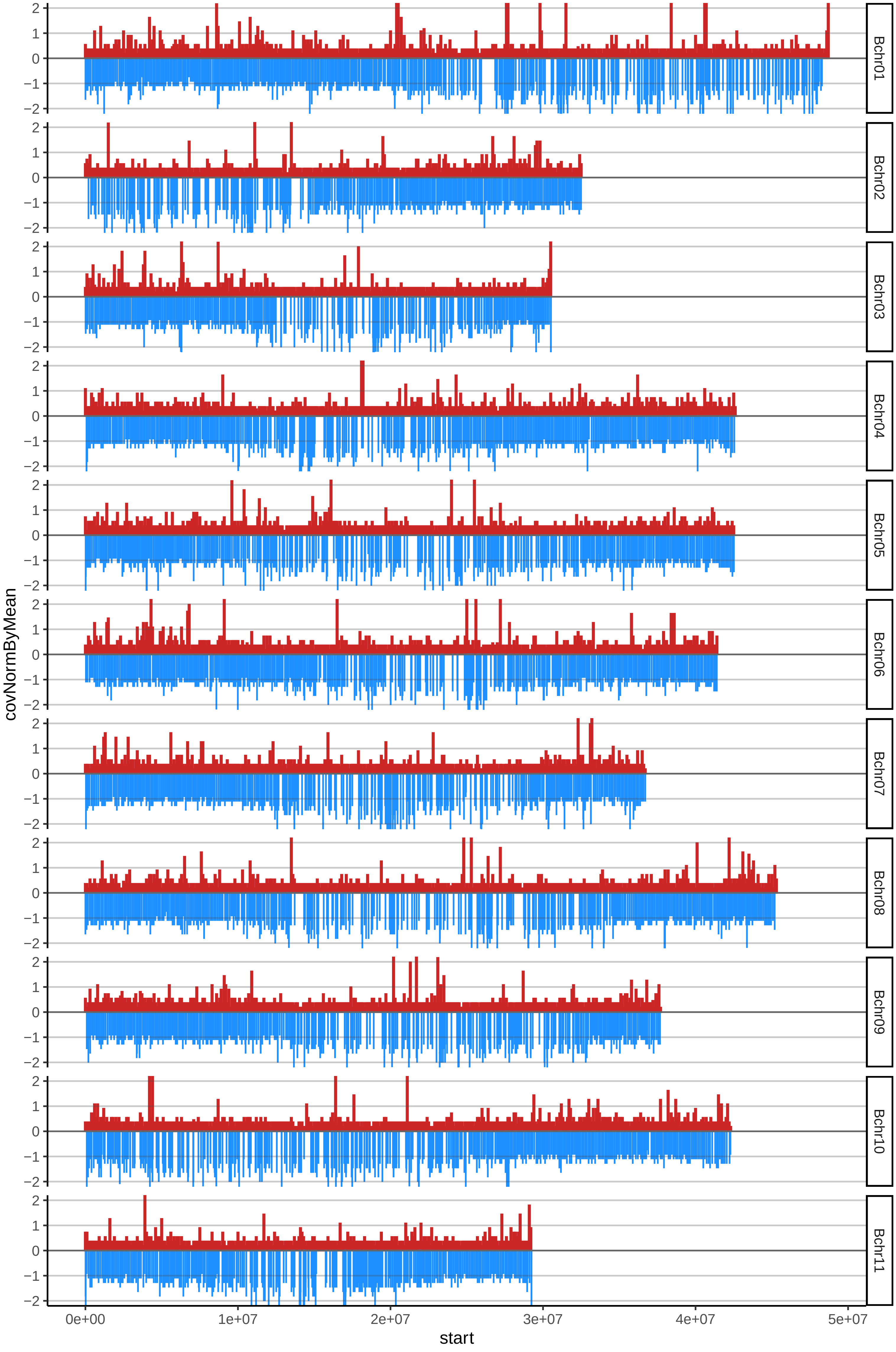

### 116GUINEONEGROcov_byA.pdf

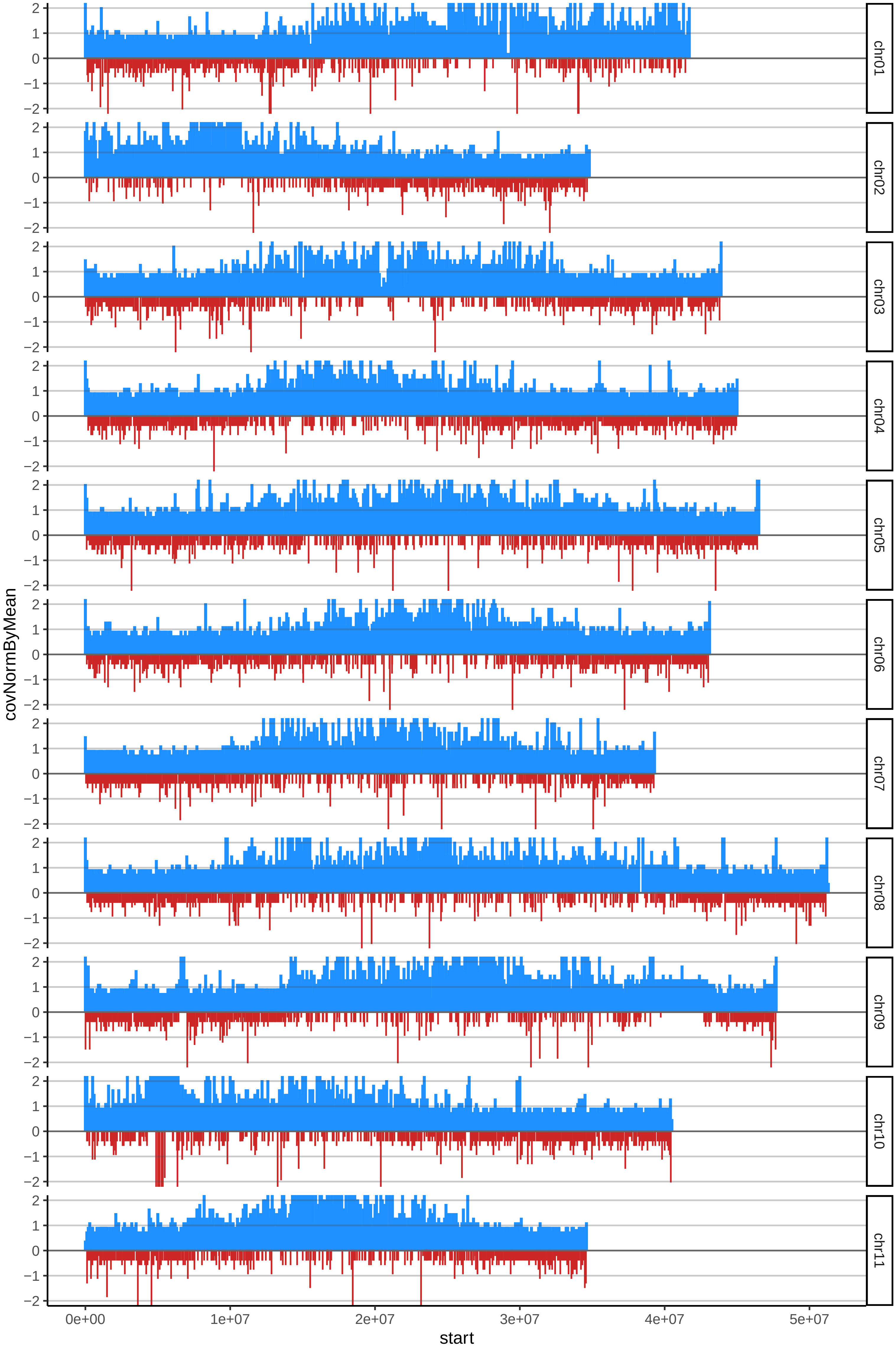

### 116GUINEONEGROcov_byB.pdf

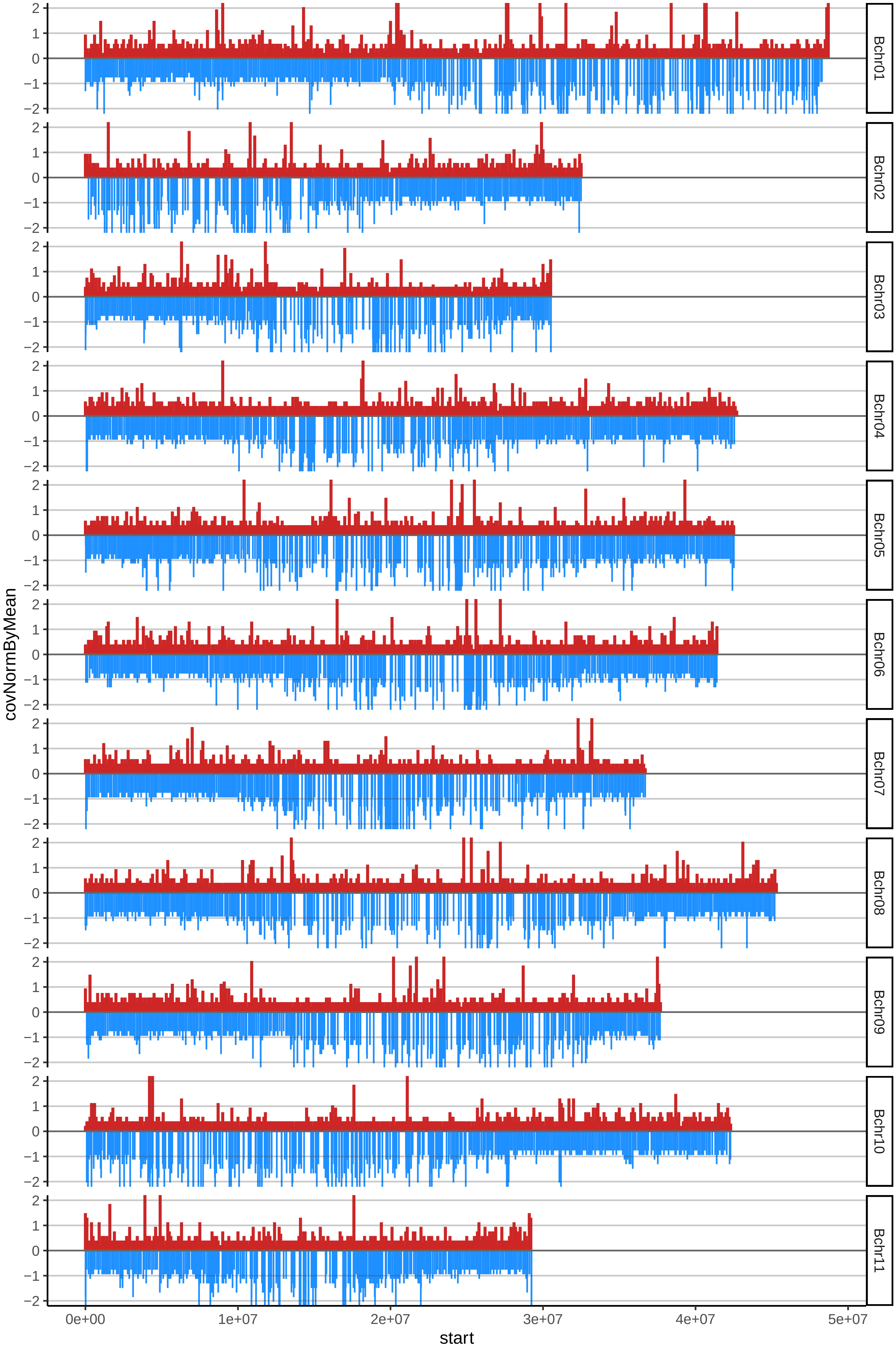

### 117NAKITENGWAcov_byA.pdf

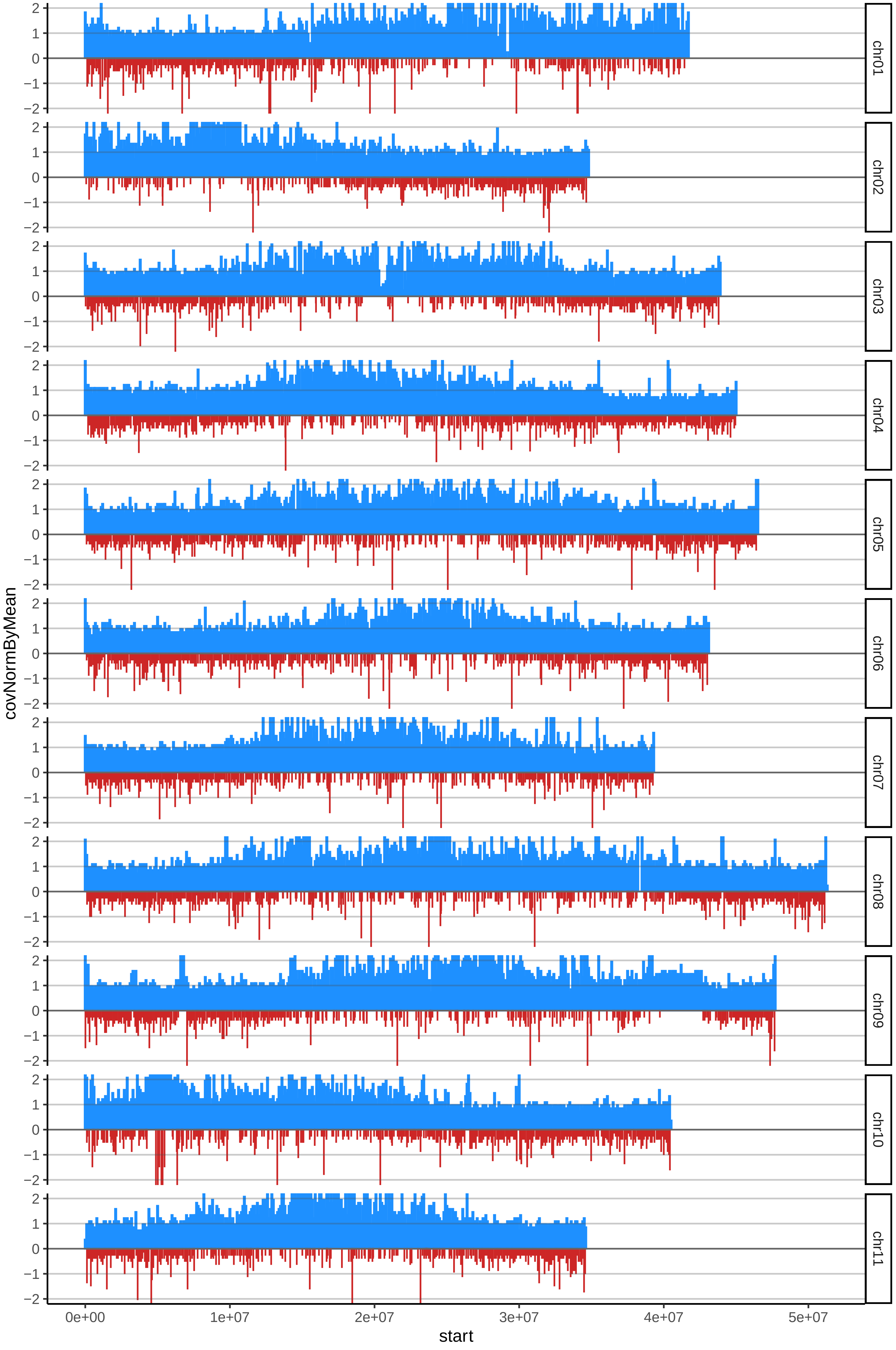

### 117NAKITENGWAcov_byB.pdf

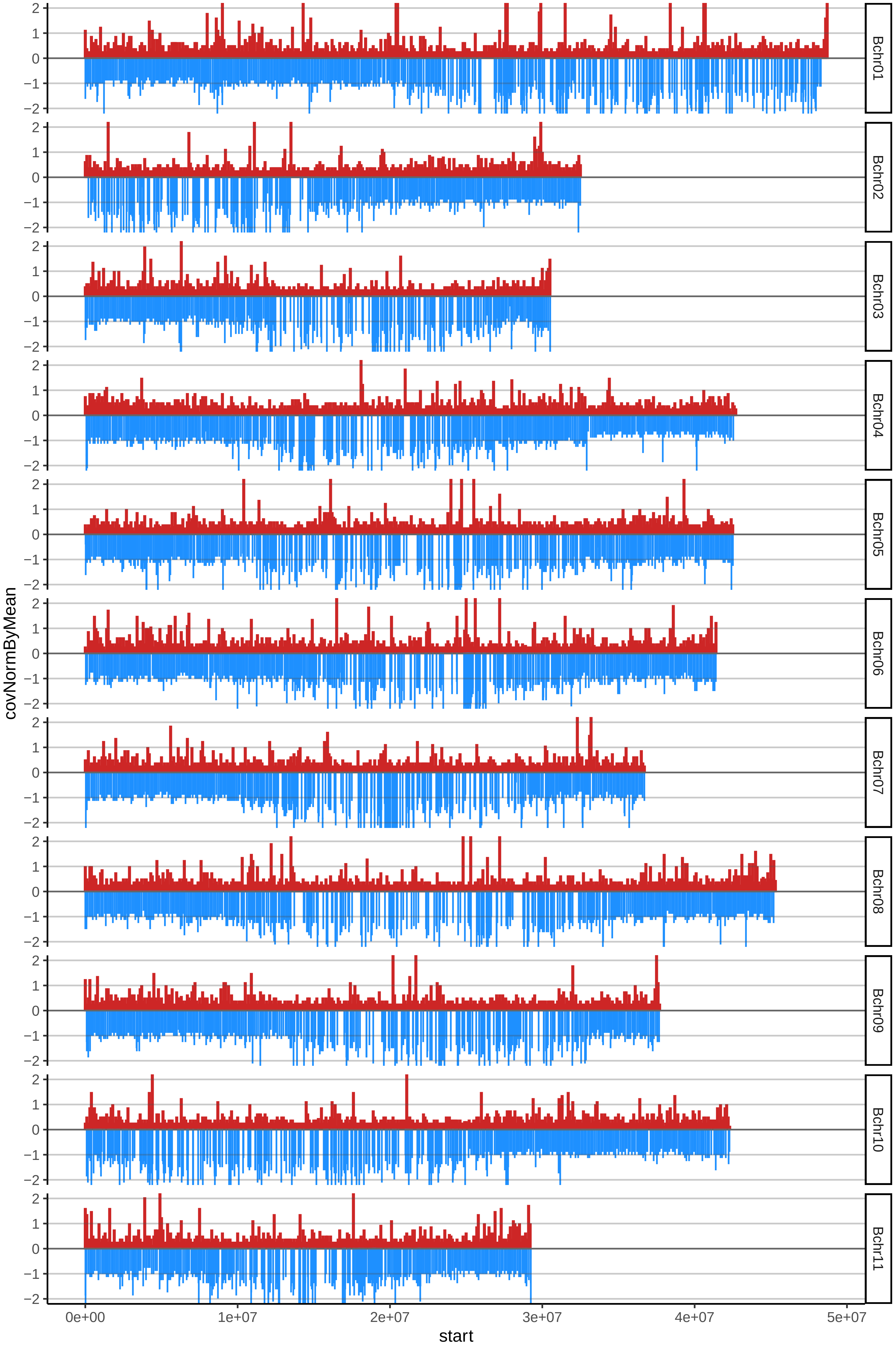

### 118IGITSIRIINTUNTUcov_byA.pdf

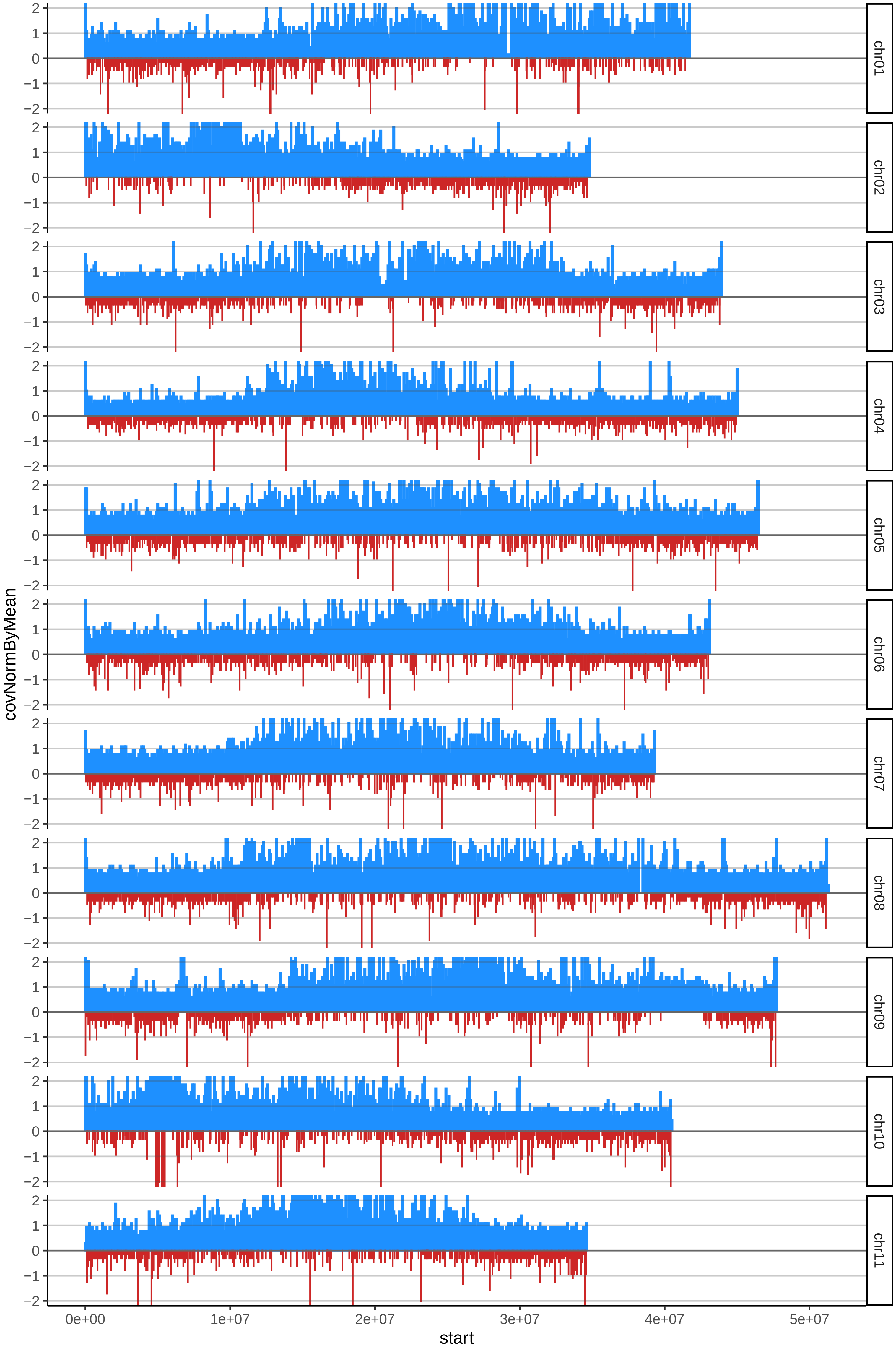

### 118IGITSIRIINTUNTUcov_byB.pdf

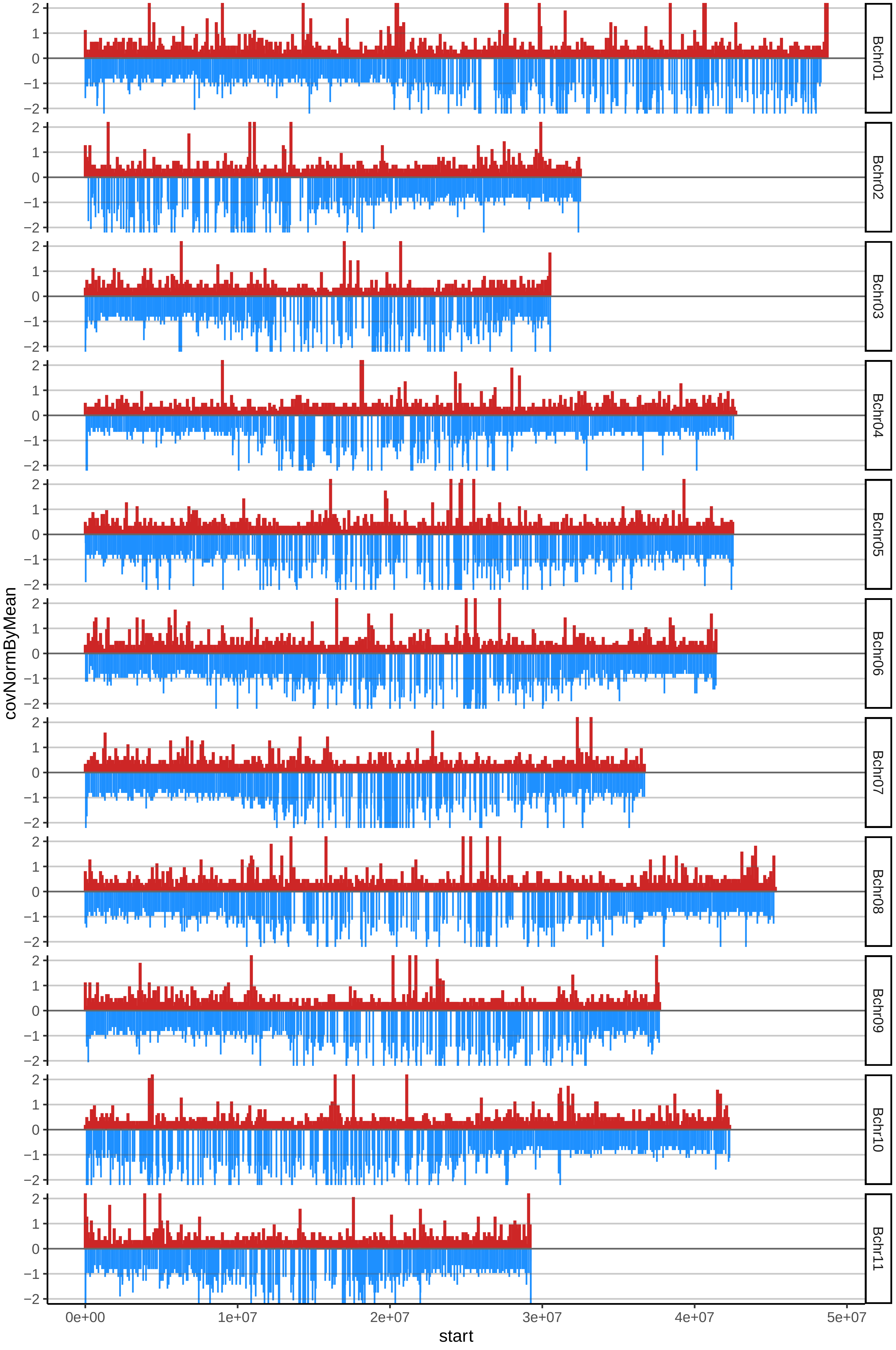

### 127IITAcov_byA.pdf

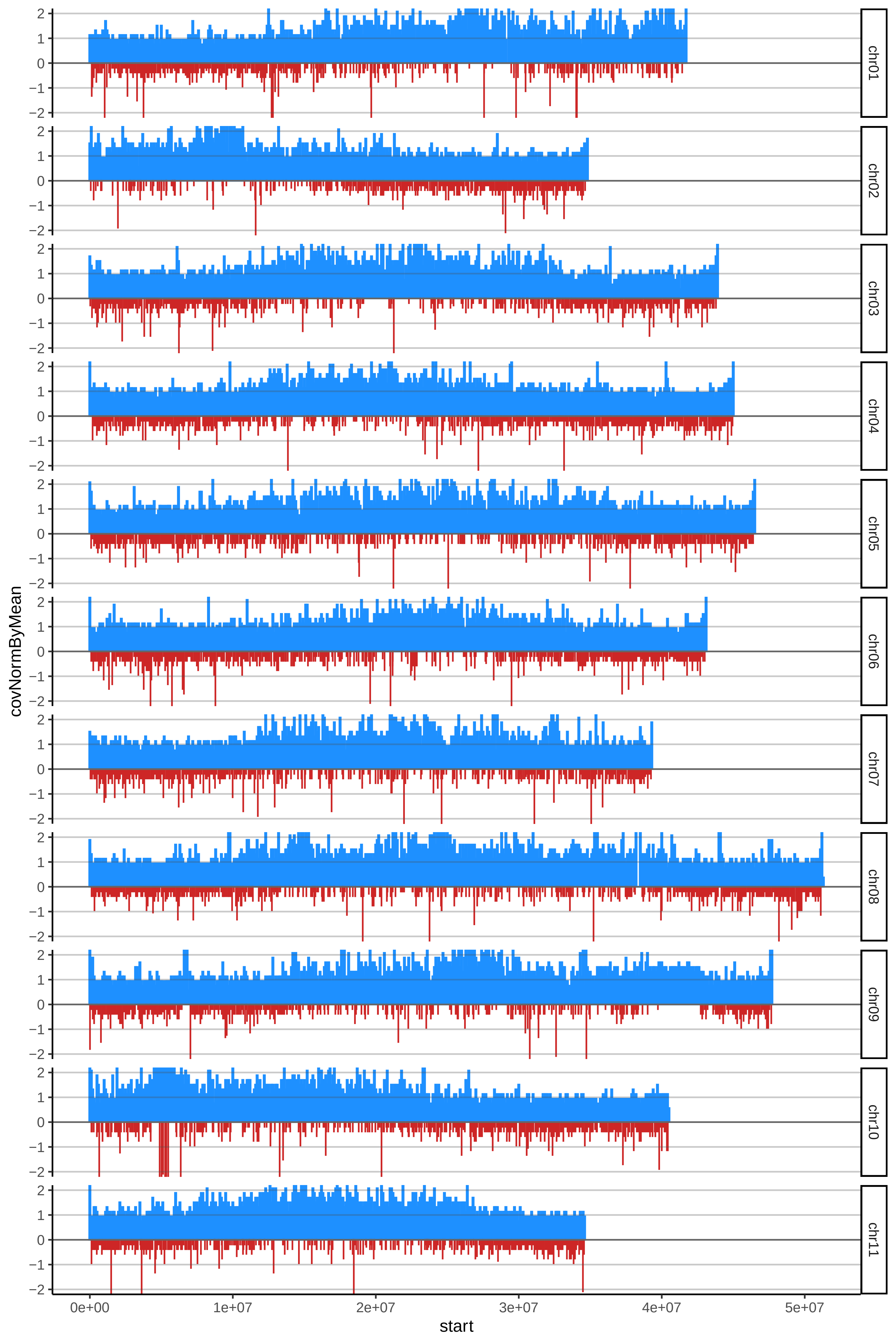

### 127IITAcov_byB.pdf

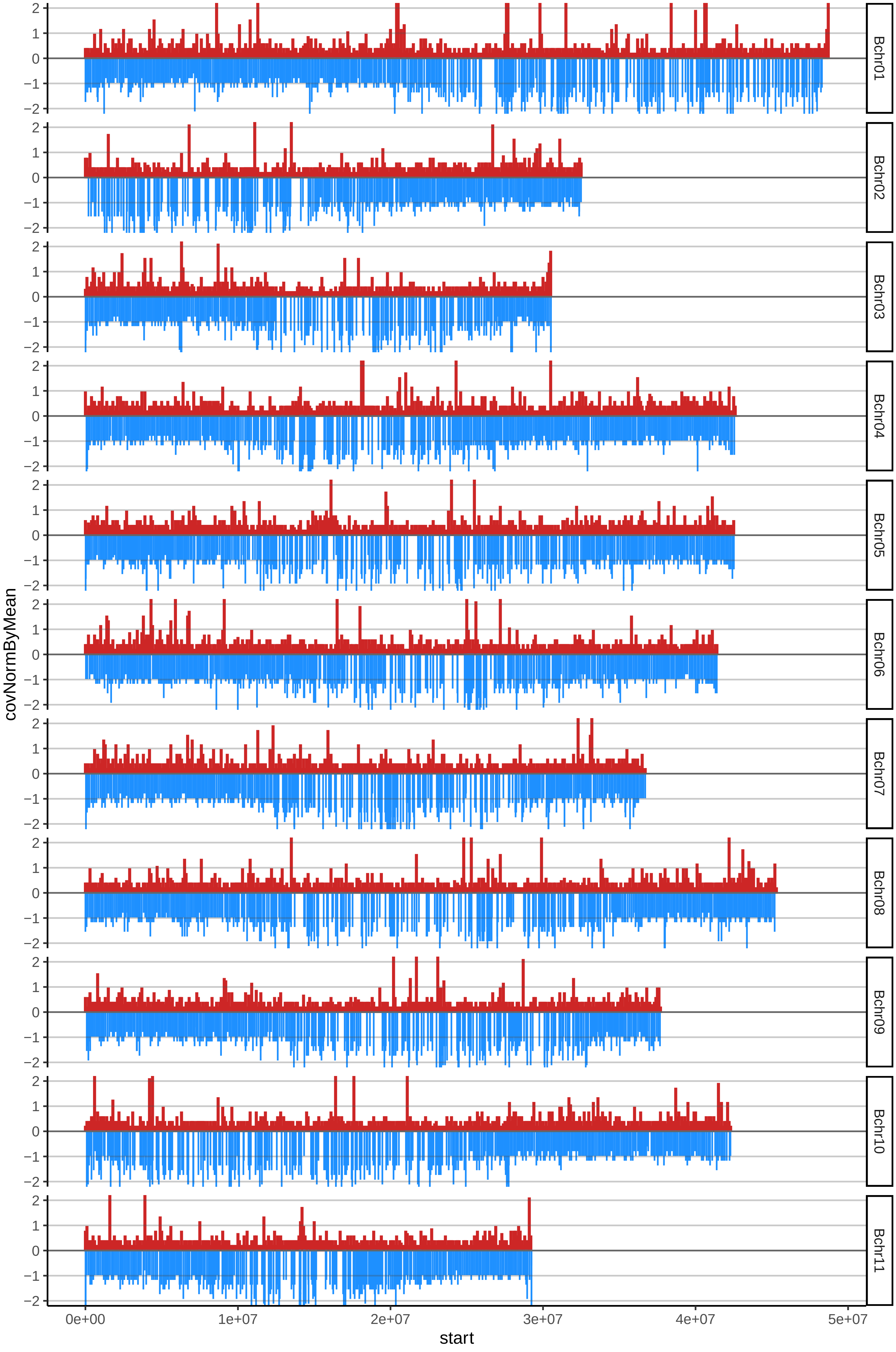

### 136DOMINICOCAOBAcov_byA.pdf

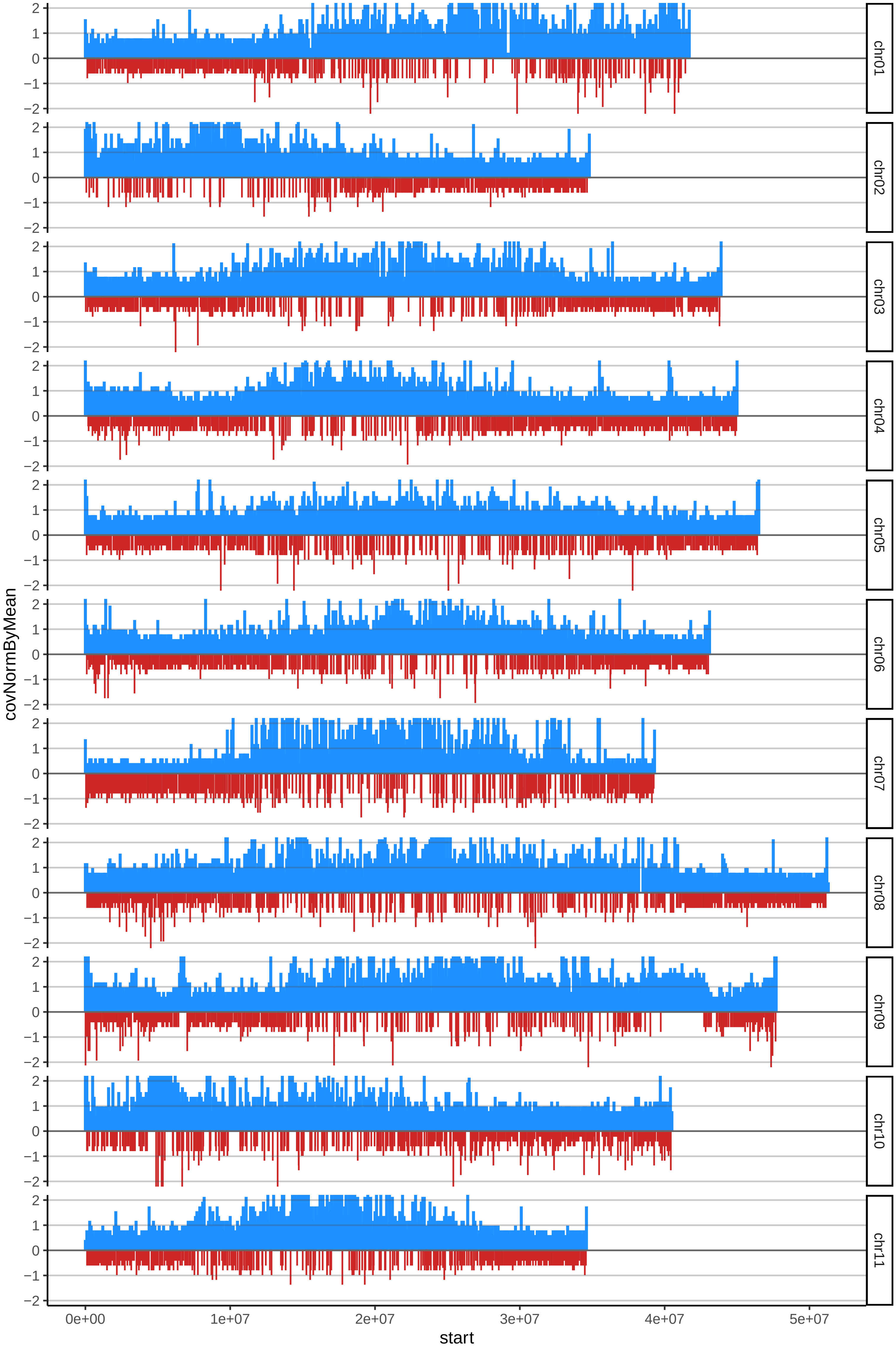

### 136DOMINICOCAOBAcov_byB.pdf

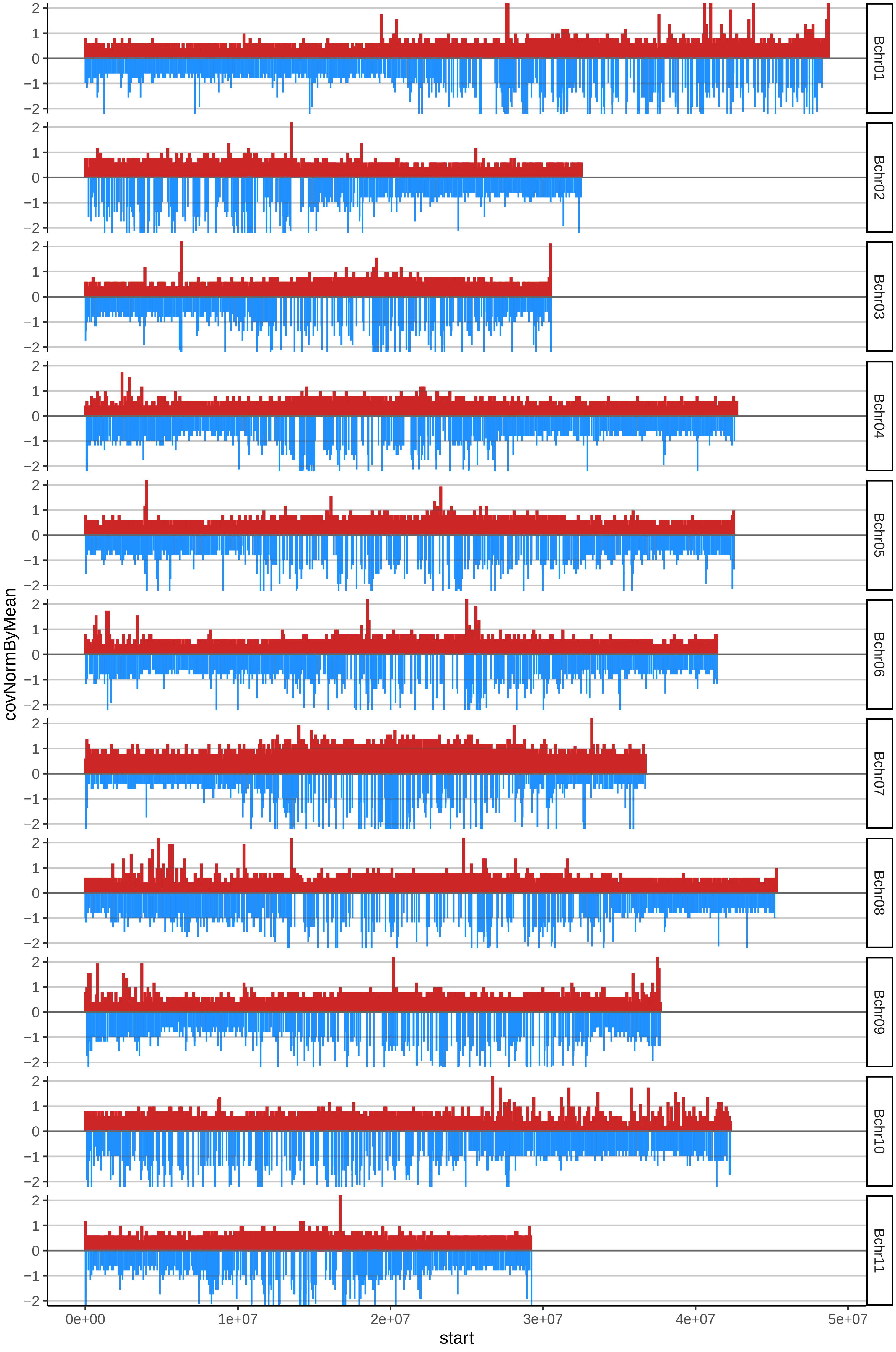

### 138HARTONROJODELMETAcov_byA.pdf

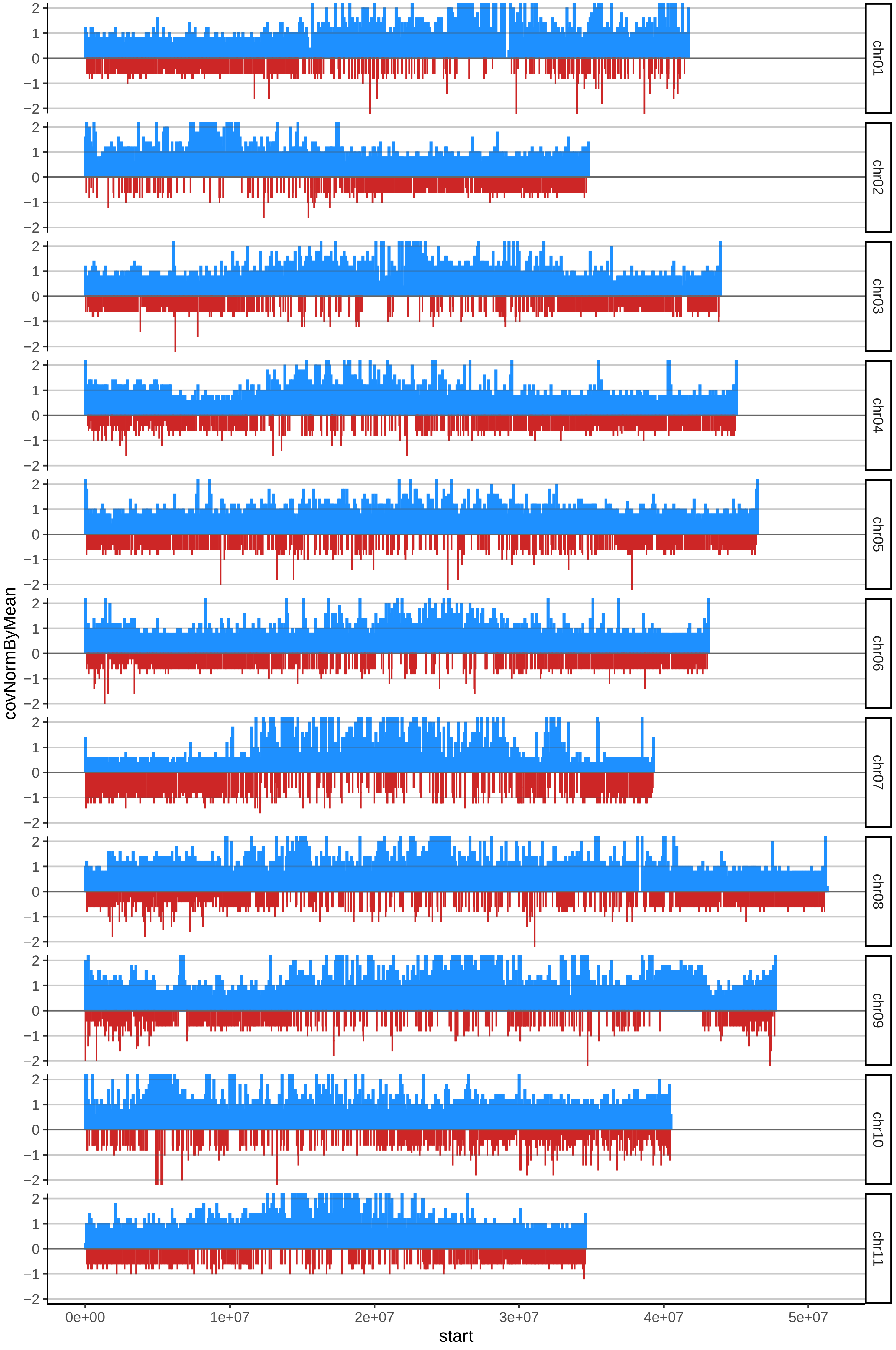

### 138HARTONROJODELMETAcov_byB.pdf

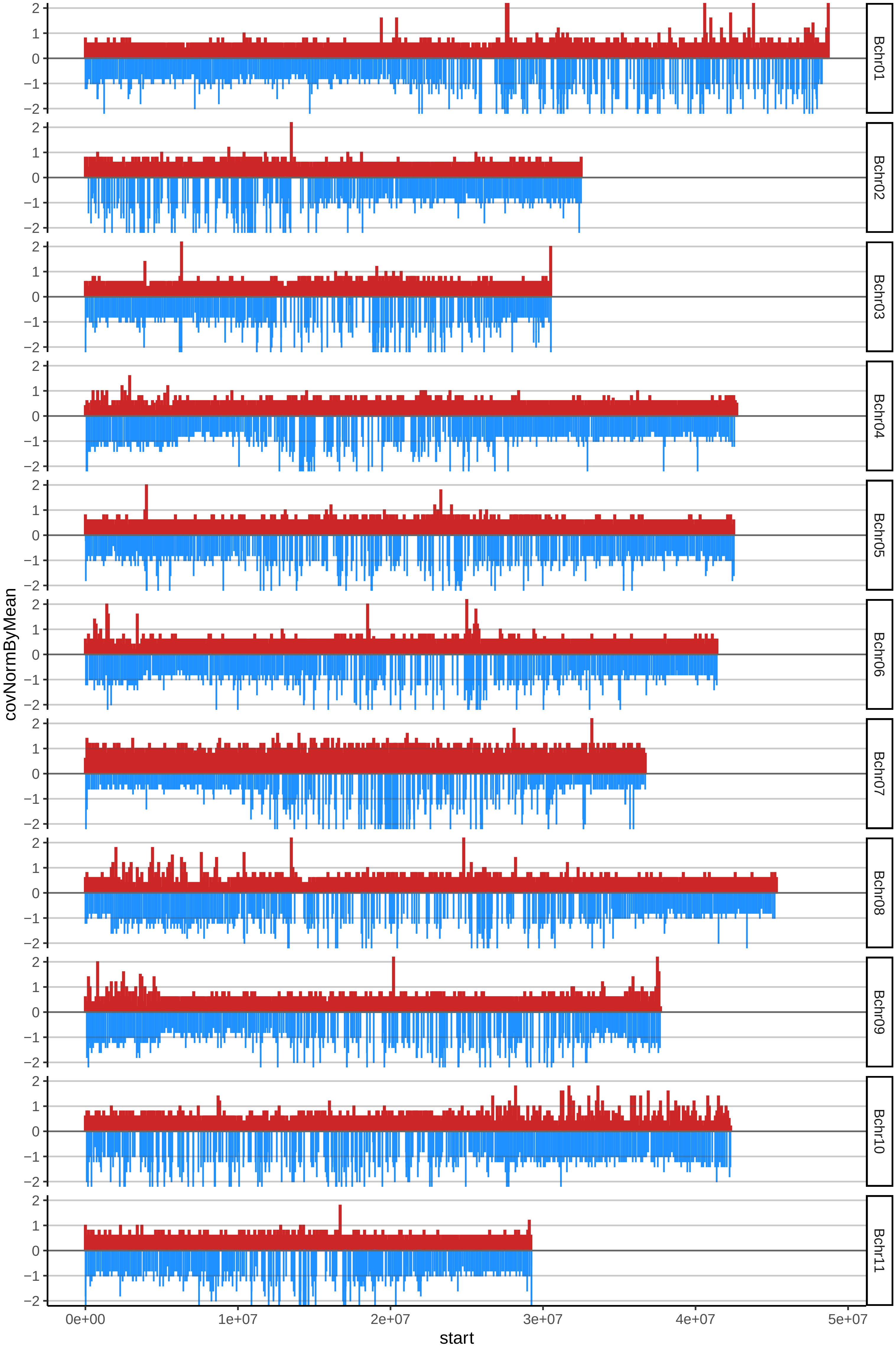

### 140HARTONLIBERALcov_byA.pdf

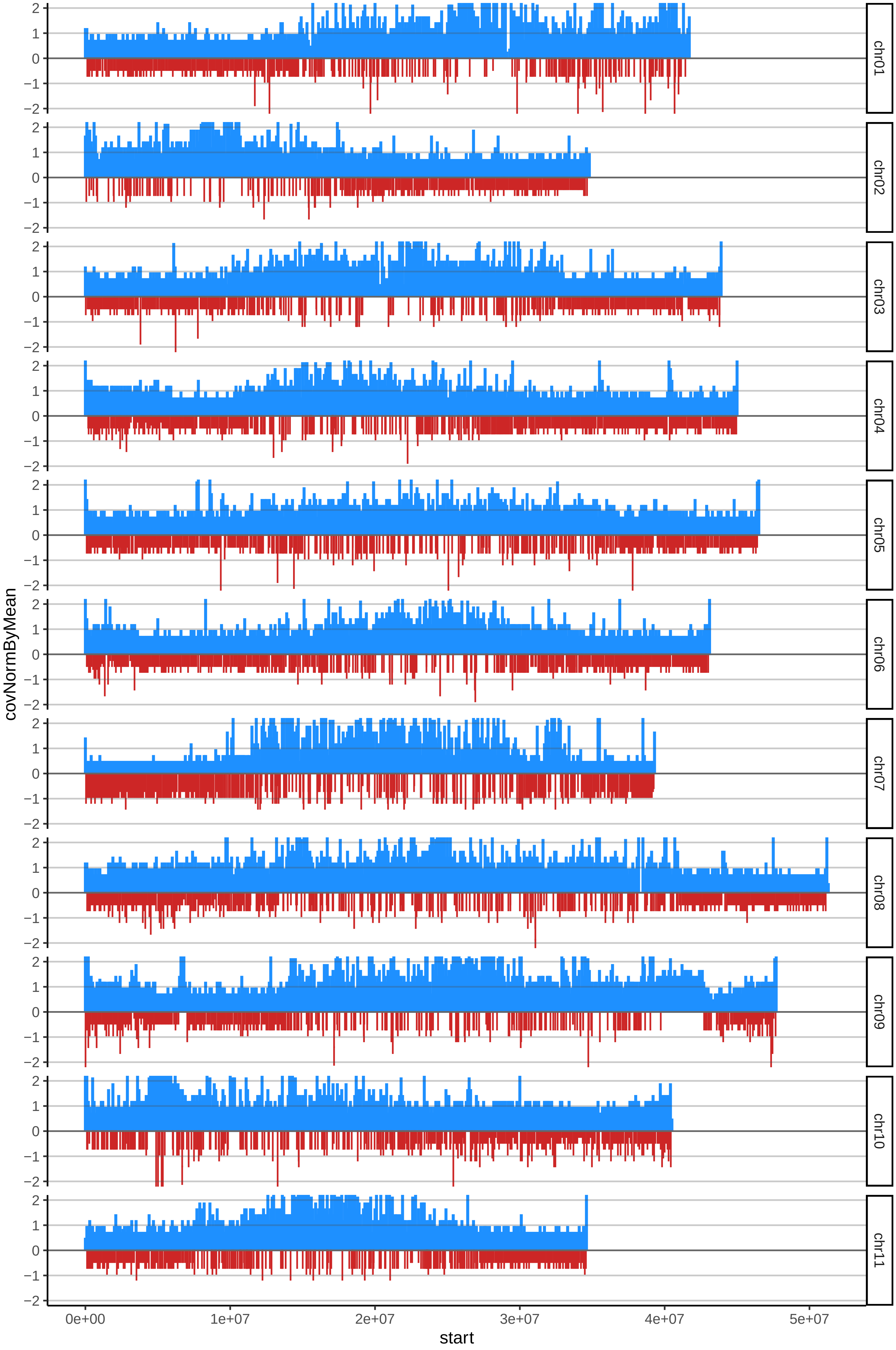

### 140HARTONLIBERALcov_byB.pdf

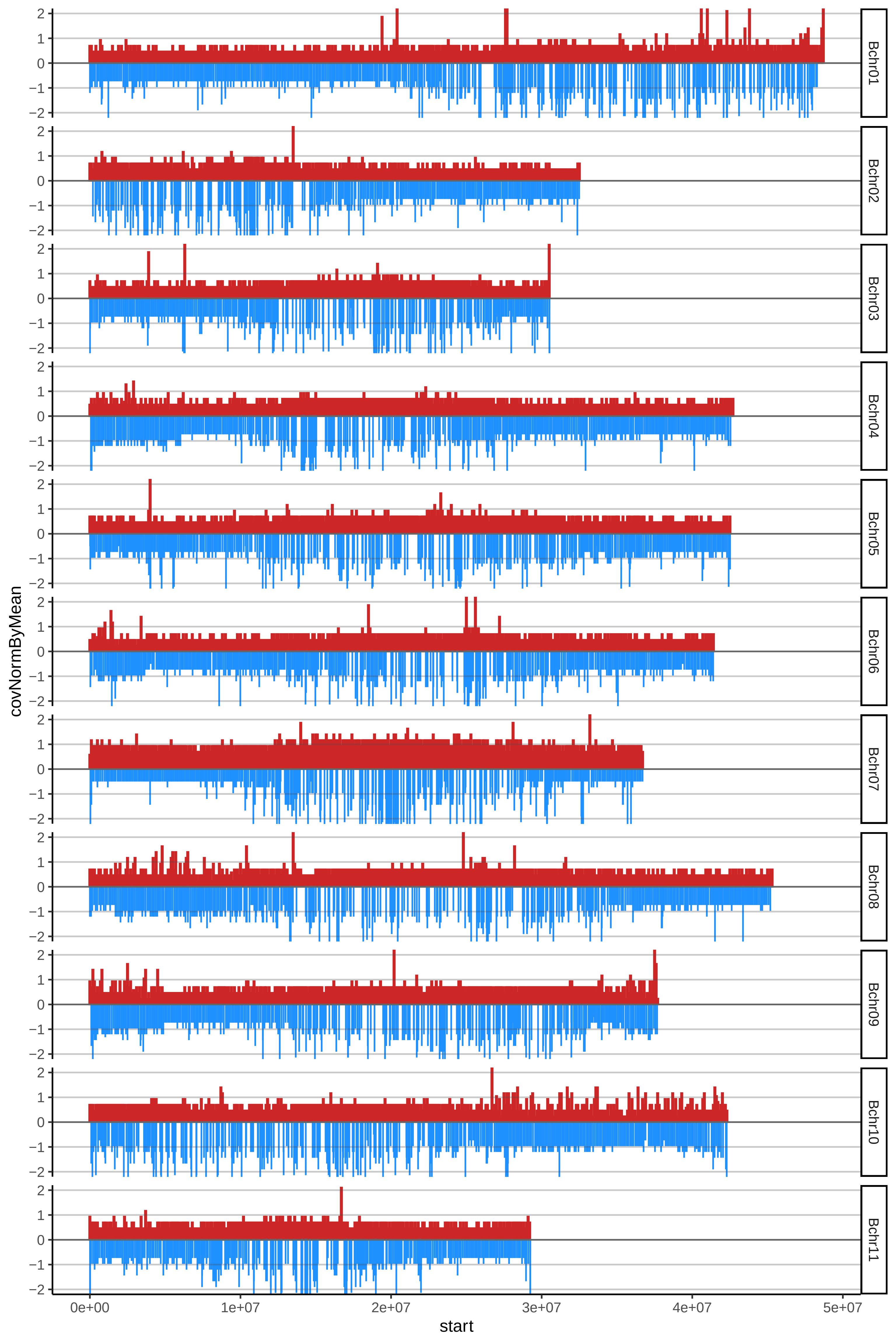

### 146GUAYABOBcov_byA.pdf

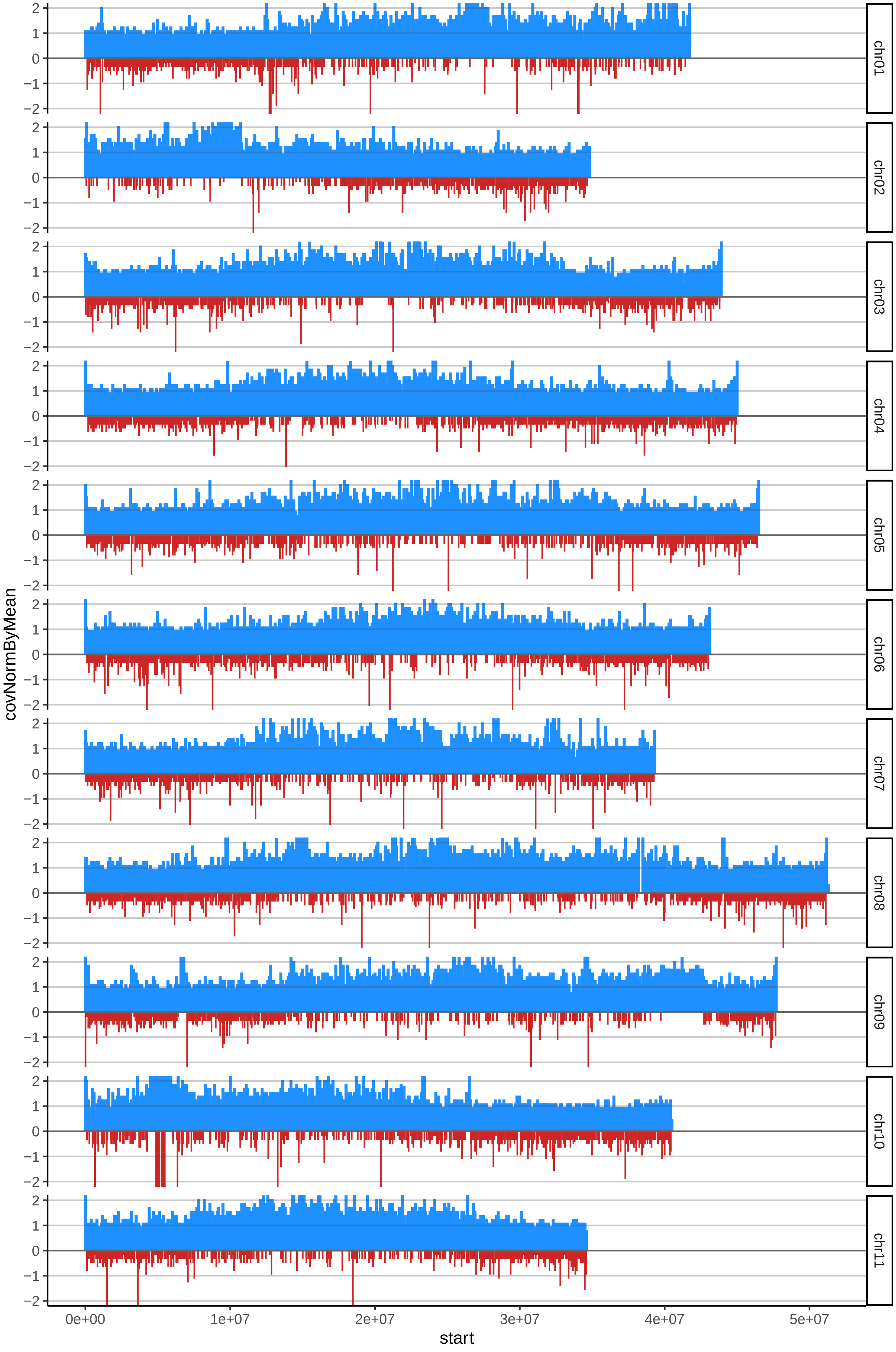

### 146GUAYABOBcov_byB.pdf

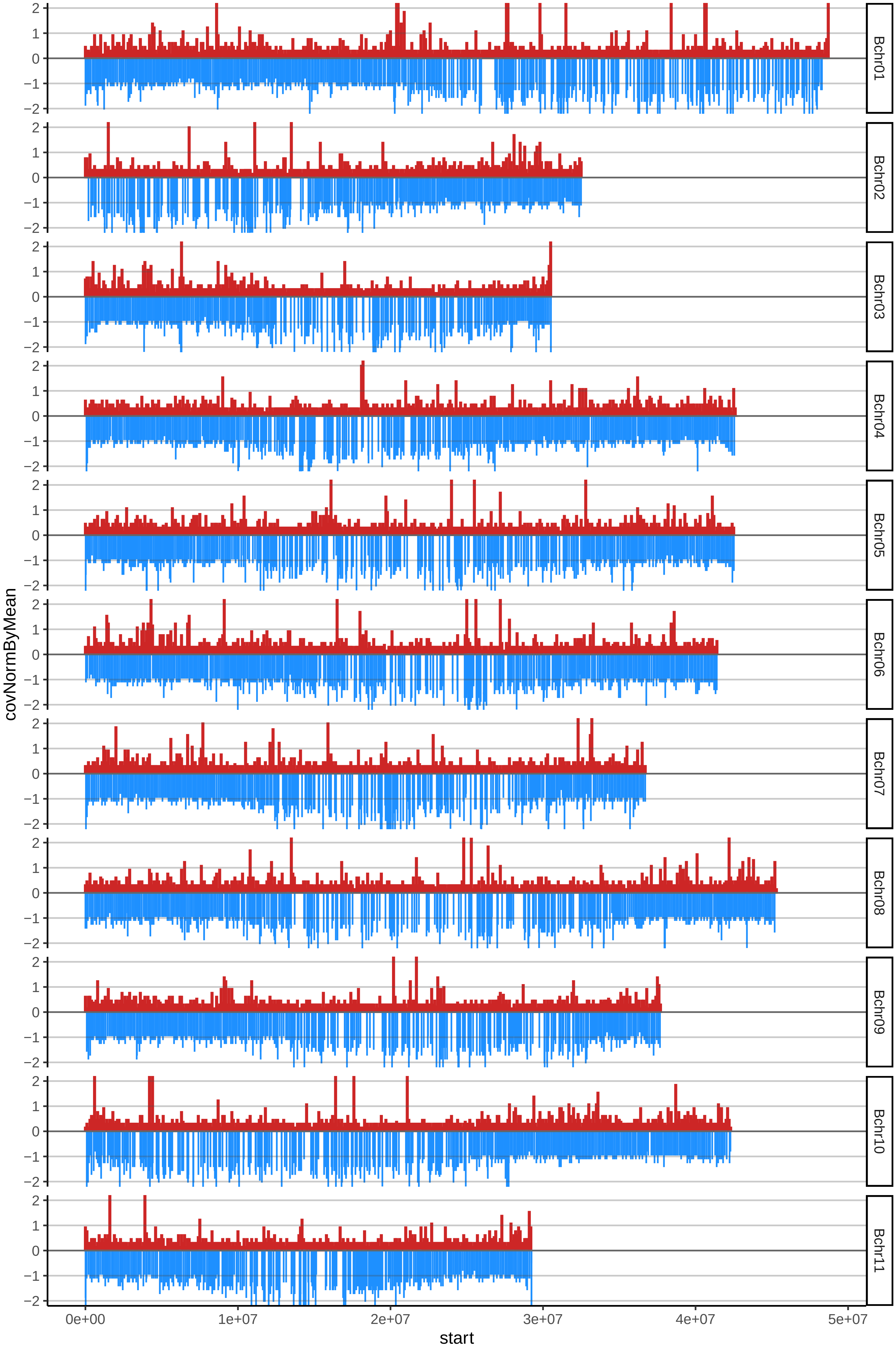

### 150HAWAIANOcov_byA.pdf

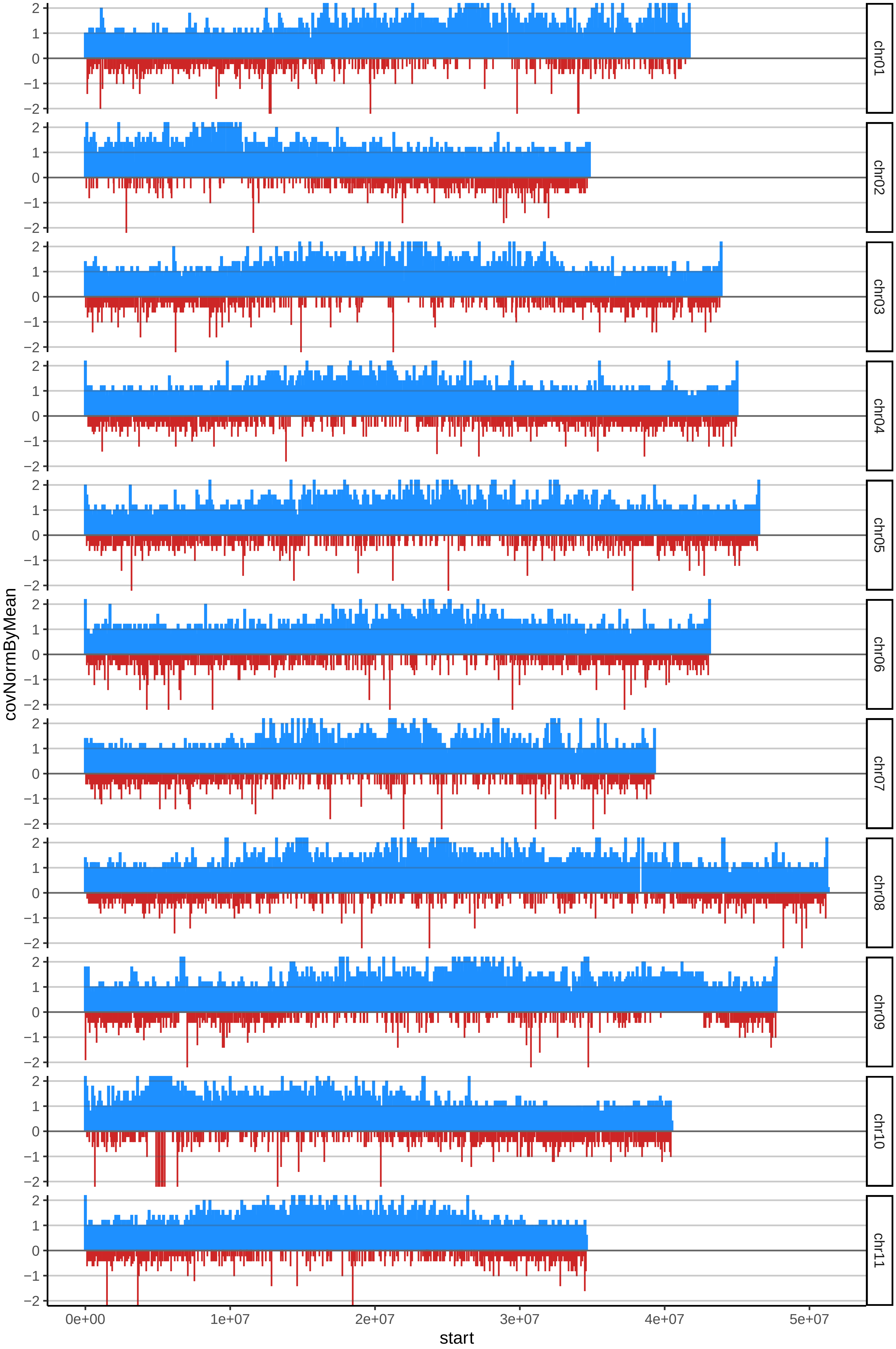

### 150HAWAIANOcov_byB.pdf

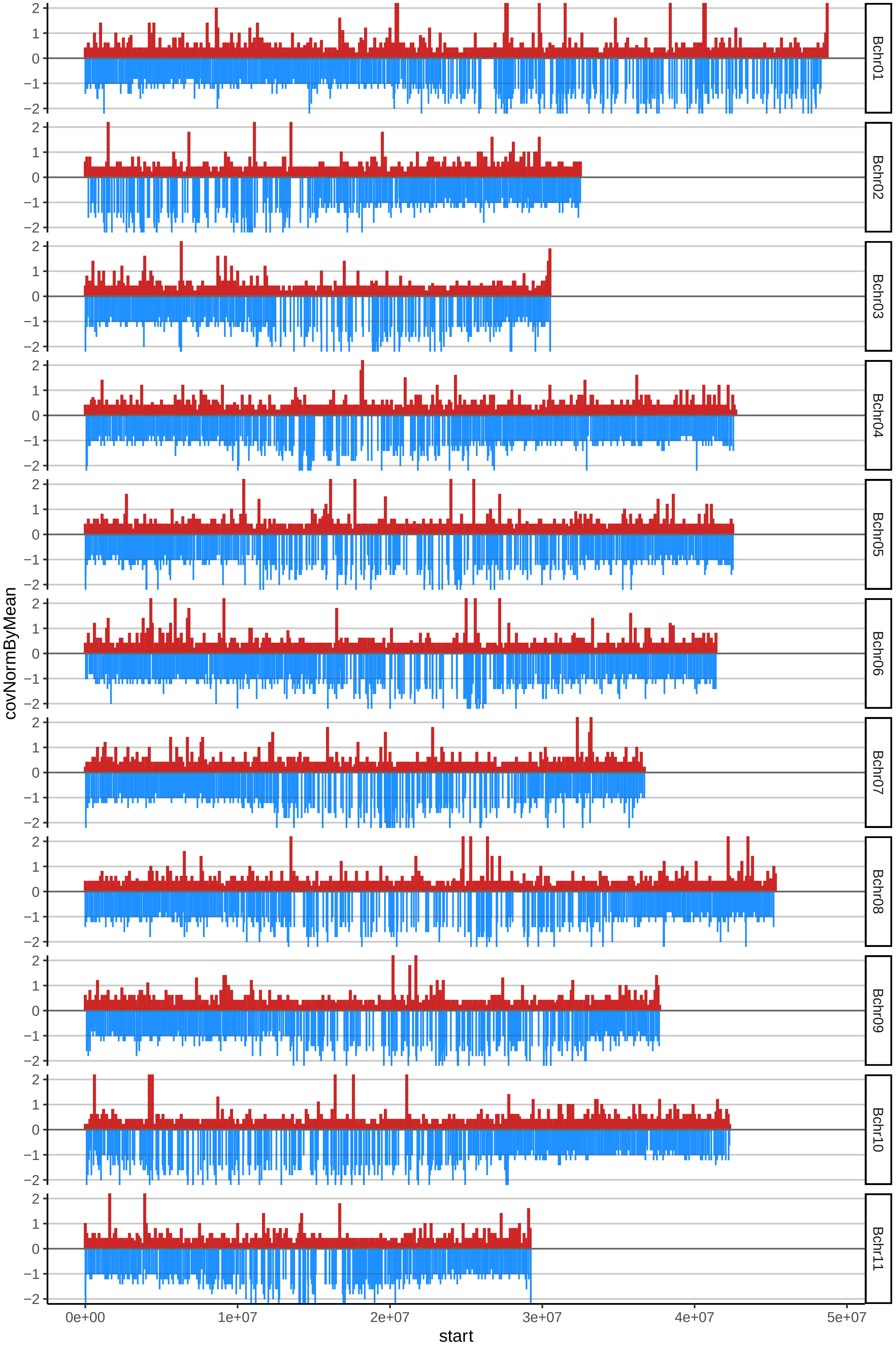

### 151BLANCOcov_byA.pdf

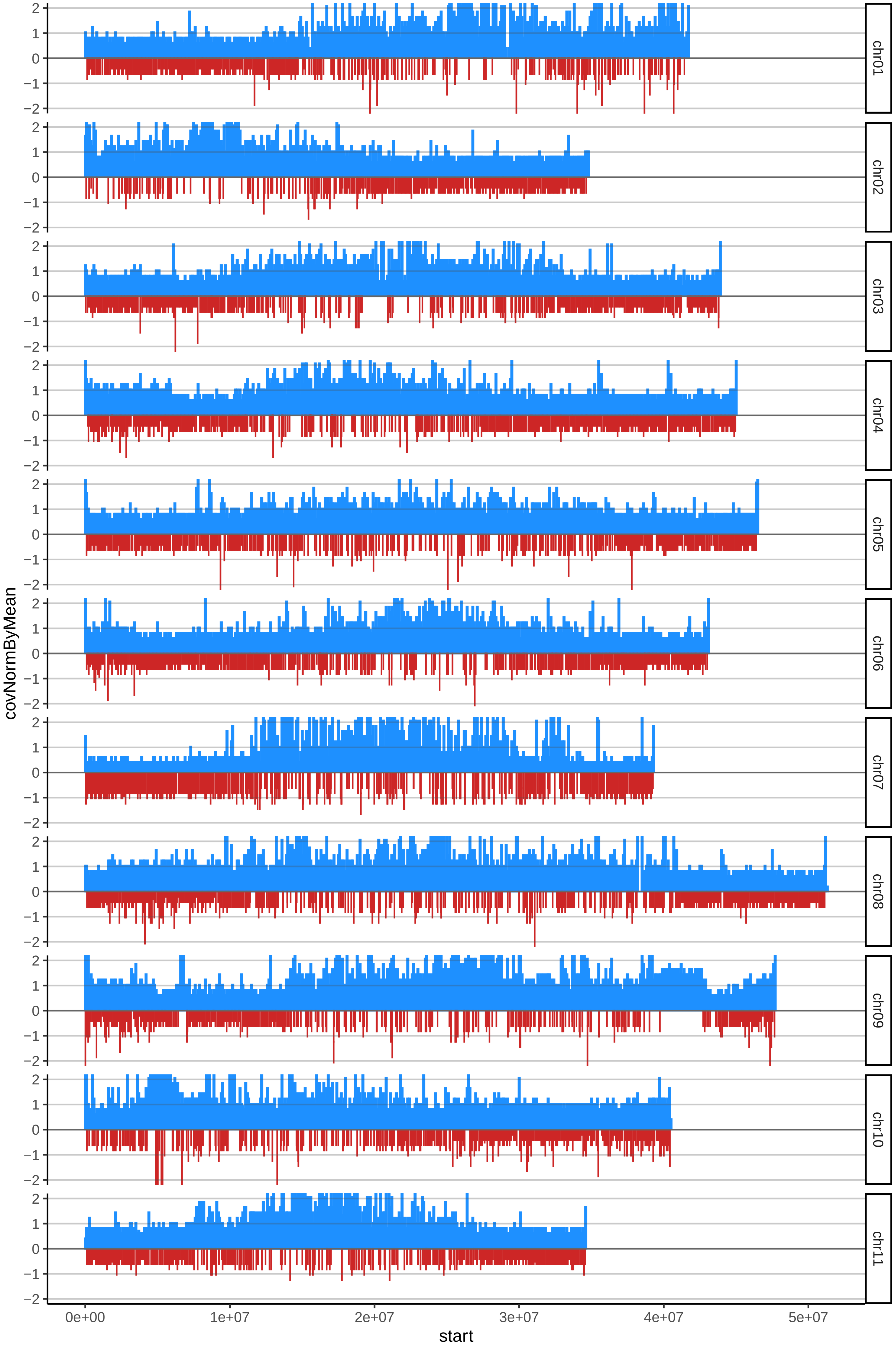

### 151BLANCOcov_byB.pdf

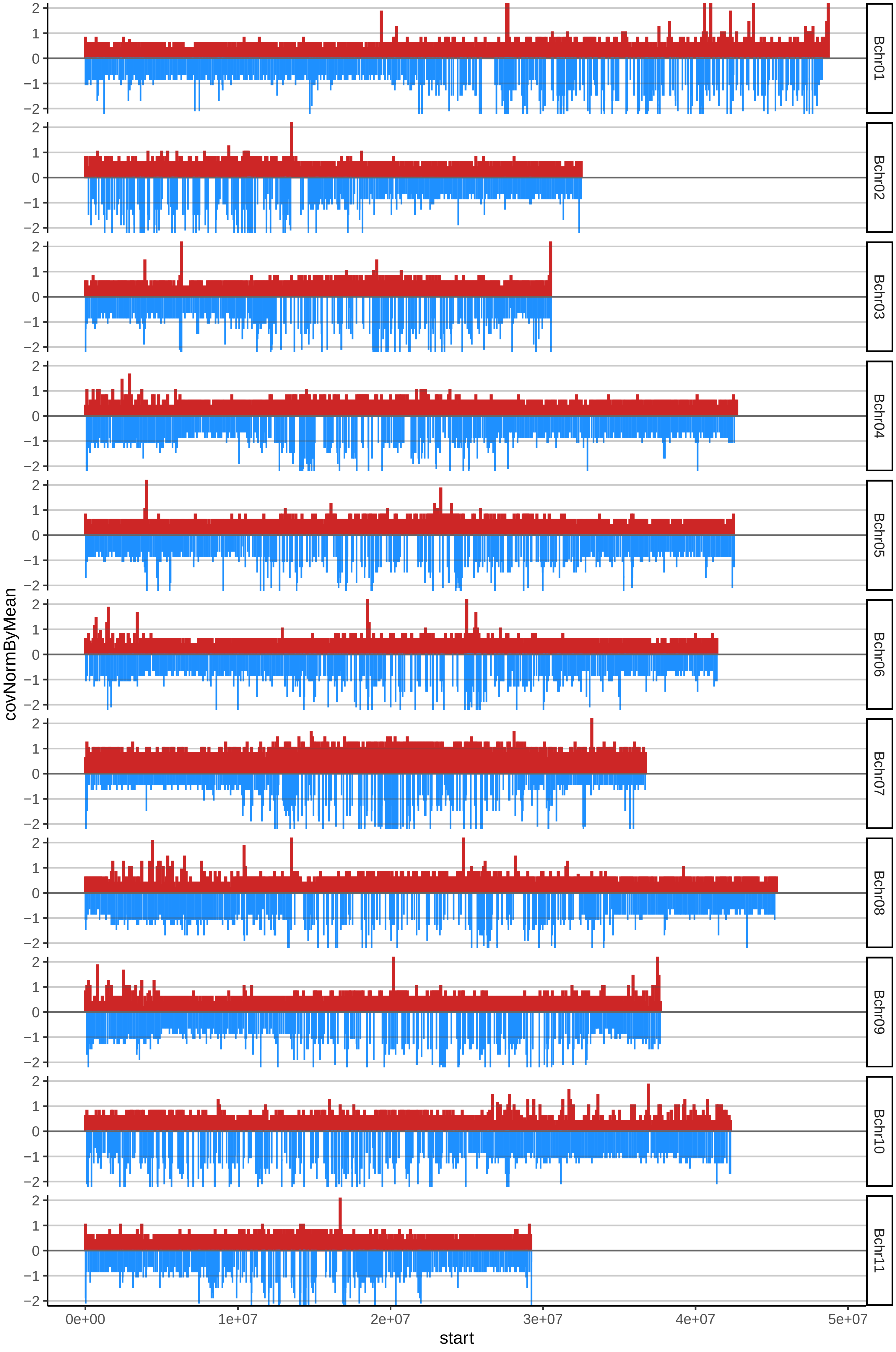

### 153MAQUENNOcov_byA.pdf

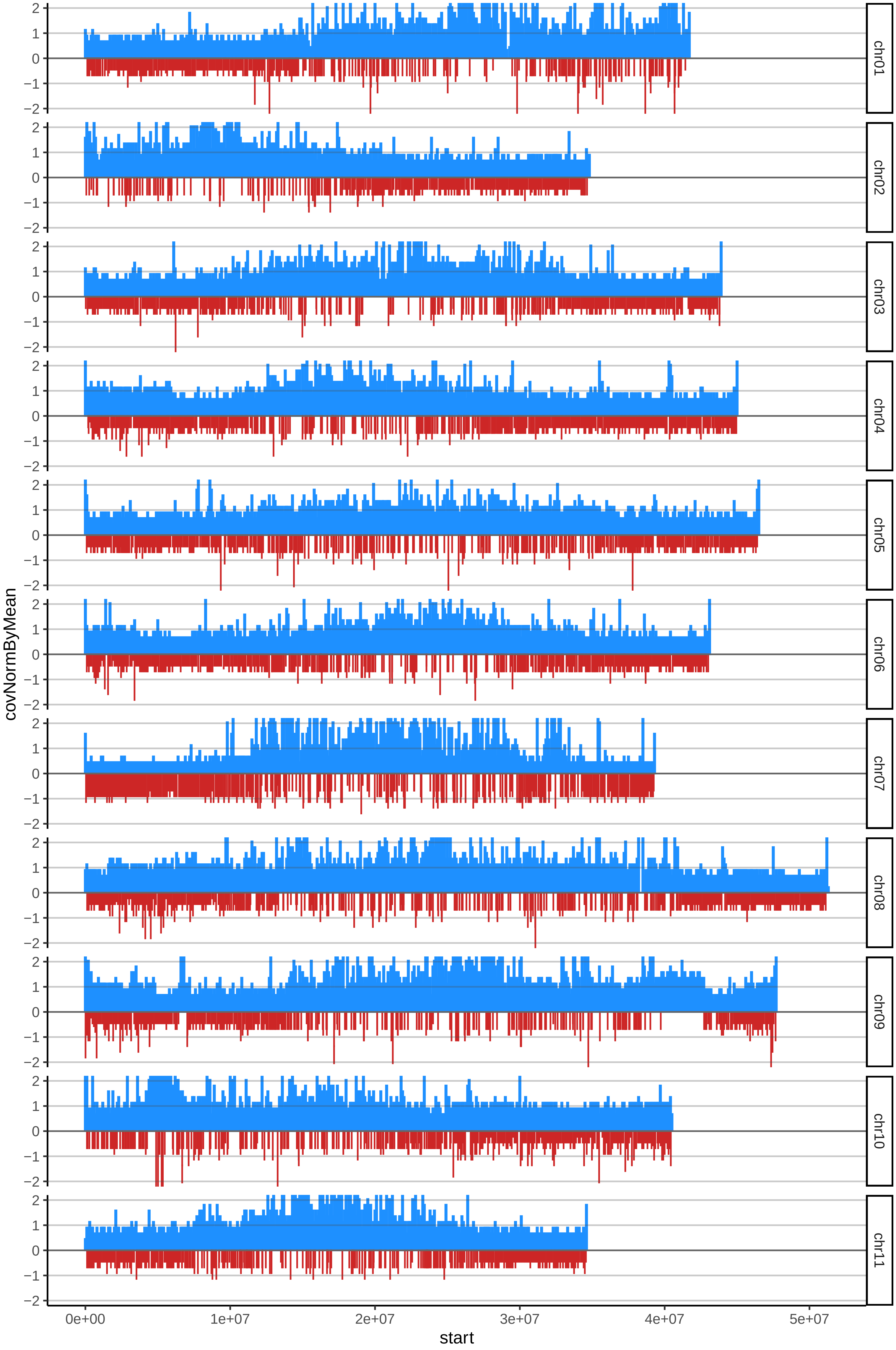

### 153MAQUENNOcov_byB.pdf

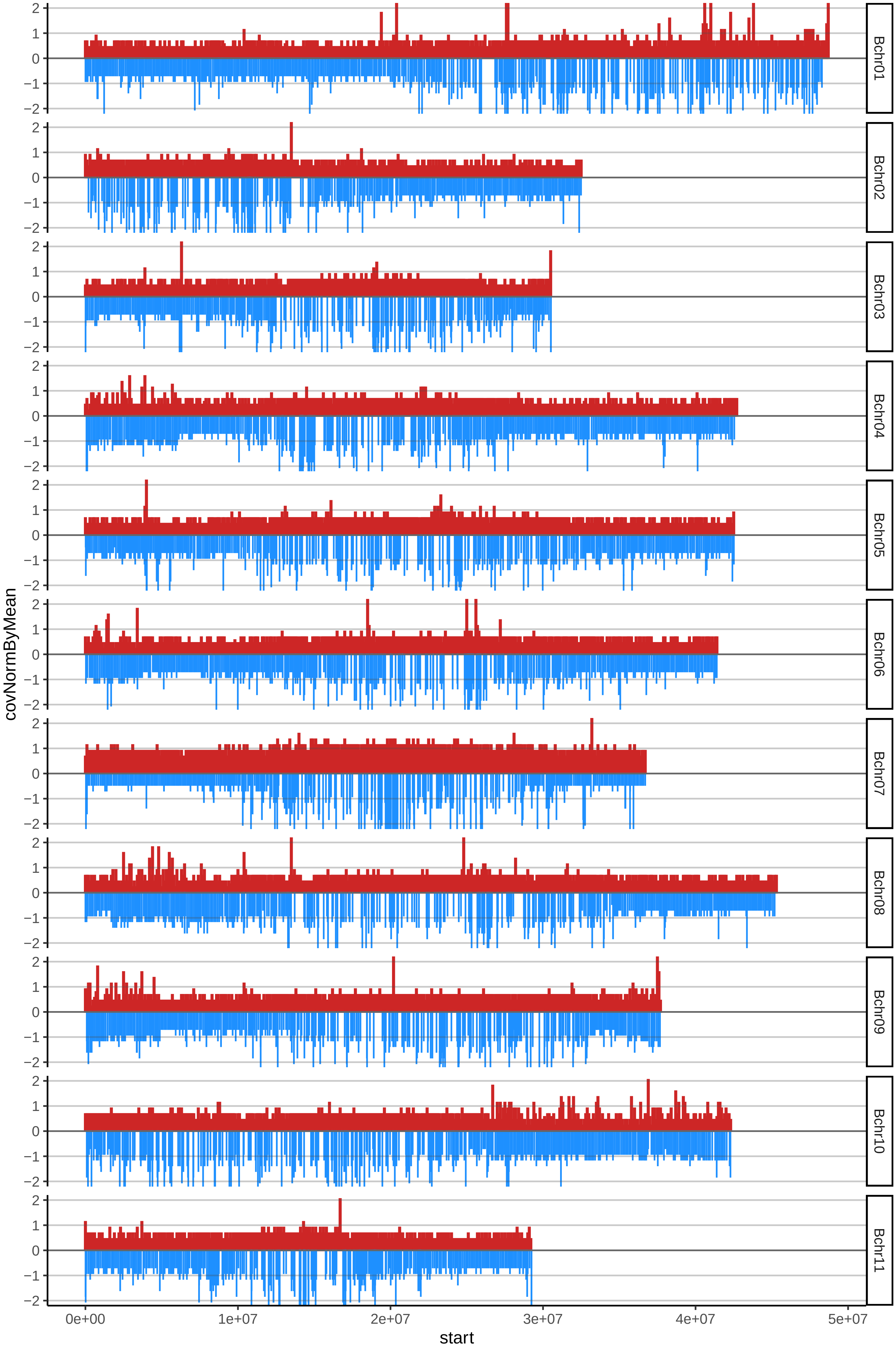

### 157ANVA03cov_byA.pdf

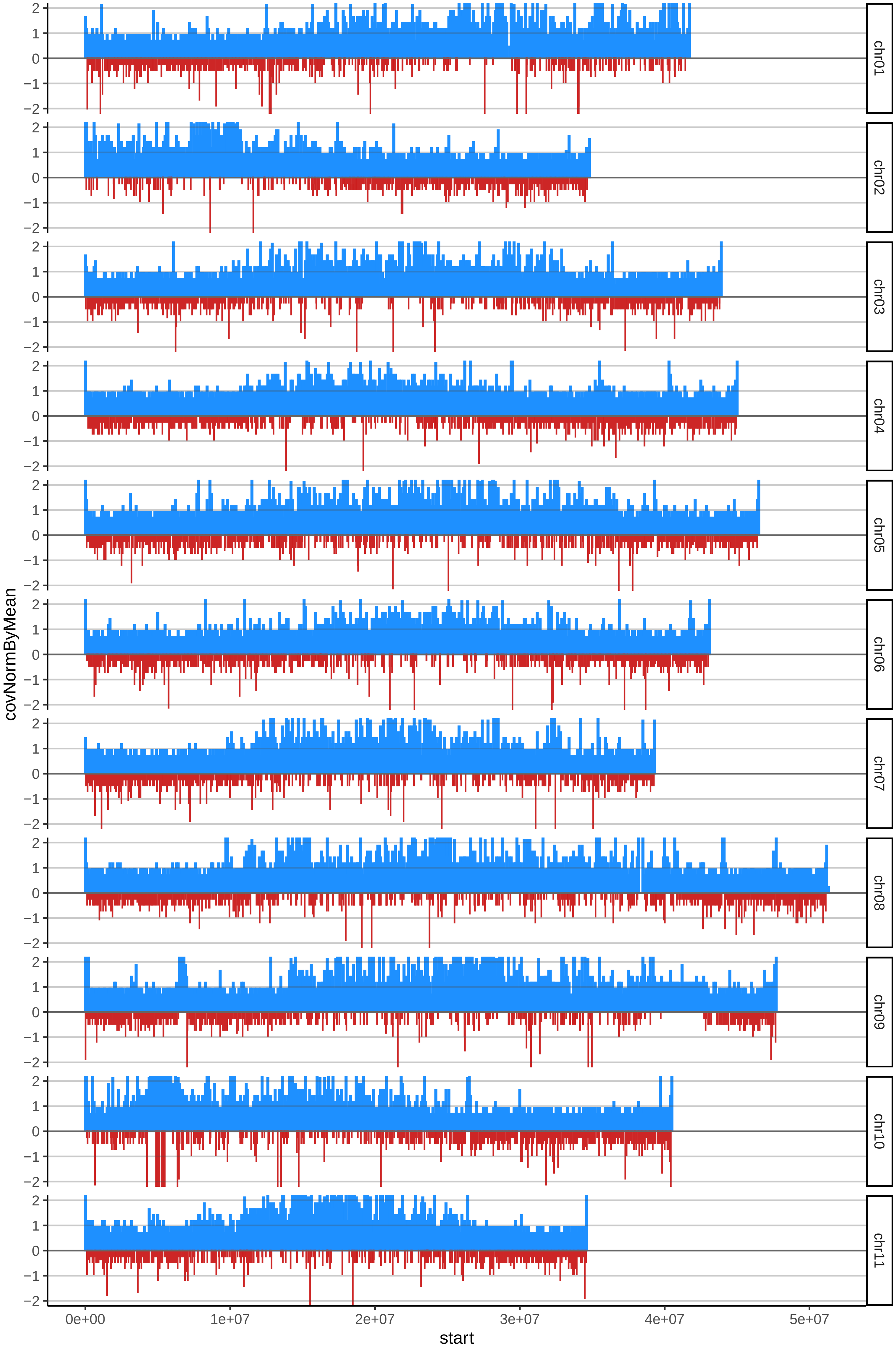

### 157ANVA03cov_byB.pdf

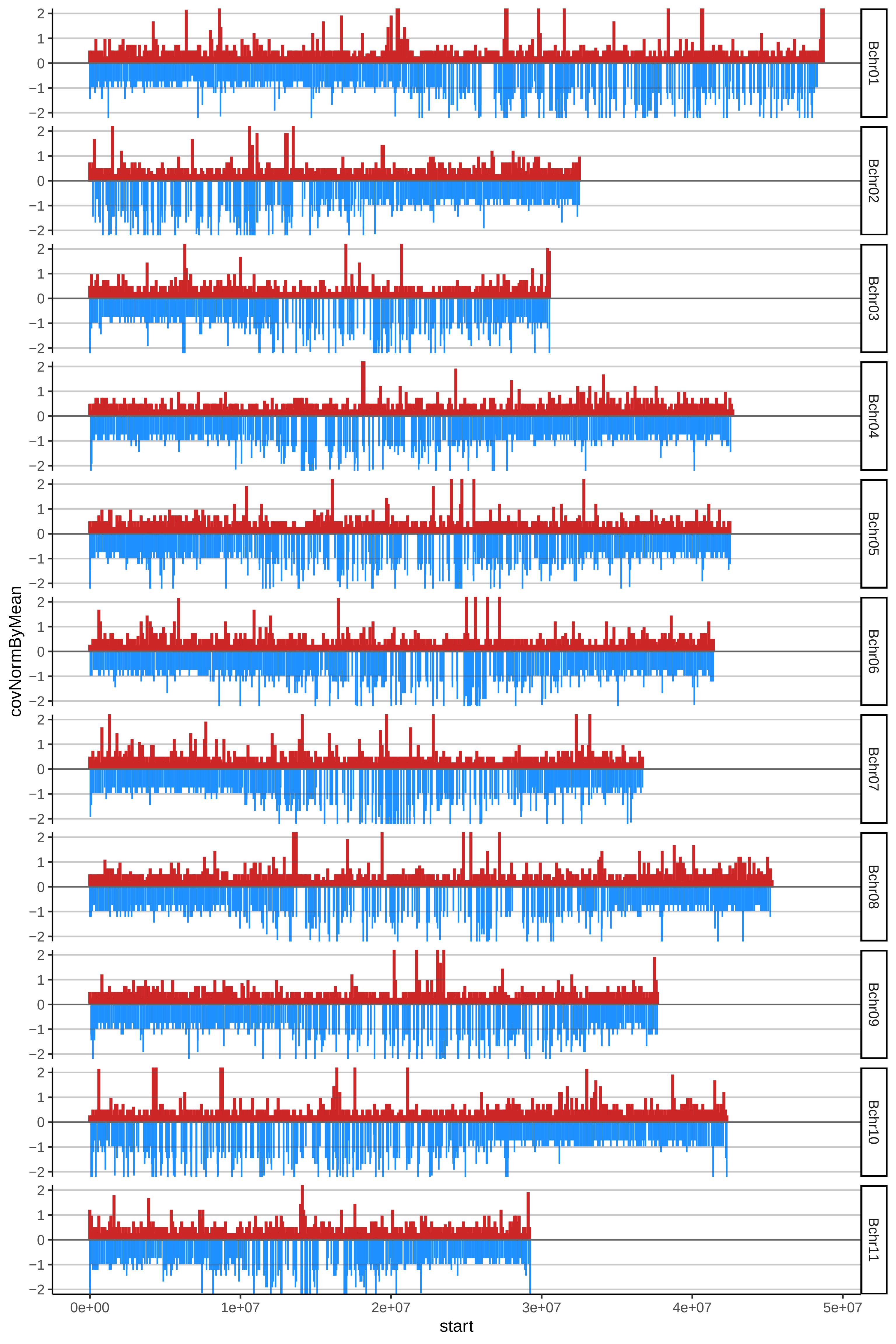

### 158NALLO04cov_byA.pdf

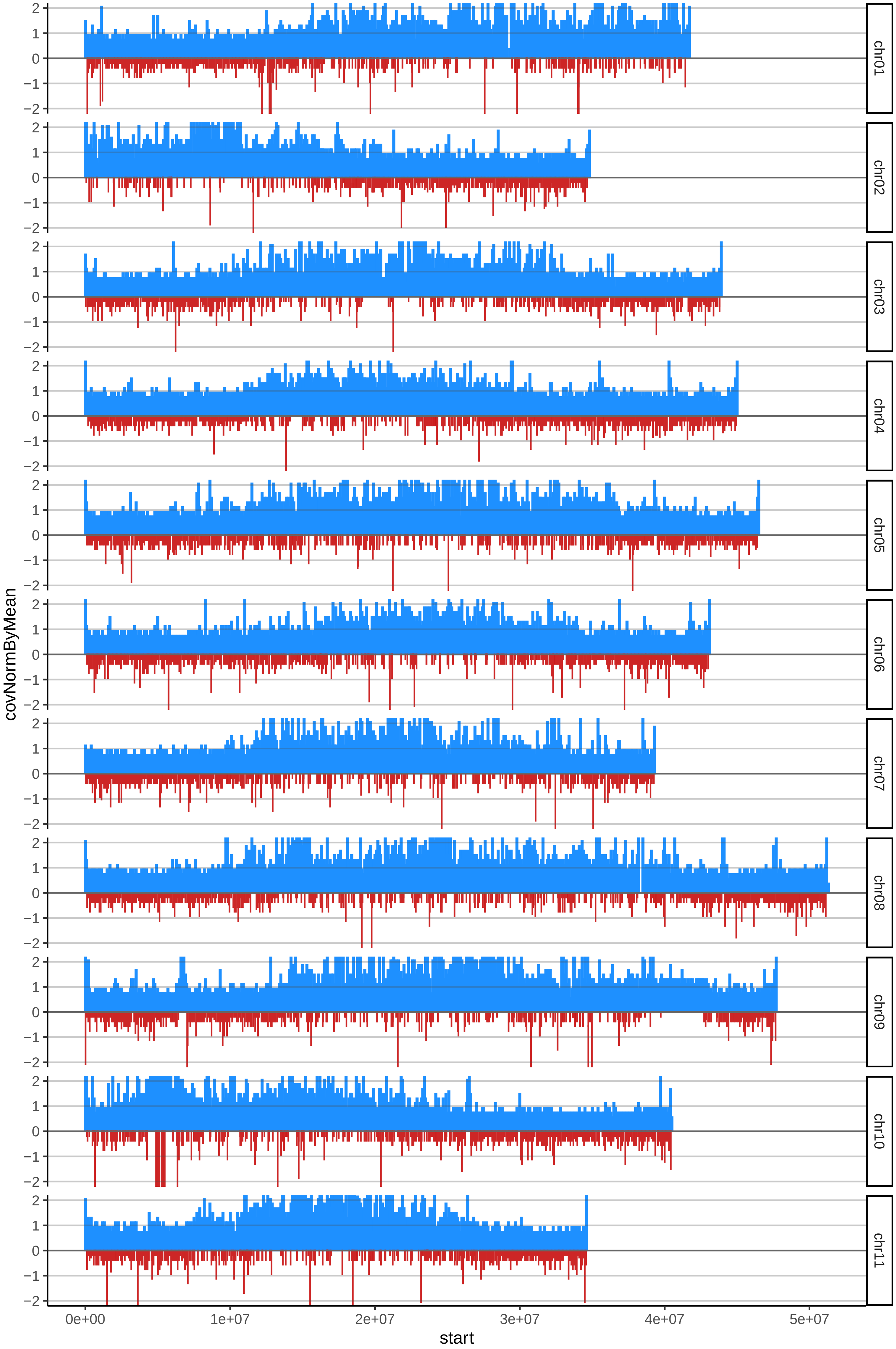

### 158NALLO04cov_byB.pdf

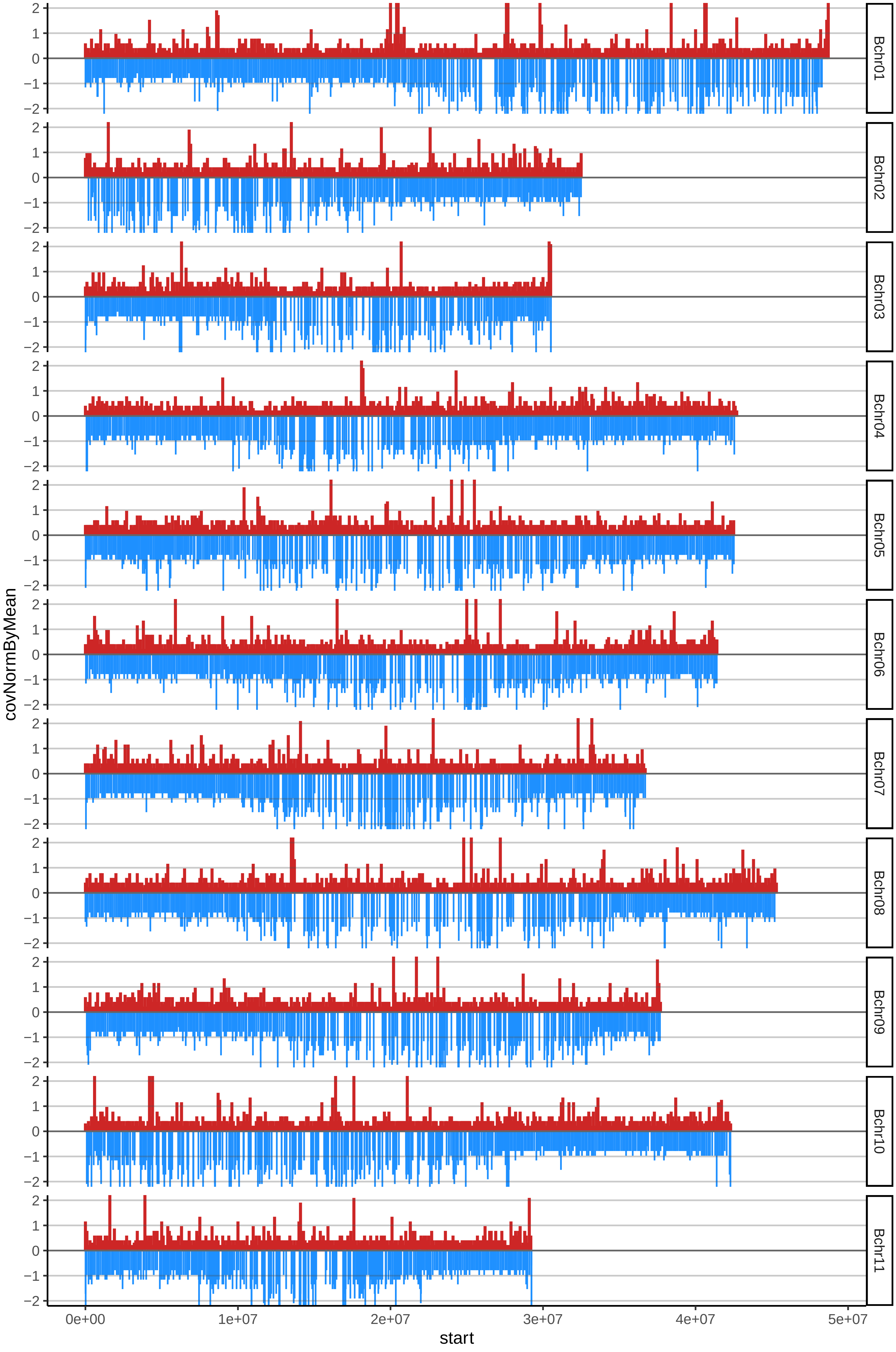

### 160NALLO06cov_byA.pdf

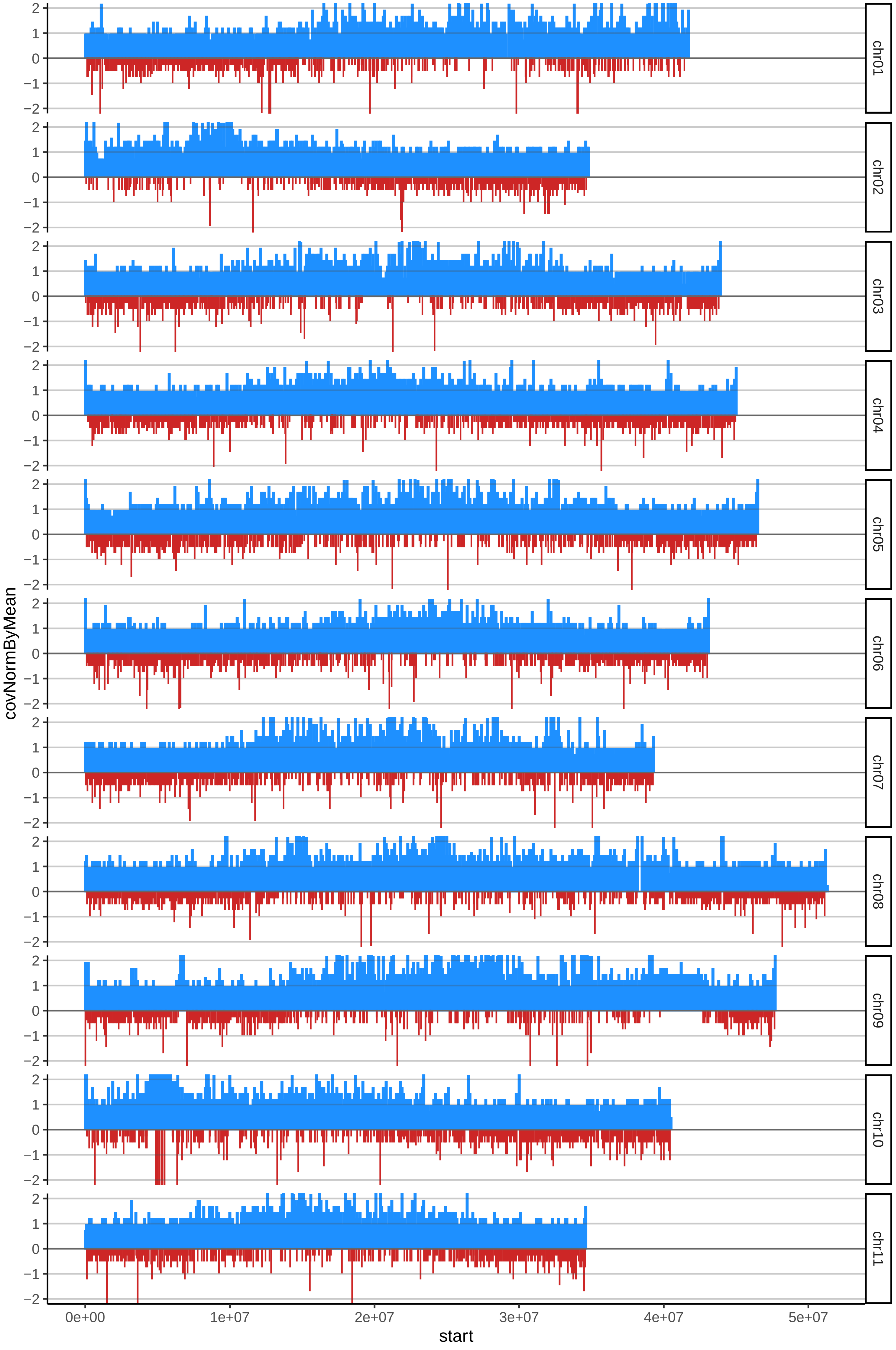

### 160NALLO06cov_byB.pdf

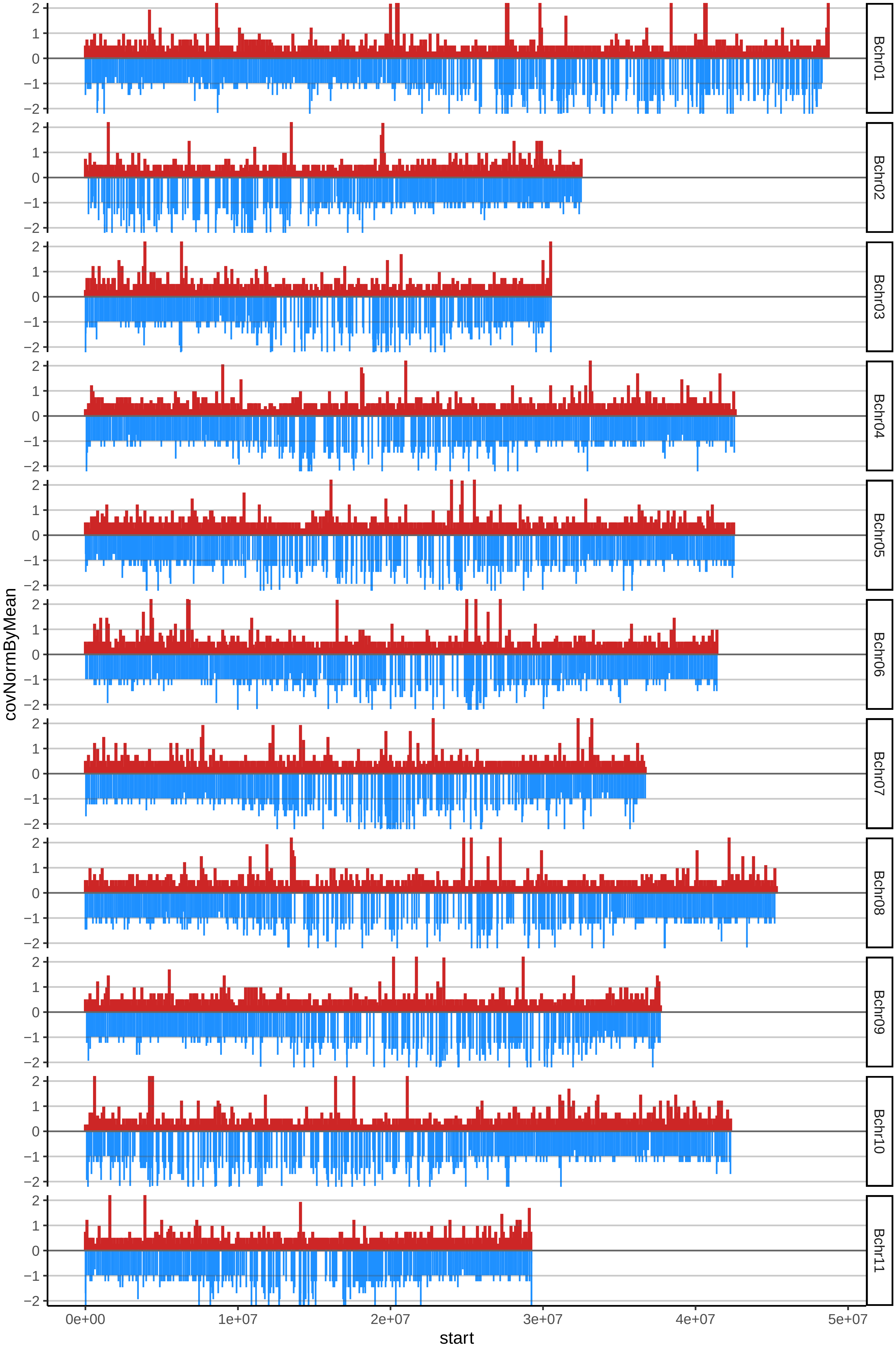

### 161NARI01cov_byA.pdf

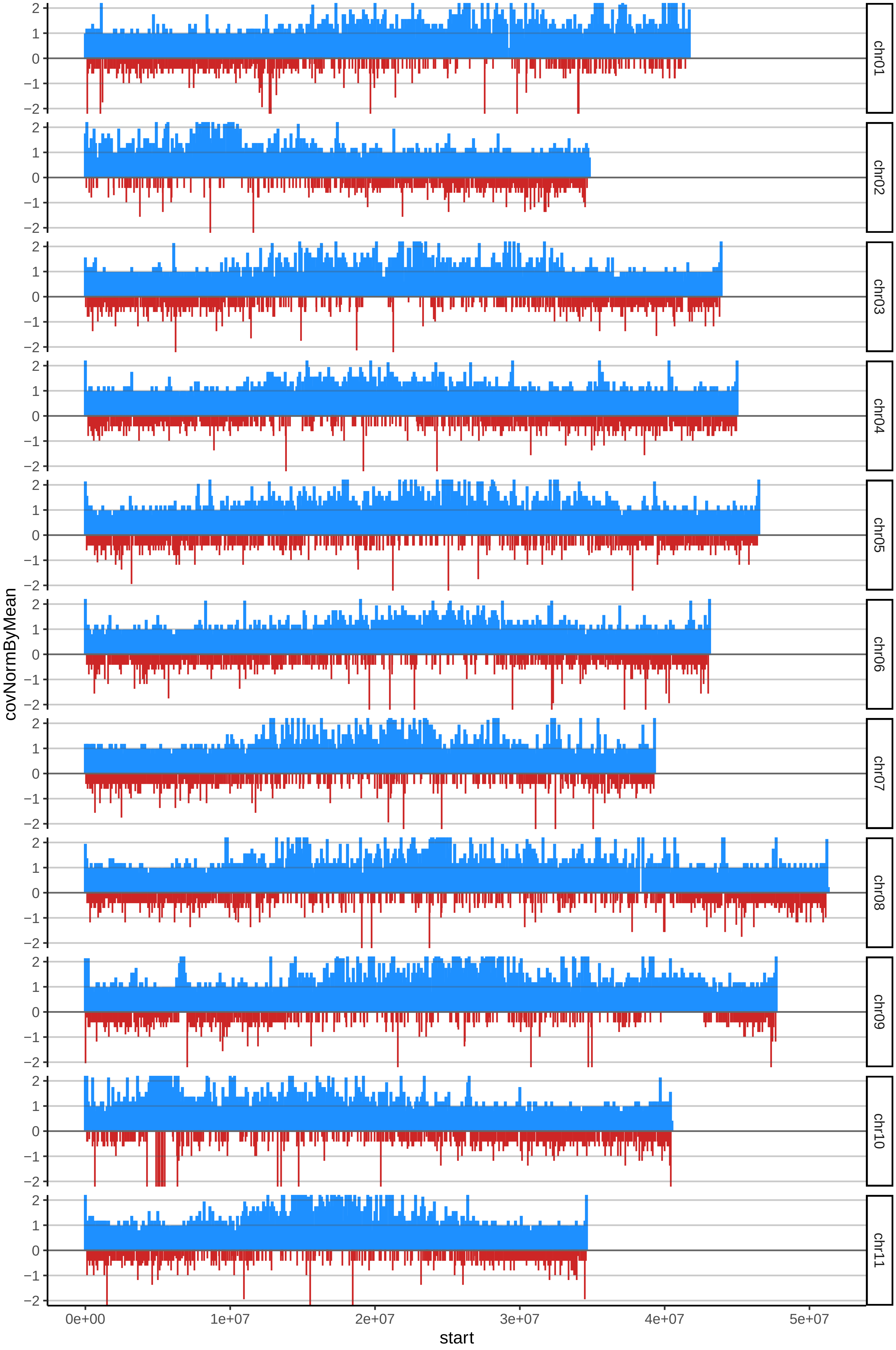

### 161NARI01cov_byB.pdf

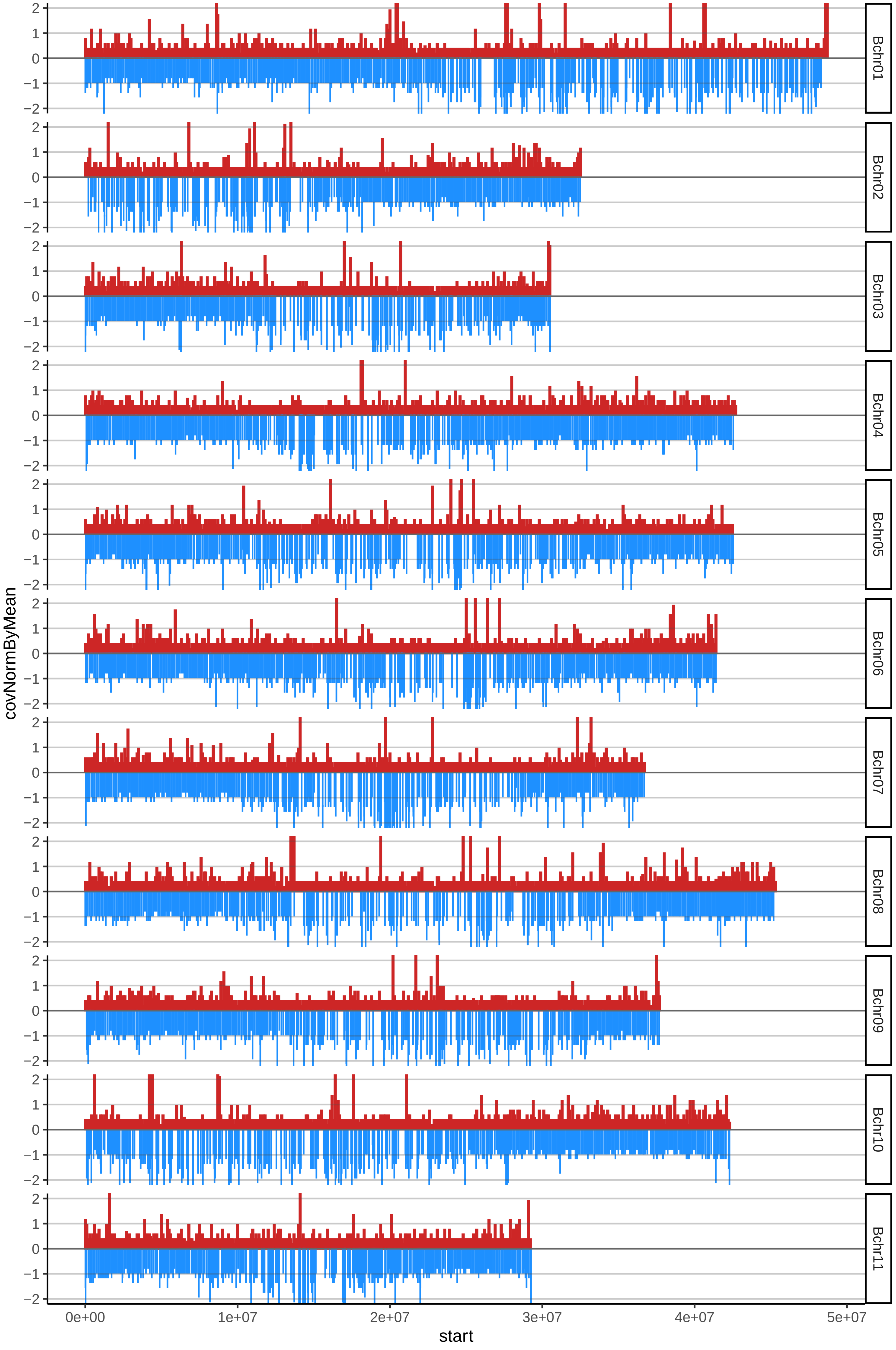

### 169SABO04cov_byA.pdf

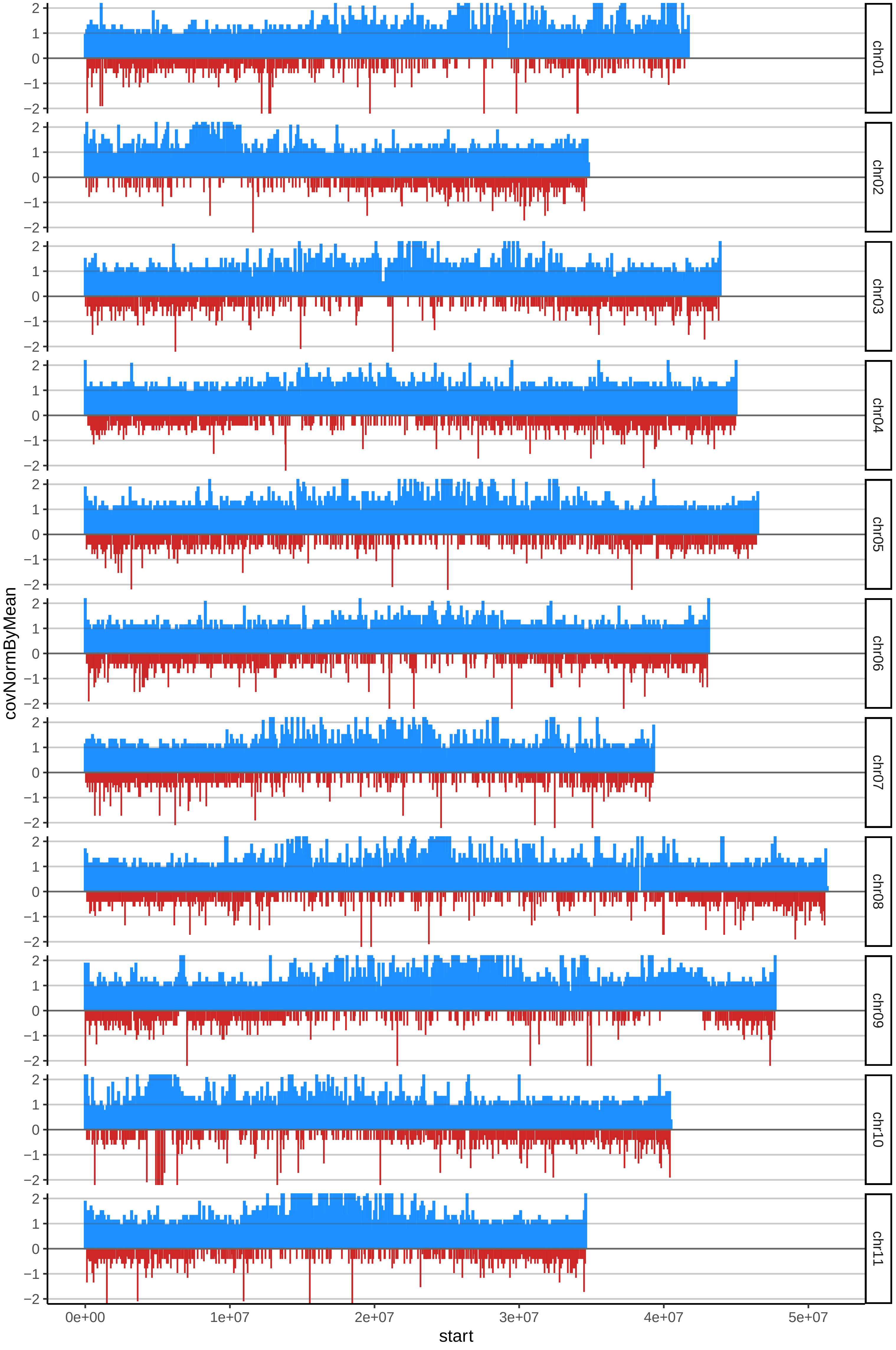

### 169SABO04cov_byB.pdf

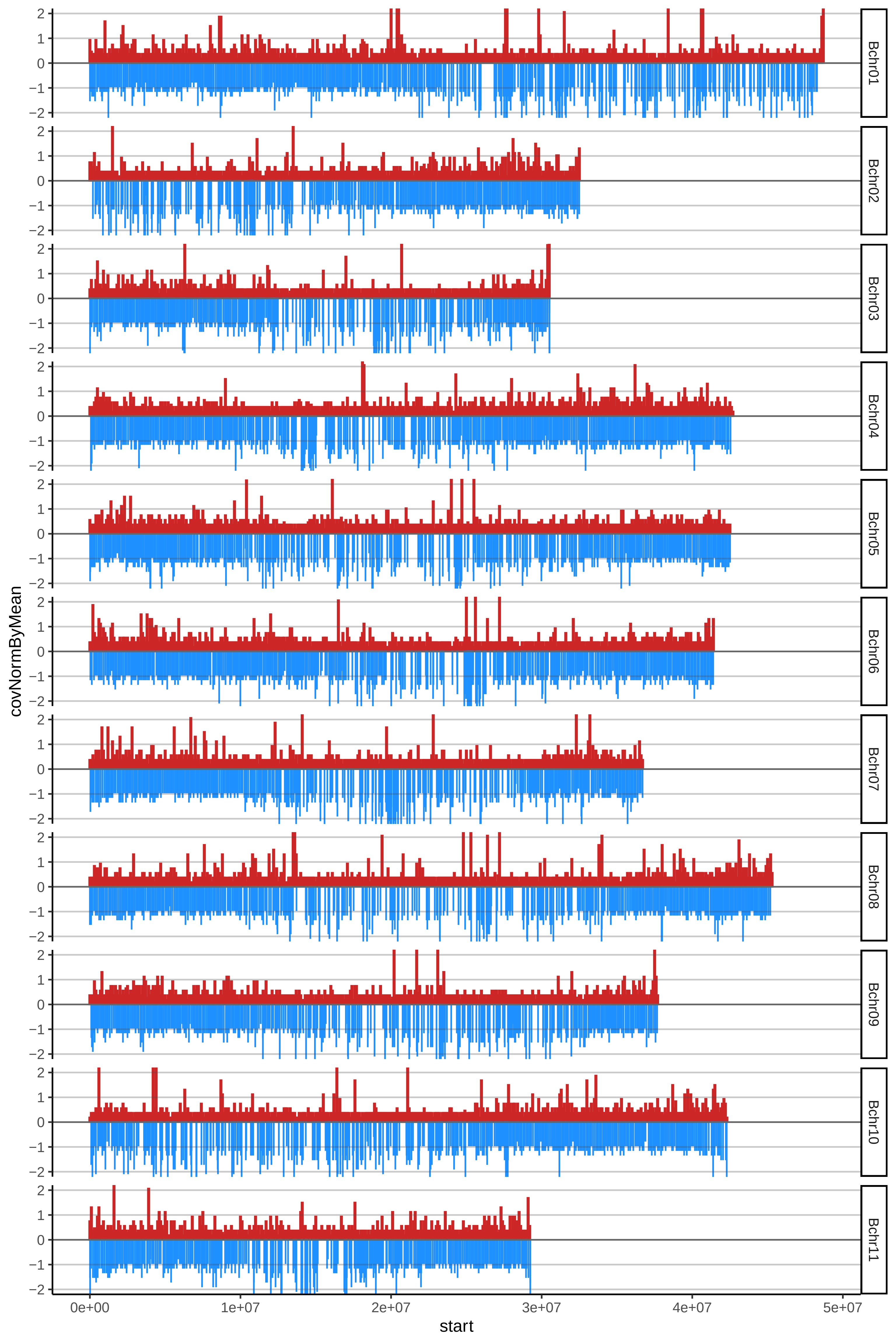

### 173SAPI09cov_byA.pdf

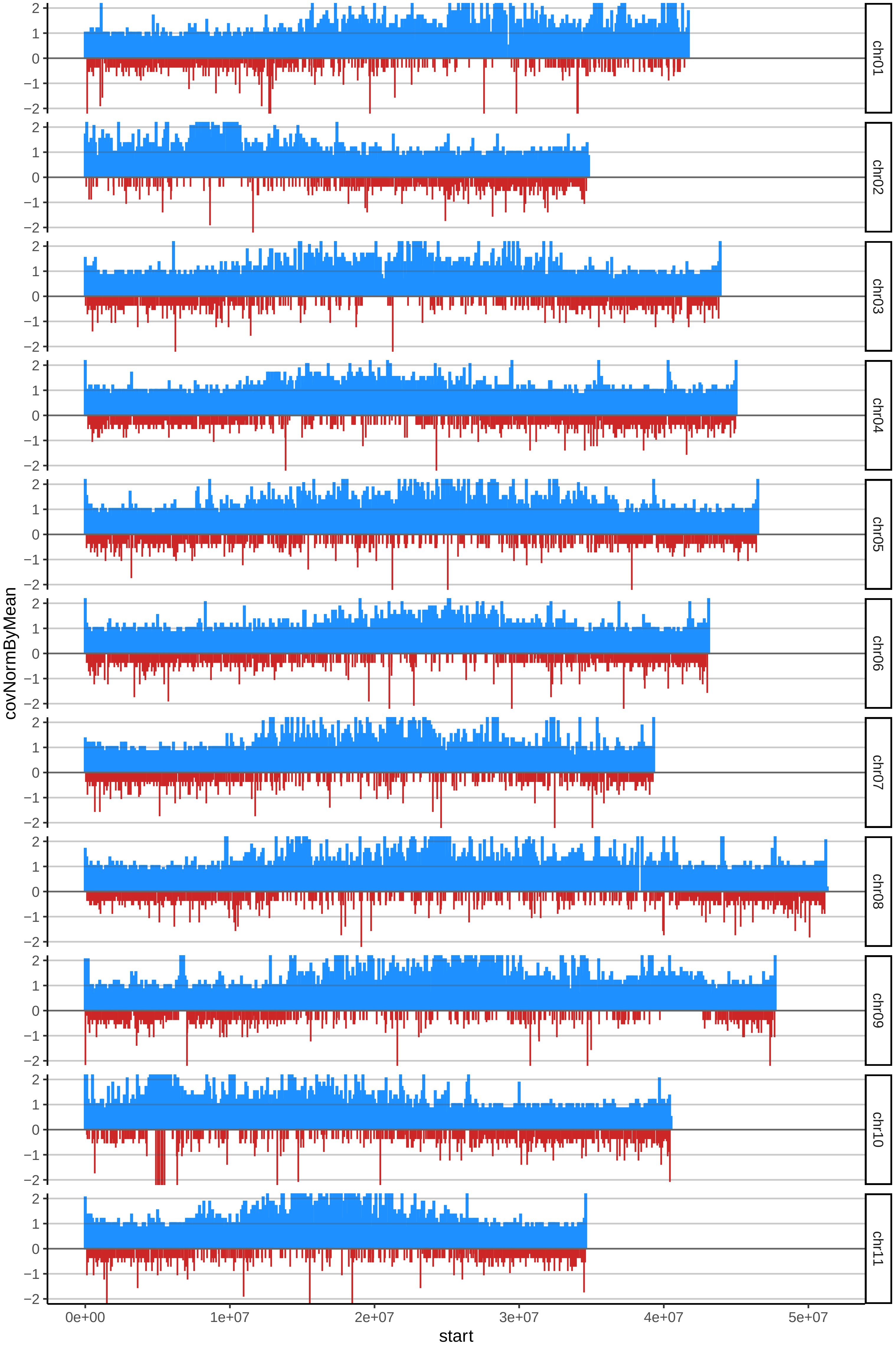

### 173SAPI09cov_byB.pdf

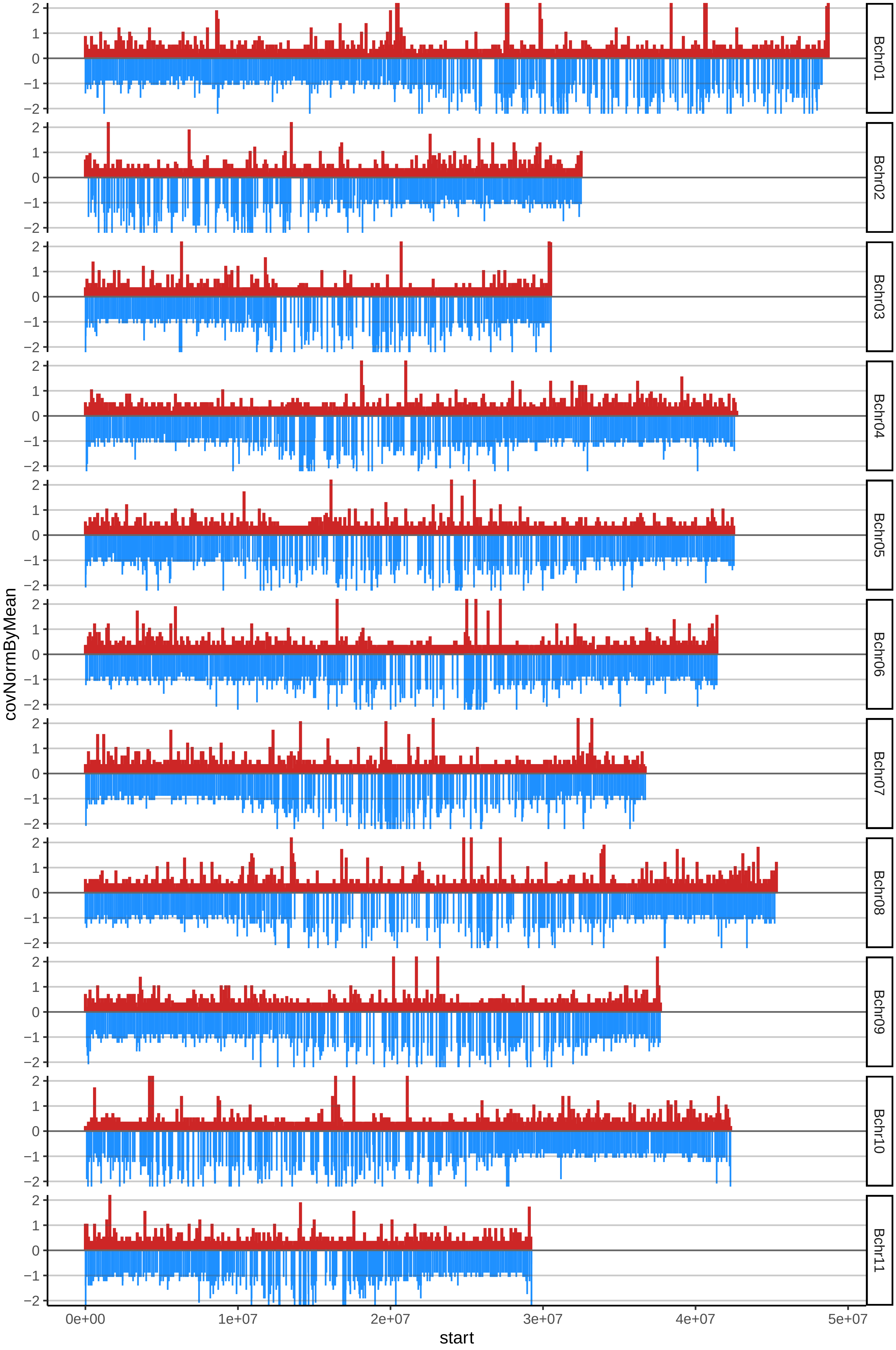

### 174SARI10cov_byA.pdf

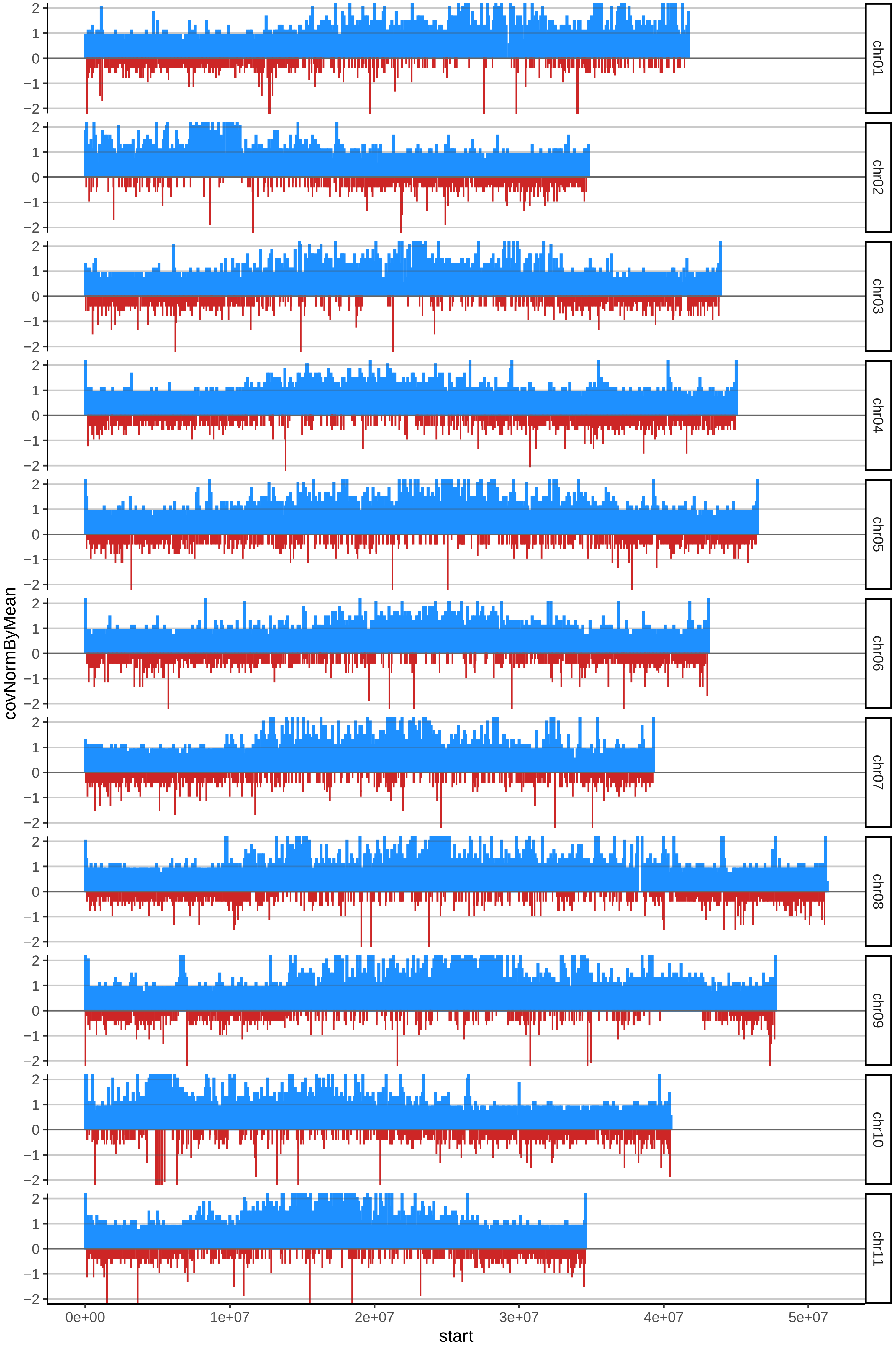

### 174SARI10cov_byB.pdf

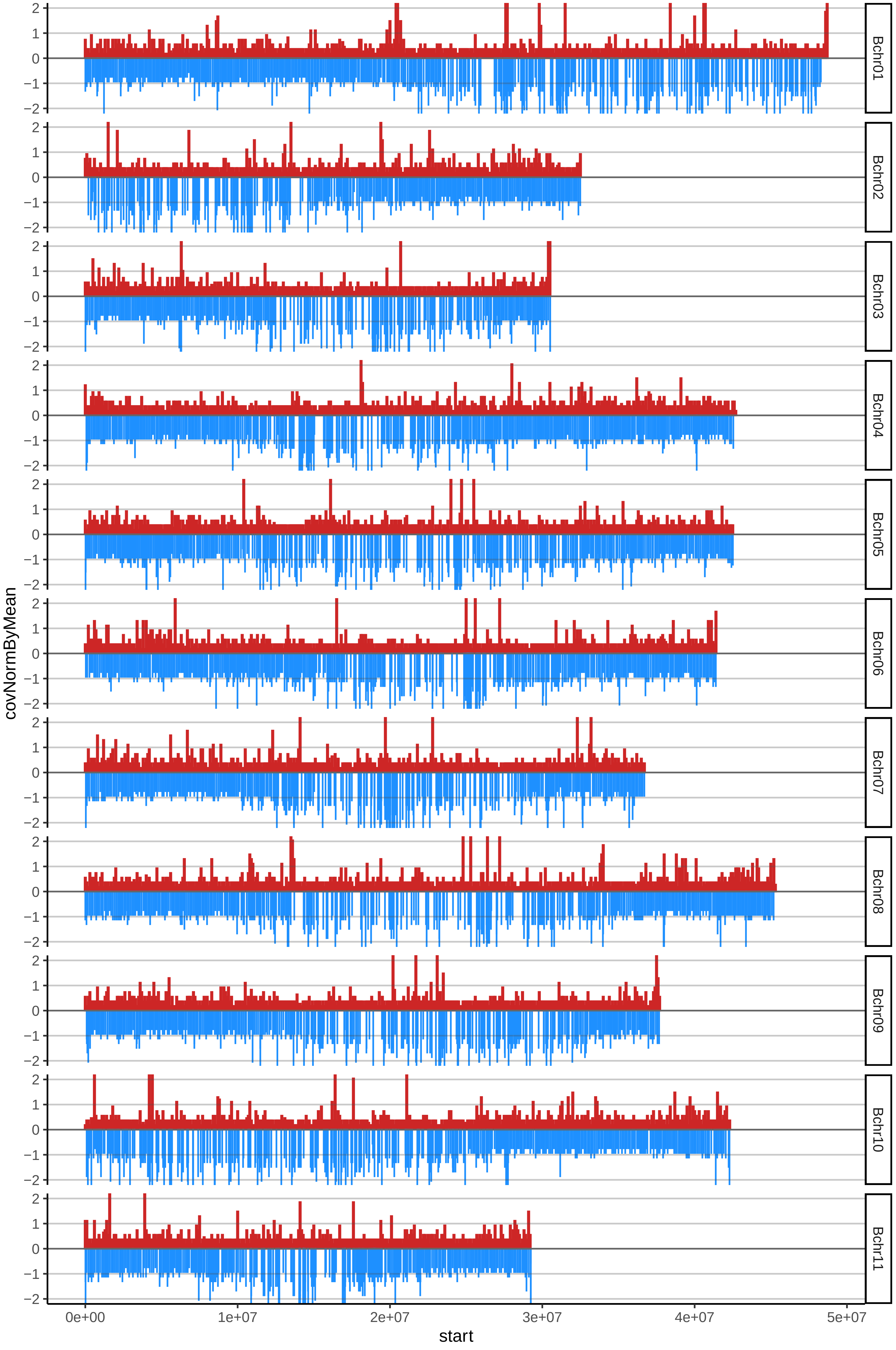

### 175SAPI08cov_byA.pdf

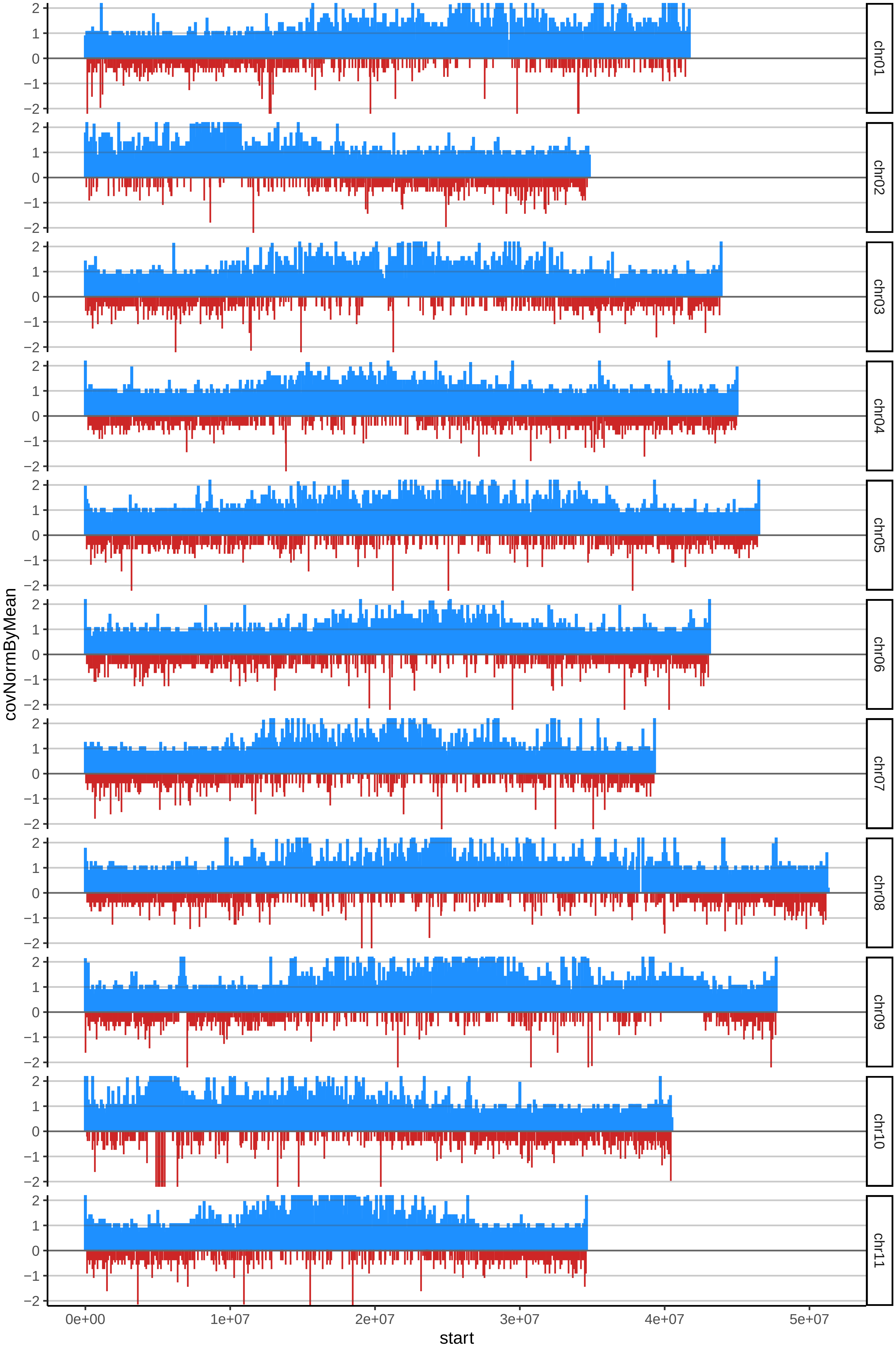

### 175SAPI08cov_byB.pdf

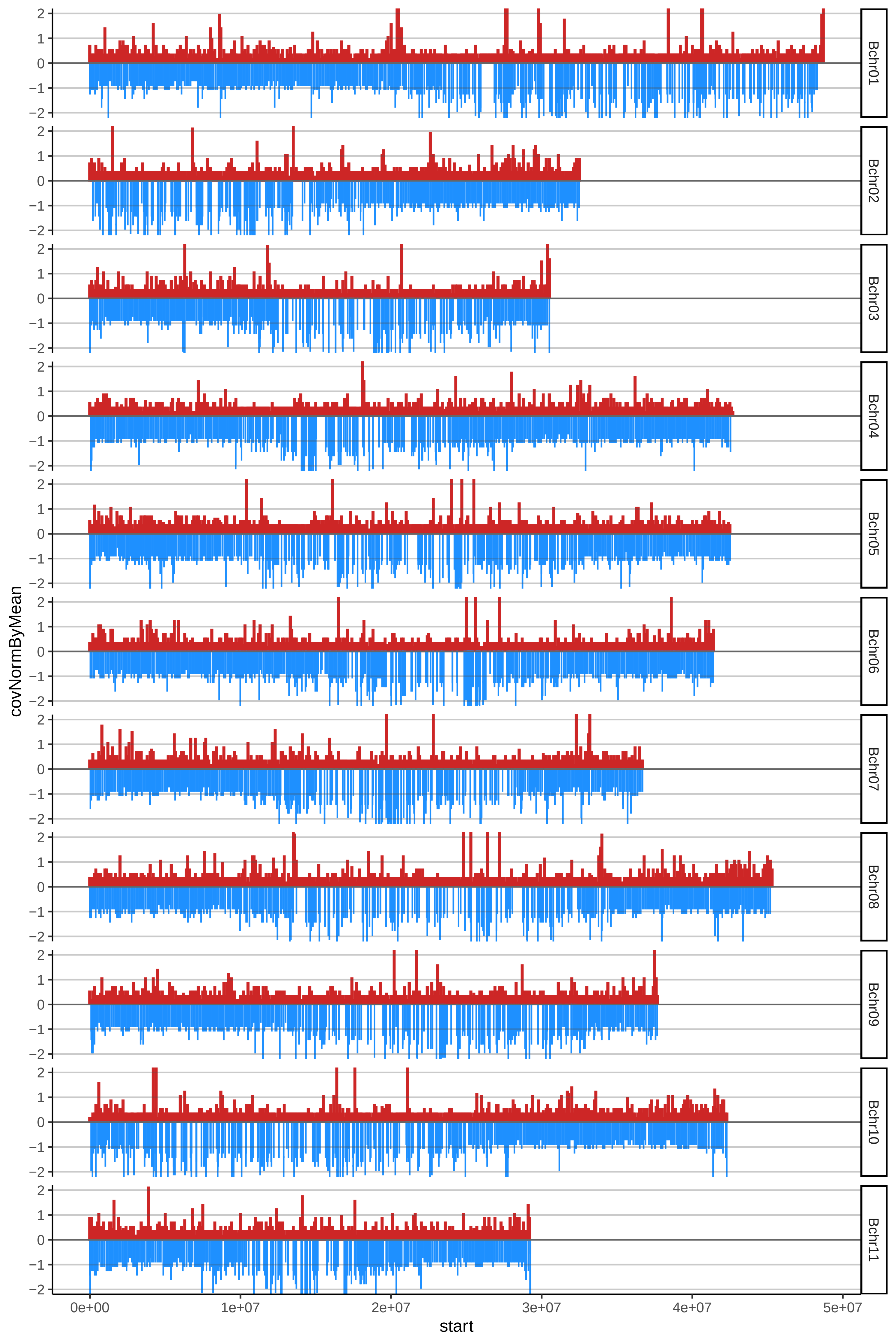

### 178VABU04cov_byA.pdf

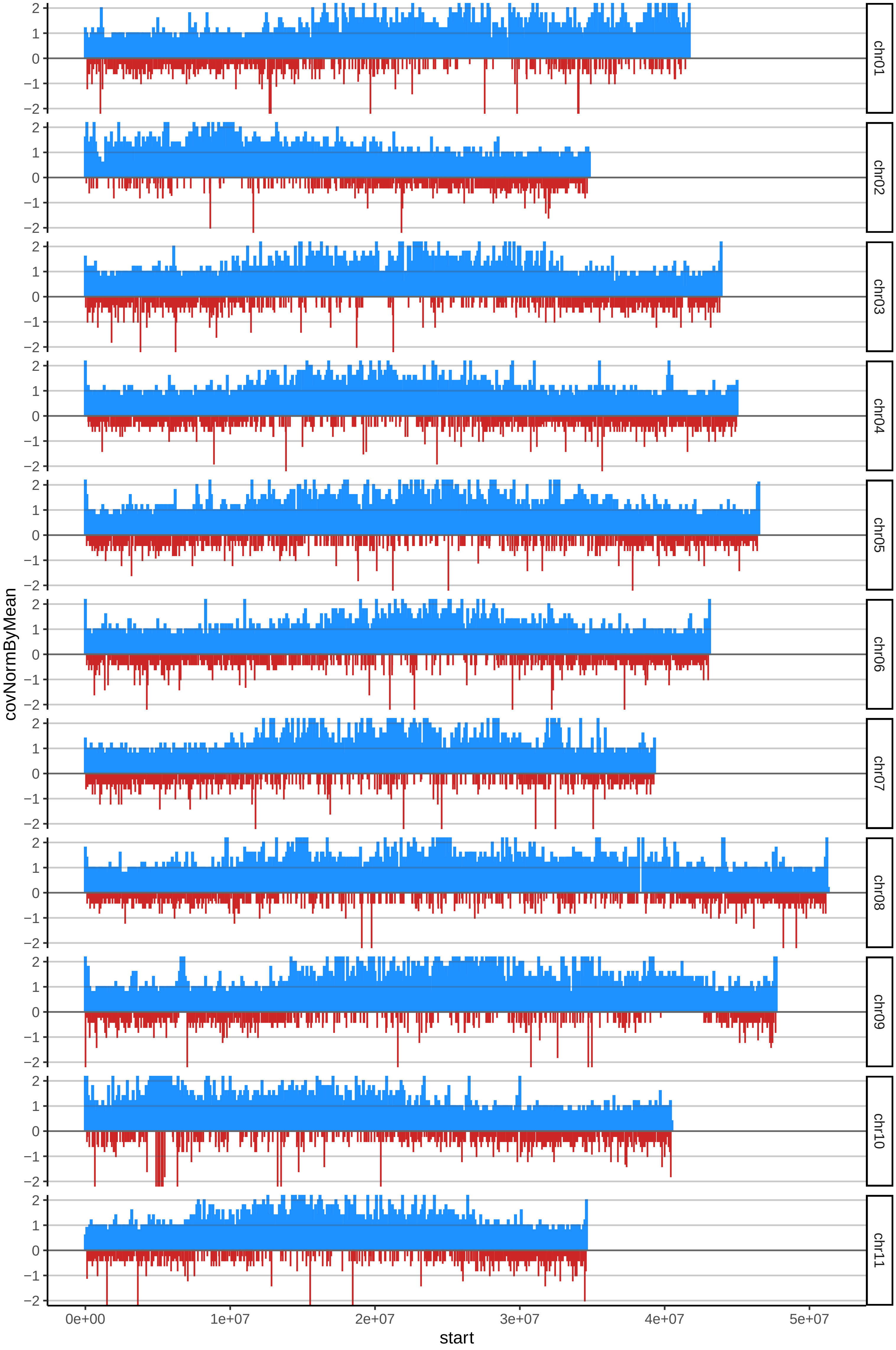

### 178VABU04cov_byB.pdf

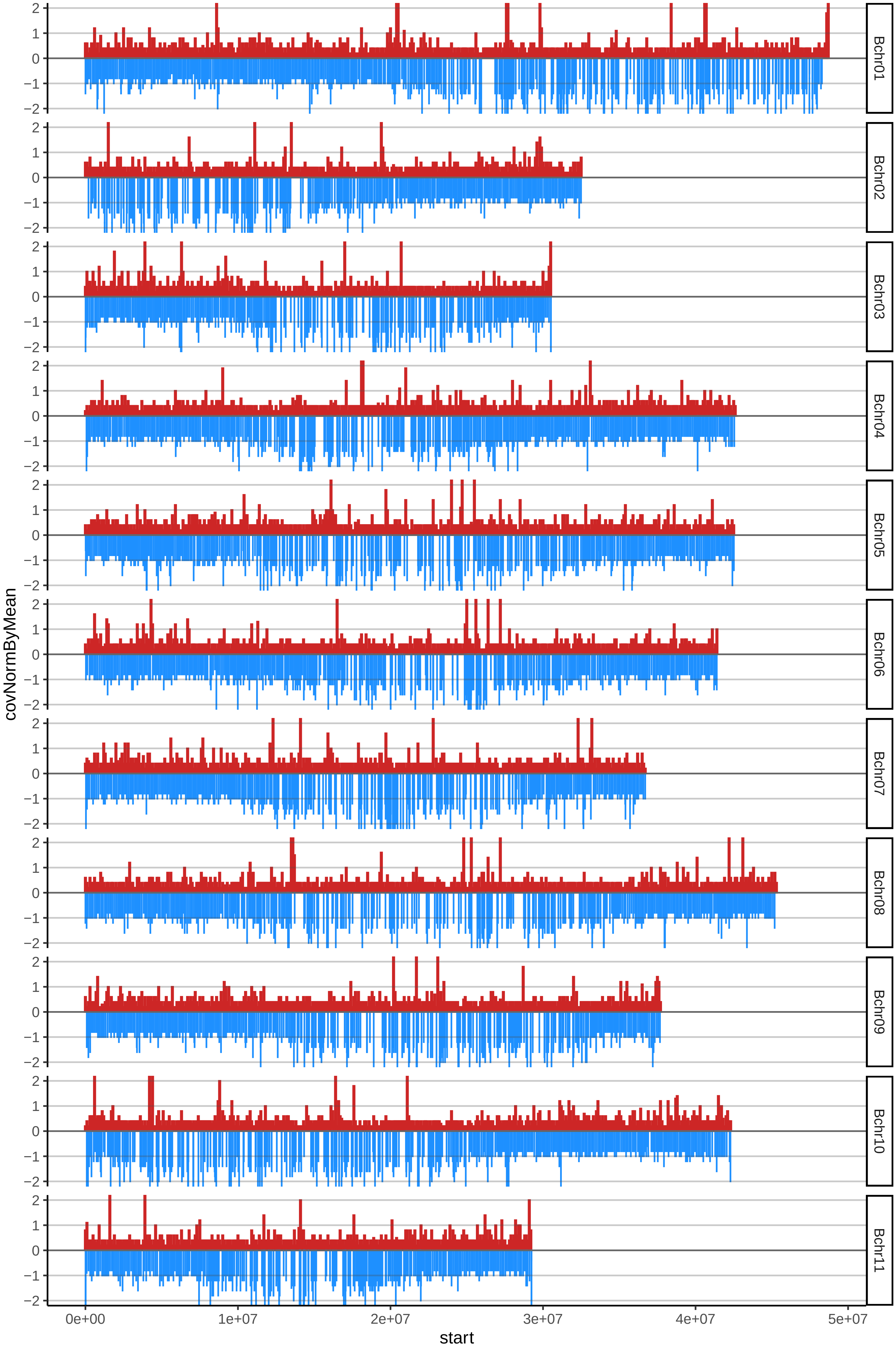

### 188RESTREPO01cov_byA.pdf

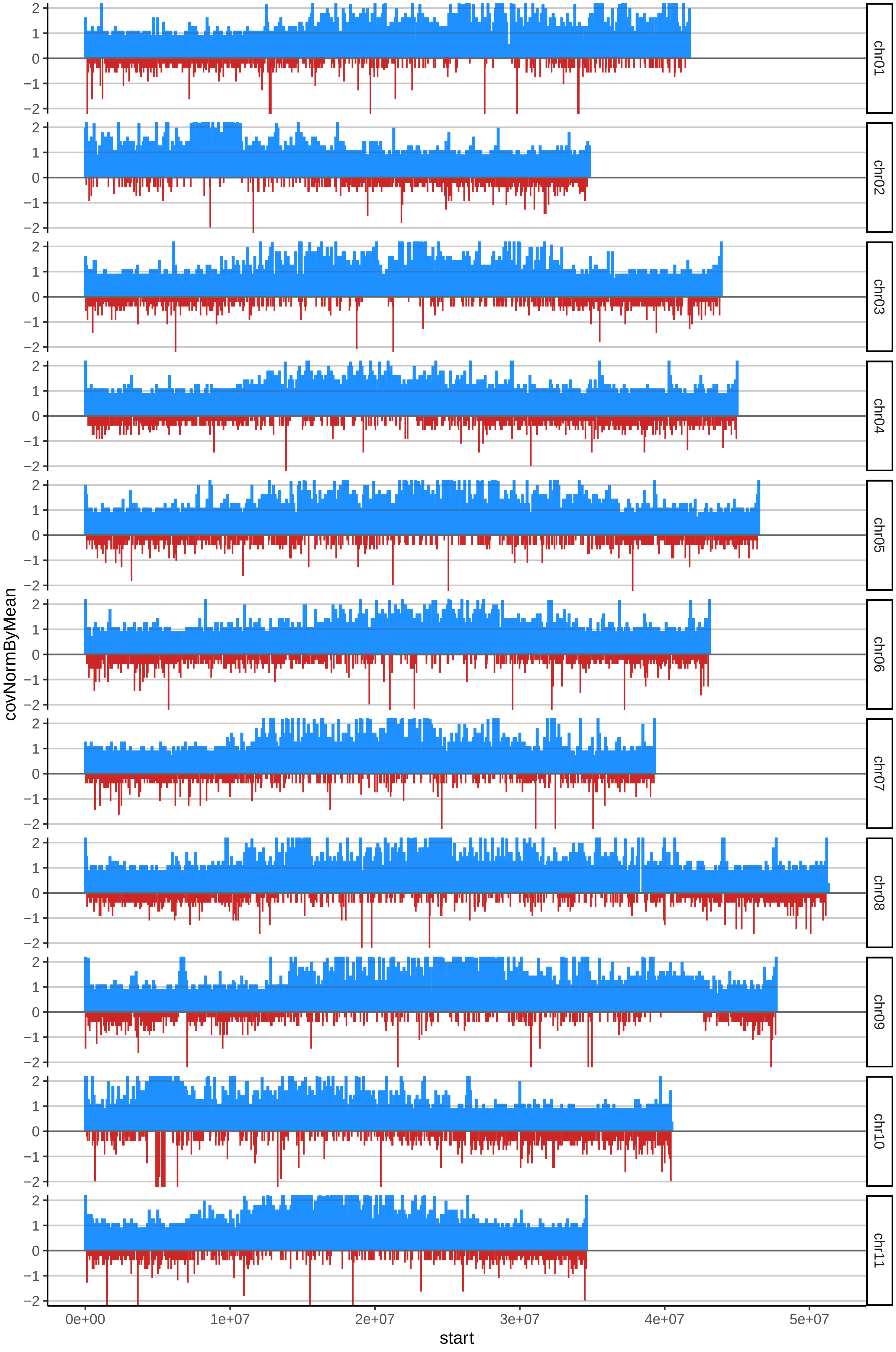

### 188RESTREPO01cov_byB.pdf

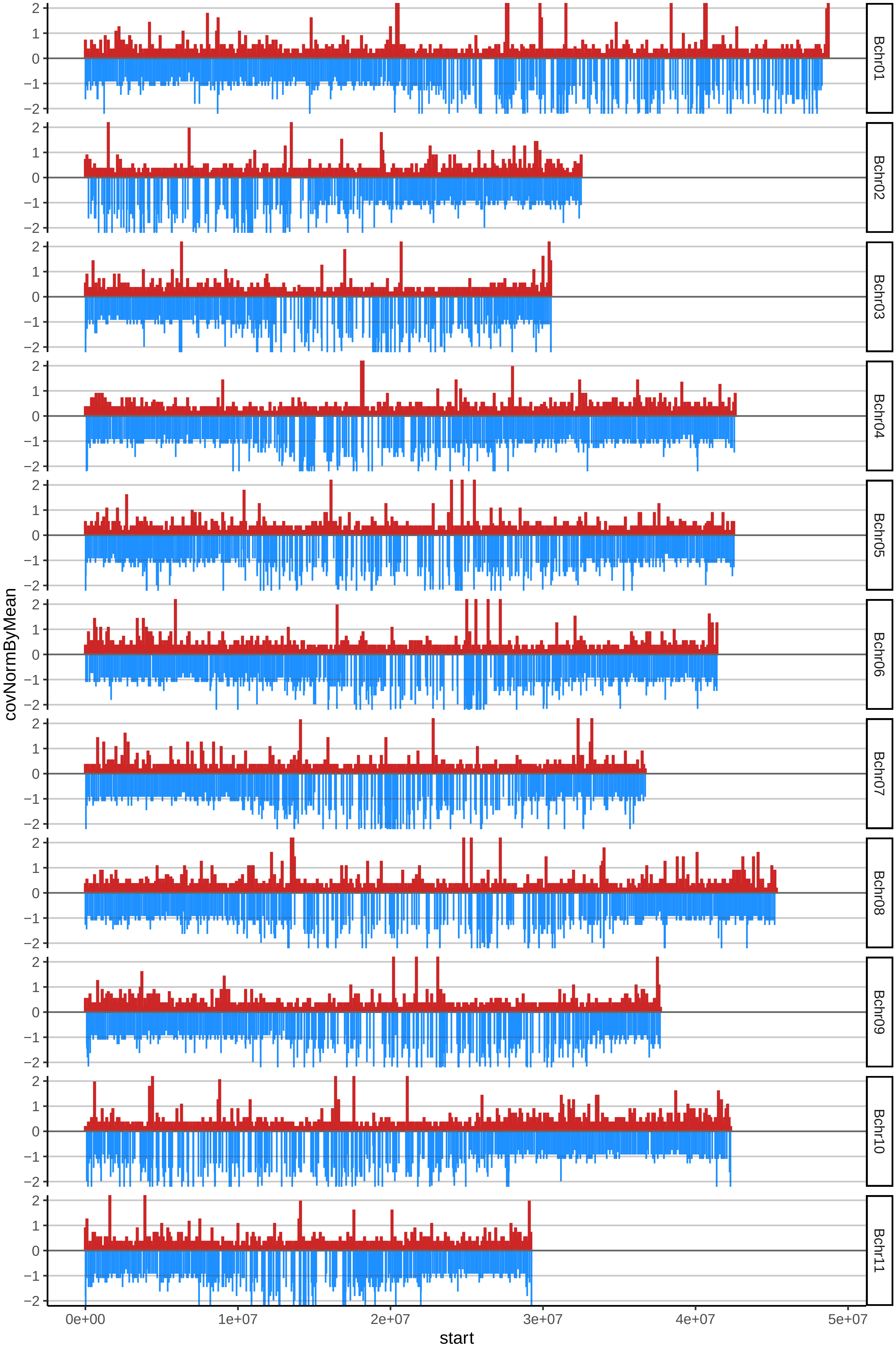

### 190LIBERALcov_byA.pdf

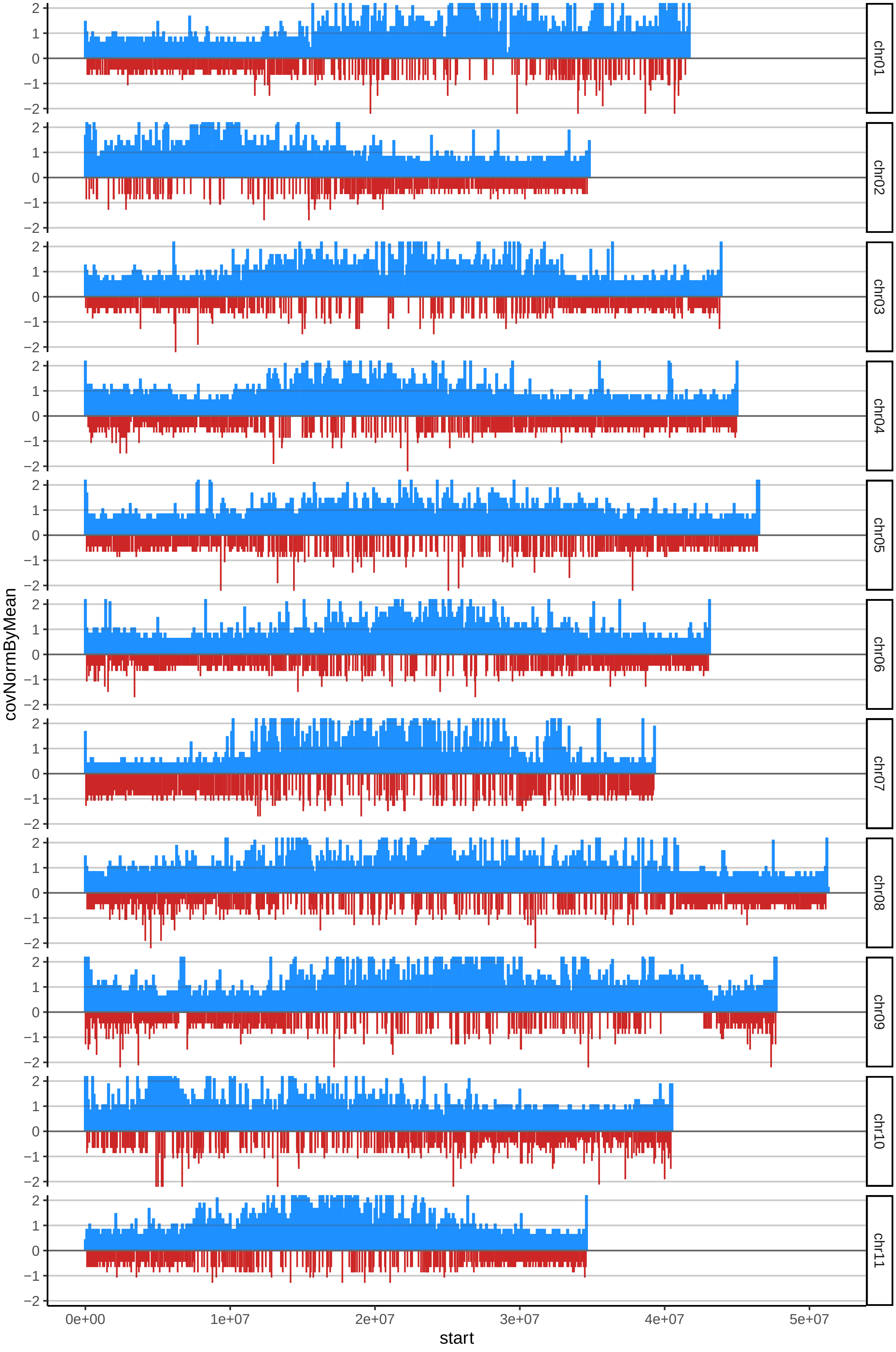

### 190LIBERALcov_byB.pdf

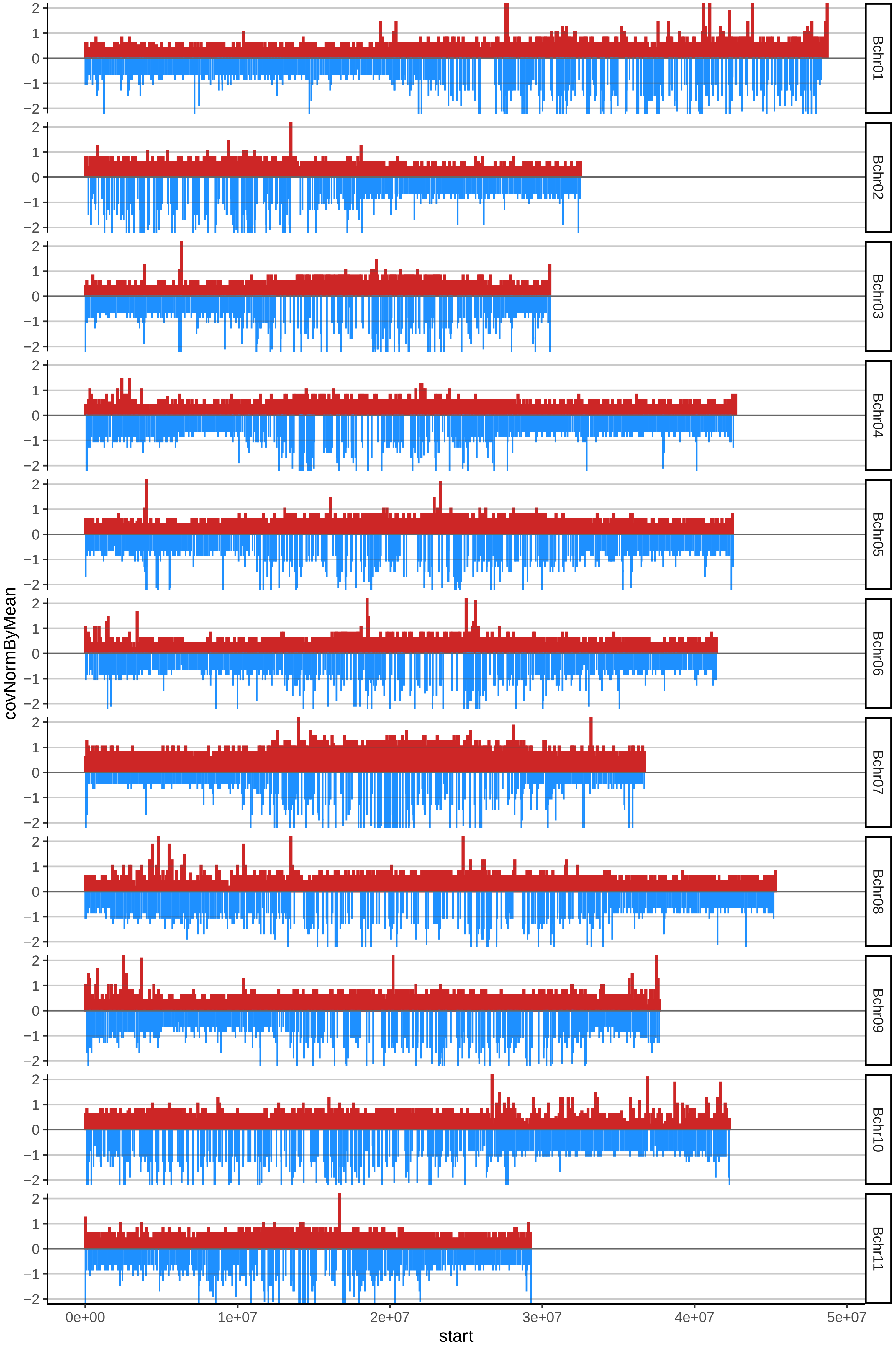

### 193BANANOINDIOCOMESENTAOcov_byA.pdf

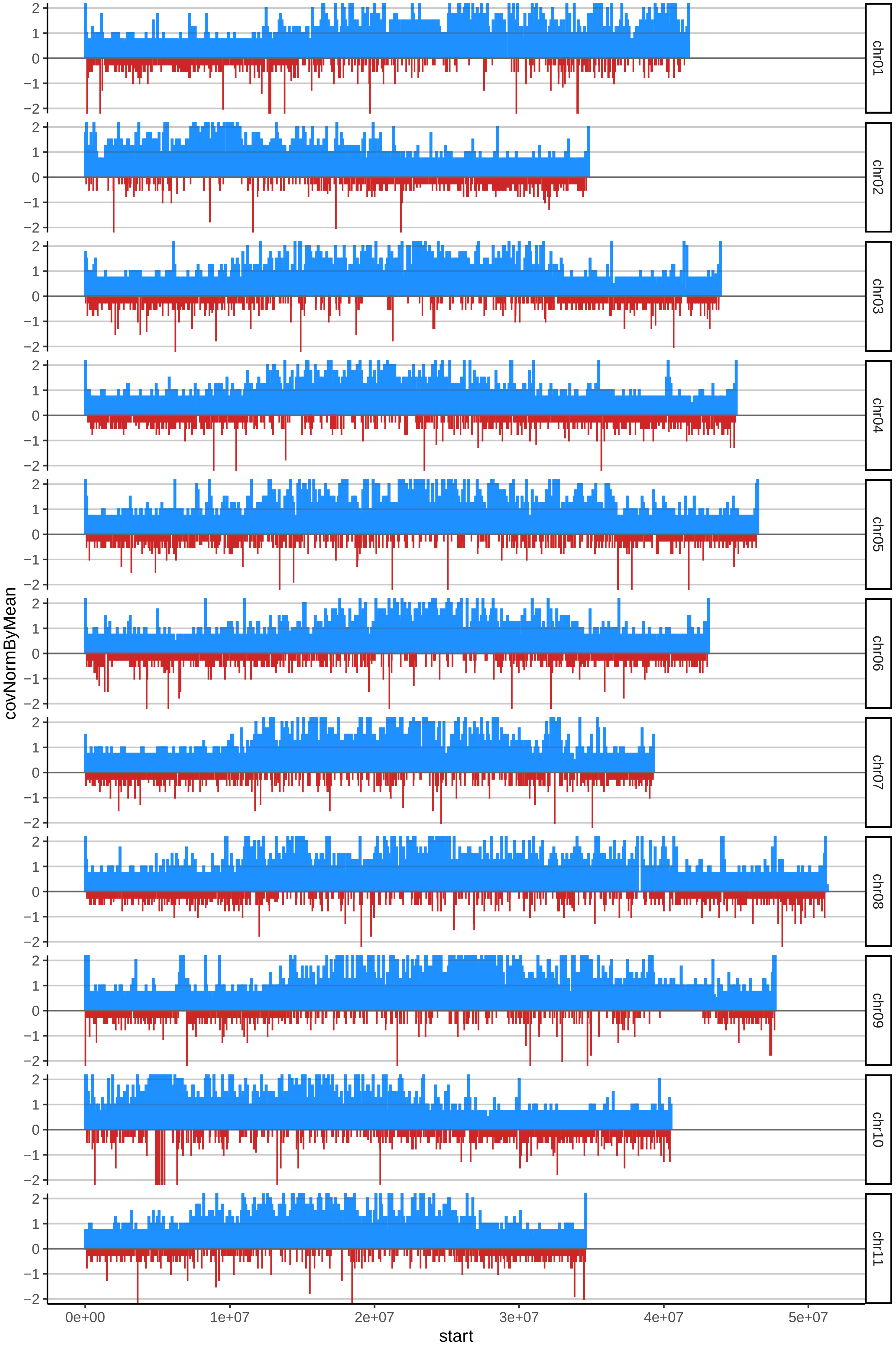

### 193BANANOINDIOCOMESENTAOcov_byB.pdf

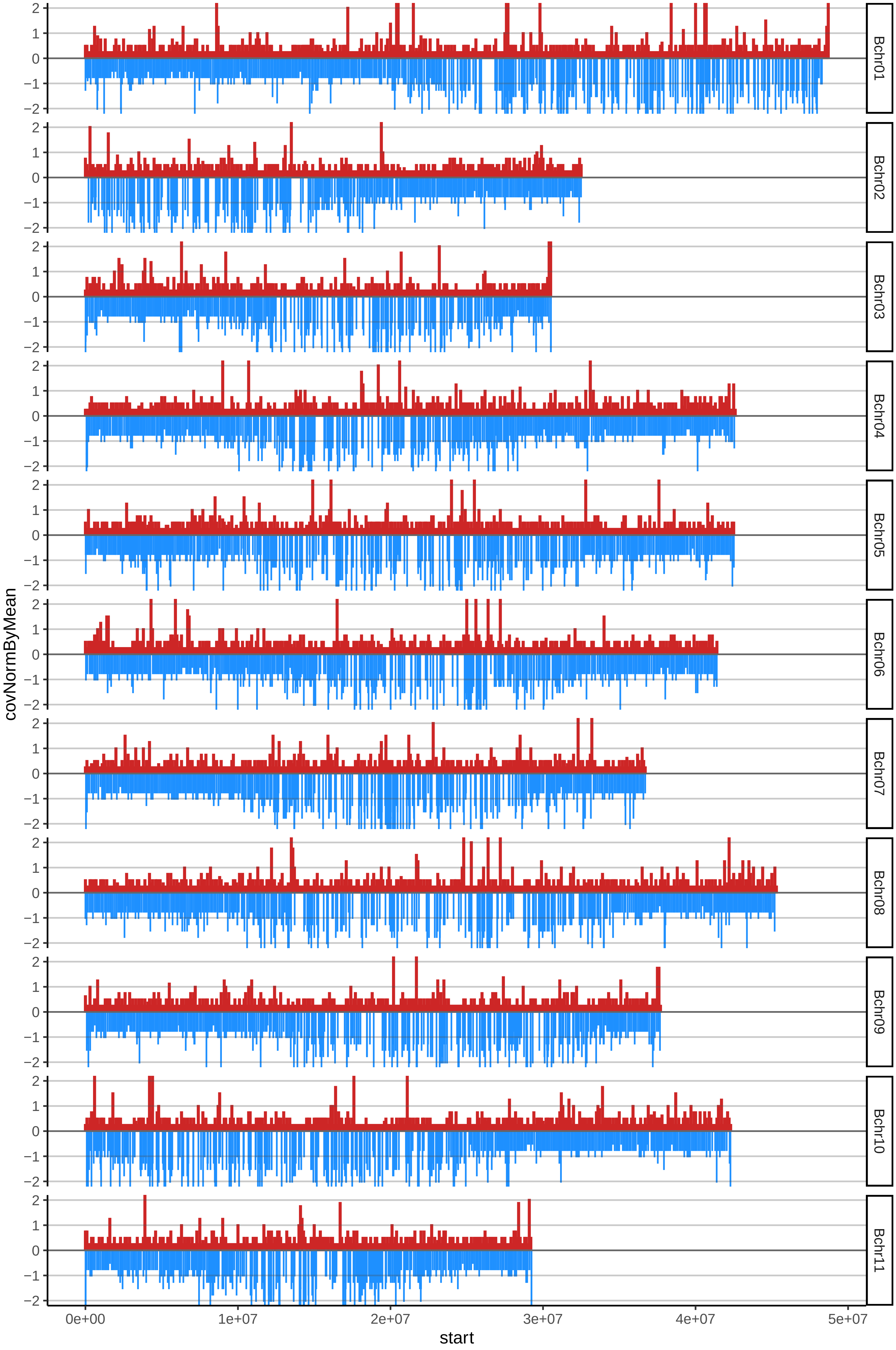

### 200TRIRACIMOcov_byA.pdf

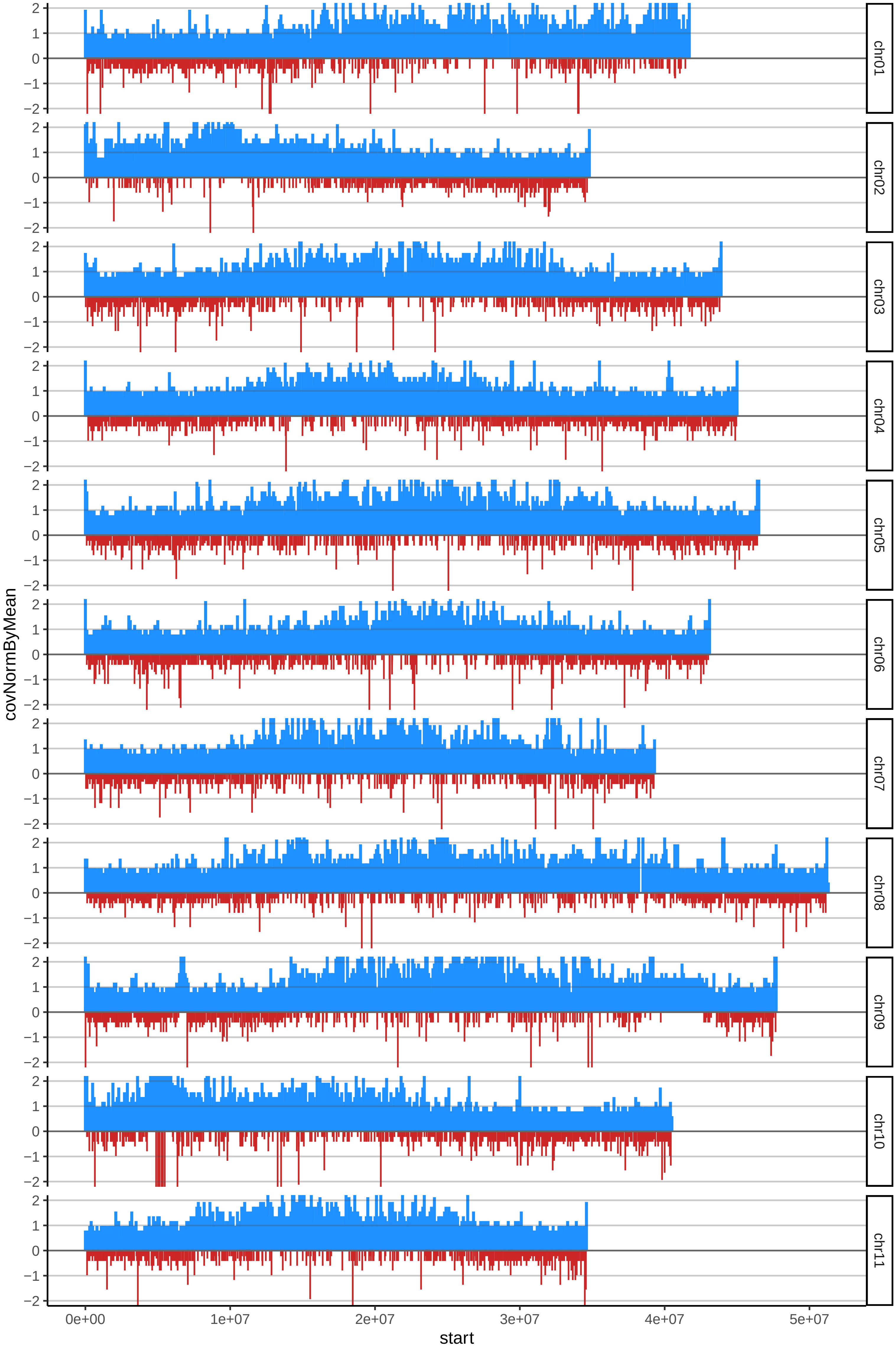

### 200TRIRACIMOcov_byB.pdf

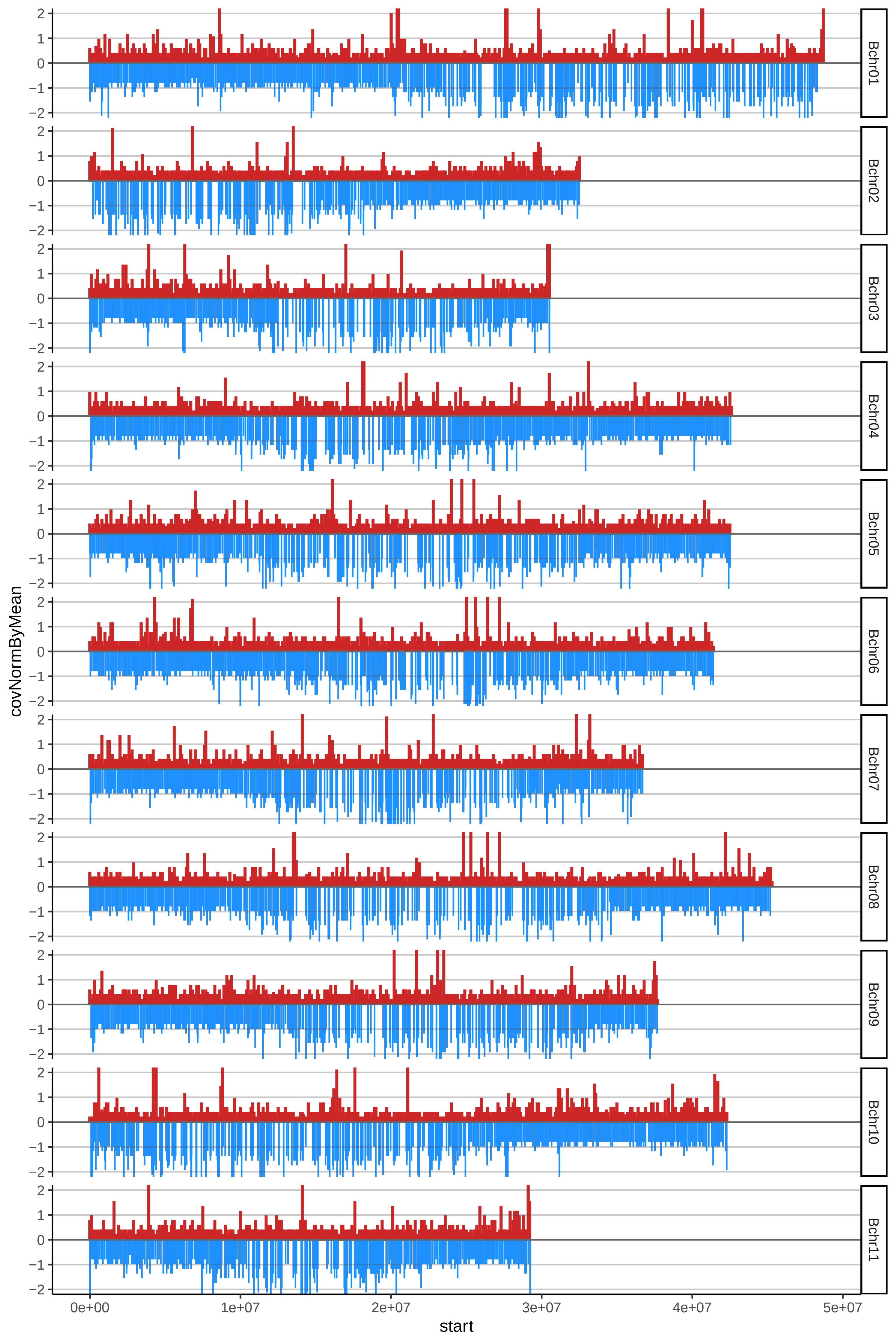
