## additional file 6 for "Characterising genome composition and large structural variation in banana varietal groups"

### Sucrier_AAA

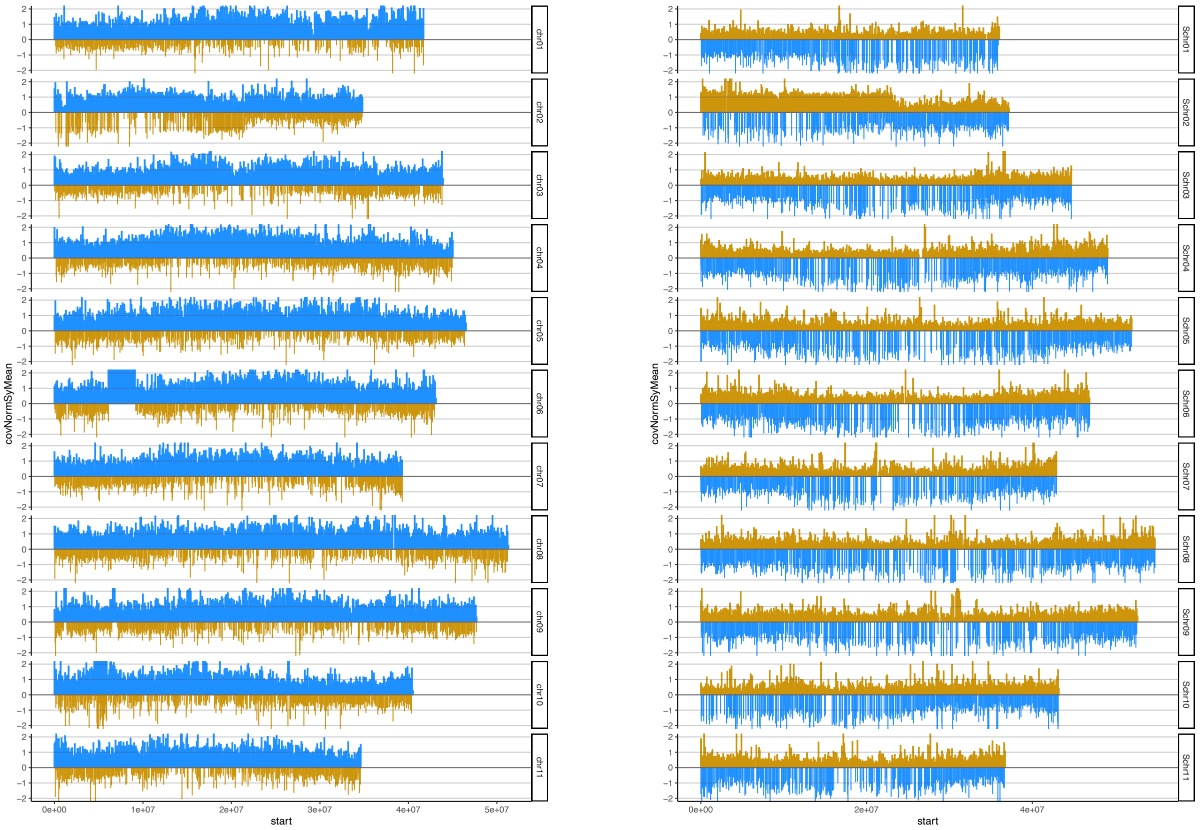


### Cavendish_AAA

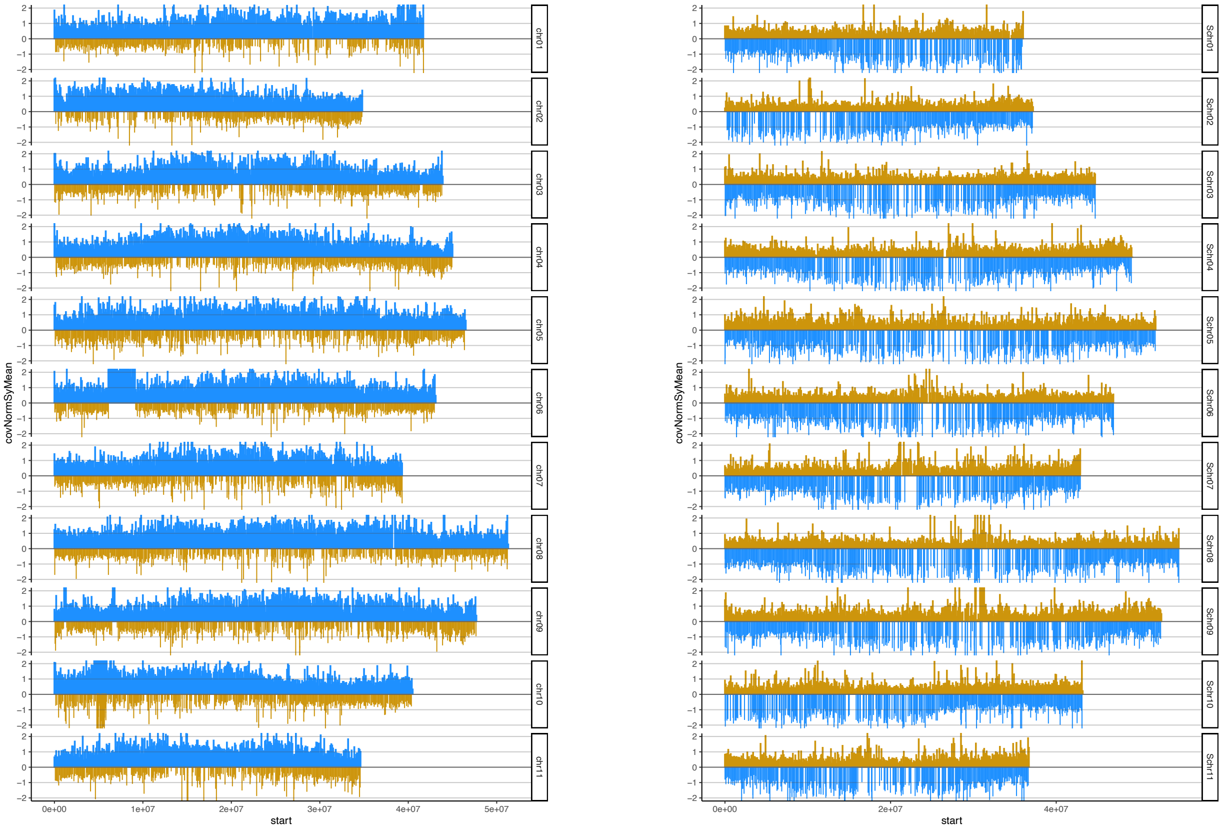

### Gros Michel_AAA

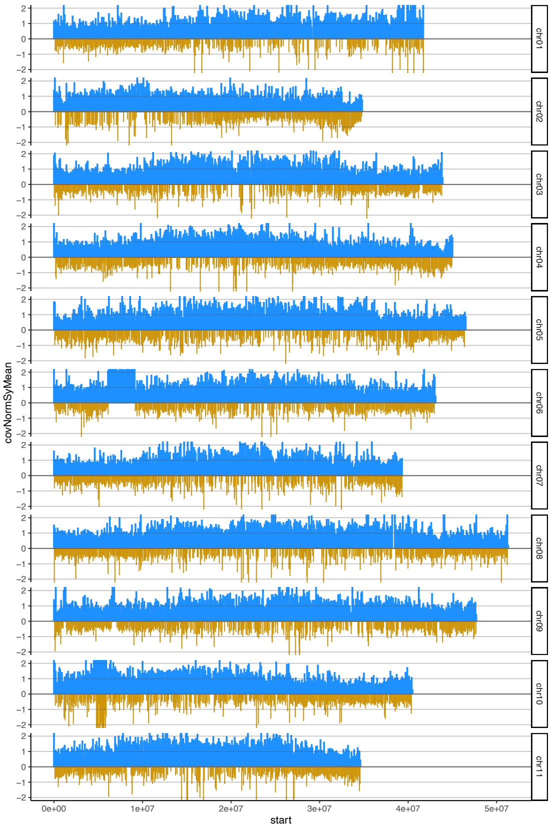


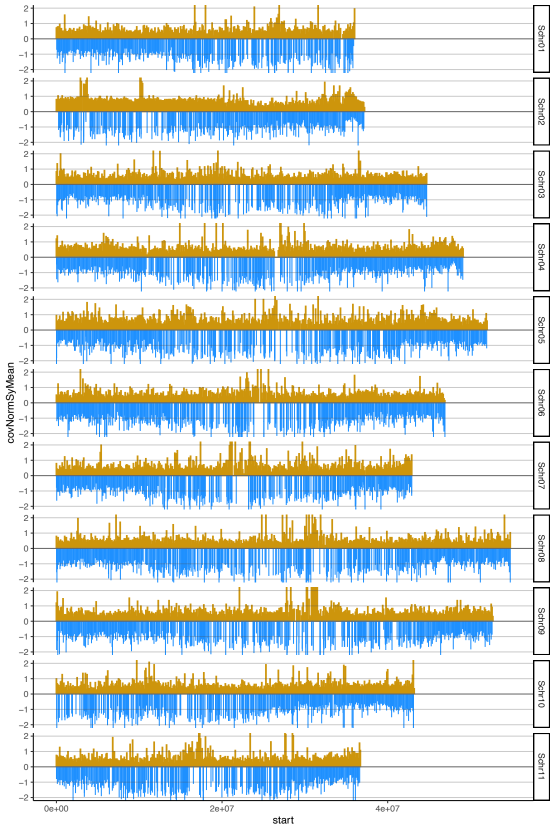


### unknown_AAA

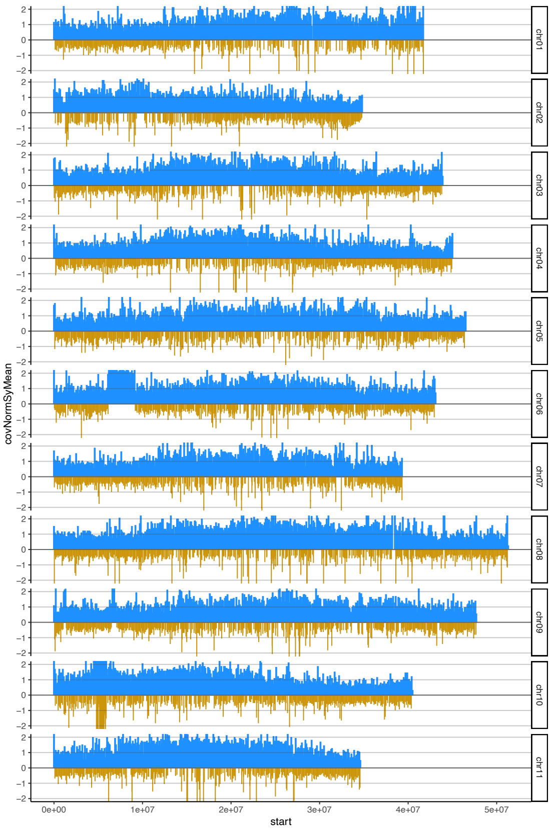


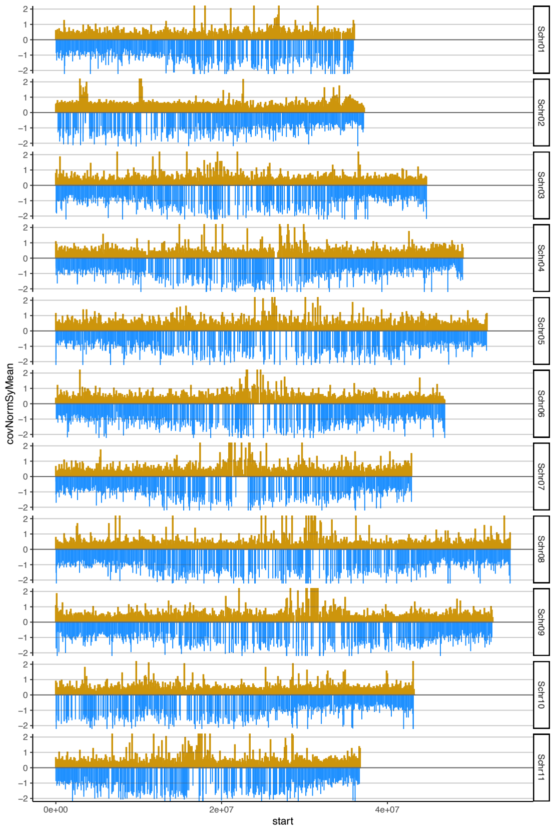

### Red_AAA

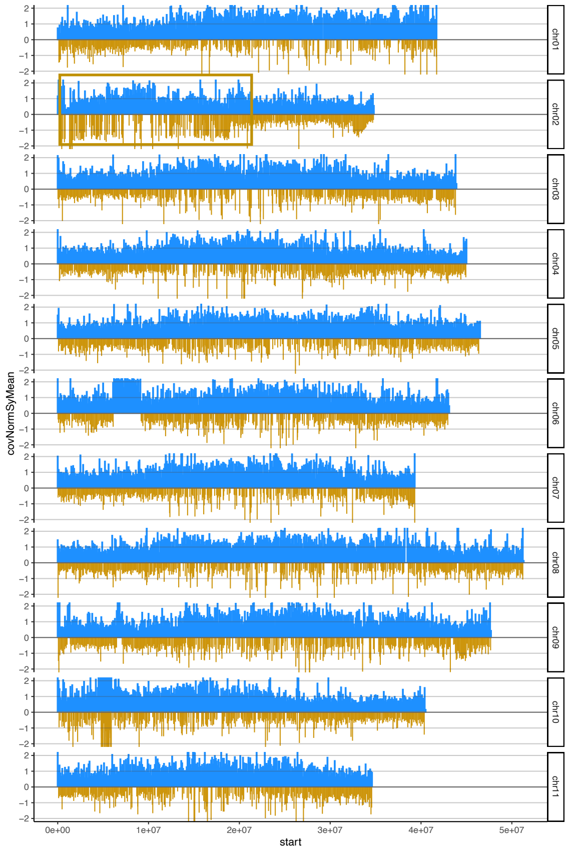


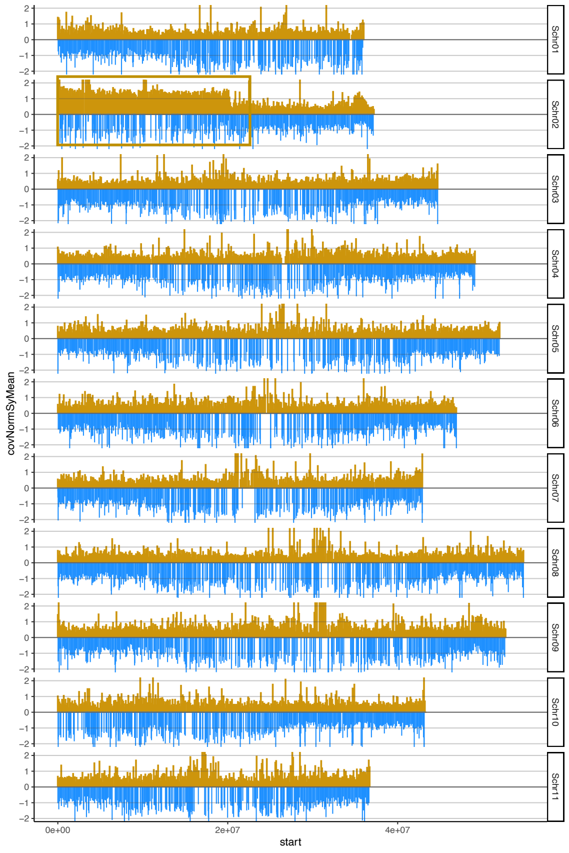


### Mutika AAA

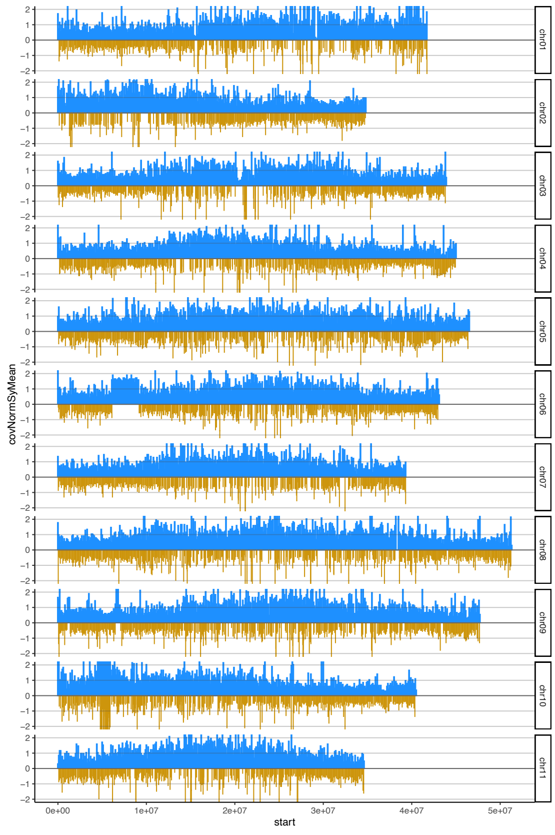


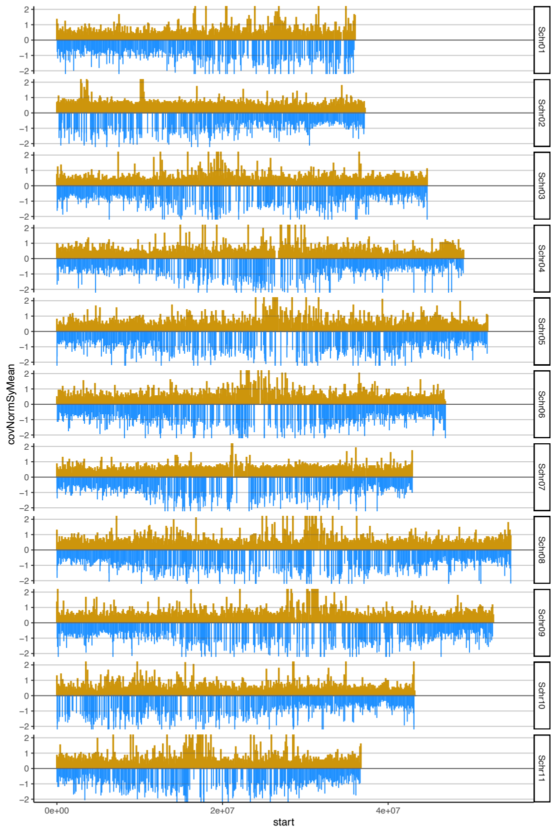

### Popoulu AAB

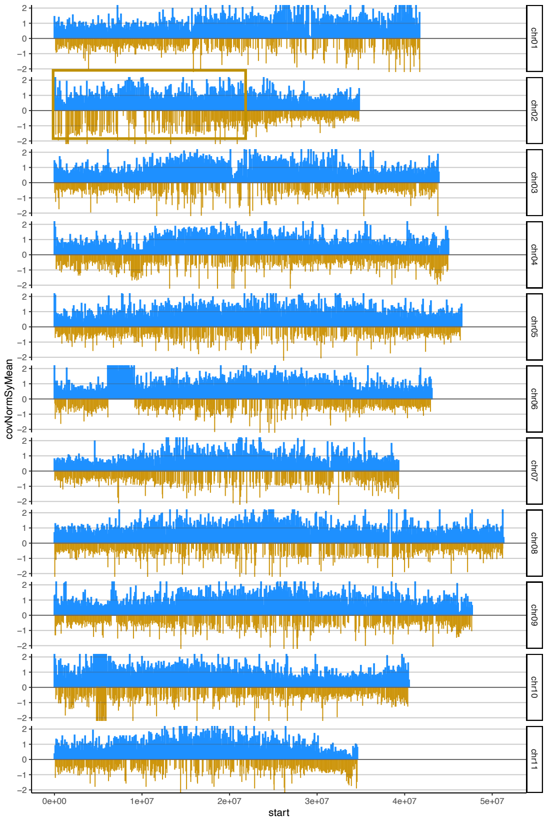


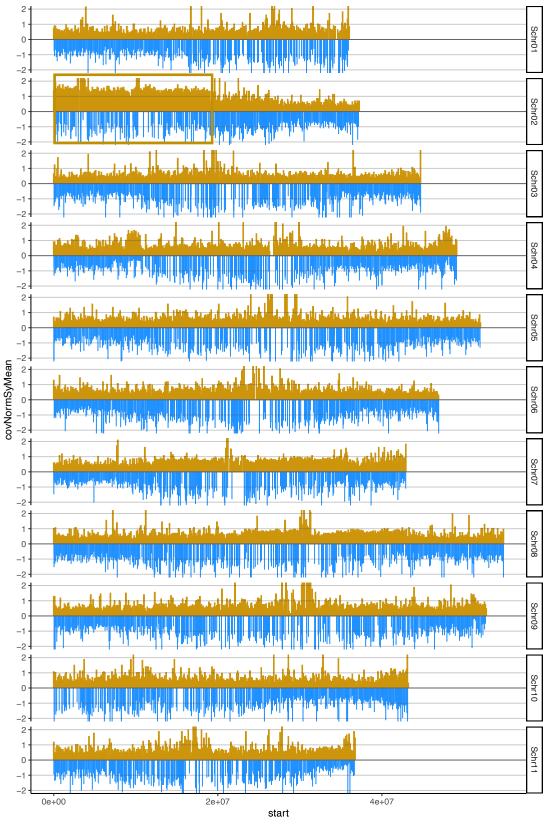


### Plantain AAB

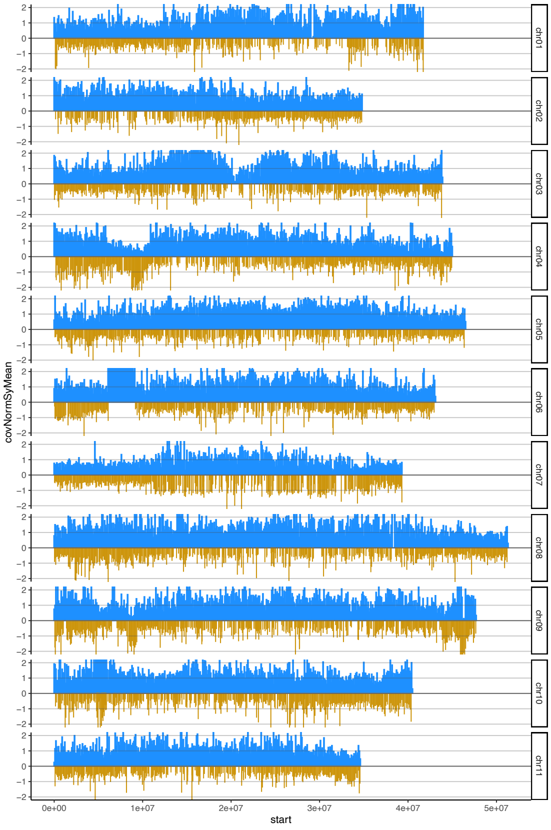


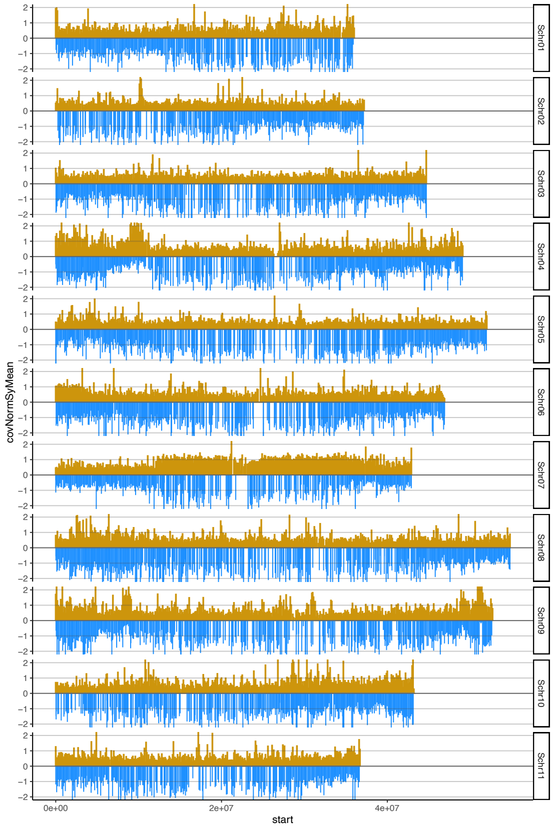

### Bluggoe_ABB

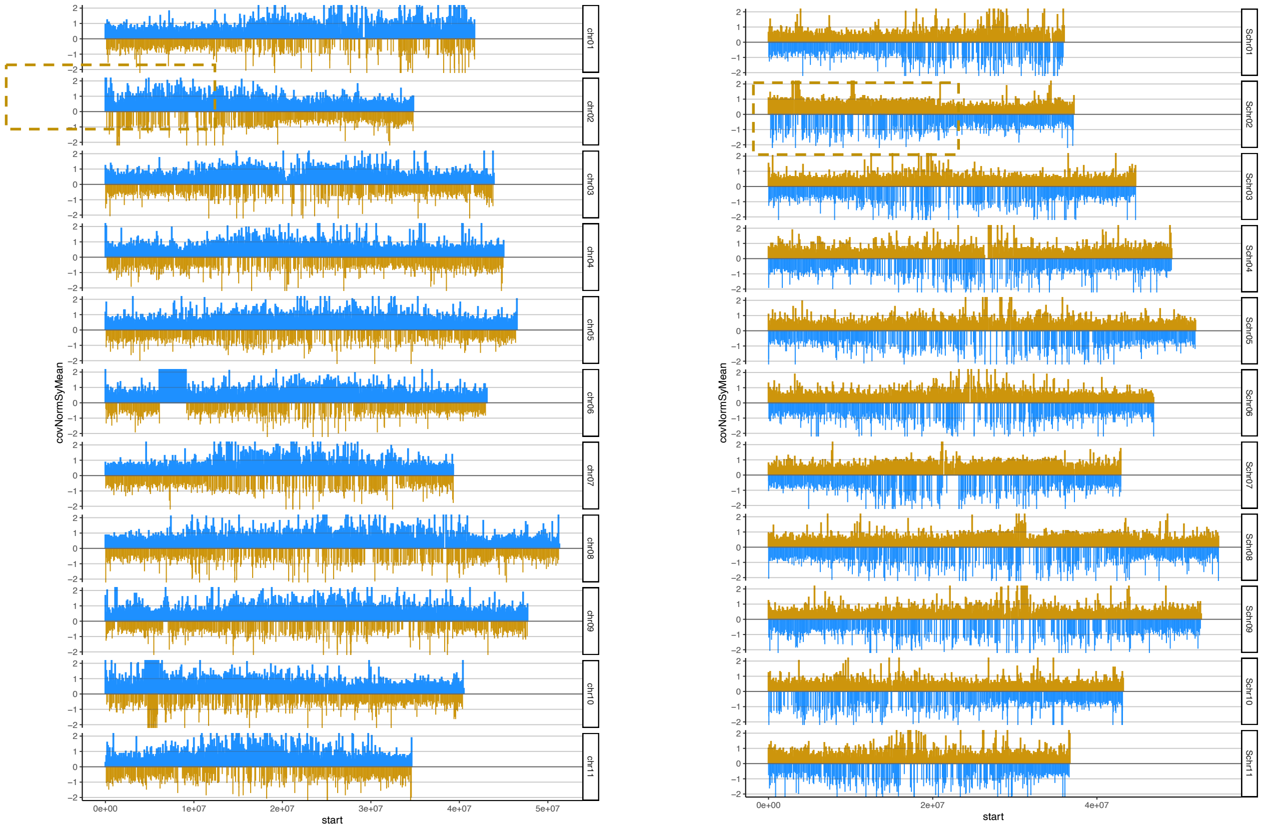


### Pelipita_ABB

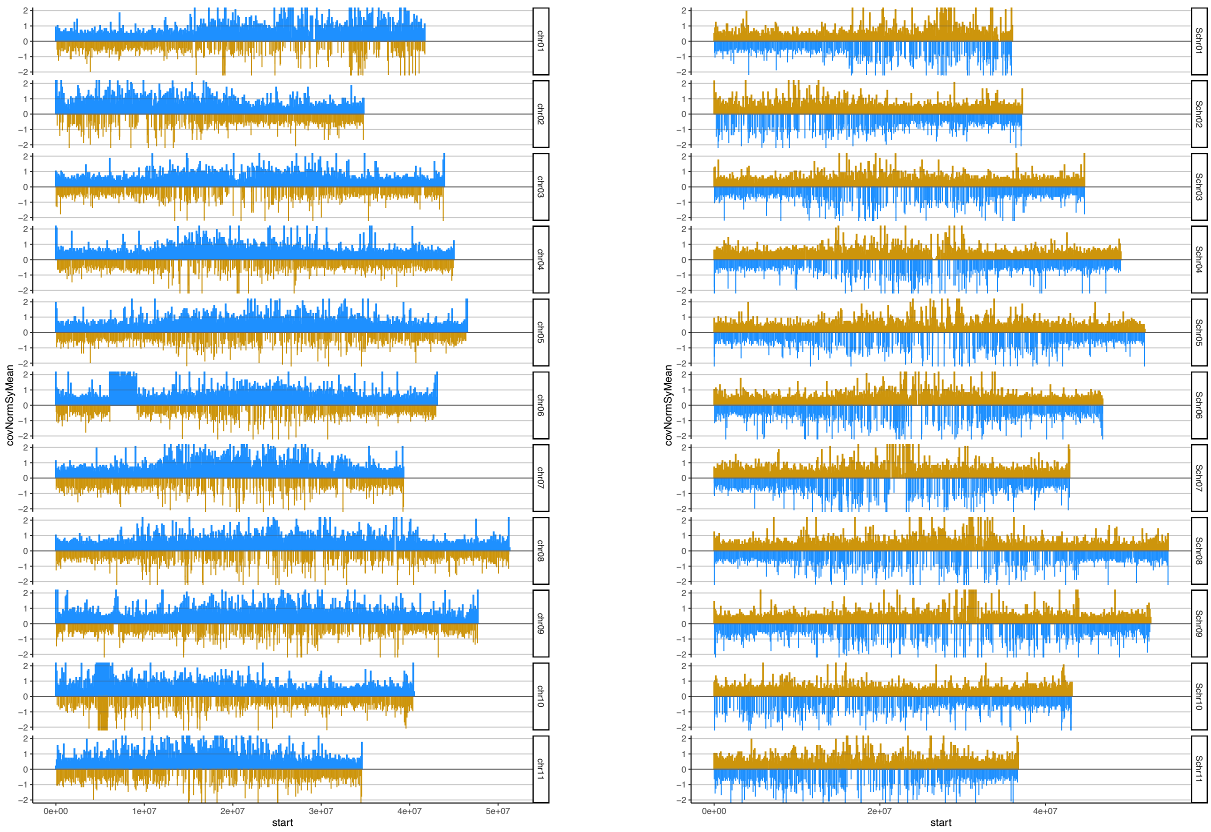

### AAAA

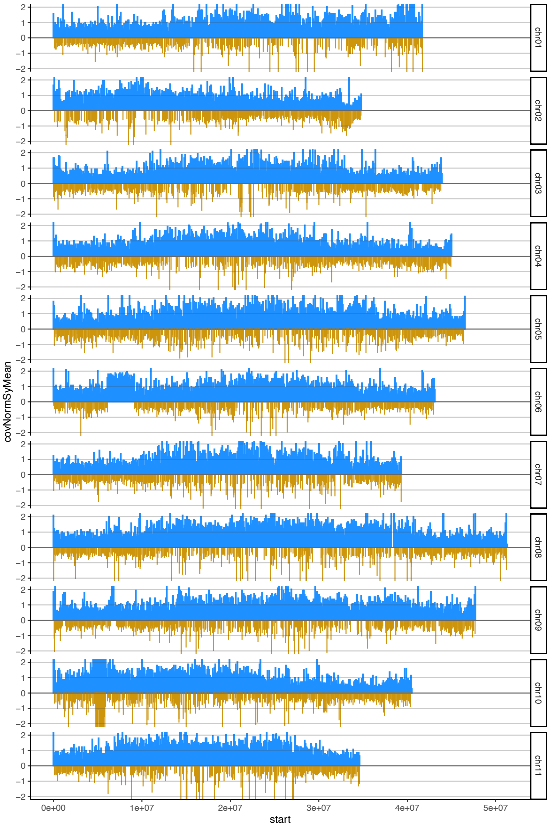
covNormSyMean
start
chr01
chr02
chr03
chr04
chr05
chr06
chr07
chr08
chr09
chr10
chr11
0e+00
1e+07
2e+07
3e+07
4e+07
5e+07

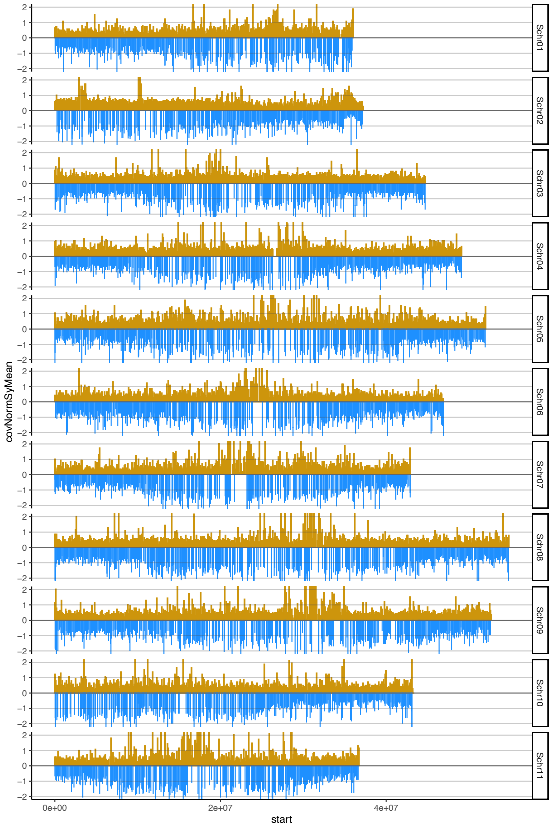
covNormSyMean
start
Schr01
Schr02
Schr03
Schr04
Schr05
Schr06
Schr07
Schr08
Schr09
Schr10
Schr11
0e+00
2e+07
4e+07

### AAAB_Pome

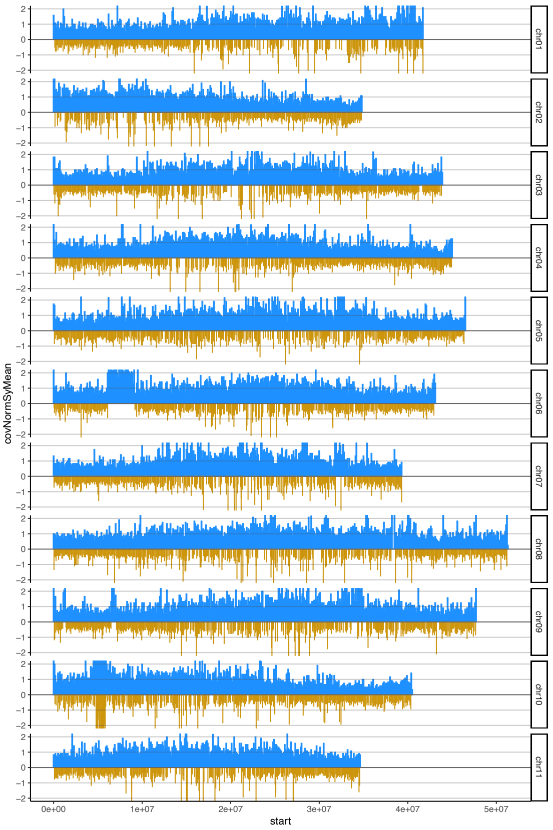


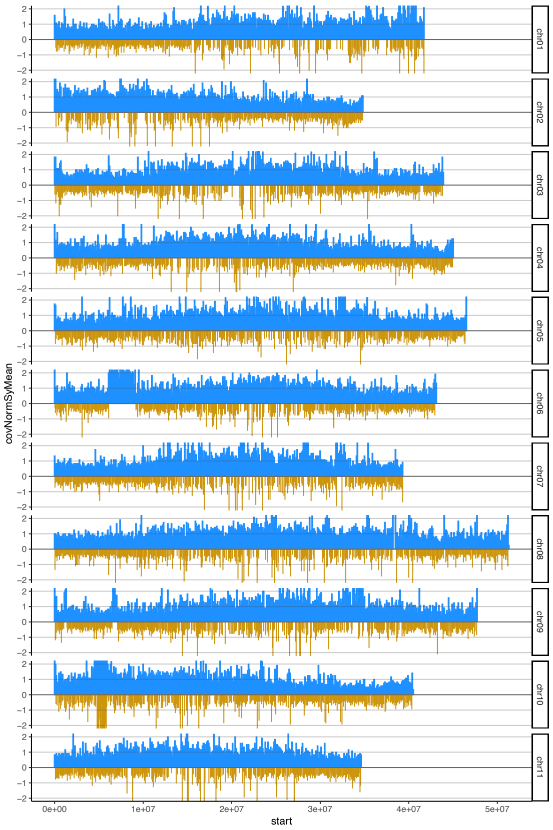


### AAAB_Africa

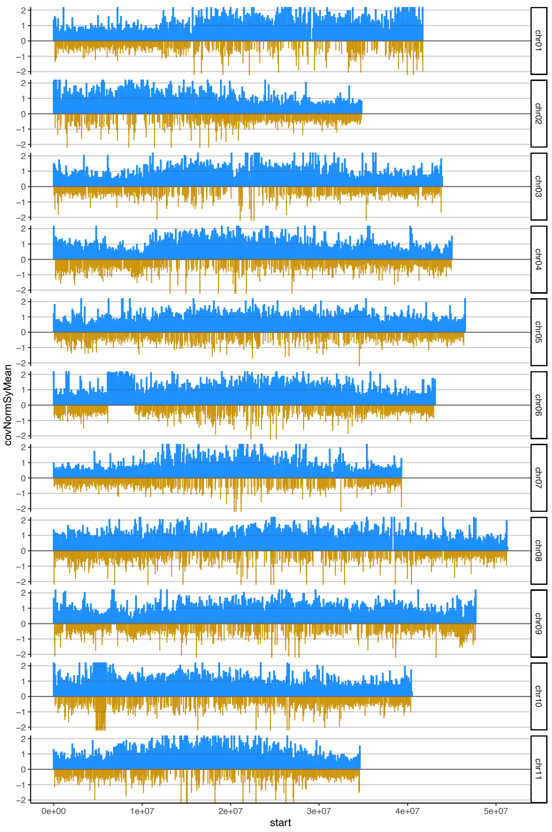


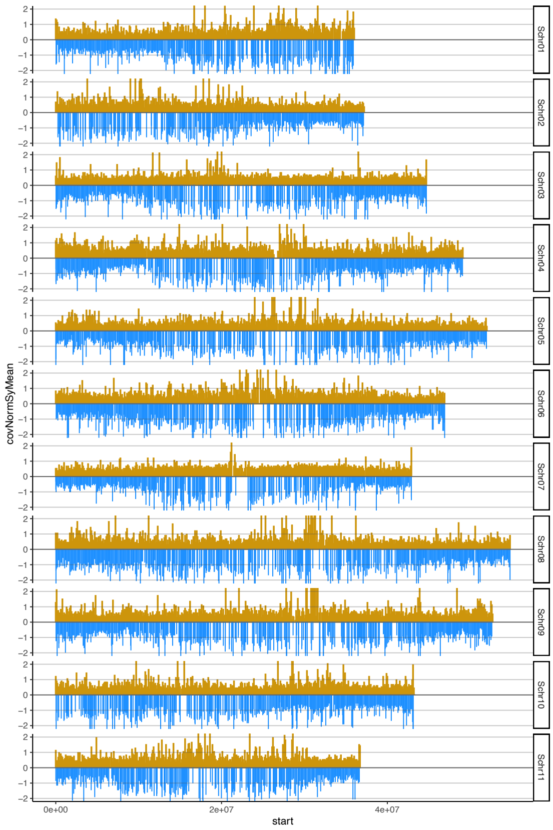
